## Supplemental Figures 1-14, Supplemental Text and Bibliography for "ASCOT identifies key regulators of neuronal subtype-specific splicing"

### Supplemental File Table of Contents


Associated with Figure 1. MESA

Supplemental Figure 1. Diagram of "junction-walking" strategy for alternative splicing analysis.

Supplemental Figure 2. Binary decision splicing events represent most local splicing variation.

Supplemental Figure 3. Rods do not express many of the common neuronal splicing factors

Supplemental Figure 4. Examples of mouse neuronal subtype-specific exons (MESA)

Associated with Figure 2. GTEx

Supplemental Figure 5. Examples of human tissue-specific exons (GTEx)

Supplemental Figure 6. Rod-specific exons, both previously identified and novel

Supplemental Figure 7. Alternative splicing analysis of single-cell RNA-Seq data derived from full-length transcripts.

Associated with Figure 4. HepG2 overexpression & ENCODE

Supplemental Figure 8. Activation of rod-specific exons in HepG2 cells requires FACS isolation of the most strongly expressing cells

Supplemental Figure 9. Increased *MSI1* expression and *PTBP1* downregulation may interact synergistically to activate rod/neuronal alternative exons

Supplemental Figure 10. Upstream and downstream flanking sequences for rod-specific exons

Supplemental Figure 11. *Msi1* is necessary for rod photoreceptor-specific splicing

Associated with Discussion

Supplemental Figure 12. Many of the exons detected by ASCOT are unannotated

Supplemental Figure 13. Dataset clustering can improve the detection of binary splicing events

Supplemental Figure 14. Rod-specific exons in the SRAv2 Snaptron compilation

Personal clarification statement from Christopher Wilks

Bibliography

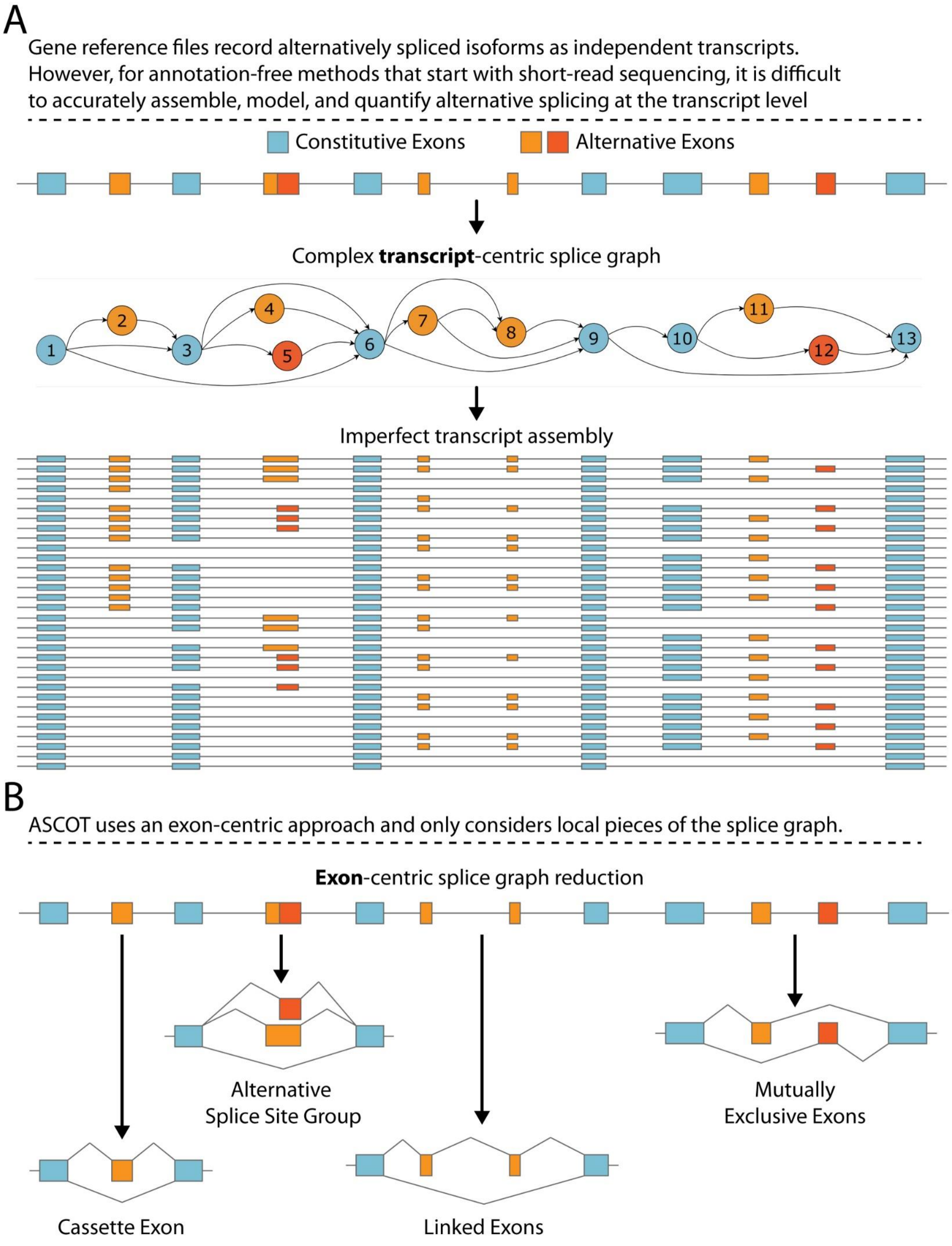
A
Gene reference files record alternatively spliced isoforms as independent transcripts. However, for annotation-free methods that start with short-read sequencing, it is difficult to accurately assemble, model, and quantify alternative splicing at the transcript level
Constitutive Exons
Alternative Exons
Complex transcript-centric splice graph
1
2
3
4
5
6
7
8
9
10
11
12
13
Imperfect transcript assembly
B
ASCOT uses an exon-centric approach and only considers local pieces of the splice graph.
Exon-centric splice graph reduction
Alternative Splice Site Group
Mutually Exclusive Exons
Cassette Exon
Linked Exons

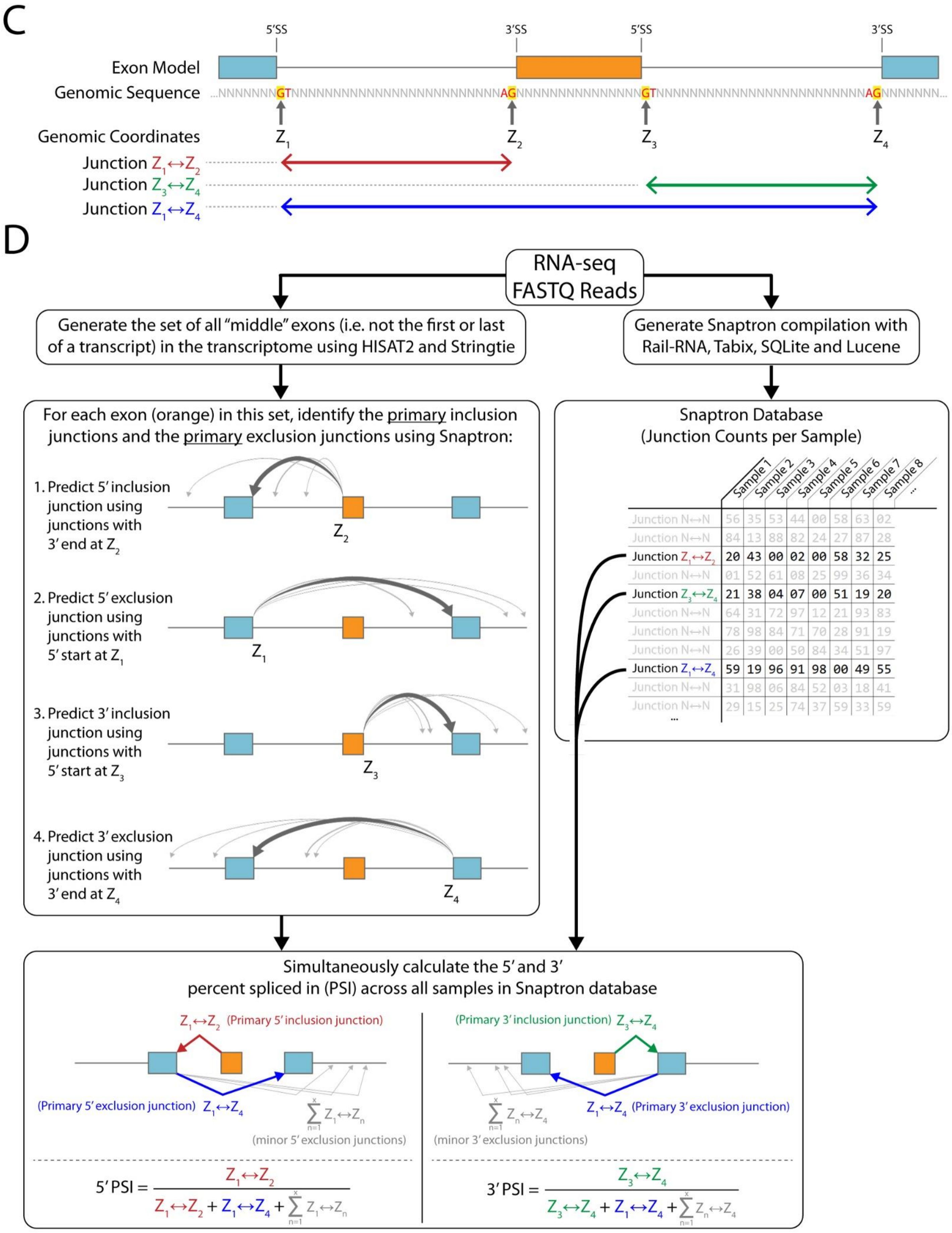
C
5'SS
3'SS
5'SS
3'SS
Exon Model
Genomic Sequence
...NNNNNNNGTNNNNNNNNNNNNNNNNNNNNNNNNNNNAGNNNNNNNNNNNNNNNGTNNNNNNNNNNNNNNNNNNNNNNNNNNNAGNNNNNNN...
Genomic Coordinates
Z1
Z2
Z3
Z4
Junction Z1↔Z2
Junction Z3↔Z4
Junction Z1↔Z4
D
RNA-seq FASTQ Reads
Generate the set of all "middle" exons (i.e. not the first or last of a transcript) in the transcriptome using HISAT2 and Stringtie
Generate Snaptron compilation with Rail-RNA, Tabix, SQLite and Lucene
For each exon (orange) in this set, identify the primary inclusion junctions and the primary exclusion junctions using Snaptron:
1. Predict 5' inclusion junction using junctions with 3' end at Z2
2. Predict 5' exclusion junction using junctions with 5' start at Z1
3. Predict 3' inclusion junction using junctions with 5' start at Z3
4. Predict 3' exclusion junction using junctions with 3' end at Z4
Snaptron Database (Junction Counts per Sample)
Sample 1
Sample 2
Sample 3
Sample 4
Sample 5
Sample 6
Sample 7
Sample 8
...
Junction N↔N 56 35 53 44 00 58 63 02
Junction N↔N 84 13 88 82 24 27 87 28
Junction Z1↔Z2 20 43 00 02 00 58 32 25
Junction N↔N 01 52 61 08 25 99 36 34
Junction Z3↔Z4 21 38 04 07 00 51 19 20
Junction N↔N 64 31 72 97 12 21 93 83
Junction N↔N 78 98 84 71 70 28 91 19
Junction N↔N 26 39 00 50 84 34 51 97
Junction Z1↔Z4 59 19 96 91 98 00 49 55
Junction N↔N 31 98 06 84 52 03 18 41
Junction N↔N 29 15 25 74 37 59 33 59
...
Simultaneously calculate the 5' and 3' percent spliced in (PSI) across all samples in Snaptron database
Z1↔Z2 (Primary 5' inclusion junction)
(Primary 5' exclusion junction) Z1↔Z4
$\sum_{n=1}^{x} Z_1 \leftrightarrow Z_n$
(minor 5' exclusion junctions)
(Primary 3' inclusion junction) Z3↔Z4
Z1↔Z4 (Primary 3' exclusion junction)
$\sum_{n=1}^{x} Z_n \leftrightarrow Z_4$
(minor 3' exclusion junctions)
$5'\ PSI = \frac{Z_1 \leftrightarrow Z_2}{Z_1 \leftrightarrow Z_2 + Z_1 \leftrightarrow Z_4 + \sum_{n=1}^{x} Z_1 \leftrightarrow Z_n}$
$3'\ PSI = \frac{Z_3 \leftrightarrow Z_4}{Z_3 \leftrightarrow Z_4 + Z_1 \leftrightarrow Z_4 + \sum_{n=1}^{x} Z_n \leftrightarrow Z_4}$

E

#### Classification Conditions

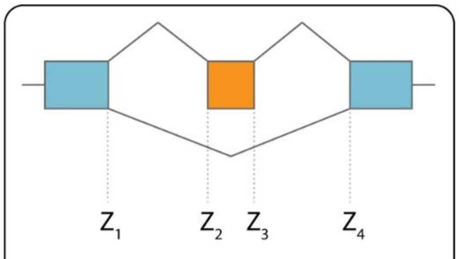

Cassette Exon

A cassette is defined as an exon that has equivalent 5′ and 3′ exclusion junctions ( $Z_1$-$Z_4$ )

Junction walking:
$Z_2 \rightarrow Z_1 \rightarrow Z_4$ and $Z_3 \rightarrow Z_4 \rightarrow Z_1$

---

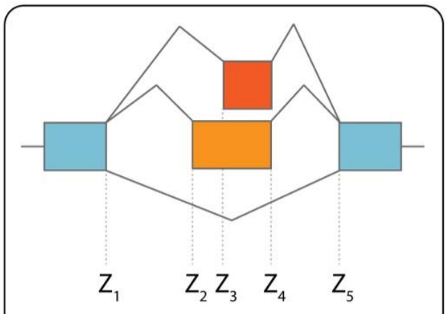

Alternative Splice Site Group

A group of exons define an alternative splite site exon group if they share the same 3′ inclusion junction ( $Z_4$-$Z_5$ ) and have equivalent 5′ and 3′ exclusion junctions ( $Z_1$-$Z_5$ )

Junction walking:
$Z_3 \rightarrow Z_1 \rightarrow Z_5$ and $Z_4 \rightarrow Z_5 \rightarrow Z_1$
$Z_2 \rightarrow Z_1 \rightarrow Z_5$ and $Z_4 \rightarrow Z_5 \rightarrow Z_1$

---

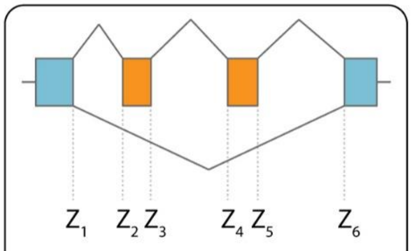

Linked Exons

Exons are linked if they share equivalent 5′ and 3′ exclusion junctions ( $Z_1$-$Z_6$ ) but have inclusion junctions that end between $Z_3$ and $Z_4$

Junction walking:
$Z_2 \rightarrow Z_1 \rightarrow Z_6$ and $Z_5 \rightarrow Z_6 \rightarrow Z_1$
$Z_3 \rightarrow Z_4$ and $Z_4 \rightarrow Z_3$ , otherwise complex splicing between $Z_2$ and $Z_5$

---

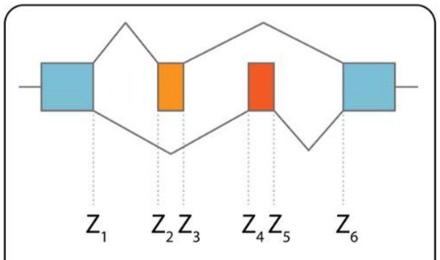

Mutually Exclusive Exons

Exons are mutually exclusive if they have high confidence inclusion junctions that extend to the same boundary splice sites ( $Z_1$ and $Z_6$ )

Junction walking:
$Z_2 \rightarrow Z_1$ and $Z_4 \rightarrow Z_1$
$Z_3 \rightarrow Z_6$ and $Z_5 \rightarrow Z_6$

**Supplemental Figure 1. Diagram of "junction-walking" strategy for identifying alternative splicing patterns using a splice junction count table**.

**(A)** Alternative splicing introduces a large amount transcriptomic and proteomic diversity. Most exons can be generally classified as either <u>constitutive</u> (spliced into mRNA for nearly all cellular environments) or <u>alternative</u> (spliced in only under certain conditions). Constitutive exons are labeled blue and alternative exons are labeled as shades of orange. Complex splice graphs can be derived from split-reads and a variety of mRNA transcripts are generated from these graphs. When analyzing alternative splicing with a gene reference file, spliced isoforms are recorded and quantified as independent transcripts. However, it is difficult for annotation-free methods to accurately assemble, model, and quantify alternative splicing with short-read sequencing. This is especially true for transcripts with distant alternative exons. Low abundance, spurious isoforms can also be reported as alternatively spliced and it is often difficult for biologists to interpret which transcripts are biologically relevant.

**(B)** ASCOT uses an exon-centric approach to reduce this complexity by only considering local regions of the splice graph and analyzing these elements independently from one another. In this work, we focus on four binary splicing decisions: cassette exons, alternative splice site exon groups that share the same exclusion junction, linked exons, and mutually exclusive pairs of exons.

**(C)** Our method for splicing analysis relies on evidence from RNA-Seq split-read alignments (i.e. splice junctions), as opposed to coverage. Split-reads are generally very reliable measurements, since sequences that contain two different regions of the genome are likely to have been joined via the spliceosome. Split-read alignment accuracy also benefits from requiring known splice motifs at either end of an intron. We can therefore easily identify which sets of junctions share the same start or end coordinates. In this diagram, inclusion junctions are shown in red ( $Z_1 \leftrightarrow Z_2$ ) and green ( $Z_3 \leftrightarrow Z_4$ ) while the exclusion junction is shown in blue ( $Z_1 \leftrightarrow Z_4$ ). For these inclusion junctions, $Z_2$ and $Z_3$ are adjacent to the alternative exon (orange), but no information links $Z_1$ and $Z_4$ unless we know that $Z_1 \leftrightarrow Z_4$ is the exclusion junction. This forms a closed loop since we can start from any coordinate and trace a path through an alternating series of exons and introns that leads to the starting coordinate, e.g. start at $Z_1$, follow the 5' inclusion junction to get to $Z_2$, follow the alternative exon to get to $Z_3$, follow the 3' inclusion junction to get to $Z_4$, then follow the exclusion junction to get back to $Z_1$. If identical loops are derived from the 5' and 3' inclusion junctions, there is high confidence that the alternative exon is a cassette exon. We developed a "junction-walking" strategy to identify these closed junction loops both across the transcriptome and across tens of thousands of summarized RNA-Seq samples by using the coordinates and count tables of a Snaptron[1] splice junction database. Because we use Snaptron queries in this process, the strategy is informed by junction evidence in all samples, and not just a single simple.

**(D)** The flowchart of our method begins with a set of raw RNA-Seq fastq reads. First, all the raw reads from all the input accessions were analyzed using the Rail-RNA spliced alignment program. Rail-RNA outputs a few summaries, the relevant here being a table of splice-junction evidence. In this table, each row is a splice junction and each column is an individual run accession. The elements of the table give the number of times a spliced alignment from an individual (column) spanned a junction (row). This summary was then indexed using Tabix and SQLite, and all the associated metadata for the run accessions was indexed using Lucene, to form a complete Snaptron compilation, a queryable database. In parallel with this process, we also analyzed every sample of raw RNA-SEQ Fastq reads with StringTie and identified the set of all "middle" exons by excluding all first and last exons. In theory, this set of middle exons contains the coordinates for all possible internal alternative exons that would be suitable queries for our junction-walking strategy. Next, for each middle exon, we predict the 5' and 3' inclusion junctions by querying Snaptron for all junctions ending at $Z_2$ (for the 5' inclusion junction) or beginning at $Z_3$ (for the 3' inclusion junction). We must also establish that if this is an alternative exon, there is only one primary 5' or 3' inclusion junction. For the 5' inclusion junction, we set a threshold where the primary inclusion junction must represent >70% of all junction counts (across all samples) that end at $Z_2$; a similar threshold is set for the 3' inclusion junction and junctions that begin at $Z_3$. We then calculate the primary 5' and 3' exclusion junctions by identifying all junctions that begin at $Z_1$ (for the 5' exclusion junction) and all junctions that end at $Z_4$ (for the 3' exclusion junction). We then test whether a junction loop has been closed using logic outlined in section (**E**). For exons that have consistent 5' and 3' closed junction loops, we can then easily calculate 5' and 3' percent spliced-in (PSI) values for each alternative exon with Snaptron per the formulas shown in the figure. For example, for the 5'PSI, $Z_1 \leftrightarrow Z_2$ represents a vector of 5' inclusion junction counts for each sample in the Snaptron database. Similarly, $Z_1 \leftrightarrow Z_4$ represents an equivalent count vector for the exclusion junction, and the sum of $Z_1 \leftrightarrow Z_n$ represents the sum of vectors for every other possible exclusion junction that begins at $Z_1$. This sum of minor exclusion junctions is important to ensure an accurate PSI calculation. Equivalent calculations are also performed to find the 3' PSI for each sample.

**(E)** In this work, we identify four closed junction loops that are essentially decisions between two possible splicing outcomes (binary decisions): 1. Cassette Exons, 2. Alternative Splice Site Exon

Groups, 3. Linked Exons, and 4. Mutually Exclusive Exons. In the simplest case, cassette exons are identified when the primary 5' and 3' exclusion junctions are identical. However, in some cases multiple exons can share the exclusion junction but possess alternative 5' or 3' splice sites. By treating these exons as a group, we can again model the splicing as a binary decision: either using any of the exons or using none. Further analysis can determine which specific splice sites are used within the group, but for the current work we focus on the primary splicing decision because measuring alternative splice site usage within this exon group would be treated as a nested decision. We also identify linked exons by finding an exon pair where the 5' exon's 5' exclusion junction is identical the 3' exon's 3' exclusion junction. It is important to note that our junction walking strategy is unable to determine what lies in between the exons. Given the possibility that nested splicing decisions exists between $Z_3$ and $Z_4$, we do not currently analyze splicing between linked exons. Finally, we also identify mutually exclusive exons by finding exon pairs where both exons inclusion junctions terminate at the same coordinates, but the 5' exon's inclusion junctions are the 3' exon's exclusion junctions and vice versa.

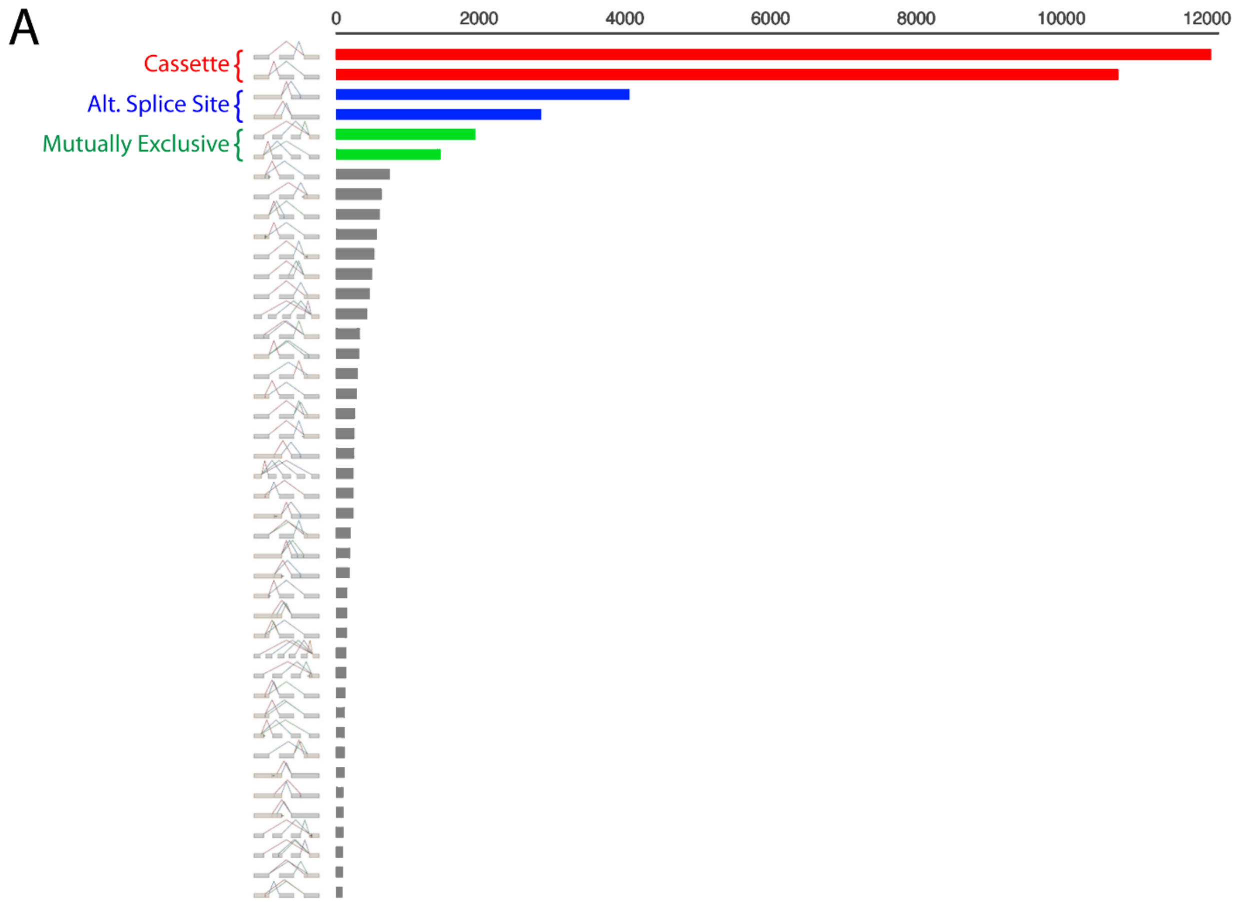


**Supplemental Figure 2. Binary decision splicing events represent most local splicing variation.** (**A**) Exonic splicing variation across the genome as detected by MAJIQ, adapted from Figure 4 of Vaquero-Garcia et al.[2] Simple alternative splicing decisions such as cassette exons, alternative splice sites, and mutual exclusive exons represent most local splicing variation.

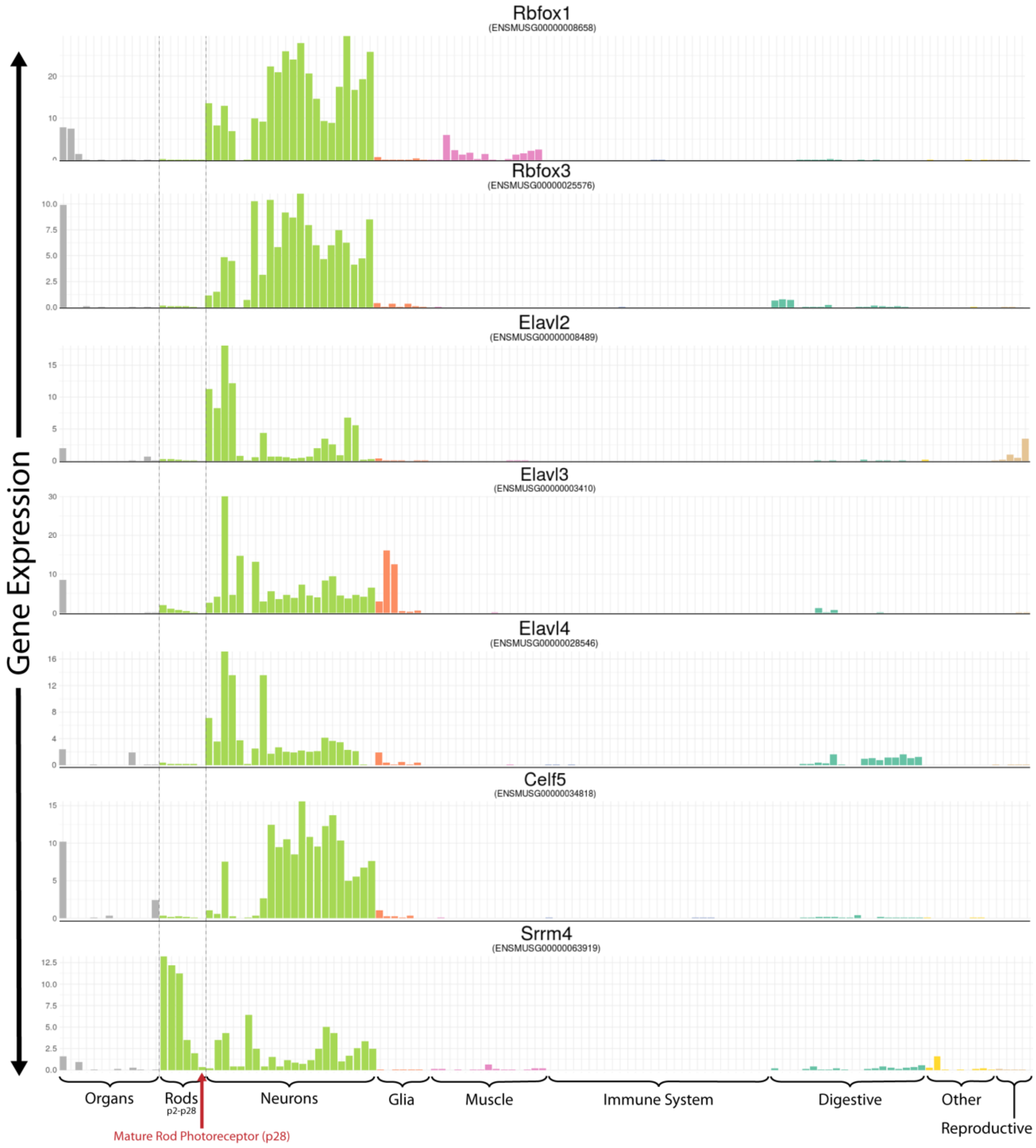


**Supplemental Figure 3. Rods do not express many of the common neuronal splicing factors**. *Rbfox1*, *Rbfox3*, *Elavl2*, *Elavl3*, *Elavl4*, *Celf5*, and *Srrm4* are splicing factors that are selectively expressed in most neurons. Rod photoreceptors, however, do not express these genes.

SFig4, pg1

| Gene Symbol | Exon Location (mm10, mouse) | Brain | Skeletal Muscle | Heart | Lung | Stomach | Spleen | Colon | Liver | Kidney | Pancreas islet | Mammary gland | Ovary | Testes | Rod Photoreceptor (p28) | SST interneurons (cortex) | PV interneurons (hippocampus) | Motor neurons (Hb9+, embryo) | DRG neurons | Olfactory sensory neurons (Omp+) | Hair cells (Atoh1+, cochlea) | Purkinje cells (cerebellum) | Pyramidal neurons (Thy1+, cortex) | Pyramidal neurons (hippo - proximal CA1) | Pyramidal neurons (hippo - distal CA1) | Pyramidal neurons (hippo - superficial CA1) | Pyramidal neurons (hippo - dorsal CA1) | Pyramidal neurons (hippo - dorsal CA1, postsubiculum-projecting) | Pyramidal neurons (hippo - intermediate CA1) | Pyramidal neurons (hippo - ventral CA1) | Pyramidal neurons (hippo - ventral CA1, amygdala-projecting) | Pyramidal neurons (hippo - ventral CA1, nucleus accumbens-projecting | Pyramidal neurons (hippo - deep CA1) | Pyramidal neurons (hippo - dorsal CA3) | Pyramidal neurons (hippo - ventral CA3) | Granule cells (hippo - dorsal DG) | Granule cells (hippo - ventral DG) | Oligodendrocyte precursor cells (cortex) | Newly formed oligodendrocytes (cortex) | Myelinating oligodendrocytes (cortex) | Astrocytes (cortex) | Microglia (cortex) | Microglia (spinal cord) | Endothelial cells (cortex) | Quadriceps | Gastrocnemius | Heart ventricle (p90) | Hematopoietic stem cell (bone marrow) | Dendritic cells (spleen) | Neutrophil | Mast cell | NKT cell | NK cell | Macrophage | Eosinophils | B cell, spleen | CD4+ T-cell (Th17) | CD4+ T-cell (Th1) | CD4+ T-cell (Th2) | CD8+ T-cell | CD4+/CD8+ double positive T-cell | Gamma delta T-cell | Regulatory T-cell (Rag2+, spleen) | Hepatocyte (liver) | Liver sinusoidal endothelial cells (liver) | Kupffer cells (liver) | Hepatic stellate cells (liver) | Smooth muscle cells (colon) | Interstitial Cell of Cajal (colon) | Gastrointestinal PDGFRa+ cells (colon) | Gastrointestinal PDGFRa+ cells (colon mucosa) | Pancreatic alpha cells (7-16wks) | Pancreatic beta cells (7-16wks) | Alveolar type 1 (lung) | Podocytes (kidney) | Classical brown adipocyte | Inguinal white adipocyte | Sertoli cells (testis) | Leydig cells (testis) | Interstitial cells (testis) | Spermatogonial stem cells (testis) | Female germline stem cells (ovary) |
|---|---|---|---|---|---|---|---|---|---|---|---|---|---|---|---|---|---|---|---|---|---|---|---|---|---|---|---|---|---|---|---|---|---|---|---|---|---|---|---|---|---|---|---|---|---|---|---|---|---|---|---|---|---|---|---|---|---|---|---|---|---|---|---|---|---|---|---|---|---|---|---|---|---|---|---|---|---|---|---|---|---|---|
| Unc13b | chr4:43254307-43254345 | 0 | 0 | 0 | 0 | 0 | 0 | 0 | 0 | 0 | 0 | 0 | 0 | 0 | 82 | -1 | -1 | 0 | 0 | 0 | 0 | 0 | -1 | 0 | 0 | 0 | 0 | 0 | 0 | 0 | 0 | 0 | 0 | 0 | 0 | 0 | -1 | 0 | 0 | 0 | 1 | 0 | 0 | 0 | 0 | 0 | 0 | 0 | -1 | -1 | 0 | -1 | -1 | 0 | 0 | -1 | 0 | 0 | 0 | 0 | -1 | -1 | -1 | 0 | 0 | 0 | 0 | 0 | 0 | 0 | 0 | 0 | 0 | 0 | 0 | -1 | 0 | 0 | 0 | 0 | 0 | 0 |
| Adgrv1 | chr13:81505652-81505678 | 0 | -1 | -1 | -1 | -1 | -1 | -1 | -1 | 0 | 0 | -1 | 0 | -1 | 95 | -1 | -1 | 0 | -1 | -1 | 0 | -1 | -1 | -1 | -1 | -1 | -1 | -1 | -1 | -1 | -1 | -1 | -1 | -1 | -1 | -1 | -1 | 0 | 0 | -1 | -1 | -1 | -1 | -1 | -1 | -1 | -1 | -1 | -1 | -1 | -1 | -1 | -1 | -1 | -1 | -1 | -1 | -1 | -1 | -1 | -1 | -1 | -1 | -1 | -1 | -1 | -1 | -1 | 0 | -1 | -1 | 0 | 0 | -1 | 0 | -1 | -1 | 0 | 0 | 0 | 0 | 0 |
| Amph | chr13:19101902-19101910 | 0 | 0 | -1 | 0 | 0 | 0 | 0 | -1 | 0 | 0 | 0 | 0 | 0 | 96 | 0 | 0 | 0 | 0 | 0 | 0 | 0 | 0 | 0 | 0 | 0 | 0 | 0 | 0 | 0 | 0 | 0 | 0 | 0 | 0 | 0 | 0 | 0 | 0 | 0 | 0 | 0 | 0 | 0 | 0 | -1 | 0 | -1 | -1 | -1 | 0 | -1 | -1 | -1 | -1 | -1 | 0 | -1 | -1 | -1 | -1 | -1 | -1 | -1 | -1 | -1 | 0 | 0 | 0 | 0 | 0 | 0 | 0 | -1 | 0 | -1 | -1 | 0 | 0 | 0 | 0 | 0 |
| Hcn1 | chr13:117971926-117971974 | 0 | 0 | 0 | 0 | -1 | -1 | 0 | -1 | -1 | 0 | 0 | 0 | 0 | 65 | 0 | 0 | 0 | 0 | 0 | 0 | 0 | 1 | 0 | 0 | 0 | 0 | 0 | 0 | 0 | 0 | 0 | 0 | 0 | 0 | 0 | 0 | -1 | -1 | 0 | 0 | -1 | 0 | -1 | -1 | -1 | 0 | -1 | -1 | -1 | -1 | -1 | -1 | -1 | -1 | -1 | 0 | -1 | -1 | -1 | 0 | -1 | -1 | -1 | -1 | -1 | -1 | -1 | 0 | -1 | -1 | 0 | 0 | 0 | -1 | -1 | -1 | 0 | -1 | 0 | -1 | 0 |
| Doc2b | chr11:75785517-75785543 | 1 | 0 | -1 | -1 | -1 | 0 | 0 | 0 | 0 | 0 | 0 | 0 | 0 | 95 | -1 | -1 | 0 | 0 | -1 | 0 | 0 | -1 | 0 | 0 | 0 | 0 | 0 | 0 | 0 | 0 | 0 | 0 | -1 | 0 | 0 | 0 | 0 | -1 | -1 | -1 | 0 | -1 | -1 | 0 | -1 | -1 | -1 | -1 | -1 | -1 | -1 | -1 | -1 | -1 | -1 | -1 | -1 | -1 | -1 | -1 | -1 | -1 | -1 | -1 | -1 | -1 | 0 | 0 | 0 | -1 | 0 | 0 | -1 | 0 | -1 | -1 | 0 | 1 | 0 | 0 | 0 |
| Cep290 | chr10:100495641-100495661 | 0 | 0 | -1 | 0 | -1 | 0 | 0 | 0 | 0 | 0 | 0 | 0 | 0 | 99 | 0 | 0 | 0 | 0 | 0 | 0 | 1 | 0 | 0 | 0 | 0 | 0 | 0 | 0 | 0 | 0 | 0 | 0 | 0 | 2 | 0 | -1 | 0 | 0 | 0 | 0 | -1 | 1 | 0 | -1 | -1 | 0 | -1 | 0 | -1 | 0 | -1 | -1 | -1 | 0 | -1 | 0 | 0 | 0 | 0 | 0 | -1 | -1 | -1 | -1 | 0 | 0 | 0 | 0 | 0 | 0 | 0 | 0 | 0 | 0 | 0 | 0 | 0 | 0 | 0 | 0 | 0 |
| Atp1b2 | chr11:69601924-69602330 | 0 | 1 | 0 | 0 | 0 | 1 | 0 | -1 | 0 | 0 | 3 | 0 | 0 | 96 | 0 | 0 | 0 | 0 | 0 | 0 | 0 | 0 | 0 | 0 | 0 | 0 | 0 | 0 | 0 | 0 | 0 | 0 | 0 | 0 | 0 | 0 | 0 | 0 | 0 | 0 | 0 | 0 | 0 | 0 | 0 | 0 | 0 | -1 | -1 | 0 | -1 | -1 | -1 | -1 | -1 | 0 | 0 | 0 | -1 | 0 | -1 | -1 | -1 | 0 | -1 | 0 | 0 | 0 | 0 | 0 | 0 | 1 | 0 | 0 | -1 | 0 | 0 | 0 | 0 | 0 | 0 |
| Erp44 | chr4:48220044-48220100 | 0 | 0 | 0 | 0 | 0 | 0 | 0 | 0 | 0 | 0 | 0 | 0 | 0 | 37 | 0 | 0 | 1 | 0 | 2 | 0 | 0 | 0 | 0 | 0 | 0 | 0 | 0 | -1 | 0 | 0 | 0 | 0 | 0 | 0 | 0 | 0 | 0 | 0 | 0 | 0 | 0 | 0 | 0 | 0 | 0 | 0 | 0 | 0 | 0 | 0 | 0 | 0 | 0 | 0 | 0 | 0 | 0 | 0 | 0 | 0 | 0 | 0 | 0 | 0 | 0 | 0 | 0 | 0 | 0 | 0 | 0 | 0 | 0 | 0 | 0 | 0 | 0 | 0 | 0 | 0 | 0 |
| Man2a2 | chr7:80355988-80356062 | 0 | 0 | 0 | 0 | 0 | 0 | 0 | 0 | 0 | 0 | 0 | 0 | 0 | 83 | 0 | 0 | 0 | 0 | 0 | 0 | 0 | 0 | 0 | 0 | 0 | 0 | 0 | 0 | 0 | 0 | 0 | 0 | 0 | 0 | 0 | 0 | 0 | 0 | 0 | 0 | 0 | 0 | 0 | 0 | 0 | 0 | 0 | 0 | 0 | 0 | 0 | 0 | 0 | 0 | 0 | 0 | 0 | 0 | 0 | 0 | 0 | 0 | 0 | 0 | 0 | 0 | 0 | 0 | 0 | 0 | 0 | 0 | 0 | 0 | 0 | 0 | 0 | 0 | 0 | 0 | 0 |
| Dmd | chrX:84526910-84526945 | 0 | 0 | 0 | -1 | -1 | -1 | 0 | -1 | 0 | 0 | 0 | 0 | 0 | 94 | 0 | 0 | 0 | 0 | 8 | -1 | 0 | -1 | 0 | 0 | 0 | 0 | 0 | 0 | 0 | 0 | 0 | 0 | 0 | 0 | 0 | 0 | 0 | 0 | 0 | 0 | -1 | 0 | -1 | 0 | 0 | 0 | -1 | -1 | -1 | -1 | -1 | -1 | -1 | -1 | -1 | -1 | -1 | -1 | -1 | -1 | -1 | -1 | -1 | -1 | -1 | -1 | 0 | 0 | 0 | 0 | 0 | -1 | -1 | 0 | 0 | -1 | 0 | -1 | 0 | 0 | 0 |
| Cask | chrX:13524825-13524851 | 0 | 0 | 0 | 0 | 0 | 0 | 0 | 0 | 0 | 0 | 0 | 0 | 0 | 97 | 0 | 0 | 1 | 0 | 0 | 12 | 0 | 0 | 0 | 0 | 0 | 0 | 1 | 0 | 0 | 0 | 0 | 0 | 0 | 0 | 0 | 0 | 0 | 0 | 0 | 0 | 0 | 0 | 0 | 0 | 0 | 0 | 0 | 0 | 0 | 0 | 0 | -1 | 0 | 0 | 0 | 0 | 0 | 0 | 0 | 0 | 0 | 0 | 0 | 0 | 0 | 0 | 0 | 0 | 0 | 0 | 2 | 0 | 0 | 0 | 0 | 0 | 0 | 0 | 0 | 0 | 1 |
| Efr3a | chr15:65857901-65857951 | 0 | 0 | 0 | 0 | 0 | 0 | 0 | 0 | 0 | 0 | 0 | 1 | 0 | 95 | 0 | 0 | 0 | 0 | 0 | 0 | 0 | 0 | 0 | 0 | 1 | 0 | 0 | 0 | 0 | 0 | 0 | 0 | 0 | 0 | 0 | 0 | 0 | 0 | -1 | 3 | 0 | 0 | 0 | 0 | -1 | 0 | 0 | 0 | 0 | 0 | 0 | 0 | 0 | 0 | 0 | 0 | 0 | 0 | 0 | 0 | 0 | 0 | 0 | 0 | 0 | 0 | 0 | 0 | 0 | 0 | 0 | 0 | 0 | 0 | 0 | 0 | 0 | 1 | 0 | 0 | 1 |
| Prom1 | chr5:44012499-44012516 | 0 | 0 | -1 | 0 | 0 | -1 | 1 | 0 | 0 | 1 | 3 | 0 | 0 | 98 | -1 | -1 | 7 | 1 | 0 | 0 | 0 | -1 | -1 | -1 | -1 | -1 | -1 | -1 | -1 | -1 | -1 | -1 | -1 | -1 | -1 | -1 | 0 | 0 | 0 | 0 | 0 | 0 | 0 | 0 | 0 | 0 | -1 | -1 | 0 | 0 | -1 | -1 | -1 | -1 | -1 | -1 | -1 | -1 | -1 | -1 | -1 | 0 | -1 | 0 | -1 | 0 | -1 | 0 | 0 | 0 | 2 | 3 | 0 | 0 | -1 | -1 | 0 | 0 | 0 | 0 | 1 |
| Wdr47 | chr3:108633762-108633794 | 0 | 0 | 0 | 0 | -1 | 0 | 0 | 0 | 0 | 0 | 0 | 0 | 0 | 68 | 0 | 0 | 0 | 0 | 8 | 0 | 0 | 0 | 0 | 0 | 0 | 0 | 0 | 1 | 0 | 0 | 0 | 0 | 0 | 0 | 0 | 0 | 0 | 0 | 0 | 0 | 0 | 1 | 0 | 0 | 0 | 0 | 0 | 0 | 0 | 0 | 0 | 0 | 0 | 0 | 0 | 1 | 0 | 0 | 0 | 3 | 0 | 0 | 0 | 0 | 0 | 0 | 0 | 0 | 0 | 0 | 0 | 0 | 0 | 0 | 0 | 0 | 0 | 0 | 0 | 1 | 0 |
| Nap1l4 | chr7:143541579\|143541460-143541603 | 0 | 0 | 0 | 0 | 0 | 0 | 0 | 0 | 0 | 0 | 0 | 0 | 0 | 30 | 0 | 0 | 0 | 1 | 0 | 0 | 4 | 0 | 0 | 0 | 0 | 0 | 0 | 0 | 0 | 0 | 0 | -1 | 0 | 1 | 0 | 0 | 0 | 0 | 0 | 0 | 0 | 0 | 0 | 0 | 0 | 0 | 0 | 0 | 0 | 0 | 0 | 0 | 0 | 0 | 0 | 0 | 0 | 0 | 0 | 0 | 0 | 0 | 0 | 0 | 0 | 0 | 0 | 0 | 0 | 0 | 0 | 0 | 0 | 0 | 0 | 0 | 0 | 0 | 0 | 0 | 0 |
| Slc17a7 | chr7:45169511-45169585 | 0 | 0 | 0 | 0 | -1 | -1 | -1 | -1 | 0 | 0 | 0 | -1 | -1 | 85 | -1 | 0 | 0 | 1 | -1 | 0 | 0 | 0 | 0 | 2 | 0 | 0 | 0 | 0 | 1 | 0 | 0 | 0 | 0 | 0 | 0 | 1 | 0 | 0 | 0 | 0 | 0 | -1 | 0 | -1 | -1 | 3 | -1 | -1 | -1 | -1 | -1 | -1 | -1 | 0 | -1 | 0 | 0 | 0 | -1 | 0 | -1 | -1 | -1 | -1 | -1 | -1 | -1 | -1 | -1 | 0 | -1 | -1 | 0 | -1 | -1 | -1 | -1 | -1 | -1 | 2 | 0 |
| Ykt6 | chr11:5960427-5960441 | 2 | 0 | 0 | 0 | 0 | 0 | 0 | 0 | 0 | 0 | 0 | 0 | 0 | 44 | 0 | 0 | 1 | 2 | 0 | 0 | 3 | 0 | 0 | 0 | 0 | 2 | 0 | 0 | 0 | 0 | 0 | 0 | 4 | 2 | 0 | 0 | 0 | 0 | 0 | 0 | 0 | 0 | 0 | 0 | 0 | 0 | 0 | 0 | 0 | 0 | 0 | 0 | 0 | 0 | 0 | 0 | 0 | 0 | 0 | 0 | 0 | 0 | 0 | 0 | 0 | 0 | 0 | 0 | 0 | 0 | 0 | 0 | 0 | 0 | 0 | 0 | 0 | 0 | 0 | 0 | 0 |
| Slc4a7 | chr14:14747733-14747792 | 6 | 0 | 0 | 0 | 0 | 0 | 0 | 0 | 0 | 0 | 0 | 0 | 0 | 69 | -1 | -1 | 6 | 0 | 0 | 7 | 2 | 0 | 3 | 0 | 0 | 0 | 0 | 0 | 0 | 0 | 0 | 0 | 0 | 2 | 0 | 1 | 1 | 1 | 0 | 1 | 0 | 0 | 0 | 0 | 0 | 0 | 0 | 0 | 0 | 0 | 0 | 0 | 0 | 0 | 0 | 0 | 0 | 0 | 0 | 0 | 0 | 0 | 0 | 0 | -1 | 0 | 0 | 0 | 0 | 0 | 1 | 1 | 2 | 0 | 0 | 0 | 0 | 0 | 0 | 0 | 0 |
| Kctd5 | chr17:24053272-24053330 | 0 | 2 | 0 | 1 | 0 | 2 | 1 | 1 | 1 | 0 | 1 | 0 | 0 | 74 | -1 | 0 | 0 | 1 | 17 | 0 | 4 | 0 | 0 | 0 | 0 | 0 | 0 | 0 | 0 | 0 | -1 | -1 | -1 | 0 | 0 | 0 | 0 | 0 | 0 | 0 | 0 | 1 | 0 | 0 | 0 | 0 | 0 | 0 | 0 | 0 | 0 | 0 | 0 | 0 | 0 | 0 | 0 | 0 | 0 | 0 | 0 | 0 | 0 | 0 | 0 | 0 | 0 | 0 | 1 | 0 | 0 | 0 | 0 | 0 | 0 | 0 | 0 | 0 | 0 | 0 | 0 |
| Madd | chr2:91172327\|91170835-91172335 | 4 | 0 | 0 | -1 | 0 | 1 | 0 | 1 | 0 | 0 | 1 | 0 | 0 | 99 | -1 | 0 | 0 | 0 | 3 | 0 | 2 | 0 | 7 | 0 | 0 | 0 | 0 | 0 | 0 | 7 | 0 | 0 | 0 | 0 | 0 | -1 | 1 | 0 | -1 | 0 | 2 | 0 | 2 | 0 | 0 | 0 | -1 | 0 | 0 | 0 | 0 | 0 | 0 | 0 | 0 | 0 | 1 | 1 | 0 | 0 | 4 | 0 | 0 | 0 | 1 | -1 | 0 | 0 | 0 | 0 | 0 | 0 | 1 | 1 | -1 | -1 | 1 | 3 | 1 | 0 | 0 |
| Brms1l | chr12:55840469-55840494 | 0 | 0 | 0 | 0 | 0 | 0 | 1 | 0 | 0 | 0 | 0 | 0 | 0 | 63 | 0 | 0 | 0 | 0 | 16 | 0 | 2 | 0 | 0 | 0 | 0 | 0 | 13 | 0 | 0 | 0 | 0 | 0 | 0 | 0 | 0 | 0 | 0 | 0 | 0 | 0 | 0 | 1 | 0 | 0 | 0 | 0 | 0 | 0 | 0 | 0 | 0 | 0 | 0 | 0 | 0 | 0 | 1 | 0 | 0 | 0 | 0 | 0 | -1 | 0 | 0 | 0 | 0 | 0 | 0 | 0 | 0 | 0 | 0 | 0 | 0 | 0 | 0 | 0 | 0 | 0 | 0 |
| Nup85 | chr11:115572736-115572759 | 4 | 0 | 0 | 0 | 0 | 0 | 0 | 0 | 0 | 0 | 0 | 0 | 0 | 46 | 15 | 0 | 2 | 1 | 0 | 0 | 6 | -1 | 0 | 0 | 0 | 0 | 0 | 0 | 0 | 0 | 0 | 0 | 0 | 1 | 0 | 0 | 0 | 0 | 0 | 0 | 0 | 0 | 0 | 0 | 0 | 0 | 0 | 0 | 0 | 0 | 0 | 0 | 0 | 0 | 0 | 0 | 0 | 0 | 0 | 0 | 0 | 0 | 0 | 0 | 0 | 0 | 0 | 0 | 0 | 0 | 0 | 0 | 0 | 0 | 0 | 0 | 0 | 0 | 0 | 0 | 0 |
| Fn3krp | chr11:121423310-121423411 | 1 | 0 | 0 | 0 | 0 | 0 | 0 | 0 | 0 | 0 | 0 | 0 | 0 | 34 | 0 | 0 | 2 | 0 | 2 | 0 | 3 | 0 | 0 | 0 | 0 | 0 | 0 | -1 | 0 | 0 | 0 | 0 | 0 | 0 | 0 | 0 | 0 | 2 | 0 | 0 | 0 | 0 | 0 | 0 | -1 | 1 | 0 | 0 | -1 | -1 | 0 | 5 | 0 | 0 | 0 | 0 | 0 | 0 | -1 | 0 | 0 | -1 | -1 | 0 | 0 | 0 | 0 | 0 | 0 | 0 | 0 | 0 | 0 | 0 | 0 | 0 | 0 | 0 | 0 | 0 | 0 |
| Farp2 | chr1:93617298-93617318 | 2 | 2 | 0 | 0 | 0 | 0 | 2 | 2 | 0 | 1 | 0 | 0 | 0 | 99 | -1 | -1 | 0 | 1 | 21 | 0 | 0 | -1 | -1 | -1 | -1 | -1 | -1 | 0 | -1 | -1 | -1 | -1 | 0 | 0 | 0 | -1 | 0 | 0 | 0 | 0 | 0 | 0 | 0 | 0 | 0 | 3 | -1 | 0 | 0 | 0 | -1 | 0 | -1 | 0 | 0 | 0 | -1 | 0 | -1 | -1 | 0 | -1 | 4 | 0 | 0 | 0 | 0 | 0 | 0 | 0 | 0 | 0 | 0 | 1 | 0 | -1 | 0 | 0 | 0 | 0 | 0 |
| Rims2 | chr15:39586534-39586785 | 0 | -1 | -1 | -1 | -1 | -1 | -1 | -1 | 0 | 0 | 0 | -1 | 3 | 86 | 0 | 0 | 0 | 6 | 0 | 1 | 0 | -1 | -1 | 0 | 0 | 0 | 0 | 0 | 0 | 0 | 0 | 0 | 0 | 0 | 0 | 0 | 0 | 4 | 0 | 0 | 0 | 0 | -1 | 0 | -1 | -1 | -1 | -1 | -1 | -1 | -1 | -1 | -1 | -1 | -1 | 0 | -1 | 0 | 0 | -1 | -1 | -1 | -1 | 0 | -1 | -1 | -1 | -1 | -1 | -1 | 0 | 1 | 0 | 0 | -1 | -1 | 0 | 0 | 1 | 4 | 9 |
| Kif1b | chr4:149195473-149195499 | 0 | 1 | 0 | 0 | 0 | 0 | 0 | 0 | 0 | 0 | 0 | 0 | 0 | 86 | 0 | 0 | 2 | 0 | 0 | 1 | 0 | -1 | 1 | 1 | 0 | 1 | 0 | 6 | 0 | 0 | 0 | 0 | 0 | 0 | 5 | 21 | 0 | 0 | 0 | 0 | 0 | 0 | 0 | 3 | 1 | 2 | 0 | 0 | 0 | 0 | 0 | 0 | 0 | 0 | 0 | 1 | 0 | 0 | 0 | 0 | 0 | 0 | 0 | 0 | 0 | 0 | 2 | 0 | 0 | 1 | 1 | 0 | 0 | 0 | 0 | 1 | 0 | 0 | 0 | 0 | 0 |
| Camsap1 | chr2:25953915\|25953903-25953974 | 0 | 0 | 0 | 0 | 0 | 0 | 0 | 0 | 0 | 0 | 1 | 1 | 0 | 94 | -1 | 0 | 3 | 1 | 39 | 4 | 0 | 0 | 0 | 0 | 0 | 0 | 0 | 0 | 0 | 0 | 0 | 0 | 1 | 0 | 0 | 0 | 0 | 0 | 0 | 9 | 0 | 0 | 0 | 0 | 0 | 0 | 0 | 0 | 0 | 0 | 0 | 0 | 0 | 0 | -1 | 1 | 0 | 0 | 0 | 0 | 0 | 0 | 0 | 0 | -1 | 0 | 0 | 0 | 0 | 0 | 0 | 0 | 0 | 0 | 0 | 0 | 0 | 0 | 0 | 12 | 0 |
| Kif1b | chr4:149205686-149205886 | 1 | 1 | 11 | 0 | 0 | 0 | 0 | 0 | 0 | 0 | 0 | 2 | 4 | 73 | -1 | 0 | 1 | 0 | 2 | 0 | 0 | -1 | 0 | 0 | 0 | 0 | 0 | 1 | 0 | 0 | 0 | 0 | 0 | 0 | 0 | 0 | 2 | 2 | 1 | 4 | 0 | 0 | 0 | 0 | 0 | 2 | 0 | 0 | 0 | 1 | -1 | 0 | 0 | 0 | 0 | 0 | 0 | 0 | 0 | 0 | 0 | -1 | 0 | 0 | 0 | 0 | 0 | 0 | 0 | 0 | 0 | 0 | 0 | 0 | 0 | 0 | 0 | 0 | 0 | 0 | 5 |
| Amph | chr13:19118424-19118912\|19119875 | 1 | 0 | -1 | 0 | 0 | 0 | 0 | -1 | 0 | 0 | 0 | -1 | 1 | 93 | 8 | 12 | 0 | 1 | 11 | 0 | 1 | 0 | 0 | 3 | 1 | 1 | 0 | 0 | 0 | 0 | 1 | 10 | 1 | 0 | -1 | 0 | 1 | 1 | 0 | 4 | 0 | 0 | 0 | 0 | -1 | 0 | -1 | 0 | 0 | 0 | -1 | -1 | -1 | -1 | -1 | 0 | -1 | -1 | -1 | -1 | -1 | -1 | -1 | -1 | -1 | 0 | 0 | 0 | 0 | 0 | 0 | 0 | 0 | 0 | -1 | 0 | 0 | 0 | -1 | 0 | -1 |
| Actr3 | chr1:125408782-125408901 | 1 | 0 | 0 | 3 | 0 | 0 | 1 | 0 | 0 | 1 | -1 | 1 | 0 | 72 | 1 | 1 | 1 | 0 | 15 | 0 | 0 | 0 | 0 | 0 | 0 | 0 | 0 | 0 | 0 | 1 | 1 | 0 | 0 | 0 | 0 | 0 | 1 | 0 | 0 | 0 | 0 | -1 | 0 | -1 | 0 | 0 | 0 | -1 | -1 | 0 | -1 | -1 | 0 | 4 | -1 | 1 | 0 | 0 | 0 | 1 | 0 | 0 | 0 | -1 | -1 | 1 | 0 | 0 | 0 | 0 | -1 | -1 | -1 | 0 | 1 | 1 | 1 | 0 | 0 | 0 | 0 |
| Bdh1 | chr16:31445346-31445433 | 4 | 1 | 1 | 0 | 0 | 0 | 0 | 0 | 1 | 1 | 2 | 0 | 1 | 45 | 1 | -1 | 1 | 4 | 1 | 0 | 2 | 2 | 0 | 0 | 0 | -1 | 0 | 0 | 0 | 0 | 0 | 0 | 0 | 0 | -1 | -1 | 1 | 4 | 6 | 0 | 0 | 0 | 0 | 0 | 1 | 1 | -1 | 0 | -1 | -1 | 0 | 0 | -1 | 0 | 0 | 0 | 0 | 0 | 0 | 0 | 0 | 0 | 0 | 0 | 0 | 0 | 1 | 0 | 0 | 0 | 1 | 0 | 0 | 0 | -1 | 0 | 0 | 0 | 2 | 0 | 0 |
| Gss | chr2:155576955-155576971 | 1 | 1 | 0 | 0 | 1 | 1 | 0 | 0 | 0 | 1 | 1 | 0 | 0 | 26 | 0 | 0 | 4 | 0 | 1 | 0 | 0 | 0 | 0 | 0 | 0 | -1 | -1 | -1 | -1 | 0 | -1 | 0 | 0 | 0 | 0 | -1 | 0 | 1 | 0 | 0 | 0 | 0 | 0 | 0 | 0 | 1 | -1 | -1 | 0 | 0 | -1 | -1 | 0 | 0 | 0 | 1 | 0 | 1 | -1 | 1 | 0 | 0 | 0 | 0 | 0 | 0 | 0 | 1 | 0 | 0 | 0 | 1 | 0 | 1 | 0 | 0 | 0 | 0 | 0 | 0 | 3 |
| Cltc | chr11:86713425-86713436 | 1 | 2 | 1 | 1 | 0 | 1 | 3 | 1 | 1 | 0 | 1 | 0 | 1 | 47 | 0 | 3 | 1 | 1 | 0 | 3 | 1 | 0 | 0 | 0 | 0 | 1 | 10 | 1 | 1 | 0 | 0 | 2 | 0 | 0 | 0 | 0 | 0 | 0 | 0 | 0 | 0 | 0 | 0 | 0 | 0 | 0 | 0 | 1 | 0 | 0 | 0 | 1 | 0 | 0 | 0 | 0 | 0 | 0 | 1 | 0 | 0 | 0 | 0 | 0 | 0 | 0 | 0 | 0 | 0 | 0 | 1 | 1 | 0 | 0 | 0 | 2 | 0 | 0 | 0 | 0 | 0 |
| Gramd1b | chr9:40330585-40332683 | 2 | 1 | 1 | 0 | 0 | 1 | 0 | 0 | 0 | 0 | 1 | 0 | 0 | 97 | 2 | 3 | 1 | 29 | 0 | 0 | 31 | 0 | 0 | 0 | 0 | 0 | 0 | 0 | 1 | 0 | 0 | 0 | 1 | 2 | 0 | 1 | 0 | 0 | 0 | 0 | 0 | 1 | 0 | 0 | 0 | 0 | 0 | 0 | -1 | 1 | 2 | 0 | 0 | 0 | -1 | 0 | 1 | 1 | 0 | 0 | 0 | -1 | -1 | 0 | 0 | 0 | 0 | 0 | 0 | 1 | -1 | 0 | 0 | 0 | 0 | 0 | 0 | 0 | 0 | 0 | -1 |
| Ints13 | chr6:146561594-146561617 | 8 | 0 | 0 | 0 | 0 | 0 | 0 | 0 | 0 | 1 | 0 | 0 | 1 | 89 | 4 | -1 | 5 | 4 | 29 | 19 | 18 | 0 | -1 | -1 | 0 | 0 | 0 | 0 | 0 | 0 | -1 | 0 | 0 | -1 | 0 | 0 | 1 | 0 | 0 | 0 | 0 | 0 | 1 | 0 | 0 | 0 | 0 | 0 | 0 | 0 | 0 | 0 | 0 | 0 | 3 | 0 | 0 | 0 | 0 | 0 | 0 | 0 | 0 | 0 | 0 | 0 | 0 | 0 | 0 | 0 | 3 | 1 | 0 | 0 | 0 | 0 | 0 | 0 | 0 | 0 | 0 |
| Plekhg3 | chr12:76563500-76563521 | 2 | 0 | -1 | 0 | 0 | 0 | 0 | 0 | 0 | 2 | 0 | 0 | 0 | 99 | -1 | -1 | -1 | 46 | 0 | 0 | 16 | -1 | 0 | -1 | -1 | -1 | -1 | -1 | -1 | -1 | -1 | -1 | -1 | -1 | -1 | -1 | 0 | 0 | 0 | 0 | 0 | -1 | 0 | 0 | 0 | -1 | 0 | 0 | 0 | 0 | 0 | 0 | 0 | 0 | 0 | 0 | 0 | 0 | 0 | 0 | 0 | -1 | 0 | 0 | 0 | 0 | 0 | 0 | 0 | 0 | 3 | 2 | 0 | 0 | 0 | 0 | 0 | 0 | 0 | 0 | 0 |
| 2210016L21Rik | chr5:114946610-114946618 | 3 | 1 | 0 | 0 | 0 | 0 | 0 | 0 | 0 | 1 | 0 | 0 | 0 | 45 | 1 | 16 | 4 | 3 | 4 | 1 | 4 | 0 | 1 | 0 | 1 | 0 | 0 | 0 | 0 | 0 | 0 | 1 | 4 | 5 | 0 | 0 | 3 | 0 | 1 | 0 | 0 | 0 | 0 | 1 | 1 | 0 | -1 | 0 | 0 | 0 | 0 | 0 | 0 | 1 | 0 | 1 | 1 | 1 | 0 | 0 | 0 | 0 | 0 | 2 | 0 | 0 | 0 | 0 | 0 | 0 | 0 | 1 | 0 | 0 | 0 | 0 | 0 | 1 | 0 | 1 | 0 |
| Bsg | chr10:79708697-79709044 | 0 | 0 | 0 | 4 | 1 | 2 | 1 | 2 | 2 | 3 | 1 | 0 | 0 | 96 | 0 | 0 | 0 | 7 | 11 | 1 | 1 | 0 | 0 | 0 | 0 | 0 | 0 | 0 | 1 | 0 | 0 | 0 | 0 | 0 | 1 | 0 | 0 | 0 | 0 | 0 | 0 | 0 | 0 | 0 | 0 | 0 | 10 | 3 | 11 | 4 | 9 | 7 | 3 | 2 | 2 | 1 | 6 | 2 | 6 | 2 | 4 | -1 | 1 | 0 | 0 | 0 | 0 | 1 | 1 | 1 | 6 | 6 | 1 | 0 | 2 | 1 | 0 | 0 | 0 | 0 | 1 |
| Stxbp5 | chr10:9796938-9797021 | 3 | 0 | -1 | 0 | 0 | 0 | 0 | 0 | 1 | 0 | 0 | 0 | 11 | 52 | 0 | 0 | 30 | 3 | -1 | 0 | 2 | 0 | 0 | 0 | 0 | 0 | 0 | 1 | 1 | 0 | 0 | 0 | 0 | 0 | 0 | 0 | 1 | 34 | -1 | 1 | 0 | 0 | 0 | 0 | 0 | 3 | 0 | 0 | 0 | 0 | -1 | 0 | 0 | 0 | 0 | 1 | 0 | 0 | 0 | 0 | 0 | 0 | 0 | 0 | 0 | -1 | 0 | 1 | 0 | 0 | 0 | 0 | 0 | 0 | 0 | 0 | 3 | 0 | 1 | 2 | 4 |
| Ttc8 | chr12:98937120-98937149 | 3 | 6 | 0 | 0 | -1 | 0 | 0 | 3 | 1 | 2 | 0 | 8 | 0 | 99 | 3 | 1 | 3 | 1 | 90 | 0 | 5 | 0 | 1 | 1 | 2 | 0 | 3 | 0 | 0 | -1 | 0 | 0 | 0 | 1 | 0 | 2 | 2 | 0 | -1 | 1 | 0 | 3 | 2 | 0 | 0 | -1 | 0 | 0 | -1 | 0 | 0 | -1 | -1 | 0 | -1 | 0 | 0 | 0 | -1 | -1 | -1 | 0 | 0 | 0 | 0 | 0 | 0 | 1 | 0 | 0 | 4 | 3 | 0 | 1 | 0 | 0 | 0 | 1 | 1 | 0 | 0 |
| Cc2d2a | chr5:43728271-43728291 | 12 | 38 | 0 | 0 | 0 | 0 | 0 | 0 | 0 | 0 | 0 | 0 | -1 | 99 | 0 | -1 | 0 | 4 | 58 | 0 | 18 | 0 | 0 | 0 | 0 | 0 | 0 | 0 | 0 | 0 | 0 | 0 | 0 | -1 | 5 | 0 | 0 | 0 | 0 | 2 | 0 | 0 | 0 | 26 | -1 | 13 | -1 | 0 | -1 | 0 | -1 | -1 | 0 | -1 | -1 | 0 | -1 | -1 | -1 | -1 | -1 | -1 | 0 | 0 | -1 | 0 | 0 | 0 | 0 | 0 | 0 | 0 | 0 | 0 | -1 | 0 | 0 | 0 | 0 | 0 | 0 |
| Ttc7b | chr12:100445956-100445979\|100445974 | 9 | 0 | 1 | 1 | 0 | 0 | 0 | 0 | 0 | 0 | 0 | 0 | 0 | 92 | -1 | 54 | 0 | 37 | 38 | 0 | 33 | -1 | 0 | 0 | 0 | 1 | 0 | 2 | 8 | 0 | 0 | 2 | 1 | 1 | 5 | 0 | 0 | 0 | 0 | 0 | 0 | -1 | 0 | 0 | 0 | 1 | 1 | 0 | 1 | 0 | 0 | 0 | 0 | 0 | 0 | 0 | 0 | 0 | 0 | 1 | 0 | 0 | 0 | 0 | 0 | 0 | 0 | 0 | 1 | 0 | 1 | 1 | 0 | 0 | 0 | 0 | 0 | 0 | 0 | 0 | 1 |
| Impdh1 | chr6:29201888-29201904 | 4 | 0 | 0 | 0 | 0 | 1 | 1 | 0 | 2 | 4 | 0 | 0 | 0 | 96 | 23 | -1 | 4 | 7 | 82 | 12 | 11 | 0 | 0 | 0 | 0 | 0 | 0 | 0 | 0 | 0 | 0 | -1 | -1 | 0 | 0 | 0 | 0 | 0 | 0 | 0 | 0 | 0 | 0 | 0 | 0 | 0 | 3 | 0 | 0 | 2 | 0 | 0 | 0 | 0 | 0 | 0 | 0 | 1 | 1 | 2 | 0 | 0 | 0 | 0 | 0 | 0 | 1 | 3 | 0 | 1 | 13 | 12 | 1 | 1 | 0 | 0 | 0 | 0 | 0 | 1 | 0 |
| Msi2 | chr11:88368212-88368235 | 14 | 1 | 1 | 2 | 0 | 0 | 0 | 0 | 0 | 2 | 1 | 1 | 0 | 93 | -1 | 1 | 1 | 4 | 95 | 12 | 4 | 0 | 0 | 0 | -1 | 0 | 0 | -1 | 0 | 0 | 0 | -1 | 0 | 3 | 0 | 0 | 2 | 0 | 4 | 8 | 3 | -1 | 2 | 1 | 3 | 1 | 9 | 0 | 0 | 0 | 0 | 0 | -1 | 0 | 0 | 0 | 0 | 0 | 0 | 0 | 0 | 0 | 0 | 0 | 0 | 0 | 0 | 2 | 0 | 0 | 4 | 1 | 0 | 0 | 0 | 0 | 0 | 1 | 0 | 2 | 0 |
| 5530601H04Rik | chrX:105069114-105069146 | 2 | 0 | 0 | 0 | 0 | 0 | 0 | 0 | 0 | 0 | 0 | 0 | 0 | 40 | 17 | 0 | 1 | 5 | 16 | 0 | 21 | 9 | 2 | -1 | -1 | 0 | -1 | 0 | 0 | 0 | 0 | -1 | -1 | -1 | 0 | 0 | 0 | 4 | 1 | 0 | 0 | 1 | 0 | 9 | 0 | 4 | 0 | 0 | 0 | 0 | 0 | 0 | 0 | 0 | 0 | 0 | 0 | 0 | 0 | 0 | 0 | 0 | 0 | 0 | 0 | 0 | 0 | 0 | 0 | 0 | 1 | 0 | 0 | 0 | 0 | 0 | 0 | 1 | 0 | 0 | 0 |
| Gpbp1l1 | chr4:116578193-116578261 | 1 | 0 | 0 | 1 | 0 | 1 | 0 | 1 | 0 | 0 | 0 | 0 | 0 | 21 | 0 | 0 | 3 | 0 | 1 | 2 | 1 | -1 | 0 | 0 | 0 | 0 | 0 | 1 | 0 | 0 | 0 | 0 | 9 | 0 | 4 | 0 | 0 | 0 | 0 | 2 | 0 | 0 | 1 | 0 | 0 | 0 | 0 | 0 | 2 | 1 | 0 | 0 | 0 | 2 | 2 | 1 | 2 | 1 | 0 | 1 | 0 | 2 | 0 | 0 | 0 | 1 | 0 | 0 | 1 | 0 | 0 | 0 | 0 | 0 | 0 | 0 | 0 | 1 | 1 | 1 | 0 |
| Coa7 | chr4:108329188-108329218 | 1 | 5 | 0 | 0 | 0 | 0 | 2 | 1 | 0 | 1 | 2 | 0 | 2 | 33 | 7 | 0 | 1 | 0 | 1 | 0 | 0 | 0 | 0 | 0 | 0 | 0 | 0 | 0 | -1 | -1 | 0 | -1 | 0 | 0 | 0 | 0 | 0 | 0 | 0 | 1 | 4 | 1 | 1 | 4 | -1 | 1 | 0 | -1 | 0 | 0 | 0 | 0 | 0 | 0 | 2 | 1 | 1 | 2 | 0 | 4 | 0 | 0 | 0 | 0 | -1 | 0 | 4 | 0 | 0 | 0 | 3 | 1 | 3 | 1 | 0 | 0 | 0 | 1 | 2 | 2 | 1 |
| Atp11a | chr8:12851633-12851656 | 6 | 3 | 0 | 0 | 0 | 1 | -1 | 2 | 1 | 1 | 0 | 2 | 0 | 99 | -1 | -1 | 4 | 2 | 9 | 56 | 52 | -1 | -1 | -1 | -1 | -1 | -1 | -1 | -1 | -1 | 0 | -1 | -1 | -1 | -1 | 0 | 5 | 2 | 5 | 3 | 0 | 3 | 2 | 0 | 2 | 0 | 0 | 0 | 0 | 2 | 3 | 0 | 0 | 2 | 0 | 0 | 1 | 1 | 0 | 1 | 0 | -1 | -1 | 0 | 0 | 0 | 0 | 1 | 2 | 1 | 0 | 1 | 0 | 1 | 0 | 3 | 1 | 1 | 1 | 0 | 0 |
| Kmt2d | chr15:98836581-98836628 | 0 | 0 | 0 | 2 | 0 | 2 | 4 | 2 | 1 | 2 | 2 | 8 | 32 | 59 | 0 | 0 | 0 | 1 | 0 | 8 | 0 | -1 | 0 | 0 | 0 | 0 | 0 | 0 | 0 | 0 | -1 | 0 | 0 | 0 | 0 | 0 | 1 | 2 | 0 | 2 | 0 | 0 | 1 | 0 | 0 | 0 | 3 | 3 | 1 | 1 | 4 | 0 | 0 | 2 | 2 | 3 | 1 | 1 | 0 | 1 | 3 | 0 | 3 | 0 | 1 | 0 | 1 | 1 | 0 | 0 | 1 | 1 | 1 | 1 | 0 | 0 | 3 | 3 | 2 | 2 | 11 |
| Taf1 | chrX:101540555-101540569 | 1 | 0 | 0 | -1 | -1 | 5 | -1 | 0 | -1 | 1 | 3 | 0 | 0 | 77 | 0 | 0 | 0 | 2 | 37 | 0 | 1 | 0 | 0 | 0 | 0 | 0 | 0 | 0 | 0 | 0 | 0 | -1 | 4 | 0 | 0 | 0 | 2 | 1 | -1 | 4 | -1 | 3 | 0 | -1 | 0 | 2 | -1 | 0 | -1 | -1 | 0 | 5 | -1 | -1 | 1 | 3 | 10 | 10 | 3 | 4 | -1 | 0 | -1 | 0 | 0 | 0 | 0 | 3 | -1 | 0 | 0 | 2 | 0 | 3 | -1 | 4 | 0 | 0 | 0 | 8 | 0 |
| Cc2d2a | chr5:43716937-43716951 | 12 | 7 | 0 | 0 | -1 | 0 | 0 | 0 | 0 | 0 | 0 | 0 | 0 | 97 | 0 | 0 | -1 | 0 | 0 | 0 | 12 | 0 | 0 | 13 | 27 | 28 | 0 | -1 | -1 | 22 | -1 | 54 | 17 | 2 | 24 | 0 | 1 | 1 | 0 | 0 | 0 | 0 | 0 | 5 | 0 | 1 | 0 | -1 | -1 | -1 | -1 | -1 | 0 | -1 | -1 | -1 | -1 | -1 | -1 | -1 | -1 | -1 | -1 | 0 | 0 | 0 | 0 | 0 | 0 | 0 | 1 | 0 | 0 | 0 | 0 | 0 | 1 | 0 | 0 | 0 | 0 |
| Unc13b | chr4:43169968\|43166668-43178334 | 0 | 0 | 0 | 0 | 0 | 0 | 0 | 0 | 0 | 2 | 2 | -1 | -1 | 94 | -1 | -1 | 1 | 5 | 10 | 1 | -1 | -1 | -1 | -1 | -1 | -1 | -1 | -1 | -1 | -1 | -1 | -1 | -1 | -1 | -1 | -1 | 0 | -1 | -1 | 0 | -1 | 0 | 4 | -1 | 0 | -1 | 0 | 0 | -1 | -1 | -1 | -1 | 0 | 0 | -1 | -1 | -1 | -1 | -1 | -1 | -1 | -1 | -1 | -1 | -1 | -1 | -1 | -1 | -1 | 1 | 7 | 5 | 3 | 6 | 0 | 0 | 2 | 1 | 1 | 2 | 14 |
| Rnf145 | chr11:44562074-44562082 | 5 | 2 | 0 | 1 | 0 | 2 | 1 | 1 | 2 | 0 | 1 | 0 | 2 | 22 | 0 | 6 | 0 | 4 | 0 | 0 | 4 | 0 | 2 | 2 | 0 | 2 | 0 | 0 | 0 | 0 | -1 | 2 | 3 | 0 | 0 | -1 | 0 | 0 | 0 | 0 | 1 | 1 | 1 | 0 | 0 | 0 | 0 | 1 | 1 | 0 | 4 | 2 | 4 | 0 | 1 | 1 | 0 | 0 | 3 | 1 | 1 | 0 | 0 | 0 | 0 | 0 | 0 | 0 | 0 | 0 | 1 | 1 | 0 | 0 | 0 | 0 | 0 | 0 | 0 | 0 | 0 |
| March1 | chr8:66386809-66387561 | 3 | -1 | -1 | 8 | 0 | 10 | -1 | -1 | -1 | 4 | -1 | -1 | -1 | 83 | -1 | -1 | 1 | -1 | -1 | -1 | -1 | 0 | 0 | 1 | 2 | 1 | 1 | 0 | 0 | 0 | -1 | -1 | 9 | 3 | 0 | 0 | 1 | -1 | 0 | -1 | 17 | 6 | -1 | 7 | 0 | -1 | -1 | 6 | 10 | -1 | 0 | -1 | 0 | 3 | 8 | -1 | -1 | -1 | -1 | 0 | -1 | -1 | -1 | 0 | 2 | 4 | -1 | 0 | -1 | -1 | 0 | -1 | 0 | -1 | 0 | 0 | -1 | 0 | 0 | 0 | -1 |
| Gm42902 | chr5:107396266-107396362 | 0 | 0 | -1 | 0 | -1 | -1 | -1 | -1 | -1 | 0 | -1 | 0 | 1 | -1 | 31 | 0 | 0 | 0 | 0 | 0 | 0 | 0 | 0 | 0 | 0 | 0 | 0 | 0 | 0 | 0 | 0 | 0 | 0 | 1 | 0 | 0 | 0 | 0 | 0 | -1 | -1 | 0 | 0 | -1 | -1 | -1 | -1 | -1 | -1 | 0 | -1 | -1 | -1 | -1 | -1 | 0 | 0 | -1 | -1 | 0 | -1 | -1 | -1 | -1 | -1 | -1 | -1 | 0 | -1 | -1 | 0 | 0 | 0 | 0 | -1 | -1 | 0 | 0 | -1 | 0 | 0 |
| Gabbr2 | chr4:46734882-46735017 | 0 | 0 | -1 | -1 | -1 | -1 | -1 | 0 | -1 | 0 | 0 | -1 | 0 | 0 | 30 | 0 | 0 | 0 | 0 | -1 | 0 | 0 | 0 | 1 | 0 | 1 | 0 | 1 | 0 | 0 | 0 | 0 | 0 | 0 | 0 | 0 | 0 | 0 | 0 | 0 | 0 | 0 | 0 | -1 | -1 | -1 | -1 | -1 | 0 | -1 | -1 | -1 | -1 | -1 | -1 | -1 | -1 | -1 | -1 | 0 | -1 | -1 | 0 | -1 | -1 | 0 | 0 | 0 | -1 | -1 | 0 | 0 | -1 | -1 | 0 | 0 | 0 | 0 | -1 | -1 | -1 |
| Mir99ahg | chr16:77686265-77686404 | 0 | 2 | 0 | -1 | -1 | -1 | 0 | -1 | 0 | -1 | 0 | 0 | 2 | -1 | 51 | -1 | 0 | 0 | -1 | 0 | 0 | 0 | 0 | -1 | -1 | -1 | -1 | 0 | -1 | -1 | -1 | -1 | 0 | 0 | -1 | -1 | -1 | 0 | 0 | 4 | -1 | 0 | 0 | -1 | 0 | 6 | -1 | -1 | -1 | -1 | -1 | -1 | 0 | -1 | -1 | -1 | -1 | -1 | -1 | -1 | -1 | -1 | -1 | -1 | -1 | 0 | -1 | 0 | -1 | 0 | 0 | -1 | 0 | 0 | 0 | 0 | -1 | 0 | 0 | 0 | -1 |
| B130034C11Rik | chr16:87498972-87499017 | -1 | 0 | -1 | 0 | 0 | 0 | 0 | 0 | -1 | 8 | 0 | 0 | -1 | 0 | 37 | 0 | -1 | 0 | 1 | 1 | 1 | 0 | -1 | 0 | -1 | -1 | -1 | -1 | -1 | -1 | -1 | 0 | 0 | 0 | -1 | -1 | 0 | 0 | 0 | 0 | -1 | 0 | -1 | 0 | 0 | -1 | -1 | 0 | -1 | -1 | -1 | 0 | -1 | 0 | -1 | 0 | 0 | 0 | -1 | -1 | 0 | 0 | -1 | -1 | -1 | -1 | -1 | 0 | 0 | -1 | -1 | 0 | 0 | 0 | -1 | 0 | 0 | -1 | 0 | -1 | -1 |
| Ift88 | chr14:57495367-57495499 | 0 | 1 | 0 | 6 | 0 | 0 | 1 | 0 | 1 | 0 | 0 | 1 | 1 | 0 | 32 | 0 | 0 | 0 | 0 | 0 | 0 | 0 | 0 | 0 | -1 | 0 | -1 | -1 | -1 | 0 | 0 | 0 | 0 | 0 | 0 | 0 | 0 | 3 | 0 | 0 | 0 | 0 | 0 | 0 | 0 | 0 | 0 | 0 | -1 | 0 | 0 | -1 | 0 | 0 | -1 | 0 | 0 | 0 | 0 | 5 | -1 | 0 | 0 | 0 | 0 | 0 | 0 | 0 | 0 | 0 | 1 | 0 | 0 | 1 | 0 | 0 | 0 | 0 | 0 | 1 | 0 |
| Cutc | chr19:43766099-43766233 | 0 | 0 | 0 | 0 | 0 | 0 | 1 | 0 | 1 | 0 | 0 | 0 | 0 | 0 | 26 | 0 | 0 | 0 | 5 | 2 | 0 | 0 | -1 | 0 | 0 | -1 | 0 | 0 | 0 | 0 | 0 | -1 | -1 | 0 | 0 | 0 | 0 | 0 | 0 | 0 | 0 | 0 | 0 | 2 | 0 | 0 | 0 | 0 | 0 | 0 | 0 | 0 | 0 | 0 | 1 | 0 | 0 | 0 | 0 | 0 | 0 | 0 | 0 | 0 | 0 | 0 | 0 | 0 | 0 | 0 | 0 | 0 | 0 | 1 | 2 | 0 | 1 | 0 | 0 | 1 | 0 |
| Mthfd1 | chr12:76265533-76265640 | 0 | 0 | 0 | 2 | 0 | 0 | 0 | 0 | 0 | 0 | 1 | 0 | 0 | 0 | 26 | -1 | 0 | 0 | 0 | 0 | 0 | 0 | 0 | 0 | 0 | 0 | 0 | 0 | 0 | 0 | -1 | 0 | 0 | 0 | 0 | 0 | 0 | 0 | 0 | 1 | 0 | 0 | 1 | 1 | 0 | 0 | 0 | 0 | 0 | 0 | 0 | 0 | 0 | 0 | 2 | 0 | 0 | 0 | 1 | 0 | 0 | 0 | 0 | 0 | 0 | 0 | 0 | 0 | 1 | 1 | 0 | 0 | 0 | 1 | 0 | 2 | 0 | 0 | 0 | 1 | 1 |
| Stx17 | chr4:48153476-48153570 | 4 | 4 | -1 | 0 | 0 | 0 | 1 | 0 | 0 | 0 | 2 | 0 | 0 | 0 | 51 | 0 | 0 | 2 | 9 | 0 | 6 | 0 | 0 | -1 | 0 | 0 | -1 | -1 | 0 | 0 | 0 | 0 | 0 | 16 | -1 | 0 | 1 | 1 | 0 | 0 | 0 | 0 | 0 | 3 | 0 | 0 | 0 | 1 | 2 | 0 | 0 | -1 | 0 | 0 | 0 | 0 | 0 | 0 | 3 | 0 | -1 | 0 | -1 | -1 | 0 | 0 | 0 | 0 | 0 | 0 | 1 | 1 | 0 | 0 | 0 | 0 | 0 | 0 | 0 | 0 | 0 |
| Prkce | chr17:86635065-86635094 | 4 | 0 | 0 | 0 | 0 | 0 | 0 | 0 | 0 | 0 | 0 | 0 | 0 | 0 | 23 | 20 | 2 | 1 | 0 | 0 | 13 | 4 | 1 | 0 | 2 | 0 | 0 | 0 | 0 | 0 | 0 | 0 | 0 | 0 | 1 | 0 | 4 | 0 | 0 | 0 | 0 | 4 | 0 | 0 | 0 | 0 | 0 | 0 | 0 | 0 | 0 | 0 | 0 | 0 | 0 | 0 | 0 | 0 | 0 | 0 | -1 | 0 | 0 | 0 | 0 | 0 | 0 | 0 | 0 | 0 | 0 | 0 | 0 | 0 | 0 | 0 | 0 | 0 | 0 | 0 | -1 |
| Dhfr | chr13:92381204-92381323 | 3 | 1 | 0 | 0 | 0 | 1 | 1 | 1 | 1 | 1 | 0 | 0 | 3 | 0 | 35 | 0 | -1 | 1 | 2 | 1 | 20 | 0 | -1 | 0 | 0 | 0 | 0 | -1 | 0 | -1 | 0 | -1 | 0 | 0 | 0 | -1 | 0 | 1 | 1 | 2 | 0 | 0 | 1 | 2 | 0 | 1 | -1 | 0 | 0 | 0 | 0 | 0 | 0 | 0 | 0 | 1 | 0 | 0 | -1 | 0 | 0 | 1 | 0 | 0 | 0 | 0 | 0 | 0 | 0 | 1 | 2 | 0 | 1 | 2 | 0 | 0 | 1 | 1 | 0 | 1 | 0 |
| Tmem19 | chr10:115348907-115349062 | 1 | 26 | 0 | 0 | 0 | 0 | 1 | 0 | 2 | 0 | 0 | 0 | 0 | 0 | 30 | 0 | 0 | 1 | 0 | 2 | 1 | 0 | 0 | 0 | 0 | 0 | 0 | 0 | 0 | 0 | 0 | 0 | 0 | -1 | 0 | -1 | 0 | 0 | 0 | 0 | 0 | 0 | 1 | 17 | -1 | 0 | 0 | 0 | 3 | 0 | 0 | 0 | 0 | 0 | 0 | 0 | 0 | 0 | 0 | 0 | 0 | 0 | 0 | 0 | -1 | 0 | 2 | 0 | 0 | 0 | 0 | 0 | 0 | 0 | 0 | 0 | 0 | 0 | 0 | 0 | 1 |

### SFig4, pg2

| Gene Symbol | Exon Location (mm10, mouse) | Brain | Skeletal Muscle | Heart | Lung | Stomach | Spleen | Colon | Liver | Kidney | Pancreas islet | Mammary gland | Ovary | Testes | Rod Photoreceptor (p28) | SST interneurons (cortex) | PV interneurons (hippocampus) | Motor neurons (Hb9+, embryo) | DRG neurons | Olfactory sensory neurons (Omp+) | Hair cells (Atoh1+, cochlea) | Purkinje cells (cerebellum) | Pyramidal neurons (Thy1+, cortex) | Pyramidal neurons (hippo - proximal CA1) | Pyramidal neurons (hippo - distal CA1) | Pyramidal neurons (hippo - superficial CA1) | Pyramidal neurons (hippo - dorsal CA1) | Pyramidal neurons (hippo - dorsal CA1, postsubiculum-projecting) | Pyramidal neurons (hippo - intermediate CA1) | Pyramidal neurons (hippo - ventral CA1) | Pyramidal neurons (hippo - ventral CA1, amygdala-projecting) | Pyramidal neurons (hippo - ventral CA1, nucleus accumbens-projecting | Pyramidal neurons (hippo - deep CA1) | Pyramidal neurons (hippo - dorsal CA3) | Pyramidal neurons (hippo - ventral CA3) | Granule cells (hippo - dorsal DG) | Granule cells (hippo - ventral DG) | Oligodendrocyte precursor cells (cortex) | Newly formed oligodendrocytes (cortex) | Myelinating oligodendrocytes (cortex) | Astrocytes (cortex) | Microglia (cortex) | Microglia (spinal cord) | Endothelial cells (cortex) | Quadriceps | Gastrocnemius | Heart ventricle (p90) | Hematopoietic stem cell (bone marrow) | Dendritic cells (spleen) | Neutrophil | Mast cell | NKT cell | NK cell | Macrophage | Eosinophils | B cell, spleen | CD4+ T-cell (Th17) | CD4+ T-cell (Th1) | CD4+ T-cell (Th2) | CD8+ T-cell | CD4+/CD8+ double positive T-cell | Gamma delta T-cell | Regulatory T-cell (Rag2+, spleen) | Hepatocyte (liver) | Liver sinusoidal endothelial cells (liver) | Kupffer cells (liver) | Hepatic stellate cells (liver) | Smooth muscle cells (colon) | Interstitial Cell of Cajal (colon) | Gastrointestinal PDGFRa+ cells (colon) | Gastrointestinal PDGFRa+ cells (colon mucosa) | Pancreatic alpha cells (7-16wks) | Pancreatic beta cells (7-16wks) | Alveolar type 1 (lung) | Podocytes (kidney) | Classical brown adipocyte | Inguinal white adipocyte | Sertoli cells (testis) | Leydig cells (testis) | Interstitial cells (testis) | Spermatogonial stem cells (testis) | Female germline stem cells (ovary) |
|---|---|---|---|---|---|---|---|---|---|---|---|---|---|---|---|---|---|---|---|---|---|---|---|---|---|---|---|---|---|---|---|---|---|---|---|---|---|---|---|---|---|---|---|---|---|---|---|---|---|---|---|---|---|---|---|---|---|---|---|---|---|---|---|---|---|---|---|---|---|---|---|---|---|---|---|---|---|---|---|---|---|---|
| Sybu | chr15:44729101-44729202 | 3 | 0 | -1 | 0 | 0 | -1 | 0 | 0 | 0 | 2 | 0 | 0 | 0 | -1 | 28 | 0 | 0 | 1 | 2 | 4 | 1 | 0 | 1 | 1 | 1 | 1 | 0 | 0 | 0 | 1 | 0 | 3 | 0 | 1 | 12 | 0 | 0 | 0 | 0 | 2 | 0 | 0 | 0 | -1 | -1 | 0 | -1 | -1 | -1 | -1 | -1 | -1 | -1 | -1 | -1 | -1 | -1 | -1 | -1 | -1 | -1 | -1 | -1 | -1 | -1 | 0 | 0 | -1 | 0 | 1 | 0 | 0 | 0 | -1 | -1 | 0 | -1 | 0 | -1 | 0 |
| Mettl16 | chr11:74782035-74782144 | 0 | 1 | 0 | 0 | 0 | 2 | 0 | 2 | 4 | 1 | 1 | 1 | 0 | 0 | 35 | 0 | 0 | 0 | 4 | 1 | 1 | 0 | 0 | 0 | 0 | 1 | 0 | 0 | 0 | 0 | 0 | 0 | 0 | 0 | -1 | 0 | 0 | 1 | 3 | 3 | -1 | 0 | 3 | 0 | 0 | 0 | 0 | 3 | 0 | 0 | 0 | 0 | 0 | 1 | 1 | 2 | 3 | 3 | 3 | 4 | 0 | 9 | -1 | 0 | -1 | 0 | 1 | 0 | 2 | 0 | 2 | 2 | 0 | 1 | 2 | 4 | 2 | 1 | 1 | 0 | 0 |
| Ascc3 | chr10:50596037-50596127 | -1 | 0 | 0 | 0 | -1 | 0 | 0 | 0 | 0 | 3 | 2 | 0 | 3 | 0 | 38 | 0 | 2 | 6 | 4 | 0 | 1 | -1 | 0 | 0 | 8 | -1 | 0 | -1 | -1 | 0 | 0 | 3 | -1 | 0 | -1 | 0 | -1 | 0 | 0 | 0 | 0 | 0 | 0 | 0 | 1 | 1 | 0 | -1 | 0 | 3 | 0 | -1 | -1 | 2 | 0 | 1 | 3 | 0 | 0 | 4 | 0 | 0 | 0 | 0 | -1 | 0 | 0 | 0 | -1 | 0 | 2 | 2 | 0 | 1 | 0 | 0 | 0 | 1 | 2 | 1 | 1 |
| Rwdd3 | chr3:121171112-121171182 | 2 | -1 | -1 | -1 | 0 | 0 | 0 | 2 | 0 | 2 | 6 | -1 | -1 | 0 | 21 | 0 | 0 | 0 | 2 | 3 | 0 | 0 | 0 | 0 | 0 | 0 | -1 | 0 | 0 | 0 | 0 | 0 | 0 | 0 | 0 | -1 | 0 | 0 | 0 | 4 | 0 | 2 | 0 | -1 | 0 | 1 | 0 | 0 | -1 | 0 | 0 | -1 | -1 | -1 | -1 | -1 | -1 | 0 | 0 | 10 | -1 | 0 | 0 | 0 | 0 | -1 | 0 | 0 | 0 | 0 | 0 | 0 | -1 | 0 | -1 | 0 | 0 | 2 | 1 | 2 | 3 |
| Fcho2 | chr13:98796921-98797016 | 0 | 2 | 0 | 0 | 0 | 2 | 2 | 1 | 2 | 1 | 0 | 3 | 27 | 0 | 62 | 0 | 0 | 0 | 0 | 5 | 0 | 0 | 0 | 0 | 0 | 0 | 0 | 0 | 0 | 0 | 0 | 0 | 0 | 0 | 0 | -1 | 5 | 0 | 0 | 1 | 0 | 0 | 2 | 2 | -1 | 2 | 0 | -1 | 8 | 1 | 0 | -1 | 0 | 2 | 0 | 0 | 1 | 1 | -1 | 1 | 0 | 0 | -1 | 0 | 0 | 2 | 3 | 1 | 2 | 1 | 1 | 2 | 2 | 1 | 0 | 7 | 2 | 1 | 1 | 5 | 19 |
| Rapgef1 | chr2:29717716-29717790 | 0 | 0 | 0 | -1 | -1 | 0 | -1 | 0 | 0 | 0 | 0 | -1 | 0 | 0 | -1 | 61 | 0 | 0 | 1 | 0 | 0 | -1 | 0 | 0 | 0 | 0 | 0 | 0 | 0 | 0 | 0 | 0 | 0 | 0 | 0 | 0 | 0 | 0 | 0 | 0 | 0 | 0 | 0 | 0 | 0 | 0 | -1 | -1 | -1 | -1 | -1 | -1 | -1 | -1 | -1 | -1 | -1 | -1 | -1 | 0 | -1 | -1 | -1 | -1 | -1 | -1 | 0 | 0 | 0 | -1 | -1 | -1 | -1 | 0 | -1 | -1 | 0 | -1 | -1 | 0 | 0 |
| Dsc2 | chr18:20048783-20048817 | -1 | -1 | -1 | 0 | 0 | 0 | 0 | 1 | 0 | 0 | 0 | 0 | -1 | -1 | -1 | 99 | -1 | -1 | 0 | 0 | -1 | -1 | -1 | -1 | 0 | -1 | -1 | -1 | -1 | -1 | -1 | -1 | -1 | -1 | -1 | -1 | 0 | -1 | -1 | 0 | -1 | -1 | -1 | 0 | -1 | 1 | -1 | -1 | -1 | -1 | -1 | -1 | -1 | 0 | -1 | -1 | -1 | -1 | -1 | -1 | -1 | -1 | 0 | 0 | -1 | -1 | -1 | -1 | -1 | 0 | 1 | 0 | 0 | 0 | 0 | 0 | 0 | 0 | 0 | 0 | -1 |
| Herc1 | chr9:66424940-66425005 | 8 | -1 | 0 | 0 | -1 | 0 | 0 | 0 | 0 | 0 | 0 | 0 | 0 | 0 | 8 | 43 | 0 | 1 | 0 | 0 | 12 | 0 | 0 | 0 | 0 | 0 | 0 | 0 | 0 | 0 | 0 | 0 | 0 | 0 | 0 | 0 | 1 | 0 | 0 | 0 | 0 | 1 | 0 | 0 | 0 | 0 | 0 | 0 | 0 | 0 | 0 | 0 | 0 | 0 | 0 | 0 | 0 | 0 | 0 | 0 | 0 | 0 | 0 | 0 | -1 | 0 | 0 | 0 | 0 | 0 | 0 | 0 | 0 | 0 | 0 | 0 | 0 | 0 | 0 | 0 | 0 |
| Ccdc138 | chr10:58537924-58538002 | -1 | 0 | 0 | 0 | -1 | 0 | 0 | 1 | 0 | 2 | 0 | 0 | 0 | 0 | 1 | 35 | 0 | 2 | 1 | 0 | 1 | 0 | 0 | -1 | 0 | -1 | 0 | 0 | 0 | -1 | -1 | -1 | 0 | 0 | -1 | -1 | 0 | 0 | -1 | 0 | -1 | 0 | 0 | 0 | 0 | -1 | 0 | 0 | 0 | 0 | -1 | 0 | -1 | 0 | 0 | 0 | 0 | 0 | 0 | 6 | 0 | 0 | 0 | -1 | 0 | -1 | 0 | 0 | 0 | 2 | 0 | 0 | -1 | 0 | 0 | 0 | 3 | 2 | 1 | 1 | 1 |
| Krit1 | chr5:3815336-3815435 | 0 | 0 | 0 | 0 | -1 | 1 | 1 | 0 | 1 | 0 | 0 | 0 | 1 | 1 | 0 | 31 | 1 | 0 | 1 | 0 | 0 | 0 | 0 | 0 | 1 | 0 | 1 | 0 | 0 | 0 | 0 | 0 | 0 | 0 | 0 | 0 | 0 | 1 | 0 | 2 | 0 | 1 | 0 | 1 | 1 | 0 | 0 | 0 | -1 | 0 | 0 | -1 | 0 | 0 | 0 | 0 | 1 | 0 | 0 | 0 | 0 | 0 | 0 | 0 | 0 | 0 | 0 | 0 | 0 | 1 | 0 | 1 | 0 | 0 | 0 | 0 | 0 | 0 | 0 | 0 | 0 |
| Eya3 | chr4:132664966-132665002 | 1 | 1 | 0 | -1 | 0 | 0 | 1 | 1 | 0 | 1 | 0 | 0 | 0 | 3 | -1 | 32 | 0 | 1 | 0 | 1 | 1 | 0 | 0 | 0 | 0 | 2 | 0 | 0 | 1 | 0 | 0 | 0 | 0 | 0 | 0 | 0 | 0 | 0 | 0 | 0 | 0 | 0 | 0 | 1 | 0 | 0 | 1 | 0 | 0 | 0 | 0 | 0 | 0 | 0 | 0 | 0 | 1 | 1 | 0 | 0 | 0 | -1 | -1 | 0 | 0 | 0 | 0 | 0 | 0 | 0 | 1 | 1 | 0 | 0 | 0 | 2 | 0 | 0 | 1 | 2 | 1 |
| Ift43 | chr12:86085743-86085839 | 0 | 1 | 0 | 0 | 0 | 1 | 0 | 1 | 1 | 0 | 1 | 0 | 0 | 0 | 5 | 30 | 1 | 2 | 0 | 0 | 0 | 4 | 0 | 0 | 0 | -1 | 0 | -1 | 0 | 0 | 0 | 0 | 0 | 0 | 0 | 0 | 0 | 3 | 0 | 0 | 0 | 0 | 0 | 0 | 0 | 1 | 0 | 0 | -1 | -1 | 0 | -1 | 0 | -1 | -1 | 1 | 2 | 0 | -1 | -1 | -1 | 0 | 0 | 0 | 0 | 6 | 1 | 0 | 0 | 0 | 1 | 2 | 0 | 1 | 0 | -1 | 0 | 0 | 1 | 1 | 1 |
| Twf1 | chr15:94583624-94583644 | 1 | 0 | 0 | 0 | 0 | 0 | 0 | 0 | 0 | 0 | 0 | 0 | 0 | 0 | 0 | 23 | 0 | 1 | 0 | 0 | 15 | 0 | 0 | 1 | 1 | 7 | 0 | 0 | 0 | 0 | 0 | 0 | 6 | 1 | 2 | 0 | 0 | 0 | 0 | 0 | 0 | 0 | 0 | 1 | 0 | 1 | 0 | 0 | 0 | 0 | 0 | 0 | 0 | 0 | 0 | 0 | 0 | 0 | 0 | 1 | -1 | 0 | 0 | 0 | 0 | 0 | 0 | 0 | 0 | 0 | 0 | 0 | 0 | 0 | 0 | 0 | 0 | 0 | 0 | 0 | 0 |
| Ranbp17 | chr11:33303919-33304010 | 8 | 0 | -1 | -1 | -1 | 0 | 0 | -1 | 0 | 4 | 0 | -1 | 1 | -1 | 0 | 74 | 0 | 1 | 0 | 1 | 5 | -1 | 0 | -1 | 0 | 0 | 0 | 0 | 0 | -1 | 0 | -1 | 0 | 0 | 13 | 0 | -1 | -1 | 0 | 4 | -1 | -1 | -1 | 0 | -1 | -1 | 0 | -1 | -1 | 0 | -1 | -1 | -1 | -1 | -1 | 0 | 0 | -1 | -1 | -1 | -1 | -1 | -1 | -1 | -1 | -1 | 0 | -1 | 0 | 0 | -1 | -1 | 4 | 0 | -1 | -1 | 2 | 2 | 3 | 4 | 20 |
| Drg2 | chr11:60456222-60456438 | 1 | 0 | 0 | 3 | 0 | 0 | 0 | 1 | 1 | 1 | 2 | 0 | 1 | 0 | 0 | 25 | 0 | 1 | 1 | 1 | 1 | 0 | 1 | 0 | 0 | 0 | 0 | 0 | 0 | 0 | 0 | 0 | 1 | 0 | 0 | 0 | 3 | 1 | 1 | 3 | 1 | 1 | 1 | 0 | 0 | 0 | 0 | 3 | 0 | 0 | 0 | 0 | 0 | 0 | 0 | 0 | 2 | 0 | 0 | 0 | 0 | 0 | 0 | 0 | 0 | 0 | 1 | 1 | 1 | 0 | 1 | 0 | 0 | 1 | 0 | 0 | 0 | 0 | 0 | 1 | 1 |
| Cr1l | chr1:195129334-195129452 | 2 | 0 | 0 | 0 | 0 | 2 | 0 | 0 | 0 | 1 | 1 | 1 | 5 | 0 | 0 | 35 | 1 | 0 | 0 | 0 | 0 | 0 | 0 | 0 | 0 | 7 | 0 | 0 | -1 | 0 | 0 | 0 | 2 | 0 | 4 | 0 | 1 | 1 | 0 | 2 | 0 | 0 | 2 | 0 | 2 | 0 | 3 | 1 | 1 | 1 | 1 | 1 | 1 | 0 | 0 | 1 | 0 | 0 | 0 | 1 | 1 | 4 | 0 | 1 | 0 | 0 | 0 | 0 | 0 | 1 | 1 | 2 | 4 | 0 | 0 | 1 | 2 | 1 | 1 | 0 | 1 |
| Cyfip2 | chr11:46258676-46258750 | 9 | -1 | 4 | 0 | -1 | 0 | 0 | 0 | 0 | 0 | 0 | -1 | 0 | 0 | 0 | 33 | 0 | 24 | 0 | 0 | 13 | 2 | 2 | 3 | 2 | 1 | 3 | 1 | 1 | 1 | 0 | 2 | 3 | 2 | 0 | 1 | 1 | 1 | 3 | 0 | 0 | 11 | 0 | 0 | -1 | -1 | 0 | 0 | 0 | 0 | 0 | 0 | -1 | 0 | 0 | 0 | 0 | 0 | 0 | 0 | 0 | 0 | -1 | 0 | 0 | 0 | 1 | 0 | 0 | 0 | 0 | 0 | 0 | 0 | -1 | 0 | 0 | 0 | 1 | 0 | 0 |
| Cdkl5 | chrX:160812928-160813050 | 8 | 0 | 0 | 0 | -1 | 0 | 0 | -1 | 0 | 0 | 3 | 10 | 4 | 0 | -1 | 62 | 17 | 0 | 0 | -1 | 30 | 0 | 1 | 0 | 0 | 0 | 1 | 0 | 0 | 8 | 0 | 17 | 3 | 1 | 2 | 0 | 1 | -1 | 0 | -1 | -1 | -1 | -1 | -1 | 0 | 0 | 0 | 0 | -1 | -1 | -1 | -1 | 0 | 0 | -1 | -1 | 0 | 0 | -1 | 0 | -1 | -1 | -1 | 0 | 0 | 0 | -1 | 0 | 0 | -1 | 0 | 0 | 5 | 2 | 0 | -1 | 4 | 3 | 0 | 7 | 12 |
| Ptprd | chr4:76041380-76041499 | 0 | 0 | 0 | 0 | -1 | 0 | 0 | 0 | 0 | 0 | 0 | 0 | 0 | 0 | 0 | 1 | 37 | 0 | 0 | 0 | 0 | 0 | 0 | 0 | 0 | 0 | 0 | 0 | 1 | 0 | 0 | 0 | 1 | 0 | 0 | 0 | 1 | 0 | 0 | 1 | 0 | 0 | 0 | 0 | 0 | 0 | -1 | -1 | -1 | -1 | -1 | -1 | -1 | -1 | -1 | -1 | -1 | -1 | -1 | -1 | -1 | -1 | 0 | -1 | -1 | -1 | 0 | 0 | 0 | 0 | 0 | 0 | 0 | 0 | 0 | 0 | 0 | 0 | 0 | 0 | 0 |
| Itga6 | chr2:71824323-71824439 | 3 | 0 | 0 | 0 | 0 | 0 | 0 | 0 | 0 | 0 | 0 | 2 | 0 | 0 | -1 | -1 | 78 | 1 | 0 | 0 | 12 | -1 | -1 | -1 | 0 | 0 | -1 | -1 | -1 | -1 | -1 | -1 | -1 | 0 | 0 | -1 | 1 | -1 | 0 | 1 | 1 | 0 | 1 | 0 | 0 | 0 | 0 | 0 | 0 | 1 | 0 | 0 | 0 | 0 | 0 | 2 | 0 | 0 | 0 | 1 | -1 | 0 | -1 | -1 | -1 | 0 | 0 | 0 | -1 | 0 | 0 | 1 | 3 | 0 | 2 | 1 | 1 | 0 | 0 | 1 | 1 |
| Pcnx2 | chr8:125860773-125860837 | 4 | -1 | -1 | -1 | -1 | -1 | -1 | -1 | -1 | -1 | 0 | -1 | -1 | -1 | -1 | 0 | 78 | 0 | 3 | -1 | 1 | 0 | 0 | 0 | 0 | 0 | 2 | 0 | -1 | -1 | 0 | 3 | 1 | 0 | 0 | 0 | -1 | -1 | 0 | -1 | 0 | -1 | -1 | -1 | -1 | -1 | -1 | -1 | -1 | -1 | -1 | -1 | -1 | -1 | -1 | -1 | -1 | -1 | -1 | -1 | -1 | -1 | -1 | -1 | -1 | -1 | -1 | 0 | -1 | -1 | 0 | -1 | -1 | -1 | -1 | -1 | -1 | -1 | -1 | -1 | -1 |
| Cpne3 | chr4:19539152-19539180 | 16 | 0 | 0 | 0 | -1 | 0 | 0 | 0 | 0 | 0 | 0 | 0 | 0 | -1 | 0 | -1 | 27 | 0 | 0 | 0 | 9 | -1 | -1 | -1 | -1 | -1 | -1 | -1 | -1 | -1 | -1 | -1 | -1 | -1 | -1 | -1 | 0 | 0 | 0 | 0 | 0 | 0 | 0 | 0 | 0 | 0 | 0 | 0 | 0 | 0 | 0 | 0 | 0 | 0 | 0 | 0 | 0 | 0 | 0 | 0 | 0 | 0 | 0 | 0 | 0 | 0 | 0 | 0 | 0 | 0 | 0 | 0 | 0 | 0 | 0 | 0 | 0 | 0 | 0 | 0 | 0 |
| Stx2 | chr5:128989379-128989406 | 39 | 0 | 0 | 1 | 0 | 0 | 0 | 0 | 0 | 0 | 0 | 0 | 9 | 0 | -1 | 0 | 52 | 0 | 0 | 0 | -1 | -1 | -1 | -1 | -1 | -1 | -1 | -1 | -1 | -1 | -1 | -1 | -1 | -1 | -1 | -1 | 0 | 0 | 0 | 0 | 0 | 0 | 0 | 1 | 0 | 0 | 0 | 0 | 2 | 0 | 0 | 0 | 0 | 0 | 0 | 0 | 0 | 0 | 0 | 0 | 0 | 0 | 0 | 0 | 0 | 0 | 0 | 1 | 0 | 0 | 1 | 0 | 0 | 0 | 0 | 0 | 0 | 0 | 0 | 2 | 0 |
| Carmil1 | chr13:24049643-24049660 | 21 | 0 | 0 | 1 | 0 | 0 | 0 | 1 | 0 | 3 | 0 | -1 | 0 | -1 | -1 | -1 | 97 | -1 | 38 | 0 | 22 | -1 | -1 | -1 | 0 | 0 | -1 | -1 | -1 | -1 | -1 | -1 | -1 | -1 | 0 | 2 | 0 | 0 | 0 | 0 | -1 | 1 | 0 | 0 | 0 | 0 | 0 | -1 | -1 | 0 | -1 | -1 | -1 | 0 | 0 | 0 | 0 | -1 | -1 | 0 | -1 | -1 | 0 | -1 | -1 | 0 | 0 | 0 | 0 | 0 | 3 | 4 | 0 | 0 | 0 | 0 | 0 | 0 | 0 | 1 | 1 |
| Clip1 | chr5:123586334-123586354 | 9 | 0 | 0 | 0 | 0 | 0 | 0 | 0 | 0 | 2 | 0 | 0 | 0 | 0 | 24 | -1 | 67 | 5 | 5 | 4 | 14 | 0 | 0 | 0 | 16 | 0 | -1 | -1 | 0 | 9 | 0 | -1 | 6 | 3 | -1 | 0 | 0 | 0 | 0 | 0 | 0 | 0 | 0 | 0 | 0 | 0 | 0 | 0 | 0 | 0 | 0 | 0 | 0 | 0 | 0 | 0 | 0 | 0 | 0 | 0 | 0 | 0 | 0 | 0 | 0 | 0 | 0 | 0 | 0 | 0 | 2 | 1 | 0 | 0 | 0 | 0 | 0 | 0 | 0 | 0 | 0 |
| Mettl25 | chr10:105800227-105800384 | 4 | 9 | 0 | 0 | 0 | 0 | 0 | 0 | 0 | 0 | 1 | 0 | 0 | 0 | 0 | 0 | 31 | 4 | 0 | 0 | 3 | -1 | 0 | 0 | -1 | -1 | -1 | 0 | 0 | 0 | -1 | -1 | 0 | -1 | -1 | -1 | 1 | 2 | -1 | 0 | -1 | 0 | 0 | 10 | 0 | 0 | 0 | 0 | -1 | 0 | 0 | 0 | 0 | 3 | 0 | 0 | 0 | 0 | 0 | 0 | 0 | 0 | 0 | 0 | -1 | 0 | 0 | 0 | 0 | 0 | 0 | 0 | 0 | 0 | 0 | 0 | 0 | 1 | 0 | 1 | 2 |
| Sema5a | chr15:32640717-32640734 | 15 | 0 | -1 | 0 | 0 | 0 | 0 | -1 | 0 | 0 | 0 | 0 | 0 | -1 | -1 | -1 | 21 | 5 | -1 | 0 | 7 | 0 | -1 | -1 | -1 | -1 | -1 | -1 | -1 | -1 | -1 | 0 | 0 | 0 | 8 | 0 | 0 | 0 | 0 | 0 | 0 | 0 | 0 | 0 | 0 | 0 | 0 | -1 | -1 | 0 | -1 | -1 | 0 | -1 | 0 | -1 | -1 | -1 | 0 | -1 | -1 | -1 | -1 | 0 | -1 | 0 | 0 | 0 | 0 | 0 | 0 | 0 | 0 | 0 | -1 | 0 | 0 | 0 | 0 | 0 | 0 |
| Ptprf | chr4:118252076-118252093 | 38 | 0 | 0 | 0 | 0 | 0 | 0 | 0 | 0 | 0 | 0 | 0 | 0 | 0 | -1 | -1 | 55 | 0 | 0 | 0 | 21 | -1 | 0 | -1 | -1 | 0 | 0 | -1 | -1 | 3 | -1 | 0 | -1 | 31 | -1 | -1 | 3 | 3 | 2 | 0 | 0 | -1 | 0 | 0 | 0 | 0 | 0 | -1 | -1 | 0 | -1 | -1 | -1 | -1 | 0 | -1 | 0 | -1 | 0 | 0 | 0 | 0 | 0 | -1 | -1 | 0 | 0 | 0 | 0 | 0 | 0 | 0 | 0 | 0 | 0 | 0 | 0 | 0 | 0 | 0 | 0 |
| Dgki | chr6:36994859-36994921 | 6 | -1 | -1 | -1 | -1 | 0 | 0 | -1 | 0 | 0 | -1 | -1 | -1 | -1 | -1 | -1 | 99 | 15 | 0 | 0 | 12 | -1 | -1 | 0 | 2 | 3 | 0 | -1 | 0 | 0 | 0 | 0 | 0 | 0 | -1 | 0 | 4 | -1 | -1 | 5 | -1 | 0 | 1 | 0 | 0 | 0 | -1 | -1 | -1 | 0 | 0 | -1 | -1 | -1 | -1 | 0 | -1 | -1 | -1 | -1 | -1 | -1 | -1 | -1 | -1 | -1 | -1 | 0 | 0 | -1 | 0 | -1 | -1 | -1 | -1 | -1 | -1 | -1 | -1 | 56 | -1 |
| Piezo2 | chr18:63099053-63099094 | -1 | 0 | 0 | 0 | 0 | -1 | -1 | 0 | 0 | 0 | 0 | 0 | -1 | -1 | -1 | -1 | -1 | 73 | -1 | 0 | 0 | -1 | -1 | -1 | -1 | -1 | -1 | -1 | -1 | -1 | -1 | -1 | -1 | -1 | -1 | -1 | 0 | 0 | 0 | 0 | -1 | -1 | 0 | 0 | -1 | 0 | -1 | 0 | -1 | -1 | -1 | -1 | -1 | -1 | -1 | -1 | -1 | -1 | -1 | -1 | -1 | -1 | -1 | 0 | -1 | 0 | 0 | 0 | 0 | 0 | -1 | 0 | 0 | 0 | 0 | 0 | 0 | 0 | 0 | 0 | 0 |
| Pkn1 | chr8:83698008-83698072 | 0 | 3 | 0 | 0 | 0 | 0 | 0 | 0 | 0 | 0 | 0 | 0 | 0 | 0 | -1 | 0 | 0 | 62 | 0 | 0 | 0 | -1 | 0 | -1 | -1 | 0 | 0 | -1 | -1 | -1 | -1 | -1 | -1 | 0 | 0 | 0 | 0 | 0 | 0 | 1 | 0 | 0 | 0 | 1 | 0 | 0 | 0 | 0 | 0 | 0 | 0 | 0 | 0 | 0 | 0 | 0 | 0 | 0 | 0 | 0 | 0 | -1 | 0 | 2 | 0 | 0 | 1 | 1 | 1 | 0 | 0 | 0 | 0 | 0 | 0 | 0 | 0 | 0 | 0 | 0 | 0 |
| Sh3kbp1 | chrX:159767951-159767972 | 1 | -1 | 0 | 0 | 0 | 0 | 0 | 0 | 0 | 0 | 0 | 0 | 0 | 0 | 0 | 1 | 1 | 29 | 0 | 0 | 1 | -1 | -1 | -1 | -1 | -1 | -1 | 0 | 0 | 0 | 0 | 0 | 0 | 0 | 0 | 0 | 0 | 0 | 0 | 0 | 0 | 0 | 0 | 6 | 0 | 0 | 0 | 0 | 0 | 1 | 0 | 0 | 0 | 0 | 0 | 0 | 0 | 0 | 0 | 1 | 0 | 0 | -1 | 0 | 0 | 0 | 0 | 0 | 0 | 0 | 1 | 1 | 0 | 0 | 0 | 1 | 0 | 1 | 0 | 0 | 0 |
| Ptprf | chr4:118221571-118221582 | 3 | 0 | 0 | 0 | 0 | 0 | 0 | 0 | 0 | 0 | 0 | 0 | 0 | 0 | -1 | 0 | 4 | 80 | 0 | 0 | 17 | 0 | 0 | 0 | 0 | 0 | -1 | -1 | 0 | 0 | 0 | 0 | 3 | 1 | 0 | 0 | 0 | 0 | 0 | 0 | -1 | -1 | -1 | 0 | 0 | 0 | -1 | 0 | 0 | -1 | -1 | 0 | -1 | -1 | 0 | 0 | 0 | 0 | 0 | 0 | -1 | 0 | 0 | 0 | 0 | 0 | 1 | 1 | 1 | 0 | 0 | 0 | 0 | 0 | -1 | 0 | 0 | 0 | 0 | 0 | 0 |
| Epb41l1 | chr2:156517671-156523771 | 10 | 0 | 0 | -1 | 0 | 0 | 0 | 0 | 0 | 0 | 1 | 0 | 0 | -1 | -1 | 24 | 1 | 83 | 8 | 2 | 20 | 0 | 0 | 3 | 0 | 0 | 0 | 0 | 0 | 0 | 0 | 5 | 1 | 0 | 1 | 0 | 2 | 0 | 2 | 1 | 1 | -1 | 0 | 0 | 0 | 0 | 0 | -1 | -1 | 1 | -1 | -1 | 0 | -1 | 0 | -1 | 0 | 0 | -1 | -1 | -1 | -1 | -1 | 0 | 0 | 0 | 1 | 1 | 2 | 0 | 0 | 0 | 0 | 0 | 0 | 0 | 0 | 0 | 1 | 0 | 0 |
| Opcml | chr9:28904271-28904297 | 2 | 0 | 0 | -1 | -1 | 0 | 0 | -1 | 0 | 0 | 0 | -1 | 0 | -1 | 0 | 4 | 48 | 92 | 0 | 0 | 7 | 0 | 0 | 0 | 0 | 0 | 5 | 0 | 0 | 0 | 0 | 0 | 1 | 1 | 1 | 0 | 0 | 0 | 0 | 0 | 0 | -1 | -1 | 0 | 0 | -1 | -1 | -1 | -1 | -1 | -1 | -1 | -1 | -1 | -1 | -1 | -1 | -1 | -1 | -1 | -1 | -1 | -1 | -1 | -1 | -1 | 0 | 0 | -1 | 0 | 0 | -1 | -1 | 0 | -1 | -1 | 0 | 0 | 0 | 0 | -1 |
| Elmo1 | chr13:20285315-20285359 | 10 | 0 | 0 | 0 | 0 | 0 | 0 | 0 | 0 | 0 | 0 | 0 | 0 | 0 | 0 | -1 | 0 | 80 | -1 | 0 | 95 | -1 | 0 | -1 | -1 | -1 | -1 | -1 | -1 | -1 | -1 | -1 | -1 | -1 | 0 | -1 | 0 | 1 | 0 | 0 | 0 | 0 | 0 | 0 | 2 | 0 | 1 | 0 | 0 | 0 | -1 | 0 | 0 | 0 | 0 | 0 | 0 | 0 | 0 | 0 | 0 | -1 | 0 | 0 | 0 | 0 | 0 | 2 | 0 | 0 | 0 | 1 | 0 | 0 | 0 | 0 | 0 | 0 | 0 | 0 | 0 |
| Dysf | chr6:84147315-84147362 | -1 | 2 | 0 | 0 | 0 | 1 | -1 | 0 | 3 | 2 | 0 | -1 | 0 | -1 | -1 | -1 | -1 | 74 | -1 | 0 | -1 | -1 | -1 | -1 | -1 | -1 | -1 | -1 | -1 | -1 | -1 | -1 | -1 | -1 | -1 | -1 | 0 | -1 | 0 | 0 | -1 | -1 | 0 | 7 | 2 | 2 | -1 | -1 | -1 | -1 | 0 | -1 | -1 | -1 | -1 | -1 | -1 | -1 | -1 | -1 | -1 | -1 | -1 | 0 | 0 | 1 | 1 | 1 | 0 | 3 | -1 | 0 | -1 | 1 | 0 | 0 | 0 | 0 | 0 | 0 | 3 |
| Tacc2 | chr7:130731531-130731566 | 2 | 1 | 0 | 0 | 6 | 0 | 0 | 0 | 0 | 0 | 2 | 1 | 1 | 0 | 0 | -1 | 15 | 33 | 0 | 0 | 0 | 0 | 0 | 0 | 0 | 0 | 0 | 0 | 0 | 0 | 0 | 0 | 0 | 0 | 0 | 0 | 0 | 1 | 0 | 0 | 0 | -1 | 3 | 2 | 4 | 0 | 0 | -1 | -1 | 0 | 0 | 0 | -1 | -1 | -1 | 0 | 0 | 0 | 0 | 0 | 0 | 0 | 0 | 0 | 0 | 0 | 0 | 0 | 0 | 0 | 0 | 0 | 0 | 1 | 0 | 0 | 1 | 2 | 1 | 0 | 0 |
| Kcnq4 | chr4:120708854-120708910 | -1 | 59 | 0 | 0 | -1 | 0 | -1 | 0 | -1 | 0 | 0 | -1 | 0 | -1 | -1 | -1 | 0 | 84 | -1 | 2 | 10 | -1 | -1 | -1 | -1 | -1 | -1 | -1 | -1 | -1 | -1 | -1 | -1 | -1 | -1 | -1 | 0 | 0 | 0 | 0 | 0 | -1 | -1 | -1 | -1 | 0 | -1 | -1 | -1 | -1 | -1 | -1 | -1 | -1 | -1 | -1 | 0 | -1 | -1 | -1 | -1 | -1 | -1 | -1 | -1 | 0 | 3 | 0 | 0 | 0 | 0 | -1 | 0 | -1 | -1 | 0 | 0 | 0 | 0 | 0 | 0 |
| Col11a1 | chr3:114094861-114095013 | 1 | 0 | -1 | -1 | -1 | -1 | -1 | -1 | 0 | 0 | 1 | 2 | -1 | -1 | -1 | -1 | 1 | 23 | -1 | 0 | 0 | -1 | 0 | 0 | 0 | 0 | 0 | -1 | -1 | 0 | 0 | 0 | 0 | 0 | 0 | -1 | 0 | 0 | -1 | 2 | -1 | 0 | -1 | 1 | 0 | 0 | -1 | -1 | -1 | -1 | -1 | -1 | -1 | -1 | -1 | -1 | -1 | -1 | -1 | -1 | -1 | -1 | -1 | -1 | 0 | -1 | 0 | -1 | 0 | -1 | 0 | -1 | 0 | 0 | 0 | 1 | 0 | -1 | 0 |
| Kdm2b | chr5:122937628-122937650 | 2 | 0 | 0 | 0 | 0 | 0 | 0 | 0 | 0 | 0 | 0 | 0 | 0 | 0 | -1 | -1 | 4 | 45 | 0 | 0 | 15 | -1 | -1 | 0 | 7 | 0 | 0 | 0 | 40 | 0 | 0 | 0 | 14 | 1 | -1 | 0 | 2 | -1 | 0 | 0 | 0 | 2 | 0 | 0 | 0 | 0 | 0 | 0 | 0 | 0 | 0 | 0 | 0 | 0 | 0 | 0 | 0 | 0 | 0 | 0 | 0 | -1 | 0 | 0 | 0 | 0 | 0 | 0 | 0 | 0 | 0 | 0 | 0 | 0 | 0 | 0 | 0 | 0 | 0 | 0 | 0 |
| Tbc1d4 | chr14:101471428-101471451 | 17 | 1 | 0 | 0 | 0 | 0 | 0 | 0 | 0 | 0 | 0 | 0 | 0 | -1 | 0 | -1 | 0 | 49 | 0 | 0 | 41 | -1 | -1 | -1 | 0 | -1 | 0 | -1 | -1 | 0 | -1 | -1 | -1 | -1 | 0 | -1 | 0 | 0 | 0 | 0 | 1 | 22 | 1 | 0 | 1 | 0 | 0 | 0 | -1 | 0 | 0 | -1 | -1 | 0 | -1 | 0 | 0 | 0 | 0 | 0 | -1 | 0 | -1 | 0 | -1 | 0 | -1 | 0 | 0 | 0 | 0 | 0 | 0 | 0 | 0 | 2 | 0 | 0 | 0 | 0 | 0 |
| Fez2 | chr17:78379870-78379898 | 3 | 0 | 0 | 0 | 0 | 0 | 0 | 0 | 0 | 0 | 0 | 0 | 0 | 4 | 0 | 1 | 0 | 31 | 0 | 0 | 34 | 17 | 0 | 0 | 0 | 1 | -1 | -1 | 0 | 0 | 0 | -1 | 8 | 0 | 0 | 0 | 0 | 0 | 0 | 0 | 0 | 0 | 0 | 0 | 0 | 0 | 0 | 0 | -1 | 0 | 0 | 0 | 0 | 0 | 0 | 0 | 0 | 0 | 0 | 0 | 0 | 0 | 0 | 0 | 0 | 0 | 0 | 0 | 0 | 0 | 0 | 0 | 0 | 0 | 0 | 0 | 0 | 0 | 0 | 0 | 0 |
| Ryr2 | chr13:11746985-11747014 | 3 | 0 | 1 | -1 | -1 | -1 | -1 | -1 | 0 | -1 | 0 | -1 | 0 | 0 | -1 | 0 | 0 | 30 | -1 | 0 | 12 | 0 | 4 | 0 | 0 | 0 | 3 | 0 | 0 | 0 | 1 | 1 | 3 | 3 | 3 | 0 | 0 | 0 | 0 | 0 | -1 | -1 | -1 | -1 | -1 | 0 | 0 | -1 | -1 | -1 | -1 | -1 | 0 | -1 | -1 | -1 | -1 | -1 | -1 | -1 | -1 | -1 | -1 | -1 | -1 | -1 | 0 | 0 | 0 | -1 | -1 | -1 | 0 | -1 | 0 | 0 | 0 | -1 | 0 | 0 | 0 |
| Trim36 | chr18:46192360-46192389 | 8 | 0 | -1 | 0 | -1 | 0 | 0 | -1 | 0 | 0 | 0 | 0 | 0 | 0 | -1 | -1 | 0 | 39 | 0 | 46 | 5 | -1 | -1 | 0 | -1 | 0 | -1 | 0 | -1 | 0 | -1 | 0 | -1 | 0 | -1 | -1 | 0 | 0 | 1 | 0 | 0 | 0 | 0 | -1 | 0 | 0 | 0 | 0 | -1 | 0 | 0 | 0 | 0 | 0 | 0 | 0 | 0 | 0 | -1 | -1 | 0 | -1 | -1 | -1 | 0 | 0 | 0 | -1 | -1 | 0 | 0 | -1 | 0 | 0 | -1 | -1 | 0 | -1 | 0 | 0 | 0 |
| Nxpe4 | chr9:48359328-48359412 | 5 | -1 | 0 | -1 | -1 | -1 | -1 | -1 | -1 | 3 | 0 | 0 | 0 | 0 | -1 | -1 | -1 | 30 | 0 | 4 | 11 | -1 | 0 | 0 | -1 | -1 | -1 | -1 | -1 | 0 | -1 | -1 | -1 | -1 | 0 | -1 | -1 | -1 | -1 | 8 | 0 | -1 | -1 | 1 | 0 | 0 | -1 | -1 | -1 | -1 | -1 | -1 | -1 | -1 | -1 | -1 | -1 | -1 | -1 | -1 | -1 | 0 | 0 | 0 | 0 | 0 | -1 | 0 | 0 | -1 | 0 | 3 | 0 | 0 | 0 | 0 | -1 | 0 | -1 |
| Cd99l2 | chrX:71425415-71425432 | 12 | 2 | 5 | 1 | 0 | 0 | 1 | 1 | 1 | 0 | 0 | 0 | 0 | -1 | 15 | 22 | 2 | 71 | 6 | 0 | 53 | 0 | 0 | 0 | 3 | 4 | 0 | -1 | 0 | 0 | 0 | 2 | 4 | 0 | 3 | 0 | 0 | 0 | 0 | 1 | 0 | 2 | 0 | 0 | 0 | 6 | 0 | 0 | -1 | 0 | 0 | 0 | -1 | 0 | 0 | 1 | 0 | 1 | 0 | 0 | 0 | 0 | 0 | 0 | 0 | 0 | 2 | 0 | 1 | 0 | 1 | 1 | 0 | 0 | 2 | 0 | 0 | 0 | 0 | 0 | 6 |
| Slc15a2 | chr16:36756865-36756892 | 4 | -1 | -1 | 0 | -1 | 0 | 0 | -1 | 0 | 6 | 1 | -1 | 0 | 0 | -1 | -1 | -1 | 33 | 0 | 5 | 0 | -1 | -1 | -1 | -1 | -1 | -1 | -1 | -1 | -1 | -1 | -1 | -1 | -1 | -1 | -1 | 0 | 0 | 0 | 0 | 0 | 1 | 0 | 37 | -1 | 4 | 0 | 0 | 0 | 2 | -1 | 0 | 0 | 0 | 0 | 1 | 0 | 2 | 0 | 0 | 0 | 0 | 0 | 0 | 0 | 0 | 0 | 0 | 0 | 0 | 1 | 0 | 0 | 0 | -1 | -1 | 0 | 0 | 0 | 0 | 0 |
| Nrxn3 | chr12:89191153-89191176\|89191197 | 3 | 0 | -1 | -1 | -1 | -1 | 0 | -1 | 0 | 0 | 0 | 0 | 0 | -1 | 0 | 0 | 0 | 88 | -1 | 94 | 4 | 0 | 0 | 0 | 0 | 3 | 0 | 1 | 0 | 0 | 0 | 0 | 0 | 3 | 0 | 0 | 0 | 0 | -1 | 0 | -1 | 0 | 0 | -1 | 0 | -1 | -1 | -1 | -1 | -1 | -1 | -1 | -1 | -1 | -1 | -1 | -1 | -1 | -1 | -1 | -1 | -1 | -1 | -1 | -1 | -1 | -1 | -1 | -1 | -1 | -1 | -1 | -1 | 0 | -1 | -1 | 0 | 0 | 0 | 0 | -1 |
| Kcnab2 | chr4:152400221-152400268 | 2 | 0 | -1 | -1 | -1 | 0 | 0 | 0 | 0 | 0 | 0 | 0 | 1 | 6 | 0 | 2 | 0 | 27 | 0 | 1 | 6 | -1 | 0 | 0 | 0 | 2 | 0 | 2 | 0 | 1 | 0 | 0 | 2 | 1 | 4 | 3 | 1 | -1 | 7 | 0 | 2 | 1 | 0 | 0 | 0 | -1 | 0 | 2 | 5 | 0 | 3 | 1 | 2 | 2 | 1 | 0 | 0 | 0 | 1 | 1 | 1 | 0 | -1 | 0 | 0 | -1 | 0 | 0 | 0 | 0 | 0 | 0 | 0 | 0 | -1 | 0 | 0 | 0 | 0 | 3 | 0 |
| Cask | chrX:13531866-13531985 | 14 | 7 | -1 | 0 | 0 | 0 | 0 | 0 | 0 | 0 | 0 | 3 | 0 | 0 | -1 | 1 | 12 | 61 | 0 | 0 | 16 | 8 | 11 | 0 | 15 | 3 | 4 | -1 | 5 | 0 | 0 | 0 | 2 | 2 | 0 | 0 | 4 | 4 | 3 | 1 | 0 | 0 | 1 | 0 | 4 | 0 | 0 | -1 | 0 | 0 | 0 | 0 | 0 | 0 | -1 | 0 | -1 | 0 | 0 | 0 | -1 | 0 | 0 | 0 | 0 | 0 | 13 | 1 | 1 | 0 | 1 | 0 | 0 | 1 | 0 | 0 | 0 | 0 | 0 | 1 | 7 |
| Dopey2 | chr16:93781689-93781724 | 28 | 26 | 0 | 3 | 0 | 0 | 0 | 1 | 1 | 0 | 2 | 0 | 0 | 0 | -1 | -1 | 1 | 91 | 0 | 1 | 26 | -1 | 5 | 25 | 19 | 20 | -1 | -1 | 4 | 0 | 0 | -1 | 38 | 15 | -1 | -1 | 5 | 0 | 0 | 0 | 0 | 2 | 0 | -1 | -1 | 0 | -1 | 0 | 0 | 0 | 2 | 0 | 0 | 0 | 0 | 1 | 1 | 1 | 0 | 0 | 0 | 0 | 0 | 0 | 0 | 0 | 8 | 6 | 7 | 1 | 2 | 1 | 0 | 0 | 0 | 0 | 2 | 2 | 0 | 0 | 1 |
| Ano4 | chr10:89256969-89257137 | 7 | -1 | -1 | -1 | -1 | -1 | -1 | -1 | -1 | -1 | -1 | 0 | -1 | -1 | 0 | 0 | 2 | 22 | -1 | 2 | 3 | 0 | 0 | -1 | -1 | 0 | 0 | 0 | 0 | 0 | 0 | 0 | -1 | 0 | -1 | -1 | 1 | 3 | 0 | 0 | -1 | -1 | -1 | 0 | -1 | -1 | -1 | -1 | -1 | -1 | -1 | -1 | -1 | -1 | -1 | -1 | -1 | -1 | -1 | -1 | -1 | -1 | -1 | -1 | -1 | -1 | 3 | -1 | 0 | -1 | -1 | -1 | -1 | -1 | -1 | -1 | 0 | 0 | -1 | -1 | -1 |
| Pwp2 | chr10:78172633-78172681 | 15 | 6 | -1 | 0 | 0 | 1 | 0 | 0 | 1 | 1 | 0 | 0 | 0 | 0 | 0 | 0 | 0 | 21 | 0 | 0 | 5 | 12 | -1 | 0 | -1 | -1 | 0 | -1 | 0 | 0 | -1 | 0 | 0 | -1 | 0 | 0 | 0 | 0 | 0 | 0 | 0 | 1 | 0 | 3 | 0 | 0 | 0 | 0 | 0 | 0 | 0 | 0 | 0 | 0 | 0 | 0 | 1 | 0 | 0 | 0 | 0 | 0 | 0 | 0 | 0 | -1 | 2 | 3 | 1 | 2 | 1 | 0 | 0 | 0 | 0 | 0 | 0 | 1 | 1 | 0 | 0 |
| Ppfia2 | chr10:106835177-106835263 | 10 | 0 | -1 | 0 | -1 | 0 | -1 | 0 | 0 | 0 | 0 | 0 | 0 | 1 | 1 | 11 | 0 | 37 | 0 | -1 | 13 | 9 | 2 | 10 | 5 | 7 | 4 | 5 | 3 | 0 | 0 | 2 | 5 | 3 | 0 | 1 | 0 | -1 | -1 | 0 | 0 | -1 | 0 | 0 | -1 | 0 | -1 | -1 | -1 | -1 | -1 | -1 | -1 | -1 | -1 | -1 | -1 | -1 | -1 | -1 | -1 | -1 | -1 | 0 | 0 | 0 | 0 | 0 | -1 | 0 | 1 | 0 | -1 | 0 | -1 | -1 | -1 | 0 | 0 | 0 | -1 |
| Fam184a | chr10:53662824-53662865 | 17 | 0 | -1 | -1 | -1 | -1 | -1 | 0 | 0 | 0 | 0 | 0 | 0 | 0 | 0 | 66 | 2 | 79 | 0 | 0 | 25 | -1 | 18 | -1 | -1 | 7 | -1 | 0 | -1 | 0 | 0 | -1 | -1 | -1 | 1 | -1 | 0 | 6 | -1 | 0 | -1 | -1 | 0 | -1 | -1 | -1 | 0 | 0 | -1 | -1 | -1 | -1 | -1 | -1 | -1 | 0 | -1 | -1 | -1 | 0 | -1 | -1 | -1 | -1 | -1 | -1 | -1 | 0 | 0 | 0 | 0 | 0 | 0 | 0 | -1 | 0 | 0 | 0 | 0 | 0 | 0 |
| Slc8a1 | chr17:81429227-81429250 | 0 | 0 | 0 | 0 | 0 | 0 | 0 | 0 | 0 | 0 | 0 | 0 | 0 | -1 | 0 | 0 | 0 | 0 | 32 | 0 | 0 | 0 | 0 | 0 | 0 | 0 | 0 | 0 | 0 | 0 | 0 | 0 | 0 | 0 | 0 | 0 | 0 | 0 | 0 | 0 | 0 | 0 | 0 | 0 | 0 | 0 | 0 | 0 | 0 | 0 | 0 | -1 | 0 | 0 | -1 | 0 | -1 | -1 | 0 | 0 | -1 | -1 | -1 | -1 | 0 | 0 | 0 | 0 | 0 | 0 | 0 | 0 | 0 | 0 | -1 | -1 | 0 | 0 | 0 | 0 | 0 |
| Fam19a4 | chr6:96929987-96930139 | 0 | 0 | -1 | -1 | 0 | -1 | -1 | -1 | -1 | -1 | -1 | 0 | 0 | -1 | -1 | -1 | 0 | 0 | 37 | 0 | -1 | -1 | -1 | -1 | -1 | -1 | -1 | -1 | -1 | -1 | -1 | -1 | -1 | -1 | -1 | -1 | 0 | 0 | -1 | -1 | -1 | 0 | -1 | 0 | 0 | -1 | -1 | -1 | -1 | -1 | -1 | -1 | -1 | -1 | -1 | -1 | -1 | -1 | -1 | -1 | -1 | -1 | -1 | -1 | -1 | -1 | -1 | 0 | -1 | -1 | -1 | -1 | 0 | -1 | -1 | -1 | -1 | -1 | -1 | -1 | -1 |
| Thsd1 | chr8:22249792-22249979 | 0 | 0 | 0 | 1 | 0 | -1 | 0 | 0 | 0 | 0 | -1 | 0 | 0 | -1 | -1 | -1 | 0 | 0 | 44 | 0 | 0 | -1 | -1 | -1 | -1 | -1 | -1 | -1 | -1 | -1 | -1 | -1 | -1 | -1 | -1 | 0 | 0 | 0 | -1 | -1 | 0 | 0 | 0 | 0 | 0 | 0 | 0 | -1 | 0 | -1 | 0 | -1 | -1 | -1 | -1 | 0 | 0 | 0 | -1 | 0 | 0 | -1 | 0 | 0 | 0 | 0 | -1 | 0 | 0 | 0 | -1 | -1 | 0 | 1 | 0 | 0 | 0 | 0 | 0 | 0 | 0 |
| Gm38250 | chr8:105580154-105580384 | -1 | -1 | -1 | 0 | -1 | 0 | -1 | -1 | -1 | 5 | -1 | 0 | -1 | -1 | -1 | -1 | 0 | 0 | 53 | 0 | -1 | -1 | -1 | -1 | -1 | -1 | -1 | -1 | -1 | -1 | -1 | -1 | -1 | -1 | -1 | -1 | -1 | -1 | -1 | -1 | -1 | 0 | 0 | -1 | -1 | -1 | 0 | -1 | -1 | 0 | -1 | -1 | -1 | -1 | 0 | 0 | 0 | 0 | 0 | 0 | -1 | -1 | -1 | -1 | -1 | -1 | -1 | 0 | -1 | -1 | 0 | 0 | -1 | -1 | -1 | -1 | -1 | -1 | 0 | -1 | -1 |
| Cngb1 | chr8:95275556-95275823 | -1 | -1 | -1 | -1 | -1 | -1 | -1 | -1 | -1 | -1 | -1 | -1 | -1 | 3 | -1 | -1 | -1 | -1 | 99 | -1 | -1 | -1 | -1 | -1 | -1 | -1 | -1 | -1 | -1 | -1 | -1 | -1 | -1 | -1 | -1 | -1 | -1 | -1 | -1 | -1 | -1 | -1 | -1 | -1 | -1 | -1 | -1 | -1 | -1 | -1 | -1 | -1 | -1 | -1 | -1 | -1 | -1 | -1 | -1 | -1 | -1 | -1 | -1 | 0 | -1 | 0 | -1 | -1 | -1 | -1 | -1 | -1 | -1 | -1 | -1 | -1 | -1 | -1 | -1 | -1 | -1 |
| Spice1 | chr16:44356506-44356615 | 0 | 0 | -1 | 0 | 0 | 4 | 0 | 0 | 0 | 0 | 0 | 0 | 0 | 0 | 0 | 0 | 0 | 0 | 21 | 0 | 0 | 0 | 0 | 0 | 0 | 0 | 0 | 0 | 0 | 0 | 0 | 0 | 0 | 0 | 0 | 0 | 0 | 0 | 0 | 0 | -1 | 0 | 2 | 0 | 0 | 0 | 0 | 0 | 0 | 0 | 0 | 0 | 0 | -1 | 0 | 0 | 0 | 1 | 0 | 0 | 0 | 0 | -1 | -1 | 0 | -1 | 0 | 0 | 0 | 0 | 0 | 0 | 0 | 0 | 0 | 0 | 0 | 0 | 0 | 0 | 0 |
| Myt1 | chr2:181803930-181804022 | 0 | -1 | -1 | 0 | -1 | -1 | 0 | -1 | -1 | 0 | -1 | -1 | 0 | 0 | -1 | 0 | 0 | 1 | 29 | 1 | 0 | -1 | 0 | -1 | 0 | 0 | -1 | 0 | 0 | 0 | -1 | 0 | 0 | 0 | 0 | 0 | 0 | 0 | 0 | 0 | -1 | 0 | 0 | -1 | -1 | -1 | -1 | -1 | -1 | -1 | -1 | -1 | -1 | -1 | -1 | -1 | -1 | -1 | -1 | -1 | -1 | -1 | -1 | -1 | -1 | -1 | -1 | 0 | -1 | -1 | 1 | 1 | -1 | -1 | -1 | -1 | -1 | -1 | -1 | -1 | -1 |
| Efna5 | chr17:62612253-62612294 | 1 | 0 | 0 | 0 | 0 | -1 | 0 | 0 | 0 | 1 | 0 | 0 | 0 | -1 | -1 | 0 | 1 | 0 | 71 | 4 | 9 | -1 | 0 | -1 | 0 | -1 | -1 | -1 | 0 | 0 | 0 | -1 | -1 | 0 | 0 | 0 | 0 | -1 | -1 | 0 | 0 | -1 | -1 | -1 | 0 | 0 | -1 | -1 | -1 | 0 | -1 | -1 | -1 | -1 | -1 | 0 | -1 | 0 | -1 | -1 | -1 | -1 | 0 | -1 | -1 | 0 | 1 | 0 | 0 | 0 | 2 | 0 | -1 | 1 | 0 | 0 | 0 | 0 | 0 | 0 | 0 |
| Prom1 | chr5:44031733-44031853 | 0 | 0 | -1 | 0 | 0 | 0 | 0 | 0 | 0 | 0 | 0 | 0 | 0 | 1 | -1 | -1 | 0 | 1 | 40 | 0 | 13 | -1 | -1 | 0 | -1 | -1 | -1 | -1 | -1 | -1 | -1 | -1 | -1 | -1 | -1 | -1 | 0 | 0 | 0 | 0 | 0 | 0 | 0 | 0 | 0 | 0 | 0 | -1 | 0 | -1 | -1 | -1 | -1 | -1 | -1 | -1 | -1 | 0 | -1 | 0 | -1 | -1 | -1 | 0 | -1 | -1 | 0 | 0 | 0 | 0 | 0 | 0 | 0 | 0 | -1 | 0 | 0 | 0 | 0 | 0 | 0 |

#### SFig4, pg3

| Gene Symbol | Exon Location (mm10, mouse) | Brain | Skeletal Muscle | Heart | Lung | Stomach | Spleen | Colon | Liver | Kidney | Pancreas islet | Mammary gland | Ovary | Testes | Rod Photoreceptor (p28) | SST interneurons (cortex) | PV interneurons (hippocampus) | Motor neurons (Hb9+, embryo) | DRG neurons | Olfactory sensory neurons (Omp+) | Hair cells (Atoh1+, cochlea) | Purkinje cells (cerebellum) | Pyramidal neurons (Thy1+, cortex) | Pyramidal neurons (hippo - proximal CA1) | Pyramidal neurons (hippo - distal CA1) | Pyramidal neurons (hippo - superficial CA1) | Pyramidal neurons (hippo - dorsal CA1) | Pyramidal neurons (hippo - dorsal CA1, postsubiculum-projecting) | Pyramidal neurons (hippo - intermediate CA1) | Pyramidal neurons (hippo - ventral CA1) | Pyramidal neurons (hippo - ventral CA1, amygdala-projecting) | Pyramidal neurons (hippo - ventral CA1, nucleus accumbens-projecting | Pyramidal neurons (hippo - deep CA1) | Pyramidal neurons (hippo - dorsal CA3) | Pyramidal neurons (hippo - ventral CA3) | Granule cells (hippo - dorsal DG) | Granule cells (hippo - ventral DG) | Oligodendrocyte precursor cells (cortex) | Newly formed oligodendrocytes (cortex) | Myelinating oligodendrocytes (cortex) | Astrocytes (cortex) | Microglia (cortex) | Microglia (spinal cord) | Endothelial cells (cortex) | Quadriceps | Gastrocnemius | Heart ventricle (p90) | Hematopoietic stem cell (bone marrow) | Dendritic cells (spleen) | Neutrophil | Mast cell | NKT cell | NK cell | Macrophage | Eosinophils | B cell, spleen | CD4+ T-cell (Th17) | CD4+ T-cell (Th1) | CD4+ T-cell (Th2) | CD8+ T-cell | CD4+/CD8+ double positive T-cell | Gamma delta T-cell | Regulatory T-cell (Rag2+, spleen) | Hepatocyte (liver) | Liver sinusoidal endothelial cells (liver) | Kupffer cells (liver) | Hepatic stellate cells (liver) | Smooth muscle cells (colon) | Interstitial Cell of Cajal (colon) | Gastrointestinal PDGFRa+ cells (colon) | Gastrointestinal PDGFRa+ cells (colon mucosa) | Pancreatic alpha cells (7-16wks) | Pancreatic beta cells (7-16wks) | Alveolar type 1 (lung) | Podocytes (kidney) | Classical brown adipocyte | Inguinal white adipocyte | Sertoli cells (testis) | Leydig cells (testis) | Interstitial cells (testis) | Spermatogonial stem cells (testis) | Female germline stem cells (ovary) |
|---|---|---|---|---|---|---|---|---|---|---|---|---|---|---|---|---|---|---|---|---|---|---|---|---|---|---|---|---|---|---|---|---|---|---|---|---|---|---|---|---|---|---|---|---|---|---|---|---|---|---|---|---|---|---|---|---|---|---|---|---|---|---|---|---|---|---|---|---|---|---|---|---|---|---|---|---|---|---|---|---|---|---|
| Zfp207 | chr11:80396169-80396189 | 2 | -1 | 0 | 0 | 0 | 2 | 0 | 2 | 0 | 0 | 7 | 0 | 1 | 0 | 0 | 0 | 0 | 0 | 35 | 0 | 1 | -1 | 0 | 0 | 0 | 0 | 0 | 0 | 0 | 1 | 0 | 0 | 0 | 0 | 0 | 0 | 0 | 0 | 0 | 0 | 0 | 2 | 0 | 0 | 0 | 0 | 0 | 0 | -1 | 0 | 0 | 0 | 0 | 0 | 0 | 0 | 0 | 0 | -1 | 0 | -1 | 0 | -1 | 0 | -1 | 0 | 0 | 0 | 0 | 0 | 5 | 2 | 0 | 0 | 0 | 0 | 0 | 0 | 0 | 0 | 0 |
| Gm14022 | chr2:129061745-129061847 | 0 | -1 | -1 | -1 | -1 | -1 | 0 | -1 | -1 | -1 | -1 | -1 | 0 | -1 | -1 | -1 | 0 | -1 | 51 | 0 | 0 | -1 | -1 | -1 | 0 | -1 | -1 | -1 | -1 | 0 | -1 | -1 | -1 | -1 | -1 | -1 | 0 | 0 | 0 | 0 | -1 | 0 | 0 | -1 | 0 | 0 | -1 | -1 | -1 | -1 | -1 | -1 | -1 | -1 | -1 | -1 | -1 | -1 | -1 | -1 | -1 | -1 | -1 | -1 | -1 | -1 | -1 | 0 | -1 | -1 | -1 | -1 | -1 | -1 | -1 | 0 | -1 | 0 | -1 | 0 | -1 |
| Gm16794 | chr9:95944645-95944880\|95944848 | -1 | -1 | -1 | -1 | -1 | -1 | -1 | -1 | -1 | 8 | 0 | -1 | -1 | -1 | 0 | -1 | -1 | 0 | 45 | 0 | 0 | -1 | 0 | -1 | -1 | -1 | -1 | 0 | -1 | -1 | -1 | -1 | 0 | -1 | 0 | -1 | -1 | -1 | -1 | 0 | -1 | 0 | 0 | 0 | 0 | 0 | 0 | -1 | -1 | 0 | 0 | -1 | -1 | -1 | 0 | 0 | -1 | -1 | -1 | 7 | -1 | -1 | -1 | -1 | -1 | -1 | -1 | -1 | 0 | 0 | -1 | 0 | -1 | -1 | -1 | 0 | 0 | -1 | 0 | 0 | -1 |
| Unc79 | chr12:103022105-103022223 | 0 | -1 | -1 | -1 | -1 | -1 | -1 | -1 | -1 | 2 | -1 | 0 | 0 | 0 | -1 | 1 | 1 | -1 | 53 | 0 | 1 | 0 | 3 | 1 | 0 | 0 | 0 | 0 | 0 | -1 | 0 | 0 | 6 | 1 | 1 | 0 | 1 | 0 | -1 | 0 | -1 | 0 | -1 | 0 | -1 | -1 | -1 | -1 | -1 | -1 | -1 | -1 | -1 | -1 | -1 | -1 | -1 | -1 | -1 | -1 | -1 | -1 | -1 | -1 | -1 | -1 | -1 | 0 | 0 | 0 | 1 | 1 | -1 | 0 | -1 | -1 | 0 | 0 | 0 | 0 | 0 |
| Clstn2 | chr9:97690867-97690955 | -1 | -1 | -1 | -1 | -1 | -1 | -1 | -1 | -1 | -1 | -1 | -1 | -1 | -1 | -1 | -1 | 0 | -1 | 51 | -1 | -1 | -1 | -1 | -1 | -1 | 0 | -1 | -1 | -1 | -1 | -1 | -1 | -1 | -1 | -1 | -1 | 4 | 0 | 0 | -1 | 0 | -1 | -1 | -1 | -1 | -1 | -1 | -1 | -1 | -1 | -1 | -1 | -1 | -1 | -1 | -1 | -1 | -1 | -1 | -1 | -1 | -1 | -1 | -1 | -1 | -1 | -1 | -1 | -1 | -1 | -1 | -1 | -1 | -1 | -1 | -1 | -1 | -1 | 0 | -1 | -1 |
| Zfc3h1 | chr10:115428498-115428604 | 1 | 0 | 0 | 0 | 0 | 0 | 0 | 1 | 3 | 0 | 1 | 2 | 0 | 5 | 0 | 0 | 0 | 0 | 31 | 0 | 0 | 0 | 0 | 0 | 0 | 0 | 0 | 0 | 0 | 0 | 0 | 0 | 0 | 0 | 0 | 0 | 0 | 0 | 0 | 0 | 0 | 0 | 0 | 1 | 0 | 1 | 0 | 0 | 0 | 0 | 0 | 0 | 0 | 0 | 0 | 1 | 0 | 0 | 0 | 0 | 0 | 0 | 0 | 0 | 0 | 0 | 1 | 0 | 0 | 0 | 0 | 0 | 0 | 1 | 0 | 0 | 0 | 0 | 0 | 0 | 0 |
| Vars | chr17:35009796-35009994 | 1 | 1 | 0 | 0 | 0 | 0 | 0 | 0 | 0 | 0 | 0 | 0 | 0 | 2 | -1 | 0 | 1 | 0 | 22 | 0 | 2 | -1 | 0 | 0 | 0 | 0 | -1 | 0 | 0 | -1 | 0 | 0 | 0 | 0 | -1 | -1 | 3 | 2 | 2 | 2 | 1 | 0 | 0 | 0 | 0 | 0 | 0 | 0 | 0 | 0 | 0 | 0 | 0 | 0 | 0 | 0 | 0 | 0 | 0 | 1 | 0 | 0 | 0 | 0 | 0 | 0 | 0 | 0 | 0 | 0 | 0 | 0 | 0 | 1 | 0 | 0 | 0 | 0 | 0 | 0 | 0 |
| Skp2 | chr15:9114921-9115203 | 0 | 0 | 0 | 0 | 0 | 0 | 0 | 0 | 0 | 0 | 0 | 0 | 0 | 0 | 0 | 0 | 2 | -1 | 27 | 1 | 0 | -1 | 0 | 0 | 0 | 0 | -1 | -1 | -1 | -1 | 0 | -1 | 0 | 0 | -1 | 0 | 7 | 0 | 0 | 4 | 0 | 3 | 0 | 0 | 0 | 0 | 0 | 0 | 0 | 0 | 0 | 0 | 0 | 0 | 0 | 1 | 0 | 0 | 0 | 0 | -1 | 0 | 0 | 0 | 0 | 0 | 0 | 0 | 0 | 0 | -1 | 0 | 0 | 1 | -1 | 0 | 0 | 1 | 1 | 1 | 2 |
| Dmd | chrX:84405493-84405604 | 0 | 0 | 0 | -1 | -1 | 0 | 0 | -1 | -1 | 0 | 0 | -1 | 0 | 1 | 0 | 0 | 0 | 0 | 22 | -1 | 0 | -1 | 0 | 0 | 0 | 0 | 0 | 0 | -1 | 3 | 0 | 0 | 0 | 4 | 0 | 0 | 0 | 0 | 0 | 4 | -1 | 0 | -1 | 0 | 0 | 0 | -1 | -1 | -1 | -1 | -1 | -1 | -1 | -1 | -1 | -1 | -1 | -1 | -1 | -1 | -1 | -1 | -1 | -1 | -1 | -1 | 2 | 0 | 0 | 0 | -1 | -1 | -1 | -1 | 0 | -1 | -1 | -1 | -1 | -1 | 0 |
| Ttc23 | chr7:67711195-67711239 | 0 | 0 | -1 | 1 | 0 | 0 | 2 | 2 | 0 | 0 | 1 | 1 | 1 | -1 | -1 | -1 | -1 | 4 | 47 | 8 | 0 | -1 | -1 | -1 | -1 | -1 | -1 | -1 | -1 | -1 | -1 | -1 | 0 | -1 | -1 | -1 | 0 | -1 | 0 | 2 | 0 | 6 | 1 | 0 | 0 | 1 | 0 | -1 | -1 | 0 | -1 | -1 | -1 | 0 | 0 | 0 | 0 | -1 | -1 | 0 | -1 | 0 | 0 | 0 | -1 | 0 | 0 | 0 | 0 | 0 | 3 | 0 | 3 | 1 | 0 | 1 | 1 | 1 | 1 | 0 | 0 |
| Chn1 | chr2:73762063-73762227 | 3 | -1 | 0 | 0 | 0 | 0 | 0 | -1 | 0 | 1 | 0 | 0 | 0 | 0 | 0 | 0 | 1 | 1 | 23 | 0 | 0 | 0 | 0 | 0 | 0 | 0 | -1 | -1 | 0 | 0 | 0 | -1 | 0 | 0 | 0 | 0 | 1 | 0 | 0 | 0 | -1 | -1 | 0 | -1 | -1 | 0 | -1 | -1 | -1 | -1 | -1 | -1 | -1 | -1 | -1 | -1 | 0 | -1 | -1 | -1 | -1 | -1 | -1 | -1 | -1 | -1 | 0 | 0 | 0 | -1 | -1 | 0 | 0 | 0 | -1 | -1 | 0 | 0 | 2 | 0 | 0 |
| Agpat5 | chr8:18856262-18856385 | 0 | 0 | 0 | 0 | 0 | 1 | 0 | 0 | 0 | 0 | 1 | 0 | 3 | 1 | 0 | 0 | 1 | 0 | 25 | 0 | 0 | 0 | 0 | 0 | 0 | 0 | 0 | 0 | 0 | 0 | 0 | 0 | 0 | 1 | 0 | 0 | 0 | 0 | 0 | 1 | 0 | 0 | 3 | 0 | 0 | 0 | 0 | -1 | 0 | 0 | 0 | 0 | 0 | 1 | -1 | 1 | 1 | 1 | 0 | 1 | 0 | 0 | 0 | 0 | 0 | 0 | 0 | 0 | 0 | 1 | 0 | 0 | 1 | 0 | 0 | 0 | 1 | 0 | 1 | 1 | 1 |
| Mbnl3 | chrX:51126126-51126161 | 0 | 0 | -1 | 0 | 0 | 4 | 1 | 0 | 0 | 0 | 0 | 0 | -1 | -1 | -1 | -1 | -1 | -1 | 55 | 3 | -1 | -1 | -1 | -1 | -1 | -1 | -1 | -1 | -1 | -1 | -1 | -1 | -1 | -1 | -1 | -1 | -1 | -1 | -1 | 0 | -1 | 0 | 0 | 0 | -1 | 0 | 0 | 0 | -1 | 0 | 0 | 0 | -1 | 3 | 0 | 1 | 1 | 1 | 0 | 3 | -1 | 2 | -1 | 0 | 0 | -1 | -1 | 0 | -1 | 0 | -1 | 0 | 0 | 0 | 0 | -1 | 0 | 1 | 1 | 2 | 7 |
| Shpk | chr11:73206316-73206492 | 3 | 0 | -1 | -1 | 0 | 3 | 0 | 1 | 0 | 0 | 4 | -1 | -1 | 0 | -1 | 0 | 0 | 6 | 34 | 0 | 0 | -1 | 0 | -1 | -1 | -1 | -1 | 0 | -1 | -1 | -1 | 0 | -1 | -1 | 0 | -1 | 0 | -1 | 0 | -1 | 0 | 0 | 0 | -1 | 0 | 0 | 0 | 0 | 0 | -1 | -1 | 0 | 0 | 0 | 0 | 0 | 0 | 0 | 0 | 0 | -1 | 0 | 0 | 0 | -1 | 0 | 0 | -1 | -1 | -1 | 0 | 2 | 0 | 2 | 0 | 0 | 0 | 0 | 0 | 0 | 8 |
| Pgpep1l | chr7:68245773-68245915 | 0 | 0 | -1 | 0 | -1 | -1 | -1 | 0 | 0 | 0 | 0 | 0 | 0 | -1 | -1 | -1 | -1 | 0 | 26 | -1 | -1 | -1 | -1 | 0 | -1 | 0 | -1 | -1 | -1 | -1 | -1 | 0 | 0 | 0 | -1 | -1 | -1 | -1 | -1 | -1 | -1 | 0 | -1 | 1 | -1 | 0 | -1 | -1 | -1 | -1 | -1 | -1 | -1 | -1 | -1 | 0 | 6 | -1 | 5 | 0 | 0 | -1 | -1 | 0 | -1 | -1 | 0 | 0 | 0 | 0 | -1 | -1 | -1 | 2 | -1 | -1 | -1 | -1 | -1 | 0 | -1 |
| Nin | chr12:70079979-70080068 | 0 | 0 | 0 | 0 | -1 | 0 | -1 | 0 | 0 | 14 | 1 | 1 | 0 | 0 | -1 | 0 | 0 | 3 | 62 | 27 | 20 | -1 | 0 | 0 | 0 | 0 | 0 | 0 | 0 | 0 | 0 | 0 | 0 | 0 | 0 | 0 | 0 | 0 | 0 | 0 | 0 | 0 | 0 | 0 | 0 | 0 | 0 | 0 | 0 | 0 | 1 | 0 | 0 | 0 | 0 | 0 | 0 | 0 | 0 | 0 | 0 | -1 | 0 | 0 | 0 | 0 | 6 | 0 | 0 | 0 | 18 | 13 | 0 | 0 | 0 | 0 | 0 | 1 | 0 | 4 | 0 |
| Rab36 | chr10:75043031-75043158 | 0 | 0 | -1 | 0 | -1 | 0 | 0 | 0 | 0 | 1 | 0 | 0 | 2 | 0 | 0 | 0 | 0 | 2 | 27 | 3 | 7 | 0 | 0 | 0 | 0 | 0 | 0 | 0 | 0 | 0 | 0 | -1 | 0 | 0 | 0 | 0 | 6 | -1 | 0 | 1 | 0 | 0 | 0 | -1 | -1 | -1 | 0 | 0 | -1 | 0 | -1 | 0 | -1 | -1 | -1 | 0 | 0 | -1 | -1 | -1 | 0 | -1 | -1 | -1 | -1 | 0 | 0 | 1 | 0 | 0 | 1 | 0 | 0 | 2 | -1 | 0 | 0 | 0 | 1 | 12 | 1 |
| Slc30a7 | chr3:115958094-115958197 | 0 | 0 | 0 | 0 | -1 | 0 | 2 | 0 | 0 | 0 | 0 | 2 | 0 | -1 | -1 | 0 | 2 | 3 | 35 | 0 | 1 | 0 | -1 | 0 | 0 | 0 | 0 | 0 | 0 | 0 | 0 | 0 | 0 | 0 | 0 | -1 | 0 | 4 | -1 | 1 | 4 | 1 | 0 | 0 | 0 | 3 | 0 | -1 | 0 | 1 | 0 | 0 | -1 | 3 | 0 | 1 | 2 | 0 | 0 | 7 | 0 | 0 | 0 | 0 | 0 | 0 | 0 | 3 | 1 | 0 | 0 | 1 | 1 | 1 | 0 | 0 | 0 | 0 | 1 | 1 | 0 |
| Mid2 | chrX:140736124-140736333 | 3 | 0 | 0 | 0 | -1 | 0 | 0 | -1 | 0 | 0 | 2 | -1 | 0 | 0 | -1 | 1 | 1 | 1 | 58 | 0 | 1 | -1 | -1 | 0 | 1 | 0 | 0 | 0 | 0 | 0 | 0 | 0 | 0 | 0 | 3 | 0 | 0 | -1 | 0 | 8 | -1 | 0 | 0 | 2 | -1 | 1 | 0 | 0 | -1 | -1 | -1 | -1 | -1 | -1 | -1 | -1 | 0 | -1 | -1 | -1 | -1 | -1 | 0 | 0 | -1 | 0 | 4 | 3 | 3 | 1 | 0 | 0 | 0 | 1 | 0 | 2 | 1 | 3 | -1 | 1 | 10 |
| 4930578M01Rik | chr15:98985543-98985619 | -1 | -1 | -1 | -1 | -1 | -1 | 0 | -1 | -1 | 5 | -1 | -1 | 0 | -1 | -1 | -1 | -1 | -1 | 40 | -1 | 0 | -1 | 0 | 0 | -1 | 0 | -1 | -1 | -1 | -1 | -1 | -1 | 0 | 0 | -1 | -1 | -1 | -1 | -1 | -1 | -1 | -1 | -1 | -1 | -1 | -1 | -1 | -1 | -1 | -1 | -1 | -1 | -1 | -1 | -1 | -1 | -1 | -1 | -1 | -1 | -1 | -1 | -1 | -1 | -1 | -1 | -1 | -1 | -1 | -1 | -1 | 6 | -1 | -1 | -1 | -1 | -1 | -1 | 0 | -1 | -1 |
| 2810039B14Rik | chr15:75643553\|75643130-75643601 | -1 | -1 | 0 | -1 | -1 | -1 | -1 | -1 | -1 | -1 | -1 | 2 | -1 | -1 | -1 | -1 | -1 | -1 | 24 | -1 | 8 | -1 | -1 | -1 | -1 | -1 | -1 | -1 | -1 | -1 | -1 | -1 | -1 | -1 | -1 | -1 | -1 | -1 | -1 | -1 | 0 | -1 | -1 | -1 | 0 | 0 | -1 | -1 | -1 | -1 | -1 | -1 | -1 | 0 | -1 | -1 | -1 | -1 | -1 | -1 | -1 | -1 | 0 | -1 | 0 | -1 | -1 | 0 | -1 | 0 | -1 | -1 | -1 | -1 | -1 | -1 | -1 | -1 | -1 | -1 | -1 |
| Gm44696 | chr6:48376270-48376427 | 0 | -1 | 0 | -1 | 0 | -1 | -1 | -1 | -1 | 0 | -1 | -1 | 21 | -1 | 0 | -1 | -1 | -1 | 95 | 0 | 9 | -1 | -1 | -1 | -1 | 0 | -1 | -1 | -1 | -1 | -1 | -1 | -1 | 0 | -1 | -1 | -1 | -1 | -1 | -1 | -1 | -1 | -1 | 0 | 0 | -1 | 0 | -1 | -1 | -1 | -1 | -1 | -1 | -1 | -1 | -1 | -1 | -1 | 0 | -1 | -1 | -1 | -1 | -1 | -1 | -1 | -1 | -1 | 0 | -1 | 0 | 0 | -1 | -1 | -1 | -1 | -1 | -1 | -1 | -1 | 25 |
| Slc1a2 | chr2:102780717-102780787 | 1 | 0 | -1 | -1 | -1 | 0 | 0 | 0 | 0 | 0 | 0 | 0 | 27 | -1 | -1 | 0 | 16 | -1 | 94 | 0 | 1 | 0 | 6 | 1 | 0 | 9 | 0 | 4 | 1 | 8 | 16 | 2 | 6 | 5 | 5 | 0 | 0 | 0 | 1 | 1 | 0 | 0 | 0 | -1 | -1 | 0 | -1 | -1 | -1 | -1 | -1 | -1 | -1 | -1 | -1 | -1 | -1 | -1 | -1 | -1 | -1 | -1 | 0 | -1 | 0 | 0 | -1 | -1 | -1 | -1 | -1 | -1 | -1 | 0 | -1 | -1 | 0 | 0 | 0 | -1 | -1 |
| Spata6 | chr4:111749539-111749643 | -1 | 0 | 0 | 0 | 0 | 2 | 0 | 0 | 0 | 1 | 0 | 0 | 0 | -1 | -1 | -1 | 0 | 1 | 20 | 1 | 0 | -1 | 0 | -1 | 0 | -1 | -1 | -1 | 0 | 0 | 0 | 0 | -1 | -1 | -1 | -1 | 0 | 0 | 0 | 0 | 0 | 0 | 1 | 0 | 0 | 0 | 0 | 0 | 0 | -1 | 0 | 0 | 0 | 0 | 0 | 0 | 1 | 1 | 0 | 2 | 0 | 0 | -1 | 0 | 0 | 0 | 0 | 0 | 0 | 0 | 0 | 2 | 0 | 1 | 0 | 0 | 2 | 4 | 2 | 0 | 0 |
| Myh10 | chr11:68745935-68745964 | 15 | 0 | 0 | 2 | -1 | 0 | -1 | 0 | 0 | 0 | 0 | 0 | 1 | 50 | -1 | 14 | 0 | 31 | 70 | 33 | 6 | 0 | 2 | 1 | 0 | 1 | -1 | 1 | 0 | 0 | 0 | 0 | 0 | 2 | 2 | 0 | 0 | 0 | 0 | 0 | 0 | 0 | 0 | 0 | -1 | 0 | 0 | 0 | -1 | -1 | 0 | 0 | -1 | -1 | -1 | 0 | 0 | 0 | -1 | 0 | 0 | -1 | 0 | 0 | 0 | 0 | 0 | 0 | 0 | 0 | 0 | 2 | 0 | 0 | 0 | 0 | 0 | 0 | 0 | 0 | 0 |
| Arhgef12 | chr9:43083516-43083678 | 0 | 0 | 0 | 0 | -1 | 2 | 1 | 0 | 1 | 0 | 0 | 0 | 1 | 1 | 0 | 2 | 2 | 1 | 23 | 3 | 1 | 0 | 1 | 1 | 1 | 5 | 3 | 1 | 0 | 1 | 4 | 1 | 0 | 0 | 1 | -1 | 0 | 1 | 1 | 0 | 0 | 0 | 1 | 0 | 1 | 0 | 0 | 0 | 0 | 0 | 0 | -1 | 0 | -1 | 0 | 1 | 0 | 0 | 0 | 0 | -1 | -1 | 0 | 0 | 0 | 0 | 0 | 1 | 0 | 1 | 1 | 0 | 0 | 1 | 1 | 2 | 1 | 1 | 1 | 1 | 0 |
| Pgpep1l | chr7:68262158-68262349\|68262336 | -1 | -1 | -1 | -1 | -1 | -1 | -1 | -1 | -1 | -1 | -1 | -1 | -1 | -1 | -1 | -1 | -1 | -1 | 71 | -1 | -1 | -1 | 0 | -1 | -1 | -1 | -1 | -1 | -1 | 0 | -1 | -1 | 0 | 0 | -1 | -1 | -1 | -1 | -1 | -1 | -1 | -1 | -1 | -1 | -1 | -1 | 0 | -1 | -1 | -1 | -1 | -1 | -1 | -1 | -1 | -1 | -1 | -1 | 0 | -1 | -1 | -1 | -1 | -1 | -1 | 0 | -1 | -1 | -1 | -1 | -1 | -1 | 0 | 19 | -1 | -1 | -1 | -1 | -1 | -1 | -1 |
| Atp5g1 | chr11:96074589-96074761 | 1 | 0 | 0 | 0 | 1 | 2 | 1 | 0 | 1 | 0 | -1 | 0 | -1 | 1 | 1 | 0 | 0 | 0 | 22 | 0 | 0 | 0 | 0 | 0 | 0 | 0 | 0 | 0 | 0 | 0 | 0 | 0 | 0 | 0 | 0 | 0 | 4 | 6 | 1 | 4 | 0 | 0 | 2 | 0 | 0 | 0 | 0 | -1 | -1 | 2 | -1 | -1 | -1 | 1 | -1 | 1 | 2 | 4 | -1 | 1 | -1 | 0 | 0 | -1 | -1 | 0 | 2 | 0 | 0 | 0 | -1 | -1 | 0 | 0 | 0 | 2 | 0 | 0 | 0 | 1 | 0 |
| Zfp950 | chr19:61121615-61122027 | 9 | -1 | -1 | 0 | 0 | 1 | 0 | 3 | 0 | 0 | 1 | 0 | 0 | -1 | 12 | 0 | 0 | 0 | 39 | 2 | 0 | 0 | 0 | 0 | 0 | 0 | 0 | 0 | -1 | 0 | -1 | 0 | 0 | 0 | 0 | 0 | 11 | -1 | -1 | -1 | 0 | 0 | 4 | 0 | 0 | 0 | 0 | -1 | -1 | -1 | 0 | 0 | 0 | 0 | 0 | 0 | 0 | 0 | 0 | 0 | 0 | 0 | 0 | 0 | 2 | 0 | 2 | -1 | 0 | 0 | 2 | -1 | 1 | 0 | 0 | 0 | 1 | 1 | 23 | 16 |
| Ahcy | chr2:155069759-155069825 | 0 | 0 | 0 | -1 | -1 | -1 | -1 | 54 | -1 | 13 | -1 | -1 | -1 | -1 | -1 | -1 | -1 | 6 | 68 | 2 | 4 | 0 | -1 | -1 | 0 | -1 | -1 | -1 | -1 | 0 | -1 | 0 | -1 | -1 | -1 | -1 | 0 | -1 | -1 | 0 | 0 | -1 | 0 | 0 | -1 | 0 | 0 | -1 | -1 | -1 | -1 | -1 | -1 | 0 | -1 | 4 | -1 | -1 | -1 | 14 | -1 | -1 | -1 | -1 | -1 | -1 | 0 | 0 | 0 | 0 | -1 | -1 | -1 | -1 | -1 | 0 | -1 | -1 | -1 | -1 | 2 |
| Zfp229 | chr17:21734799-21735601 | 0 | 0 | 0 | 0 | -1 | -1 | -1 | 0 | -1 | 1 | -1 | -1 | -1 | 0 | -1 | 0 | -1 | 1 | 21 | 2 | 0 | -1 | -1 | -1 | -1 | -1 | -1 | -1 | 0 | 0 | -1 | -1 | -1 | 0 | -1 | -1 | -1 | 0 | 0 | 0 | -1 | 0 | 0 | 1 | 0 | 0 | 0 | 0 | -1 | 0 | -1 | -1 | -1 | -1 | -1 | 0 | -1 | -1 | -1 | 8 | -1 | -1 | -1 | 0 | -1 | 0 | 0 | 0 | -1 | 0 | 1 | 0 | -1 | 2 | 0 | -1 | 1 | 3 | 9 | 2 | 6 |
| BC065397 | chrX:136765160-136765764 | -1 | -1 | -1 | -1 | -1 | -1 | -1 | -1 | -1 | 7 | -1 | -1 | 9 | -1 | -1 | -1 | 4 | -1 | 91 | -1 | -1 | 0 | -1 | -1 | -1 | 0 | -1 | 0 | 0 | 0 | 0 | 0 | 0 | -1 | 0 | 0 | -1 | 13 | 0 | 6 | -1 | -1 | -1 | -1 | 0 | 0 | -1 | 0 | -1 | -1 | 4 | -1 | -1 | 0 | 0 | -1 | -1 | -1 | 0 | 0 | -1 | -1 | 0 | -1 | 0 | 0 | -1 | 9 | -1 | -1 | 7 | -1 | -1 | 6 | 0 | -1 | 5 | 10 | 10 | 8 | 6 |
| Olfr558 | chr7:102707381-102707469 | -1 | 0 | 0 | -1 | -1 | 0 | -1 | -1 | -1 | 0 | -1 | 0 | -1 | -1 | -1 | -1 | -1 | 0 | 26 | -1 | -1 | -1 | -1 | -1 | -1 | -1 | -1 | -1 | -1 | -1 | -1 | -1 | -1 | -1 | -1 | -1 | -1 | -1 | -1 | -1 | -1 | -1 | -1 | -1 | 0 | 5 | -1 | -1 | -1 | -1 | -1 | -1 | -1 | -1 | -1 | -1 | -1 | -1 | -1 | -1 | -1 | -1 | -1 | -1 | -1 | -1 | -1 | -1 | -1 | -1 | -1 | -1 | -1 | 5 | -1 | -1 | -1 | -1 | -1 | -1 | -1 |
| Cep57l1 | chr10:41802212-41802346 | 1 | -1 | 0 | 0 | -1 | 2 | 0 | 4 | 2 | 2 | -1 | 0 | 1 | -1 | -1 | 0 | 0 | 0 | 28 | 3 | 3 | -1 | 0 | 0 | 0 | 0 | -1 | 0 | 0 | 0 | 0 | 0 | 0 | 0 | 0 | 0 | 1 | 0 | -1 | -1 | -1 | 0 | 3 | 0 | -1 | 0 | -1 | -1 | -1 | 0 | 0 | 0 | -1 | 13 | 2 | 7 | 2 | 3 | 0 | 1 | 0 | -1 | 0 | 0 | 0 | 0 | -1 | 0 | 0 | 0 | 4 | 0 | -1 | 2 | 0 | -1 | 3 | 4 | 3 | 2 | 4 |
| Mgll | chr6:88813880-88813963 | 0 | 0 | 0 | 2 | 34 | 0 | 10 | 7 | 5 | 16 | 11 | 0 | 0 | 0 | 0 | 0 | 0 | 0 | 35 | 0 | 0 | 0 | 0 | 0 | 0 | 0 | 0 | 0 | 0 | 0 | 0 | 0 | 0 | 0 | 0 | 0 | 0 | 0 | 0 | 0 | 0 | 1 | 0 | 0 | 0 | 0 | -1 | 0 | 0 | 0 | 0 | 0 | -1 | 0 | 0 | 0 | 1 | 0 | 0 | 0 | 0 | 0 | 2 | 0 | 0 | 1 | 0 | 0 | 0 | 2 | -1 | 0 | -1 | 0 | 0 | 0 | 0 | 0 | 0 | 1 | 0 |
| Limch1 | chr5:67025305-67025373 | 0 | 0 | 1 | 15 | 0 | 0 | 0 | 0 | 14 | 26 | 2 | 0 | 0 | 0 | 0 | 0 | 0 | 0 | 69 | 22 | 0 | -1 | 0 | 0 | 0 | 0 | 0 | 0 | 0 | 0 | 0 | 1 | 0 | 0 | 0 | 0 | 0 | 0 | 0 | 0 | 0 | 0 | 0 | 0 | 0 | 0 | -1 | -1 | -1 | -1 | -1 | -1 | -1 | -1 | -1 | -1 | -1 | -1 | -1 | -1 | -1 | -1 | -1 | 0 | -1 | 0 | 0 | 0 | 0 | 0 | -1 | 28 | 19 | 0 | -1 | 0 | 1 | 0 | 0 | -1 | -1 |
| Pcdh15 | chr10:74321438-74321458 | 0 | -1 | 0 | -1 | -1 | 0 | 0 | -1 | -1 | 0 | 0 | 0 | -1 | 0 | -1 | 0 | 0 | 0 | -1 | 55 | 0 | 0 | -1 | -1 | -1 | -1 | -1 | 0 | -1 | 0 | 0 | -1 | -1 | -1 | -1 | -1 | 0 | 0 | 0 | -1 | -1 | -1 | -1 | -1 | -1 | -1 | -1 | -1 | -1 | -1 | -1 | -1 | -1 | -1 | -1 | -1 | -1 | -1 | -1 | -1 | -1 | -1 | -1 | -1 | -1 | -1 | -1 | -1 | -1 | -1 | 0 | 1 | -1 | -1 | -1 | -1 | 0 | 0 | 0 | 0 | 0 |
| Pclo | chr5:14764297-14764349 | 0 | 0 | -1 | 0 | -1 | -1 | 0 | 0 | 0 | 0 | -1 | 0 | 0 | 0 | 0 | 0 | 0 | 0 | 0 | 26 | 0 | 0 | 0 | 0 | 0 | 0 | 0 | 0 | 0 | 0 | 0 | 0 | 0 | 0 | 0 | 0 | 0 | -1 | 0 | 0 | 0 | 0 | -1 | -1 | -1 | -1 | 0 | -1 | -1 | -1 | -1 | -1 | -1 | -1 | -1 | -1 | 0 | -1 | -1 | -1 | -1 | -1 | -1 | -1 | -1 | 0 | 0 | 0 | -1 | -1 | 0 | 0 | -1 | 1 | -1 | -1 | 0 | 0 | 0 | 0 | 0 |
| Sptan1 | chr2:29982166-29982234 | 0 | 0 | 2 | 0 | 0 | 0 | 0 | 0 | 0 | 0 | 0 | 0 | 0 | 0 | 0 | 0 | 1 | 0 | 0 | 85 | 0 | 0 | 0 | 0 | 0 | 1 | 0 | 0 | 0 | 0 | 0 | 0 | 1 | 1 | 1 | 0 | 0 | 0 | 0 | 0 | 0 | 0 | 0 | 0 | 0 | 0 | 0 | 0 | 0 | 0 | 0 | 0 | 0 | 0 | 0 | 0 | 0 | 0 | 0 | 0 | 0 | 0 | 0 | 0 | 0 | 0 | 0 | 0 | 0 | 0 | 0 | 0 | 0 | 0 | 0 | 0 | 0 | 0 | 0 | 0 | 0 |
| Eps8 | chr6:137530542-137530592 | 0 | 0 | 0 | 0 | 0 | 0 | 0 | 0 | 0 | 1 | 0 | 0 | 0 | 0 | -1 | -1 | 0 | 0 | -1 | 29 | 0 | -1 | -1 | -1 | 0 | -1 | -1 | 0 | -1 | -1 | 0 | 0 | 0 | 0 | 0 | -1 | 0 | 0 | 0 | 0 | 0 | 0 | 0 | 0 | 0 | 0 | 0 | 0 | 0 | 0 | 0 | -1 | 0 | 0 | 0 | 0 | 0 | 0 | -1 | 0 | -1 | -1 | -1 | 0 | -1 | 0 | 0 | 0 | 0 | 0 | 0 | 0 | 0 | 0 | 0 | 0 | 0 | 0 | 0 | 0 | 0 |
| Cyfip2 | chr11:46228486-46228551 | 0 | 0 | 1 | 0 | 0 | 0 | 0 | 0 | 0 | 0 | 0 | 0 | 0 | 0 | 0 | 0 | 0 | 0 | 0 | 56 | 0 | 3 | 0 | 0 | 0 | 0 | 0 | 0 | 0 | 0 | 0 | 0 | 0 | 0 | 0 | 0 | 0 | 0 | 0 | 0 | 0 | 0 | 0 | 0 | 0 | 0 | 0 | 0 | 0 | 0 | 0 | 0 | -1 | 0 | 0 | 0 | 0 | 0 | 0 | 0 | 0 | 0 | -1 | 0 | 0 | 0 | 0 | 0 | 0 | 0 | 0 | 0 | 0 | 0 | -1 | -1 | 0 | 0 | 0 | 3 | 0 |
| Erc2 | chr14:28039421-28039450 | 0 | -1 | -1 | 0 | -1 | -1 | -1 | 0 | 0 | 0 | -1 | 0 | 0 | 0 | 0 | 0 | 0 | 0 | -1 | 37 | 0 | -1 | 0 | 3 | 0 | 1 | 0 | 1 | 1 | 0 | 0 | 1 | 0 | 1 | 0 | 2 | 0 | 0 | 0 | 0 | -1 | 0 | 0 | 0 | 0 | 0 | 0 | -1 | -1 | -1 | -1 | -1 | -1 | -1 | -1 | -1 | -1 | -1 | -1 | -1 | -1 | -1 | 0 | -1 | -1 | -1 | 0 | 0 | 0 | -1 | 1 | 1 | -1 | 0 | 0 | -1 | -1 | 0 | 0 | 0 | 0 |
| Fam49b | chr15:64023096-64023244 | 0 | 0 | 0 | 0 | 0 | 0 | 0 | 0 | 0 | 0 | 0 | 0 | 0 | -1 | 0 | 0 | 0 | 0 | -1 | 28 | 0 | 0 | 2 | 4 | 0 | 0 | 2 | 0 | 0 | 0 | 0 | 0 | 0 | 0 | 0 | 0 | 1 | 0 | 0 | 0 | 0 | -1 | 1 | 0 | 0 | 0 | 0 | 0 | 0 | 0 | 0 | 0 | 0 | 0 | 0 | 0 | 0 | 0 | 0 | 0 | 0 | -1 | -1 | 0 | 0 | 0 | 0 | 0 | 0 | 0 | 1 | 0 | 0 | 0 | -1 | -1 | 0 | 0 | 0 | 1 | 3 |
| Ap3b2 | chr7:81466031-81466087 | 1 | 0 | -1 | -1 | 0 | -1 | 0 | -1 | -1 | 0 | 0 | 0 | 0 | 0 | 0 | 0 | 0 | 0 | 0 | 71 | 0 | 1 | 0 | 0 | 0 | 0 | 0 | 0 | 3 | 0 | 0 | 0 | 0 | 0 | 1 | 0 | 1 | 0 | 0 | 6 | 0 | 1 | 0 | 0 | -1 | -1 | -1 | -1 | -1 | -1 | -1 | -1 | -1 | -1 | -1 | -1 | -1 | -1 | -1 | -1 | -1 | -1 | -1 | -1 | -1 | -1 | 0 | 0 | 0 | -1 | 1 | 1 | -1 | -1 | -1 | -1 | 0 | -1 | 2 | 0 | 1 |
| Cdh23 | chr10:60304424-60304528 | 0 | 0 | -1 | -1 | -1 | -1 | -1 | -1 | -1 | -1 | 0 | -1 | 9 | -1 | -1 | -1 | 0 | -1 | 0 | 95 | -1 | -1 | -1 | -1 | -1 | -1 | -1 | -1 | -1 | -1 | -1 | -1 | -1 | -1 | -1 | -1 | 0 | -1 | -1 | 0 | 0 | -1 | -1 | 0 | 0 | 0 | -1 | -1 | -1 | -1 | -1 | -1 | -1 | -1 | -1 | 9 | -1 | 0 | -1 | 0 | -1 | -1 | -1 | 0 | -1 | -1 | -1 | -1 | -1 | -1 | 0 | 0 | -1 | 0 | -1 | -1 | 3 | 2 | 0 | 4 | -1 |
| Adgrv1 | chr13:81570483-81570515 | -1 | -1 | -1 | -1 | -1 | -1 | -1 | -1 | -1 | 3 | -1 | 1 | -1 | 8 | -1 | -1 | -1 | -1 | -1 | 97 | -1 | -1 | -1 | -1 | -1 | -1 | -1 | -1 | -1 | -1 | -1 | -1 | -1 | -1 | -1 | -1 | -1 | 0 | -1 | 0 | -1 | -1 | -1 | -1 | -1 | -1 | -1 | -1 | -1 | -1 | -1 | -1 | -1 | -1 | -1 | -1 | -1 | -1 | -1 | -1 | -1 | -1 | -1 | -1 | -1 | -1 | -1 | 0 | -1 | -1 | 0 | 0 | -1 | -1 | -1 | -1 | -1 | -1 | -1 | -1 | 0 |
| Cacna1d | chr14:30137041-30137124 | 4 | -1 | -1 | 0 | 0 | 0 | 0 | 0 | 0 | 4 | 0 | 0 | 0 | -1 | 0 | 0 | 0 | 0 | 0 | 85 | 2 | 0 | -1 | 0 | 0 | 4 | -1 | -1 | -1 | 0 | 0 | -1 | 0 | 10 | 1 | 0 | 0 | 0 | -1 | 0 | 0 | 0 | 0 | -1 | -1 | -1 | 0 | -1 | -1 | -1 | -1 | -1 | -1 | -1 | -1 | -1 | -1 | -1 | -1 | -1 | -1 | -1 | 0 | -1 | -1 | -1 | 0 | 1 | 0 | 0 | 3 | 3 | 0 | 0 | -1 | -1 | 0 | 0 | 0 | 5 | 3 |
| Rasa2 | chr9:96577131-96577143 | 0 | -1 | 0 | 0 | 0 | 0 | 0 | 0 | 0 | 0 | 0 | 0 | 0 | 0 | -1 | 0 | 0 | 0 | 0 | 63 | 0 | 0 | 0 | 0 | 0 | 0 | 0 | 0 | 0 | -1 | -1 | 0 | 0 | 0 | -1 | 0 | 0 | 0 | 0 | 0 | 0 | 0 | -1 | -1 | 15 | 0 | 0 | 0 | 0 | 0 | 0 | 0 | 0 | 0 | 0 | 0 | 0 | 0 | 0 | 0 | 0 | -1 | 0 | -1 | 0 | 0 | 0 | 0 | 0 | 0 | 0 | 0 | 0 | -1 | 0 | 0 | 0 | 0 | 0 | 0 |
| C530008M17Rik | chr5:76770235-76770316 | 0 | -1 | -1 | 0 | -1 | -1 | -1 | -1 | -1 | 0 | 0 | 0 | 0 | -1 | 0 | 0 | 1 | 0 | 9 | 25 | 0 | -1 | 0 | -1 | 0 | 0 | 0 | 0 | 0 | 0 | 0 | 0 | -1 | 1 | -1 | 0 | 0 | 3 | 0 | 0 | -1 | 0 | 0 | -1 | -1 | 0 | 1 | -1 | -1 | 0 | -1 | -1 | -1 | -1 | -1 | 0 | -1 | -1 | -1 | -1 | -1 | -1 | -1 | -1 | -1 | -1 | 0 | -1 | 0 | -1 | -1 | -1 | 0 | -1 | -1 | 0 | 1 | 1 | 0 | 0 | -1 |
| Nwd1 | chr8:72668322-72668351 | 7 | -1 | -1 | -1 | -1 | -1 | -1 | -1 | -1 | 0 | -1 | 0 | 0 | 0 | -1 | -1 | -1 | 4 | 4 | 50 | 8 | -1 | -1 | -1 | -1 | -1 | -1 | -1 | -1 | -1 | -1 | -1 | -1 | -1 | -1 | -1 | 0 | -1 | 0 | 3 | 0 | 0 | -1 | -1 | -1 | -1 | 0 | -1 | -1 | -1 | -1 | -1 | -1 | -1 | -1 | -1 | -1 | -1 | -1 | 2 | -1 | -1 | -1 | -1 | -1 | -1 | 0 | -1 | -1 | -1 | 0 | -1 | 0 | 0 | -1 | -1 | 0 | 0 | -1 | 0 | -1 |
| Cfap44 | chr16:44461458-44461709 | -1 | -1 | -1 | 9 | -1 | -1 | -1 | -1 | -1 | -1 | -1 | 0 | 1 | -1 | -1 | -1 | -1 | -1 | -1 | 76 | -1 | -1 | -1 | -1 | 0 | -1 | -1 | -1 | -1 | -1 | -1 | -1 | -1 | -1 | -1 | -1 | -1 | -1 | -1 | -1 | -1 | -1 | -1 | -1 | -1 | -1 | -1 | -1 | -1 | -1 | -1 | -1 | -1 | -1 | -1 | -1 | -1 | -1 | -1 | -1 | -1 | -1 | -1 | -1 | -1 | -1 | -1 | -1 | -1 | -1 | -1 | -1 | -1 | -1 | -1 | -1 | -1 | -1 | -1 | -1 | 0 |
| Nuak2 | chr1:132326206-132326229 | -1 | 0 | 0 | 1 | 0 | 2 | 1 | 0 | 1 | 0 | 1 | 0 | 0 | -1 | -1 | -1 | 23 | -1 | 2 | 31 | -1 | -1 | -1 | -1 | -1 | -1 | -1 | -1 | -1 | -1 | -1 | -1 | -1 | -1 | 0 | -1 | 0 | -1 | -1 | 0 | 0 | 0 | 0 | 0 | 0 | 0 | 0 | 0 | 0 | 1 | 0 | 0 | 0 | 0 | -1 | 0 | 0 | 0 | -1 | 0 | -1 | 0 | 0 | 1 | 0 | 0 | -1 | 0 | 0 | 1 | 1 | 1 | 1 | 0 | -1 | 0 | 0 | 0 | 0 | 0 | 0 |
| Rimbp2 | chr5:128811983-128812003 | 2 | 0 | -1 | -1 | -1 | -1 | 0 | -1 | -1 | 0 | -1 | 0 | 0 | -1 | 0 | 0 | 12 | 4 | -1 | 27 | 0 | 0 | 0 | 6 | 0 | 0 | 0 | 0 | 0 | 0 | 0 | 0 | 1 | 0 | 0 | 0 | 0 | 0 | 0 | 0 | 0 | -1 | -1 | -1 | -1 | -1 | -1 | -1 | -1 | -1 | -1 | -1 | -1 | -1 | -1 | 0 | -1 | -1 | -1 | -1 | -1 | -1 | -1 | -1 | -1 | -1 | 0 | 0 | 0 | 0 | 0 | 0 | -1 | 0 | -1 | -1 | 0 | 0 | 0 | 0 | -1 |
| Vps13d | chr4:145136680-145136727 | 2 | 2 | -1 | 0 | 0 | 0 | 0 | 0 | 0 | 1 | 1 | 0 | 0 | 0 | -1 | 0 | 0 | 1 | 0 | 21 | 0 | -1 | 3 | 0 | 0 | 0 | -1 | 0 | 0 | 0 | -1 | -1 | 0 | -1 | -1 | 0 | 0 | 0 | 0 | 0 | 0 | 3 | 0 | 1 | 2 | 1 | 0 | -1 | 0 | 0 | 1 | 0 | 0 | 0 | 0 | 0 | 1 | 0 | 0 | 0 | 0 | -1 | 0 | 0 | 0 | 0 | 1 | 1 | 0 | 0 | 1 | 0 | 0 | 0 | 0 | 4 | 0 | 0 | 0 | 1 | 0 |
| Piezo2 | chr18:63063324-63063422 | -1 | 0 | 0 | 0 | -1 | 0 | -1 | 0 | 0 | 1 | 16 | 0 | 0 | -1 | -1 | -1 | -1 | 54 | -1 | 99 | -1 | -1 | -1 | -1 | -1 | -1 | -1 | -1 | -1 | -1 | -1 | -1 | -1 | -1 | -1 | -1 | -1 | 0 | 0 | 0 | -1 | 0 | -1 | 0 | 0 | 0 | -1 | 0 | -1 | -1 | -1 | -1 | -1 | -1 | -1 | -1 | -1 | -1 | -1 | -1 | -1 | -1 | -1 | 0 | -1 | -1 | -1 | 3 | 1 | 7 | -1 | -1 | 0 | 0 | -1 | 0 | 0 | 2 | 1 | -1 | -1 |
| Mindy4b-ps | chr3:58809718-58809783 | -1 | -1 | -1 | -1 | -1 | -1 | -1 | -1 | -1 | -1 | -1 | -1 | -1 | 0 | -1 | -1 | -1 | -1 | -1 | 83 | 6 | -1 | -1 | -1 | -1 | -1 | -1 | -1 | -1 | -1 | -1 | -1 | -1 | -1 | -1 | -1 | -1 | -1 | -1 | -1 | -1 | 0 | -1 | -1 | -1 | -1 | -1 | -1 | -1 | -1 | -1 | -1 | -1 | -1 | -1 | -1 | -1 | -1 | -1 | -1 | -1 | -1 | -1 | -1 | -1 | -1 | -1 | -1 | -1 | -1 | -1 | -1 | -1 | -1 | -1 | -1 | -1 | -1 | -1 | -1 | -1 |
| Sptan1 | chr2:29968776-29968826 | 5 | 11 | 4 | 0 | 0 | 0 | 0 | 2 | 0 | 0 | 0 | 0 | 0 | 0 | -1 | 25 | 0 | 2 | 0 | 46 | 11 | 0 | 1 | 1 | 0 | 2 | 1 | 1 | 0 | 0 | 1 | 5 | 2 | 4 | 1 | 0 | 0 | 0 | 0 | 0 | 0 | -1 | 0 | 4 | 6 | 4 | 0 | 0 | 0 | 1 | 0 | 0 | -1 | 3 | 0 | 0 | 0 | 0 | 0 | 0 | 0 | -1 | 0 | 1 | 0 | -1 | 0 | 0 | 0 | 0 | 0 | 0 | 0 | 0 | 0 | 0 | 0 | 0 | 0 | 0 | 0 |
| Ddx11 | chr17:66133436-66133522 | -1 | -1 | -1 | 0 | 0 | 3 | -1 | 1 | -1 | 0 | 6 | 1 | 1 | -1 | -1 | -1 | -1 | 0 | 0 | 77 | 0 | -1 | -1 | -1 | -1 | 0 | 0 | -1 | -1 | -1 | -1 | -1 | -1 | -1 | -1 | -1 | 0 | 0 | -1 | 0 | 0 | -1 | 0 | -1 | 0 | 26 | 0 | 11 | -1 | 0 | 8 | 4 | 0 | 0 | 6 | 1 | 2 | 2 | 4 | 0 | 2 | 0 | -1 | 0 | -1 | 0 | -1 | 0 | 0 | 0 | -1 | 0 | -1 | 1 | -1 | -1 | 0 | 1 | 2 | 5 | 1 |
| Pnpla3 | chr15:84178854-84179057 | 3 | 4 | 0 | 0 | 0 | 0 | 0 | 0 | 3 | 2 | 0 | -1 | 0 | 0 | -1 | 0 | 0 | 5 | 4 | 24 | 0 | -1 | 0 | 0 | 0 | -1 | 0 | -1 | 0 | 0 | -1 | 0 | 0 | 0 | 0 | -1 | -1 | 1 | -1 | 5 | 0 | 0 | 0 | -1 | 8 | -1 | 0 | -1 | -1 | -1 | -1 | -1 | -1 | -1 | -1 | 0 | -1 | -1 | -1 | -1 | -1 | -1 | -1 | 0 | -1 | 0 | 0 | 0 | -1 | -1 | 0 | 0 | -1 | 0 | 1 | 1 | 0 | 0 | 0 | 0 | -1 |
| Cep135 | chr5:76599326-76599419 | 8 | 0 | -1 | 0 | -1 | 0 | 0 | 0 | 0 | 1 | 0 | 1 | 1 | -1 | -1 | 0 | 0 | -1 | 0 | 22 | 5 | -1 | 0 | 0 | 0 | 0 | 0 | -1 | 0 | 0 | 0 | -1 | -1 | 0 | 0 | -1 | -1 | -1 | -1 | 13 | 0 | -1 | 3 | -1 | -1 | 0 | 0 | 2 | 4 | 0 | 0 | 0 | 0 | 0 | 0 | 2 | 3 | 1 | 0 | 0 | 0 | 0 | 0 | -1 | 0 | -1 | 0 | 0 | 0 | 0 | 0 | 0 | 0 | 0 | 0 | 0 | 1 | 0 | 1 | 2 | 2 |
| Eea1 | chr10:96000351-96000365 | 17 | 0 | 0 | 0 | -1 | 1 | 0 | 0 | 0 | 4 | 0 | 0 | 0 | -1 | 0 | -1 | 34 | 94 | -1 | 67 | 8 | 3 | 1 | 0 | 0 | 1 | 0 | -1 | -1 | 0 | 2 | 4 | 0 | 5 | 2 | 0 | 0 | 0 | 0 | 0 | 0 | 1 | 0 | 0 | 0 | 1 | 0 | 0 | 1 | 0 | 0 | 0 | 0 | 0 | 0 | 2 | 0 | 0 | 0 | 0 | 0 | 0 | 0 | 0 | 0 | 0 | 1 | 2 | 0 | 0 | 5 | 5 | 0 | 0 | 0 | 0 | 0 | 0 | 0 | 1 | 0 |
| Nek5 | chr8:22078456-22078573 | -1 | -1 | -1 | 15 | -1 | -1 | 0 | 0 | -1 | 11 | -1 | 0 | -1 | -1 | -1 | -1 | -1 | 0 | -1 | 55 | -1 | -1 | -1 | -1 | -1 | -1 | -1 | -1 | -1 | -1 | -1 | -1 | -1 | -1 | -1 | -1 | -1 | -1 | -1 | -1 | -1 | -1 | 0 | -1 | -1 | -1 | -1 | -1 | -1 | -1 | -1 | -1 | -1 | -1 | -1 | -1 | -1 | -1 | -1 | -1 | -1 | -1 | -1 | -1 | -1 | -1 | -1 | -1 | -1 | -1 | -1 | -1 | -1 | -1 | -1 | -1 | -1 | -1 | -1 | -1 | 0 |
| Ush2a | chr1:188961968-188962039 | -1 | -1 | -1 | -1 | -1 | -1 | -1 | -1 | 0 | 11 | -1 | -1 | 0 | 0 | -1 | -1 | -1 | -1 | -1 | 99 | -1 | -1 | -1 | -1 | -1 | -1 | -1 | -1 | -1 | -1 | -1 | -1 | -1 | -1 | -1 | -1 | -1 | -1 | -1 | -1 | -1 | -1 | -1 | -1 | -1 | -1 | -1 | -1 | -1 | -1 | -1 | -1 | -1 | -1 | -1 | -1 | -1 | -1 | -1 | -1 | -1 | -1 | -1 | -1 | -1 | -1 | -1 | -1 | -1 | -1 | -1 | 4 | -1 | -1 | -1 | -1 | -1 | -1 | -1 | -1 | -1 |
| Pls1 | chr9:95796275-95796369 | 7 | 0 | -1 | 0 | 4 | 0 | 1 | 0 | 0 | 2 | 0 | 10 | -1 | 0 | -1 | -1 | 0 | 37 | 10 | 59 | 32 | -1 | -1 | -1 | -1 | -1 | -1 | -1 | -1 | -1 | -1 | -1 | -1 | -1 | -1 | -1 | 0 | 0 | 0 | 0 | -1 | -1 | -1 | -1 | -1 | 0 | 0 | -1 | -1 | 0 | 0 | 0 | -1 | -1 | -1 | 0 | 0 | 0 | -1 | -1 | -1 | -1 | -1 | -1 | -1 | -1 | -1 | 0 | -1 | 0 | 2 | 2 | 0 | -1 | -1 | -1 | -1 | -1 | 3 | -1 | 2 |
| Wasl | chr6:24623334-24623351 | 1 | 75 | -1 | 0 | 0 | 0 | 0 | 0 | 0 | 0 | 0 | 1 | 0 | 0 | 0 | 0 | 0 | 0 | 0 | 32 | 0 | 0 | 0 | 0 | 0 | 0 | 0 | 0 | 0 | 0 | 0 | 0 | 0 | 0 | 0 | 0 | 0 | 0 | 0 | 0 | 0 | 0 | 0 | -1 | -1 | -1 | 0 | 0 | 0 | 0 | 0 | 0 | 0 | 0 | 0 | 0 | 0 | 0 | 0 | 0 | 0 | 0 | 0 | 0 | 0 | 0 | 3 | 0 | 0 | 0 | 0 | 0 | 0 | 0 | 0 | 0 | 0 | 0 | 0 | 0 | 0 |

SFig4, pg4

| Gene Symbol | Exon Location (mm10, mouse) | Brain | Skeletal Muscle | Heart | Lung | Stomach | Spleen | Colon | Liver | Kidney | Pancreas islet | Mammary gland | Ovary | Testes | Rod Photoreceptor (p28) | SST interneurons (cortex) | PV interneurons (hippocampus) | Motor neurons (Hb9+, embryo) | DRG neurons | Olfactory sensory neurons (Omp+) | Hair cells (Atoh1+, cochlea) | Purkinje cells (cerebellum) | Pyramidal neurons (Thy1+, cortex) | Pyramidal neurons (hippo - proximal CA1) | Pyramidal neurons (hippo - distal CA1) | Pyramidal neurons (hippo - superficial CA1) | Pyramidal neurons (hippo - dorsal CA1) | Pyramidal neurons (hippo - dorsal CA1, postsubiculum-projecting) | Pyramidal neurons (hippo - intermediate CA1) | Pyramidal neurons (hippo - ventral CA1) | Pyramidal neurons (hippo - ventral CA1, amygdala-projecting) | Pyramidal neurons (hippo - ventral CA1, nucleus accumbens-projecting) | Pyramidal neurons (hippo - deep CA1) | Pyramidal neurons (hippo - dorsal CA3) | Pyramidal neurons (hippo - ventral CA3) |
|---|---|---|---|---|---|---|---|---|---|---|---|---|---|---|---|---|---|---|---|---|---|---|---|---|---|---|---|---|---|---|---|---|---|---|---|
| Casq2 | chr3:102133090-102133098 | 3 | 0 | 0 | 0 | 0 | 0 | 0 | -1 | 0 | 3 | 0 | 0 | 0 | -1 | -1 | -1 | 0 | 0 | -1 | -1 | 99 | -1 | -1 | -1 | -1 | -1 | -1 | -1 | -1 | -1 | -1 | -1 | -1 | -1 |
| Vav3 | chr3:109680686-109680769 | 44 | 0 | 0 | 0 | 0 | 0 | 0 | 0 | 0 | 0 | 0 | 0 | 0 | -1 | 0 | -1 | 2 | 1 | 0 | 0 | 98 | -1 | -1 | -1 | -1 | 0 | -1 | -1 | -1 | -1 | -1 | -1 | 0 | -1 |
| Rps6ka6 | chrX:111446379-111446393 | 8 | 0 | 0 | 0 | -1 | 0 | -1 | -1 | 0 | 2 | 0 | 0 | 0 | -1 | -1 | 0 | 5 | -1 | 0 | 1 | 32 | -1 | -1 | -1 | 0 | -1 | -1 | -1 | -1 | -1 | -1 | -1 | 0 | 0 |
| Stam2 | chr2:52736131-52736319 | 5 | 1 | -1 | 0 | 0 | 0 | 0 | 0 | 0 | 0 | 0 | 0 | 0 | 0 | 0 | 0 | 1 | 0 | 0 | 0 | 30 | 7 | -1 | -1 | -1 | 0 | 0 | -1 | -1 | 0 | 0 | -1 | -1 | -1 |
| Arhgap6 | chrX:169295042-169298469 | 89 | 2 | 0 | 0 | 0 | 0 | 0 | 0 | 0 | 0 | 0 | 0 | -1 | -1 | -1 | -1 | -1 | 0 | 0 | -1 | 91 | -1 | -1 | -1 | -1 | -1 | -1 | -1 | -1 | -1 | -1 | -1 | -1 | -1 |
| Dock7 | chr4:98992016-98992051 | 22 | 2 | 0 | 0 | 0 | 2 | 0 | 0 | 0 | 1 | 0 | 0 | 0 | -1 | -1 | -1 | 1 | 0 | 0 | 0 | 31 | -1 | -1 | 0 | -1 | -1 | -1 | -1 | -1 | 0 | -1 | -1 | -1 | -1 |
| P3h1 | chr4:119239706\|119239709-119239761 | 30 | 1 | 0 | 0 | 0 | 0 | 0 | 0 | 0 | 1 | 0 | 0 | 0 | -1 | -1 | -1 | -1 | 17 | -1 | 2 | 54 | -1 | -1 | -1 | -1 | -1 | -1 | -1 | -1 | -1 | -1 | -1 | -1 | -1 |
| Morc4 | chrX:139835432-139835487 | -1 | 0 | 0 | 0 | -1 | 0 | 0 | 0 | 0 | 0 | 0 | 0 | 0 | -1 | -1 | 0 | -1 | 2 | 0 | -1 | 21 | -1 | 0 | -1 | 0 | -1 | 0 | 0 | -1 | -1 | -1 | -1 | -1 | -1 |
| Patj | chr4:98469220-98469239 | -1 | -1 | 0 | 3 | 0 | 0 | 1 | 1 | 0 | 2 | 2 | 0 | 1 | -1 | -1 | -1 | -1 | -1 | 28 | 19 | 83 | -1 | -1 | -1 | -1 | -1 | -1 | -1 | -1 | -1 | -1 | -1 | 0 | -1 |
| Poc1b | chr10:99164586-99164607 | 9 | 0 | 0 | 0 | 0 | 0 | 0 | 0 | 0 | 0 | 0 | 0 | 0 | -1 | -1 | -1 | 2 | 11 | 0 | 0 | 37 | -1 | 0 | 0 | 0 | -1 | 33 | 0 | -1 | -1 | -1 | 0 | 0 | 0 |
| Afap1l2 | chr19:56933717-56933800 | 17 | -1 | 0 | 0 | -1 | 1 | -1 | 0 | 0 | 5 | 1 | 2 | 0 | -1 | -1 | -1 | 0 | 0 | 0 | 0 | 82 | -1 | -1 | -1 | -1 | -1 | -1 | -1 | -1 | -1 | -1 | -1 | -1 | -1 |
| Myh10 | chr11:68775365-68775427 | 12 | 0 | 0 | 0 | -1 | 0 | 0 | 0 | 0 | 0 | 0 | 0 | 0 | 0 | 0 | 27 | 0 | 15 | 0 | 0 | 59 | 18 | 2 | 4 | 2 | 11 | 1 | 4 | 1 | 0 | 0 | 6 | 3 | 2 |
| Grb7 | chr11:98449112-98449162 | -1 | -1 | 0 | 0 | 0 | 0 | 1 | 0 | 1 | 1 | 0 | -1 | 0 | -1 | -1 | -1 | -1 | -1 | 0 | 1 | 26 | -1 | -1 | -1 | -1 | -1 | -1 | -1 | -1 | -1 | -1 | -1 | -1 | -1 |
| Tmem2 | chr19:21842686-21842789 | 15 | 2 | 0 | 0 | 0 | 0 | 0 | 0 | 0 | 0 | 0 | 0 | 3 | 0 | -1 | 0 | 8 | 8 | 3 | 0 | 23 | 0 | -1 | -1 | 0 | -1 | 0 | -1 | 0 | 0 | -1 | -1 | 0 | 0 |
| Lpin1 | chr12:16558644-16558721 | 46 | 0 | 0 | 0 | 0 | 0 | 0 | 0 | 0 | 0 | 0 | 0 | 0 | 0 | -1 | -1 | 12 | 22 | 17 | 0 | 42 | -1 | -1 | 0 | -1 | -1 | 0 | 0 | -1 | -1 | -1 | 0 | -1 | -1 |
| Prkd2 | chr7:16861731-16861829 | 17 | 0 | 1 | 0 | 0 | 2 | 2 | 1 | 0 | 0 | 1 | 0 | 0 | -1 | -1 | -1 | -1 | 11 | -1 | 1 | 67 | -1 | -1 | -1 | -1 | -1 | -1 | -1 | -1 | -1 | -1 | -1 | -1 | -1 |
| Padi2 | chr4:140930977-140931042 | 5 | 0 | 0 | 0 | 0 | 0 | 0 | 0 | 0 | 1 | 1 | 0 | 0 | 5 | -1 | -1 | -1 | -1 | 0 | 0 | 23 | -1 | -1 | -1 | -1 | -1 | -1 | -1 | 0 | -1 | -1 | -1 | -1 | -1 |
| Dennd4c | chr4:86780293-86780514 | -1 | 6 | 0 | 0 | 0 | 0 | 0 | 0 | 0 | 1 | 2 | 3 | 0 | -1 | -1 | -1 | -1 | -1 | 0 | 0 | 42 | -1 | -1 | 0 | -1 | -1 | -1 | 0 | -1 | 0 | -1 | -1 | -1 | 0 |
| Igsf10 | chr3:59334113-59334188 | -1 | 0 | -1 | -1 | -1 | 6 | 0 | 0 | 0 | 4 | 22 | 1 | 0 | -1 | -1 | -1 | -1 | -1 | 0 | 0 | 90 | -1 | -1 | -1 | -1 | -1 | -1 | -1 | -1 | -1 | -1 | -1 | -1 | -1 |
| Carmil1 | chr13:24020047-24020071 | 43 | 0 | 0 | 0 | 0 | 0 | 2 | 1 | 0 | 2 | 2 | 0 | 1 | -1 | 0 | -1 | 29 | 30 | 19 | 1 | 55 | -1 | 0 | -1 | -1 | -1 | 0 | -1 | -1 | -1 | -1 | -1 | -1 | -1 |
| Lima1 | chr15:99787325-99787383 | 5 | 0 | -1 | 0 | 0 | 0 | 0 | 1 | 0 | 0 | 0 | 0 | 0 | 4 | -1 | -1 | -1 | 0 | 1 | 0 | 74 | -1 | -1 | -1 | -1 | -1 | -1 | -1 | -1 | -1 | -1 | -1 | -1 | -1 |
| Ptbp1 | chr10:79860447-79860502 | 34 | 13 | 0 | 0 | 0 | 0 | 0 | 0 | 0 | 0 | 0 | 0 | 0 | 0 | 0 | 0 | 2 | 12 | 0 | 0 | 25 | -1 | -1 | -1 | -1 | -1 | -1 | -1 | -1 | 0 | 0 | -1 | -1 | -1 |
| Celsr3 | chr9:108837781-108837818 | 43 | -1 | -1 | -1 | -1 | -1 | 0 | -1 | -1 | 0 | 0 | -1 | 0 | 0 | -1 | -1 | 0 | 3 | 0 | 0 | 30 | -1 | -1 | -1 | -1 | -1 | -1 | -1 | -1 | 0 | -1 | -1 | -1 | -1 |
| Wrn | chr8:33326586-33326640 | 0 | 0 | 0 | 0 | -1 | 0 | 0 | 0 | 0 | 0 | 0 | 0 | 0 | 0 | 0 | 0 | 0 | 0 | 0 | 0 | 0 | 20 | 0 | 0 | 0 | 2 | 0 | 0 | 0 | 0 | -1 | 0 | 0 | 0 |
| Tgds | chr14:118122739-118122765 | 2 | 7 | 0 | 0 | 0 | 0 | 0 | 0 | 0 | 0 | 1 | 0 | 0 | 0 | 0 | 0 | 0 | 8 | 0 | 0 | 5 | 64 | -1 | -1 | 1 | 0 | 0 | 0 | 0 | -1 | 0 | 0 | 0 | -1 |
| Cul3 | chr1:80320314-80320341 | 0 | 0 | 0 | 0 | -1 | 0 | 0 | 0 | 1 | 0 | 1 | 1 | 1 | 2 | 0 | 0 | 1 | 0 | 9 | 0 | 0 | 37 | 1 | 0 | 0 | 1 | 0 | 0 | -1 | 7 | 0 | 4 | 0 | 0 |
| Ccl27b | chr4:42652879\|42651450-42653052 | 28 | -1 | 0 | 0 | 0 | -1 | 0 | -1 | 0 | 0 | -1 | -1 | 33 | 0 | 0 | 0 | 1 | 2 | 5 | 1 | 9 | 59 | 0 | 0 | 0 | 0 | 0 | -1 | 0 | 0 | 0 | 6 | -1 | 0 |
| Brd7 | chr8:88359784-88359925 | 3 | 0 | 1 | 1 | 0 | 0 | 0 | 1 | 0 | 0 | 1 | 0 | 0 | 1 | 0 | 0 | 0 | 1 | 0 | 0 | 0 | 24 | 0 | 0 | 0 | 0 | 0 | 0 | 0 | 13 | 1 | 0 | 2 | 0 |
| Rab11fip2 | chr19:59939157-59939253 | 2 | 0 | 0 | 2 | -1 | 0 | 0 | 0 | 1 | 2 | 2 | 1 | 0 | 1 | 1 | 0 | 2 | 3 | 4 | 1 | 4 | 27 | 1 | 0 | 10 | 2 | 1 | 0 | 0 | 0 | 0 | 0 | 1 | 0 |
| Agfg1 | chr1:82866408-82866513 | 2 | 1 | 0 | 0 | 0 | 0 | 0 | 1 | 0 | 0 | 2 | 1 | 3 | 1 | -1 | 2 | 0 | 1 | 0 | 1 | 2 | 33 | 1 | 3 | 1 | 1 | 1 | 1 | 0 | 1 | 1 | 0 | 1 | 0 |
| Kcnc2 | chr10:112456929-112457032 | 20 | 0 | 0 | 0 | -1 | -1 | -1 | -1 | 0 | 0 | 0 | 0 | 0 | 0 | 0 | 1 | 0 | 0 | 0 | -1 | 1 | -1 | 77 | 76 | 65 | 75 | 76 | 50 | -1 | -1 | 96 | 16 | 58 | 62 |
| Fnbp1l | chr3:122552136-122556570 | 81 | 0 | 0 | 0 | 0 | 0 | 0 | 0 | 0 | 0 | 0 | 0 | 0 | -1 | -1 | -1 | 1 | 5 | 6 | 19 | 37 | -1 | 96 | 99 | 97 | 97 | 99 | 98 | -1 | -1 | -1 | -1 | -1 | 99 |
| Cnih1 | chr14:46783826-46783873 | 22 | 0 | 0 | 0 | 0 | 0 | 0 | 0 | 0 | 0 | 0 | 0 | 0 | 0 | 7 | 1 | 0 | 1 | 0 | 0 | 8 | 50 | 22 | 16 | 26 | 32 | 35 | -1 | -1 | 27 | 89 | -1 | 18 | 40 |
| Clasp1 | chr1:118565143-118565259 | 59 | 0 | 2 | 0 | 0 | 0 | 0 | 0 | 0 | 0 | 0 | 0 | 0 | 0 | 0 | -1 | 13 | 1 | 0 | 0 | 42 | -1 | 66 | 75 | 65 | 40 | -1 | -1 | 22 | -1 | -1 | -1 | 31 | 46 |
| Cdc42bpa | chr1:180118736-180118783 | 32 | 0 | 0 | 0 | 0 | 1 | 0 | 0 | 0 | 0 | 0 | 0 | 0 | 18 | 0 | 3 | 1 | 1 | 0 | 0 | 63 | 0 | 73 | 85 | 65 | 71 | 74 | 75 | -1 | 15 | 69 | 59 | 60 | 48 |
| Map4k3 | chr17:80622398-80622430 | 17 | 0 | 0 | 0 | 0 | 0 | 0 | 0 | 0 | 0 | 0 | 0 | 0 | 6 | 0 | 18 | 0 | 9 | 7 | 0 | 18 | 0 | 36 | 32 | 22 | 26 | 18 | 34 | 23 | 19 | 53 | 35 | 20 | 26 |
| Mindy3 | chr2:12402200-12402362 | 9 | 0 | 0 | 0 | 1 | 1 | 1 | 1 | 1 | 2 | 2 | 0 | 2 | 0 | 1 | -1 | 2 | 1 | 0 | 0 | 2 | 0 | 4 | 20 | 31 | 41 | -1 | -1 | 25 | -1 | -1 | -1 | -1 | 28 |
| Gyg | chr3:20123122-20123172 | 29 | 1 | 1 | 0 | 0 | 0 | 0 | 3 | 0 | 1 | 0 | 0 | 0 | 0 | 49 | 16 | 37 | 19 | 9 | 1 | 17 | 60 | 76 | 17 | 48 | 69 | -1 | -1 | -1 | 77 | 91 | 17 | 43 | 59 |
| Enah | chr1:181931174-181931185 | 64 | 1 | 0 | 0 | 0 | 0 | 0 | 0 | 0 | 0 | 0 | 0 | 0 | 1 | -1 | 72 | 2 | 6 | 0 | 0 | 40 | -1 | 86 | 67 | 88 | 86 | 92 | 89 | 74 | 59 | -1 | 83 | 72 | 84 |
| Nf1 | chr11:79414241-79414270 | 46 | -1 | 0 | 0 | -1 | 0 | 0 | 3 | 0 | 0 | 0 | 0 | 0 | 0 | -1 | 64 | 35 | 1 | 0 | 12 | 15 | 0 | 88 | 84 | 94 | 94 | 89 | 91 | -1 | 77 | 88 | 73 | 77 | 89 |
| Rnf32 | chr5:29196914-29197017 | 1 | 0 | -1 | 0 | -1 | 0 | 0 | 0 | 0 | 0 | 0 | 0 | 0 | 0 | 0 | 0 | 0 | 0 | 0 | 0 | 0 | 0 | 0 | 0 | -1 | 0 | 0 | 0 | 0 | 0 | 0 | 0 | 0 | 0 |
| Rrn3 | chr16:13811939-13812023 | 0 | 0 | 1 | 0 | 0 | 0 | 0 | 0 | 0 | 0 | 0 | 0 | 0 | 0 | 0 | 0 | 0 | 0 | 0 | 0 | 0 | 0 | 0 | 0 | 0 | 0 | 0 | 0 | 0 | 0 | 0 | -1 | 1 | 0 |
| Arfgef2 | chr2:166892284-166892399 | 2 | 3 | 0 | 0 | 0 | 1 | 0 | 1 | 1 | 1 | 0 | 2 | 1 | 0 | 0 | 0 | 0 | 2 | 0 | 3 | 2 | 0 | -1 | -1 | -1 | -1 | 0 | 0 | -1 | 0 | 0 | 0 | 4 | -1 |
| Rab18 | chr18:6777667-6777697 | 2 | 1 | 0 | 1 | 0 | 1 | 0 | 0 | 0 | 1 | 1 | 0 | 1 | 2 | 0 | 0 | 1 | 0 | 2 | 1 | 1 | 0 | 1 | 0 | 0 | 1 | 0 | 0 | -1 | 0 | 13 | 1 | 2 | 10 |
| Aim2 | chr1:173462756-173462870 | -1 | 0 | -1 | 0 | -1 | 3 | 1 | 0 | -1 | 2 | 0 | 0 | 0 | 0 | -1 | 0 | 0 | -1 | -1 | -1 | -1 | -1 | 0 | 0 | -1 | 0 | 0 | -1 | -1 | -1 | -1 | -1 | 0 | 0 |
| Otof | chr5:30370076-30370278 | 85 | -1 | -1 | 0 | -1 | 0 | -1 | -1 | 0 | 0 | -1 | 0 | -1 | -1 | -1 | -1 | -1 | -1 | -1 | 0 | 0 | -1 | -1 | -1 | -1 | -1 | -1 | -1 | -1 | -1 | -1 | -1 | -1 | -1 |
| Gm10010 | chr6:128196769-128196833 | -1 | 0 | -1 | -1 | -1 | -1 | -1 | -1 | -1 | 8 | 0 | -1 | 3 | 0 | -1 | 0 | -1 | 2 | -1 | 0 | 10 | 0 | -1 | 6 | 11 | 11 | 0 | 0 | -1 | 21 | -1 | -1 | 20 | 25 |
| Slit3 | chr11:35595297-35595329 | 0 | 0 | 0 | 0 | 0 | 0 | 0 | 0 | 0 | 0 | 0 | 0 | 0 | -1 | -1 | 0 | 1 | 0 | -1 | 0 | 0 | -1 | 0 | 0 | 0 | 0 | 0 | 0 | 0 | 0 | 0 | 0 | 0 | 0 |
| Atrx | chrX:105881169-105881255 | 1 | 0 | 0 | 0 | -1 | 0 | 0 | 0 | 0 | 0 | 0 | 0 | 0 | 0 | 5 | 1 | 1 | 0 | 0 | 0 | 1 | 0 | 0 | 0 | 0 | 0 | 0 | 0 | 0 | 0 | 1 | 0 | 0 | 0 |
| Pld3 | chr7:27551510-27551565 | 0 | 0 | 0 | 0 | 0 | 2 | 1 | 0 | 0 | 0 | 0 | 0 | 0 | 14 | 0 | 0 | 3 | 0 | 3 | 0 | 0 | 0 | 0 | 0 | 0 | 0 | 0 | 0 | 0 | 0 | 0 | 0 | -1 | 8 |
| Lyst | chr13:13632545-13632674 | 0 | 0 | 0 | -1 | -1 | 0 | -1 | 0 | -1 | 0 | 0 | 0 | -1 | 0 | 0 | 1 | 5 | 3 | 0 | 0 | 2 | -1 | 1 | 0 | 0 | 8 | 0 | 0 | 0 | 0 | 0 | 0 | 5 | 0 |
| Srrm1 | chr4:135338549-135338586 | 2 | 2 | 2 | 1 | 0 | 1 | 1 | 0 | 0 | 1 | 0 | 0 | 1 | 0 | 0 | 1 | 1 | 2 | 0 | 1 | 1 | 0 | 1 | 0 | 0 | 0 | 0 | 1 | 0 | 0 | 0 | 0 | 0 | 1 |
| Nalcn | chr14:123603733-123603834 | 4 | -1 | 0 | -1 | -1 | -1 | -1 | -1 | -1 | 1 | 0 | -1 | -1 | -1 | 0 | 0 | 1 | 2 | 0 | 0 | 4 | -1 | 1 | 0 | 2 | 3 | -1 | -1 | 11 | 7 | 0 | 4 | 6 | 5 |
| Nek9 | chr12:85334357-85334496 | 4 | 0 | 3 | 0 | -1 | 0 | 0 | 0 | 0 | 0 | 0 | 0 | 0 | 9 | -1 | -1 | 2 | 2 | -1 | 0 | 3 | -1 | 2 | -1 | 2 | 1 | 0 | 0 | 0 | 0 | 16 | 0 | 0 | 22 |
| Arhgap12 | chr18:6073211-6073286 | 1 | 9 | 0 | 0 | 0 | 1 | 1 | 1 | 1 | 1 | 1 | 1 | 0 | 0 | 0 | 0 | 2 | 1 | 0 | 2 | 1 | 0 | 2 | 5 | 2 | 3 | 4 | 0 | 3 | 0 | 9 | 0 | 8 | 2 |
| Irak1bp1 | chr9:82839139-82839274 | 4 | 2 | 0 | 0 | 0 | 0 | 0 | 0 | 0 | 1 | 1 | 2 | 5 | 2 | 0 | 2 | 2 | 1 | 1 | 0 | 1 | 4 | 5 | 4 | 1 | 5 | 4 | 2 | 6 | 1 | -1 | 1 | 2 | 1 |

| Gene Symbol | Granule cells (hippo - dorsal DG) | Granule cells (hippo - ventral DG) | Oligodendrocyte precursor cells (cortex) | Newly formed oligodendrocytes (cortex) | Myelinating oligodendrocytes (cortex) | Astrocytes (cortex) | Microglia (cortex) | Microglia (spinal cord) | Endothelial cells (cortex) | Quadriceps | Gastrocnemius | Heart ventricle (p90) | Hematopoietic stem cell (bone marrow) | Dendritic cells (spleen) | Neutrophil | Mast cell | NKT cell | NK cell | Macrophage | Eosinophils | B cell, spleen | CD4+ T-cell (Th17) | CD4+ T-cell (Th1) | CD4+ T-cell (Th2) | CD8+ T-cell | CD4+/CD8+ double positive T-cell | Gamma delta T-cell | Regulatory T-cell (Rag2+, spleen) | Hepatocyte (liver) | Liver sinusoidal endothelial cells (liver) | Kupffer cells (liver) | Hepatic stellate cells (liver) | Smooth muscle cells (colon) | Interstitial Cell of Cajal (colon) | Gastrointestinal PDGFRa+ cells (colon) | Gastrointestinal PDGFRa+ cells (colon mucosa) | Pancreatic alpha cells (7-16wks) | Pancreatic beta cells (7-16wks) | Alveolar type 1 (lung) | Podocytes (kidney) | Classical brown adipocyte | Inguinal white adipocyte | Sertoli cells (testis) | Leydig cells (testis) | Interstitial cells (testis) | Spermatogonial stem cells (testis) | Female germline stem cells (ovary) |
|---|---|---|---|---|---|---|---|---|---|---|---|---|---|---|---|---|---|---|---|---|---|---|---|---|---|---|---|---|---|---|---|---|---|---|---|---|---|---|---|---|---|---|---|---|---|---|
| Casq2 | -1 | -1 | -1 | -1 | -1 | -1 | 0 | 0 | 0 | 0 | 0 | 0 | -1 | -1 | -1 | -1 | -1 | -1 | -1 | -1 | -1 | -1 | -1 | -1 | -1 | -1 | -1 | -1 | -1 | -1 | -1 | -1 | 0 | 0 | 0 | 0 | 0 | -1 | -1 | 0 | 0 | 0 | 0 | 0 | 0 | 0 | 0 |
| Vav3 | -1 | -1 | 2 | 0 | 0 | 0 | 0 | -1 | 0 | 0 | 0 | 0 | 0 | 0 | 0 | 0 | 0 | 0 | 0 | 0 | 0 | 0 | 0 | 0 | 0 | 0 | 0 | 0 | -1 | 0 | -1 | 0 | 0 | 0 | 0 | 0 | 0 | -1 | -1 | 0 | 0 | 0 | 0 | 0 | 0 | 0 | 0 |
| Rps6ka6 | 0 | -1 | 0 | 0 | 0 | 0 | -1 | 0 | -1 | 0 | -1 | 0 | 0 | -1 | -1 | -1 | -1 | -1 | -1 | -1 | -1 | -1 | -1 | -1 | -1 | 0 | -1 | -1 | -1 | -1 | -1 | 0 | 0 | 0 | 0 | -1 | 0 | 0 | 0 | 0 | -1 | -1 | 0 | 0 | 0 | 0 | 0 |
| Stam2 | 0 | 0 | 0 | 0 | 0 | 0 | 0 | 0 | 0 | 3 | 0 | 1 | 0 | 0 | 0 | 0 | 0 | 0 | 0 | 0 | 0 | 0 | 0 | 0 | 0 | 0 | 0 | 0 | 0 | 0 | 0 | 0 | 0 | 0 | 0 | 0 | 0 | 0 | 0 | 0 | 0 | 0 | 0 | 0 | 0 | 0 | 0 |
| Arhgap6 | -1 | -1 | 0 | -1 | -1 | 0 | -1 | -1 | 0 | 0 | 0 | 0 | 0 | 0 | -1 | 0 | 0 | 0 | 0 | 0 | 0 | -1 | 0 | 0 | -1 | -1 | -1 | -1 | 0 | 0 | -1 | 0 | 0 | 0 | 0 | 0 | 0 | 0 | 0 | 0 | 0 | 0 | 0 | 0 | 0 | 0 | 0 |
| Dock7 | -1 | -1 | 0 | 0 | 0 | 1 | 0 | 0 | 0 | 0 | -1 | 0 | 0 | -1 | -1 | 0 | -1 | -1 | 0 | -1 | -1 | 0 | 0 | 0 | 0 | -1 | -1 | -1 | 0 | 0 | 0 | 0 | 0 | 0 | 0 | 0 | 3 | 0 | 0 | 0 | 0 | 0 | 0 | 0 | 0 | 0 | 1 |
| P3h1 | -1 | -1 | 0 | 0 | 2 | 3 | 0 | 0 | 0 | 0 | 0 | 0 | 0 | 0 | -1 | 0 | -1 | -1 | 0 | 0 | 0 | 0 | 0 | 0 | -1 | 1 | -1 | 0 | 0 | 0 | 0 | 0 | 0 | 0 | 0 | 0 | 3 | 1 | 0 | 0 | 0 | 0 | 0 | 0 | 0 | 0 | 0 |
| Morc4 | -1 | 0 | 0 | 0 | 0 | 0 | 0 | -1 | 0 | 0 | 0 | 1 | 0 | -1 | -1 | -1 | -1 | -1 | 0 | -1 | -1 | -1 | -1 | -1 | -1 | -1 | -1 | -1 | 0 | 0 | 0 | 0 | 0 | 6 | 0 | 0 | -1 | 0 | 0 | 1 | 0 | 0 | 3 | 1 | 1 | 0 | 0 |
| Patj | -1 | -1 | 0 | 0 | -1 | 0 | -1 | 0 | 0 | -1 | -1 | 0 | 0 | 0 | 0 | 0 | 0 | 0 | 0 | -1 | 0 | 0 | 0 | 0 | 0 | 0 | 0 | 0 | 0 | 0 | 0 | 0 | 0 | -1 | 0 | 0 | 10 | 5 | 0 | 2 | 0 | 0 | 0 | 1 | 1 | 0 | 1 |
| Poc1b | -1 | -1 | 1 | 0 | 0 | 1 | 0 | 1 | 0 | 0 | 0 | 0 | 0 | 0 | 0 | 0 | 0 | 0 | 0 | 0 | 0 | 0 | 0 | 0 | 0 | 0 | 0 | 0 | 0 | 0 | 0 | 0 | 0 | 0 | 0 | 0 | 2 | 0 | 0 | 0 | 0 | 0 | 0 | 0 | 0 | 0 | 0 |
| Afap1l2 | -1 | -1 | 1 | 0 | -1 | 0 | -1 | 0 | 0 | -1 | 0 | 0 | -1 | -1 | -1 | -1 | -1 | -1 | -1 | -1 | -1 | -1 | -1 | -1 | -1 | -1 | -1 | -1 | -1 | -1 | 0 | 0 | -1 | 7 | 2 | 3 | -1 | -1 | 1 | 0 | 0 | 0 | 0 | 2 | 8 | 4 | -1 |
| Myh10 | 0 | 0 | 0 | 0 | 0 | 0 | 0 | 4 | 0 | 0 | 0 | 0 | 0 | -1 | -1 | -1 | 0 | 0 | 0 | -1 | -1 | 0 | 0 | 0 | 0 | 0 | -1 | -1 | 0 | 0 | 0 | 0 | 0 | 0 | 0 | 0 | 0 | 0 | 0 | 0 | 0 | 0 | 0 | 0 | 0 | 1 | 0 |
| Grb7 | -1 | -1 | -1 | -1 | -1 | -1 | 0 | -1 | 0 | -1 | -1 | -1 | -1 | -1 | -1 | -1 | -1 | 0 | -1 | 0 | 0 | 0 | -1 | 7 | -1 | 0 | -1 | -1 | 1 | 0 | -1 | 0 | -1 | 0 | 0 | 0 | 0 | 2 | 0 | 0 | -1 | 0 | 0 | 0 | 0 | 1 | -1 |
| Tmem2 | -1 | -1 | 0 | 1 | 1 | 0 | 0 | -1 | 0 | 3 | -1 | 0 | 0 | 0 | 0 | 0 | 0 | 0 | 0 | 0 | 0 | 0 | 0 | 0 | -1 | 0 | 0 | 0 | 0 | 0 | 0 | 0 | 0 | 0 | 0 | 0 | 0 | 0 | 0 | 0 | 0 | 0 | 0 | 0 | 0 | 0 | 0 |
| Lpin1 | -1 | -1 | 0 | 2 | 0 | 1 | 0 | 1 | 0 | 0 | 0 | 0 | 0 | 0 | -1 | 0 | 0 | 0 | 0 | 0 | 0 | 0 | 0 | 0 | 0 | 0 | 0 | 0 | 0 | 0 | 0 | 0 | 0 | 0 | 0 | 1 | 0 | 0 | 0 | 0 | 0 | 0 | 0 | 0 | 0 | 0 | 0 |
| Prkd2 | -1 | -1 | 31 | -1 | 0 | 12 | 0 | -1 | 0 | 0 | 0 | 0 | 1 | 1 | 1 | 2 | 0 | 0 | 0 | 2 | 1 | 1 | 0 | 0 | 1 | 2 | 0 | 1 | 0 | 1 | 0 | 1 | 0 | 1 | 1 | 0 | 1 | 0 | -1 | 1 | -1 | 0 | 1 | 3 | 0 | 0 | -1 |
| Padi2 | -1 | -1 | 0 | 7 | 7 | 0 | -1 | 4 | -1 | 0 | 0 | -1 | 0 | 0 | 0 | 0 | 0 | 0 | 0 | 0 | -1 | 1 | 0 | 0 | 0 | 0 | 3 | -1 | -1 | -1 | -1 | -1 | -1 | 0 | -1 | -1 | 0 | -1 | -1 | 0 | 0 | -1 | 1 | 1 | 0 | 0 | 0 |
| Dennd4c | -1 | -1 | -1 | -1 | 0 | 9 | 0 | -1 | 1 | 5 | 3 | 1 | 0 | 0 | 0 | 1 | 0 | 0 | 0 | 0 | 0 | 0 | 0 | 0 | 0 | 1 | 0 | -1 | 0 | 0 | -1 | 0 | 2 | 1 | 0 | 1 | 2 | 1 | 0 | 1 | 0 | 1 | 4 | 2 | 3 | 5 | 9 |
| Igsf10 | -1 | -1 | -1 | -1 | -1 | -1 | -1 | -1 | 0 | 0 | -1 | 1 | 0 | -1 | -1 | -1 | -1 | -1 | -1 | -1 | -1 | -1 | 0 | -1 | -1 | 0 | -1 | -1 | -1 | -1 | -1 | 0 | -1 | 0 | 2 | 3 | 0 | -1 | -1 | 0 | 0 | 5 | 7 | 8 | 7 | 4 | 0 |
| Carmil1 | -1 | 0 | 3 | 1 | 2 | 1 | 0 | 1 | 0 | 0 | -1 | 0 | 0 | 0 | -1 | 1 | 0 | 0 | 0 | 0 | -1 | 0 | 0 | 0 | -1 | 0 | -1 | 0 | 0 | 0 | 0 | 0 | -1 | 0 | 0 | 0 | 3 | 5 | 1 | 0 | 0 | 0 | 0 | 0 | 0 | 0 | 0 |
| Lima1 | -1 | -1 | 12 | 22 | 17 | -1 | 2 | 1 | 0 | 0 | 0 | 27 | -1 | 0 | 0 | 0 | 0 | -1 | 0 | 0 | -1 | -1 | 0 | -1 | -1 | -1 | -1 | -1 | 0 | 0 | 2 | 0 | 0 | 0 | 0 | 0 | 0 | 0 | 0 | 0 | 0 | 0 | 0 | 0 | 1 | 1 | 23 |
| Ptbp1 | -1 | -1 | 0 | 0 | 1 | 0 | 0 | 0 | 0 | 4 | -1 | 1 | 0 | 0 | 0 | 0 | 0 | 0 | 0 | 0 | 0 | 0 | 0 | 0 | 0 | 0 | 0 | 0 | 0 | 0 | 0 | 0 | 3 | 0 | 0 | 0 | 0 | 0 | 0 | 0 | 1 | 1 | 0 | 0 | 0 | 0 | 0 |
| Celsr3 | -1 | -1 | -1 | 0 | -1 | 0 | -1 | -1 | -1 | -1 | -1 | -1 | -1 | -1 | 0 | 0 | -1 | -1 | -1 | 0 | -1 | -1 | -1 | -1 | -1 | -1 | -1 | -1 | -1 | -1 | -1 | -1 | -1 | 0 | -1 | -1 | 0 | 0 | 0 | -1 | -1 | -1 | -1 | -1 | 0 | 0 | 0 |
| Wrn | 0 | 0 | 0 | 0 | 0 | 0 | -1 | 0 | 0 | 0 | 0 | 0 | 0 | 0 | -1 | 0 | 0 | 0 | 0 | 0 | 0 | 0 | 0 | 0 | 0 | 0 | 0 | 0 | -1 | 0 | 0 | 0 | 0 | 0 | 0 | 0 | 0 | 0 | 0 | 0 | 0 | 0 | 0 | 0 | 0 | 2 | 0 |
| Tgds | 0 | -1 | 0 | 0 | 0 | 0 | 0 | 0 | 0 | 6 | 0 | 3 | 0 | 0 | 0 | 1 | 0 | 0 | 0 | 0 | 0 | 0 | 0 | 0 | 0 | 0 | 0 | 0 | 0 | 0 | 0 | 0 | 11 | 4 | 2 | 0 | 0 | 0 | 0 | 0 | 0 | 0 | 0 | 1 | 0 | 0 | 2 |
| Cul3 | 0 | 0 | 0 | 0 | 3 | 1 | 0 | 0 | 0 | 1 | 1 | 1 | 0 | 0 | 0 | 1 | 0 | 0 | 0 | 1 | 1 | 1 | 0 | 0 | 0 | 1 | 0 | 0 | 0 | 0 | 0 | 0 | 0 | 0 | 1 | 0 | 1 | 0 | 4 | 0 | 0 | 0 | 0 | 0 | 0 | 1 | 1 |
| Ccl27b | 0 | 15 | 0 | 0 | -1 | 0 | 0 | -1 | 0 | -1 | 0 | 0 | 0 | -1 | -1 | -1 | -1 | -1 | -1 | -1 | -1 | -1 | 0 | 0 | -1 | 0 | -1 | 0 | 0 | -1 | 0 | 0 | 0 | 0 | 0 | -1 | -1 | 0 | 0 | 0 | 0 | 0 | -1 | -1 | -1 | -1 | -1 |
| Brd7 | 2 | 0 | 1 | 0 | 2 | 2 | 1 | 0 | 2 | 0 | 1 | 1 | 2 | 0 | 0 | 1 | 0 | 0 | 0 | 0 | 0 | 0 | 0 | 1 | 1 | 0 | 0 | 0 | 0 | 0 | 0 | 0 | 0 | 1 | 0 | 0 | 1 | 1 | 1 | 0 | 0 | 2 | 0 | 0 | 1 | 0 | 0 |
| Rab11fip2 | 2 | 0 | 2 | 1 | 0 | 1 | 0 | 0 | 0 | 1 | -1 | 1 | 0 | 0 | 0 | 0 | -1 | 0 | -1 | 0 | -1 | 3 | 0 | -1 | -1 | 6 | 0 | 0 | -1 | 0 | -1 | 0 | 0 | 0 | 0 | 0 | 1 | 2 | 4 | 1 | 0 | 0 | 4 | 3 | 2 | 0 | 1 |
| Agfg1 | 2 | 0 | 5 | 5 | 1 | 4 | 2 | 0 | 3 | 0 | 1 | 0 | 0 | 0 | -1 | 1 | 0 | 0 | 0 | 0 | 0 | 1 | 2 | 1 | 0 | 1 | 0 | 5 | 0 | 0 | 0 | 0 | 0 | 1 | 0 | 2 | 2 | 2 | 4 | 1 | 0 | 0 | 2 | 2 | 1 | 1 | 0 |
| Kcnc2 | -1 | -1 | 0 | -1 | -1 | 0 | -1 | 0 | -1 | 0 | 0 | 0 | -1 | -1 | -1 | -1 | -1 | -1 | -1 | -1 | -1 | -1 | -1 | -1 | -1 | -1 | -1 | -1 | -1 | -1 | -1 | -1 | 0 | 0 | -1 | -1 | 0 | 0 | -1 | 0 | 0 | 0 | 0 | 0 | 0 | 1 | 0 |
| Fnbp1l | -1 | -1 | 0 | 0 | -1 | 1 | -1 | 0 | 1 | 0 | 0 | 0 | 0 | 0 | 0 | 0 | -1 | -1 | -1 | 0 | -1 | 0 | 0 | 0 | -1 | 0 | -1 | -1 | 0 | 0 | 0 | 0 | 0 | 0 | 1 | 0 | 0 | 0 | 0 | 0 | 0 | 0 | 0 | 0 | 0 | 0 | 0 |
| Cnih1 | 25 | 5 | 0 | 0 | 0 | 0 | 0 | 0 | 0 | 0 | 0 | 0 | 0 | 0 | 0 | 0 | 0 | 0 | 0 | 0 | 0 | 0 | 0 | 0 | 0 | 0 | 0 | 0 | 0 | 0 | 0 | 0 | 0 | 0 | 0 | 0 | 0 | 0 | 0 | 0 | 0 | 0 | 0 | 0 | 0 | 0 | 0 |
| Clasp1 | -1 | -1 | 4 | 2 | 0 | 1 | 4 | 1 | 0 | 0 | 0 | 1 | 0 | 1 | 0 | 0 | 0 | 0 | 0 | 0 | 0 | 0 | 0 | 0 | 0 | 0 | 0 | -1 | -1 | 0 | 0 | 0 | 1 | 0 | 0 | 0 | 0 | 0 | 0 | 0 | 0 | 0 | 0 | 0 | 0 | 0 | 0 |
| Cdc42bpa | 54 | -1 | 1 | 0 | 1 | 1 | 0 | 5 | 0 | 1 | 0 | 0 | 0 | 0 | -1 | 0 | 0 | 0 | 0 | 0 | 0 | 0 | 0 | 0 | 0 | 0 | -1 | -1 | 0 | 0 | 0 | 0 | 0 | 0 | 0 | 0 | 0 | 0 | 0 | 0 | 0 | 0 | 0 | 0 | 0 | 0 | 0 |
| Map4k3 | 6 | -1 | 1 | 0 | 0 | 0 | 0 | 0 | 0 | 0 | 0 | 0 | 0 | 0 | -1 | 0 | -1 | -1 | 0 | 0 | 0 | 0 | 0 | 0 | -1 | 0 | -1 | 0 | 0 | 0 | 0 | 0 | 0 | 0 | 0 | 0 | 1 | 1 | 0 | 0 | 0 | 0 | 0 | 0 | 0 | 0 | 0 |
| Mindy3 | -1 | 21 | 1 | 0 | 1 | 0 | 0 | 0 | 1 | 0 | 0 | 1 | 0 | 0 | 0 | 1 | 2 | 0 | 0 | 0 | 0 | 1 | 2 | 0 | 1 | 0 | 0 | 1 | 0 | 0 | -1 | 0 | 0 | 1 | 0 | 0 | 1 | 2 | 5 | 1 | 0 | 1 | 1 | 1 | 2 | 0 | 1 |
| Gyg | -1 | -1 | 1 | 1 | 2 | 0 | 0 | 0 | 0 | 1 | 0 | 3 | 1 | 1 | 0 | 0 | 0 | 1 | 0 | 0 | 0 | 0 | 0 | 0 | 4 | 0 | 0 | 0 | 0 | 0 | 0 | 0 | 0 | 2 | 1 | 0 | 3 | 1 | 1 | 0 | 0 | 0 | 0 | 0 | 1 | 0 | 1 |
| Enah | 72 | -1 | 4 | 1 | 0 | 0 | -1 | -1 | 0 | 1 | 0 | 1 | -1 | -1 | -1 | 0 | -1 | -1 | -1 | 0 | -1 | 0 | 0 | -1 | -1 | -1 | -1 | -1 | -1 | 0 | -1 | 0 | 1 | 0 | 0 | 0 | 2 | 1 | 0 | 0 | 0 | 0 | 0 | 0 | 0 | 0 | 1 |
| Nf1 | 98 | -1 | 0 | 0 | 0 | 1 | 0 | 0 | 0 | 19 | 49 | 5 | 0 | 0 | 0 | 0 | 0 | 0 | 0 | 0 | 0 | 0 | 0 | 0 | 0 | 0 | 0 | -1 | 0 | 0 | 0 | 0 | 0 | 0 | 0 | 0 | 0 | 0 | 0 | 0 | 0 | 0 | 0 | 0 | 0 | 0 | 0 |
| Rnf32 | 30 | 0 | 0 | 0 | 0 | 0 | -1 | 0 | 0 | -1 | 0 | -1 | -1 | -1 | -1 | -1 | -1 | -1 | -1 | -1 | -1 | 0 | 0 | 0 | 0 | -1 | -1 | 0 | -1 | -1 | -1 | -1 | 0 | 0 | -1 | 0 | 0 | 0 | 0 | 0 | -1 | 0 | 0 | 0 | 0 | 0 | 0 |
| Rrn3 | 44 | 0 | 0 | 0 | 0 | 0 | 0 | 0 | 0 | 0 | 1 | 0 | 0 | 0 | 0 | 0 | 0 | 0 | 0 | 0 | 0 | 0 | 0 | 0 | 0 | 0 | 0 | 0 | 0 | 0 | 0 | 0 | 0 | 0 | 3 | 0 | 1 | 0 | 0 | 0 | 0 | 1 | 0 | 0 | 0 | 0 | 0 |
| Arfgef2 | 90 | 0 | 1 | 1 | 0 | 1 | 0 | 1 | 0 | 1 | 4 | 1 | 0 | 1 | 0 | 1 | 0 | 0 | 0 | 0 | 0 | 1 | 2 | 1 | 0 | 1 | 0 | 1 | 0 | 1 | 0 | 1 | 1 | 2 | 1 | 1 | 0 | 1 | 1 | 1 | 0 | 0 | 1 | 2 | 1 | 2 | 1 |
| Rab18 | 21 | 0 | 1 | 0 | 1 | 0 | 0 | 0 | 0 | 0 | 1 | 0 | 0 | 0 | 0 | 1 | 1 | 0 | 0 | 0 | 0 | 1 | 0 | 0 | 0 | 0 | 0 | 0 | 0 | 0 | 0 | 0 | 1 | 0 | 0 | 1 | 0 | 0 | 0 | 1 | 0 | 0 | 0 | 1 | 0 | 2 | 6 |
| Aim2 | 92 | 0 | 0 | 0 | -1 | -1 | 0 | 11 | 0 | 0 | 0 | 0 | 6 | 6 | 5 | 9 | 14 | 5 | 0 | 5 | 11 | 1 | 4 | 3 | 7 | 1 | 3 | -1 | -1 | -1 | -1 | 4 | 0 | 0 | 0 | 8 | -1 | -1 | -1 | 3 | -1 | 0 | 0 | 7 | 2 | 11 | -1 |
| Otof | 93 | -1 | -1 | -1 | -1 | -1 | -1 | -1 | -1 | -1 | -1 | 0 | -1 | 0 | -1 | -1 | -1 | -1 | -1 | -1 | -1 | -1 | -1 | -1 | -1 | -1 | -1 | -1 | -1 | -1 | -1 | -1 | -1 | 0 | -1 | -1 | 0 | -1 | -1 | 0 | -1 | 0 | 0 | 0 | 0 | 0 | -1 |
| Gm10010 | 79 | -1 | 0 | -1 | -1 | 0 | -1 | -1 | -1 | 0 | -1 | -1 | 0 | -1 | -1 | 0 | -1 | -1 | -1 | -1 | -1 | 0 | -1 | -1 | -1 | -1 | -1 | -1 | -1 | -1 | -1 | -1 | -1 | 0 | -1 | -1 | -1 | -1 | -1 | 0 | -1 | 3 | 0 | 0 | -1 | 3 | 2 |
| Slit3 | 0 | 22 | -1 | 0 | -1 | -1 | -1 | 0 | 0 | 0 | 0 | 0 | -1 | -1 | 0 | -1 | -1 | -1 | -1 | -1 | -1 | 0 | -1 | -1 | 0 | 0 | -1 | -1 | -1 | -1 | -1 | -1 | 0 | 0 | 0 | 0 | 0 | -1 | 0 | 0 | 0 | 0 | 0 | 0 | 0 | 0 | 0 |
| Atrx | 0 | 65 | 0 | 0 | 0 | 0 | 0 | 0 | 0 | 2 | 0 | 0 | 1 | 0 | 0 | 0 | 0 | 0 | 0 | 0 | 0 | 0 | 0 | 0 | 0 | 0 | 0 | 0 | 0 | 0 | 0 | 0 | 0 | 0 | 0 | 0 | 0 | 0 | 0 | 0 | 0 | 0 | 0 | 0 | 0 | 0 | 0 |
| Pld3 | 0 | 26 | 0 | 1 | 3 | 1 | 0 | 0 | 0 | 0 | 0 | 0 | 0 | 0 | 0 | 0 | 0 | 0 | 0 | 0 | 0 | 0 | 0 | 0 | 0 | 0 | 0 | -1 | 0 | 0 | 0 | 0 | 0 | 0 | 0 | 0 | 1 | 0 | 0 | 0 | 0 | 0 | 0 | 1 | 0 | 0 | 0 |
| Lyst | 0 | 27 | 0 | 0 | -1 | 0 | 0 | 0 | 0 | 1 | 3 | 1 | 0 | 0 | 0 | 0 | 0 | 0 | 0 | -1 | 0 | 0 | 0 | 0 | 1 | 0 | 0 | 0 | 0 | 0 | 0 | 0 | -1 | 0 | 0 | 0 | 0 | 0 | 1 | 1 | 2 | 0 | -1 | 1 | 0 | 1 | 0 |
| Srrm1 | 2 | 40 | 1 | 2 | 2 | 2 | 1 | 0 | 0 | 4 | 0 | -1 | 0 | 0 | 0 | 1 | 3 | 0 | 0 | 1 | 0 | 1 | 1 | 1 | 5 | 1 | 0 | 0 | 0 | 1 | 0 | 1 | 1 | 0 | 1 | 0 | 1 | 0 | 1 | 0 | 1 | 1 | 1 | 1 | 1 | 5 | 2 |
| Nalcn | 5 | 64 | 3 | 1 | 0 | 0 | 0 | -1 | 0 | 0 | 0 | -1 | -1 | -1 | -1 | -1 | -1 | -1 | -1 | -1 | -1 | -1 | -1 | -1 | -1 | -1 | -1 | -1 | -1 | -1 | -1 | -1 | 0 | 0 | -1 | -1 | 3 | 5 | -1 | -1 | -1 | -1 | -1 | -1 | 0 | 0 | 5 |
| Nek9 | -1 | 32 | 0 | 1 | 8 | 0 | 0 | 0 | 0 | 0 | 0 | 0 | 0 | 0 | 0 | 0 | 0 | 0 | 0 | 0 | 0 | 0 | 0 | 0 | 0 | 0 | 0 | 0 | 0 | 0 | 0 | 0 | 0 | 0 | 0 | 0 | 0 | 0 | 0 | 0 | 0 | 0 | 0 | 0 | 0 | 0 | 0 |
| Arhgap12 | 7 | 63 | 1 | 0 | 0 | 0 | 0 | 1 | 1 | 9 | 9 | 7 | 0 | -1 | -1 | 0 | 0 | 0 | -1 | -1 | -1 | 2 | 4 | 3 | -1 | 6 | 0 | 0 | -1 | 0 | 1 | 1 | 0 | 1 | 5 | 1 | 2 | 2 | 0 | 1 | 0 | 1 | 1 | 3 | 2 | 3 | 1 |
| Irak1bp1 | 1 | 27 | 0 | 0 | 0 | 0 | 0 | 0 | 0 | 0 | -1 | 0 | 0 | -1 | -1 | 0 | -1 | -1 | -1 | 0 | -1 | 1 | 0 | 0 | -1 | 1 | -1 | -1 | -1 | 0 | 0 | 0 | 0 | 0 | 2 | 0 | 0 | 0 | 1 | 0 | 0 | 0 | 0 | 0 | 0 | 1 | 3 |

**Supplemental Figure 4. Examples of mouse neuronal subtype-specific exons (MESA)**

Alternative exons can be highly cell type specific. Some examples of exons that are selectively enriched in a single neuronal subtype (e.g. rod photoreceptors, SST/PV inhibitory neurons, motor neurons, DRG neurons, olfactory sensory neurons, cochlear hair cells, Purkinje cells, pyramidal neurons, hippocampal granule cells) are shown here. These exons are derived from mouse (MESA compilation).

SFig5, pg1

| Gene Symbol | Exon Location (hg38, human) | Adipose - Subcutaneous | Adipose - Visceral (Omentum) | Adrenal Gland | Artery - Aorta | Artery - Coronary | Artery - Tibial | Bladder | Breast - Mammary Tissue | Brain - Amygdala | Brain - Anterior cingulate cortex (BA24) | Brain - Caudate (basal ganglia) | Brain - Cerebellar Hemisphere | Brain - Cerebellum | Brain - Cortex | Brain - Frontal Cortex (BA9) | Brain - Hippocampus | Brain - Hypothalamus | Brain - Nucleus accumbens (basal ganglia) | Brain - Putamen (basal ganglia) | Brain - Spinal cord (cervical c-1) | Brain - Substantia nigra | Cervix - Ectocervix | Cervix - Endocervix | Colon - Sigmoid | Colon - Transverse | Esophagus - Gastroesophageal Junction | Esophagus - Mucosa | Esophagus - Muscularis | Fallopian Tube | Heart - Atrial Appendage | Heart - Left Ventricle | Kidney - Cortex | Liver | Lung | Minor Salivary Gland | Muscle - Skeletal | Nerve - Tibial | Ovary | Pancreas | Pituitary | Prostate | Skin - Not Sun Exposed (Suprapubic) | Skin - Sun Exposed (Lower leg) | Small Intestine - Terminal Ileum | Spleen | Stomach | Testis | Thyroid | Uterus | Vagina | Whole Blood |
|---|---|---|---|---|---|---|---|---|---|---|---|---|---|---|---|---|---|---|---|---|---|---|---|---|---|---|---|---|---|---|---|---|---|---|---|---|---|---|---|---|---|---|---|---|---|---|---|---|---|---|---|---|
| CCNDBP1 | chr15:43194081-43194137 | 3 | 5 | 34 | 3 | 4 | 2 | 3 | 4 | 11 | 7 | 10 | 5 | 7 | 6 | 4 | 10 | 5 | 8 | 10 | 15 | 9 | 4 | 4 | 4 | 7 | 3 | 4 | 3 | 10 | 3 | 2 | 25 | 10 | 8 | 8 | 1 | 5 | 6 | 8 | 6 | 10 | 2 | 1 | 9 | 8 | 10 | 14 | 19 | 4 | 4 | 5 |
| ATP6V1C2 | chr2:10757427-10757456 | 3 | 4 | 2 | 56 | 19 | 14 | 0 | 3 | -1 | 80 | 11 | 36 | -1 | 62 | 60 | 56 | 27 | 49 | -1 | 5 | -1 | 5 | 3 | 8 | 4 | 3 | 3 | 4 | -1 | 7 | 4 | 1 | -1 | 8 | 2 | 7 | 4 | 5 | 3 | 48 | 3 | 3 | 2 | 4 | 6 | 14 | 1 | 5 | 8 | 2 | 1 |
| CASQ2 | chr1:115718186-115718194 | 0 | 0 | 0 | 0 | 0 | 0 | 0 | 0 | 1 | 1 | 1 | 15 | 54 | 0 | 2 | 1 | 4 | 0 | 0 | 1 | 0 | 0 | 0 | 0 | 0 | 0 | 0 | 0 | 0 | 0 | 0 | 0 | 0 | 0 | 0 | 0 | 0 | 0 | 0 | 0 | 0 | 0 | 0 | 0 | 0 | 0 | 0 | 0 | 0 | 0 | 0 |
| ANK3 | chr10:60108188-60108286 | 0 | 0 | 0 | 0 | 0 | 0 | 0 | 0 | 4 | 17 | 6 | 26 | 27 | 17 | 21 | 8 | 12 | 10 | 4 | 4 | 15 | 0 | 0 | 0 | 0 | 0 | 0 | 0 | 0 | 0 | 0 | 0 | 0 | 0 | 0 | 0 | 0 | 0 | 0 | 0 | 0 | 0 | 0 | 0 | 0 | 0 | 0 | 0 | 0 | 0 | 0 |
| VAV3 | chr1:107576392-107576475 | 0 | 0 | 0 | 0 | 0 | 0 | 0 | 0 | 11 | 22 | 61 | 77 | 66 | 24 | 29 | 23 | 61 | 58 | 31 | 41 | 53 | 0 | 0 | 0 | 0 | 0 | 0 | 0 | 0 | 0 | 0 | 0 | 0 | 0 | 0 | 0 | 0 | 0 | 0 | 1 | 0 | 0 | 0 | 0 | 0 | 0 | 0 | 0 | 0 | 0 | 0 |
| CTNNAL1 | chr9:108983417-108983459 | 0 | 0 | 0 | 0 | 0 | 0 | 0 | 0 | 4 | 11 | 2 | 19 | 24 | 21 | 16 | 4 | 2 | 5 | 3 | 0 | 1 | 0 | 0 | 0 | 0 | 0 | 0 | 0 | 0 | 0 | 0 | 0 | 0 | 0 | 0 | 0 | 0 | 0 | 0 | 0 | 0 | 0 | 0 | 0 | 0 | 0 | 0 | 0 | 0 | 0 | 0 |
| LPIN1 | chr2:11791708-11791785 | 0 | 0 | 0 | 0 | 0 | 0 | 0 | 0 | 5 | 22 | 8 | 51 | 54 | 25 | 30 | 11 | 19 | 14 | 9 | 4 | 6 | 0 | 0 | 1 | 0 | 0 | 0 | 0 | 0 | 0 | 0 | 0 | 0 | 0 | 0 | 0 | 0 | 0 | 0 | 0 | 0 | 0 | 0 | 0 | 0 | 0 | 0 | 0 | 0 | 0 | 0 |
| CPNE3 | chr8:86542046-86542074 | 0 | 0 | 0 | 0 | 0 | 0 | 0 | 0 | 1 | 3 | 2 | 18 | 23 | 5 | 5 | 2 | 3 | 2 | 1 | 1 | 3 | 0 | 0 | 0 | 0 | 0 | 0 | 0 | 0 | 0 | 0 | 0 | 0 | 0 | 0 | 0 | 0 | 0 | 0 | 0 | 0 | 0 | 0 | 0 | 0 | 0 | 0 | 0 | 0 | 0 | 0 |
| MYH14 | chr19:50255220-50255318 | 0 | 0 | 0 | 0 | 0 | 0 | 0 | 0 | 4 | 16 | 7 | 64 | 61 | 17 | 23 | 9 | 18 | 8 | 6 | 3 | 10 | 0 | 0 | 0 | 0 | 0 | 0 | 0 | 0 | 0 | 0 | 0 | 0 | 0 | 0 | 0 | 0 | 1 | 0 | 0 | 0 | 0 | 0 | 0 | 0 | 0 | 0 | 0 | 0 | 0 | 0 |
| MACF1 | chr1:39459821\|39459818-39459832 | 0 | 0 | 0 | 0 | 0 | 0 | 0 | 0 | 7 | 16 | 7 | 38 | 33 | 18 | 24 | 8 | 10 | 10 | 7 | 2 | 5 | 0 | 0 | 0 | 0 | 0 | 0 | 0 | 0 | 0 | 0 | 0 | 0 | 0 | 0 | 0 | 0 | 0 | 0 | 1 | 0 | 0 | 0 | 0 | 0 | 0 | 0 | 0 | 0 | 0 | 0 |
| TAF1 | chrX:71455007-71456117 | 0 | 0 | 0 | 0 | 0 | 0 | 0 | 0 | 0 | 5 | 2 | 45 | 67 | 9 | 5 | 4 | 2 | 2 | 2 | 2 | 1 | 0 | 0 | 0 | 0 | 0 | 0 | 0 | 0 | 1 | 1 | 0 | 0 | 0 | 0 | 1 | 0 | 0 | 0 | 0 | 0 | 0 | 0 | 0 | 0 | 0 | 0 | 0 | 0 | 0 | 0 |
| RBBP8 | chr18:22969596-22969629 | 0 | 0 | 0 | 0 | 0 | 0 | 0 | 0 | 4 | 8 | 4 | 39 | 46 | 15 | 16 | 6 | 5 | 5 | 2 | 2 | 4 | 0 | 0 | 0 | 0 | 0 | 0 | 0 | 0 | 0 | 0 | 0 | 0 | 0 | 0 | 1 | 0 | 0 | 0 | 0 | 0 | 0 | 0 | 0 | 0 | 0 | 0 | 0 | 0 | 0 | 0 |
| TRAPPC8 | chr18:31891699-31891719 | 0 | 0 | 0 | 0 | 0 | 0 | 0 | 0 | 4 | 14 | 3 | 23 | 23 | 14 | 15 | 5 | 2 | 4 | 4 | 1 | 1 | 0 | 0 | 0 | 0 | 0 | 0 | 0 | 1 | 0 | 0 | 0 | 0 | 0 | 0 | 0 | 0 | 0 | 0 | 0 | 0 | 0 | 0 | 0 | 0 | 0 | 0 | 0 | 0 | 0 | 0 |
| PTCH1 | chr9:95450124-95450139 | 0 | 0 | 0 | 0 | 0 | 0 | 0 | 0 | 1 | 5 | 2 | 53 | 44 | 6 | 7 | 2 | 3 | 2 | 3 | 3 | 1 | 0 | 0 | 0 | 0 | 0 | 0 | 0 | 0 | 0 | 0 | 0 | 0 | 0 | 0 | 0 | 0 | 0 | 0 | 1 | 0 | 0 | 1 | 0 | 0 | 0 | 0 | 0 | 0 | 0 | 1 |
| LAMB1 | chr7:107936279-107936373 | 0 | 0 | 0 | 0 | 0 | 0 | 0 | 0 | 18 | 19 | 20 | 37 | 42 | 28 | 22 | 23 | 16 | 20 | 17 | 7 | 14 | 1 | 0 | 0 | 0 | 0 | 0 | 0 | 0 | 0 | 1 | 1 | 0 | 0 | 0 | 0 | 1 | 0 | 0 | 1 | 0 | 0 | 0 | 0 | 0 | 0 | 0 | 0 | 0 | 0 | 1 |
| DAAM1 | chr14:59338385-59338414 | 0 | 0 | 0 | 0 | 0 | 0 | 0 | 0 | 18 | 34 | 11 | 77 | 77 | 32 | 32 | 24 | 49 | 16 | 11 | 10 | 29 | 0 | 0 | 0 | 0 | 0 | 0 | 0 | 0 | 0 | 0 | 0 | 0 | 0 | 0 | 0 | 0 | 0 | 0 | 2 | 0 | 0 | 0 | 0 | 0 | 0 | 0 | 0 | 0 | 0 | 1 |
| DLG1 | chr3:197068502-197068537 | 0 | 0 | 0 | 0 | 0 | 0 | 0 | 0 | 18 | 37 | 19 | 52 | 52 | 36 | 51 | 11 | 12 | 30 | 15 | 2 | 5 | 0 | 0 | 0 | 0 | 0 | 0 | 0 | 0 | 0 | 0 | 0 | 0 | 0 | 0 | 0 | 0 | 0 | 0 | 1 | 0 | 0 | 0 | 0 | 0 | 0 | 0 | 0 | 0 | 0 | 0 |
| PLEKHA5 | chr12:19270188-19270205 | 0 | 1 | 0 | 0 | 0 | 0 | 1 | 1 | 8 | 30 | 8 | 62 | 59 | 36 | 37 | 14 | 25 | 9 | 8 | 3 | 17 | 0 | 0 | 0 | 0 | 0 | 0 | 0 | 0 | 0 | 0 | 0 | 0 | 0 | 0 | 1 | 0 | 0 | 0 | 0 | 0 | 0 | 0 | 0 | 0 | 0 | 1 | 0 | 0 | 0 | 1 |
| CLASP1 | chr2:121382208-121382324 | 0 | 0 | 0 | 0 | 0 | 0 | 0 | 0 | 9 | 21 | 7 | 14 | 13 | 23 | 24 | 12 | 17 | 12 | 7 | 2 | 6 | 0 | 0 | 0 | 0 | 0 | 0 | 0 | 0 | 0 | 0 | 0 | 0 | 0 | 0 | 0 | 0 | 0 | 0 | 0 | 0 | 0 | 0 | 0 | 0 | 0 | 0 | 0 | 0 | 0 | 0 |
| ZYX | chr7:143387762-143387787 | 0 | 0 | 0 | 0 | 0 | 0 | 0 | 0 | 19 | 33 | 20 | 16 | 16 | 39 | 36 | 24 | 12 | 34 | 21 | 4 | 4 | 0 | 0 | 0 | 0 | 0 | 0 | 0 | 0 | 0 | 0 | 0 | 0 | 0 | 0 | 0 | 0 | 0 | 0 | 0 | 0 | 0 | 0 | 0 | 0 | 0 | 0 | 0 | 0 | 0 | 0 |
| DYM | chr18:49209551-49209715 | 0 | 0 | 0 | 0 | 0 | 0 | 0 | 0 | 13 | 26 | 13 | 39 | 40 | 32 | 33 | 15 | 21 | 14 | 12 | 3 | 14 | 0 | 0 | 0 | 0 | 0 | 0 | 0 | 0 | 0 | 0 | 0 | 0 | 0 | 0 | 0 | 0 | 0 | 0 | 1 | 0 | 0 | 0 | 0 | 0 | 0 | 1 | 0 | 0 | 0 | 0 |
| EXOC1 | chr4:55866872-55866892 | 0 | 0 | 0 | 0 | 0 | 0 | 0 | 0 | 7 | 21 | 5 | 47 | 44 | 19 | 27 | 11 | 22 | 12 | 7 | 6 | 8 | 0 | 0 | 1 | 0 | 0 | 0 | 0 | 0 | 0 | 1 | 0 | 0 | 0 | 0 | 1 | 0 | 0 | 0 | 1 | 0 | 0 | 0 | 0 | 0 | 0 | 0 | 0 | 0 | 0 | 0 |
| KIAA0141 | chr5:141933951-141933961 | 0 | 0 | 0 | 0 | 0 | 0 | 0 | 0 | 2 | 5 | 1 | 38 | 39 | 7 | 9 | 2 | 3 | 1 | 1 | 1 | 1 | 0 | 0 | 0 | 0 | 0 | 0 | 0 | 1 | 0 | 1 | 0 | 0 | 0 | 0 | 1 | 0 | 0 | 0 | 1 | 0 | 0 | 0 | 0 | 0 | 0 | 0 | 0 | 0 | 0 | 0 |
| DENND4A | chr15:65665199-65665225 | 0 | 1 | 1 | 0 | 0 | 0 | 0 | 0 | 14 | 37 | 10 | 67 | 63 | 45 | 51 | 21 | 34 | 13 | 10 | 10 | 23 | 0 | 0 | 1 | 1 | 0 | 0 | 0 | 0 | 0 | 0 | 0 | 0 | 0 | 0 | 0 | 0 | 0 | 0 | 2 | 0 | 1 | 1 | 1 | 1 | 0 | 0 | 0 | 0 | 1 | 1 |
| PLEC | chr8:143937749-143937784 | 0 | 0 | 0 | 0 | 0 | 0 | 1 | 0 | 9 | 25 | 18 | 60 | 61 | 32 | 33 | 17 | 20 | 19 | 24 | 5 | 12 | 0 | 0 | 1 | 1 | 0 | 0 | 1 | 0 | 1 | 1 | 0 | 1 | 0 | 0 | 0 | 0 | 0 | 0 | 1 | 1 | 0 | 0 | 1 | 0 | 0 | 2 | 0 | 0 | 0 | 0 |
| FBN1 | chr15:48434108-48434212 | 0 | 0 | 0 | 0 | 0 | 0 | 0 | 0 | 4 | 24 | 6 | 14 | 14 | 36 | 43 | 10 | 12 | 8 | 4 | 4 | 7 | 0 | 0 | 0 | 0 | 0 | 0 | 0 | 0 | 0 | 0 | 0 | 0 | 0 | 0 | 0 | 0 | 0 | 0 | 0 | 0 | 0 | 0 | 0 | 0 | 0 | 0 | 0 | 0 | 0 | 0 |
| SH3PXD2B | chr5:172356691-172356774 | 0 | 0 | 1 | 1 | 0 | 1 | 1 | 1 | 5 | 12 | 5 | 81 | 81 | 14 | 17 | 8 | 18 | 8 | 3 | 3 | 6 | 2 | 0 | 0 | 1 | 0 | 0 | 0 | 0 | 1 | 0 | 0 | 1 | 1 | 0 | 1 | 1 | 1 | 0 | 3 | 1 | 0 | 1 | 0 | 0 | 0 | 2 | 0 | 0 | 1 | 0 |
| FCHSD2 | chr11:72900257-72900328 | 0 | 0 | 0 | 0 | 0 | 0 | 0 | 0 | 2 | 6 | 2 | 28 | 27 | 6 | 7 | 2 | 10 | 2 | 1 | 1 | 4 | 0 | 0 | 0 | 0 | 0 | 0 | 0 | 0 | 0 | 0 | 0 | 1 | 0 | 0 | 0 | 0 | 0 | 0 | 1 | 0 | 0 | 0 | 0 | 0 | 0 | 1 | 0 | 0 | 0 | 0 |
| ENAH | chr1:225531046-225531057 | 0 | 0 | 0 | 0 | 0 | 0 | 0 | 0 | 3 | 7 | 2 | 18 | 20 | 6 | 13 | 5 | 10 | 6 | 3 | 1 | 4 | 0 | 0 | 0 | 0 | 0 | 0 | 0 | 0 | 0 | 0 | 0 | 0 | 0 | 0 | 0 | 0 | 0 | 0 | 0 | 0 | 0 | 0 | 0 | 0 | 0 | 0 | 0 | 0 | 0 | 1 |
| USP24 | chr1:55084393-55084421 | 0 | 0 | 0 | 0 | 0 | 0 | 0 | 0 | 1 | 5 | 5 | 28 | 30 | 8 | 6 | 3 | 5 | 12 | 5 | 1 | 1 | 0 | 0 | 0 | 0 | 0 | 0 | 0 | 0 | 0 | 0 | 0 | 0 | 0 | 0 | 0 | 0 | 0 | 0 | 1 | 0 | 0 | 0 | 0 | 0 | 0 | 0 | 0 | 0 | 0 | 0 |
| TBC1D31 | chr8:123126354-123126378 | 0 | 0 | 0 | 0 | 0 | 0 | 1 | 0 | 3 | 15 | 4 | 35 | 32 | 12 | 15 | 4 | 3 | 3 | 3 | 2 | 2 | 0 | 0 | 0 | 0 | 0 | 0 | 0 | 0 | 0 | 0 | 0 | 0 | 0 | 0 | 0 | 0 | 0 | 0 | 1 | 0 | 0 | 0 | 0 | 0 | 0 | 0 | 0 | 0 | 0 | 0 |
| AFAP1L2 | chr10:114319562-114319645 | 1 | 1 | 1 | 2 | 2 | 2 | 1 | 1 | 5 | 14 | 5 | 80 | 78 | 22 | 24 | 4 | 5 | 5 | 5 | 5 | 4 | 2 | 0 | 2 | 1 | 2 | 1 | 2 | 2 | 2 | 3 | 1 | 1 | 1 | 2 | 2 | 2 | 1 | 1 | 1 | 1 | 1 | 1 | 1 | 1 | 2 | 1 | 1 | 2 | 1 | 2 |
| SORBS2 | chr4:185622914-185624494 | 1 | 0 | 0 | 0 | 0 | 0 | 0 | 0 | 47 | 66 | 62 | 41 | 57 | 59 | 73 | 44 | 40 | 78 | 64 | 12 | 17 | 0 | 0 | 0 | 0 | 0 | 0 | 0 | 0 | 0 | 0 | 0 | 0 | 1 | 0 | 0 | 0 | 0 | 0 | 2 | 0 | 1 | 0 | 0 | 0 | 0 | 1 | 0 | 0 | 0 | 2 |
| CALCOCO2 | chr17:48856579-48856616 | 1 | 0 | 0 | 0 | 0 | 0 | 0 | 1 | 9 | 22 | 3 | 62 | 59 | 27 | 30 | 9 | 19 | 7 | 3 | 2 | 5 | 0 | 0 | 0 | 0 | 0 | 0 | 0 | 0 | 0 | 0 | 0 | 0 | 0 | 0 | 0 | 0 | 0 | 0 | 3 | 0 | 0 | 0 | 0 | 0 | 0 | 0 | 0 | 0 | 0 | 0 |
| PTBP1 | chr19:805832-805902 | 0 | 0 | 0 | 1 | 1 | 0 | 1 | 0 | 3 | 7 | 6 | 58 | 53 | 13 | 13 | 4 | 4 | 10 | 7 | 1 | 1 | 0 | 0 | 1 | 0 | 1 | 0 | 0 | 1 | 0 | 0 | 0 | 0 | 0 | 0 | 3 | 0 | 0 | 0 | 1 | 0 | 0 | 0 | 0 | 0 | 0 | 0 | 0 | 1 | 0 | 0 |
| STIM1 | chr11:4088702-4088738 | 0 | 0 | 0 | 0 | 0 | 0 | 0 | 0 | 4 | 11 | 8 | 19 | 22 | 12 | 13 | 7 | 8 | 6 | 8 | 4 | 10 | 0 | 0 | 0 | 0 | 0 | 0 | 0 | 0 | 0 | 0 | 0 | 0 | 0 | 0 | 0 | 0 | 0 | 0 | 1 | 0 | 0 | 0 | 0 | 0 | 0 | 0 | 0 | 0 | 0 | 0 |
| CKAP5 | chr11:46760352-46760375 | 0 | 0 | 0 | 0 | 0 | 0 | 0 | 0 | 1 | 2 | 1 | 34 | 32 | 2 | 3 | 1 | 4 | 1 | 1 | 1 | 2 | 0 | 0 | 1 | 1 | 0 | 0 | 0 | 0 | 0 | 0 | 0 | 0 | 0 | 0 | 0 | 0 | 0 | 0 | 1 | 0 | 0 | 0 | 1 | 0 | 1 | 0 | 0 | 0 | 0 | 0 |
| EPB41L3 | chr18:5415818-5416378 | 0 | 0 | 0 | 0 | 0 | 0 | 0 | 0 | 6 | 18 | 12 | 10 | 12 | 18 | 20 | 6 | 8 | 15 | 8 | 1 | 3 | 0 | 0 | 0 | 0 | 0 | 0 | 0 | 0 | 0 | 0 | 0 | 0 | 0 | 0 | 0 | 0 | 0 | 0 | 0 | 0 | 0 | 0 | 0 | 0 | 0 | 0 | 0 | 0 | 0 | 0 |
| MAPKBP1 | chr15:41813330-41813347 | 0 | 0 | 0 | 0 | 0 | 0 | 0 | 0 | 19 | 53 | 31 | 91 | 91 | 65 | 65 | 20 | 30 | 38 | 26 | 12 | 17 | 0 | 0 | 1 | 0 | 0 | 0 | 0 | 0 | 2 | 5 | 0 | 0 | 0 | 0 | 3 | 0 | 0 | 0 | 0 | 0 | 0 | 1 | 0 | 0 | 0 | 0 | 0 | 0 | 0 | 1 |
| SEMA4B | chr15:90214937-90214964 | 0 | 0 | 0 | 0 | 0 | 0 | 0 | 0 | 6 | 11 | 4 | 55 | 45 | 17 | 14 | 8 | 13 | 5 | 4 | 2 | 4 | 0 | 0 | 0 | 0 | 0 | 0 | 0 | 0 | 1 | 1 | 0 | 0 | 0 | 0 | 1 | 0 | 0 | 0 | 1 | 0 | 0 | 0 | 0 | 0 | 0 | 0 | 0 | 0 | 0 | 0 |
| ZFYVE27 | chr10:97752857-97752877 | 1 | 1 | 1 | 1 | 0 | 0 | 1 | 1 | 31 | 68 | 32 | 81 | 76 | 71 | 77 | 45 | 40 | 28 | 31 | 10 | 27 | 1 | 0 | 1 | 1 | 1 | 1 | 1 | 0 | 2 | 2 | 1 | 1 | 1 | 1 | 4 | 2 | 0 | 1 | 2 | 1 | 1 | 1 | 1 | 1 | 1 | 0 | 1 | 0 | 1 | 1 |
| CLASP2 | chr3:33574473\|33573735-33574496 | 0 | 0 | 1 | 0 | 0 | 0 | 0 | 0 | 12 | 34 | 8 | 61 | 56 | 26 | 37 | 13 | 18 | 11 | 8 | 4 | 9 | 0 | 0 | 1 | 1 | 0 | 0 | 0 | 0 | 0 | 0 | 0 | 0 | 0 | 0 | 0 | 0 | 0 | 0 | 2 | 0 | 0 | 1 | 1 | 0 | 1 | 3 | 0 | 0 | 0 | 1 |
| SH3GLB2 | chr9:129013008-129013019 | 0 | 0 | 0 | 0 | 0 | 0 | 0 | 0 | 36 | 60 | 27 | 71 | 70 | 60 | 65 | 28 | 39 | 36 | 24 | 3 | 20 | 0 | 0 | 1 | 0 | 0 | 1 | 0 | 0 | 0 | 0 | 0 | 0 | 0 | 0 | 0 | 0 | 0 | 0 | 4 | 0 | 1 | 1 | 0 | 0 | 0 | 1 | 0 | 0 | 1 | 0 |
| EDEM3 | chr1:184701497-184701544 | 0 | 0 | 0 | 0 | 0 | 0 | 0 | 0 | 4 | 25 | 6 | 67 | 56 | 22 | 40 | 10 | 18 | 12 | 5 | 2 | 6 | 0 | 0 | 0 | 0 | 0 | 0 | 0 | 0 | 0 | 0 | 0 | 0 | 0 | 0 | 0 | 0 | 0 | 0 | 4 | 0 | 0 | 0 | 0 | 0 | 0 | 0 | 0 | 0 | 0 | 0 |
| LRCH1 | chr13:46715560\|46715607-46715664 | 2 | 2 | 1 | 1 | 1 | 1 | 1 | 3 | 10 | 32 | 13 | 92 | 90 | 47 | 50 | 19 | 36 | 24 | 14 | 9 | 13 | 2 | 1 | 1 | 2 | 1 | 1 | 1 | 3 | 2 | 2 | 1 | 1 | 2 | 1 | 5 | 2 | 1 | 2 | 2 | 1 | 2 | 2 | 4 | 1 | 2 | 1 | 1 | 1 | 1 | 1 |
| NRBP1 | chr2:27435723-27435746 | 0 | 0 | 0 | 0 | 0 | 0 | 0 | 0 | 6 | 12 | 7 | 32 | 30 | 14 | 18 | 8 | 15 | 9 | 7 | 5 | 10 | 0 | 0 | 0 | 0 | 0 | 0 | 0 | 0 | 0 | 0 | 0 | 0 | 0 | 0 | 1 | 0 | 0 | 0 | 2 | 0 | 0 | 0 | 0 | 0 | 0 | 0 | 0 | 0 | 0 | 0 |
| KCNMA1 | chr10:76914057-76914137 | 0 | 0 | 1 | 0 | 0 | 0 | 0 | 0 | 30 | 60 | 41 | 30 | 25 | 62 | 71 | 49 | 56 | 53 | 42 | 7 | 30 | 0 | 0 | 0 | 0 | 0 | 0 | 0 | 0 | 1 | 0 | 0 | 0 | 0 | 0 | 0 | 0 | 0 | 0 | 1 | 0 | 0 | 0 | 1 | 0 | 0 | 0 | 0 | 0 | 0 | 1 |
| STX2 | chr12:130799168-130799190 | 0 | 0 | 0 | 0 | 0 | 0 | 0 | 0 | 22 | 34 | 10 | 73 | 68 | 34 | 46 | 22 | 22 | 20 | 13 | 1 | 8 | 0 | 0 | 0 | 0 | 0 | 0 | 0 | 0 | 0 | 0 | 0 | 0 | 0 | 0 | 0 | 0 | 0 | 0 | 5 | 0 | 0 | 0 | 0 | 0 | 0 | 0 | 0 | 0 | 0 | 0 |
| ANK3 | chr10:60263126-60263149 | 0 | 0 | 1 | 0 | 0 | 0 | 0 | 0 | 3 | 17 | 6 | 30 | 29 | 20 | 22 | 11 | 11 | 5 | 4 | 7 | 22 | 0 | 0 | 0 | 0 | 0 | 0 | 0 | 0 | 0 | 0 | 0 | 0 | 0 | 0 | 0 | 0 | 0 | 0 | 2 | 0 | 0 | 0 | 0 | 0 | 0 | 0 | 0 | 0 | 0 | 0 |
| MAPK9 | chr5:180247855-180247926 | 1 | 1 | 2 | 1 | 1 | 1 | 0 | 1 | 34 | 58 | 30 | 55 | 49 | 55 | 62 | 41 | 54 | 32 | 30 | 13 | 38 | 0 | 0 | 1 | 1 | 1 | 1 | 1 | 1 | 1 | 1 | 2 | 1 | 2 | 1 | 1 | 0 | 1 | 1 | 3 | 1 | 1 | 1 | 1 | 2 | 1 | 1 | 1 | 1 | 1 | 2 |
| TPD52L2 | chr20:63886016-63887605 | 0 | 0 | 1 | 0 | 0 | 0 | 0 | 0 | 26 | 51 | 29 | 76 | 76 | 57 | 62 | 30 | 44 | 42 | 28 | 8 | 20 | 0 | 0 | 1 | 1 | 0 | 2 | 0 | 0 | 0 | 0 | 0 | 0 | 0 | 0 | 0 | 0 | 0 | 0 | 4 | 0 | 0 | 0 | 1 | 0 | 1 | 4 | 0 | 0 | 1 | 0 |
| ABCA6 | chr17:69079958-69080027\|69080137 | 0 | 0 | 0 | 0 | 0 | 0 | 0 | 0 | 25 | 28 | 26 | 24 | 13 | 17 | 19 | 40 | 35 | 28 | 27 | 42 | 44 | 0 | 0 | 0 | 0 | 0 | 0 | 0 | 0 | 0 | 1 | 0 | 0 | 0 | 0 | 1 | 0 | 0 | 0 | 0 | 0 | 0 | 0 | 0 | 0 | 0 | 0 | 0 | 0 | 0 | 0 |
| FMNL2 | chr2:152622537-152622557 | 0 | 0 | 0 | 0 | 0 | 0 | 0 | 0 | 1 | 4 | 1 | 32 | 36 | 5 | 6 | 1 | 2 | 3 | 1 | 0 | 0 | 0 | 0 | 0 | 0 | 0 | 0 | 0 | 0 | 0 | 0 | 0 | 0 | 0 | 0 | 0 | 0 | 0 | 0 | 3 | 0 | 0 | 0 | 0 | 0 | 0 | 0 | 0 | 0 | 0 | 0 |
| DNAJC13 | chr3:132537158-132537266 | 0 | 0 | 0 | 0 | 0 | 0 | 0 | 0 | 8 | 17 | 7 | 28 | 30 | 18 | 20 | 10 | 16 | 8 | 7 | 2 | 6 | 0 | 0 | 0 | 0 | 0 | 0 | 0 | 0 | 1 | 1 | 0 | 0 | 0 | 0 | 0 | 0 | 0 | 0 | 1 | 0 | 0 | 0 | 0 | 0 | 0 | 2 | 0 | 0 | 0 | 0 |
| PLS3 | chrX:115640110-115640136 | 0 | 0 | 0 | 0 | 0 | 0 | 0 | 0 | 13 | 38 | 2 | 20 | 19 | 33 | 53 | 13 | 9 | 2 | 2 | 1 | 7 | 0 | 0 | 1 | 1 | 0 | 0 | 0 | 0 | 0 | 0 | 0 | 0 | 0 | 0 | 0 | 0 | 0 | 0 | 1 | 0 | 0 | 0 | 1 | 0 | 0 | 0 | 0 | 0 | 0 | 0 |
| CFAP43 | chr10:104130738-104130830 | 1 | 1 | 0 | 0 | 0 | 1 | 0 | 0 | 10 | 42 | 10 | 37 | 36 | 46 | 47 | 9 | 8 | 27 | 46 | 3 | 10 | 0 | 0 | 0 | 0 | 0 | 1 | 1 | 0 | 2 | 1 | 0 | 0 | 0 | 0 | 1 | 0 | 0 | 0 | 0 | 0 | 0 | 0 | 0 | 0 | 1 | 0 | 0 | 0 | 0 | 1 |
| DCTN4 | chr5:150731886-150731906 | 0 | 0 | 0 | 0 | 0 | 0 | 0 | 0 | 6 | 11 | 4 | 33 | 34 | 9 | 12 | 6 | 5 | 6 | 4 | 1 | 1 | 0 | 0 | 0 | 0 | 0 | 0 | 0 | 0 | 0 | 0 | 0 | 0 | 0 | 0 | 0 | 0 | 0 | 0 | 1 | 0 | 0 | 0 | 0 | 0 | 0 | 3 | 0 | 0 | 0 | 0 |
| GOLT1B | chr12:21515238-21515270 | 0 | 0 | 0 | 0 | 0 | 0 | 0 | 0 | 18 | 32 | 16 | 14 | 12 | 32 | 37 | 20 | 26 | 20 | 16 | 3 | 16 | 0 | 0 | 1 | 0 | 0 | 0 | 0 | 0 | 0 | 0 | 0 | 0 | 0 | 0 | 0 | 0 | 0 | 0 | 1 | 0 | 0 | 0 | 0 | 0 | 0 | 0 | 0 | 0 | 0 | 0 |
| SIRT7 | chr17:81913207-81913293 | 0 | 1 | 2 | 1 | 1 | 1 | 1 | 1 | 11 | 19 | 10 | 67 | 66 | 25 | 25 | 12 | 28 | 12 | 8 | 6 | 11 | 1 | 1 | 1 | 1 | 1 | 1 | 2 | 1 | 2 | 3 | 1 | 1 | 1 | 1 | 4 | 1 | 1 | 1 | 4 | 2 | 0 | 0 | 1 | 1 | 1 | 4 | 1 | 1 | 1 | 1 |
| SYNJ1 | chr21:32642095-32645006 | 2 | 2 | 3 | 2 | 1 | 3 | 3 | 2 | 62 | 86 | 65 | 91 | 88 | 83 | 90 | 72 | 65 | 75 | 66 | 26 | 66 | 1 | 1 | 5 | 4 | 2 | 1 | 2 | 5 | 4 | 3 | 4 | 3 | 3 | 2 | 6 | 2 | 2 | 2 | 6 | 1 | 3 | 4 | 3 | 1 | 3 | 3 | 1 | 1 | 2 | 4 |
| DYSF | chr2:71608576-71608617 | 1 | 1 | 1 | 0 | 1 | 1 | 1 | 0 | 5 | 13 | 6 | 42 | 39 | 13 | 17 | 5 | 16 | 8 | 6 | 6 | 15 | 3 | 0 | 1 | 1 | 1 | 1 | 1 | 1 | 2 | 2 | 0 | 1 | 1 | 2 | 2 | 0 | 1 | 1 | 1 | 1 | 1 | 1 | 1 | 1 | 1 | 1 | 1 | 1 | 1 | 1 |
| EXOC7 | chr17:76086854-76086892 | 0 | 0 | 0 | 0 | 0 | 0 | 0 | 0 | 37 | 60 | 36 | 87 | 87 | 67 | 70 | 48 | 57 | 43 | 36 | 51 | 44 | 0 | 0 | 1 | 1 | 0 | 0 | 0 | 0 | 5 | 7 | 0 | 0 | 0 | 0 | 1 | 0 | 0 | 0 | 1 | 0 | 0 | 0 | 1 | 0 | 0 | 1 | 0 | 0 | 0 | 0 |
| PLCH1 | chr3:155483349-155483376 | 2 | 0 | 0 | 0 | 0 | 2 | 0 | 0 | 16 | 87 | 22 | 86 | 67 | 75 | 88 | 32 | 51 | 34 | 39 | -1 | 69 | 0 | 0 | 0 | 0 | 0 | 1 | 0 | 0 | 0 | 1 | 0 | -1 | 0 | 0 | 0 | 0 | 0 | 0 | 0 | 0 | 1 | 6 | 0 | 0 | 0 | 0 | 0 | 0 | 0 | -1 |
| PLEKHG3 | chr14:64729960-64729981 | 0 | 0 | 1 | 0 | 0 | 0 | 0 | 0 | 1 | 2 | 1 | 21 | 24 | 1 | 2 | 1 | 2 | 2 | 1 | 1 | 1 | 0 | 0 | 0 | 0 | 0 | 0 | 0 | 0 | 0 | 0 | 0 | 0 | 0 | 0 | 0 | 0 | 0 | 0 | 2 | 0 | 0 | 0 | 0 | 0 | 0 | 1 | 0 | 0 | 0 | 0 |
| PHTF1 | chr1:113719876-113719929\|113719926 | 1 | 1 | 1 | 0 | 0 | 0 | 0 | 1 | 6 | 11 | 11 | 36 | 42 | 17 | 12 | 8 | 5 | 13 | 12 | 5 | 3 | 0 | -1 | 1 | 1 | 1 | 1 | 1 | 0 | 1 | 1 | 1 | 0 | 1 | 1 | 3 | 2 | 1 | 1 | 2 | 1 | 1 | 1 | 2 | 2 | 1 | 2 | 0 | 1 | 1 | 2 |
| BICD2 | chr9:92713879-92713924 | 1 | 1 | 0 | 1 | 1 | 1 | 1 | 1 | 3 | 5 | 4 | 35 | 33 | 5 | 6 | 4 | 2 | 7 | 3 | 1 | 1 | 0 | 0 | 1 | 1 | 1 | 0 | 1 | 2 | 1 | 1 | 1 | 0 | 1 | 1 | 1 | 1 | 1 | 1 | 0 | 1 | 0 | 0 | 1 | 0 | 1 | 2 | 1 | 1 | 1 | 0 |
| LIMA1 | chr12:50185390-50185448 | 0 | 0 | 0 | 0 | 0 | 0 | 0 | 0 | 4 | 6 | 9 | 58 | 59 | 15 | 11 | 4 | 8 | 11 | 9 | 5 | 6 | 0 | 0 | 0 | 0 | 0 | 0 | 0 | 0 | 1 | 1 | 0 | 0 | 0 | 0 | 6 | 0 | 0 | 0 | 0 | 0 | 0 | 0 | 0 | 0 | 0 | 0 | 0 | 0 | 0 | 1 |
| MLLT4 | chr6:167954458-167954493 | 0 | 0 | 1 | 0 | 0 | 0 | 0 | 0 | 16 | 38 | 16 | 68 | 67 | 35 | 46 | 24 | 34 | 29 | 14 | 5 | 14 | 0 | 0 | 1 | 0 | 0 | 0 | 0 | 0 | 2 | 2 | 0 | 0 | 0 | 0 | 5 | 0 | 0 | 0 | 5 | 0 | 0 | 0 | 0 | 0 | 0 | 0 | 0 | 0 | 0 | 1 |
| ARHGEF7 | chr13:111292118-111292294 | 0 | 0 | 0 | 0 | 0 | 0 | 0 | 0 | 33 | 43 | 34 | 85 | 81 | 48 | 53 | 46 | 43 | 40 | 37 | 48 | 41 | 0 | 0 | 1 | 0 | 0 | 0 | 0 | 0 | 2 | 1 | 0 | 0 | 0 | 0 | 5 | 2 | 0 | 0 | 3 | 0 | 0 | 0 | 0 | 0 | 0 | 7 | 0 | 0 | 0 | 0 |
| PATJ | chr1:61833358-61833377 | 0 | 0 | 0 | 0 | 0 | 0 | 0 | 0 | 3 | 13 | 13 | 85 | 82 | 27 | 32 | 1 | 10 | 16 | 8 | 7 | 0 | 0 | 0 | 0 | 0 | 0 | 0 | 0 | 0 | 0 | 0 | 0 | 0 | 0 | 0 | 0 | 0 | 0 | 0 | 9 | 0 | 0 | 0 | 0 | 0 | 0 | 0 | 0 | 0 | 0 | 0 |
| SYNE1 | chr6:152511571-152511591 | 1 | 2 | 4 | 0 | 3 | 2 | 0 | 1 | 9 | 29 | 16 | 90 | 86 | 46 | 43 | 23 | 26 | 21 | 15 | 25 | 19 | 0 | 0 | 3 | 5 | 0 | 1 | 1 | 0 | 3 | 3 | 0 | -1 | 2 | 0 | 2 | 2 | 1 | 0 | 6 | 0 | 0 | 1 | -1 | 0 | 0 | 0 | 1 | 1 | 2 | 0 |
| VTI1A | chr10:112533530-112533550 | 0 | 0 | 1 | 0 | 0 | 0 | 1 | 0 | 8 | 16 | 5 | 35 | 40 | 16 | 16 | 10 | 12 | 8 | 4 | 2 | 6 | 1 | 0 | 0 | 0 | 0 | 0 | 0 | 0 | 1 | 1 | 2 | 0 | 0 | 0 | 4 | 0 | 0 | 0 | 2 | 0 | 0 | 0 | 0 | 0 | 0 | 0 | 0 | 0 | 0 | 0 |
| REEP6 | chr19:1496591-1496671 | 0 | 0 | 0 | 0 | 1 | 0 | 0 | 0 | 47 | 50 | 57 | 78 | 73 | 59 | 57 | 61 | 34 | 55 | 59 | 10 | 33 | 0 | 0 | 4 | 2 | 1 | 0 | 1 | 0 | 1 | 3 | 0 | 0 | 1 | 0 | 7 | 0 | 0 | 0 | 3 | 0 | 0 | 1 | 0 | 0 | 1 | 0 | 0 | 0 | 0 | 1 |
| MANBA | chr4:102672009-102672054 | 1 | 1 | 0 | 1 | 1 | 1 | 0 | 1 | 3 | 7 | 2 | 19 | 25 | 11 | 5 | 1 | 3 | 4 | 2 | 2 | 3 | 1 | 0 | 1 | 1 | 1 | 1 | 1 | 0 | 1 | 1 | 1 | 1 | 1 | 0 | 2 | 1 | 1 | 1 | 2 | 1 | 1 | 2 | 1 | 1 | 0 | 1 | 1 | 2 | 1 | 1 |
| SBF1 | chr22:50466978-50467001 | 0 | 0 | 2 | 0 | 0 | 0 | 0 | 0 | 10 | 18 | 10 | 55 | 55 | 24 | 23 | 10 | 40 | 10 | 10 | 13 | 38 | 0 | 0 | 2 | 2 | 1 | 0 | 1 | 0 | 0 | 0 | 0 | 0 | 1 | 0 | 0 | 0 | 0 | 0 | 6 | 0 | 0 | 0 | 1 | 0 | 1 | 1 | 0 | 0 | 1 | 0 |
| CLASP2 | chr3:33577202-33577264 | 1 | 0 | 2 | 0 | 0 | 0 | 0 | 1 | 44 | 32 | 35 | 59 | 52 | 35 | 32 | 49 | 56 | 27 | 43 | 76 | 68 | 1 | 0 | 4 | 4 | 1 | 0 | 2 | 0 | 0 | 0 | 0 | 0 | 0 | 0 | 0 | 1 | 0 | 0 | 1 | 0 | 0 | 2 | 2 | 0 | 2 | 3 | 0 | 0 | 1 | 1 |
| TMEM39A | chr3:119437588-119437649 | 2 | 1 | 4 | 2 | 2 | 1 | 1 | 1 | 5 | 17 | 10 | 78 | 77 | 23 | 28 | 9 | 20 | 17 | 11 | 3 | 6 | 1 | 1 | 2 | 2 | 2 | 1 | 2 | 1 | 1 | 2 | 3 | 6 | 2 | 2 | 2 | 1 | 1 | 3 | 9 | 1 | 1 | 1 | 2 | 4 | 2 | 2 | 1 | 2 | 1 | 3 |
| ITPR1 | chr3:4717367-4727173 | 2 | 2 | 4 | 1 | 1 | 1 | 1 | 3 | 67 | 94 | 80 | 93 | 98 | 93 | 95 | 74 | 45 | 88 | 80 | 40 | 26 | 1 | 2 | 1 | 1 | 1 | 3 | 1 | 1 | 2 | 2 | 3 | 9 | 2 | 2 | 10 | 3 | 1 | 3 | 5 | 1 | 2 | 3 | 2 | 3 | 2 | 2 | 2 | 1 | 1 | 4 |
| DGKB | chr7:14753928-14753948 | 1 | 1 | -1 | 0 | 0 | 1 | 0 | 0 | 24 | 30 | 33 | 12 | 33 | 34 | 34 | 21 | 10 | 30 | 45 | 0 | 6 | 0 | 0 | 1 | 1 | 0 | 1 | 0 | 0 | 1 | 0 | 0 | 0 | -1 | 0 | 3 | 0 | 0 | 0 | 1 | 0 | 0 | 0 | 0 | 0 | 0 | 0 | 0 | 0 | 0 | -1 |
| RELN | chr7:103478389-103478524\|103482270\|103478394 | 0 | 0 | 1 | 0 | 0 | 0 | 0 | 0 | 9 | 19 | 25 | 40 | 43 | 18 | 19 | 7 | 17 | 30 | 21 | 6 | 8 | 0 | 0 | 0 | 0 | 0 | 0 | 0 | 0 | 0 | 0 | 0 | 0 | 0 | 0 | 1 | 0 | 0 | 1 | 0 | 0 | 0 | 1 | 0 | 0 | 0 | 4 | 0 | 0 | 0 | 3 |
| ADGRL2 | chr1:81942034\|81942031\|81941193-81942045 | 1 | 1 | 0 | 0 | 1 | 0 | 2 | 1 | 40 | 70 | 33 | 52 | 72 | 77 | 78 | 26 | 40 | 55 | 47 | 7 | 37 | 0 | 0 | 1 | 1 | 1 | 3 | 0 | 0 | 3 | 3 | 2 | 1 | 0 | 2 | 3 | 1 | 0 | 3 | 3 | 1 | 6 | 4 | 1 | 0 | 2 | 2 | 3 | 1 | 3 | -1 |
| CHPT1 | chr12:101719490-101719516 | 0 | 0 | 0 | 0 | 0 | 0 | 0 | 0 | 7 | 11 | 4 | 44 | 42 | 13 | 17 | 7 | 7 | 4 | 4 | 1 | 3 | 0 | 0 | 0 | 0 | 0 | 0 | 0 | 0 | 0 | 0 | 0 | 0 | 0 | 0 | 0 | 0 | 0 | 0 | 6 | 0 | 0 | 0 | 0 | 0 | 0 | 1 | 0 | 0 | 0 | 0 |
| NECAP2 | chr1:16450125-16450241 | 1 | 2 | 1 | 1 | 1 | 1 | 2 | 1 | 5 | 8 | 4 | 47 | 45 | 10 | 9 | 4 | 8 | 6 | 4 | 2 | 3 | 2 | 2 | 3 | 1 | 2 | 1 | 2 | 2 | 3 | 3 | 1 | 1 | 1 | 1 | 3 | 1 | 4 | 1 | 4 | 2 | 1 | 1 | 2 | 1 | 1 | 3 | 2 | 3 | 2 | 1 |
| MYH10 | chr17:8530623-8530685 | 0 | 0 | 0 | 0 | 0 | 0 | 0 | 0 | 15 | 47 | 27 | 22 | 20 | 50 | 59 | 26 | 35 | 32 | 28 | 11 | 35 | 0 | 1 | 0 | 0 | 0 | 0 | 0 | 0 | 1 | 2 | 0 | 0 | 0 | 0 | 2 | 0 | 0 | 0 | 0 | 1 | 0 | 0 | 0 | 0 | 0 | 0 | 0 | 0 | 0 | 0 |
| ASAP2 | chr2:9336036-9336044 | 0 | 0 | 1 | 0 | 0 | 0 | 0 | 0 | 8 | 27 | 19 | 85 | 72 | 26 | 44 | 12 | 29 | 31 | 18 | 3 | 13 | 0 | 0 | 1 | 0 | 0 | 0 | 0 | 0 | 0 | 0 | 0 | 0 | 0 | 0 | 0 | 0 | 0 | 0 | 10 | 0 | 0 | 0 | 0 | 0 | 0 | 0 | 0 | 0 | 0 | 1 |
| SSR1 | chr6:7295088-7295102 | 0 | 0 | 1 | 0 | 0 | 0 | 1 | 0 | 9 | 28 | 14 | 64 | 62 | 33 | 42 | 13 | 28 | 20 | 14 | 3 | 11 | 0 | 0 | 1 | 0 | 0 | 0 | 0 | 0 | 1 | 2 | 0 | 0 | 0 | 0 | 4 | 0 | 0 | 1 | 8 | 0 | 0 | 0 | 0 | 0 | 1 | 1 | 0 | 0 | 0 | 0 |
| MYO1D | chr17:32745522-32745689 | 0 | 0 | 0 | 0 | 0 | 0 | 0 | 0 | 5 | 16 | 5 | 45 | 73 | 23 | 22 | 2 | 5 | 7 | 3 | 0 | 1 | 0 | 0 | 0 | 0 | 0 | 0 | 0 | 0 | 9 | 4 | 0 | 0 | 0 | 0 | 5 | 0 | 0 | 0 | 0 | 0 | 0 | 0 | 0 | 0 | 0 | 1 | 0 | 0 | 0 | 0 |
| SHANK2 | chr11:70489328-70489348 | 1 | 0 | 0 | -1 | 0 | 0 | 0 | 0 | 48 | 77 | 41 | 89 | 93 | 78 | 77 | 61 | 39 | 33 | 40 | 41 | 48 | -1 | 0 | 2 | 2 | 0 | 0 | 0 | 0 | 0 | 0 | 0 | 0 | 0 | 0 | -1 | 0 | 0 | 0 | 2 | 0 | 1 | 1 | 1 | 0 | 0 | 13 | 0 | 0 | -1 | -1 |
| DOPEY1 | chr6:83125166-83125195 | 0 | 0 | 1 | 0 | 0 | 0 | 0 | 0 | 19 | 67 | 37 | 95 | 93 | 67 | 77 | 35 | 54 | 58 | 47 | 7 | 17 | 0 | 0 | 1 | 1 | 0 | 0 | 0 | 0 | 2 | 4 | 0 | 0 | 0 | 0 | 13 | 0 | 0 | 0 | 3 | 0 | 0 | 0 | 0 | 0 | 0 | 1 | 0 | 0 | 1 | 0 |
| MFSD6 | chr2:190499832-190499856 | 0 | 0 | 1 | 0 | 0 | 0 | 0 | 0 | 34 | 79 | 34 | 81 | 86 | 77 | 83 | 34 | 45 | 45 | 29 | 3 | 33 | 0 | 1 | 4 | 2 | 1 | 0 | 1 | 0 | 0 | 0 | 0 | 0 | 0 | 0 | 0 | 0 | 0 | 1 | 12 | 0 | 0 | 0 | 2 | 0 | 1 | 0 | 0 | 0 | 1 | 1 |
| CLTB | chr5:176396479-176396532 | 0 | 0 | 1 | 0 | 0 | 0 | 1 | 1 | 58 | 73 | 50 | 91 | 89 | 79 | 82 | 60 | 69 | 58 | 49 | 20 | 51 | 0 | 1 | 2 | 2 | 1 | 1 | 1 | 1 | 1 | 1 | 1 | 1 | 0 | 1 | 0 | 0 | 1 | 3 | 10 | 1 | 1 | 1 | 2 | 0 | 2 | 9 | 2 | 0 | 1 | 0 |
| CNTRL | chr9:121110515-121110571 | 0 | 0 | 1 | 0 | 0 | 0 | 0 | 0 | 7 | 18 | 10 | 80 | 76 | 30 | 25 | 8 | 21 | 13 | 8 | 5 | 1 | 0 | 0 | 0 | 0 | 0 | 0 | 0 | 0 | 3 | 5 | 0 | 0 | 0 | 0 | 10 | 0 | 0 | 1 | 3 | 0 | 0 | 0 | 0 | 0 | 0 | 2 | 0 | 0 | 0 | 0 |
| EIF4G1 | chr3:184316713-184316733 | 0 | 0 | 1 | 0 | 0 | 0 | 0 | 0 | 16 | 42 | 14 | 82 | 78 | 43 | 50 | 24 | 28 | 25 | 12 | 5 | 13 | 0 | 0 | 1 | 0 | 0 | 0 | 0 | 0 | 0 | 0 | 0 | 0 | 0 | 0 | 2 | 0 | 0 | 0 | 12 | 0 | 0 | 0 | 0 | 0 | 1 | 0 | 0 | 0 | 0 | 0 |
| GTF2I | chr7:74714857\|74714860-74714916 | 11 | 8 | 5 | 5 | 5 | 5 | 4 | 9 | 39 | 54 | 36 | 86 | 79 | 52 | 64 | 42 | 58 | 46 | 40 | 34 | 37 | 5 | 3 | 6 | 5 | 5 | 5 | 5 | 6 | 7 | 8 | 6 | 4 | 8 | 6 | 8 | 8 | 5 | 5 | 11 | 6 | 6 | 7 | 6 | 6 | 6 | 11 | 5 | 6 | 5 | 7 |
| HERC1 | chr15:63702138-63702203 | 0 | 0 | 0 | 0 | 0 | 0 | 0 | 0 | 7 | 17 | 8 | 9 | 10 | 26 | 31 | 7 | 15 | 6 | 8 | 1 | 8 | 0 | 0 | 0 | 0 | 0 | 0 | 0 | 0 | 0 | 0 | 0 | 0 | 0 | 0 | 2 | 0 | 0 | 0 | 0 | 0 | 0 | 0 | 0 | 0 | 0 | 0 | 0 | 0 | 0 | 0 |
| TACC1 | chr8:38823990-38824025 | 1 | 1 | 1 | 2 | 1 | 1 | 2 | 1 | 5 | 23 | 5 | 45 | 43 | 26 | 36 | 8 | 13 | 14 | 7 | 1 | 5 | 1 | 1 | 2 | 2 | 2 | 4 | 2 | 1 | 2 | 3 | 0 | 2 | 2 | 2 | 1 | 0 | 1 | 3 | 5 | 2 | 1 | 1 | 1 | 0 | 2 | 0 | 1 | 1 | 2 | 0 |
| PPP6R3 | chr11:68609611\|68609473-68609628 | 0 | 0 | 2 | 0 | 0 | 0 | 0 | 0 | 20 | 55 | 29 | 93 | 93 | 53 | 67 | 28 | 55 | 40 | 22 | 8 | 25 | 0 | 0 | 1 | 1 | 0 | 0 | 0 | 0 | 0 | 0 | 0 | 0 | 0 | 0 | 0 | 0 | 0 | 0 | 15 | 0 | 0 | 1 | 0 | 0 | 0 | 0 | 0 | 0 | 0 | 0 |
| SH3KBP1 | chrX:19687617-19687691 | 0 | 0 | 0 | 0 | 0 | 0 | 0 | 0 | 36 | 58 | 37 | 34 | 46 | 65 | 69 | 36 | 55 | 43 | 34 | 3 | 30 | 0 | 0 | 3 | 1 | 1 | 0 | 1 | 0 | 1 | 1 | 0 | 0 | 0 | 0 | 5 | 1 | 0 | 0 | 5 | 0 | 0 | 0 | 1 | 0 | 1 | 1 | 0 | 0 | 0 | 0 |
| PHTF1 | chr1:113719876\|113713611-113719929 | 0 | 2 | 1 | 1 | 0 | 0 | 0 | 1 | 5 | 9 | 10 | 30 | 34 | 13 | 11 | 7 | 4 | 11 | 11 | 4 | 3 | -1 | -1 | 1 | 1 | 0 | 0 | 1 | 3 | 1 | 1 | 0 | 0 | 2 | 1 | 2 | 2 | 1 | 1 | 2 | 1 | 1 | 1 | 2 | 2 | 1 | 1 | 0 | 1 | 1 | 3 |
| SGCE | chr7:94587778-94587812 | 0 | 0 | 1 | 0 | 0 | 0 | 0 | 1 | 20 | 39 | 30 | 76 | 73 | 43 | 50 | 25 | 34 | 38 | 32 | 7 | 16 | 1 | 0 | 1 | 1 | 0 | 0 | 0 | 1 | 0 | 0 | 0 | 1 | 0 | 0 | 1 | 1 | 0 | 2 | 12 | 0 | 1 | 1 | 1 | 0 | 1 | 1 | 0 | 0 | 0 | 3 |
| PLEKHA5 | chr12:19290677\|19290512-19320056 | 1 | 1 | 0 | 1 | 1 | 1 | 1 | 1 | 37 | 65 | 55 | 45 | 44 | 72 | 72 | 43 | 32 | 68 | 65 | 8 | 21 | 0 | 0 | 1 | 1 | 1 | 0 | 1 | 1 | 3 | 2 | 1 | 0 | 1 | 1 | 1 | 1 | 0 | 0 | 2 | 1 | 1 | 1 | 0 | 1 | 1 | 6 | 0 | 0 | 1 | 4 |
| SHANK3 | chr22:50711615-50711638\|50714149 | 2 | 2 | 2 | 3 | 2 | 3 | 4 | 2 | 72 | 81 | 63 | 94 | 92 | 79 | 82 | 73 | 42 | 75 | 60 | 15 | 27 | 4 | 4 | 6 | 4 | 3 | 2 | 3 | 2 | 3 | 3 | 1 | 2 | 3 | 2 | 5 | 3 | 3 | 3 | 9 | 2 | 3 | 3 | 2 | 1 | 3 | 10 | 2 | 3 | 3 | 13 |
| ADGRL3 | chr4:62028855-62028881 | 0 | 0 | 0 | 0 | 0 | 0 | 0 | 0 | 29 | 40 | 25 | 51 | 41 | 42 | 50 | 31 | 43 | 34 | 27 | 24 | 29 | 0 | 0 | 2 | 1 | 0 | 0 | 0 | 0 | 2 | 2 | 0 | -1 | 0 | 1 | 0 | 1 | 0 | 1 | 6 | 0 | 0 | 0 | 1 | 0 | 0 | 1 | 0 | 0 | 0 | -1 |
| ASAP1 | chr8:130160785-130160793 | 0 | 0 | 2 | 0 | 0 | 0 | 0 | 0 | 24 | 53 | 23 | 63 | 58 | 50 | 62 | 23 | 35 | 37 | 25 | 7 | 13 | 0 | 0 | 1 | 1 | 0 | 0 | 0 | 0 | 2 | 2 | 0 | 0 | 0 | 0 | 1 | 1 | 0 | 1 | 10 | 0 | 0 | 0 | 1 | 0 | 1 | 0 | 0 | 0 | 0 | 1 |
| RYR1 | chr19:38544286-38546330 | 1 | 3 | 8 | 1 | 1 | 1 | 0 | 1 | 22 | 35 | 22 | 94 | 95 | 47 | 46 | 26 | 76 | 32 | 28 | 27 | 24 | -1 | 0 | 4 | 3 | 2 | 1 | 2 | 0 | 2 | 2 | 3 | 4 | 2 | 2 | 1 | 2 | 1 | 4 | 11 | 1 | 2 | 11 | 2 | 3 | 7 | 1 | 2 | 3 | 1 | 2 |
| TFG | chr3:100747414-100747454 | 0 | 0 | 1 | 0 | 0 | 0 | 0 | 0 | 10 | 19 | 11 | 28 | 29 | 21 | 23 | 12 | 11 | 11 | 11 | 2 | 9 | 0 | 0 | 1 | 0 | 0 | 0 | 0 | 0 | 0 | 0 | 0 | 0 | 0 | 0 | 0 | 0 | 0 | 0 | 5 | 0 | 0 | 0 | 0 | 0 | 0 | 1 | 0 | 0 | 0 | 0 |
| GTF2I | chr7:74714857-74714916\|74715166 | 11 | 8 | 5 | 5 | 5 | 5 | 4 | 9 | 36 | 49 | 33 | 77 | 71 | 47 | 57 | 38 | 52 | 42 | 36 | 31 | 34 | 5 | 2 | 6 | 5 | 5 | 4 | 5 | 7 | 7 | 7 | 6 | 4 | 8 | 6 | 8 | 8 | 4 | 5 | 10 | 5 | 7 | 7 | 6 | 5 | 6 | 10 | 4 | 5 | 4 | 7 |

SFig5, pg2

| Gene Symbol | Exon Location (hg38, human) | Adipose - Subcutaneous | Adipose - Visceral (Omentum) | Adrenal Gland | Artery - Aorta | Artery - Coronary | Artery - Tibial | Bladder | Breast - Mammary Tissue | Brain - Amygdala | Brain - Anterior cingulate cortex (BA24) | Brain - Caudate (basal ganglia) | Brain - Cerebellar Hemisphere | Brain - Cerebellum | Brain - Cortex | Brain - Frontal Cortex (BA9) | Brain - Hippocampus | Brain - Hypothalamus | Brain - Nucleus accumbens (basal ganglia) | Brain - Putamen (basal ganglia) | Brain - Spinal cord (cervical c-1) | Brain - Substantia nigra | Cervix - Ectocervix | Cervix - Endocervix | Colon - Sigmoid | Colon - Transverse | Esophagus - Gastroesophageal Junction | Esophagus - Mucosa | Esophagus - Muscularis | Fallopian Tube | Heart - Atrial Appendage | Heart - Left Ventricle | Kidney - Cortex | Liver | Lung | Minor Salivary Gland | Muscle - Skeletal | Nerve - Tibial | Ovary | Pancreas | Pituitary | Prostate | Skin - Not Sun Exposed (Suprapubic) | Skin - Sun Exposed (Lower leg) | Small Intestine - Terminal Ileum | Spleen | Stomach | Testis | Thyroid | Uterus | Vagina | Whole Blood |
|---|---|---|---|---|---|---|---|---|---|---|---|---|---|---|---|---|---|---|---|---|---|---|---|---|---|---|---|---|---|---|---|---|---|---|---|---|---|---|---|---|---|---|---|---|---|---|---|---|---|---|---|---|
| LZTR1 | chr22:20980747-20980767 | 1 | 0 | 1 | 0 | 1 | 2 | 0 | 1 | 2 | 9 | 1 | 56 | 54 | 19 | 19 | 5 | 5 | 1 | 1 | 6 | 5 | 0 | 0 | 0 | 0 | 1 | 0 | 1 | 0 | 0 | 1 | 0 | 1 | 1 | 1 | 1 | 1 | 0 | 1 | 7 | 0 | 1 | 8 | 0 | 0 | 1 | 1 | 0 | 1 | 1 | 1 |
| DOCK4 | chr7:111784097-111784123 | 0 | 0 | 1 | 0 | 0 | 0 | 0 | 0 | 15 | 40 | 16 | 87 | 81 | 35 | 51 | 35 | 34 | 30 | 16 | 5 | 17 | 0 | 0 | 3 | 1 | 1 | 0 | 1 | 0 | 0 | 0 | 0 | 0 | 0 | 0 | 0 | 0 | 0 | 1 | 14 | 0 | 0 | 1 | 2 | 1 | 3 | 0 | 0 | 0 | 0 | 0 |
| CADPS2 | chr7:122436360-122436380 | 4 | 4 | 1 | 4 | 4 | 4 | 2 | 5 | 8 | 34 | 4 | 55 | 57 | 41 | 47 | 19 | 24 | 5 | 4 | 14 | 16 | 0 | 5 | 4 | 3 | 4 | 2 | 3 | 3 | 5 | 6 | 2 | 2 | 2 | 2 | 8 | 5 | 2 | 1 | 4 | 2 | 4 | 6 | 3 | 3 | 2 | 6 | 2 | 4 | 3 | 4 |
| GYG1 | chr3:149026452-149026502 | 1 | 4 | 6 | 2 | 1 | 1 | 1 | 2 | 69 | 86 | 74 | 63 | 77 | 82 | 70 | 80 | 83 | 80 | 72 | 50 | 67 | 0 | 3 | 3 | 3 | 2 | 2 | 2 | 1 | 3 | 1 | 5 | 6 | 1 | 4 | 1 | 2 | 1 | 1 | 13 | 2 | 1 | 1 | 5 | 2 | 3 | 1 | 1 | 2 | 3 | 1 |
| FMNL1 | chr17:45246505-45246604 | 5 | 4 | 5 | 7 | 4 | 6 | 5 | 5 | 69 | 85 | 67 | 91 | 90 | 90 | 89 | 74 | 63 | 69 | 71 | 25 | 49 | 1 | 2 | 5 | 4 | 3 | 3 | 3 | 5 | 7 | 14 | 4 | 3 | 7 | 3 | 13 | 7 | 4 | 3 | 3 | 3 | 7 | 7 | 3 | 3 | 4 | 8 | 3 | 4 | 3 | 6 |
| GALNT5 | chr2:157266776-157266856 | 2 | 0 | 0 | 0 | 0 | 0 | -1 | 0 | 0 | 0 | 0 | 16 | 20 | -1 | 0 | 0 | 0 | 0 | 0 | -1 | 0 | 0 | 0 | 1 | 0 | 0 | 0 | 1 | -1 | 1 | 0 | 0 | -1 | 0 | 0 | -1 | 0 | 0 | 1 | 0 | 1 | 0 | 0 | 0 | -1 | 0 | 1 | 1 | 0 | 1 | 0 |
| STXBP1 | chr9:127684368-127684493\|127689907 | 5 | 4 | 1 | 4 | 5 | 5 | 6 | 5 | 11 | 17 | 13 | 55 | 52 | 21 | 22 | 11 | 17 | 6 | 13 | 19 | 48 | 5 | 7 | 6 | 6 | 5 | 3 | 5 | 5 | 5 | 3 | 2 | 5 | 3 | 3 | 6 | 4 | 4 | 2 | 5 | 5 | 3 | 5 | 5 | 1 | 5 | 2 | 3 | 5 | 7 | 7 |
| TNNI3K | chr1:74345878-74347756\|74346801 | 1 | 1 | 1 | 1 | 1 | 2 | 1 | 1 | 15 | 43 | 29 | 75 | 76 | 54 | 54 | 19 | 33 | 42 | 35 | 6 | 8 | 0 | 0 | 2 | 2 | 2 | 1 | 2 | 0 | 11 | 5 | 1 | 0 | 1 | 1 | 8 | 1 | 0 | 1 | 7 | 2 | 1 | 1 | 1 | 1 | 3 | 5 | 1 | 1 | 0 | 2 |
| SLC12A2 | chr5:128177574-128177639 | 1 | 3 | 2 | 2 | 3 | 2 | 3 | 2 | 2 | 6 | 3 | 40 | 54 | 8 | 5 | 2 | 3 | 5 | 2 | 1 | 2 | 1 | 3 | 4 | 4 | 3 | 3 | 3 | 3 | 4 | 4 | 1 | 6 | 2 | 2 | 6 | 1 | 3 | 3 | 7 | 4 | 1 | 1 | 3 | 4 | 2 | 8 | 3 | 4 | 4 | 2 |
| ELMO1 | chr7:37217712-37217756 | 0 | 0 | 2 | 0 | 0 | 0 | 0 | 0 | 3 | 8 | 2 | 24 | 32 | 5 | 7 | 2 | 15 | 4 | 2 | 4 | 6 | 0 | 0 | 0 | 0 | 0 | 0 | 0 | 0 | 1 | 1 | 0 | 0 | 0 | 0 | 0 | 1 | 0 | 2 | 1 | 0 | 0 | 0 | 0 | 0 | 0 | 6 | 0 | 0 | 0 | 0 |
| SLC25A25 | chr9:128102080-128102115 | 1 | 2 | 1 | 1 | 1 | 0 | 6 | 1 | 13 | 33 | 13 | 58 | 52 | 36 | 46 | 23 | 39 | 10 | 15 | 10 | 34 | 0 | 0 | 3 | 2 | 1 | 1 | 1 | 1 | 1 | 1 | 2 | 2 | 1 | 1 | 3 | 0 | 0 | 5 | 6 | 2 | 2 | 1 | 3 | 0 | 3 | 6 | 1 | 0 | 0 | 0 |
| COL15A1 | chr9:99043594-99043619 | 0 | 0 | 0 | 0 | 0 | 0 | 0 | 0 | 36 | 61 | 6 | 28 | 51 | 50 | 49 | 19 | 56 | 26 | 7 | 8 | 20 | 0 | 0 | 0 | 0 | 0 | 0 | 0 | 0 | 0 | 0 | 0 | 0 | 0 | 0 | 0 | 0 | 0 | 0 | 0 | 0 | 0 | 0 | 0 | 0 | 0 | 10 | 0 | 0 | 0 | 1 |
| ANK3 | chr10:60082150-60082176 | 0 | 0 | 1 | 0 | 0 | 0 | 0 | 1 | 4 | 16 | 4 | 66 | 68 | 15 | 19 | 4 | 25 | 6 | 3 | 8 | 30 | 0 | 0 | 0 | 0 | 0 | 0 | 0 | 1 | 2 | 2 | 0 | 1 | 0 | 0 | 1 | 0 | 0 | 1 | 12 | 1 | 0 | 0 | 0 | 0 | 1 | 7 | 0 | 1 | 1 | 2 |
| GNAI3 | chr1:109584622-109584713 | 0 | 1 | 0 | 0 | 0 | 0 | 0 | 0 | 6 | 12 | 9 | 23 | 29 | 22 | 18 | 8 | 12 | 11 | 9 | 4 | 5 | 1 | 0 | 1 | 0 | 0 | 0 | 0 | 0 | 2 | 2 | 1 | 1 | 0 | 0 | 2 | 1 | 1 | 1 | 4 | 1 | 0 | 0 | 0 | 1 | 0 | 4 | 1 | 1 | 0 | 0 |
| PTPRU | chr1:29307129-29307146 | 0 | 0 | 0 | 0 | 0 | 0 | 0 | 0 | 9 | 14 | 5 | 26 | 26 | 19 | 14 | 10 | 13 | 8 | 4 | 2 | 8 | 0 | 0 | 0 | 0 | 0 | 0 | 0 | 0 | 2 | 3 | 0 | 0 | 0 | 0 | 4 | 0 | 0 | 0 | 0 | 0 | 0 | 0 | 0 | 0 | 0 | 0 | 0 | 0 | 0 | 1 |
| AGAP1 | chr2:236014805-236014879 | 0 | 0 | 1 | 0 | 0 | 0 | 0 | 0 | 17 | 28 | 11 | 35 | 32 | 24 | 27 | 14 | 16 | 21 | 8 | 3 | 8 | 0 | 0 | 0 | 0 | 0 | 0 | 0 | 0 | 2 | 2 | 0 | 0 | 0 | 0 | 1 | 0 | 0 | 0 | 6 | 0 | 0 | 0 | 0 | 0 | 0 | 0 | 0 | 0 | 0 | 1 |
| CDC14B | chr9:96504098-96504163 | 0 | 0 | 2 | 0 | 1 | 0 | 1 | 0 | 7 | 15 | 7 | 24 | 24 | 17 | 19 | 11 | 12 | 7 | 6 | 5 | 5 | 0 | 0 | 2 | 1 | 0 | 1 | 0 | 0 | 0 | 0 | 0 | 0 | 0 | 0 | 0 | 0 | 0 | 0 | 4 | 0 | 0 | 0 | 0 | 0 | 0 | 1 | 0 | 0 | 0 | 0 |
| ERGIC3 | chr20:35554372-35554386 | 0 | 0 | 1 | 0 | 0 | 0 | 0 | 0 | 23 | 42 | 27 | 75 | 69 | 49 | 54 | 25 | 35 | 36 | 29 | 5 | 15 | 0 | 0 | 1 | 0 | 0 | 0 | 0 | 0 | 0 | 1 | 0 | 0 | 0 | 0 | 3 | 1 | 0 | 1 | 14 | 0 | 0 | 0 | 0 | 0 | 1 | 1 | 0 | 0 | 0 | 0 |
| DNM1L | chr12:32705825-32705863\|32706814\|32706642 | 2 | 1 | 2 | 1 | 0 | 1 | 0 | 2 | 28 | 58 | 16 | 74 | 68 | 56 | 67 | 34 | 62 | 28 | 13 | 9 | 32 | 1 | 0 | 4 | 3 | 1 | 0 | 1 | 1 | 1 | 1 | 1 | 1 | 1 | 1 | 14 | 2 | 1 | 1 | 3 | 1 | 2 | 2 | 4 | 1 | 2 | 2 | 1 | 1 | 1 | 1 |
| SSPO | chr7:149794165-149794255 | 1 | 0 | 0 | 0 | 0 | 0 | 0 | 0 | 10 | 25 | 30 | 55 | 53 | 28 | 30 | 18 | 14 | 36 | 30 | 9 | 5 | 0 | 0 | 0 | 0 | 0 | 0 | 0 | 0 | 5 | 6 | 0 | 0 | 0 | 0 | 7 | 0 | 0 | 0 | 0 | 0 | 0 | 2 | 0 | 0 | 0 | 2 | 0 | 0 | 0 | 0 |
| DOCK9 | chr13:98808640-98808675\|98809128 | 0 | 0 | 1 | 1 | 0 | 1 | 0 | 0 | 56 | 59 | 52 | 69 | 66 | 64 | 59 | 64 | 61 | 54 | 55 | 77 | 68 | 0 | 0 | 1 | 0 | 0 | 0 | 0 | 2 | 2 | 2 | 0 | 0 | 1 | 0 | 8 | 1 | 0 | 0 | 7 | 0 | 0 | 1 | 1 | 0 | 0 | 10 | 0 | 0 | 0 | 2 |
| SRC | chr20:37386460-37393015 | 1 | 1 | 0 | 1 | 0 | 1 | 1 | 1 | 30 | 46 | 34 | 65 | 63 | 57 | 60 | 39 | 41 | 45 | 33 | 12 | 23 | 1 | 0 | 1 | 0 | 1 | 0 | 1 | 1 | 3 | 4 | 1 | 1 | 0 | 0 | 13 | 1 | 0 | 0 | 3 | 0 | 0 | 0 | 0 | 0 | 0 | 0 | 0 | 0 | 0 | 0 |
| NSMAF | chr8:58637234-58637389\|58640102 | 1 | 4 | 1 | 2 | 2 | 2 | 4 | 1 | 12 | 22 | 14 | 34 | 30 | 21 | 25 | 20 | 20 | 16 | 12 | 9 | 13 | 5 | 2 | 3 | 2 | 2 | 2 | 2 | 1 | 3 | 2 | 1 | 4 | 3 | 2 | 5 | 2 | 2 | 1 | 3 | 3 | 1 | 1 | 2 | 1 | 2 | 2 | 2 | 3 | 2 | 2 |
| CD2AP | chr6:47603759-47603832 | 1 | 1 | 1 | 1 | 1 | 1 | 2 | 1 | 8 | 18 | 7 | 54 | 59 | 20 | 22 | 5 | 12 | 19 | 9 | 3 | 3 | 1 | 1 | 2 | 1 | 1 | 1 | 1 | 3 | 6 | 6 | 2 | 1 | 1 | 1 | 6 | 2 | 1 | 1 | 4 | 1 | 1 | 1 | 2 | 1 | 2 | 9 | 1 | 1 | 1 | 0 |
| POC1B | chr12:89460016-89461306\|89460038 | 3 | 4 | 3 | 6 | 5 | 5 | 6 | 4 | 7 | 15 | 8 | 36 | 42 | 17 | 22 | 8 | 7 | 11 | 7 | 4 | 5 | 5 | 2 | 4 | 3 | 3 | 3 | 3 | 4 | 4 | 3 | 2 | 2 | 3 | 3 | 3 | 4 | 4 | 2 | 9 | 4 | 3 | 2 | 3 | 3 | 4 | 4 | 4 | 5 | 4 | 2 |
| MAPK8 | chr10:48424088-48424159 | 5 | 4 | 4 | 4 | 4 | 5 | 6 | 4 | 50 | 75 | 40 | 90 | 87 | 73 | 84 | 59 | 65 | 50 | 38 | 23 | 48 | 6 | 4 | 6 | 5 | 8 | 4 | 8 | 6 | 4 | 4 | 1 | 1 | 5 | 4 | 6 | 5 | 7 | 2 | 18 | 3 | 5 | 5 | 4 | 4 | 4 | 11 | 2 | 5 | 4 | 6 |
| CNNM1 | chr10:99368618-99368680 | 0 | 1 | 0 | 1 | 1 | 2 | -1 | 1 | 18 | 34 | 24 | 46 | 44 | 45 | 49 | 26 | 22 | 15 | 27 | 44 | 42 | 0 | 0 | 1 | 0 | 2 | 0 | 3 | 0 | 8 | -1 | 0 | 0 | 1 | 0 | -1 | -1 | 0 | 0 | 1 | 2 | 0 | 1 | 2 | -1 | 2 | 0 | 2 | 0 | 0 | -1 |
| FBN1 | chr15:48427882-48427943 | 0 | 0 | 0 | 0 | 1 | 1 | 1 | 1 | 71 | 87 | 77 | 66 | 63 | 87 | 88 | 75 | 78 | 81 | 78 | 80 | 86 | 1 | 1 | 1 | 1 | 2 | 1 | 3 | 0 | 0 | 0 | 1 | 0 | 0 | 1 | 1 | 8 | 1 | 0 | 4 | 4 | 0 | 0 | 0 | 0 | 1 | 11 | 0 | 1 | 1 | 1 |
| PLEKHA5 | chr12:19346994\|19347107\|19346972-19347182 | 2 | 2 | 1 | 2 | 1 | 3 | 3 | 3 | 39 | 58 | 33 | 61 | 58 | 61 | 66 | 38 | 43 | 36 | 33 | 24 | 45 | 6 | 2 | 4 | 3 | 1 | 2 | 1 | 2 | 4 | 3 | 2 | 2 | 4 | 4 | 2 | 3 | 3 | 2 | 3 | 2 | 3 | 3 | 4 | 1 | 3 | 13 | 2 | 1 | 3 | 3 |
| ZDHHC20 | chr13:21392959-21392994 | 1 | 1 | 3 | 1 | 1 | 1 | 1 | 3 | 19 | 38 | 19 | 45 | 64 | 32 | 32 | 10 | 21 | 24 | 12 | 2 | 7 | 0 | 0 | 1 | 1 | 0 | 0 | 1 | 0 | 0 | 2 | 2 | 1 | 3 | 1 | 1 | 2 | 1 | 1 | 14 | 0 | 4 | 1 | 2 | 1 | 1 | 0 | 0 | 1 | 0 | 2 |
| MIER1 | chr1:66940186-66940222 | 1 | 2 | 2 | 2 | 1 | 2 | 1 | 1 | 5 | 9 | 7 | 21 | 21 | 9 | 17 | 6 | 10 | 8 | 6 | 4 | 7 | 1 | 0 | 1 | 2 | 2 | 2 | 2 | 2 | 3 | 2 | 2 | 1 | 1 | 1 | 3 | 2 | 2 | 1 | 4 | 1 | 1 | 1 | 2 | 3 | 2 | 3 | 1 | 2 | 1 | 1 |
| MINK1 | chr17:4893286-4893309 | 0 | 1 | 1 | 0 | 0 | 0 | 1 | 0 | 35 | 50 | 42 | 85 | 81 | 51 | 58 | 45 | 33 | 61 | 41 | 3 | 15 | 0 | 0 | 3 | 2 | 1 | 1 | 1 | 0 | 5 | 9 | 1 | 1 | 1 | 1 | 5 | 0 | 0 | 2 | 16 | 1 | 0 | 0 | 2 | 1 | 2 | 4 | 1 | 0 | 1 | 2 |
| KIF1C | chr17:5019849-5019871 | 0 | 0 | 0 | 0 | 0 | 0 | 0 | 0 | 1 | 4 | 1 | 25 | 25 | 5 | 5 | 1 | 2 | 1 | 1 | 0 | 1 | 1 | 0 | 1 | 0 | 0 | 0 | 0 | 0 | 0 | 0 | 0 | 0 | 0 | 0 | 0 | 0 | 0 | 0 | 6 | 0 | 0 | 0 | 0 | 0 | 0 | 1 | 0 | 0 | 0 | 0 |
| MTMR1 | chrX:150714479-150714529\|150714532 | 0 | 0 | 0 | 1 | 0 | 0 | 0 | 0 | 56 | 66 | 60 | 81 | 82 | 70 | 70 | 45 | 66 | 56 | 59 | 25 | 52 | 0 | 0 | 2 | 0 | 0 | 0 | 0 | 1 | 6 | 5 | 0 | 0 | 0 | 0 | 4 | 1 | 0 | 0 | 9 | 1 | 0 | 0 | 0 | 0 | 0 | 16 | 0 | 0 | 0 | 0 |
| CASK | chrX:41622617-41622634 | 0 | 0 | 0 | 0 | 0 | 0 | 0 | 0 | 2 | 5 | 2 | 30 | 27 | 5 | 9 | 2 | 6 | 6 | 3 | 1 | 2 | 0 | 0 | 0 | 0 | 0 | 0 | 0 | 0 | 0 | 0 | 0 | 0 | 0 | 0 | 0 | 0 | 0 | 0 | 6 | 0 | 0 | 0 | 0 | 0 | 0 | 0 | 0 | 0 | 0 | 0 |
| CSPP1 | chr8:67114329-67114370 | 0 | 0 | 0 | 0 | 0 | 0 | 0 | 0 | 3 | 9 | 5 | 39 | 35 | 14 | 16 | 5 | 4 | 5 | 7 | 3 | 10 | 0 | 0 | 0 | 0 | 0 | 0 | 0 | 0 | 0 | 0 | 0 | 1 | 0 | 0 | 0 | 0 | 0 | 1 | 1 | 1 | 0 | 0 | 1 | 1 | 0 | 8 | 0 | 0 | 0 | 1 |
| LCP1 | chr13:46148782-46148808 | 0 | 0 | 1 | 0 | 0 | 0 | 0 | 0 | 1 | 3 | 0 | 40 | 75 | 7 | 4 | 1 | 5 | 0 | 0 | 3 | 6 | 0 | 0 | 1 | 0 | 0 | 0 | 0 | 0 | 0 | 0 | 0 | 0 | 0 | 0 | 0 | 0 | 0 | 0 | 1 | 0 | 0 | 0 | 0 | 0 | 0 | 18 | 0 | 0 | 0 | 0 |
| NF1 | chr17:31203102-31203131 | 0 | 0 | 0 | 0 | 0 | 0 | 0 | 0 | 8 | 28 | 15 | 47 | 43 | 29 | 35 | 12 | 8 | 17 | 12 | 2 | 9 | 0 | 0 | 0 | 0 | 0 | 0 | 0 | 0 | 3 | 5 | 0 | 0 | 0 | 0 | 9 | 0 | 0 | 1 | 1 | 0 | 0 | 0 | 0 | 0 | 0 | 0 | 0 | 0 | 0 | 0 |
| PXK | chr3:58420536-58423494\|58420568 | 3 | 5 | 2 | 3 | 4 | 3 | 4 | 4 | 32 | 46 | 35 | 68 | 68 | 48 | 51 | 32 | 41 | 43 | 35 | 26 | 30 | 2 | 0 | 3 | 3 | 3 | 4 | 3 | 5 | 4 | 5 | 3 | 5 | 6 | 4 | 5 | 5 | 3 | 6 | 8 | 3 | 6 | 4 | 7 | 10 | 3 | 4 | 3 | 2 | 2 | 15 |
| CLASP1 | chr2:121418622-121418729 | 0 | 0 | 1 | 0 | 0 | 0 | 0 | 0 | 12 | 27 | 13 | 46 | 42 | 32 | 39 | 16 | 21 | 17 | 13 | 5 | 19 | 0 | 0 | 2 | 1 | 0 | 0 | 0 | 0 | 0 | 0 | 0 | 2 | 0 | 0 | 0 | 0 | 0 | 1 | 10 | 0 | 0 | 0 | 1 | 0 | 1 | 0 | 0 | 0 | 0 | 1 |
| CSTF2 | chrX:100830808-100830867\|100831136 | 0 | 1 | 2 | 1 | 0 | 0 | 2 | 0 | 29 | 57 | 39 | 89 | 86 | 61 | 67 | 34 | 55 | 52 | 39 | 15 | 28 | 1 | 0 | 2 | 2 | 2 | 7 | 1 | 0 | 7 | 11 | 1 | 3 | 1 | 2 | 13 | 1 | 1 | 2 | 11 | 1 | 1 | 1 | 1 | 1 | 2 | 8 | 0 | 0 | 3 | 0 |
| UHRF2 | chr9:6498611\|6498640-6498925 | 3 | 4 | 2 | 2 | 3 | 1 | 2 | 4 | 13 | 20 | 14 | 48 | 61 | 24 | 14 | 18 | 24 | 21 | 16 | 19 | 15 | 6 | 3 | 3 | 3 | 3 | 2 | 2 | 7 | 4 | 2 | 3 | 1 | 2 | 2 | 12 | 11 | 5 | 1 | 4 | 2 | 2 | 1 | 4 | 5 | 3 | 2 | 2 | 4 | 3 | 4 |
| PTPRB | chr12:70565800-70565904 | 5 | 6 | 6 | 4 | 4 | 4 | 6 | 7 | 5 | 7 | 4 | 42 | 52 | 12 | 7 | 5 | 5 | 8 | 4 | 5 | 3 | 5 | 6 | 5 | 5 | 6 | 5 | 6 | 4 | 5 | 5 | 4 | 7 | 4 | 5 | 5 | 6 | 4 | 4 | 8 | 7 | 6 | 6 | 5 | 8 | 5 | 15 | 5 | 5 | 6 | 5 |
| ADAM17 | chr2:9543696-9543760 | 1 | 1 | 1 | 1 | 0 | 1 | 1 | 1 | 4 | 8 | 4 | 30 | 28 | 10 | 7 | 4 | 6 | 4 | 3 | 1 | 3 | 0 | 0 | 1 | 3 | 1 | 1 | 1 | 2 | 0 | 1 | 0 | 1 | 1 | 1 | 1 | 1 | 1 | 2 | 3 | 1 | 2 | 2 | 4 | 2 | 3 | 4 | 1 | 1 | 1 | 2 |
| KCTD3 | chr1:215618088-215618154 | 0 | 1 | 0 | 1 | 1 | 1 | 1 | 1 | 2 | 4 | 3 | 22 | 24 | 8 | 8 | 3 | 3 | 3 | 3 | 1 | 2 | 1 | 0 | 1 | 1 | 0 | 0 | 0 | 1 | 1 | 3 | 1 | 1 | 0 | 0 | 4 | 1 | 1 | 1 | 2 | 1 | 1 | 0 | 1 | 0 | 1 | 5 | 1 | 1 | 0 | 1 |
| KIF1B | chr1:10273014-10273031 | 0 | 0 | 1 | 0 | 0 | 0 | 0 | 0 | 38 | 67 | 29 | 90 | 86 | 62 | 71 | 35 | 37 | 48 | 28 | 7 | 16 | 0 | 0 | 2 | 1 | 0 | 0 | 0 | 0 | 0 | 0 | 0 | 0 | 0 | 0 | 0 | 0 | 0 | 0 | 21 | 0 | 0 | 1 | 1 | 0 | 1 | 0 | 0 | 0 | 1 | 0 |
| ADGRL2 | chr1:81980795-81980839 | 0 | 0 | 0 | 0 | 0 | 0 | 0 | 0 | 11 | 21 | 13 | 16 | 19 | 23 | 23 | 12 | 13 | 20 | 18 | 4 | 8 | 0 | 1 | 0 | 0 | 0 | 0 | 1 | 0 | 1 | 1 | 0 | 0 | 0 | 1 | 1 | 0 | 0 | 1 | 1 | 0 | 0 | 0 | 0 | 0 | 0 | 4 | 0 | 0 | 0 | 2 |
| CPEB3 | chr10:92158049-92158072 | 0 | 0 | 1 | 0 | 0 | 0 | 0 | 0 | 10 | 32 | 13 | 48 | 46 | 30 | 39 | 15 | 23 | 16 | 10 | 8 | 21 | 0 | 0 | 1 | 1 | 0 | 0 | 0 | 0 | 0 | 0 | 0 | 0 | 0 | 0 | 0 | 0 | 0 | 0 | 11 | 0 | 0 | 0 | 0 | 0 | 1 | 0 | 0 | 0 | 0 | 2 |
| IFFO2 | chr1:18917797-18917859 | 1 | 1 | 0 | 2 | 2 | 2 | 3 | 1 | 7 | 19 | 5 | 38 | 38 | 26 | 31 | 10 | 7 | 10 | 4 | 2 | 2 | 0 | 1 | 5 | 4 | 3 | 0 | 2 | 2 | 0 | 1 | 0 | 0 | 1 | 1 | 7 | 1 | 1 | 0 | 0 | 2 | 0 | 0 | 1 | 0 | 2 | 0 | 0 | 2 | 0 | 0 |
| DOCK10 | chr2:224768872-224768947 | 4 | 2 | 2 | 4 | 2 | 6 | 2 | 3 | 72 | 79 | 70 | 81 | 82 | 85 | 83 | 73 | 75 | 57 | 77 | 85 | 83 | 2 | 2 | 12 | 4 | 2 | 1 | 3 | 1 | 1 | 2 | 0 | 2 | 1 | 1 | 9 | 9 | 1 | 1 | 14 | 0 | 3 | 4 | 1 | 1 | 2 | 9 | 2 | 1 | 1 | 0 |
| VPS13C | chr15:62023780-62028420 | 0 | 1 | 5 | 0 | 1 | 0 | 2 | 0 | 18 | 47 | 29 | 92 | 90 | 45 | 56 | 18 | 39 | 38 | 28 | 4 | 12 | 0 | 0 | 1 | 2 | 0 | 0 | 1 | 0 | 0 | 0 | 1 | 0 | 0 | 0 | 1 | 0 | 0 | 1 | 23 | 1 | 1 | 1 | 1 | 0 | 1 | 1 | 0 | 0 | 1 | 0 |
| CLUAP1 | chr16:3533099-3533155 | 0 | 0 | 0 | 0 | 0 | 0 | 0 | 0 | 18 | 39 | 19 | 59 | 53 | 41 | 48 | 23 | 22 | 24 | 22 | 9 | 13 | 0 | 0 | 0 | 0 | 0 | 0 | 0 | 0 | 5 | 9 | 0 | 0 | 0 | 0 | 10 | 2 | 0 | 0 | 1 | 0 | 0 | 0 | 0 | 0 | 0 | 2 | 0 | 0 | 0 | 0 |
| GANC | chr15:42344428-42344911\|42344522 | 1 | 2 | 4 | 1 | 2 | 1 | 1 | 1 | 18 | 20 | 21 | 55 | 63 | 29 | 20 | 24 | 22 | 21 | 22 | 17 | 18 | 1 | 0 | 3 | 2 | 2 | 2 | 2 | 1 | 4 | 4 | 2 | 13 | 2 | 2 | 3 | 2 | 1 | 5 | 6 | 3 | 1 | 1 | 2 | 1 | 3 | 11 | 1 | 2 | 2 | 1 |
| ARVCF | chr22:19976706-19976723\|19976876 | 3 | 1 | 12 | 1 | 1 | 2 | 3 | 2 | 23 | 36 | 21 | 88 | 86 | 47 | 48 | 24 | 47 | 28 | 22 | 27 | 28 | 1 | 0 | 8 | 9 | 2 | 1 | 2 | 2 | 2 | 1 | 2 | 1 | 2 | 2 | 3 | 3 | 1 | 8 | 17 | 2 | 3 | 6 | 8 | 2 | 7 | 4 | 2 | 1 | 4 | 4 |
| SLC43A2 | chr17:1586949-1586960 | 0 | 0 | 3 | 0 | 0 | 0 | 0 | 0 | 25 | 50 | 17 | 89 | 83 | 56 | 64 | 35 | 51 | 25 | 17 | 9 | 27 | 0 | 0 | 2 | 1 | 0 | 0 | 1 | 0 | 1 | 1 | 0 | 0 | 0 | 0 | 0 | 0 | 0 | 2 | 21 | 0 | 0 | 1 | 1 | 0 | 1 | 1 | 0 | 0 | 1 | 0 |
| SNRPC | chr6:34772951-34773007 | 0 | 2 | 2 | 1 | 1 | 1 | 2 | 1 | 16 | 18 | 16 | 9 | 18 | 17 | 7 | 14 | 9 | 22 | 13 | 2 | 4 | 2 | 1 | 1 | 1 | 1 | 2 | 1 | 0 | 1 | 0 | 2 | 3 | 1 | 1 | 1 | 1 | 1 | 1 | 1 | 1 | 1 | 0 | 2 | 4 | 1 | 0 | 0 | 1 | 1 | 2 |
| RAB3GAP1 | chr2:135167693-135167713 | 0 | 0 | 0 | 0 | 0 | 0 | 0 | 0 | 16 | 40 | 21 | 66 | 61 | 37 | 52 | 20 | 43 | 26 | 20 | 6 | 26 | 0 | 0 | 0 | 0 | 0 | 0 | 0 | 0 | 1 | 2 | 0 | 0 | 0 | 0 | 16 | 0 | 0 | 0 | 1 | 0 | 0 | 0 | 0 | 0 | 0 | 0 | 0 | 0 | 0 | 0 |
| FGFR4 | chr5:177089858-177089940 | 1 | 1 | 0 | 1 | 2 | 3 | 1 | 1 | 6 | 5 | 3 | 45 | 44 | 8 | 4 | 6 | 11 | 10 | 2 | 5 | 0 | -1 | 0 | 1 | 0 | 1 | 0 | 1 | -1 | 2 | 1 | 1 | 0 | 1 | 1 | 10 | 2 | 1 | 1 | 2 | 1 | 1 | 3 | 1 | 0 | 0 | 5 | 1 | 2 | 2 | 5 |
| ARVCF | chr22:19976706\|19976568-19976723 | 5 | 3 | 14 | 4 | 3 | 3 | 5 | 5 | 25 | 37 | 21 | 88 | 86 | 48 | 48 | 26 | 48 | 29 | 24 | 27 | 28 | 3 | 2 | 10 | 11 | 3 | 4 | 4 | 3 | 4 | 3 | 5 | 4 | 5 | 5 | 4 | 5 | 2 | 11 | 19 | 4 | 6 | 8 | 9 | 6 | 9 | 9 | 5 | 3 | 6 | 7 |
| WNK1 | chr12:901561-906343 | 0 | 0 | 0 | 0 | 0 | 0 | 0 | 0 | 3 | 10 | 5 | 38 | 39 | 13 | 16 | 5 | 10 | 8 | 5 | 4 | 5 | 0 | 0 | 1 | 1 | 0 | 0 | 0 | 0 | 3 | 2 | 0 | 0 | 0 | 0 | 11 | 0 | 0 | 0 | 1 | 0 | 0 | 0 | 0 | 0 | 0 | 0 | 0 | 0 | 0 | 0 |
| GRAMD1B | chr11:123618345-123618356 | 0 | 0 | 0 | 1 | 0 | 2 | 0 | 0 | 29 | 53 | 34 | 83 | 81 | 64 | 67 | 45 | 46 | 43 | 32 | 21 | 30 | 0 | 0 | 4 | 2 | 2 | 1 | 4 | 0 | 0 | 0 | 0 | 1 | 1 | 1 | 1 | 0 | 0 | 1 | 5 | 2 | 1 | 22 | 0 | 0 | 0 | 0 | 0 | 0 | 0 | 0 |
| C2CD5 | chr12:22502739\|22497579\|22494445-22502804 | 2 | 1 | 1 | 0 | 1 | 0 | 0 | 1 | 11 | 38 | 18 | 24 | 25 | 44 | 56 | 20 | 16 | 18 | 16 | 4 | 13 | 0 | 0 | 1 | 1 | 0 | 0 | 1 | 0 | 2 | 3 | 1 | 1 | 1 | 0 | 6 | 3 | 0 | 0 | 1 | 0 | 1 | 1 | 1 | 1 | 1 | 1 | 0 | 0 | 0 | 1 |
| SHANK2 | chr11:70500570-70501931 | 0 | 0 | 4 | 0 | -1 | -1 | 0 | 0 | 34 | 37 | 29 | 53 | 61 | 41 | 32 | 33 | 38 | 37 | 39 | -1 | 35 | -1 | 0 | 2 | 1 | 0 | 0 | 0 | 0 | -1 | 0 | 0 | 0 | 0 | 0 | 0 | 0 | 0 | 0 | 16 | 0 | 0 | 0 | 3 | 0 | 0 | 0 | 0 | 0 | 0 | -1 |
| NPHP3-ACAD11 | chr3:132621132-132621292 | 1 | 1 | 1 | 1 | 1 | 1 | 1 | 1 | 12 | 17 | 12 | 45 | 44 | 20 | 19 | 17 | 22 | 12 | 13 | 25 | 21 | 1 | 2 | 1 | 1 | 1 | 1 | 1 | 0 | 9 | 8 | 1 | 1 | 1 | 1 | 7 | 2 | 1 | 1 | 3 | 1 | 1 | 1 | 1 | 1 | 1 | 1 | 1 | 1 | 1 | 2 |
| NPTN | chr15:73597022-73597369 | 0 | 0 | 1 | 0 | 0 | 0 | 1 | 0 | 21 | 23 | 32 | 24 | 25 | 20 | 22 | 23 | 12 | 40 | 32 | 5 | 9 | 0 | 0 | 1 | 1 | 1 | 0 | 1 | 0 | 1 | 2 | 1 | 1 | 1 | 0 | 4 | 0 | 0 | 0 | 5 | 1 | 1 | 1 | 1 | 1 | 1 | 2 | 1 | 0 | 0 | 3 |
| AKAP9 | chr7:92043282-92043377 | 3 | 3 | 3 | 4 | 3 | 3 | 1 | 4 | 3 | 2 | 4 | 63 | 64 | 7 | 4 | 2 | 1 | 1 | 1 | 2 | 3 | 2 | -1 | 3 | 2 | 4 | 1 | 4 | 5 | 5 | 1 | 1 | 0 | 3 | 1 | 1 | 2 | 6 | 1 | 1 | 3 | 2 | 2 | 1 | 2 | 6 | 17 | 3 | 2 | 2 | 2 |
| SNX2 | chr5:122826638-122826649 | 0 | 0 | 1 | 0 | 0 | 0 | 0 | 0 | 22 | 45 | 24 | 75 | 77 | 49 | 56 | 28 | 34 | 36 | 24 | 4 | 16 | 0 | 0 | 1 | 0 | 0 | 0 | 0 | 0 | 0 | 0 | 0 | 0 | 0 | 0 | 0 | 0 | 0 | 1 | 22 | 0 | 0 | 0 | 0 | 0 | 1 | 0 | 0 | 0 | 0 | 0 |
| KLHDC4 | chr16:87765066-87765340 | 2 | 3 | 1 | 3 | 3 | 2 | 3 | 3 | 3 | 3 | 2 | 39 | 40 | 4 | 3 | 4 | 2 | 2 | 3 | 3 | 2 | 5 | 3 | 5 | 3 | 3 | 2 | 4 | 3 | 3 | 2 | 2 | 4 | 2 | 3 | 11 | 3 | 3 | 2 | 3 | 3 | 3 | 3 | 2 | 2 | 2 | 6 | 2 | 6 | 5 | 2 |
| P3H3 | chr12:6830870-6831033 | 0 | 0 | 1 | 0 | 1 | 0 | 1 | 0 | 82 | 92 | 87 | 99 | 99 | 94 | 93 | 86 | 92 | 94 | 86 | 71 | 80 | 0 | 1 | 2 | 1 | 1 | 1 | 2 | 0 | 16 | 23 | 0 | 0 | 0 | 0 | 2 | 7 | 1 | 0 | 7 | 1 | 1 | 1 | 1 | 1 | 1 | 8 | 1 | 0 | 0 | 3 |
| POLE | chr12:132658692-132658743 | 0 | 0 | 0 | 2 | 1 | 1 | 4 | 0 | 7 | 22 | 8 | 46 | 47 | 32 | 22 | 10 | 6 | 16 | 6 | 7 | 7 | 2 | 1 | 6 | 2 | 4 | 0 | 4 | 1 | 2 | 3 | 0 | 0 | 0 | 0 | 11 | 0 | 0 | 0 | 0 | 1 | 0 | 0 | 1 | 0 | 1 | 0 | 0 | 1 | 1 | 0 |
| ENPP2 | chr8:119572176-119572250 | 2 | 3 | 2 | 2 | 3 | 2 | 3 | 2 | 88 | 85 | 87 | 92 | 84 | 87 | 86 | 90 | 86 | 80 | 90 | 96 | 94 | 3 | 2 | 5 | 4 | 5 | 4 | 6 | 5 | 5 | 7 | 3 | 4 | 5 | 3 | 4 | 11 | 3 | 4 | 16 | 3 | 2 | 2 | 4 | 15 | 3 | 15 | 5 | 2 | 3 | 12 |
| ASCC3 | chr6:100770432-100770473 | 2 | 2 | 1 | 2 | 1 | 2 | 0 | 2 | 6 | 3 | 6 | 22 | 23 | 9 | 3 | 1 | 2 | 3 | 2 | 2 | 1 | 2 | -1 | 1 | 1 | 1 | 2 | 1 | -1 | 2 | 2 | 1 | 3 | 2 | 2 | 1 | 1 | 3 | 3 | 4 | 2 | 2 | 2 | 3 | 4 | 2 | 2 | 2 | 1 | 2 | 2 |
| ADGRL3 | chr4:62068166\|62063505-62068217 | 0 | 1 | 1 | 0 | 0 | 0 | 0 | 1 | 10 | 27 | 12 | 46 | 36 | 33 | 39 | 13 | 24 | 14 | 12 | 5 | 11 | 1 | 0 | 7 | 4 | 2 | 0 | 2 | 0 | 0 | 1 | 1 | -1 | 0 | 1 | 0 | 0 | 0 | 0 | 6 | 0 | 1 | 1 | 3 | 0 | 1 | 4 | 1 | 1 | 1 | -1 |
| FRY | chr13:32287867-32287884 | 0 | 0 | 2 | 0 | 0 | 0 | 0 | 0 | 15 | 34 | 21 | 69 | 71 | 37 | 39 | 17 | 25 | 37 | 21 | 7 | 12 | 0 | 0 | 0 | 0 | 0 | 0 | 1 | 0 | 12 | 15 | 0 | 0 | 0 | 1 | 9 | 0 | 0 | 0 | 2 | 0 | 0 | 0 | 1 | 0 | 0 | 7 | 0 | 0 | 0 | 1 |
| LMAN2L | chr2:96713107-96713139 | 7 | 8 | 5 | 5 | 6 | 5 | 8 | 7 | 22 | 44 | 27 | 86 | 84 | 46 | 54 | 29 | 68 | 41 | 25 | 8 | 21 | 5 | 2 | 14 | 10 | 13 | 4 | 14 | 7 | 5 | 6 | 2 | 4 | 6 | 2 | 9 | 4 | 1 | 2 | 18 | 4 | 2 | 2 | 9 | 5 | 8 | 1 | 1 | 6 | 5 | 7 |
| SPIRE1 | chr18:12485959-12486000 | 2 | 2 | 2 | 1 | 1 | 1 | 0 | 2 | 11 | 21 | 12 | 48 | 42 | 22 | 25 | 12 | 25 | 15 | 8 | 8 | 10 | 0 | 0 | 1 | 2 | 1 | 2 | 1 | 0 | 8 | 6 | 1 | 5 | 2 | 2 | 6 | 2 | 0 | 5 | 6 | 1 | 2 | 3 | 2 | 0 | 1 | 2 | 2 | 1 | 1 | 5 |
| DNAH10 | chr12:123932618-123932731 | 0 | 1 | 1 | 0 | 0 | 2 | -1 | 1 | 3 | 19 | 6 | 35 | 31 | 24 | 21 | 4 | 10 | 12 | 9 | 3 | 6 | 0 | 0 | 7 | 2 | 0 | 0 | 0 | 0 | 0 | 0 | -1 | 0 | 0 | 0 | 0 | 0 | -1 | 2 | 5 | 0 | 1 | 2 | 0 | 0 | 0 | 3 | 1 | 0 | 0 | 0 |
| MAGI1 | chr3:65387143-65387226 | 14 | 13 | 2 | 4 | 5 | 5 | 5 | 12 | 52 | 80 | 47 | 94 | 94 | 78 | 87 | 50 | 73 | 58 | 43 | 15 | 38 | 7 | -1 | 8 | 7 | 6 | 3 | 9 | 16 | 8 | 6 | 3 | 3 | 7 | 3 | 10 | 12 | 10 | 2 | 17 | 3 | 5 | 3 | 8 | 3 | 3 | 11 | 2 | 5 | 4 | 15 |
| FAM92A1 | chr8:93702520-93702543 | 0 | 0 | 2 | 0 | 0 | 0 | 0 | 0 | 37 | 56 | 35 | 51 | 54 | 57 | 57 | 45 | 39 | 35 | 38 | 19 | 37 | 0 | 0 | 3 | 2 | 1 | 0 | 0 | 0 | 0 | 0 | 0 | 0 | 0 | 0 | 0 | 0 | 0 | 1 | 17 | 0 | 0 | 1 | 1 | 0 | 1 | 0 | 0 | 0 | 1 | 3 |
| SCN8A | chr12:51780649-51780771 | 2 | 0 | 0 | 4 | -1 | -1 | -1 | 1 | 60 | 85 | 78 | 71 | 66 | 89 | 92 | 80 | 48 | 78 | 82 | 49 | 72 | 0 | 0 | 4 | 1 | -1 | 1 | 2 | -1 | -1 | 5 | 0 | 1 | 7 | 0 | -1 | 2 | 1 | 0 | 1 | 0 | 3 | 17 | 0 | 1 | 5 | 0 | 0 | 0 | -1 | 4 |
| TNFSF13B | chr13:108302776-108303016 | 1 | 2 | 1 | 1 | 2 | 1 | 0 | 2 | 6 | 6 | 4 | 15 | 24 | 9 | 8 | 6 | 5 | 5 | 4 | 4 | 5 | 0 | 0 | 2 | 2 | 2 | 2 | 2 | -1 | 2 | 3 | 3 | 3 | 2 | 1 | 3 | 2 | 3 | 2 | 4 | 2 | 1 | 1 | 2 | 3 | 2 | 5 | 1 | 4 | 2 | 1 |
| PACS2 | chr14:105385685-105385717\|105388036 | 1 | 1 | 1 | 9 | 7 | 9 | 3 | 1 | 28 | 46 | 27 | 75 | 73 | 48 | 51 | 26 | 22 | 37 | 22 | 2 | 8 | 8 | 5 | 8 | 5 | 6 | 2 | 4 | 6 | 1 | 1 | 1 | 1 | 2 | 1 | 7 | 1 | 2 | 0 | 18 | 4 | 1 | 2 | 4 | 0 | 4 | 0 | 1 | 8 | 5 | 1 |
| PTPRM | chr18:8371760-8371810 | 7 | 8 | 6 | 6 | 6 | 5 | 5 | 8 | 14 | 26 | 18 | 25 | 28 | 26 | 29 | 18 | 24 | 20 | 20 | 11 | 20 | 6 | 4 | 5 | 7 | 6 | 6 | 6 | 7 | 8 | 8 | 6 | 7 | 9 | 7 | 10 | 7 | 5 | 7 | 7 | 6 | 7 | 7 | 6 | 6 | 6 | 8 | 7 | 7 | 6 | 6 |
| SLC39A8 | chr4:102285949-102285973 | 0 | 0 | 0 | 0 | 0 | 0 | 0 | 0 | 1 | 5 | 8 | 45 | 35 | 12 | 17 | 2 | 3 | 7 | 8 | 1 | 2 | 0 | 0 | 0 | 0 | 0 | 0 | 0 | 0 | 0 | 0 | 0 | 0 | 0 | 0 | 0 | 0 | 0 | 0 | 11 | 0 | 0 | 0 | 0 | 0 | 0 | 0 | 0 | 0 | 0 | 0 |
| TNRC6B | chr22:40281119-40281289 | 1 | 0 | 0 | 0 | 0 | 1 | 1 | 1 | 17 | 22 | 15 | 29 | 28 | 21 | 28 | 17 | 32 | 22 | 14 | 2 | 7 | -1 | 0 | 1 | 2 | 0 | 1 | 1 | 0 | 1 | 1 | 1 | 1 | 1 | 2 | 3 | 0 | 0 | 1 | 4 | 1 | 1 | 1 | 3 | 1 | 1 | 7 | 0 | 0 | 0 | 1 |
| PTPRM | chr18:8122242-8122316 | 5 | 5 | 5 | 2 | 3 | 3 | 3 | 5 | 21 | 39 | 28 | 37 | 46 | 35 | 40 | 20 | 42 | 40 | 29 | 9 | 22 | 5 | 3 | 4 | 4 | 4 | 4 | 4 | 6 | 4 | 4 | 4 | 3 | 5 | 5 | 3 | 5 | 4 | 5 | 12 | 4 | 4 | 5 | 4 | 4 | 5 | 14 | 4 | 4 | 4 | 5 |
| CAMK2D | chr4:113547644-113547716 | 1 | 1 | 1 | 0 | 1 | 0 | 0 | 1 | 45 | 42 | 10 | 29 | 31 | 31 | 37 | 26 | 25 | 35 | 9 | 8 | 6 | 2 | 4 | 1 | 2 | 1 | 1 | 0 | 0 | 0 | 0 | 1 | 4 | 1 | 2 | 0 | 1 | 1 | 3 | 3 | 2 | 1 | 1 | 3 | 6 | 1 | 2 | 2 | 1 | 1 | 7 |
| CAMTA1 | chr1:7749771-7749791 | 0 | 0 | 7 | 0 | 0 | 0 | 0 | 0 | 11 | 20 | 13 | 33 | 32 | 19 | 19 | 13 | 23 | 12 | 15 | 7 | 12 | 0 | 0 | 2 | 1 | 1 | 0 | 1 | 0 | 0 | 0 | 0 | 0 | 0 | 0 | 0 | 0 | 0 | 1 | 6 | 0 | 0 | 0 | 1 | 0 | 1 | 0 | 0 | 0 | 1 | 5 |
| BPTF | chr17:67928959-67929120 | 15 | 17 | 12 | 11 | 11 | 13 | 14 | 15 | 15 | 19 | 17 | 68 | 70 | 20 | 18 | 13 | 23 | 17 | 14 | 11 | 12 | 14 | 11 | 16 | 13 | 16 | 11 | 16 | 12 | 12 | 9 | 15 | 12 | 10 | 13 | 5 | 8 | 17 | 10 | 19 | 13 | 11 | 13 | 10 | 10 | 12 | 20 | 16 | 19 | 14 | 14 |
| DNAJB5 | chr9:34990671\|34990499-34990812 | 4 | 8 | 6 | 6 | 5 | 4 | 3 | 5 | 30 | 32 | 28 | 46 | 49 | 33 | 29 | 32 | 36 | 29 | 26 | 19 | 22 | 4 | 9 | 6 | 6 | 5 | 6 | 5 | 9 | 4 | 3 | 7 | 8 | 10 | 4 | 2 | 4 | 6 | 6 | 15 | 7 | 4 | 3 | 7 | 6 | 6 | 7 | 8 | 7 | 6 | 5 |
| C2CD5 | chr12:22519115-22519239 | 0 | 0 | 1 | 0 | 0 | 0 | 1 | 1 | 23 | 58 | 27 | 78 | 72 | 56 | 72 | 29 | 45 | 30 | 26 | 12 | 30 | 2 | 0 | 1 | 1 | 1 | 1 | 1 | 2 | 1 | 1 | 1 | 0 | 1 | 1 | 10 | 0 | 1 | 1 | 8 | 2 | 2 | 3 | 1 | 1 | 2 | 22 | 2 | 0 | 1 | 1 |
| ZNF382 | chr19:36609468-36609501 | 1 | 2 | 0 | 1 | 0 | 1 | 0 | 1 | 19 | 21 | 15 | 44 | 32 | 23 | 27 | 10 | 8 | 19 | 14 | 3 | 5 | -1 | 0 | 1 | 3 | 0 | 0 | 1 | 0 | 1 | 1 | 0 | -1 | 1 | 0 | 1 | 0 | 1 | 4 | 2 | 1 | 6 | 4 | 1 | 1 | 1 | 7 | 1 | 1 | 2 | 0 |
| MADD | chr11:47292543-47292596 | 1 | 1 | 2 | 1 | 1 | 0 | 2 | 1 | 48 | 73 | 49 | 72 | 74 | 77 | 79 | 56 | 44 | 54 | 47 | 20 | 43 | 0 | 2 | 3 | 3 | 1 | 2 | 2 | 3 | 13 | 20 | 1 | 2 | 1 | 2 | 7 | 1 | 1 | 2 | 6 | 1 | 1 | 1 | 2 | 2 | 2 | 10 | 1 | 1 | 2 | 2 |
| ATP6V0A2 | chr12:123742029-123742093 | 0 | 1 | 0 | 0 | 0 | 0 | 1 | 0 | 3 | 3 | 2 | 19 | 21 | 7 | 6 | 2 | 3 | 4 | 3 | 4 | 1 | 0 | 0 | 0 | 0 | 0 | 0 | 0 | 0 | 2 | 3 | 0 | 0 | 0 | 0 | 6 | 0 | 0 | 0 | 0 | 1 | 0 | 0 | 0 | 0 | 0 | 2 | 0 | 0 | 0 | 0 |
| RELN | chr7:103473803-103478524\|103482270\|103473912\|103478394 | 1 | 1 | 3 | 1 | 0 | 1 | 3 | 1 | 9 | 17 | 12 | 36 | 39 | 16 | 15 | 5 | 10 | 16 | 10 | 5 | 5 | 0 | 0 | 2 | 0 | 1 | 1 | 1 | 1 | 0 | 2 | 1 | 1 | 1 | 1 | 2 | 0 | 1 | 1 | 2 | 1 | 1 | 2 | 1 | 1 | 1 | 13 | 1 | 1 | 1 | 4 |
| ENO4 | chr10:116854086-116854199 | 0 | 0 | 0 | 4 | 1 | 1 | 0 | 1 | 3 | 9 | 5 | 19 | 25 | 8 | 13 | 5 | 6 | 0 | 3 | 23 | 15 | 0 | 0 | 3 | 1 | 0 | -1 | 1 | 0 | 3 | -1 | 0 | 0 | 0 | 0 | 0 | 2 | 0 | 0 | 3 | -1 | 0 | 2 | -1 | 0 | 0 | 5 | 1 | 0 | 0 | 0 |
| CCDC149 | chr4:24821055-24821087 | 0 | 0 | 3 | 0 | 0 | 0 | 0 | 0 | 38 | 64 | 32 | 83 | 79 | 66 | 71 | 44 | 44 | 49 | 27 | 8 | 32 | 0 | 0 | 1 | 1 | 0 | 0 | 0 | 0 | 0 | 0 | 0 | 0 | 0 | 0 | 0 | 0 | 0 | 1 | 27 | 0 | 0 | 0 | 1 | 0 | 1 | 0 | 0 | 0 | 0 | 0 |
| USP40 | chr2:233491032\|233489662-233491071 | 6 | 5 | 4 | 10 | 7 | 9 | 5 | 7 | 16 | 25 | 17 | 73 | 71 | 35 | 39 | 23 | 31 | 22 | 16 | 18 | 22 | 6 | 8 | 9 | 6 | 6 | 6 | 6 | 7 | 6 | 7 | 5 | 5 | 6 | 5 | 25 | 7 | 5 | 5 | 7 | 5 | 13 | 13 | 8 | 5 | 6 | 4 | 6 | 8 | 8 | 9 |
| GABBR1 | chr6:29609229-29609379 | 3 | 2 | 6 | 6 | 5 | 6 | 7 | 2 | 94 | 96 | 94 | 99 | 97 | 94 | 97 | 94 | 95 | 96 | 95 | 88 | 93 | 4 | 6 | 9 | 8 | 7 | 3 | 9 | 3 | 24 | 23 | 3 | 2 | 2 | 2 | 5 | 6 | 3 | 2 | 17 | 5 | 2 | 4 | 5 | 1 | 6 | 7 | 3 | 4 | 4 | 2 |
| MEF2C | chr5:88730211-88730234 | 0 | 0 | 0 | 0 | 0 | 0 | 0 | 0 | 57 | 89 | 28 | 64 | 70 | 88 | 94 | 58 | 38 | 36 | 17 | 2 | 23 | 0 | 0 | 0 | 0 | 0 | 0 | 0 | 0 | 0 | 0 | 1 | 0 | 0 | 0 | 0 | 0 | 0 | 0 | 24 | 0 | 0 | 0 | 0 | 0 | 0 | 0 | 0 | 0 | 0 | 0 |
| SH3GLB2 | chr9:129009453-129009467 | 1 | 1 | 2 | 1 | 1 | 1 | 1 | 1 | 45 | 70 | 35 | 82 | 81 | 68 | 76 | 36 | 39 | 47 | 32 | 4 | 16 | 1 | 0 | 2 | 1 | 1 | 1 | 1 | 1 | 1 | 1 | 0 | 1 | 1 | 1 | 0 | 1 | 1 | 1 | 28 | 1 | 1 | 1 | 1 | 1 | 1 | 8 | 0 | 0 | 1 | 1 |
| C9orf3 | chr9:95080972-95082235 | 1 | 2 | 6 | 1 | 1 | 0 | 1 | 2 | 3 | 4 | 3 | 23 | 30 | 4 | 4 | 4 | 5 | 4 | 3 | 2 | 2 | 2 | 2 | 1 | 1 | 1 | 2 | 1 | 2 | 1 | 1 | 5 | 7 | 2 | 2 | 0 | 1 | 2 | 4 | 5 | 2 | 1 | 1 | 2 | 3 | 2 | 2 | 5 | 2 | 2 | 2 |
| LRCH2 | chrX:115123558-115123756 | 1 | 0 | 0 | 0 | 1 | 1 | 0 | 1 | 3 | 5 | 3 | 18 | 21 | 7 | 7 | 9 | 4 | 4 | 5 | 12 | 7 | 0 | 0 | 0 | 1 | 0 | 0 | 0 | 0 | 4 | 0 | 0 | 5 | 1 | 1 | 1 | 3 | 0 | 0 | 1 | 0 | 1 | 2 | 0 | 0 | 0 | 2 | 0 | 0 | 0 | -1 |
| KIAA0586 | chr14:58495483-58495614 | 4 | 3 | 2 | 3 | 3 | 2 | 2 | 4 | 3 | 4 | 3 | 22 | 24 | 4 | 4 | 2 | 4 | 3 | 3 | 6 | 4 | 5 | 4 | 3 | 2 | 2 | 3 | 2 | 5 | 3 | 2 | 3 | 5 | 3 | 3 | 4 | 4 | 3 | 4 | 1 | 3 | 4 | 5 | 3 | 3 | 4 | 8 | 4 | 3 | 4 | 2 |
| DYNC1I2 | chr2:171706547-171706564 | 0 | 0 | 0 | 0 | 0 | 0 | 2 | 0 | 5 | 13 | 12 | 9 | 12 | 15 | 23 | 5 | 3 | 16 | 8 | 1 | 2 | 0 | 0 | 2 | 1 | 2 | 0 | 2 | 0 | 1 | 1 | 0 | 0 | 0 | 0 | 2 | 0 | 0 | 0 | 1 | 1 | 0 | 0 | 0 | 0 | 1 | 1 | 0 | 1 | 0 | 0 |
| PUS7L | chr12:43739228\|43739632-43739776 | 3 | 4 | 3 | 4 | 4 | 3 | 1 | 5 | 8 | 14 | 12 | 36 | 46 | 20 | 15 | 13 | 22 | 15 | 13 | 17 | 11 | 0 | -1 | 5 | 4 | 3 | 5 | 4 | 0 | 3 | 3 | 4 | 5 | 4 | 5 | 10 | 5 | 5 | 8 | 10 | 9 | 6 | 4 | 3 | 3 | 6 | 4 | 6 | 5 | 9 | 2 |
| RNF219 | chr13:78643026-78643168\|78643135 | 7 | 4 | 2 | 5 | 4 | 4 | 2 | 7 | 4 | 7 | 3 | 35 | 32 | 4 | 5 | 3 | 4 | 3 | 3 | 3 | 7 | 2 | -1 | 3 | 2 | 3 | 2 | 3 | 4 | 3 | 1 | 4 | 2 | 5 | 4 | 2 | 5 | 4 | 4 | 4 | 4 | 7 | 6 | 5 | 3 | 4 | 2 | 3 | 3 | 3 | 3 |

SFig5, pg3

| Gene Symbol | Exon Location (hg38, human) | Adipose - Subcutaneous | Adipose - Visceral (Omentum) | Adrenal Gland | Artery - Aorta | Artery - Coronary | Artery - Tibial | Bladder | Breast - Mammary Tissue | Brain - Amygdala | Brain - Anterior cingulate cortex (BA24) | Brain - Caudate (basal ganglia) | Brain - Cerebellar Hemisphere | Brain - Cerebellum | Brain - Cortex | Brain - Frontal Cortex (BA9) | Brain - Hippocampus | Brain - Hypothalamus | Brain - Nucleus accumbens (basal ganglia) | Brain - Putamen (basal ganglia) | Brain - Spinal cord (cervical c-1) | Brain - Substantia nigra | Cervix - Ectocervix | Cervix - Endocervix | Colon - Sigmoid | Colon - Transverse | Esophagus - Gastroesophageal Junction | Esophagus - Mucosa | Esophagus - Muscularis | Fallopian Tube | Heart - Atrial Appendage | Heart - Left Ventricle | Kidney - Cortex | Liver | Lung | Minor Salivary Gland | Muscle - Skeletal | Nerve - Tibial | Ovary | Pancreas | Pituitary | Prostate | Skin - Not Sun Exposed (Suprapubic) | Skin - Sun Exposed (Lower leg) | Small Intestine - Terminal Ileum | Spleen | Stomach | Testis | Thyroid | Uterus | Vagina | Whole Blood |
|---|---|---|---|---|---|---|---|---|---|---|---|---|---|---|---|---|---|---|---|---|---|---|---|---|---|---|---|---|---|---|---|---|---|---|---|---|---|---|---|---|---|---|---|---|---|---|---|---|---|---|---|---|
| MLLT4 | chr6:167893862-167893906 | 0 | 0 | 5 | 0 | 0 | 0 | 1 | 0 | 19 | 40 | 20 | 88 | 85 | 42 | 48 | 22 | 55 | 35 | 21 | 14 | 22 | 1 | 6 | 1 | 1 | 1 | 1 | 1 | 21 | 3 | 3 | 2 | 2 | 3 | 1 | 2 | 0 | 1 | 1 | 14 | 1 | 0 | 0 | 1 | 0 | 2 | 21 | 1 | 0 | 0 | 1 |
| ROBO2 | chr3:77602935-77603060 | 1 | 1 | 7 | 1 | 2 | 2 | 0 | 1 | 61 | 78 | 75 | 72 | 74 | 73 | 72 | 67 | 79 | 84 | 76 | 47 | 67 | -1 | 0 | 12 | 10 | 0 | 0 | 1 | -1 | -1 | -1 | 0 | -1 | 0 | 0 | -1 | 1 | 1 | 6 | 11 | 3 | 1 | 0 | 7 | 0 | 0 | 19 | 3 | 0 | 1 | -1 |
| PTPRA | chr20:2975215-2975241 | 8 | 10 | 9 | 9 | 9 | 9 | 8 | 8 | 37 | 50 | 40 | 53 | 53 | 52 | 52 | 38 | 41 | 47 | 37 | 15 | 23 | 7 | 8 | 10 | 8 | 10 | 6 | 10 | 9 | 10 | 11 | 7 | 7 | 11 | 6 | 12 | 13 | 9 | 6 | 20 | 8 | 7 | 6 | 9 | 12 | 8 | 5 | 7 | 11 | 8 | 14 |
| TNS2 | chr12:53056635-53056739 | 0 | 1 | 1 | 0 | 0 | 0 | 0 | 0 | 5 | 5 | 5 | 22 | 23 | 6 | 6 | 6 | 6 | 5 | 4 | 5 | 4 | 0 | 0 | 1 | 0 | 0 | 0 | 1 | 0 | 2 | 4 | 0 | 3 | 0 | 1 | 7 | 1 | 0 | 0 | 1 | 1 | 0 | 0 | 0 | 0 | 0 | 2 | 0 | 0 | 0 | 1 |
| SLF1 | chr5:94712689-94713118 | 0 | 1 | 0 | 2 | 1 | 3 | -1 | 1 | 23 | 50 | 22 | 52 | 55 | 49 | 48 | 37 | 36 | 36 | 11 | 32 | 17 | 0 | -1 | 3 | 2 | 2 | 1 | 1 | 0 | 15 | 5 | 0 | 1 | 1 | 0 | 12 | 1 | 7 | 0 | 3 | 1 | 1 | 1 | 0 | 1 | 1 | 7 | 2 | 2 | 1 | 2 |
| PLEKHH2 | chr2:43700971-43701081 | 4 | 3 | 1 | 4 | 2 | 4 | 2 | 3 | 33 | 35 | 26 | 36 | 31 | 33 | 35 | 31 | 33 | 29 | 25 | 38 | 35 | 7 | 1 | 4 | 4 | 4 | 2 | 4 | 2 | 7 | 3 | 1 | 1 | 3 | 2 | 4 | 6 | 4 | 2 | 4 | 3 | 4 | 4 | 2 | 2 | 4 | 6 | 2 | 4 | 3 | 8 |
| EPB41L1 | chr20:36200873-36207780\|36200977 | 0 | 0 | 0 | 5 | 2 | 2 | 0 | 0 | 12 | 22 | 15 | 35 | 31 | 25 | 28 | 13 | 29 | 13 | 15 | 15 | 35 | 1 | 1 | 6 | 5 | 2 | 0 | 2 | 1 | 1 | 1 | 0 | 0 | 1 | 1 | 4 | 0 | 1 | 1 | 5 | 1 | 1 | 1 | 5 | 0 | 2 | 1 | 0 | 1 | 1 | 7 |
| ATP1B3 | chr3:141916376-141916574\|141916440 | 5 | 8 | 2 | 4 | 3 | 3 | 3 | 5 | 22 | 23 | 34 | 37 | 46 | 22 | 14 | 22 | 10 | 40 | 39 | 7 | 7 | 2 | 2 | 3 | 2 | 2 | 1 | 2 | 7 | 5 | 2 | 2 | 3 | 3 | 1 | 4 | 8 | 6 | 1 | 15 | 3 | 5 | 4 | 3 | 1 | 4 | 2 | 2 | 3 | 2 | 1 |
| ANKS6 | chr9:98772156-98772338 | 0 | 0 | 1 | 0 | 1 | 0 | 0 | 0 | 7 | 15 | 9 | 16 | 18 | 20 | 18 | 8 | 15 | 9 | 10 | 9 | 12 | 1 | 0 | 0 | 0 | 0 | 0 | 0 | 0 | 1 | 1 | 1 | 1 | 0 | 0 | 5 | 0 | 0 | 1 | 1 | 0 | 0 | 1 | 0 | 0 | 0 | 5 | 1 | 0 | 0 | 0 |
| CLTA | chr9:36209267-36210660 | 0 | 0 | 1 | 0 | 0 | 0 | 0 | 0 | 42 | 58 | 37 | 68 | 67 | 59 | 63 | 41 | 45 | 44 | 34 | 7 | 26 | 0 | 0 | 2 | 1 | 1 | 0 | 1 | 1 | 6 | 10 | 0 | 0 | 0 | 0 | 17 | 1 | 1 | 1 | 18 | 0 | 0 | 0 | 1 | 1 | 1 | 0 | 0 | 0 | 0 | 1 |
| ARL6 | chr3:97791953-97791965 | 0 | 0 | 0 | 1 | 0 | 1 | 0 | 0 | 10 | 19 | 20 | 25 | 27 | 18 | 20 | 14 | 5 | 15 | 19 | 2 | 6 | -1 | 0 | 0 | 1 | 0 | 0 | 0 | 0 | 0 | 1 | 0 | 0 | 1 | 0 | 2 | 0 | 0 | 0 | 10 | 0 | 0 | 1 | 0 | 0 | 1 | 0 | 1 | 1 | 1 | 0 |
| SNAP23 | chr15:42521614-42521708\|42522014 | 0 | 0 | 1 | 0 | 0 | 1 | 1 | 1 | 5 | 13 | 6 | 64 | 67 | 19 | 20 | 6 | 10 | 11 | 7 | 2 | 3 | 1 | 1 | 2 | 1 | 2 | 0 | 2 | 1 | 3 | 4 | 1 | 1 | 1 | 1 | 5 | 1 | 2 | 1 | 24 | 2 | 1 | 1 | 1 | 1 | 1 | 9 | 2 | 1 | 1 | 1 |
| LY6G5C | chr6:31678422-31678628\|31678604 | 0 | 0 | 0 | 0 | 0 | 0 | 0 | 0 | 16 | 25 | 47 | 73 | 74 | 37 | 32 | 16 | 22 | 40 | 54 | 13 | 12 | 0 | 0 | 0 | 0 | 0 | 0 | 0 | 0 | 15 | 15 | 0 | 0 | 0 | 0 | 7 | 0 | 0 | 0 | 0 | 0 | 0 | 1 | 0 | 0 | 0 | 19 | 0 | 1 | 0 | 0 |
| TWF1 | chr12:43798565-43798585 | 0 | 0 | 2 | 0 | 0 | 0 | 1 | 1 | 50 | 72 | 57 | 55 | 78 | 73 | 66 | 55 | 44 | 74 | 50 | 7 | 24 | 0 | -1 | 2 | 1 | 1 | 0 | 2 | 2 | 24 | 18 | 0 | 0 | 0 | 0 | 5 | 0 | 0 | 0 | 9 | 1 | 0 | 0 | 0 | 0 | 1 | 0 | 0 | 1 | 0 | 1 |
| ACAP2 | chr3:195295196-195295300 | 1 | 1 | 1 | 0 | 0 | 0 | 2 | 1 | 18 | 35 | 17 | 77 | 86 | 35 | 47 | 22 | 38 | 33 | 19 | 3 | 13 | 1 | 0 | 2 | 5 | 1 | 2 | 2 | 2 | 2 | 1 | 1 | 1 | 2 | 1 | 2 | 0 | 0 | 2 | 15 | 1 | 1 | 1 | 4 | 1 | 3 | 31 | 1 | 0 | 1 | 2 |
| MAPK8IP3 | chr16:1758978\|1758301-1758995 | 4 | 7 | 7 | 4 | 6 | 4 | 9 | 6 | 64 | 86 | 71 | 88 | 88 | 83 | 86 | 67 | 65 | 76 | 66 | 32 | 58 | 6 | 12 | 8 | 7 | 8 | 6 | 7 | 4 | 25 | 26 | 6 | 6 | 4 | 5 | 19 | 10 | 11 | 7 | 14 | 6 | 6 | 7 | 5 | 3 | 6 | 5 | 7 | 7 | 6 | 3 |
| PHTF1 | chr1:113701553\|113701547-113701821 | 1 | 2 | 0 | 3 | 3 | 1 | 0 | 0 | 9 | 10 | 11 | 22 | 27 | 13 | 11 | 11 | 6 | 14 | 9 | 6 | 4 | 0 | 2 | 2 | 2 | 2 | 1 | 2 | 0 | 8 | 3 | 1 | 1 | 3 | 0 | 7 | 2 | 4 | 0 | 2 | 2 | 1 | 4 | 1 | 1 | 1 | 1 | 2 | 3 | 2 | 3 |
| EHBP1 | chr2:62858406-62858510 | 0 | 0 | 2 | 0 | 0 | 0 | 1 | 0 | 31 | 64 | 32 | 90 | 85 | 62 | 73 | 37 | 55 | 47 | 32 | 13 | 41 | 0 | 0 | 1 | 1 | 0 | 0 | 0 | 0 | 1 | 3 | 0 | 1 | 0 | 0 | 15 | 0 | 0 | 0 | 13 | 0 | 0 | 1 | 1 | 0 | 1 | 29 | 0 | 0 | 1 | 1 |
| CMC2 | chr16:80998285-81001334\|80998351 | 8 | 7 | 4 | 3 | 5 | 3 | 2 | 7 | 38 | 43 | 38 | 44 | 44 | 41 | 42 | 30 | 37 | 42 | 34 | 16 | 28 | 1 | 1 | 4 | 4 | 3 | 2 | 4 | 3 | 2 | 2 | 8 | 7 | 6 | 3 | 2 | 11 | 2 | 4 | 12 | 6 | 2 | 2 | 5 | 8 | 4 | 7 | 4 | 2 | 2 | 7 |
| AGAP2 | chr12:57728318-57728377 | 2 | 2 | 6 | 6 | 2 | 16 | 0 | 3 | 95 | 97 | 95 | 90 | 90 | 97 | 97 | 95 | 79 | 96 | 94 | 74 | 78 | 0 | 0 | 4 | 2 | 2 | 4 | 1 | 0 | 6 | 1 | 5 | 5 | 2 | 3 | 2 | 5 | 14 | 5 | 21 | 3 | 6 | 27 | 1 | 0 | 2 | 5 | 3 | 5 | 4 | 0 |
| HERC4 | chr10:67959113-67959136 | 3 | 8 | 6 | 3 | 4 | 2 | 3 | 3 | 32 | 50 | 39 | 74 | 74 | 56 | 64 | 44 | 50 | 46 | 38 | 29 | 34 | 3 | 3 | 8 | 4 | 5 | 4 | 6 | 2 | 5 | 6 | 4 | 11 | 3 | 2 | 4 | 8 | 5 | 5 | 31 | 4 | 2 | 2 | 3 | 5 | 4 | 6 | 4 | 5 | 5 | 5 |
| DGKQ | chr4:964080\|964073\|963438-964093 | 2 | 3 | 1 | 1 | 2 | 1 | 1 | 2 | 4 | 4 | 15 | 55 | 58 | 9 | 5 | 4 | 4 | 17 | 14 | 5 | 3 | 3 | 0 | 2 | 6 | 1 | 3 | 1 | 5 | 2 | 2 | 4 | 4 | 5 | 2 | 11 | 3 | 1 | 2 | 5 | 3 | 4 | 4 | 8 | 13 | 3 | 14 | 4 | 2 | 3 | 15 |
| RPGRIP1L | chr16:53659215-53659226 | 0 | 0 | 2 | 0 | 0 | 0 | 0 | 0 | 28 | 60 | 30 | 80 | 77 | 65 | 75 | 18 | 34 | 44 | 43 | 24 | 52 | 0 | 0 | 2 | 1 | 1 | 0 | 0 | 0 | 1 | 0 | 0 | 0 | 0 | 0 | 0 | 0 | 0 | 2 | 31 | 0 | 0 | 0 | 2 | 0 | 4 | 4 | 0 | 0 | 2 | 0 |
| TAOK3 | chr12:118235795\|118236164-118236693 | 23 | 17 | 4 | 9 | 10 | 9 | 5 | 24 | 51 | 63 | 35 | 86 | 90 | 67 | 58 | 53 | 55 | 40 | 35 | 27 | 42 | 17 | 12 | 9 | 5 | 10 | 3 | 9 | 12 | 9 | 10 | 9 | 5 | 13 | 5 | 7 | 17 | 12 | 7 | 14 | 7 | 7 | 7 | 7 | 5 | 6 | 19 | 8 | 16 | 8 | 1 |
| ANK3 | chr10:60069978-60076454\|60076448 | 0 | 0 | 2 | 0 | 0 | 1 | 1 | 0 | 27 | 48 | 28 | 52 | 53 | 46 | 51 | 34 | 45 | 39 | 26 | 40 | 54 | 1 | 0 | 1 | 0 | 1 | 1 | 0 | 0 | 1 | 2 | 0 | 0 | 0 | 0 | 3 | 0 | 0 | 0 | 17 | 0 | 0 | 1 | 0 | 0 | 1 | 14 | 0 | 1 | 1 | 3 |
| UNC13B | chr9:35401951\|35401841-35402007 | 0 | 0 | 0 | 0 | 0 | 0 | 0 | 0 | 17 | 41 | 29 | 78 | 78 | 61 | 64 | 25 | 28 | 35 | 29 | 8 | 16 | 0 | 0 | 0 | 0 | 0 | 0 | 0 | 3 | 3 | 0 | 0 | 0 | 0 | 1 | 0 | 0 | 0 | 0 | 0 | 0 | 0 | 0 | 0 | 0 | 0 | 32 | 0 | 0 | 0 | 3 |
| FMNL1 | chr17:45234832-45234849 | 17 | 15 | 12 | 13 | 12 | 13 | 17 | 20 | 53 | 79 | 42 | 90 | 95 | 78 | 76 | 65 | 44 | 50 | 47 | 16 | 48 | 38 | 17 | 21 | 16 | 21 | 16 | 21 | 14 | 19 | 20 | 12 | 10 | 13 | 13 | 21 | 18 | 16 | 16 | 28 | 16 | 17 | 18 | 12 | 10 | 15 | 8 | 12 | 18 | 21 | 10 |
| NEO1 | chr15:73274692-73274724 | 6 | 5 | 6 | 4 | 4 | 5 | 7 | 5 | 25 | 34 | 27 | 74 | 71 | 34 | 44 | 26 | 34 | 32 | 25 | 14 | 23 | 4 | 3 | 7 | 3 | 7 | 5 | 8 | 7 | 8 | 10 | 5 | 5 | 5 | 3 | 10 | 7 | 4 | 6 | 20 | 6 | 5 | 4 | 4 | 6 | 6 | 27 | 3 | 5 | 5 | 5 |
| PTPRU | chr1:29289683-29289712 | 0 | 0 | 22 | 0 | 0 | 0 | 0 | 0 | 50 | 77 | 37 | 90 | 87 | 80 | 85 | 64 | 73 | 51 | 36 | 9 | 60 | 0 | 0 | 1 | 1 | 0 | 0 | 0 | 0 | 0 | 0 | 1 | 0 | 0 | 0 | 0 | 0 | 0 | 2 | 30 | 0 | 0 | 0 | 2 | 0 | 0 | 2 | 0 | 0 | 0 | 1 |
| ABI2 | chr2:203382189-203382206 | 0 | 0 | 0 | 0 | 0 | 0 | 0 | 0 | 5 | 16 | 9 | 35 | 27 | 17 | 19 | 8 | 8 | 16 | 8 | 1 | 4 | 0 | 0 | 0 | 0 | 0 | 0 | 0 | 0 | 6 | 5 | 0 | 0 | 0 | 1 | 0 | 0 | 0 | 0 | 9 | 0 | 0 | 0 | 0 | 0 | 0 | 3 | 0 | 0 | 0 | 0 |
| MPP3 | chr17:43816677-43816697 | 1 | 1 | 2 | 1 | 0 | 1 | 1 | 1 | 56 | 69 | 47 | 97 | 96 | 67 | 76 | 50 | 74 | 55 | 31 | 32 | 50 | 4 | 1 | 2 | 3 | 1 | 3 | 2 | 3 | 0 | 0 | 2 | 2 | 2 | 1 | 0 | 0 | 1 | 2 | 42 | 1 | 1 | 3 | 2 | 2 | 3 | 2 | 1 | 1 | 1 | 3 |
| SYNJ1 | chr21:32676332-32676355 | 0 | 1 | 4 | 0 | 0 | 0 | 0 | 0 | 32 | 76 | 41 | 93 | 88 | 64 | 82 | 46 | 70 | 59 | 40 | 12 | 41 | 0 | 0 | 1 | 1 | 1 | 0 | 1 | 0 | 0 | 0 | 0 | 1 | 1 | 0 | 2 | 1 | 2 | 1 | 38 | 0 | 0 | 1 | 2 | 0 | 1 | 3 | 0 | 0 | 2 | 1 |
| ATL1 | chr14:50629995-50630009 | 2 | 2 | 5 | 2 | 2 | 3 | 3 | 3 | 52 | 78 | 65 | 86 | 85 | 80 | 85 | 58 | 65 | 71 | 63 | 17 | 46 | 4 | 5 | 7 | 4 | 3 | 2 | 3 | 1 | 3 | 3 | 3 | 4 | 2 | 2 | 4 | 3 | 3 | 4 | 38 | 2 | 2 | 2 | 4 | 2 | 4 | 2 | 2 | 3 | 3 | 5 |
| NCKAP1 | chr2:183024978-183024995 | 0 | 0 | 0 | 0 | 0 | 0 | 0 | 0 | 10 | 23 | 10 | 45 | 36 | 24 | 32 | 16 | 17 | 17 | 11 | 2 | 11 | 0 | 0 | 0 | 0 | 0 | 0 | 0 | 0 | 0 | 0 | 0 | 0 | 0 | 0 | 0 | 0 | 0 | 1 | 16 | 0 | 0 | 0 | 0 | 0 | 0 | 0 | 0 | 0 | 0 | 1 |
| DVL1 | chr1:1337018-1337055 | 2 | 2 | 2 | 3 | 4 | 4 | 4 | 3 | 21 | 28 | 18 | 33 | 31 | 29 | 31 | 22 | 14 | 17 | 18 | 7 | 14 | 3 | 1 | 4 | 5 | 3 | 3 | 3 | 5 | 2 | 2 | 6 | 4 | 2 | 3 | 3 | 3 | 2 | 3 | 3 | 4 | 4 | 5 | 6 | 2 | 4 | 13 | 4 | 3 | 3 | 1 |
| BAK1 | chr6:33574506-33574525 | 2 | 2 | 1 | 5 | 4 | 6 | 4 | 3 | 44 | 70 | 63 | 90 | 93 | 79 | 75 | 53 | 62 | 65 | 65 | 16 | 46 | 0 | -1 | 6 | 2 | 4 | 1 | 5 | 0 | 3 | 3 | 0 | 1 | 1 | 1 | 3 | 4 | 1 | 2 | 41 | 4 | 2 | 1 | 2 | 2 | 2 | 1 | 1 | 5 | 1 | 1 |
| COL11A2 | chr6:33183634\|33182932-33183698 | 3 | 6 | 2 | 4 | 8 | 8 | -1 | 3 | 20 | 33 | 37 | 78 | 79 | 45 | 47 | 21 | 33 | 54 | 35 | 22 | 18 | -1 | -1 | 10 | 6 | 8 | 6 | 6 | 0 | 5 | 3 | 1 | 16 | 5 | 5 | 6 | 26 | 5 | 2 | 11 | 5 | 5 | 5 | 7 | 7 | 6 | 12 | 4 | 4 | 10 | 17 |
| EML1 | chr14:99874931-99874987 | 1 | 1 | 1 | 0 | 0 | 0 | 0 | 0 | 15 | 48 | 65 | 29 | 46 | 49 | 60 | 20 | 34 | 72 | 62 | 2 | 9 | 0 | 0 | 0 | 0 | 0 | 0 | 0 | 1 | 1 | 1 | 1 | 1 | 1 | 0 | 0 | 0 | 0 | 1 | 3 | 1 | 1 | 0 | 0 | 0 | 0 | 19 | 0 | 0 | 0 | 9 |
| INO80C | chr18:35487386-35489385 | 13 | 9 | 10 | 9 | 9 | 9 | 10 | 11 | 41 | 57 | 50 | 61 | 60 | 61 | 64 | 46 | 46 | 56 | 50 | 23 | 34 | 8 | 15 | 10 | 10 | 12 | 7 | 11 | 17 | 13 | 11 | 12 | 13 | 11 | 9 | 20 | 19 | 10 | 15 | 24 | 12 | 10 | 10 | 13 | 13 | 11 | 8 | 9 | 11 | 9 | 12 |
| TMEM39A | chr3:119435755-119435935 | 22 | 16 | 17 | 8 | 11 | 10 | 12 | 22 | 66 | 59 | 55 | 64 | 65 | 66 | 55 | 73 | 63 | 52 | 59 | 72 | 68 | 10 | 15 | 10 | 11 | 10 | 11 | 11 | 15 | 10 | 17 | 22 | 20 | 13 | 15 | 13 | 19 | 11 | 12 | 12 | 12 | 14 | 15 | 13 | 17 | 12 | 16 | 16 | 14 | 12 | 9 |
| FHL1 | chrX:136209243\|136209340-136209442 | 3 | 10 | 13 | 11 | 9 | 5 | 12 | 8 | 63 | 76 | 58 | 74 | 90 | 83 | 74 | 74 | 63 | 64 | 58 | 51 | 50 | 17 | 21 | 11 | 11 | 8 | 11 | 7 | 16 | 12 | 6 | 19 | 12 | 10 | 11 | 4 | 11 | 26 | 10 | 27 | 17 | 9 | 7 | 12 | 10 | 10 | 21 | 18 | 20 | 16 | 6 |
| AGRN | chr1:1051993-1052016\|1053493 | 0 | 0 | 1 | 0 | 0 | 0 | 0 | 0 | 18 | 37 | 24 | 20 | 5 | 49 | 53 | 14 | 23 | 18 | 19 | 8 | 11 | 0 | 0 | 1 | 0 | 0 | 0 | 0 | 0 | 1 | 1 | 0 | 0 | 0 | 0 | 1 | 0 | 0 | 0 | 1 | 0 | 0 | 0 | 1 | 0 | 0 | 1 | 0 | 0 | 1 | 1 |
| CAMSAP2 | chr1:200828573-200828605 | 1 | 1 | 1 | 1 | 1 | 1 | 1 | 1 | 21 | 48 | 28 | 72 | 69 | 50 | 63 | 25 | 28 | 41 | 25 | 9 | 17 | 3 | 1 | 2 | 2 | 1 | 1 | 1 | 1 | 3 | 6 | 0 | 0 | 1 | 1 | 30 | 1 | 1 | 1 | 8 | 1 | 1 | 1 | 2 | 1 | 2 | 4 | 1 | 1 | 1 | 7 |
| MTFR1 | chr8:65689566-65689599 | 2 | 2 | 1 | 4 | 4 | 6 | 9 | 2 | 13 | 17 | 12 | 64 | 62 | 21 | 25 | 13 | 12 | 12 | 12 | 7 | 10 | 4 | 2 | 7 | 4 | 7 | 1 | 6 | 9 | 11 | 12 | 1 | 1 | 2 | 1 | 23 | 4 | 3 | 1 | 4 | 4 | 1 | 1 | 4 | 2 | 3 | 0 | 2 | 6 | 2 | 1 |
| AP3S1 | chr5:115906821-115906861\|115912136 | 0 | 0 | 0 | 0 | 1 | 0 | 0 | 0 | 47 | 59 | 39 | 43 | 43 | 62 | 63 | 52 | 34 | 34 | 37 | 20 | 37 | 1 | 0 | 1 | 1 | 1 | 0 | 0 | 1 | 8 | 12 | 0 | 0 | 0 | 0 | 14 | 1 | 0 | 0 | 4 | 0 | 0 | 0 | 0 | 0 | 1 | 1 | 0 | 0 | 0 | 0 |
| MYO9A | chr15:71847415-71847468 | 2 | 4 | 2 | 3 | 3 | 3 | 3 | 2 | 10 | 16 | 16 | 33 | 32 | 17 | 21 | 10 | 14 | 28 | 16 | 3 | 4 | 5 | 2 | 4 | 8 | 3 | 5 | 4 | 3 | 1 | 1 | 4 | 5 | 4 | 6 | 0 | 1 | 2 | 10 | 4 | 4 | 5 | 3 | 8 | 6 | 6 | 3 | 3 | 2 | 4 | 8 |
| GOLT1B | chr12:21507117\|21507944-21508109 | 6 | 6 | 5 | 6 | 5 | 5 | 10 | 7 | 29 | 32 | 27 | 33 | 43 | 32 | 26 | 29 | 28 | 27 | 29 | 19 | 29 | 9 | 9 | 7 | 8 | 7 | 9 | 8 | 15 | 9 | 8 | 12 | 6 | 8 | 9 | 13 | 8 | 6 | 4 | 11 | 9 | 7 | 7 | 11 | 13 | 7 | 11 | 6 | 5 | 6 | 12 |
| RP4-614O4.11 | chr20:35223389-35224952\|35225997 | 5 | 4 | 3 | 4 | 4 | 5 | 2 | 3 | 4 | 9 | 6 | 29 | 19 | 8 | 13 | 8 | 7 | 9 | 8 | 6 | 7 | 3 | 4 | 3 | 3 | 3 | 4 | 3 | 1 | 4 | 4 | 3 | 5 | 4 | 3 | 5 | 3 | 3 | 1 | 3 | 2 | 4 | 6 | 3 | 3 | 3 | 2 | 3 | 3 | 4 | 4 |
| KIF21B | chr1:200974004-200974194 | 0 | 0 | 0 | 0 | 0 | 1 | 0 | 0 | 64 | 72 | 38 | 57 | 48 | 73 | 77 | 73 | 65 | 56 | 39 | 25 | 50 | 0 | 0 | 1 | 0 | 0 | 1 | 0 | 0 | 6 | 5 | 0 | 0 | 0 | 0 | 18 | 0 | 0 | 0 | 12 | 0 | 0 | 1 | 0 | 0 | 0 | 0 | 0 | 1 | 0 | 0 |
| AGBL4 | chr1:49896091-49896119 | 0 | 0 | 1 | 0 | 0 | 0 | 0 | 0 | 3 | 4 | 1 | 21 | 22 | 3 | 2 | 2 | 14 | 1 | 2 | 5 | 19 | -1 | 0 | 6 | 4 | 6 | 0 | -1 | 0 | 0 | 0 | 0 | 0 | 0 | 0 | 0 | 0 | 0 | 2 | 2 | 2 | 0 | 0 | 0 | 0 | 1 | 0 | 0 | 0 | 0 | 0 |
| STYXL1 | chr7:76016775-76016808 | 3 | 3 | 2 | 3 | 4 | 3 | 1 | 2 | 3 | 6 | 5 | 20 | 19 | 8 | 8 | 4 | 3 | 6 | 6 | 2 | 4 | 2 | 2 | 2 | 2 | 3 | 3 | 3 | 2 | 8 | 3 | 2 | 3 | 4 | 3 | 3 | 3 | 2 | 2 | 3 | 2 | 3 | 3 | 4 | 5 | 2 | 1 | 3 | 3 | 3 | 6 |
| PPFIA1 | chr11:70365373-70365967 | 0 | 1 | 2 | 0 | 0 | 0 | 1 | 0 | 9 | 23 | 13 | 78 | 72 | 31 | 41 | 13 | 22 | 21 | 14 | 5 | 8 | 1 | 0 | 2 | 1 | 1 | 1 | 1 | 1 | 1 | 0 | 1 | 1 | 0 | 0 | 17 | 0 | 1 | 1 | 30 | 2 | 0 | 0 | 1 | 0 | 1 | 1 | 1 | 0 | 1 | 0 |
| ALAS1 | chr3:52198672-52198848 | 10 | 14 | 19 | 14 | 12 | 11 | 11 | 12 | 22 | 21 | 21 | 51 | 52 | 20 | 16 | 23 | 21 | 19 | 21 | 25 | 23 | 15 | 15 | 12 | 11 | 12 | 10 | 12 | 11 | 10 | 8 | 17 | 17 | 17 | 15 | 9 | 17 | 14 | 15 | 13 | 20 | 11 | 10 | 17 | 14 | 12 | 13 | 14 | 14 | 13 | 20 |
| DPY30 | chr2:31886788-31887087 | 3 | 6 | 2 | 5 | 7 | 4 | 2 | 6 | 9 | 11 | 15 | 26 | 49 | 18 | 9 | 14 | 33 | 20 | 14 | 11 | 11 | 3 | -1 | 3 | 6 | 4 | 6 | 3 | 3 | 3 | 1 | 9 | 6 | 7 | 6 | 8 | 6 | 6 | 5 | 16 | 6 | 4 | 3 | 12 | 14 | 4 | 0 | 7 | 8 | 6 | 8 |
| PPFIA1 | chr11:70365373\|70364174-70365967 | 0 | 1 | 2 | 0 | 0 | 1 | 1 | 0 | 10 | 23 | 13 | 78 | 72 | 31 | 41 | 13 | 22 | 21 | 14 | 5 | 8 | 1 | 0 | 2 | 1 | 1 | 1 | 1 | 1 | 1 | 0 | 1 | 1 | 0 | 0 | 17 | 0 | 1 | 1 | 30 | 2 | 0 | 1 | 1 | 0 | 1 | 1 | 1 | 1 | 1 | 0 |
| PTPRS | chr19:5229316-5229342 | 1 | 0 | 3 | 1 | 1 | 1 | 3 | 1 | 31 | 45 | 38 | 95 | 94 | 62 | 63 | 39 | 62 | 49 | 42 | 30 | 22 | 1 | 0 | 5 | 4 | 3 | 1 | 4 | 1 | 7 | 12 | 0 | 0 | 1 | 0 | 17 | 0 | 1 | 1 | 40 | 1 | 1 | 2 | 2 | 1 | 2 | 1 | 0 | 1 | 1 | 3 |
| INTS10 | chr8:19849162\|19847292-19849239 | 7 | 7 | 5 | 3 | 4 | 3 | 2 | 6 | 54 | 67 | 56 | 76 | 77 | 68 | 68 | 56 | 58 | 64 | 57 | 35 | 44 | 5 | 3 | 4 | 3 | 3 | 3 | 3 | 4 | 6 | 12 | 3 | 6 | 5 | 3 | 32 | 14 | 4 | 3 | 12 | 4 | 3 | 4 | 4 | 5 | 3 | 18 | 3 | 3 | 4 | 3 |
| ARFGAP1 | chr20:63283851-63283880\|63284512 | 1 | 1 | 4 | 2 | 2 | 2 | 1 | 2 | 41 | 71 | 46 | 93 | 91 | 77 | 82 | 53 | 65 | 60 | 43 | 16 | 35 | 2 | 3 | 4 | 2 | 3 | 1 | 3 | 2 | 3 | 2 | 0 | 0 | 1 | 1 | 11 | 2 | 2 | 1 | 43 | 2 | 1 | 2 | 3 | 3 | 2 | 1 | 1 | 3 | 2 | 2 |
| PTPRM | chr18:8248150-8252499 | 1 | 1 | 5 | 1 | 1 | 1 | 1 | 1 | 31 | 58 | 42 | 64 | 73 | 55 | 60 | 37 | 57 | 50 | 44 | 26 | 43 | 1 | 1 | 2 | 3 | 2 | 1 | 2 | 2 | 13 | 17 | 1 | 2 | 1 | 2 | 15 | 1 | 1 | 1 | 11 | 2 | 1 | 1 | 1 | 1 | 1 | 25 | 1 | 1 | 1 | 3 |
| RNASET2 | chr6:166955467\|166955484\|166955495-166955627 | 3 | 4 | 4 | 4 | 4 | 4 | 5 | 5 | 9 | 15 | 12 | 18 | 26 | 16 | 14 | 8 | 6 | 13 | 12 | 4 | 6 | 3 | 5 | 4 | 3 | 5 | 3 | 5 | 6 | 3 | 6 | 4 | 4 | 5 | 3 | 7 | 10 | 5 | 2 | 5 | 5 | 3 | 3 | 8 | 5 | 3 | 4 | 5 | 7 | 4 | 3 |
| P3H1 | chr1:42756401-42756455\|42756452 | 2 | 2 | 4 | 2 | 2 | 2 | 4 | 3 | 34 | 47 | 43 | 76 | 77 | 56 | 51 | 41 | 43 | 52 | 38 | 32 | 31 | 2 | 3 | 4 | 3 | 3 | 3 | 4 | 3 | 10 | 16 | 3 | 7 | 3 | 3 | 32 | 11 | 2 | 3 | 7 | 4 | 2 | 2 | 4 | 5 | 3 | 16 | 3 | 2 | 3 | 4 |
| SAAL1 | chr11:18093702-18093965 | 2 | 3 | 2 | 2 | 3 | 2 | 1 | 3 | 15 | 14 | 13 | 28 | 30 | 15 | 12 | 15 | 11 | 12 | 16 | 18 | 14 | 3 | 3 | 3 | 2 | 3 | 1 | 3 | 5 | 6 | 4 | 2 | 1 | 3 | 2 | 7 | 7 | 5 | 2 | 6 | 4 | 2 | 2 | 2 | 2 | 2 | 11 | 3 | 3 | 3 | 2 |
| APLP2 | chr11:130137256-130137291 | 25 | 21 | 13 | 11 | 12 | 9 | 8 | 21 | 32 | 61 | 36 | 92 | 90 | 56 | 73 | 42 | 57 | 46 | 36 | 16 | 29 | 12 | 11 | 11 | 9 | 11 | 11 | 10 | 12 | 8 | 7 | 10 | 12 | 21 | 11 | 21 | 23 | 8 | 11 | 17 | 8 | 15 | 17 | 12 | 21 | 12 | 25 | 11 | 11 | 12 | 33 |
| LRRC49 | chr15:70894572-70894669\|70895735 | 2 | 3 | 1 | 1 | 1 | 2 | 3 | 2 | 4 | 5 | 5 | 37 | 38 | 6 | 4 | 2 | 5 | 3 | 5 | 7 | 5 | 0 | 0 | 1 | 2 | 1 | 1 | 1 | 1 | 2 | 2 | 2 | 5 | 3 | 2 | 3 | 2 | 2 | 5 | 3 | 2 | 3 | 4 | 2 | 0 | 2 | 15 | 2 | 2 | 2 | 11 |
| ARRB1 | chr11:75271701-75271724 | 15 | 19 | 14 | 16 | 18 | 16 | 25 | 17 | 66 | 72 | 66 | 82 | 83 | 71 | 72 | 72 | 71 | 71 | 65 | 45 | 67 | 20 | 16 | 27 | 18 | 28 | 19 | 33 | 13 | 23 | 26 | 11 | 18 | 13 | 11 | 24 | 17 | 15 | 18 | 18 | 17 | 12 | 10 | 17 | 15 | 19 | 22 | 12 | 16 | 16 | 11 |
| ZNF720 | chr16:31733567-31733627 | 6 | 4 | 2 | 4 | 4 | 4 | 5 | 5 | 11 | 16 | 12 | 18 | 18 | 19 | 21 | 11 | 9 | 13 | 8 | 4 | 7 | 6 | 3 | 4 | 4 | 4 | 3 | 4 | 1 | 3 | 3 | 3 | 2 | 4 | 3 | 5 | 6 | 3 | 3 | 2 | 3 | 3 | 4 | 4 | 3 | 4 | 2 | 3 | 4 | 3 | 2 |
| SYCP2 | chr20:59885103-59885509 | 6 | 8 | 7 | 10 | 10 | 11 | 4 | 5 | 33 | 30 | 33 | 70 | 62 | 49 | 55 | 28 | 34 | 29 | 34 | 38 | 29 | -1 | -1 | 11 | 10 | 11 | 4 | 9 | -1 | 15 | 9 | 5 | 4 | 5 | 6 | 21 | 15 | 8 | 8 | 11 | 8 | 6 | 5 | 8 | 7 | 6 | 2 | 6 | 11 | 9 | 4 |
| GTF2IP14 | chr7:65106559-65106665 | 1 | 2 | 3 | 1 | 2 | 1 | 0 | 2 | 10 | 17 | 14 | 55 | 57 | 22 | 28 | 17 | 19 | 18 | 15 | 26 | 13 | 0 | -1 | 1 | 1 | 2 | 1 | 1 | 0 | 6 | 16 | 0 | 3 | 2 | 1 | 24 | 2 | 1 | 2 | 2 | 1 | 1 | 1 | 1 | 2 | 1 | 5 | 1 | 2 | 1 | 1 |
| PTK2 | chr8:140762368-140762388 | 0 | 0 | 0 | 0 | 0 | 0 | 0 | 0 | 4 | 17 | 4 | 32 | 40 | 14 | 17 | 6 | 7 | 8 | 5 | 3 | 4 | 0 | 0 | 0 | 0 | 0 | 0 | 0 | 0 | 4 | 4 | 0 | 0 | 0 | 0 | 13 | 0 | 0 | 1 | 15 | 0 | 0 | 0 | 0 | 0 | 0 | 3 | 0 | 0 | 0 | 0 |
| RAD54L | chr1:46268566-46268588 | 1 | 1 | 0 | 2 | 0 | 1 | 0 | 1 | 12 | 19 | 28 | 17 | 17 | 14 | 22 | 20 | 9 | 25 | 31 | 0 | 2 | 0 | 0 | 4 | 0 | 0 | 0 | 0 | 0 | 2 | 1 | 1 | 0 | 2 | 0 | 0 | 0 | 1 | 0 | 7 | 1 | 1 | 3 | 1 | 2 | 0 | 3 | 1 | 0 | 1 | 1 |
| CENPC | chr4:67522201-67522512 | 2 | 2 | 3 | 2 | 2 | 2 | 3 | 3 | 13 | 32 | 22 | 46 | 57 | 36 | 32 | 17 | 19 | 26 | 19 | 6 | 12 | 0 | 0 | 3 | 2 | 1 | 2 | 3 | 0 | 14 | 18 | 0 | 2 | 1 | 1 | 21 | 5 | 1 | 1 | 2 | 2 | 2 | 2 | 3 | 2 | 2 | 2 | 2 | 2 | 2 | 1 |
| TPD52 | chr8:80044167-80050471\|80051270 | 0 | 0 | 10 | 0 | 1 | 0 | 1 | 0 | 62 | 81 | 57 | 76 | 77 | 79 | 84 | 51 | 62 | 74 | 49 | 7 | 35 | 0 | 1 | 3 | 1 | 2 | 0 | 3 | 1 | 19 | 24 | 0 | 0 | 0 | 0 | 12 | 1 | 0 | 2 | 22 | 0 | 0 | 0 | 1 | 0 | 1 | 0 | 0 | 1 | 0 | 1 |
| TPD52 | chr8:80044167-80050471\|80051270 | 0 | 0 | 10 | 0 | 1 | 0 | 1 | 0 | 62 | 81 | 57 | 76 | 77 | 79 | 84 | 51 | 62 | 74 | 49 | 7 | 35 | 0 | 1 | 3 | 1 | 2 | 0 | 3 | 1 | 19 | 24 | 0 | 0 | 0 | 0 | 12 | 1 | 0 | 2 | 22 | 0 | 0 | 0 | 1 | 0 | 1 | 0 | 0 | 1 | 0 | 1 |
| UNC13B | chr9:35299863-35308412 | 0 | 0 | 0 | 0 | 0 | 0 | 0 | 0 | 1 | 2 | 1 | 21 | 24 | 2 | 1 | 0 | 1 | 0 | 1 | 1 | 0 | 0 | 0 | 0 | 0 | 0 | 0 | 0 | 0 | 2 | 0 | 0 | 0 | 0 | 1 | 11 | 0 | 0 | 0 | 0 | 0 | 0 | 0 | 0 | 0 | 4 | 0 | 0 | 0 | 0 | 0 |
| FLNB | chr3:58099638-58099735 | 0 | 0 | 1 | 1 | 2 | 1 | 5 | 0 | 11 | 26 | 22 | 76 | 90 | 26 | 26 | 12 | 22 | 31 | 22 | 7 | 8 | 1 | 0 | 6 | 2 | 3 | 1 | 4 | 2 | 23 | 40 | 1 | 3 | 0 | 0 | 1 | 1 | 0 | 1 | 3 | 3 | 0 | 0 | 1 | 0 | 1 | 1 | 1 | 1 | 1 | 1 |
| RHBDF1 | chr16:60699-60727 | 4 | 5 | 3 | 4 | 4 | 5 | 5 | 4 | 7 | 7 | 7 | 28 | 20 | 9 | 12 | 8 | 7 | 10 | 4 | 8 | 8 | 5 | 6 | 6 | 5 | 5 | 4 | 5 | 2 | 6 | 4 | 5 | 4 | 6 | 3 | 7 | 5 | 6 | 3 | 6 | 6 | 5 | 4 | 5 | 3 | 4 | 8 | 4 | 5 | 6 | 7 |
| ACTN4 | chr19:38728316-38728381 | 3 | 5 | 6 | 3 | 3 | 2 | 4 | 4 | 45 | 61 | 60 | 73 | 79 | 68 | 69 | 41 | 41 | 60 | 59 | 29 | 31 | 5 | 5 | 4 | 3 | 4 | 3 | 4 | 5 | 26 | 26 | 9 | 6 | 4 | 3 | 19 | 7 | 7 | 5 | 12 | 5 | 3 | 2 | 4 | 7 | 4 | 20 | 6 | 5 | 4 | 4 |
| DCTN1 | chr2:74363614-74363628 | 1 | 0 | 4 | 0 | 0 | 1 | 1 | 1 | 38 | 71 | 51 | 69 | 67 | 74 | 80 | 44 | 51 | 63 | 43 | 7 | 29 | 0 | 0 | 4 | 3 | 1 | 1 | 1 | 0 | 0 | 0 | 0 | 1 | 1 | 1 | 0 | 1 | 0 | 2 | 34 | 1 | 1 | 2 | 2 | 1 | 2 | 1 | 0 | 0 | 1 | 1 |
| SLC16A4 | chr1:110390168-110390297 | 2 | 2 | 2 | 1 | 2 | 1 | 2 | 2 | 3 | 7 | 6 | 21 | 38 | 15 | 6 | 7 | 4 | 7 | 6 | 3 | 3 | -1 | -1 | 3 | 4 | 2 | 3 | 2 | 1 | 3 | 3 | 2 | 3 | 2 | 3 | 7 | 2 | 2 | 4 | 3 | 2 | 4 | 3 | 2 | 4 | 3 | 3 | 1 | 3 | 2 | 19 |
| UBAP2 | chr9:34011627-34011664 | 5 | 4 | 1 | 6 | 6 | 6 | 6 | 5 | 21 | 28 | 20 | 44 | 43 | 29 | 31 | 20 | 18 | 18 | 17 | 10 | 22 | 1 | 13 | 7 | 5 | 6 | 2 | 6 | 12 | 4 | 3 | 1 | 1 | 4 | 3 | 3 | 12 | 8 | 2 | 7 | 5 | 4 | 3 | 7 | 4 | 5 | 14 | 3 | 9 | 5 | 4 |
| ALDH1L2 | chr12:105070195-105070354 | 13 | 3 | 5 | 9 | 5 | 6 | -1 | 12 | 5 | 10 | 8 | 69 | 69 | 23 | 16 | 8 | 12 | 8 | 9 | 6 | 10 | -1 | -1 | 4 | 3 | 6 | 11 | 3 | 0 | 4 | 6 | -1 | -1 | 9 | 5 | 13 | 6 | 6 | 4 | 2 | 7 | 16 | 14 | 5 | 5 | 19 | 17 | 12 | 3 | 11 | 4 |
| VPS13C | chr15:61864641-61867947 | 5 | 8 | 5 | 5 | 6 | 5 | 5 | 7 | 63 | 63 | 60 | 86 | 91 | 72 | 68 | 64 | 64 | 64 | 58 | 59 | 60 | 12 | 11 | 10 | 6 | 7 | 6 | 7 | 10 | 9 | 9 | 5 | 4 | 6 | 4 | 10 | 38 | 17 | 4 | 29 | 8 | 6 | 5 | 7 | 7 | 5 | 23 | 4 | 11 | 12 | 6 |
| EPN2 | chr17:19285620-19285790 | 1 | 1 | 6 | 0 | 0 | 0 | 0 | 1 | 24 | 27 | 20 | 18 | 21 | 24 | 24 | 21 | 22 | 26 | 19 | 8 | 15 | 0 | 1 | 0 | 1 | 0 | 1 | 0 | 3 | 1 | 0 | 5 | 2 | 1 | 1 | 0 | 1 | 1 | 3 | 5 | 2 | 0 | 0 | 1 | 2 | 1 | 6 | 3 | 0 | 0 | 4 |
| PLEKHA6 | chr1:204243114-204243263 | 1 | 1 | 2 | 1 | 1 | 0 | 1 | 0 | 34 | 59 | 55 | 80 | 80 | 67 | 71 | 45 | 60 | 60 | 53 | 35 | 36 | 0 | 0 | 8 | 1 | 2 | 4 | 3 | 0 | 24 | 24 | 1 | 1 | 1 | 1 | 5 | 1 | 1 | 1 | 2 | 1 | 0 | 2 | 1 | 0 | 1 | 26 | 1 | 1 | 2 | 7 |
| RP11-540D14.8 | chr1:155064442-155064560 | 6 | 8 | 5 | 6 | 9 | 5 | 5 | 4 | 16 | 20 | 16 | 81 | 79 | 31 | 25 | 29 | 24 | 21 | 19 | 47 | 40 | 3 | 2 | 7 | 2 | 8 | 4 | 10 | 2 | 17 | 27 | 3 | 7 | 4 | 4 | 33 | 7 | 3 | 5 | 9 | 6 | 3 | 3 | 3 | 7 | 5 | 10 | 6 | 6 | 4 | 4 |
| DENND4C | chr9:19289091\|19289124-19289290 | 1 | 2 | 1 | 1 | 2 | 1 | 3 | 1 | 28 | 25 | 23 | 60 | 65 | 33 | 29 | 23 | 31 | 30 | 26 | 27 | 21 | 2 | 3 | 3 | 1 | 1 | 1 | 2 | 1 | 15 | 10 | 2 | 1 | 1 | 1 | 24 | 4 | 4 | 1 | 10 | 2 | 1 | 1 | 1 | 2 | 1 | 20 | 1 | 3 | 2 | 1 |
| TTYH3 | chr7:2659940-2660040 | 5 | 4 | 3 | 8 | 6 | 11 | 3 | 5 | 6 | 8 | 11 | 39 | 36 | 11 | 12 | 6 | 6 | 8 | 11 | 7 | 7 | 10 | 12 | 6 | 4 | 5 | 3 | 5 | 11 | 6 | 6 | 6 | 5 | 3 | 4 | 9 | 5 | 7 | 4 | 3 | 5 | 5 | 7 | 4 | 4 | 5 | 3 | 5 | 14 | 7 | 4 |
| OSCP1 | chr1:36444004-36444033 | 2 | 1 | 1 | 1 | 2 | 2 | 4 | 1 | 14 | 33 | 28 | 50 | 46 | 42 | 43 | 20 | 15 | 35 | 36 | 11 | 17 | 0 | 0 | 5 | 4 | 3 | 2 | 3 | 0 | 2 | 3 | 1 | 1 | 1 | 2 | 24 | 1 | 0 | 1 | 4 | 1 | 1 | 2 | 3 | 1 | 2 | 3 | 1 | 1 | 1 | 3 |
| CCDC85A | chr2:56372344-56375935 | 1 | 2 | 6 | 3 | 2 | 2 | 5 | 1 | 51 | 55 | 44 | 37 | 42 | 55 | 54 | 47 | 39 | 53 | 38 | 27 | 33 | -1 | -1 | 10 | 7 | 4 | 3 | 3 | 0 | 3 | 10 | 1 | -1 | 2 | 2 | 3 | 3 | 1 | 3 | 15 | 7 | 2 | 2 | 5 | 2 | 5 | 7 | 3 | 4 | 4 | -1 |
| RYR2 | chr1:237769587-237769619 | 0 | 0 | 15 | 0 | 0 | 0 | 0 | 0 | 11 | 14 | 10 | 30 | 30 | 12 | 13 | 10 | 40 | 14 | 9 | 6 | 14 | 0 | 0 | 1 | 1 | 0 | 0 | 0 | 0 | 0 | 0 | 0 | 0 | 0 | 0 | 1 | 0 | 0 | 0 | 2 | 0 | 0 | 2 | 1 | 0 | 1 | 0 | 0 | 0 | 1 | 3 |
| CAMTA1 | chr1:7755638-7755668 | 0 | 0 | 2 | 1 | 1 | 2 | -1 | 0 | 16 | 32 | 40 | 44 | 39 | 41 | 46 | 26 | 10 | 26 | 42 | 16 | 23 | -1 | 0 | 2 | 1 | 1 | 1 | 1 | 0 | 5 | 5 | 0 | 2 | 0 | 0 | 19 | 0 | 0 | 1 | 0 | 1 | 0 | 1 | 1 | 0 | 0 | 0 | 1 | 0 | 4 | 5 |
| EBF1 | chr5:158707568-158707636 | 1 | 1 | 1 | 1 | 1 | 1 | 2 | 1 | 1 | 1 | 18 | 26 | 27 | 3 | 1 | 1 | 9 | 16 | 9 | 15 | 8 | 1 | 0 | 1 | 1 | 1 | 1 | 1 | 1 | 1 | 1 | 2 | 0 | 1 | 0 | 1 | 1 | 1 | 0 | 1 | 1 | 1 | 1 | 1 | 1 | 1 | 15 | 1 | 1 | 1 | 2 |
| SPAG5 | chr17:28582727-28582819 | 1 | 1 | 2 | 1 | 1 | 1 | 0 | 1 | 5 | 6 | 5 | 24 | 27 | 8 | 7 | 6 | 5 | 4 | 4 | 12 | 7 | -1 | 0 | 2 | 1 | 1 | 1 | 2 | -1 | 2 | 3 | 4 | 3 | 1 | 1 | 13 | 1 | 1 | 2 | 2 | 2 | 1 | 1 | 2 | 1 | 1 | 3 | 1 | 1 | 1 | 1 |
| ZNF410 | chr14:73922493-73922697 | 3 | 3 | 2 | 4 | 4 | 5 | 6 | 3 | 7 | 11 | 10 | 39 | 38 | 15 | 14 | 9 | 10 | 11 | 9 | 5 | 5 | 5 | 5 | 7 | 4 | 5 | 2 | 5 | 4 | 12 | 9 | 2 | 2 | 3 | 2 | 11 | 4 | 8 | 2 | 9 | 6 | 2 | 2 | 3 | 2 | 3 | 13 | 3 | 7 | 5 | 1 |
| ABCC1 | chr16:16098873-16098902 | 1 | 1 | 1 | 1 | 1 | 1 | 4 | 1 | 26 | 67 | 31 | 71 | 74 | 68 | 70 | 31 | 30 | 49 | 49 | 4 | 16 | 0 | 2 | 4 | 3 | 3 | 1 | 3 | 2 | 1 | 3 | 0 | 2 | 1 | 2 | 40 | 1 | 1 | 0 | 2 | 1 | 1 | 1 | 3 | 2 | 2 | 1 | 0 | 1 | 1 | 2 |
| RYR2 | chr1:237743600\|237742451-237743623 | 7 | 6 | -1 | 4 | 7 | 5 | -1 | 3 | 63 | 83 | 90 | 92 | 95 | 86 | 91 | 64 | 59 | 93 | 86 | 24 | 36 | -1 | 0 | 10 | 10 | 11 | 10 | 12 | 0 | 25 | 22 | -1 | -1 | 11 | 3 | 17 | 5 | 5 | 2 | 35 | 8 | 10 | 9 | 8 | 4 | 11 | 14 | 3 | 7 | 5 | 16 |
| BICDL1 | chr12:120090911-120091019\|120091401 | 6 | 1 | 21 | 5 | 5 | 14 | 8 | 8 | 92 | 95 | 91 | 98 | 98 | 96 | 97 | 95 | 96 | 94 | 93 | 67 | 89 | 6 | -1 | 28 | 8 | 13 | 4 | 17 | 3 | 4 | 9 | 4 | 9 | 6 | 4 | 39 | 9 | 3 | 6 | 25 | 5 | 7 | 5 | 6 | 6 | 5 | 11 | 2 | 6 | 5 | 8 |
| TRIM9 | chr14:50982942\|50982323-50986144 | 6 | 9 | 11 | 6 | 5 | 14 | -1 | 5 | 14 | 35 | 15 | 79 | 80 | 46 | 46 | 26 | 22 | 24 | 17 | 10 | 18 | -1 | -1 | 12 | 10 | 9 | 7 | 10 | 0 | 6 | 7 | -1 | 3 | 13 | 5 | 33 | 15 | 4 | 8 | 16 | 8 | 5 | 19 | 12 | 3 | 8 | 2 | 5 | 4 | 5 | 10 |
| SYNJ1 | chr21:32634861-32634884 | 24 | 19 | 5 | 4 | 8 | 4 | 4 | 26 | 89 | 96 | 91 | 96 | 96 | 95 | 97 | 92 | 93 | 94 | 92 | 80 | 90 | 9 | 14 | 10 | 6 | 11 | 3 | 10 | 8 | 32 | 20 | 3 | 1 | 9 | 6 | 7 | 20 | 1 | 3 | 26 | 7 | 6 | 7 | 6 | 1 | 6 | 3 | 7 | 7 | 11 | 1 |
| MED12L | chr3:151402928-151402978 | 0 | 0 | 0 | 0 | 0 | 0 | 0 | 0 | 7 | 23 | 7 | 23 | 21 | 40 | 43 | 12 | 19 | 5 | 7 | 16 | 17 | 0 | 0 | 0 | 0 | 0 | 1 | 1 | 0 | 0 | 11 | 0 | 0 | 0 | 0 | 5 | 0 | 0 | 0 | 0 | 0 | 0 | 1 | -1 | 0 | 0 | 0 | 0 | 0 | 0 | 0 |
| MAP7D2 | chrX:20055734-20055832\|20055799 | 5 | 8 | 2 | -1 | -1 | 10 | 0 | 5 | 96 | 97 | 98 | 90 | 94 | 97 | 96 | 97 | 92 | 97 | 96 | 63 | 85 | 0 | -1 | 1 | 2 | 3 | 2 | 2 | 0 | 34 | 33 | 0 | -1 | 5 | -1 | 5 | 6 | 1 | 3 | 8 | 2 | 1 | 7 | 1 | 0 | 0 | 0 | 6 | 0 | 2 | 21 |
| WNK1 | chr12:901561-904494 | 1 | 1 | 0 | 1 | 0 | 1 | 0 | 1 | 8 | 14 | 8 | 34 | 34 | 15 | 18 | 9 | 15 | 10 | 8 | 10 | 9 | 0 | 1 | 1 | 1 | 0 | 0 | 0 | 1 | 4 | 3 | 0 | 0 | 0 | 0 | 18 | 1 | 0 | 0 | 1 | 1 | 0 | 0 | 1 | 0 | 0 | 0 | 0 | 1 | 0 | 0 |
| NFIX | chr19:13087989-13088136 | 8 | 6 | 4 | 5 | 5 | 7 | 7 | 9 | 16 | 18 | 14 | 26 | 26 | 18 | 18 | 18 | 17 | 14 | 16 | 18 | 17 | 7 | 5 | 5 | 5 | 5 | 7 | 5 | 9 | 5 | 3 | 7 | 6 | 7 | 8 | 4 | 9 | 6 | 7 | 8 | 6 | 10 | 10 | 6 | 5 | 6 | 10 | 7 | 7 | 7 | 7 |
| ANKS6 | chr9:98746060-98746094 | 0 | 0 | 1 | 0 | 0 | 0 | 0 | 0 | 20 | 52 | 27 | 53 | 51 | 57 | 60 | 30 | 38 | 25 | 24 | 21 | 31 | 0 | 0 | 0 | 0 | 0 | 0 | 0 | 1 | 6 | 15 | 1 | 0 | 0 | 1 | 19 | 1 | 0 | 0 | 1 | 1 | 0 | 2 | 1 | 0 | 1 | 16 | 0 | 0 | 0 | 2 |
| GDPD5 | chr11:75439340-75439354 | 0 | 0 | 0 | 0 | 0 | 0 | 0 | 0 | 52 | 64 | 53 | 80 | 80 | 67 | 69 | 44 | 49 | 55 | 54 | 21 | 37 | 0 | 0 | 1 | 1 | 0 | 0 | 0 | 1 | 1 | 10 | 0 | 0 | 0 | 1 | 43 | 0 | 0 | 0 | 0 | 0 | 0 | 1 | 0 | 0 | 1 | 0 | 0 | 0 | 0 | 0 |
| GRAMD1A | chr19:35022900-35022911 | 0 | 0 | 1 | 0 | 0 | 0 | 0 | 0 | 38 | 60 | 40 | 89 | 85 | 66 | 70 | 49 | 60 | 47 | 41 | 12 | 36 | 0 | 0 | 2 | 2 | 1 | 0 | 0 | 0 | 0 | 0 | 0 | 0 | 0 | 0 | 1 | 0 | 0 | 1 | 47 | 0 | 0 | 1 | 1 | 0 | 2 | 0 | 0 | 0 | 0 | 0 |
| AC090094.1 | chr8:61682439-61682483 | 2 | 3 | 2 | 2 | 2 | 2 | 2 | 3 | 40 | 61 | 44 | 93 | 87 | 53 | 71 | 51 | 69 | 56 | 44 | 42 | 47 | 3 | 4 | 3 | 4 | 2 | 2 | 2 | 2 | 9 | 19 | 4 | 7 | 3 | 2 | 22 | 5 | 2 | 8 | 42 | 3 | 2 | 2 | 4 | 8 | 3 | 6 | 2 | 2 | 2 | 6 |

SFig5, pg4

| Gene Symbol | Exon Location (hg38, human) | Adipose - Subcutaneous | Adipose - Visceral (Omentum) | Adrenal Gland | Artery - Aorta | Artery - Coronary | Artery - Tibial | Bladder | Breast - Mammary Tissue | Brain - Amygdala | Brain - Anterior cingulate cortex (BA24) | Brain - Caudate (basal ganglia) | Brain - Cerebellar Hemisphere | Brain - Cerebellum | Brain - Cortex | Brain - Frontal Cortex (BA9) | Brain - Hippocampus | Brain - Hypothalamus | Brain - Nucleus accumbens (basal ganglia) | Brain - Putamen (basal ganglia) | Brain - Spinal cord (cervical c-1) | Brain - Substantia nigra | Cervix - Ectocervix | Cervix - Endocervix | Colon - Sigmoid | Colon - Transverse | Esophagus - Gastroesophageal Junction | Esophagus - Mucosa | Esophagus - Muscularis | Fallopian Tube | Heart - Atrial Appendage | Heart - Left Ventricle | Kidney - Cortex | Liver | Lung | Minor Salivary Gland | Muscle - Skeletal | Nerve - Tibial | Ovary | Pancreas | Pituitary | Prostate | Skin - Not Sun Exposed (Suprapubic) | Skin - Sun Exposed (Lower leg) | Small Intestine - Terminal Ileum | Spleen | Stomach | Testis | Thyroid | Uterus | Vagina | Whole Blood |
|---|---|---|---|---|---|---|---|---|---|---|---|---|---|---|---|---|---|---|---|---|---|---|---|---|---|---|---|---|---|---|---|---|---|---|---|---|---|---|---|---|---|---|---|---|---|---|---|---|---|---|---|---|
| ANK2 | chr4:113331972-113332070 | 0 | 0 | 4 | 0 | 0 | 0 | 0 | 0 | 17 | 42 | 21 | 71 | 56 | 38 | 52 | 23 | 36 | 37 | 20 | 5 | 16 | 0 | 0 | 2 | 2 | 1 | 0 | 1 | 0 | 0 | 0 | 0 | 0 | 1 | 0 | 0 | 0 | 0 | 5 | 30 | 1 | 0 | 1 | 2 | 0 | 3 | 0 | 0 | 0 | 1 | 7 |
| WDR35 | chr2:19962295-19962327 | 0 | 0 | 3 | 0 | 0 | 0 | 0 | 0 | 9 | 28 | 10 | 60 | 51 | 27 | 48 | 13 | 32 | 17 | 11 | 3 | 9 | 0 | 0 | 1 | 0 | 0 | 0 | 0 | 0 | 0 | 0 | 0 | 0 | 0 | 0 | 0 | 0 | 0 | 1 | 29 | 0 | 0 | 0 | 1 | 0 | 1 | 1 | 0 | 0 | 0 | 0 |
| GSK3B | chr3:119866590-119866628 | 9 | 12 | 9 | 7 | 8 | 7 | 7 | 9 | 18 | 25 | 15 | 32 | 31 | 24 | 24 | 20 | 32 | 13 | 16 | 26 | 23 | 8 | 12 | 7 | 8 | 9 | 8 | 7 | 12 | 9 | 8 | 7 | 9 | 10 | 10 | 6 | 8 | 12 | 10 | 12 | 9 | 7 | 6 | 9 | 11 | 10 | 5 | 7 | 10 | 9 | 14 |
| TNFSF13B | chr13:108302776-108302893 | 3 | 3 | 2 | 3 | 3 | 2 | -1 | 4 | 9 | 8 | 8 | 15 | 24 | 15 | 11 | 7 | 10 | 7 | 7 | 6 | 8 | -1 | 0 | 4 | 4 | 4 | 3 | 4 | -1 | 3 | 4 | 6 | 5 | 3 | 2 | 5 | 3 | 4 | 4 | 7 | 5 | 3 | 2 | 4 | 4 | 4 | 7 | 2 | 6 | 4 | 2 |
| CHCHD3 | chr7:132886974-132886988 | 0 | 0 | 1 | 0 | 0 | 0 | 0 | 0 | 9 | 18 | 10 | 28 | 45 | 25 | 18 | 12 | 16 | 19 | 10 | 2 | 7 | 0 | 0 | 0 | 0 | 0 | 0 | 0 | 0 | 0 | 0 | 0 | 0 | 0 | 0 | 0 | 0 | 0 | 0 | 26 | 0 | 0 | 0 | 0 | 0 | 0 | 0 | 0 | 0 | 0 | 0 |
| CLIP2 | chr7:74372932-74373036 | 8 | 8 | 16 | 6 | 8 | 6 | 8 | 8 | 77 | 83 | 75 | 88 | 84 | 87 | 88 | 79 | 79 | 79 | 80 | 81 | 81 | 10 | 8 | 14 | 9 | 13 | 8 | 13 | 10 | 25 | 26 | 10 | 14 | 9 | 9 | 25 | 37 | 8 | 8 | 23 | 9 | 8 | 8 | 11 | 8 | 10 | 14 | 10 | 7 | 9 | 11 |
| ABLIM3 | chr5:149240676-149240774\|149241038 | 1 | 1 | 6 | 1 | 2 | 1 | 2 | 1 | 31 | 55 | 27 | 84 | 84 | 61 | 61 | 23 | 40 | 52 | 27 | 11 | 21 | 0 | 1 | 4 | 4 | 3 | 20 | 2 | 2 | 15 | 13 | 5 | 11 | 2 | 8 | 37 | 3 | 2 | 2 | 10 | 11 | 2 | 4 | 3 | 1 | 3 | 21 | 1 | 1 | 5 | 2 |
| ROCK1 | chr18:20960663-20960711 | 0 | 0 | 0 | 0 | 0 | 0 | 0 | 0 | 3 | 9 | 8 | 74 | 74 | 17 | 19 | 4 | 9 | 16 | 6 | 1 | 2 | 0 | 0 | 0 | 0 | 0 | 0 | 0 | 0 | 2 | 2 | 0 | 1 | 0 | 1 | 43 | 0 | 0 | 2 | 2 | 1 | 0 | 0 | 0 | 0 | 1 | 0 | 1 | 0 | 0 | 0 |
| ADD1 | chr4:2926642-2926675 | 5 | 5 | 6 | 2 | 2 | 1 | 2 | 5 | 29 | 28 | 28 | 55 | 53 | 32 | 30 | 40 | 47 | 21 | 35 | 59 | 49 | 3 | 2 | 3 | 4 | 2 | 6 | 2 | 3 | 9 | 13 | 7 | 5 | 4 | 5 | 17 | 12 | 2 | 5 | 7 | 4 | 5 | 5 | 4 | 4 | 4 | 24 | 5 | 2 | 4 | 2 |
| APBA2 | chr15:29094278-29094313 | 3 | 2 | 16 | 2 | 1 | 6 | 3 | 2 | 48 | 82 | 41 | 93 | 91 | 86 | 88 | 65 | 55 | 56 | 38 | 8 | 25 | 0 | 0 | 36 | 12 | 4 | 1 | 2 | 0 | 9 | 11 | 1 | 0 | 1 | 2 | 20 | 2 | 1 | 1 | 28 | 1 | 2 | 18 | 3 | 0 | 4 | 1 | 1 | 1 | 4 | 0 |
| DCTN1 | chr2:74372928-74376762 | 0 | 1 | 3 | 0 | 1 | 0 | 1 | 1 | 35 | 62 | 34 | 86 | 85 | 65 | 70 | 44 | 54 | 53 | 34 | 26 | 41 | 1 | 1 | 2 | 2 | 1 | 0 | 1 | 2 | 10 | 22 | 2 | 1 | 1 | 2 | 27 | 2 | 0 | 2 | 35 | 1 | 0 | 1 | 1 | 0 | 2 | 12 | 1 | 1 | 1 | 0 |
| MINK1 | chr17:4890222\|4890239-4890308 | 7 | 9 | 6 | 8 | 8 | 6 | 5 | 8 | 22 | 35 | 25 | 68 | 67 | 43 | 43 | 30 | 28 | 27 | 25 | 17 | 19 | 8 | 10 | 8 | 6 | 8 | 7 | 8 | 10 | 22 | 31 | 8 | 10 | 10 | 8 | 27 | 11 | 6 | 6 | 12 | 11 | 7 | 6 | 8 | 11 | 7 | 15 | 9 | 11 | 8 | 7 |
| EPB41L1 | chr20:36195329-36195364\|36197734 | 16 | 13 | 4 | 11 | 10 | 11 | 5 | 16 | 92 | 95 | 90 | 97 | 95 | 95 | 96 | 91 | 93 | 95 | 89 | 82 | 87 | 16 | 16 | 16 | 8 | 11 | 2 | 10 | 10 | 19 | 11 | 5 | 5 | 6 | 3 | 16 | 24 | 12 | 5 | 37 | 9 | 6 | 8 | 13 | 6 | 6 | 26 | 6 | 17 | 11 | 34 |
| PTPRK | chr6:127995249-127995266 | 1 | 1 | 1 | 1 | 1 | 1 | 1 | 0 | 47 | 62 | 39 | 63 | 64 | 67 | 75 | 52 | 42 | 49 | 40 | 62 | 58 | 0 | 0 | 2 | 1 | 2 | 0 | 2 | 1 | 18 | 21 | 1 | 0 | 0 | 0 | 27 | 5 | 1 | 0 | 1 | 1 | 0 | 0 | 1 | 0 | 1 | 1 | 1 | 1 | 0 | 5 |
| OFD1 | chrX:13740089-13740213 | 5 | 16 | 6 | 6 | 9 | 5 | 4 | 7 | 29 | 40 | 33 | 69 | 81 | 41 | 43 | 23 | 33 | 40 | 24 | 16 | 14 | 8 | 12 | 10 | 10 | 11 | 7 | 10 | 8 | 16 | 24 | 6 | 32 | 15 | 5 | 15 | 19 | 10 | 7 | 26 | 7 | 4 | 2 | 15 | 23 | 9 | 2 | 7 | 10 | 8 | 16 |
| CERS5 | chr12:50135433-50135616\|50135551\|50135652 | 1 | 0 | 0 | 0 | 0 | 0 | 1 | 0 | 32 | 43 | 34 | 75 | 75 | 53 | 49 | 34 | 32 | 35 | 35 | 37 | 32 | 0 | 0 | 1 | 0 | 1 | 0 | 1 | 0 | 23 | 23 | 0 | 0 | 1 | 1 | 31 | 2 | 0 | 0 | 1 | 1 | 0 | 1 | 1 | 0 | 1 | 1 | 0 | 0 | 0 | 0 |
| RECK | chr9:36056980-36057013\|36057025 | 11 | 6 | 6 | 3 | 5 | 7 | 5 | 11 | 25 | 40 | 19 | 68 | 61 | 38 | 45 | 26 | 27 | 28 | 22 | 21 | 22 | 10 | 4 | 7 | 7 | 6 | 7 | 6 | 8 | 6 | 8 | 6 | 7 | 8 | 7 | 10 | 13 | 5 | 7 | 10 | 7 | 10 | 14 | 8 | 6 | 7 | 35 | 5 | 6 | 6 | 9 |
| SMURF1 | chr7:99050915-99050992 | 1 | 1 | 1 | 1 | 1 | 1 | 1 | 1 | 6 | 16 | 6 | 16 | 21 | 16 | 16 | 7 | 9 | 7 | 7 | 10 | 13 | 0 | 1 | 1 | 1 | 1 | 1 | 1 | 0 | 2 | 2 | 1 | 1 | 1 | 1 | 3 | 2 | 1 | 1 | 1 | 1 | 1 | 1 | 1 | 1 | 1 | 12 | 1 | 1 | 1 | 1 |
| DGKA | chr12:55943297-55943418 | 6 | 6 | 5 | 6 | 6 | 6 | 5 | 8 | 13 | 13 | 14 | 20 | 22 | 15 | 13 | 11 | 11 | 17 | 15 | 10 | 10 | 8 | 9 | 8 | 5 | 7 | 4 | 7 | 8 | 7 | 7 | 7 | 7 | 7 | 4 | 6 | 11 | 9 | 7 | 9 | 7 | 7 | 7 | 6 | 7 | 6 | 7 | 7 | 9 | 7 | 7 |
| CTD-2116N17.1 | chr15:64188375-64188485 | 5 | 10 | 17 | 5 | 9 | 4 | 10 | 6 | 35 | 45 | 37 | 62 | 61 | 47 | 51 | 35 | 39 | 43 | 31 | 20 | 28 | 13 | -1 | 9 | 13 | 10 | 16 | 9 | 18 | 16 | 18 | 15 | 15 | 7 | 12 | 21 | 7 | 14 | 14 | 20 | 16 | 10 | 10 | 12 | 12 | 13 | 6 | 15 | 12 | 14 | 9 |
| PPFIA4 | chr1:203061652-203061678 | 5 | 2 | 43 | 0 | 0 | 1 | 0 | 2 | 86 | 93 | 88 | 97 | 97 | 96 | 96 | 89 | 90 | 90 | 87 | 77 | 81 | 0 | 0 | 7 | 5 | 2 | 2 | 2 | 0 | 0 | 0 | 0 | 1 | 2 | 1 | 0 | 5 | 3 | 15 | 34 | 1 | 1 | 16 | 7 | 1 | 4 | 1 | 4 | 1 | 1 | 11 |
| AZIN2 | chr1:33083018\|33083111-33083142 | 2 | 1 | 1 | 2 | 0 | 1 | -1 | 1 | 17 | 22 | 18 | 36 | 39 | 25 | 22 | 18 | 12 | 17 | 21 | 14 | 13 | 0 | 0 | 1 | 1 | 2 | 1 | 3 | 1 | 5 | 7 | 2 | 2 | 1 | 2 | 19 | 3 | 0 | 0 | 3 | 1 | 1 | 2 | 1 | 3 | 2 | 12 | 1 | 1 | 1 | 3 |
| SHF | chr15:45173576-45173716 | 5 | 4 | 6 | 3 | 5 | 6 | 8 | 4 | 76 | 84 | 74 | 96 | 96 | 86 | 86 | 75 | 82 | 83 | 71 | 71 | 64 | 3 | 2 | 13 | 10 | 8 | 5 | 10 | 3 | 33 | 35 | 4 | 2 | 3 | 2 | 37 | 9 | 3 | 4 | 6 | 5 | 2 | 6 | 6 | 3 | 8 | 14 | 2 | 3 | 3 | 8 |
| CADPS2 | chr7:122393196-122393315 | 6 | 5 | 3 | 19 | 13 | 16 | 12 | 7 | 24 | 67 | 14 | 74 | 74 | 77 | 84 | 51 | 76 | 17 | 8 | 39 | 73 | 15 | 16 | 14 | 9 | 20 | 9 | 18 | 10 | 17 | 17 | 3 | 4 | 5 | 4 | 18 | 8 | 11 | 2 | 25 | 11 | 9 | 14 | 7 | 3 | 6 | 6 | 4 | 24 | 14 | 18 |
| FBXO8 | chr4:174256027-174256145 | 0 | 0 | 0 | 0 | 0 | 0 | 1 | 0 | 4 | 6 | 3 | 42 | 38 | 8 | 9 | 4 | 5 | 6 | 4 | 1 | 2 | 1 | 0 | 2 | 1 | 1 | 0 | 1 | 0 | 4 | 4 | 0 | 0 | 0 | 0 | 22 | 0 | 0 | 0 | 4 | 1 | 0 | 0 | 1 | 0 | 1 | 0 | 0 | 1 | 0 | 0 |
| ARHGAP44 | chr17:12973302-12973319 | 2 | 0 | 14 | 8 | 7 | 9 | 8 | 2 | 55 | 57 | 56 | 58 | 58 | 64 | 68 | 59 | 43 | 61 | 63 | 37 | 62 | 0 | 3 | 6 | 1 | 6 | 2 | 5 | 5 | 0 | 1 | 2 | 1 | 0 | 1 | 1 | 4 | 1 | 3 | 30 | 2 | 1 | 5 | 3 | 1 | 2 | 1 | 0 | 4 | 5 | 9 |
| TMEM255A | chrX:120285625-120285696 | 14 | 6 | 4 | 6 | 1 | 7 | -1 | 6 | 61 | 57 | 42 | 62 | 39 | 47 | 57 | 39 | 66 | 65 | 51 | 42 | 54 | 1 | 4 | 4 | 4 | 9 | 5 | 7 | 2 | 0 | 1 | 1 | 0 | 1 | 2 | 10 | 2 | 0 | 5 | 2 | 2 | 0 | 1 | 4 | 0 | 6 | 0 | 2 | 1 | 2 | 13 |
| SORBS1 | chr10:95355627-95357105 | 0 | 0 | 0 | 0 | 0 | 0 | 0 | 0 | 4 | 12 | 22 | 17 | 14 | 12 | 16 | 6 | 3 | 33 | 25 | 0 | 2 | 0 | 0 | 0 | 0 | 0 | 0 | 0 | 0 | 0 | 0 | 0 | 0 | 0 | 0 | 0 | 0 | 0 | 0 | 2 | 0 | 0 | 0 | 0 | 0 | 0 | 8 | 0 | 0 | 0 | 1 |
| TMEM87B | chr2:112095228-112095299 | 4 | 3 | 4 | 4 | 4 | 5 | 8 | 3 | 24 | 42 | 35 | 65 | 63 | 32 | 47 | 27 | 40 | 38 | 33 | 22 | 22 | 5 | 4 | 5 | 2 | 5 | 2 | 8 | 3 | 17 | 23 | 2 | 2 | 3 | 2 | 28 | 10 | 5 | 2 | 8 | 5 | 2 | 2 | 2 | 4 | 2 | 13 | 3 | 5 | 3 | 2 |
| COL21A1 | chr6:56108788-56108812 | 16 | 17 | 9 | 7 | 9 | 8 | 19 | 13 | 42 | 65 | 39 | 83 | 86 | 63 | 60 | 30 | 45 | 39 | 25 | 32 | 31 | 17 | 10 | 23 | 22 | 20 | 16 | 19 | 25 | 12 | 7 | 4 | 6 | 15 | 18 | 25 | 25 | 15 | 11 | 16 | 20 | 24 | 21 | 20 | -1 | 26 | 9 | 11 | 14 | 16 | -1 |
| LIMK2 | chr22:31266532-31266550 | 1 | 0 | 0 | 5 | 5 | 8 | 4 | 1 | 1 | 3 | 2 | 49 | 44 | 4 | 4 | 3 | 4 | 4 | 3 | 1 | 1 | 2 | 3 | 13 | 2 | 13 | 0 | 14 | 3 | 7 | 10 | 0 | 0 | 0 | 1 | 8 | 1 | 2 | 0 | 3 | 2 | 0 | 0 | 1 | 0 | 1 | 0 | 0 | 7 | 1 | 0 |
| DOCK7 | chr1:62485659-62487412 | 1 | 2 | 1 | 5 | 5 | 8 | 5 | 2 | 11 | 12 | 13 | 35 | 39 | 13 | 11 | 11 | 19 | 17 | 12 | 11 | 12 | 7 | 2 | 9 | 3 | 9 | 1 | 10 | 8 | 4 | 8 | 2 | 1 | 1 | 1 | 12 | 1 | 3 | 1 | 12 | 4 | 0 | 0 | 3 | 1 | 3 | 1 | 2 | 8 | 4 | 0 |
| BICDL1 | chr12:120062752-120062763 | 1 | 0 | 14 | 3 | 1 | 6 | 0 | 1 | 87 | 94 | 85 | 98 | 98 | 96 | 97 | 90 | 94 | 91 | 88 | 55 | 82 | 0 | 0 | 36 | 5 | 2 | 0 | 6 | 0 | 3 | 13 | 0 | 0 | 0 | 0 | 45 | 2 | 0 | 0 | 4 | 0 | 0 | 0 | 1 | 0 | 1 | 0 | 0 | 7 | 0 | 0 |
| PHF21A | chr11:45946076-45946098 | 2 | 3 | 4 | 2 | 2 | 1 | 2 | 2 | 8 | 20 | 12 | 73 | 68 | 27 | 28 | 9 | 15 | 20 | 9 | 3 | 4 | 3 | 3 | 2 | 3 | 2 | 3 | 2 | 4 | 4 | 6 | 3 | 5 | 5 | 3 | 5 | 1 | 1 | 6 | 41 | 2 | 3 | 3 | 7 | 8 | 3 | 8 | 3 | 1 | 2 | 4 |
| CAMSAP1 | chr9:135865332\|135863788-135865364 | 1 | 1 | 1 | 1 | 2 | 1 | 8 | 0 | 45 | 70 | 55 | 86 | 86 | 71 | 77 | 44 | 59 | 69 | 52 | 11 | 27 | 1 | -1 | 6 | 4 | 7 | 1 | 11 | 4 | 21 | 32 | 0 | 2 | 1 | 2 | 37 | 2 | 1 | 1 | 9 | 7 | 0 | 1 | 4 | 1 | 3 | 2 | 1 | 0 | 1 | 2 |
| CLASP2 | chr3:33594951-33596734 | 2 | 4 | 6 | 1 | 2 | 1 | 3 | 3 | 75 | 81 | 72 | 90 | 90 | 80 | 81 | 77 | 83 | 82 | 74 | 82 | 78 | 3 | 1 | 6 | 5 | 3 | 3 | 3 | 4 | 7 | 6 | 5 | 4 | 3 | 3 | 9 | 9 | 2 | 9 | 55 | 3 | 3 | 2 | 6 | 5 | 5 | 10 | 3 | 3 | 4 | 6 |
| CFAP61 | chr20:20069745-20069901 | 0 | 0 | 0 | 0 | 0 | 0 | -1 | 0 | 25 | 74 | 19 | 63 | 65 | 64 | 73 | 39 | 32 | 28 | -1 | -1 | -1 | -1 | 0 | 0 | 0 | 0 | 0 | 0 | 0 | -1 | -1 | 0 | 0 | 0 | 0 | 40 | 0 | 0 | 0 | 2 | 0 | 0 | 0 | 0 | 0 | -1 | 2 | 0 | 0 | 0 | 0 |
| LRRC58 | chr3:120348223-120348314 | 5 | 6 | 1 | 4 | 6 | 5 | 8 | 5 | 14 | 14 | 12 | 45 | 37 | 22 | 20 | 16 | 15 | 24 | 11 | 6 | 9 | -1 | -1 | 8 | 5 | 6 | 3 | 6 | 3 | 9 | 7 | 5 | 2 | 7 | 4 | 12 | 18 | 7 | 2 | 6 | 4 | 2 | 2 | 5 | 4 | 5 | 5 | 4 | 6 | 5 | 6 |
| NTRK1 | chr1:156874383-156874400 | 0 | 0 | 28 | 1 | 2 | 0 | 0 | 0 | 22 | 40 | 88 | 95 | 58 | 47 | 45 | 39 | 72 | 83 | 90 | 38 | 74 | 0 | 0 | 1 | 2 | 0 | 0 | 1 | 0 | 1 | 0 | 0 | 0 | 1 | 0 | 2 | 1 | 1 | 5 | 1 | 1 | 1 | 1 | 2 | 0 | 1 | 4 | 0 | 0 | 22 | 0 |
| CLASP1 | chr2:121427404-121427430 | 1 | 1 | 3 | 1 | 1 | 1 | 1 | 1 | 32 | 58 | 33 | 93 | 88 | 62 | 69 | 45 | 67 | 30 | 34 | 43 | 48 | 2 | 2 | 2 | 2 | 1 | 2 | 1 | 2 | 1 | 2 | 2 | 2 | 1 | 1 | 0 | 1 | 1 | 3 | 12 | 2 | 2 | 1 | 2 | 1 | 2 | 54 | 2 | 1 | 2 | 1 |
| RIPK1 | chr6:3082547-3082660 | 1 | 1 | 1 | 1 | 1 | 1 | 1 | 1 | 15 | 28 | 19 | 81 | 78 | 37 | 38 | 28 | 40 | 23 | 18 | 31 | 27 | 1 | 1 | 2 | 1 | 1 | 0 | 1 | 0 | 11 | 19 | 1 | 2 | 1 | 1 | 45 | 5 | 1 | 1 | 2 | 2 | 1 | 1 | 0 | 1 | 1 | 8 | 1 | 1 | 1 | 0 |
| CCDC138 | chr2:108865497-108865719 | 9 | 10 | 8 | 3 | 8 | 6 | -1 | 9 | 9 | 17 | 9 | 43 | 49 | 24 | 24 | 10 | 13 | 7 | 9 | 13 | 13 | 0 | 0 | 11 | 9 | 7 | 6 | 6 | -1 | 9 | 9 | 10 | 11 | 6 | 6 | 21 | 7 | 10 | 8 | 15 | 9 | 8 | 8 | 3 | 6 | 7 | 9 | 9 | 15 | 8 | 9 |
| B4GALT4 | chr3:119219578-119219665 | 2 | 2 | 1 | 2 | 2 | 2 | 2 | 2 | 8 | 12 | 11 | 27 | 24 | 16 | 15 | 10 | 10 | 13 | 11 | 10 | 11 | 3 | 1 | 4 | 2 | 3 | 1 | 4 | 1 | 7 | 10 | 1 | 2 | 2 | 2 | 11 | 4 | 2 | 2 | 3 | 3 | 1 | 1 | 2 | 3 | 2 | 5 | 1 | 3 | 2 | 2 |
| FEZ2 | chr2:36560785-36560865 | 6 | 6 | 13 | 2 | 5 | 3 | 7 | 6 | 60 | 68 | 54 | 81 | 79 | 72 | 76 | 64 | 63 | 56 | 59 | 73 | 62 | 3 | 2 | 9 | 7 | 7 | 7 | 9 | 6 | 23 | 20 | 7 | 8 | 6 | 7 | 10 | 42 | 3 | 8 | 18 | 7 | 6 | 6 | 9 | 10 | 9 | 21 | 6 | 3 | 6 | 10 |
| PPP4R1 | chr18:9563917-9563951 | 0 | 0 | 0 | 0 | 0 | 0 | 0 | 0 | 30 | 27 | 29 | 7 | 9 | 21 | 19 | 28 | 18 | 29 | 28 | 30 | 26 | 0 | 0 | 0 | 0 | 0 | 0 | 0 | 0 | 2 | 4 | 0 | 0 | 0 | 0 | 3 | 2 | 0 | 0 | 1 | 0 | 0 | 0 | 0 | 0 | 0 | 0 | 0 | 0 | 0 | 0 |
| TTC38 | chr22:46286166-46286314 | 6 | 5 | 1 | 4 | 3 | 5 | 4 | 5 | 2 | 2 | 1 | 23 | 18 | 2 | 1 | 1 | 3 | 2 | 1 | 2 | 1 | 5 | 4 | 4 | 2 | 3 | 3 | 3 | 5 | 1 | 1 | 2 | 2 | 4 | 4 | 2 | 5 | 3 | 5 | 2 | 4 | 4 | 6 | 4 | 4 | 3 | 9 | 3 | 3 | 4 | 3 |
| ANKRD36C | chr2:95884807\|95884834-95884918 | 0 | 1 | 1 | 0 | 0 | 1 | 0 | 0 | 27 | 60 | 48 | 76 | 71 | 61 | 76 | 39 | 69 | 47 | 45 | 8 | 21 | 0 | 0 | 1 | 1 | 1 | 2 | 1 | 0 | 16 | 24 | 2 | 2 | 0 | 1 | 36 | 0 | 0 | 1 | 1 | 1 | 1 | 1 | 0 | 0 | 2 | 0 | 7 | 0 | 1 | 1 |
| DCLK1 | chr13:35788212-35788285 | 3 | 2 | 4 | 2 | 1 | 1 | 0 | 4 | 33 | 52 | 38 | 29 | 26 | 51 | 51 | 43 | 49 | 37 | 35 | 22 | 34 | 0 | 3 | 2 | 2 | 3 | 1 | 2 | 2 | 1 | 3 | 2 | -1 | 1 | 2 | 5 | 2 | 1 | 1 | 3 | 2 | 7 | 5 | 3 | 1 | 3 | 1 | 1 | 2 | 3 | 14 |
| TBC1D2 | chr9:98213785-98220253\|98217197 | 4 | 7 | 6 | 5 | 5 | 3 | 5 | 5 | 42 | 33 | 36 | 53 | 52 | 43 | 37 | 46 | 45 | 40 | 35 | 65 | 48 | 5 | 4 | 5 | 3 | 5 | 3 | 6 | 2 | 11 | 10 | 3 | 10 | 6 | 3 | 30 | 6 | 4 | 3 | 8 | 5 | 2 | 2 | 5 | 10 | 3 | 16 | 3 | 5 | 2 | 6 |
| SFT2D1 | chr6:166331385-166331407\|166332045\|166341819\|166331447 | 8 | 8 | 5 | 9 | 9 | 11 | 8 | 8 | 19 | 30 | 19 | 63 | 63 | 28 | 37 | 18 | 28 | 31 | 19 | 3 | 11 | 7 | 7 | 11 | 5 | 11 | 5 | 10 | 8 | 7 | 6 | 7 | 11 | 6 | 4 | 13 | 5 | 8 | 5 | 41 | 7 | 9 | 9 | 9 | 10 | 6 | 4 | 5 | 9 | 7 | 8 |
| R3HDM2 | chr12:57272453-57272554 | 0 | 0 | 0 | 0 | 0 | 0 | 0 | 0 | 27 | 54 | 29 | 66 | 65 | 58 | 59 | 30 | 28 | 38 | 27 | 6 | 13 | 0 | 0 | 0 | 0 | 0 | 0 | 0 | 0 | 21 | 26 | 0 | 0 | 0 | 1 | 28 | 0 | 0 | 0 | 1 | 0 | 0 | 1 | 0 | 0 | 0 | 0 | 0 | 0 | 0 | 0 |
| RPS6KB2 | chr11:67432356\|67432353-67432492 | 18 | 21 | 9 | 18 | 16 | 16 | 14 | 19 | 51 | 55 | 51 | 75 | 77 | 63 | 62 | 55 | 56 | 54 | 51 | 47 | 50 | 18 | 26 | 20 | 12 | 15 | 7 | 14 | 14 | 17 | 15 | 18 | 9 | 15 | 10 | 16 | 25 | 23 | 8 | 33 | 17 | 10 | 8 | 12 | 12 | 9 | 31 | 24 | 28 | 18 | 9 |
| PRIMPOL | chr4:184691316\|184685823-184691354 | 2 | 6 | 3 | 5 | 4 | 2 | 3 | 3 | 6 | 4 | 4 | 17 | 21 | 5 | 4 | 4 | 6 | 5 | 4 | 2 | 4 | 1 | 4 | 4 | 3 | 3 | 4 | 3 | 2 | 5 | 5 | 5 | 4 | 6 | 2 | 3 | 3 | 7 | 3 | 5 | 5 | 2 | 1 | 4 | 6 | 3 | 2 | 4 | 7 | 6 | 9 |
| KIAA0513 | chr16:85082564-85082593\|85083373 | 3 | 2 | 4 | 4 | 4 | 4 | 8 | 3 | 69 | 87 | 66 | 92 | 91 | 88 | 91 | 80 | 70 | 70 | 66 | 44 | 67 | 2 | 0 | 9 | 5 | 5 | 2 | 7 | 0 | 21 | 25 | 2 | 3 | 2 | 3 | 50 | 3 | 2 | 1 | 18 | 5 | 1 | 3 | 4 | 2 | 3 | 13 | 2 | 2 | 5 | 1 |
| SYNGAP1 | chr6:33442453-33442494 | 6 | 4 | 8 | 13 | 12 | 18 | 23 | 5 | 69 | 83 | 73 | 79 | 78 | 85 | 87 | 79 | 43 | 75 | 77 | 49 | 57 | 7 | 4 | 24 | 18 | 24 | 6 | 28 | 9 | 18 | 17 | 7 | 8 | 5 | 5 | 16 | 17 | 4 | 3 | 9 | 11 | 6 | 7 | 9 | 5 | 16 | 27 | 5 | 7 | 6 | 5 |
| RPS6KB2 | chr11:67432356-67432492\|67432488 | 14 | 16 | 7 | 14 | 12 | 12 | 11 | 15 | 38 | 41 | 40 | 56 | 58 | 49 | 48 | 42 | 44 | 42 | 40 | 36 | 39 | 14 | 19 | 16 | 10 | 11 | 5 | 11 | 12 | 13 | 11 | 14 | 7 | 11 | 7 | 12 | 19 | 18 | 6 | 25 | 13 | 7 | 7 | 9 | 10 | 7 | 24 | 18 | 22 | 13 | 7 |
| RPS6KB2 | chr11:67432353-67432492\|67432488 | 14 | 16 | 7 | 14 | 12 | 13 | 11 | 16 | 40 | 42 | 40 | 58 | 59 | 48 | 46 | 43 | 44 | 41 | 39 | 37 | 39 | 16 | 20 | 16 | 10 | 12 | 5 | 11 | 10 | 13 | 11 | 15 | 8 | 12 | 7 | 12 | 20 | 18 | 6 | 26 | 13 | 8 | 7 | 9 | 10 | 7 | 24 | 19 | 22 | 14 | 7 |
| APIP | chr11:34896761-34896855 | 6 | 9 | 7 | 5 | 5 | 4 | 10 | 9 | 35 | 42 | 29 | 33 | 40 | 39 | 35 | 30 | 34 | 33 | 30 | 19 | 25 | 11 | 6 | 7 | 7 | 7 | 6 | 8 | 2 | 18 | 14 | 7 | 6 | 7 | 8 | 14 | 11 | 8 | 7 | 20 | 10 | 6 | 5 | 7 | 6 | 8 | 8 | 5 | 8 | 8 | 5 |
| CACNA2D3 | chr3:54816853-54816870 | 1 | 0 | 9 | 0 | 1 | 1 | 0 | 1 | 15 | 42 | 52 | 41 | 35 | 46 | 52 | 44 | 35 | 54 | 45 | 12 | 13 | 0 | 0 | 2 | 2 | 1 | 0 | 3 | 0 | 16 | 15 | 0 | -1 | 0 | 0 | 2 | 0 | 0 | 0 | 4 | 3 | 0 | 0 | 2 | 0 | 1 | 0 | 0 | 2 | 1 | 3 |
| RUBCN | chr3:197699187-197699231\|197699673 | 3 | 4 | 7 | 5 | 4 | 7 | 3 | 4 | 61 | 78 | 60 | 94 | 93 | 80 | 83 | 61 | 77 | 75 | 59 | 45 | 54 | 2 | 6 | 9 | 6 | 9 | 1 | 7 | 8 | 12 | 12 | 2 | 2 | 4 | 2 | 16 | 22 | 7 | 4 | 53 | 4 | 1 | 1 | 5 | 2 | 6 | 27 | 2 | 4 | 4 | 7 |
| RIF1 | chr2:151469054-151469107 | 6 | 3 | 1 | 6 | 5 | 11 | 9 | 6 | 5 | 21 | 9 | 35 | 33 | 20 | 33 | 7 | 4 | 14 | 7 | 2 | 4 | 8 | 3 | 9 | 5 | 6 | 1 | 5 | 3 | 2 | 3 | 1 | 1 | 3 | 1 | 15 | 4 | 3 | 1 | 1 | 3 | 4 | 4 | 4 | 1 | 5 | 3 | 1 | 6 | 3 | 1 |
| CACNA2D1 | chr7:81983314-81983334 | 9 | 10 | 8 | 4 | 7 | 12 | 16 | 11 | 81 | 89 | 81 | 96 | 93 | 90 | 96 | 79 | 89 | 91 | 83 | 45 | 37 | -1 | -1 | 20 | 19 | 15 | 8 | 14 | -1 | 5 | 3 | 5 | -1 | 10 | 1 | 1 | 12 | 13 | 12 | 51 | 7 | 2 | 1 | 14 | 11 | 15 | 8 | 4 | 18 | 9 | -1 |
| AGAP1 | chr2:235718381-235718392 | 2 | 1 | 1 | 1 | 2 | 1 | 1 | 1 | 73 | 84 | 64 | 88 | 88 | 80 | 85 | 70 | 68 | 77 | 65 | 53 | 61 | 0 | 0 | 4 | 2 | 1 | 0 | 2 | 2 | 11 | 15 | 0 | 0 | 1 | 2 | 26 | 51 | 1 | 0 | 7 | 1 | 1 | 1 | 2 | 1 | 1 | 1 | 0 | 0 | 2 | 3 |
| CERS5 | chr12:50132962-50134068\|50133064 | 6 | 7 | 8 | 7 | 7 | 5 | 7 | 7 | 69 | 76 | 74 | 96 | 96 | 83 | 81 | 78 | 83 | 78 | 74 | 80 | 72 | 6 | 7 | 10 | 9 | 9 | 6 | 10 | 7 | 38 | 38 | 7 | 7 | 7 | 7 | 46 | 17 | 8 | 8 | 8 | 8 | 5 | 5 | 8 | 7 | 8 | 17 | 8 | 7 | 8 | 5 |
| ERC1 | chr12:1410389-1410453 | 0 | 0 | 1 | 0 | 0 | 0 | 0 | 0 | 18 | 35 | 25 | 87 | 82 | 36 | 38 | 28 | 41 | 44 | 22 | 6 | 15 | 0 | 0 | 1 | 1 | 0 | 0 | 0 | 0 | 8 | 7 | 0 | 0 | 0 | 1 | 0 | 0 | 0 | 5 | 54 | 1 | 0 | 0 | 1 | 0 | 1 | 9 | 1 | 0 | 0 | 1 |
| TENM3 | chr4:182744143-182744163 | 8 | 9 | 10 | 8 | 22 | 13 | -1 | 13 | 49 | 85 | 32 | 95 | 97 | 93 | 91 | 67 | 82 | 57 | 47 | 26 | 69 | 0 | -1 | 20 | 20 | 20 | 13 | 17 | -1 | 27 | 18 | -1 | -1 | 18 | 11 | 25 | 17 | 18 | 21 | 29 | 16 | 13 | 9 | 11 | 7 | 20 | 37 | 14 | 21 | 21 | -1 |
| DENND4C | chr9:19337601-19337672 | 0 | 0 | 1 | 0 | 0 | 0 | 0 | 0 | 16 | 16 | 14 | 21 | 21 | 15 | 11 | 24 | 21 | 8 | 19 | 31 | 27 | 1 | 3 | 0 | 0 | 0 | 0 | 0 | 1 | 0 | 1 | 1 | 1 | 0 | 0 | 1 | 0 | 0 | 0 | 1 | 1 | 0 | 0 | 0 | 1 | 1 | 14 | 1 | 0 | 0 | 0 |
| KDELC1 | chr13:102791994-102792136 | 7 | 6 | 10 | 11 | 9 | 13 | 23 | 11 | 31 | 47 | 32 | 53 | 60 | 53 | 49 | 26 | 38 | 51 | 32 | 19 | 25 | 18 | 17 | 12 | 13 | 12 | 10 | 12 | 17 | 8 | 11 | 14 | 9 | 7 | 13 | 15 | 21 | 19 | 14 | 27 | 20 | 11 | 12 | 12 | 6 | 14 | 27 | 9 | 15 | 14 | 8 |
| TJAP1 | chr6:43502588-43502617 | 2 | 1 | 1 | 1 | 1 | 1 | 1 | 2 | 78 | 73 | 74 | 94 | 94 | 79 | 77 | 84 | 80 | 68 | 82 | 90 | 86 | 2 | 0 | 2 | 1 | 1 | 1 | 1 | 1 | 21 | 24 | 2 | 1 | 1 | 1 | 22 | 9 | 1 | 0 | 6 | 1 | 2 | 3 | 1 | 1 | 2 | 53 | 1 | 1 | 1 | 0 |
| CCDC7 | chr10:32447108-32447167 | 3 | 4 | 3 | 5 | 7 | 6 | 0 | 3 | 6 | 14 | 14 | 46 | 43 | 11 | 18 | 11 | 11 | 17 | 16 | 6 | 8 | -1 | -1 | 4 | 4 | 6 | 4 | 6 | -1 | 10 | 15 | 5 | 1 | 5 | 2 | 23 | 6 | 9 | 3 | 7 | 5 | 1 | 2 | 4 | 3 | 3 | 0 | 4 | 6 | 5 | 2 |
| GIT1 | chr17:29578962-29578988 | 7 | 8 | 10 | 8 | 7 | 7 | 10 | 8 | 70 | 85 | 74 | 89 | 87 | 84 | 86 | 79 | 64 | 78 | 73 | 15 | 52 | 11 | 6 | 10 | 9 | 8 | 7 | 9 | 8 | 7 | 8 | 7 | 10 | 8 | 6 | 9 | 6 | 5 | 9 | 61 | 8 | 5 | 6 | 8 | 5 | 8 | 14 | 6 | 7 | 7 | 3 |
| PQLC1 | chr18:79930228\|79923912-79930311 | 8 | 10 | 8 | 18 | 15 | 18 | 16 | 9 | 55 | 62 | 44 | 47 | 47 | 61 | 62 | 64 | 40 | 50 | 45 | 22 | 34 | 13 | 11 | 18 | 11 | 18 | 7 | 17 | 19 | 10 | 8 | 8 | 6 | 11 | 9 | 14 | 13 | 13 | 7 | 13 | 14 | 8 | 7 | 12 | 9 | 13 | 4 | 9 | 18 | 11 | 10 |
| HMGXB3 | chr5:150021809-150021972 | 5 | 7 | 4 | 6 | 6 | 5 | 6 | 6 | 6 | 10 | 8 | 19 | 20 | 7 | 8 | 6 | 10 | 7 | 6 | 10 | 6 | 6 | 7 | 8 | 5 | 7 | 5 | 7 | 2 | 8 | 7 | 6 | 8 | 7 | 5 | 9 | 7 | 4 | 5 | 11 | 6 | 5 | 4 | 7 | 7 | 5 | 6 | 5 | 7 | 6 | 5 |
| LRRC39 | chr1:100171445-100171826 | 5 | 5 | 1 | 13 | 4 | 13 | -1 | 4 | 28 | 22 | 25 | 33 | 33 | 29 | 16 | 22 | 26 | 29 | 18 | 16 | 15 | -1 | 0 | 9 | 6 | 7 | 4 | 8 | -1 | 1 | 1 | 5 | 4 | 5 | 1 | 1 | 7 | 3 | 3 | 6 | 3 | 3 | 4 | 3 | 3 | 5 | 6 | 3 | 6 | 7 | 3 |
| SCYL2 | chr12:100317681-100317692 | 1 | 1 | 4 | 0 | 0 | 0 | 0 | 0 | 37 | 68 | 48 | 88 | 88 | 72 | 75 | 49 | 64 | 58 | 48 | 6 | 40 | 0 | 0 | 3 | 3 | 1 | 0 | 1 | 0 | 1 | 0 | 0 | 0 | 1 | 0 | 0 | 0 | 1 | 2 | 61 | 0 | 0 | 0 | 3 | 2 | 3 | 1 | 0 | 0 | 1 | 2 |
| POU2F2 | chr19:42116709-42117005 | 1 | 0 | 1 | 2 | 1 | 1 | 0 | 2 | 29 | 54 | 42 | 93 | 94 | 62 | 59 | 22 | 58 | 54 | 42 | 8 | 15 | 0 | -1 | 2 | 2 | 1 | 2 | 3 | 0 | 2 | 2 | 4 | 0 | 3 | 1 | 3 | 2 | 1 | 0 | 5 | 1 | 3 | 5 | 1 | 1 | 1 | 65 | 1 | 1 | 1 | 2 |
| ZFYVE21 | chr14:103729793-103729846 | 1 | 1 | 8 | 0 | 1 | 0 | 2 | 2 | 51 | 58 | 49 | 79 | 75 | 63 | 66 | 51 | 51 | 52 | 52 | 41 | 47 | 2 | 0 | 2 | 4 | 1 | 5 | 1 | 1 | 4 | 5 | 5 | 4 | 2 | 4 | 29 | 5 | 2 | 4 | 24 | 5 | 6 | 5 | 3 | 3 | 3 | 39 | 4 | 1 | 3 | 5 |
| ALG14 | chr1:95041493-95041612 | 2 | 3 | 3 | 5 | 6 | 5 | 7 | 3 | 16 | 15 | 14 | 21 | 24 | 16 | 16 | 15 | 14 | 17 | 13 | 12 | 12 | 5 | 2 | 8 | 3 | 6 | 3 | 7 | 5 | 5 | 5 | 3 | 2 | 5 | 3 | 5 | 10 | 9 | 2 | 7 | 5 | 3 | 2 | 3 | 3 | 3 | 7 | 5 | 9 | 7 | 3 |
| MVD | chr16:88660520\|88659079-88662167 | 9 | 8 | 8 | 7 | 7 | 6 | 5 | 8 | 25 | 26 | 29 | 37 | 37 | 25 | 26 | 26 | 19 | 35 | 30 | 12 | 17 | 5 | 8 | 7 | 4 | 7 | 4 | 6 | 5 | 10 | 8 | 7 | 6 | 8 | 6 | 22 | 13 | 6 | 5 | 16 | 8 | 6 | 6 | 4 | 6 | 4 | 6 | 9 | 6 | 5 | 6 |
| CADPS | chr3:62481723-62481869 | 8 | 8 | 5 | 5 | 16 | 4 | -1 | 4 | 30 | 60 | 52 | 64 | 61 | 68 | 70 | 49 | 26 | 35 | 55 | 36 | 72 | 0 | -1 | 13 | 14 | 9 | 6 | 9 | 0 | 23 | 6 | -1 | -1 | 10 | 3 | 24 | 6 | 4 | 3 | 6 | 11 | 3 | 9 | 12 | 6 | 5 | 2 | 17 | -1 | 7 | -1 |
| TRIM9 | chr14:50982942-50986144 | 6 | 9 | 11 | 6 | 5 | 14 | -1 | 5 | 13 | 34 | 13 | 62 | 61 | 43 | 44 | 25 | 21 | 22 | 15 | 9 | 18 | -1 | -1 | 12 | 10 | 8 | 7 | 10 | 0 | 5 | 6 | -1 | 2 | 12 | 5 | 32 | 15 | 4 | 7 | 15 | 7 | 5 | 16 | 12 | 2 | 8 | 2 | 5 | 3 | 5 | 10 |
| EPRS | chr1:220018126-220018146 | 0 | 0 | 2 | 0 | 0 | 0 | 1 | 0 | 11 | 32 | 18 | 63 | 62 | 33 | 41 | 18 | 36 | 25 | 18 | 6 | 17 | 1 | 0 | 1 | 1 | 0 | 0 | 1 | 0 | 0 | 0 | 0 | 0 | 0 | 0 | 1 | 0 | 0 | 1 | 44 | 0 | 0 | 0 | 0 | 0 | 0 | 1 | 1 | 0 | 0 | 0 |
| PHC1 | chr12:8915111-8916069 | 1 | 1 | 26 | 0 | 2 | 1 | 0 | 2 | 72 | 88 | 83 | 97 | 98 | 94 | 89 | 78 | 90 | 92 | 89 | 76 | 81 | 0 | 0 | 14 | 14 | 3 | 3 | 4 | 0 | 31 | 33 | 0 | -1 | 1 | 3 | 12 | 4 | 1 | 8 | 47 | 2 | 1 | 9 | 11 | 3 | 10 | 1 | 1 | 1 | 3 | 2 |
| CTNND1 | chr11:57806461-57806478 | 0 | 0 | 1 | 1 | 1 | 1 | 3 | 0 | 21 | 36 | 18 | 76 | 78 | 43 | 53 | 22 | 34 | 25 | 19 | 19 | 21 | 0 | 1 | 5 | 1 | 4 | 0 | 5 | 1 | 28 | 33 | 1 | 0 | 0 | 0 | 36 | 3 | 1 | 0 | 4 | 1 | 0 | 0 | 1 | 1 | 1 | 10 | 0 | 1 | 0 | 1 |
| ZNF304 | chr19:57355271-57355784\|57355411 | 7 | 9 | 3 | 6 | 5 | 5 | 4 | 6 | 9 | 14 | 7 | 23 | 28 | 12 | 8 | 11 | 9 | 14 | 14 | 5 | 9 | -1 | -1 | 7 | 7 | 8 | 3 | 9 | 2 | 3 | 3 | 5 | 5 | 8 | 4 | 3 | 10 | 4 | 6 | 10 | 4 | 4 | 3 | 8 | 9 | 5 | 9 | 4 | 6 | 5 | 6 |
| EPB41L3 | chr18:5398021-5398143\|5401043 | 2 | 3 | 4 | 3 | 3 | 3 | 4 | 2 | 45 | 80 | 49 | 93 | 94 | 80 | 89 | 48 | 70 | 55 | 47 | 55 | 53 | 2 | 3 | 14 | 10 | 9 | 1 | 11 | 4 | 7 | 5 | 2 | 4 | 3 | 3 | 62 | 3 | 3 | 5 | 11 | 4 | 3 | 3 | 4 | 3 | 11 | 26 | 3 | 4 | 4 | 3 |
| RP11-20I23.1 | chr16:2497710-2497727 | 1 | 3 | 4 | 5 | 5 | 5 | 3 | 1 | 36 | 62 | 39 | 96 | 96 | 65 | 75 | 56 | 66 | 45 | 33 | 32 | 45 | -1 | 7 | 6 | 4 | 3 | 2 | 3 | 4 | 15 | 19 | 1 | 5 | 1 | 1 | 65 | 3 | 3 | 5 | 12 | 3 | 3 | 4 | 3 | 1 | 3 | 1 | 1 | 4 | 4 | 1 |
| ANGEL2 | chr1:213011208-213011458 | 1 | 1 | 1 | 1 | 1 | 1 | 2 | 1 | 10 | 15 | 11 | 31 | 35 | 18 | 16 | 13 | 20 | 14 | 10 | 14 | 10 | 1 | 3 | 2 | 1 | 2 | 1 | 3 | 4 | 6 | 6 | 0 | 0 | 1 | 2 | 22 | 2 | 3 | 2 | 9 | 3 | 0 | 0 | 1 | 1 | 2 | 9 | 1 | 2 | 2 | 1 |
| N4BP2L2 | chr13:32477887-32478048 | 6 | 6 | 6 | 6 | 7 | 6 | 5 | 8 | 7 | 8 | 11 | 28 | 28 | 10 | 9 | 7 | 8 | 9 | 8 | 6 | 5 | -1 | 11 | 7 | 8 | 6 | 7 | 5 | 7 | 7 | 6 | 5 | 7 | 6 | 6 | 6 | 6 | 13 | 6 | 11 | 8 | 7 | 8 | 8 | 8 | 6 | 14 | 5 | 11 | 10 | 8 |
| EI24 | chr11:125583196-125583321 | 3 | 3 | 3 | 2 | 3 | 2 | 2 | 4 | 27 | 29 | 21 | 28 | 39 | 28 | 16 | 22 | 19 | 23 | 22 | 14 | 15 | 3 | 0 | 3 | 2 | 3 | 1 | 3 | 2 | 4 | 5 | 2 | 1 | 3 | 2 | 25 | 13 | 3 | 1 | 6 | 2 | 2 | 1 | 3 | 4 | 2 | 3 | 2 | 2 | 1 | 8 |
| EWSR1 | chr22:29274265-29274282 | 0 | 0 | 2 | 1 | 1 | 1 | 2 | 0 | 56 | 71 | 67 | 74 | 84 | 74 | 61 | 57 | 61 | 74 | 67 | 23 | 43 | 2 | 0 | 4 | 2 | 1 | 0 | 2 | 0 | 40 | 21 | 0 | 0 | 0 | 4 | 43 | 1 | 0 | 0 | 7 | 1 | 0 | 1 | 2 | 0 | 2 | 0 | 1 | 0 | 1 | 0 |
| UHRF1BP1L | chr12:100128580-100128725 | 15 | 14 | 14 | 13 | 14 | 14 | 9 | 13 | 34 | 32 | 30 | 47 | 39 | 28 | 31 | 36 | 33 | 34 | 31 | 28 | 35 | 11 | 10 | 16 | 14 | 15 | 11 | 17 | 8 | 15 | 12 | 19 | 11 | 12 | 14 | 14 | 16 | 11 | 12 | 21 | 14 | 12 | 11 | 12 | 10 | 14 | 22 | 15 | 13 | 12 | 7 |
| EMC1 | chr1:19234180-19234188 | 0 | 0 | 3 | 0 | 0 | 0 | 0 | 0 | 19 | 56 | 25 | 91 | 88 | 54 | 71 | 28 | 60 | 40 | 26 | 6 | 29 | 0 | 0 | 1 | 1 | 0 | 0 | 0 | 0 | 4 | 8 | 1 | 0 | 0 | 0 | 7 | 0 | 0 | 3 | 63 | 0 | 0 | 0 | 1 | 0 | 1 | 1 | 0 | 0 | 1 | 0 |
| VPS13D | chr1:12338231-12338305 | 7 | 9 | 9 | 8 | 10 | 9 | 10 | 6 | 43 | 62 | 46 | 91 | 85 | 65 | 71 | 53 | 64 | 54 | 49 | 64 | 48 | 10 | 6 | 10 | 10 | 9 | 9 | 10 | 5 | 27 | 28 | 15 | 15 | 8 | 8 | 13 | 12 | 6 | 14 | 14 | 10 | 6 | 6 | 9 | 7 | 11 | 56 | 11 | 7 | 9 | 6 |
| C14orf37 | chr14:58293066\|58292623-58293138 | 4 | 7 | 10 | 7 | 8 | 7 | 7 | 6 | 16 | 23 | 22 | 39 | 36 | 25 | 29 | 16 | 15 | 30 | 21 | 5 | 6 | 7 | 6 | 10 | 11 | 10 | 9 | 9 | 10 | 4 | 4 | 10 | 10 | 8 | 9 | 4 | 4 | 7 | 12 | 22 | 10 | 5 | 4 | 11 | 11 | 11 | 11 | 13 | 8 | 8 | 9 |
| KIF2A | chr5:62372438-62372551 | 0 | 0 | 4 | 0 | 0 | 0 | 0 | 0 | 23 | 48 | 23 | 77 | 66 | 43 | 63 | 31 | 47 | 36 | 23 | 22 | 28 | 0 | 0 | 2 | 2 | 1 | 1 | 1 | 0 | 1 | 1 | 0 | 1 | 0 | 0 | 2 | 0 | 0 | 2 | 48 | 1 | 0 | 0 | 1 | 0 | 1 | 2 | 0 | 0 | 1 | 0 |
| PARP6 | chr15:72249245-72249314 | 7 | 9 | 9 | 5 | 6 | 4 | 6 | 6 | 67 | 82 | 74 | 94 | 93 | 85 | 87 | 69 | 70 | 78 | 72 | 29 | 53 | 4 | 4 | 6 | 6 | 6 | 5 | 5 | 7 | 6 | 9 | 4 | 7 | 8 | 5 | 7 | 8 | 3 | 8 | 69 | 3 | 5 | 4 | 7 | 7 | 7 | 1 | 3 | 5 | 5 | 13 |
| PHKB | chr16:47633899\|47633372-47633981 | 8 | 4 | 1 | 3 | 2 | 5 | 3 | 8 | 3 | 4 | 3 | 28 | 32 | 7 | 5 | 4 | 5 | 4 | 3 | 4 | 3 | 9 | 8 | 4 | 3 | 3 | 3 | 3 | 7 | 2 | 2 | 3 | 3 | 6 | 3 | 4 | 7 | 5 | 7 | 5 | 4 | 11 | 13 | 9 | 8 | 5 | 16 | 3 | 6 | 5 | 6 |
| TXNRD3 | chr3:126617498-126617629\|126617586 | 9 | 13 | 8 | 11 | 12 | 12 | 11 | 10 | 23 | 32 | 28 | 65 | 70 | 40 | 38 | 27 | 24 | 27 | 23 | 20 | 19 | 16 | 13 | 22 | 18 | 17 | 8 | 16 | 12 | 24 | 32 | 6 | 11 | 12 | 9 | 40 | 19 | 17 | 11 | 10 | 15 | 10 | 6 | 16 | 11 | 13 | 6 | 9 | 20 | 16 | 11 |
| CASP2 | chr7:143300380-143300440 | 18 | 14 | 10 | 16 | 15 | 16 | 19 | 16 | 26 | 32 | 30 | 66 | 65 | 39 | 37 | 34 | 36 | 34 | 31 | 42 | 40 | 14 | 12 | 18 | 14 | 17 | 10 | 18 | 17 | 22 | 22 | 13 | 19 | 17 | 14 | 42 | 25 | 15 | 14 | 14 | 15 | 13 | 14 | 17 | 15 | 15 | 32 | 14 | 17 | 13 | 19 |
| GS1-57L11.1 | chr8:2799588-2799883 | 4 | 5 | 1 | 3 | -1 | 2 | 0 | 3 | 7 | 21 | 12 | 26 | 27 | 26 | 25 | 10 | 13 | 12 | 5 | 8 | -1 | -1 | 0 | 2 | 1 | 4 | 2 | 3 | 0 | 0 | 0 | 0 | 0 | 3 | 0 | 1 | 1 | 2 | -1 | 5 | 0 | 6 | 5 | 6 | 3 | 6 | 15 | 4 | 2 | 3 | 6 |
| FBLN7 | chr2:112153003-112153258 | 2 | 2 | 6 | 1 | 2 | 2 | 1 | 2 | 21 | 17 | 15 | 25 | 30 | 19 | 16 | 14 | 15 | 14 | 12 | 11 | 15 | 2 | 3 | 2 | 3 | 3 | 3 | 3 | 0 | 2 | 2 | 0 | 6 | 3 | 4 | 1 | 1 | 4 | 11 | 19 | 3 | 2 | 2 | 3 | 5 | 3 | 6 | 7 | 3 | 2 | 6 |
| FSBP | chr8:94405474-94405692 | 5 | 10 | 7 | 8 | 9 | 7 | 11 | 5 | 11 | 15 | 12 | 45 | 39 | 22 | 24 | 13 | 16 | 14 | 16 | 12 | 12 | -1 | -1 | 8 | 5 | 5 | 7 | 7 | 4 | 8 | 6 | 7 | 23 | 4 | 6 | 8 | 6 | 7 | 9 | 14 | 10 | 4 | 3 | 3 | 6 | 7 | 11 | 7 | 8 | 10 | 4 |
| XRN1 | chr3:142376697-142376795 | 12 | 10 | 6 | 10 | 9 | 12 | 9 | 14 | 18 | 25 | 17 | 34 | 47 | 24 | 18 | 27 | 14 | 23 | 17 | 29 | 21 | 8 | -1 | 12 | 9 | 10 | 6 | 8 | 18 | 13 | 9 | 7 | 7 | 10 | 10 | 11 | 16 | 23 | 11 | 24 | 11 | 12 | 12 | 9 | 10 | 10 | 12 | 12 | 18 | 15 | 8 |
| MBD2 | chr18:54187714-54187837\|54187854 | 7 | 8 | 6 | 10 | 9 | 13 | 11 | 9 | 13 | 13 | 12 | 33 | 34 | 17 | 13 | 13 | 15 | 14 | 12 | 13 | 12 | 10 | 15 | 15 | 9 | 12 | 5 | 12 | 12 | 9 | 8 | 7 | 9 | 9 | 6 | 11 | 12 | 14 | 11 | 15 | 13 | 8 | 7 | 9 | 8 | 10 | 14 | 9 | 18 | 10 | 4 |
| SNAP23 | chr15:42521448\|42521614-42521708 | 0 | 0 | 1 | 0 | 0 | 1 | 1 | 1 | 5 | 13 | 6 | 65 | 67 | 19 | 20 | 6 | 10 | 11 | 7 | 2 | 3 | 1 | 1 | 2 | 1 | 2 | 1 | 2 | 1 | 16 | 22 | 1 | 1 | 1 | 2 | 38 | 1 | 2 | 1 | 24 | 2 | 1 | 1 | 2 | 1 | 1 | 9 | 2 | 1 | 1 | 1 |
| RBBP6 | chr16:24558421-24558534 | 0 | 0 | 0 | 0 | 1 | 0 | 0 | 0 | 5 | 16 | 13 | 49 | 47 | 25 | 30 | 7 | 16 | 17 | 13 | 5 | 4 | 0 | 1 | 0 | 0 | 0 | 0 | 0 | 0 | 5 | 16 | 0 | 0 | 0 | 1 | 33 | 0 | 0 | 0 | 1 | 0 | 0 | 0 | 0 | 0 | 0 | 3 | 0 | 0 | 0 | 1 |

SFig5, pg5

| Gene Symbol | Exon Location (hg38, human) | Adipose - Subcutaneous | Adipose - Visceral (Omentum) | Adrenal Gland | Artery - Aorta | Artery - Coronary | Artery - Tibial | Bladder | Breast - Mammary Tissue | Brain - Amygdala | Brain - Anterior cingulate cortex (BA24) | Brain - Caudate (basal ganglia) | Brain - Cerebellar Hemisphere | Brain - Cerebellum | Brain - Cortex | Brain - Frontal Cortex (BA9) | Brain - Hippocampus | Brain - Hypothalamus | Brain - Nucleus accumbens (basal ganglia) | Brain - Putamen (basal ganglia) | Brain - Spinal cord (cervical c-1) | Brain - Substantia nigra | Cervix - Ectocervix | Cervix - Endocervix | Colon - Sigmoid | Colon - Transverse | Esophagus - Gastroesophageal Junction | Esophagus - Mucosa | Esophagus - Muscularis | Fallopian Tube | Heart - Atrial Appendage | Heart - Left Ventricle | Kidney - Cortex | Liver | Lung | Minor Salivary Gland | Muscle - Skeletal | Nerve - Tibial | Ovary | Pancreas | Pituitary | Prostate | Skin - Not Sun Exposed (Suprapubic) | Skin - Sun Exposed (Lower leg) | Small Intestine - Terminal Ileum | Spleen | Stomach | Testis | Thyroid | Uterus | Vagina | Whole Blood |
|---|---|---|---|---|---|---|---|---|---|---|---|---|---|---|---|---|---|---|---|---|---|---|---|---|---|---|---|---|---|---|---|---|---|---|---|---|---|---|---|---|---|---|---|---|---|---|---|---|---|---|---|---|
| WDR60 | chr7:158905166-158905283 | 4 | 9 | 9 | 8 | 8 | 7 | 8 | 6 | 21 | 20 | 25 | 27 | 32 | 20 | 15 | 20 | 20 | 25 | 22 | 13 | 14 | 2 | 8 | 12 | 8 | 10 | 6 | 10 | 15 | 12 | 9 | 9 | 13 | 7 | 4 | 10 | 10 | 10 | 12 | 12 | 9 | 4 | 2 | 7 | 5 | 7 | 4 | 7 | 9 | 10 | 4 |
| TAF4B | chr18:26388111-26388169 | 13 | 18 | 14 | 14 | 16 | 8 | -1 | 17 | 20 | 43 | 29 | 38 | 50 | 47 | 42 | 28 | 26 | 26 | 37 | 11 | 16 | 0 | 0 | 10 | 14 | 11 | 11 | 10 | -1 | 8 | 6 | 10 | -1 | 10 | 12 | 6 | 12 | 10 | 16 | 17 | 10 | 17 | 9 | 13 | 10 | 13 | 6 | 8 | 8 | 9 | 5 |
| LRRC16A | chr6:25612377\|25612740\|25612737-25612886 | 15 | 12 | 6 | 11 | 10 | 14 | 15 | 16 | 38 | 42 | 37 | 52 | 45 | 44 | 49 | 37 | 38 | 38 | 37 | 27 | 32 | 14 | 9 | 10 | 5 | 12 | 8 | 13 | 22 | 13 | 10 | 8 | 4 | 8 | 10 | 15 | 17 | 11 | 6 | 23 | 11 | 15 | 17 | 8 | 10 | 7 | 18 | 12 | 12 | 10 | 11 |
| TNRC6A | chr16:24804705-24804851 | 0 | 0 | 0 | 0 | 1 | 0 | 0 | 0 | 17 | 35 | 15 | 84 | 82 | 44 | 45 | 24 | 44 | 13 | 14 | 32 | 27 | 0 | 0 | 1 | 1 | 0 | 1 | 0 | 0 | 17 | 19 | 1 | 1 | 0 | 2 | 56 | 0 | 0 | 0 | 3 | 1 | 1 | 1 | 0 | 0 | 1 | 22 | 0 | 0 | 0 | 1 |
| PLEKHH1 | chr14:67586810\|67586368-67586949 | 4 | 3 | 4 | 5 | 4 | 6 | 5 | 3 | 9 | 11 | 9 | 22 | 22 | 11 | 12 | 10 | 9 | 9 | 9 | 8 | 9 | -1 | 4 | 4 | 2 | 4 | 2 | 4 | 4 | 8 | 6 | 3 | 2 | 3 | 3 | 14 | 6 | 6 | 2 | 4 | 3 | 4 | 4 | 3 | 3 | 4 | 3 | 4 | 8 | 4 | 7 |
| XYLB | chr3:38373115-38373279 | 2 | 1 | 1 | 1 | 1 | 1 | 3 | 1 | 19 | 14 | 16 | 20 | 23 | 22 | 10 | 18 | 12 | 14 | 14 | 29 | 16 | 0 | -1 | 2 | 0 | 2 | 0 | 3 | 0 | 5 | 10 | 0 | 1 | 1 | 1 | 14 | 5 | 1 | 0 | 3 | 1 | 0 | 1 | 0 | 1 | 1 | 1 | 0 | 1 | 1 | 0 |
| HDAC7 | chr12:47790206-47790463\|47791146 | 3 | 3 | 2 | 3 | 2 | 2 | 2 | 3 | 24 | 39 | 30 | 79 | 77 | 54 | 49 | 32 | 44 | 46 | 29 | 36 | 25 | 2 | 3 | 2 | 2 | 2 | 3 | 2 | 2 | 25 | 33 | 4 | 3 | 3 | 3 | 47 | 7 | 2 | 3 | 4 | 3 | 3 | 4 | 2 | 3 | 2 | 8 | 3 | 2 | 2 | 5 |
| TBCD | chr17:82929866-82930243 | 3 | 3 | 1 | 4 | 3 | 3 | 3 | 3 | 1 | 1 | 1 | 19 | 22 | 2 | 1 | 2 | 1 | 1 | 1 | 1 | 1 | 7 | 9 | 5 | 3 | 3 | 2 | 3 | 4 | 3 | 2 | 4 | 2 | 2 | 2 | 8 | 4 | 9 | 2 | 10 | 5 | 2 | 2 | 3 | 3 | 2 | 4 | 4 | 9 | 6 | 3 |
| RP11-2C24.9 | chr16:30722563-30722748 | 4 | 3 | 5 | 2 | 3 | 3 | 5 | 4 | 26 | 47 | 24 | 64 | 62 | 45 | 55 | 29 | 42 | 34 | 27 | 23 | 27 | 1 | 3 | 5 | 4 | 5 | 4 | 6 | 3 | 23 | 30 | 6 | 6 | 3 | 5 | 24 | 3 | 2 | 3 | 5 | 4 | 4 | 5 | 4 | 3 | 5 | 29 | 6 | 2 | 3 | 3 |
| CACNA1D | chr3:53802147-53802173 | 7 | 5 | 9 | 3 | 0 | 13 | 0 | 2 | 71 | 84 | 79 | 98 | 97 | 92 | 93 | 64 | 78 | 91 | 76 | 30 | 46 | -1 | 1 | 15 | 8 | 7 | 1 | 5 | 6 | 2 | 12 | 3 | 2 | 2 | 1 | 12 | 3 | 4 | 22 | 66 | 2 | 4 | 4 | 6 | 1 | 26 | 17 | 1 | 4 | 7 | 12 |
| CAMTA2 | chr17:4969500-4969520 | 1 | 1 | 4 | 1 | 1 | 1 | 2 | 2 | 75 | 87 | 76 | 97 | 95 | 89 | 91 | 82 | 71 | 77 | 73 | 25 | 46 | 1 | 2 | 3 | 2 | 2 | 1 | 1 | 1 | 1 | 0 | 1 | 1 | 3 | 1 | 2 | 1 | 1 | 5 | 74 | 2 | 3 | 3 | 3 | 1 | 3 | 2 | 1 | 1 | 2 | 6 |
| KIF1B | chr1:10326111-10326359 | 14 | 10 | 7 | 8 | 8 | 7 | 10 | 12 | 94 | 96 | 92 | 98 | 98 | 96 | 97 | 91 | 92 | 95 | 90 | 87 | 90 | -1 | -1 | 19 | 12 | 13 | 4 | 13 | 5 | 23 | 25 | 5 | 7 | 9 | 4 | 56 | 31 | 15 | 3 | 42 | 9 | 5 | 6 | 9 | 4 | 9 | 10 | 4 | 11 | 9 | 4 |
| CALD1 | chr7:134962820-134962934 | 0 | 0 | 0 | 1 | 0 | 0 | 0 | 0 | 16 | 17 | 10 | 52 | 70 | 19 | 20 | 18 | 18 | 15 | 11 | 16 | 13 | 0 | 0 | 0 | 0 | 0 | 0 | 0 | 0 | 43 | 34 | 0 | 0 | 0 | 0 | 5 | 1 | 0 | 0 | 4 | 1 | 0 | 0 | 0 | 0 | 0 | 1 | 0 | 0 | 0 | 2 |
| MAP3K9 | chr14:70735961-70736029 | 18 | 13 | 10 | 18 | -1 | 17 | -1 | 3 | 81 | 91 | 76 | 79 | 80 | 91 | 97 | 83 | 80 | 80 | 85 | 73 | 82 | -1 | -1 | 18 | 6 | 14 | 17 | 26 | 0 | 21 | 26 | 5 | 13 | 19 | 9 | 43 | 14 | 12 | 7 | 12 | 4 | 7 | 9 | 13 | 7 | 7 | 2 | 5 | 14 | 8 | 4 |
| SLC15A2 | chr3:121931126-121931209 | 4 | 5 | 4 | 10 | 7 | 8 | 4 | 6 | 3 | 4 | 3 | 35 | 28 | 6 | 5 | 5 | 6 | 3 | 3 | 4 | 4 | -1 | 6 | 10 | 6 | 10 | 2 | 9 | 9 | 6 | 7 | 7 | 4 | 6 | 4 | 4 | 5 | 12 | 4 | 9 | 5 | 4 | 5 | 7 | 4 | 6 | 3 | 5 | 13 | 4 | 6 |
| RFT1 | chr3:53118621-53118803 | 2 | 4 | 3 | 3 | 3 | 3 | 3 | 4 | 10 | 11 | 8 | 16 | 22 | 9 | 7 | 6 | 8 | 8 | 7 | 7 | 8 | 7 | 6 | 4 | 3 | 4 | 3 | 3 | 7 | 10 | 9 | 6 | 4 | 3 | 4 | 7 | 4 | 10 | 4 | 8 | 8 | 5 | 3 | 5 | 6 | 4 | 6 | 6 | 7 | 6 | 3 |
| GS1-259H13.13 | chr7:99563188-99563266\|99563413 | 12 | 8 | 6 | 15 | 13 | 17 | 15 | 12 | 33 | 30 | 28 | 64 | 66 | 40 | 35 | 36 | 36 | 26 | 32 | 44 | 41 | 14 | 12 | 18 | 9 | 17 | 6 | 19 | 16 | 32 | 26 | 9 | 4 | 8 | 8 | 26 | 22 | 15 | 9 | 21 | 17 | 8 | 8 | 7 | 5 | 9 | 27 | 10 | 18 | 10 | 3 |
| DZIP1 | chr13:95630714-95630813 | 10 | 9 | 12 | 16 | 14 | 16 | 14 | 9 | 17 | 20 | 23 | 51 | 52 | 25 | 26 | 20 | 13 | 16 | 30 | 23 | 21 | 13 | 8 | 14 | 13 | 16 | 10 | 23 | 8 | 17 | 23 | 6 | 10 | 12 | 8 | 27 | 17 | 6 | 9 | 24 | 14 | 12 | 12 | 15 | 12 | 17 | 18 | 9 | 8 | 9 | 9 |
| TBC1D2 | chr9:98213785\|98214083-98220253 | 4 | 8 | 8 | 5 | 5 | 4 | 5 | 14 | 42 | 31 | 36 | 46 | 48 | 43 | 36 | 42 | 44 | 37 | 32 | 60 | 50 | 6 | 5 | 5 | 4 | 5 | 3 | 6 | 6 | 9 | 9 | 4 | 26 | 8 | 9 | 25 | 5 | 4 | 4 | 12 | 9 | 4 | 4 | 6 | 14 | 4 | 15 | 4 | 5 | 3 | 10 |
| TBC1D2 | chr9:98213785\|98214083-98220253 | 4 | 8 | 8 | 5 | 5 | 4 | 5 | 14 | 42 | 31 | 36 | 46 | 48 | 43 | 36 | 42 | 44 | 37 | 32 | 60 | 50 | 6 | 5 | 5 | 4 | 5 | 3 | 6 | 6 | 9 | 9 | 4 | 26 | 8 | 9 | 25 | 5 | 4 | 4 | 12 | 9 | 4 | 4 | 6 | 14 | 4 | 15 | 4 | 5 | 3 | 10 |
| MYH9 | chr22:36308808-36308870 | 0 | 0 | 0 | 0 | 0 | 0 | 0 | 0 | 2 | 12 | 3 | 26 | 31 | 12 | 17 | 7 | 1 | 5 | 3 | 0 | 1 | 0 | 0 | 0 | 0 | 0 | 0 | 0 | 0 | 10 | 21 | 0 | 0 | 0 | 0 | 9 | 0 | 0 | 0 | 0 | 0 | 0 | 0 | 0 | 0 | 0 | 0 | 0 | 0 | 0 | 0 |
| ZNF350 | chr19:51968003\|51967258-51968071 | 2 | 4 | 3 | 3 | 3 | 3 | 2 | 2 | 10 | 11 | 12 | 24 | 25 | 15 | 12 | 10 | 9 | 12 | 16 | 9 | 10 | 2 | 0 | 3 | 3 | 3 | 3 | 3 | 2 | 6 | 7 | 4 | 4 | 2 | 3 | 19 | 2 | 3 | 4 | 3 | 4 | 2 | 1 | 3 | 2 | 3 | 2 | 2 | 2 | 3 | 4 |
| AP1S2 | chrX:15828192-15828200\|15831568\|15833524 | 3 | 2 | 11 | 3 | 3 | 3 | 3 | 3 | 63 | 70 | 51 | 90 | 87 | 66 | 78 | 45 | 77 | 75 | 50 | 21 | 45 | 2 | 2 | 4 | 5 | 3 | 9 | 3 | 2 | 2 | 1 | 8 | 9 | 3 | 7 | 1 | 1 | 2 | 22 | 65 | 3 | 7 | 8 | 5 | 3 | 5 | 11 | 10 | 2 | 4 | 1 |
| ZDHHC21 | chr9:14688247-14688307 | 1 | 1 | 1 | 2 | 2 | 2 | 1 | 1 | 35 | 53 | 27 | 54 | 53 | 51 | 59 | 40 | 33 | 30 | 26 | 19 | 28 | 3 | -1 | 3 | 2 | 2 | 1 | 2 | -1 | 30 | 28 | 0 | 1 | 1 | 2 | 17 | 5 | 2 | 1 | 4 | 2 | 1 | 1 | 2 | 2 | 2 | 5 | 1 | 3 | 1 | 2 |
| CTC-326K19.6 | chr19:49911105-49911366 | 1 | 1 | 1 | 1 | 1 | 2 | 4 | 1 | 5 | 12 | 7 | 31 | 33 | 14 | 14 | 6 | 7 | 8 | 8 | 6 | 7 | 3 | 1 | 3 | 2 | 3 | 1 | 3 | 1 | 12 | 14 | 1 | 1 | 1 | 1 | 16 | 2 | 2 | 1 | 3 | 2 | 1 | 0 | 1 | 0 | 1 | 14 | 1 | 2 | 1 | 1 |
| TLK1 | chr2:171022872-171023129\|171022997 | 14 | 14 | 7 | 12 | 9 | 12 | 10 | 15 | 9 | 7 | 11 | 37 | 44 | 12 | 8 | 12 | 8 | 10 | 10 | 14 | 11 | 14 | 19 | 10 | 9 | 10 | 10 | 9 | 14 | 12 | 10 | 9 | 15 | 16 | 10 | 30 | 18 | 17 | 11 | 14 | 12 | 16 | 15 | 11 | 13 | 10 | 16 | 11 | 17 | 16 | 17 |
| NVL | chr1:224313050\|224313034-224313155 | 7 | 9 | 7 | 9 | 9 | 8 | 9 | 9 | 14 | 13 | 10 | 20 | 24 | 13 | 13 | 11 | 12 | 15 | 11 | 16 | 9 | 6 | 7 | 12 | 9 | 11 | 7 | 11 | 9 | 11 | 11 | 7 | 9 | 7 | 6 | 12 | 9 | 10 | 8 | 16 | 8 | 8 | 6 | 9 | 7 | 7 | 9 | 7 | 11 | 9 | 6 |
| TMEM184B | chr22:38222642-38222662 | 3 | 3 | 3 | 7 | 8 | 11 | 13 | 2 | 24 | 42 | 24 | 73 | 68 | 48 | 54 | 26 | 42 | 28 | 21 | 7 | 17 | 5 | 6 | 22 | 13 | 18 | 4 | 19 | 4 | 25 | 27 | 1 | 2 | 1 | 2 | 36 | 7 | 4 | 1 | 4 | 9 | 2 | 2 | 8 | 2 | 9 | 5 | 1 | 5 | 5 | 2 |
| PIEZO2 | chr18:10774006-10780366 | 8 | 4 | 15 | 9 | 19 | 9 | -1 | 13 | 94 | 91 | 97 | 89 | 92 | 93 | 92 | 97 | 94 | 92 | 93 | 95 | 95 | -1 | -1 | 10 | 11 | 15 | 8 | 15 | 0 | 6 | -1 | 2 | 3 | 3 | -1 | 43 | 53 | 19 | 3 | 5 | 12 | 11 | 8 | 10 | 4 | 12 | 29 | 11 | 4 | 14 | -1 |
| MANBA | chr4:102658859-102659028 | 1 | 1 | 0 | 1 | 1 | 1 | 1 | 1 | 4 | 11 | 5 | 27 | 33 | 17 | 15 | 6 | 7 | 7 | 8 | 4 | 4 | 1 | 2 | 1 | 0 | 1 | 0 | 0 | 1 | 10 | 12 | 1 | 1 | 1 | 1 | 24 | 1 | 1 | 0 | 1 | 1 | 1 | 1 | 0 | 1 | 0 | 2 | 1 | 1 | 1 | 0 |
| ZCCHC7 | chr9:37127095-37127263 | 2 | 3 | 3 | 4 | 4 | 3 | 2 | 2 | 14 | 22 | 16 | 19 | 27 | 20 | 17 | 12 | 20 | 16 | 17 | 13 | 13 | -1 | 0 | 4 | 3 | 5 | 3 | 4 | 0 | 8 | 9 | 4 | 6 | 3 | 2 | 19 | 3 | 4 | 1 | 9 | 4 | 1 | 1 | 2 | 3 | 2 | 5 | 3 | 4 | 4 | 3 |
| ATP6V0A1 | chr17:42508572-42508589\|42510183 | 1 | 1 | 6 | 1 | 1 | 1 | 1 | 1 | 47 | 77 | 51 | 93 | 90 | 79 | 87 | 56 | 72 | 68 | 50 | 8 | 40 | 1 | 0 | 8 | 5 | 3 | 1 | 3 | 1 | 1 | 1 | 1 | 1 | 1 | 1 | 1 | 1 | 1 | 10 | 75 | 2 | 1 | 1 | 4 | 1 | 5 | 1 | 1 | 1 | 3 | 1 |
| ANK2 | chr4:113237599-113237622 | 1 | 1 | 3 | 0 | 1 | 1 | 0 | 2 | 38 | 60 | 43 | 86 | 70 | 63 | 71 | 54 | 52 | 52 | 48 | 37 | 54 | 0 | 0 | 3 | 5 | 2 | 3 | 2 | -1 | 1 | 1 | 1 | 0 | 1 | 1 | 4 | 3 | 0 | 10 | 57 | 2 | 1 | 4 | 4 | 0 | 5 | 1 | 0 | 0 | 1 | -1 |
| RIMKLB | chr12:8699785-8700614\|8700605 | 6 | 4 | 4 | 6 | 5 | 5 | 5 | 5 | 9 | 17 | 8 | 51 | 50 | 20 | 24 | 13 | 18 | 11 | 9 | 10 | 10 | 5 | 3 | 3 | 4 | 4 | 4 | 3 | 6 | 9 | 15 | 4 | 5 | 4 | 7 | 40 | 6 | 4 | 3 | 6 | 3 | 6 | 8 | 3 | 5 | 4 | 12 | 5 | 6 | 4 | 3 |
| FBXW11 | chr5:171957597-171957698 | 5 | 3 | 6 | 5 | 4 | 6 | 4 | 5 | 10 | 18 | 11 | 27 | 23 | 14 | 21 | 12 | 22 | 10 | 12 | 12 | 13 | 7 | 8 | 5 | 4 | 5 | 4 | 4 | 5 | 15 | 13 | 6 | 2 | 3 | 5 | 7 | 4 | 6 | 5 | 9 | 5 | 5 | 6 | 4 | 3 | 6 | 6 | 6 | 7 | 7 | 2 |
| MARCH8 | chr10:45459740-45459877 | 13 | 12 | 3 | 10 | 8 | 12 | 7 | 16 | 15 | 19 | 12 | 48 | 47 | 22 | 23 | 16 | 27 | 17 | 14 | 11 | 12 | 16 | 15 | 6 | 4 | 6 | 5 | 4 | 7 | 11 | 5 | 6 | 5 | 11 | 11 | 11 | 21 | 20 | 6 | 13 | 14 | 17 | 20 | 5 | 6 | 5 | 9 | 9 | 16 | 19 | 6 |
| HOOK2 | chr19:12765829-12765834 | 4 | 2 | 1 | 4 | 3 | 4 | 3 | 3 | 2 | 4 | 3 | 24 | 20 | 8 | 6 | 2 | 5 | 5 | 2 | 2 | 3 | 5 | 3 | 2 | 1 | 2 | 2 | 3 | 2 | 2 | 0 | 1 | 1 | 4 | 2 | 1 | 3 | 3 | 1 | 15 | 1 | 4 | 4 | 1 | 1 | 1 | 0 | 1 | 2 | 2 | 4 |
| MGRN1 | chr16:4686259-4686303 | 14 | 17 | 21 | 9 | 12 | 9 | 11 | 16 | 37 | 45 | 35 | 48 | 47 | 45 | 44 | 39 | 37 | 29 | 34 | 31 | 36 | 12 | 10 | 10 | 14 | 12 | 13 | 12 | 17 | 23 | 27 | 22 | 20 | 17 | 14 | 12 | 19 | 17 | 17 | 17 | 15 | 9 | 9 | 17 | 17 | 18 | 22 | 19 | 14 | 12 | 15 |
| TRRAP | chr7:98945747-98945950 | 2 | 3 | 4 | 4 | 4 | 3 | 3 | 3 | 19 | 33 | 27 | 80 | 78 | 35 | 44 | 21 | 38 | 42 | 29 | 11 | 11 | 0 | 4 | 4 | 3 | 4 | 2 | 4 | 5 | 10 | 11 | 2 | 6 | 2 | 3 | 34 | 6 | 3 | 4 | 57 | 3 | 3 | 2 | 3 | 2 | 3 | 5 | 2 | 3 | 3 | 2 |
| MEAF6 | chr1:37496707-37496736\|37496942 | 0 | 0 | 6 | 0 | 1 | 0 | 1 | 0 | 67 | 83 | 65 | 91 | 89 | 80 | 88 | 65 | 82 | 79 | 67 | 21 | 58 | 1 | 1 | 3 | 3 | 2 | 0 | 2 | 2 | 12 | 16 | 0 | 0 | 1 | 1 | 14 | 0 | 1 | 5 | 73 | 1 | 0 | 1 | 2 | 1 | 3 | 1 | 1 | 1 | 1 | 1 |
| NIN | chr14:50730923-50730983 | 0 | 0 | 1 | 1 | 1 | 1 | 1 | 0 | 20 | 52 | 26 | 89 | 92 | 58 | 58 | 34 | 32 | 34 | 32 | 6 | 12 | 1 | 0 | 2 | 2 | 1 | 0 | 2 | 0 | 48 | 40 | 0 | 0 | 0 | 1 | 49 | 0 | 0 | 1 | 7 | 1 | 0 | 0 | 0 | 0 | 1 | 1 | 0 | 2 | 0 | 0 |
| KCNN1 | chr19:17982568-17982576 | 3 | 1 | 1 | 2 | 0 | 6 | -1 | 2 | 8 | 11 | 10 | 29 | 31 | 13 | 13 | 9 | 11 | 10 | 11 | 23 | 13 | -1 | 0 | 11 | 7 | 3 | 1 | 11 | 0 | 3 | 12 | 10 | -1 | 4 | -1 | 5 | 2 | 1 | 4 | 6 | 2 | 1 | 5 | 12 | 0 | 3 | 0 | 3 | 1 | 6 | 9 |
| IL16 | chr15:81266654-81266801 | 6 | 5 | 5 | 5 | 6 | 8 | 3 | 7 | -1 | -1 | 10 | 23 | 24 | 23 | 10 | 2 | 11 | 21 | -1 | 17 | -1 | 5 | -1 | 5 | 5 | 5 | 5 | 4 | 4 | 5 | 5 | -1 | -1 | 5 | 5 | 7 | 9 | 8 | 4 | 10 | 5 | 5 | 6 | 5 | 3 | 5 | 11 | 5 | 8 | 6 | 9 |
| GULP1 | chr2:188569620\|188569625-188569643 | 1 | 0 | 0 | 0 | 0 | 0 | 0 | 1 | 12 | 22 | 22 | 14 | 18 | 22 | 20 | 10 | 7 | 14 | 18 | 2 | 10 | 1 | 2 | 0 | 0 | 0 | 0 | 0 | 0 | 0 | 1 | 1 | 15 | 0 | 0 | 0 | 0 | 1 | 1 | 1 | 1 | 1 | 0 | 0 | 0 | 0 | 0 | 0 | 0 | 3 | 2 |
| ANK1 | chr8:41700431-41700454 | 1 | 5 | 0 | 1 | 1 | 1 | 0 | 5 | 87 | 98 | 93 | 99 | 99 | 97 | 97 | 93 | 94 | 89 | 96 | 90 | 98 | 0 | 0 | 9 | 5 | 3 | 2 | 3 | 0 | 54 | 59 | 0 | -1 | 1 | 5 | 17 | 4 | 0 | -1 | 4 | 2 | 7 | 28 | 5 | 1 | 3 | 6 | 2 | -1 | -1 | 0 |
| EPB41L3 | chr18:5407701\|5407437-5407736 | 0 | 0 | 5 | 0 | 0 | 0 | 0 | 0 | 43 | 86 | 58 | 90 | 96 | 84 | 91 | 44 | 66 | 60 | 46 | 7 | 33 | 0 | 0 | 9 | 6 | 7 | 0 | 10 | 0 | 9 | 4 | 0 | 1 | 0 | 2 | 55 | 0 | 0 | 21 | 58 | 1 | 1 | 2 | 3 | 0 | 10 | 0 | 1 | 0 | 3 | 1 |
| EIF4A1 | chr17:7573400-7573547 | 3 | 7 | 9 | 9 | 7 | 4 | 13 | 6 | 18 | 21 | 21 | 20 | 39 | 25 | 11 | 18 | 21 | 29 | 22 | 11 | 14 | 12 | 16 | 9 | 7 | 7 | 6 | 7 | 9 | 7 | 3 | 13 | 9 | 4 | 8 | 3 | 12 | 12 | 4 | 23 | 15 | 7 | 4 | 6 | 6 | 7 | 6 | 7 | 12 | 12 | 1 |
| PAICS | chr4:56446357-56446377 | 1 | 1 | 0 | 0 | 1 | 0 | 0 | 1 | 1 | 7 | 2 | 31 | 40 | 7 | 6 | 3 | 9 | 2 | 3 | 3 | 8 | 0 | 0 | 1 | 0 | 0 | 0 | 0 | 1 | 0 | 0 | 0 | 0 | 0 | 1 | 35 | 1 | 0 | 0 | 3 | 0 | 1 | 0 | 1 | 0 | 0 | 1 | 0 | 0 | 0 | 1 |
| USP28 | chr11:113828844-113828920 | 11 | 4 | 2 | 10 | 8 | 11 | 7 | 13 | 17 | 27 | 16 | 35 | 38 | 34 | 39 | 16 | 18 | 19 | 18 | 10 | 13 | 14 | 8 | 6 | 7 | 5 | 3 | 4 | 21 | 7 | 3 | 4 | 3 | 6 | 5 | 2 | 13 | 8 | 5 | 11 | 7 | 11 | 10 | 13 | 8 | 7 | 19 | 6 | 9 | 8 | 5 |
| PHLDB1 | chr11:118644647-118644691 | 13 | 8 | 7 | 28 | 21 | 24 | 15 | 10 | 3 | 6 | 3 | 52 | 52 | 5 | 7 | 2 | 8 | 6 | 2 | 2 | 2 | 19 | 19 | 17 | 11 | 16 | 13 | 13 | 13 | 16 | 15 | 9 | 11 | 11 | 15 | 25 | 9 | 6 | 7 | 8 | 17 | 12 | 16 | 9 | 8 | 13 | 7 | 11 | 23 | 20 | 9 |
| LRRC6 | chr8:132643374\|132643624-132643686 | 3 | 3 | 1 | 3 | 1 | 2 | 4 | 3 | 17 | 26 | 16 | 22 | 23 | 37 | 33 | 17 | 12 | 24 | 22 | 7 | 13 | -1 | -1 | 2 | 3 | 3 | 1 | 2 | 2 | 12 | 14 | 3 | 0 | 2 | 2 | 9 | 3 | 3 | 2 | 4 | 3 | 2 | 4 | 4 | 4 | 4 | 3 | 1 | 2 | 3 | 4 |
| PPFIA1 | chr11:70333022-70333096 | 1 | 1 | 0 | 4 | 5 | 10 | 20 | 1 | 5 | 13 | 6 | 66 | 63 | 21 | 28 | 7 | 12 | 8 | 6 | 4 | 6 | 2 | 7 | 16 | 8 | 17 | 2 | 22 | 4 | 25 | 35 | 0 | 0 | 0 | 1 | 6 | 0 | 1 | 0 | 0 | 6 | 0 | 0 | 2 | 1 | 6 | 0 | 0 | 6 | 3 | 0 |
| PAK3 | chrX:111146521-111146565 | 3 | 2 | 8 | 2 | 2 | 1 | 0 | 2 | 41 | 46 | 41 | 14 | 19 | 39 | 44 | 37 | 40 | 43 | 34 | -1 | 29 | 0 | 0 | 10 | 7 | 0 | 2 | 2 | -1 | -1 | 0 | -1 | 0 | 3 | 0 | 3 | 5 | 2 | 1 | 4 | 2 | 2 | 2 | -1 | 1 | 1 | 3 | 1 | 1 | 2 | -1 |
| CMC1 | chr3:28318732\|28316753-28318879 | 2 | 2 | 1 | 2 | 2 | 2 | 2 | 3 | 8 | 13 | 8 | 24 | 24 | 13 | 12 | 10 | 12 | 8 | 8 | 12 | 11 | 3 | 4 | 2 | 2 | 2 | 1 | 2 | 3 | 7 | 8 | 2 | 2 | 3 | 2 | 12 | 4 | 2 | 2 | 2 | 3 | 2 | 2 | 4 | 5 | 2 | 16 | 2 | 2 | 2 | 4 |
| AP2A1 | chr19:49802534-49802599 | 1 | 1 | 5 | 0 | 1 | 1 | 2 | 2 | 57 | 77 | 45 | 93 | 91 | 81 | 82 | 67 | 72 | 47 | 47 | 56 | 63 | 1 | 1 | 4 | 3 | 2 | 1 | 2 | 2 | 44 | 59 | 3 | 0 | 1 | 4 | 37 | 3 | 1 | 9 | 6 | 3 | 1 | 1 | 3 | 0 | 2 | 3 | 1 | 1 | 1 | 1 |
| MAPK14 | chr6:36100188-36100267 | 13 | 16 | 26 | 9 | 10 | 7 | 12 | 14 | 40 | 59 | 42 | 66 | 66 | 59 | 64 | 44 | 58 | 53 | 40 | 31 | 38 | 12 | 13 | 10 | 19 | 11 | 22 | 10 | 19 | 8 | 8 | 24 | 14 | 17 | 24 | 3 | 12 | 18 | 24 | 40 | 15 | 17 | 17 | 19 | 26 | 15 | 30 | 21 | 11 | 17 | 26 |
| SPAG1 | chr8:100223662-100223753 | 0 | 0 | 0 | 0 | 0 | 0 | 0 | 0 | 4 | 12 | 8 | 23 | 27 | 17 | 17 | 6 | 3 | 12 | 14 | 3 | 3 | 0 | 0 | 0 | 0 | 0 | 0 | 0 | 0 | 14 | 19 | 0 | 0 | 0 | 0 | 3 | 0 | 0 | 0 | 0 | 0 | 0 | 0 | 0 | 0 | 0 | 2 | 0 | 0 | 0 | 0 |
| VEZF1 | chr17:57979845-57979967 | 6 | 8 | 5 | 9 | 7 | 7 | 13 | 9 | 9 | 7 | 6 | 17 | 24 | 5 | 4 | 7 | 7 | 6 | 6 | 4 | 7 | 6 | 9 | 12 | 9 | 12 | 5 | 10 | 13 | 8 | 6 | 3 | 3 | 5 | 6 | 5 | 7 | 9 | 4 | 10 | 9 | 9 | 6 | 7 | 6 | 8 | 8 | 5 | 10 | 8 | 7 |
| ITGB7 | chr12:53206674-53206804 | 3 | 1 | 0 | 4 | 3 | 7 | 0 | 4 | -1 | 0 | 6 | 15 | 22 | 6 | 4 | 0 | 3 | 10 | -1 | 2 | 2 | 0 | 0 | 3 | 1 | 2 | 2 | 5 | 0 | 1 | 4 | 1 | 0 | 1 | 2 | 5 | 10 | 4 | 1 | 2 | 2 | 7 | 12 | 0 | 1 | 0 | 7 | 1 | 3 | 3 | 0 |
| BIN1 | chr2:127068163\|127064774-127068255 | 1 | 1 | 3 | 1 | 1 | 1 | 1 | 1 | 25 | 56 | 23 | 83 | 86 | 52 | 59 | 23 | 37 | 43 | 17 | 8 | 16 | 1 | 1 | 4 | 5 | 2 | 1 | 2 | 1 | 2 | 1 | 6 | 1 | 2 | 2 | 0 | 1 | 1 | 4 | 43 | 4 | 1 | 3 | 6 | 2 | 5 | 65 | 7 | 1 | 1 | 2 |
| L1CAM | chrX:153873228-153873242 | 0 | 1 | 68 | 1 | 0 | 1 | 3 | 2 | 88 | 98 | 80 | 99 | 99 | 96 | 98 | 93 | 96 | 93 | 70 | 81 | 94 | 0 | 0 | 5 | 9 | 2 | 0 | 2 | 0 | 0 | 0 | 0 | 5 | 3 | 0 | 5 | 0 | 0 | 5 | 57 | 0 | 1 | 5 | 11 | 1 | 6 | 2 | 0 | 0 | 6 | 7 |
| CA5B | chrX:15768643-15769511\|15769293 | 6 | 6 | 4 | 4 | 4 | 4 | 5 | 8 | 7 | 10 | 9 | 39 | 39 | 13 | 12 | 11 | 9 | 9 | 10 | 13 | 7 | 7 | 11 | 8 | 10 | 7 | 8 | 8 | 6 | 7 | 15 | 8 | 8 | 9 | 8 | 32 | 12 | 7 | 13 | 11 | 10 | 9 | 9 | 9 | 10 | 11 | 6 | 8 | 10 | 7 | 6 |
| SLC38A6 | chr14:60987052-60987157 | 4 | 5 | 4 | 10 | 9 | 12 | 13 | 6 | 42 | 56 | 53 | 87 | 87 | 67 | 69 | 47 | 55 | 64 | 53 | 38 | 44 | 17 | 21 | 15 | 8 | 12 | 6 | 14 | 9 | 30 | 32 | 5 | 4 | 5 | 4 | 63 | 12 | 12 | 5 | 20 | 12 | 5 | 4 | 7 | 6 | 9 | 28 | 7 | 16 | 10 | 2 |
| NRXN3 | chr14:79692173-79692262 | 12 | 17 | 14 | 21 | 12 | 18 | 25 | 17 | 79 | 61 | 92 | 95 | 94 | 65 | 59 | 59 | 91 | 92 | 87 | 81 | 87 | -1 | -1 | 20 | 22 | 17 | 14 | 15 | 27 | 11 | 13 | 13 | -1 | 19 | 17 | 67 | 6 | 16 | 19 | 31 | 11 | 15 | 23 | 21 | 16 | 28 | 25 | 10 | 26 | 13 | -1 |
| DOCK3 | chr3:51333001-51333027 | 1 | 6 | 10 | 1 | 3 | 2 | 2 | 5 | 41 | 80 | 55 | 93 | 92 | 75 | 88 | 42 | 63 | 68 | 52 | 6 | 23 | 0 | -1 | 7 | 6 | 3 | 3 | 2 | 0 | 1 | 2 | -1 | 0 | 5 | 2 | 7 | 1 | 3 | 45 | 70 | 2 | 1 | 5 | 12 | 5 | 6 | 4 | 2 | 2 | 6 | 10 |
| NMNAT3 | chr3:139572107-139572233 | 1 | 1 | 2 | 2 | 2 | 1 | 1 | 1 | 5 | 11 | 6 | 23 | 26 | 15 | 15 | 11 | 5 | 7 | 7 | 1 | 1 | 3 | 2 | 2 | 1 | 2 | 2 | 2 | 3 | 2 | 3 | 1 | 5 | 2 | 1 | 5 | 2 | 2 | 2 | 3 | 3 | 1 | 0 | 3 | 2 | 2 | 23 | 2 | 2 | 2 | 1 |
| ANKS6 | chr9:98784380-98784500 | 1 | 1 | 1 | 1 | 1 | 2 | 1 | 1 | 22 | 49 | 25 | 81 | 82 | 60 | 61 | 25 | 39 | 27 | 23 | 16 | 28 | 0 | 4 | 4 | 3 | 5 | 1 | 6 | 1 | 34 | 46 | 1 | 1 | 2 | 3 | 49 | 1 | 1 | 2 | 5 | 3 | 1 | 3 | 2 | 1 | 3 | 19 | 1 | 2 | 2 | 3 |
| PTCH1 | chr9:95474012-95474143 | 3 | 3 | 1 | 4 | 3 | 5 | 6 | 3 | 23 | 28 | 21 | 91 | 82 | 36 | 35 | 37 | 28 | 21 | 26 | 50 | 25 | 0 | 4 | 4 | 3 | 5 | 3 | 6 | 1 | 19 | 43 | 2 | 2 | 2 | 2 | 61 | 4 | 2 | 2 | 3 | 4 | 2 | 3 | 3 | 1 | 3 | 5 | 2 | 4 | 3 | 2 |
| CFP | chrX:47628476-47628553 | 3 | 3 | 9 | 5 | 4 | 9 | -1 | 6 | 21 | 26 | 35 | 63 | 69 | 49 | 39 | 35 | 43 | 38 | 35 | 14 | 26 | -1 | -1 | 6 | 4 | 6 | 7 | 6 | 6 | 5 | 7 | 6 | 8 | 5 | 8 | 9 | 7 | 9 | 7 | 37 | 18 | 4 | 4 | 6 | 4 | 6 | 51 | 12 | 9 | 11 | 3 |
| CERCAM | chr9:128421204-128422786\|128421511\|128422354\|128421345 | 10 | 9 | 6 | 5 | 5 | 6 | 12 | 18 | 88 | 80 | 90 | 70 | 70 | 89 | 82 | 91 | 82 | 87 | 91 | 84 | 87 | 13 | 10 | 14 | 12 | 11 | 19 | 12 | 9 | 4 | 6 | 20 | 8 | 7 | 12 | 22 | 15 | 7 | 9 | 57 | 12 | 13 | 9 | 12 | 20 | 10 | 23 | 12 | 13 | 13 | 14 |
| AFTPH | chr2:64581190\|64580622-64581273 | 5 | 6 | 12 | 5 | 6 | 5 | 8 | 9 | 40 | 64 | 46 | 72 | 70 | 64 | 74 | 47 | 52 | 54 | 43 | 20 | 31 | 9 | 11 | 8 | 11 | 8 | 8 | 7 | 15 | 7 | 8 | 9 | 4 | 6 | 14 | 24 | 5 | 10 | 17 | 49 | 18 | 7 | 7 | 10 | 9 | 12 | 41 | 10 | 7 | 8 | 6 |
| COL11A1 | chr1:102979597-102979612 | 1 | 8 | 3 | 2 | 1 | 2 | -1 | 5 | 16 | 43 | 32 | 29 | 31 | 63 | 67 | 22 | 60 | 47 | 32 | 13 | 44 | 0 | 0 | 11 | 15 | 6 | -1 | 11 | 0 | -1 | -1 | -1 | 0 | 8 | 1 | 3 | 4 | 6 | 6 | 6 | 2 | 1 | 2 | 8 | 4 | 5 | 9 | 13 | 7 | 7 | 0 |
| MVD | chr16:88660520-88662167\|88662781\|88660838 | 13 | 11 | 11 | 10 | 10 | 8 | 7 | 10 | 29 | 30 | 33 | 38 | 37 | 29 | 30 | 30 | 24 | 40 | 33 | 14 | 19 | 6 | 9 | 10 | 5 | 9 | 6 | 9 | 6 | 14 | 13 | 9 | 7 | 10 | 8 | 28 | 17 | 8 | 7 | 19 | 9 | 9 | 9 | 6 | 8 | 6 | 8 | 11 | 8 | 6 | 6 |
| TBC1D9B | chr5:179867445\|179867176-179867495 | 10 | 10 | 8 | 8 | 9 | 9 | 12 | 10 | 52 | 64 | 49 | 90 | 89 | 69 | 73 | 65 | 67 | 57 | 53 | 60 | 60 | 9 | 10 | 16 | 10 | 12 | 7 | 13 | 9 | 27 | 41 | 9 | 14 | 8 | 8 | 73 | 23 | 7 | 8 | 14 | 10 | 9 | 9 | 9 | 9 | 10 | 17 | 7 | 11 | 11 | 7 |
| PABPC1L | chr20:44934501-44934563 | 5 | 3 | 1 | 2 | 2 | 2 | 1 | 1 | 24 | 28 | 14 | 32 | 30 | 30 | 31 | 22 | 43 | 20 | 15 | 10 | 17 | 1 | 1 | 4 | 2 | 3 | 1 | 4 | 1 | 6 | 10 | 0 | 1 | 2 | 1 | 24 | 12 | 1 | 1 | 2 | 1 | 0 | 1 | 1 | 2 | 1 | 5 | 0 | 1 | 1 | 1 |
| G3BP2 | chr4:75653983-75654081 | 8 | 9 | 15 | 16 | 17 | 17 | 18 | 9 | 52 | 74 | 54 | 50 | 50 | 66 | 75 | 54 | 65 | 69 | 55 | 31 | 49 | 14 | 14 | 19 | 14 | 17 | 9 | 17 | 15 | 20 | 26 | 12 | 12 | 9 | 9 | 34 | 12 | 12 | 12 | 18 | 13 | 7 | 7 | 11 | 8 | 14 | 20 | 13 | 17 | 11 | 4 |
| RP11-365O16.6 | chr1:224223414-224223497 | 1 | 2 | 3 | 1 | 2 | 1 | -1 | 1 | 38 | -1 | 36 | 62 | 54 | 43 | 32 | -1 | 42 | 47 | 45 | -1 | -1 | -1 | 0 | 3 | 4 | 4 | 1 | 5 | -1 | 33 | 15 | -1 | 0 | 1 | -1 | 36 | 4 | 2 | 0 | 2 | 1 | 1 | 1 | 4 | 1 | 2 | 0 | 1 | 1 | 1 | 1 |
| NVL | chr1:224313050\|224313034-224313151 | 8 | 11 | 8 | 11 | 11 | 9 | 12 | 10 | 15 | 15 | 12 | 21 | 25 | 14 | 14 | 12 | 15 | 17 | 12 | 16 | 12 | 10 | 7 | 14 | 10 | 13 | 8 | 13 | 11 | 15 | 13 | 7 | 10 | 9 | 7 | 13 | 11 | 11 | 9 | 17 | 9 | 9 | 6 | 10 | 9 | 9 | 11 | 9 | 13 | 11 | 7 |
| FRYL | chr4:48523488-48523496 | 0 | 0 | 1 | 0 | 0 | 0 | 0 | 0 | 1 | 8 | 2 | 56 | 54 | 5 | 9 | 2 | 10 | 2 | 1 | 1 | 2 | 0 | 0 | 0 | 0 | 0 | 0 | 0 | 0 | 0 | 0 | 0 | 0 | 0 | 0 | 0 | 0 | 0 | 4 | 51 | 0 | 0 | 0 | 0 | 1 | 0 | 0 | 0 | 0 | 0 | 0 |
| DZIP1 | chr13:95636980\|95636905-95637176 | 4 | 3 | 3 | 4 | 3 | 3 | 2 | 3 | 4 | 7 | 4 | 22 | 20 | 9 | 9 | 6 | 4 | 2 | 4 | 8 | 7 | 5 | 3 | 4 | 4 | 4 | 4 | 4 | 5 | 5 | 5 | 5 | 6 | 4 | 4 | 18 | 4 | 3 | 4 | 4 | 4 | 6 | 6 | 4 | 7 | 4 | 2 | 4 | 4 | 4 | 5 |
| ZNF554 | chr19:2833163-2833269 | 3 | 2 | 2 | 3 | 3 | 3 | 1 | 2 | 30 | 38 | 33 | 46 | 43 | 37 | 42 | 32 | 37 | 37 | 29 | 13 | 23 | 5 | 3 | 4 | 2 | 4 | 1 | 4 | 1 | 20 | 20 | 2 | 2 | 2 | 2 | 31 | 3 | 2 | 2 | 4 | 3 | 2 | 2 | 2 | 2 | 2 | 9 | 2 | 3 | 2 | 3 |
| MYCBP2 | chr13:77129156-77129230 | 17 | 17 | 16 | 18 | 19 | 20 | 21 | 18 | 25 | 41 | 25 | 54 | 49 | 45 | 50 | 30 | 23 | 26 | 28 | 26 | 30 | 17 | 17 | 20 | 21 | 23 | 14 | 25 | 27 | 21 | 25 | 12 | 14 | 12 | 17 | 35 | 18 | 15 | 15 | 11 | 15 | 13 | 12 | 16 | 13 | 20 | 13 | 14 | 18 | 15 | 10 |
| FBLN5 | chr14:91939850-91939950 | 3 | 3 | 2 | 3 | 3 | 3 | 3 | 3 | 21 | 17 | 23 | 4 | 7 | 21 | 18 | 10 | 12 | 28 | 25 | 6 | 12 | 2 | 3 | 4 | 4 | 3 | 3 | 4 | 3 | 2 | 3 | 4 | 2 | 3 | 3 | 4 | 6 | 3 | 3 | 4 | 4 | 3 | 3 | 3 | 3 | 3 | 2 | 3 | 4 | 3 | 4 |
| GRB7 | chr17:39738573-39738924\|39740181 | 3 | 2 | -1 | 4 | 2 | 4 | 3 | 2 | 0 | -1 | 18 | 18 | 18 | 25 | 29 | -1 | -1 | -1 | 12 | -1 | 0 | 2 | 3 | 3 | 2 | 4 | 3 | 4 | 3 | 2 | 5 | 4 | 1 | 1 | 3 | 10 | 3 | 3 | 2 | 3 | 2 | 5 | 5 | 3 | 13 | 2 | 5 | 2 | 4 | 3 | 1 |
| AK2 | chr1:33031544-33031661 | 33 | 30 | 18 | 27 | 30 | 27 | 27 | 33 | 61 | 62 | 54 | 71 | 78 | 66 | 61 | 63 | 65 | 56 | 60 | 70 | 65 | 22 | 21 | 28 | 16 | 29 | 9 | 29 | 29 | 34 | 30 | 21 | 11 | 20 | 17 | 37 | 45 | 33 | 11 | 40 | 27 | 19 | 17 | 16 | 17 | 18 | 22 | 21 | 29 | 18 | 11 |
| AP1B1 | chr22:29339754-29339774 | 1 | 1 | 1 | 0 | 1 | 0 | 0 | 1 | 55 | 74 | 61 | 87 | 86 | 77 | 78 | 63 | 72 | 68 | 64 | 49 | 61 | 1 | 0 | 1 | 1 | 1 | 0 | 1 | 1 | 39 | 49 | 1 | 0 | 1 | 1 | 35 | 9 | 0 | 0 | 2 | 0 | 0 | 0 | 1 | 1 | 1 | 45 | 0 | 0 | 1 | 1 |
| NTRK3 | chr15:87885694-87885735 | 15 | 15 | 10 | 17 | 17 | 13 | -1 | 17 | 59 | 57 | 45 | 53 | 55 | 57 | 56 | 50 | 40 | 43 | 45 | 39 | 40 | -1 | -1 | 18 | 10 | 16 | 15 | 15 | -1 | 21 | 10 | -1 | -1 | 12 | 17 | 10 | 24 | 13 | -1 | 9 | 12 | 15 | 11 | 17 | 10 | 16 | 3 | 13 | 9 | 12 | -1 |
| DGKI | chr7:137609879-137610001 | 0 | 0 | 1 | 0 | 0 | 0 | 0 | 0 | 9 | 13 | 21 | 2 | 2 | 6 | 7 | 9 | 3 | 34 | 16 | 0 | 2 | 0 | -1 | 0 | 0 | 0 | 0 | 0 | -1 | 0 | 0 | 0 | 0 | 0 | 0 | 0 | 0 | 0 | 0 | 0 | 0 | 0 | 0 | 0 | 0 | 0 | 0 | 0 | 0 | 0 | 0 |
| C5orf45 | chr5:179851243-179853276 | 15 | 12 | 6 | 10 | 10 | 10 | 11 | 15 | 7 | 6 | 8 | 29 | 30 | 10 | 9 | 9 | 8 | 6 | 6 | 11 | 8 | 13 | 14 | 10 | 11 | 10 | 14 | 10 | 13 | 12 | 12 | 12 | 14 | 15 | 15 | 21 | 16 | 12 | 11 | 11 | 12 | 17 | 19 | 15 | 14 | 12 | 12 | 12 | 13 | 15 | 15 |
| RECK | chr9:36066769-36066870 | 1 | 1 | 2 | 1 | 1 | 2 | 2 | 1 | 27 | 38 | 22 | 79 | 81 | 44 | 46 | 34 | 32 | 27 | 21 | 42 | 35 | 2 | 0 | 3 | 2 | 2 | 1 | 2 | 3 | 11 | 20 | 1 | 2 | 1 | 1 | 9 | 5 | 1 | 1 | 6 | 4 | 1 | 2 | 1 | 2 | 1 | 76 | 1 | 2 | 3 | 1 |
| DNAH7 | chr2:195847555-195847637 | 4 | 3 | 9 | 7 | 7 | 6 | 0 | 8 | 62 | 58 | 59 | 29 | 45 | 49 | 44 | 60 | 44 | 59 | 62 | -1 | 51 | 0 | 0 | 7 | 6 | 6 | 0 | 3 | 21 | 3 | 0 | 7 | 0 | 8 | 3 | -1 | 10 | 15 | 5 | 23 | 9 | 4 | 3 | -1 | -1 | 4 | 25 | 7 | 7 | -1 | 0 |
| SLC12A8 | chr3:125084310-125084420 | 4 | 4 | 5 | 4 | 4 | 4 | 8 | 3 | 55 | 69 | 47 | 59 | 45 | 70 | 71 | 57 | 64 | 50 | 51 | 28 | 43 | 0 | 0 | 3 | 3 | 3 | 2 | 7 | 9 | 21 | 22 | 4 | 3 | 5 | 1 | 33 | 6 | 5 | 3 | 12 | 5 | 5 | 2 | 4 | 3 | 4 | 8 | 3 | 8 | 2 | 3 |
| CYP2U1 | chr4:107935595\|107936517-107938176 | 6 | 6 | 4 | 5 | 5 | 7 | 4 | 6 | 8 | 13 | 10 | 33 | 34 | 23 | 19 | 13 | 14 | 9 | 11 | 13 | 18 | 5 | -1 | 9 | 7 | 9 | 5 | 8 | 4 | 15 | 21 | 10 | 7 | 6 | 5 | 19 | 15 | 5 | 7 | 12 | 14 | 8 | 6 | 11 | 10 | 8 | 7 | 6 | 6 | 8 | 8 |
| KYAT3 | chr1:88962932\|88964556-88964712 | 3 | 5 | 2 | 5 | 4 | 4 | 3 | 4 | 11 | 10 | 12 | 17 | 19 | 13 | 8 | 18 | 13 | 11 | 13 | 27 | 16 | 8 | 7 | 5 | 4 | 4 | 4 | 4 | 7 | 6 | 6 | 5 | 10 | 7 | 3 | 7 | 11 | 4 | 5 | 10 | 8 | 3 | 2 | 6 | 7 | 5 | 9 | 5 | 5 | 5 | 4 |
| TBC1D5 | chr3:17233685-17233750 | 9 | 7 | 6 | 5 | 6 | 5 | 8 | 7 | 18 | 36 | 15 | 37 | 36 | 32 | 42 | 19 | 28 | 22 | 11 | 6 | 14 | 2 | 1 | 7 | 7 | 5 | 7 | 5 | 7 | 6 | 8 | 6 | 7 | 7 | 8 | 14 | 6 | 3 | 6 | 5 | 5 | 9 | 9 | 6 | 6 | 6 | 33 | 5 | 4 | 5 | 6 |
| DTNB | chr2:25419515-25419535 | 3 | 3 | 3 | 12 | 8 | 9 | 0 | 1 | 39 | 44 | 42 | 64 | 60 | 55 | 52 | 41 | 23 | 44 | 40 | 28 | 21 | 11 | 4 | 8 | 5 | 7 | 1 | 4 | 5 | 28 | 32 | 1 | 0 | 2 | 0 | 39 | 5 | 5 | 0 | 10 | 1 | 1 | 1 | 2 | 1 | 1 | 5 | 0 | 16 | 4 | 4 |
| ABI2 | chr2:203396785-203396967 | 0 | 0 | 0 | 0 | 1 | 0 | 3 | 0 | 31 | 58 | 42 | 81 | 70 | 54 | 68 | 40 | 49 | 60 | 42 | 8 | 20 | 0 | 0 | 2 | 1 | 2 | 1 | 4 | 0 | 48 | 45 | 0 | 0 | 0 | 0 | 18 | 0 | 0 | 0 | 2 | 1 | 0 | 1 | 1 | 0 | 1 | 19 | 0 | 0 | 1 | 1 |
| KIAA0141 | chr5:141936854-141937014\|141936878 | 3 | 6 | 4 | 11 | 7 | 7 | 8 | 5 | 4 | 6 | 4 | 27 | 28 | 7 | 5 | 5 | 4 | 3 | 4 | 4 | 4 | 12 | 10 | 11 | 5 | 10 | 5 | 9 | 5 | 8 | 6 | 5 | 3 | 9 | 5 | 6 | 6 | 14 | 4 | 16 | 11 | 4 | 3 | 5 | 5 | 5 | 4 | 7 | 13 | 11 | 4 |
| KIFAP3 | chr1:169925787\|169923066-169925855 | 3 | 4 | 9 | 15 | 13 | 15 | 20 | 4 | 42 | 72 | 50 | 79 | 76 | 80 | 84 | 60 | 36 | 59 | 51 | 16 | 33 | 15 | 10 | 23 | 15 | 17 | 4 | 16 | 18 | 26 | 28 | 9 | 7 | 3 | 5 | 57 | 2 | 9 | 6 | 6 | 13 | 2 | 2 | 19 | 3 | 14 | 2 | 6 | 22 | 10 | 2 |
| RALGAPA2 | chr20:20598679-20598819 | 0 | 0 | 1 | 0 | 1 | 0 | 0 | 0 | 32 | 39 | 32 | 90 | 90 | 41 | 48 | 35 | 50 | 39 | 30 | 38 | 50 | 1 | 0 | 0 | 0 | 0 | 0 | 1 | 0 | 32 | 42 | 0 | 0 | 0 | 2 | 71 | 0 | 0 | 0 | 2 | 1 | 0 | 0 | 0 | 0 | 0 | 18 | 0 | 0 | 0 | 0 |
| LAS1L | chrX:65520702-65522493\|65520817\|65521287 | 16 | 15 | 15 | 20 | 19 | 25 | 28 | 15 | 55 | 62 | 61 | 83 | 85 | 71 | 69 | 53 | 57 | 68 | 63 | 36 | 44 | 19 | 20 | 35 | 21 | 35 | 9 | 37 | 31 | 30 | 36 | 11 | 8 | 11 | 11 | 54 | 22 | 30 | 10 | 19 | 27 | 7 | 8 | 17 | 9 | 18 | 25 | 14 | 29 | 18 | 4 |
| PORCN | chrX:48512835-48514141\|48512852 | 7 | 11 | 8 | 7 | 9 | 6 | 11 | 9 | 35 | 52 | 39 | 89 | 87 | 57 | 62 | 54 | 54 | 52 | 38 | 33 | 34 | 6 | 7 | 8 | 10 | 8 | 6 | 8 | 8 | 37 | 53 | 7 | 15 | 10 | 7 | 62 | 9 | 8 | 9 | 28 | 8 | 9 | 9 | 12 | 16 | 10 | 11 | 8 | 8 | 7 | 20 |
| SH3TC2 | chr5:149034391-149034520 | 2 | 3 | 0 | 3 | 5 | 6 | 0 | 3 | 80 | 90 | 89 | 85 | 72 | 86 | 83 | 89 | 85 | 89 | 84 | 90 | 89 | 0 | -1 | 11 | 4 | 5 | 0 | 3 | 0 | 51 | 49 | -1 | 0 | 2 | 0 | 6 | 6 | 0 | -1 | 6 | 4 | 1 | 2 | 4 | 0 | 2 | 2 | 4 | 7 | 1 | 2 |
| FUBP3 | chr9:130580659-130581513 | 1 | 1 | 1 | 2 | 2 | 2 | 3 | 1 | 8 | 15 | 9 | 30 | 27 | 14 | 21 | 12 | 18 | 15 | 10 | 7 | 6 | 2 | 3 | 2 | 2 | 2 | 3 | 2 | 5 | 1 | 1 | 4 | 1 | 1 | 2 | 1 | 1 | 4 | 3 | 13 | 3 | 1 | 1 | 2 | 2 | 2 | 22 | 8 | 3 | 3 | 3 |
| CELSR3 | chr3:48650275-48650289 | 3 | 2 | 45 | 22 | 23 | 41 | -1 | 3 | 68 | 85 | 80 | 95 | 95 | 90 | 91 | 65 | 82 | 88 | 77 | 22 | 50 | 0 | -1 | 44 | 17 | 25 | 4 | 33 | -1 | 14 | 18 | 2 | 0 | 1 | 3 | 28 | 3 | 10 | 8 | 33 | 16 | 2 | 7 | 5 | 1 | 13 | 0 | 1 | 27 | 11 | 2 |
| C11orf84 | chr11:63822631\|63822791-63822848 | 10 | 22 | 10 | 15 | 13 | 12 | 10 | 12 | 16 | 17 | 17 | 32 | 37 | 17 | 16 | 18 | 17 | 15 | 14 | 22 | 19 | 11 | 16 | 18 | 15 | 14 | 11 | 13 | 6 | 19 | 14 | 10 | 18 | 16 | 10 | 26 | 17 | 12 | 14 | 12 | 15 | 10 | 8 | 13 | 10 | 12 | 5 | 15 | 16 | 16 | 15 |
| INVS | chr9:100241452\|100241475-100241556 | 4 | 3 | 3 | 5 | 5 | 5 | 6 | 3 | 32 | 39 | 29 | 63 | 71 | 48 | 44 | 42 | 34 | 34 | 40 | 50 | 40 | 5 | 4 | 8 | 5 | 7 | 3 | 9 | 4 | 31 | 38 | 2 | 6 | 3 | 4 | 55 | 12 | 2 | 2 | 5 | 5 | 2 | 3 | 6 | 5 | 5 | 8 | 3 | 4 | 4 | 2 |

SFig5, pg6

| Gene Symbol | Exon Location (hg38, human) | Adipose - Subcutaneous | Adipose - Visceral (Omentum) | Adrenal Gland | Artery - Aorta | Artery - Coronary | Artery - Tibial | Bladder | Breast - Mammary Tissue | Brain - Amygdala | Brain - Anterior cingulate cortex (BA24) | Brain - Caudate (basal ganglia) | Brain - Cerebellar Hemisphere | Brain - Cerebellum | Brain - Cortex | Brain - Frontal Cortex (BA9) | Brain - Hippocampus | Brain - Hypothalamus | Brain - Nucleus accumbens (basal ganglia) | Brain - Putamen (basal ganglia) | Brain - Spinal cord (cervical c-1) | Brain - Substantia nigra | Cervix - Ectocervix | Cervix - Endocervix | Colon - Sigmoid | Colon - Transverse | Esophagus - Gastroesophageal Junction | Esophagus - Mucosa | Esophagus - Muscularis | Fallopian Tube | Heart - Atrial Appendage | Heart - Left Ventricle | Kidney - Cortex | Liver | Lung | Minor Salivary Gland | Muscle - Skeletal | Nerve - Tibial | Ovary | Pancreas | Pituitary | Prostate | Skin - Not Sun Exposed (Suprapubic) | Skin - Sun Exposed (Lower leg) | Small Intestine - Terminal Ileum | Spleen | Stomach | Testis | Thyroid | Uterus | Vagina | Whole Blood |
|---|---|---|---|---|---|---|---|---|---|---|---|---|---|---|---|---|---|---|---|---|---|---|---|---|---|---|---|---|---|---|---|---|---|---|---|---|---|---|---|---|---|---|---|---|---|---|---|---|---|---|---|---|
| HKR1 | chr19:37357962\|37357896-37358039 | 10 | 10 | 9 | 11 | 11 | 11 | 7 | 8 | 13 | 18 | 14 | 23 | 23 | 16 | 18 | 15 | 12 | 14 | 16 | 12 | 12 | 7 | 8 | 10 | 10 | 10 | 7 | 11 | 10 | 14 | 17 | 9 | 9 | 9 | 7 | 15 | 10 | 8 | 7 | 7 | 8 | 6 | 7 | 10 | 11 | 9 | 13 | 6 | 9 | 7 | 8 |
| MED17 | chr11:93794250-93794439 | 8 | 10 | 4 | 8 | 8 | 8 | 6 | 8 | 7 | 6 | 7 | 19 | 20 | 8 | 7 | 10 | 7 | 7 | 6 | 9 | 7 | 7 | 8 | 11 | 6 | 9 | 6 | 8 | 7 | 9 | 5 | 7 | 8 | 9 | 7 | 7 | 9 | 13 | 7 | 9 | 10 | 6 | 7 | 7 | 6 | 6 | 6 | 10 | 14 | 11 | 6 |
| STAG3 | chr7:100198264-100198303 | 4 | 7 | 8 | 15 | 13 | 7 | 0 | 4 | -1 | -1 | 27 | 49 | 35 | 35 | 25 | 27 | 11 | 20 | -1 | 29 | -1 | -1 | 0 | 1 | 9 | 3 | 4 | 6 | 0 | 4 | 1 | 0 | -1 | 3 | -1 | 17 | 7 | 5 | 4 | 19 | -1 | 3 | 2 | 2 | 2 | 0 | 3 | 4 | 10 | 5 | 3 |
| CACNA1C | chr12:2597437-2597496 | 4 | 3 | 14 | 3 | 4 | 5 | 7 | 3 | 58 | 71 | 58 | 88 | 89 | 76 | 83 | 65 | 76 | 69 | 54 | 37 | 42 | -1 | 0 | 7 | 7 | 6 | 5 | 5 | 3 | 2 | 2 | 4 | 1 | 6 | 5 | 4 | 6 | 4 | 29 | 53 | 6 | 5 | 6 | 8 | 4 | 7 | 68 | 6 | 4 | 5 | -1 |
| CPOX | chr3:98521653-98521703\|98521696\|98521718 | 3 | 4 | 8 | 4 | 3 | 3 | 3 | 3 | 42 | 42 | 36 | 50 | 31 | 42 | 45 | 46 | 49 | 36 | 38 | 67 | 55 | 6 | 3 | 3 | 5 | 2 | 18 | 2 | 2 | 3 | 5 | 9 | 17 | 4 | 9 | 12 | 2 | 2 | 15 | 7 | 5 | 6 | 6 | 6 | 4 | 6 | 14 | 5 | 3 | 9 | 1 |
| COL13A1 | chr10:69898697-69898762 | 6 | 6 | 4 | 3 | 4 | 6 | 5 | 4 | 28 | 69 | 41 | 96 | 96 | 69 | 71 | 35 | 48 | 45 | 20 | 66 | 22 | 3 | 7 | 5 | 4 | 9 | 4 | 13 | 10 | 35 | 33 | 2 | 4 | 4 | 7 | 81 | 6 | 7 | 4 | 7 | 3 | 7 | 11 | 4 | 4 | 8 | 37 | 4 | 7 | 2 | 8 |
| PTPRF | chr1:43602071-43602097 | 4 | 4 | 4 | 8 | 6 | 10 | 13 | 6 | 23 | 53 | 24 | 76 | 66 | 59 | 67 | 28 | 44 | 37 | 26 | 6 | 26 | 17 | 4 | 14 | 16 | 10 | 38 | 9 | 7 | 3 | 4 | 2 | 8 | 14 | 11 | 14 | 2 | 3 | 10 | 41 | 7 | 26 | 23 | 8 | 3 | 11 | 5 | 3 | 7 | 27 | 12 |
| MORC4 | chrX:106955955-106956010 | 0 | 1 | 0 | 3 | 4 | 4 | 13 | 0 | 2 | 4 | 1 | 38 | 41 | 6 | 4 | 2 | 3 | 1 | 1 | 1 | 1 | 1 | 3 | 17 | 8 | 20 | 2 | 24 | 2 | 10 | 19 | 3 | 1 | 1 | 1 | 0 | 0 | 2 | 0 | 1 | 5 | 0 | 0 | 4 | 1 | 7 | 1 | 3 | 3 | 1 | 0 |
| RPE | chr2:210022972-210029998 | 19 | 22 | 17 | 14 | 16 | 17 | -1 | 16 | 22 | 19 | 21 | 51 | 55 | 25 | 32 | 19 | 12 | 22 | 19 | 18 | 11 | -1 | -1 | 19 | 13 | 21 | 15 | 18 | -1 | 17 | 17 | 8 | 17 | 18 | 13 | 31 | 18 | 9 | 16 | 16 | 17 | 19 | 24 | 12 | 13 | 23 | 14 | 7 | 17 | 14 | 11 |
| DIP2A | chr21:46559834-46559863 | 1 | 1 | 2 | 1 | 1 | 1 | 1 | 1 | 42 | 37 | 34 | 27 | 24 | 35 | 32 | 53 | 44 | 24 | 38 | 67 | 58 | 0 | 0 | 1 | 0 | 0 | 1 | 0 | 1 | 16 | 15 | 1 | 1 | 1 | 1 | 6 | 1 | 0 | 2 | 1 | 2 | 1 | 1 | 1 | 0 | 1 | 9 | 1 | 1 | 1 | 0 |
| MTCL1 | chr18:8828908\|8827555-8828964 | 2 | 3 | 5 | 2 | 5 | 3 | 2 | 2 | 72 | 92 | 81 | 90 | 91 | 94 | 96 | 69 | 85 | 84 | 79 | 37 | 72 | 0 | 0 | 3 | 3 | 4 | 2 | 5 | 0 | 54 | 67 | 1 | 1 | 1 | 0 | 22 | 2 | 2 | 0 | 2 | 3 | 1 | 17 | 4 | 1 | 4 | 2 | 4 | 0 | 1 | 32 |
| ALDH8A1 | chr6:134940194-134940221 | 0 | 0 | 0 | 2 | 1 | 3 | -1 | 2 | 34 | 62 | 42 | 87 | 87 | 59 | 68 | 33 | 49 | 61 | 43 | 8 | 22 | 0 | 0 | 12 | 6 | 10 | 8 | 14 | 0 | 15 | 30 | 0 | 0 | 1 | 3 | 80 | 1 | 1 | 1 | 7 | 6 | 1 | 4 | 8 | 0 | 6 | 1 | 1 | 3 | 2 | 1 |
| BRAT1 | chr7:2540771\|2540830-2540894 | 9 | 6 | 6 | 8 | 7 | 8 | 13 | 10 | 6 | 10 | 7 | 19 | 20 | 11 | 10 | 8 | 7 | 8 | 7 | 5 | 5 | 9 | 8 | 10 | 8 | 9 | 7 | 7 | 10 | 6 | 6 | 8 | 7 | 7 | 9 | 11 | 9 | 12 | 8 | 11 | 12 | 10 | 11 | 9 | 6 | 7 | 6 | 8 | 10 | 9 | 5 |
| IST1 | chr16:71918141-71918258 | 3 | 4 | 3 | 5 | 4 | 5 | 8 | 4 | 9 | 12 | 10 | 34 | 43 | 16 | 11 | 10 | 14 | 12 | 8 | 12 | 13 | 6 | 8 | 6 | 5 | 7 | 2 | 7 | 5 | 19 | 22 | 4 | 5 | 3 | 4 | 36 | 5 | 7 | 3 | 7 | 7 | 4 | 3 | 4 | 3 | 4 | 6 | 4 | 8 | 5 | 2 |
| IQGAP1 | chr15:90482702-90482749 | 3 | 2 | 2 | 2 | 2 | 2 | 4 | 3 | 30 | 28 | 28 | 72 | 78 | 44 | 34 | 38 | 35 | 32 | 37 | 48 | 42 | 3 | 2 | 5 | 3 | 4 | 1 | 4 | 3 | 11 | 18 | 1 | 3 | 2 | 2 | 77 | 6 | 2 | 1 | 5 | 4 | 2 | 2 | 4 | 3 | 3 | 13 | 2 | 3 | 2 | 1 |
| RANBP1 | chr22:20122580-20122697 | 3 | 6 | 7 | 7 | 6 | 4 | 8 | 5 | 5 | 5 | 7 | 16 | 30 | 7 | 3 | 7 | 4 | 8 | 5 | 5 | 4 | 8 | 9 | 10 | 6 | 9 | 4 | 8 | 5 | 6 | 5 | 5 | 7 | 5 | 7 | 6 | 8 | 11 | 4 | 6 | 8 | 4 | 3 | 7 | 7 | 7 | 27 | 6 | 10 | 8 | 6 |
| ATP2B1 | chr12:89598657\|89598590\|89598630\|89598592-89598743 | 0 | 0 | 0 | 0 | 0 | 0 | 1 | 0 | 62 | 89 | 89 | 87 | 86 | 92 | 94 | 78 | 62 | 90 | 89 | 28 | 47 | 0 | 1 | 1 | 0 | 0 | 0 | 1 | 0 | 18 | 36 | 0 | 1 | 0 | 3 | 80 | 1 | 0 | 0 | 3 | 1 | 0 | 1 | 0 | 0 | 0 | 1 | 0 | 0 | 0 | 0 |
| DOCK11 | chrX:118685520-118685537 | 0 | 1 | 19 | 1 | 0 | 1 | 0 | 0 | 39 | 60 | 80 | 66 | 75 | 66 | 70 | 55 | 71 | 90 | 82 | 11 | 45 | 5 | 1 | 2 | 3 | 1 | 0 | 1 | 1 | 1 | 1 | 0 | 1 | 1 | 1 | 0 | 0 | 0 | 3 | 76 | 0 | 1 | 1 | 2 | 0 | 2 | 4 | 0 | 1 | 1 | 1 |
| ARHGAP6 | chrX:11145072\|11148630-11148719 | 8 | 4 | 3 | 3 | 3 | 4 | 3 | 7 | 61 | 57 | 25 | 26 | 18 | 36 | 42 | 45 | 65 | 84 | 35 | 11 | 33 | 2 | 2 | 4 | 4 | 4 | 3 | 3 | 3 | 3 | 4 | 4 | 2 | 5 | 5 | 14 | 7 | 8 | 3 | 5 | 5 | 7 | 7 | 3 | 3 | 3 | 4 | 3 | 5 | 4 | 3 |
| SH3PXD2A | chr10:103668608-103668652\|103669720\|103673556 | 9 | 14 | 15 | 10 | 11 | 12 | 9 | 8 | 17 | 46 | 31 | 77 | 84 | 60 | 60 | 20 | 30 | 42 | 24 | 9 | 14 | 12 | -1 | 11 | 8 | 9 | 3 | 12 | 9 | 23 | 37 | -1 | 5 | 19 | 5 | 59 | 7 | 9 | 9 | 55 | 12 | 3 | 3 | 9 | 12 | 9 | 20 | 8 | 11 | 6 | 11 |
| KMT5B | chr11:68179470-68179830\|68179616\|68179537 | 9 | 11 | 16 | 11 | 10 | 7 | 5 | 11 | 13 | 16 | 14 | 35 | 42 | 19 | 17 | 16 | 18 | 17 | 15 | 15 | 16 | 10 | 11 | 10 | 9 | 12 | 8 | 10 | 11 | 21 | 20 | 16 | 17 | 12 | 10 | 22 | 10 | 12 | 11 | 28 | 16 | 6 | 6 | 15 | 23 | 13 | 17 | 17 | 16 | 11 | 8 |
| ABLIM2 | chr4:8032648-8032680 | 10 | 15 | 17 | 4 | 7 | 4 | 19 | 12 | 43 | 52 | 33 | 64 | 57 | 53 | 52 | 47 | 63 | 31 | 32 | 39 | 53 | 13 | -1 | 24 | 25 | 10 | 14 | 9 | -1 | 5 | 3 | 11 | 21 | 8 | 8 | 4 | 10 | 8 | 26 | 32 | 7 | 12 | 10 | 23 | 17 | 22 | 3 | 14 | 14 | 14 | 5 |
| RP11-582E3.4 | chr12:27145414-27145506 | 0 | 0 | 0 | 0 | 2 | 0 | 0 | 0 | -1 | 68 | 30 | 74 | 83 | 79 | 63 | 46 | 59 | 60 | -1 | -1 | -1 | 0 | 0 | 1 | 0 | 2 | 0 | 1 | 0 | 25 | 31 | 0 | 0 | 0 | 0 | 79 | 0 | 0 | 0 | 1 | 0 | 0 | 0 | 0 | 3 | 0 | 1 | 0 | 0 | 0 | 0 |
| IQSEC2 | chrX:53235783-53235832 | 9 | 8 | 8 | 9 | 9 | 8 | 7 | 9 | 76 | 88 | 80 | 87 | 85 | 91 | 90 | 84 | 68 | 87 | 82 | 35 | 51 | 5 | 9 | 9 | 7 | 9 | 5 | 8 | 18 | 19 | 29 | 10 | 10 | 7 | 9 | 54 | 13 | 14 | 7 | 17 | 10 | 7 | 8 | 9 | 8 | 9 | 70 | 12 | 11 | 7 | 7 |
| EDEM2 | chr20:35134105-35134155 | 20 | 13 | 10 | 14 | 15 | 18 | 15 | 19 | 12 | 11 | 10 | 43 | 43 | 16 | 16 | 11 | 9 | 11 | 11 | 10 | 11 | 22 | 15 | 21 | 17 | 19 | 14 | 19 | 21 | 10 | 9 | 15 | 15 | 14 | 18 | 12 | 17 | 17 | 18 | 11 | 21 | 19 | 20 | 25 | 21 | 19 | 33 | 16 | 20 | 18 | 12 |
| MADD | chr11:47308980-47309042 | 1 | 1 | 4 | 0 | 2 | 1 | 0 | 1 | 61 | 87 | 59 | 98 | 98 | 89 | 91 | 70 | 77 | 71 | 59 | 25 | 53 | 1 | 1 | 7 | 4 | 2 | 1 | 3 | 0 | 44 | 58 | 0 | 1 | 1 | 3 | 70 | 2 | 1 | 2 | 32 | 1 | 0 | 2 | 3 | 1 | 2 | 3 | 1 | 1 | 2 | 1 |
| SMG7 | chr1:183547103-183547252 | 13 | 9 | 18 | 14 | 13 | 19 | 31 | 15 | 44 | 66 | 48 | 95 | 91 | 69 | 76 | 53 | 56 | 60 | 51 | 46 | 45 | 21 | 17 | 26 | 17 | 24 | 15 | 27 | 24 | 44 | 47 | 18 | 17 | 14 | 18 | 35 | 11 | 11 | 29 | 31 | 21 | 15 | 15 | 14 | 8 | 25 | 73 | 22 | 14 | 15 | 8 |
| AGRN | chr1:1051032-1051043 | 0 | 0 | 1 | 0 | 1 | 0 | 0 | 0 | 59 | 69 | 56 | 43 | 35 | 72 | 75 | 53 | 56 | 56 | 51 | 55 | 49 | 1 | 1 | 2 | 1 | 1 | 1 | 2 | 1 | 17 | 15 | 1 | 1 | 1 | 0 | 5 | 4 | 0 | 0 | 2 | 1 | 0 | 0 | 1 | 0 | 1 | 29 | 0 | 0 | 1 | 2 |
| CCDC66 | chr3:56598190-56598300 | 8 | 11 | 10 | 10 | 11 | 9 | 7 | 8 | 13 | 14 | 15 | 19 | 21 | 14 | 13 | 12 | 12 | 17 | 13 | 10 | 7 | 10 | 9 | 13 | 10 | 10 | 9 | 11 | 10 | 8 | 9 | 12 | 14 | 10 | 7 | 9 | 10 | 8 | 12 | 13 | 10 | 7 | 7 | 7 | 10 | 10 | 5 | 12 | 12 | 10 | 11 |
| OCRL | chrX:129584344-129584367 | 11 | 14 | 23 | 15 | 16 | 16 | 22 | 13 | 50 | 73 | 48 | 89 | 86 | 71 | 82 | 60 | 77 | 65 | 50 | 36 | 54 | 11 | 10 | 21 | 13 | 23 | 13 | 26 | 19 | 24 | 37 | 6 | 8 | 12 | 12 | 64 | 11 | 10 | 18 | 62 | 15 | 11 | 11 | 12 | 12 | 18 | 24 | 9 | 16 | 13 | 16 |
| MICAL3 | chr22:17827644-17827781 | 7 | 5 | 18 | 5 | 8 | 5 | 7 | 8 | 73 | 78 | 77 | 96 | 96 | 89 | 88 | 81 | 81 | 81 | 85 | 89 | 86 | 5 | 9 | 12 | 8 | 9 | 6 | 9 | 11 | 31 | 27 | 10 | 7 | 5 | 11 | 52 | 51 | 7 | 9 | 44 | 13 | 9 | 11 | 7 | 7 | 6 | 65 | 12 | 5 | 9 | 10 |
| SEC24C | chr10:73762098-73762166 | 5 | 6 | 5 | 7 | 8 | 7 | 12 | 6 | 24 | 31 | 25 | 38 | 38 | 33 | 38 | 18 | 22 | 29 | 24 | 7 | 15 | 6 | 8 | 17 | 11 | 14 | 4 | 16 | 10 | 13 | 15 | 6 | 7 | 5 | 5 | 16 | 3 | 5 | 5 | 29 | 9 | 3 | 3 | 8 | 4 | 9 | 5 | 5 | 8 | 6 | 4 |
| MAP2 | chr2:209692625\|209692637-209697051 | 25 | 19 | 39 | 12 | 12 | 13 | -1 | 20 | 78 | 88 | 67 | 83 | 77 | 89 | 90 | 82 | 82 | 80 | 67 | 56 | 74 | -1 | -1 | 38 | 37 | 10 | 4 | 9 | -1 | 20 | 20 | 8 | 5 | 13 | 6 | 26 | 15 | 17 | 6 | 35 | 9 | 3 | 3 | 24 | 12 | 11 | 11 | 5 | 18 | 7 | 35 |
| BMP1 | chr8:22177567-22177732 | 5 | 4 | 2 | 4 | 3 | 4 | 5 | 5 | 10 | 10 | 12 | 22 | 21 | 12 | 12 | 14 | 11 | 12 | 10 | 18 | 13 | 3 | 3 | 5 | 4 | 5 | 3 | 5 | 3 | 6 | 7 | 3 | 3 | 4 | 4 | 22 | 6 | 2 | 2 | 2 | 4 | 3 | 3 | 4 | 3 | 4 | 3 | 2 | 3 | 3 | 3 |
| MICALL2 | chr7:1443577-1443724 | 0 | 0 | 0 | 0 | 0 | 0 | 0 | 0 | 13 | 26 | 9 | 55 | 46 | 34 | 33 | 23 | 38 | 15 | 10 | 21 | 21 | 0 | 0 | 0 | 0 | 0 | 0 | 0 | 0 | 7 | 13 | 0 | 0 | 0 | 0 | 48 | 0 | 0 | 0 | 0 | 0 | 0 | 0 | 0 | 0 | 0 | 2 | 0 | 0 | 0 | 0 |
| ALG14 | chr1:94983519-94983633 | 2 | 3 | 2 | 5 | 5 | 5 | 4 | 3 | 6 | 8 | 8 | 23 | 26 | 11 | 11 | 7 | 4 | 8 | 7 | 2 | 4 | 6 | 6 | 8 | 2 | 5 | 2 | 5 | 4 | 6 | 3 | 4 | 4 | 5 | 2 | 9 | 7 | 10 | 2 | 4 | 6 | 3 | 3 | 3 | 2 | 2 | 23 | 4 | 10 | 9 | 2 |
| NCAPG2 | chr7:158641431-158641554 | 5 | 5 | 3 | 5 | 5 | 4 | 7 | 5 | 16 | 18 | 18 | 22 | 18 | 16 | 19 | 17 | 11 | 18 | 21 | 9 | 10 | 6 | 2 | 5 | 6 | 5 | 5 | 5 | 8 | 7 | 6 | 4 | 4 | 5 | 6 | 16 | 5 | 4 | 3 | 4 | 4 | 7 | 8 | 7 | 6 | 6 | 5 | 3 | 5 | 5 | 9 |
| RRAGB | chrX:55727307-55727390 | 12 | 9 | 7 | 12 | 10 | 14 | 10 | 11 | 63 | 76 | 58 | 56 | 54 | 73 | 78 | 65 | 66 | 64 | 60 | 58 | 64 | 11 | 10 | 15 | 11 | 13 | 6 | 13 | 14 | 29 | 33 | 9 | 7 | 8 | 7 | 29 | 24 | 12 | 7 | 17 | 12 | 8 | 9 | 10 | 5 | 11 | 36 | 8 | 10 | 9 | 5 |
| UNC13A | chr19:17616408-17616464 | 1 | 0 | 0 | 0 | 0 | 0 | 0 | -1 | 1 | 2 | 2 | 25 | 24 | 3 | 4 | 1 | 2 | 1 | 3 | 19 | 8 | 0 | 0 | 14 | 6 | -1 | 0 | 17 | -1 | 0 | 3 | 0 | -1 | 1 | -1 | 0 | 2 | 0 | 0 | 0 | 0 | -1 | 5 | 11 | 0 | 4 | 1 | 0 | 0 | 2 | -1 |
| ACBD5 | chr10:27223207-27223239 | 9 | 12 | 11 | 10 | 11 | 8 | 9 | 10 | 11 | 20 | 12 | 49 | 43 | 20 | 28 | 12 | 17 | 13 | 11 | 9 | 12 | 12 | 8 | 11 | 13 | 10 | 23 | 11 | 5 | 7 | 10 | 10 | 24 | 13 | 13 | 11 | 7 | 6 | 19 | 20 | 11 | 15 | 13 | 17 | 34 | 15 | 6 | 10 | 10 | 14 | 18 |
| BCAS3 | chr17:61026866-61026910 | 10 | 7 | 3 | 20 | 18 | 22 | 15 | 7 | 17 | 38 | 22 | 59 | 57 | 45 | 51 | 21 | 24 | 27 | 21 | 3 | 17 | 12 | 12 | 24 | 16 | 19 | 5 | 17 | 14 | 17 | 19 | 4 | 4 | 7 | 6 | 44 | 7 | 8 | 3 | 7 | 8 | 4 | 5 | 11 | 3 | 11 | 2 | 5 | 18 | 9 | 2 |
| L1CAM | chrX:153863368-153863379 | 3 | 4 | 77 | 4 | 2 | 3 | 6 | 3 | 97 | ## | 95 | ## | ## | 99 | ## | 98 | 99 | 98 | 91 | 91 | 99 | 0 | 1 | 14 | 20 | 5 | 3 | 5 | 0 | 2 | 3 | 1 | 3 | 9 | 3 | 9 | 2 | 5 | 14 | 72 | 2 | 3 | 10 | 18 | 3 | 11 | 6 | 3 | 3 | 9 | 24 |
| ABLIM3 | chr5:149237448-149239277 | 1 | 1 | 2 | 1 | 2 | 1 | 0 | 1 | 37 | 58 | 30 | 91 | 91 | 61 | 62 | 33 | 51 | 58 | 32 | 32 | 40 | 1 | 1 | 4 | 4 | 2 | 1 | 2 | 1 | 64 | 61 | 1 | 1 | 1 | 4 | 40 | 5 | 1 | 2 | 4 | 13 | 1 | 2 | 3 | 1 | 2 | 31 | 1 | 1 | 1 | 1 |
| AFAP1 | chr4:7781376\|7781574\|7781551-7781627 | 1 | 2 | 3 | 1 | 1 | 1 | 1 | 2 | 40 | 62 | 53 | 86 | 92 | 65 | 69 | 45 | 43 | 69 | 46 | 12 | 20 | 0 | 1 | 2 | 4 | 2 | 2 | 2 | 1 | 3 | 2 | 2 | 2 | 2 | 5 | 1 | 2 | 3 | 4 | 45 | 4 | 2 | 2 | 5 | 2 | 3 | 91 | 5 | 1 | 1 | 3 |
| GPC2 | chr7:100175572-100175894 | 11 | 12 | 14 | 8 | 8 | 9 | -1 | 7 | 91 | 94 | 86 | 96 | 97 | 90 | 92 | 92 | 83 | 89 | 87 | 89 | 87 | 5 | 0 | 14 | 7 | 10 | 5 | 12 | -1 | 25 | 11 | 7 | 13 | 14 | 3 | 53 | 34 | 12 | 8 | 32 | 7 | 7 | 8 | 7 | 4 | 10 | 82 | 6 | 5 | 6 | 19 |
| EVC | chr4:5820402\|5819899-5822834 | 28 | 23 | 5 | 15 | 12 | 15 | 16 | 31 | 55 | 53 | 47 | 59 | 56 | 44 | 47 | 53 | 61 | 57 | 46 | 50 | 54 | 22 | 28 | 16 | 18 | 16 | 10 | 14 | 15 | 9 | 10 | 5 | 9 | 11 | 10 | 28 | 9 | 18 | 5 | 19 | 17 | 10 | 9 | 18 | 4 | 13 | 11 | 10 | 24 | 32 | 8 |
| OMA1 | chr1:58522827-58522912\|58523000 | 11 | 6 | 4 | 8 | 8 | 11 | 7 | 9 | 10 | 10 | 10 | 28 | 23 | 13 | 14 | 10 | 12 | 9 | 10 | 14 | 11 | 10 | 15 | 9 | 6 | 9 | 4 | 9 | 9 | 7 | 5 | 4 | 2 | 8 | 6 | 8 | 11 | 8 | 5 | 15 | 10 | 8 | 10 | 8 | 8 | 6 | 5 | 6 | 9 | 8 | 6 |
| WDFY3 | chr4:84808334-84808417 | 3 | 2 | 12 | 2 | 3 | 3 | 10 | 5 | 26 | 38 | 25 | 89 | 81 | 44 | 46 | 35 | 52 | 21 | 29 | 65 | 54 | -1 | -1 | 4 | 11 | 1 | 14 | 1 | 0 | 2 | 2 | 15 | 7 | 2 | 31 | 4 | 1 | 2 | 46 | 61 | 20 | 8 | 9 | 17 | 1 | 20 | 26 | 12 | 5 | 8 | 0 |
| NT5DC3 | chr12:103785621\|103785571-103785747 | 16 | 16 | 8 | 17 | 14 | 15 | 22 | 12 | 4 | 2 | 7 | 5 | 4 | 3 | 1 | 1 | 3 | 5 | 3 | 3 | 3 | 36 | 15 | 10 | 9 | 8 | 9 | 7 | 12 | 16 | 9 | 6 | 11 | 18 | 12 | 5 | 13 | 16 | 10 | 11 | 10 | 12 | 14 | 9 | 12 | 8 | 11 | 12 | 12 | 12 | 19 |
| NT5DC3 | chr12:103785621-103785747\|103787253 | 12 | 13 | 6 | 12 | 10 | 11 | 15 | 9 | 4 | 1 | 6 | 3 | 2 | 3 | 0 | 1 | 2 | 3 | 3 | 4 | 2 | 27 | 11 | 7 | 6 | 5 | 6 | 5 | 10 | 9 | 6 | 5 | 10 | 13 | 9 | 5 | 10 | 13 | 7 | 6 | 6 | 10 | 11 | 7 | 11 | 6 | 8 | 8 | 8 | 9 | 15 |
| NT5DC3 | chr12:103785571-103785747\|103787253 | 11 | 12 | 8 | 13 | 11 | 11 | 15 | 9 | 2 | 2 | 8 | 5 | 4 | 2 | 1 | 1 | 3 | 4 | 2 | 5 | 2 | 27 | 10 | 8 | 7 | 7 | 7 | 5 | 9 | 15 | 8 | 7 | 11 | 15 | 10 | 4 | 10 | 12 | 11 | 10 | 9 | 9 | 10 | 7 | 11 | 7 | 11 | 11 | 10 | 9 | 19 |
| WWC2 | chr4:183269762\|183269995-183270047 | 11 | 8 | 5 | 14 | 12 | 15 | 17 | 13 | 7 | 5 | 8 | 6 | 10 | 9 | 8 | 5 | 8 | 6 | 9 | 7 | 4 | 29 | 11 | 13 | 11 | 12 | 9 | 13 | 9 | 7 | 6 | 8 | 8 | 7 | 8 | 13 | 12 | 11 | 12 | 8 | 14 | 13 | 10 | 16 | 12 | 14 | 20 | 7 | 16 | 13 | 7 |
| MGLL | chr3:127715853-127715930 | 0 | 0 | 0 | 0 | 0 | 0 | 2 | 0 | 0 | 0 | 0 | 0 | 0 | 0 | 0 | 0 | 0 | 0 | 0 | 0 | 0 | 6 | 4 | 4 | 25 | 1 | 15 | 1 | 2 | 0 | 0 | 6 | 2 | 9 | 15 | 0 | 0 | 1 | 3 | 0 | 1 | 2 | 1 | 11 | 0 | 8 | 0 | 0 | 0 | 9 | 0 |
| AOAH | chr7:36597635-36608478 | 7 | 6 | 4 | 4 | 5 | 6 | 6 | 7 | 4 | 5 | 4 | 5 | 5 | 5 | 4 | 7 | 5 | 4 | 3 | 4 | 4 | -1 | -1 | 7 | 7 | 5 | 8 | 8 | 23 | 7 | 6 | 9 | 10 | 8 | 9 | 6 | 7 | 6 | 7 | 7 | 10 | 10 | 7 | 8 | 5 | 8 | 10 | 10 | 8 | 7 | 6 |
| SCIN | chr7:12577542-12577759 | 1 | 0 | 2 | 1 | 0 | 1 | 1 | 2 | 6 | 2 | 8 | 3 | 3 | 0 | 1 | 4 | 11 | 4 | 2 | 2 | 1 | 0 | -1 | 2 | 2 | 0 | 1 | 1 | 32 | 4 | -1 | 2 | -1 | 4 | 2 | 1 | 1 | 0 | 0 | 1 | 1 | 1 | 2 | 4 | -1 | 1 | 28 | 2 | 2 | 1 | -1 |
| ARHGAP21 | chr10:24609430-24610365\|24609531 | 8 | 13 | 9 | 11 | 12 | 11 | 15 | 11 | 8 | 11 | 10 | 5 | 6 | 11 | 9 | 11 | 8 | 13 | 9 | 7 | 5 | 12 | 10 | 11 | 12 | 9 | 12 | 10 | 22 | 10 | 8 | 7 | 11 | 10 | 11 | 7 | 9 | 12 | 9 | 9 | 12 | 12 | 8 | 12 | 13 | 12 | 10 | 8 | 10 | 11 | 13 |
| UBE3D | chr6:83049709-83049772 | 9 | 8 | 5 | 8 | 8 | 8 | 5 | 12 | 7 | 5 | 11 | 8 | 6 | 9 | 7 | 9 | 11 | 9 | 7 | 7 | 6 | -1 | -1 | 9 | 8 | 9 | 7 | 10 | 28 | 7 | 5 | 5 | 12 | 13 | 6 | 8 | 13 | 7 | 9 | 7 | 9 | 12 | 13 | 7 | 6 | 8 | 18 | 9 | 10 | 8 | 6 |
| NFASC | chr1:204962108-204962158 | 1 | 1 | 2 | 1 | 2 | 1 | 0 | 1 | 62 | 32 | 60 | 10 | 14 | 39 | 28 | 57 | 39 | 49 | 60 | 63 | 56 | 2 | 0 | 1 | 1 | 1 | 1 | 1 | 0 | 48 | 6 | 5 | 1 | 1 | 1 | 2 | 13 | 1 | 1 | 2 | 1 | 0 | 1 | 1 | 1 | 3 | 1 | 2 | 1 | 1 | -1 |
| SYCE1L | chr16:77201067-77201142 | 11 | 10 | 6 | 7 | 7 | 10 | -1 | 8 | 1 | 2 | 5 | 2 | 4 | 5 | 1 | 3 | 3 | 4 | 2 | 5 | 2 | 0 | 0 | 9 | 12 | 10 | 8 | 10 | -1 | 57 | 34 | -1 | 12 | 10 | 1 | 12 | 9 | 8 | 9 | 2 | 7 | 10 | 9 | 11 | 13 | 9 | 5 | 2 | 5 | 4 | 12 |
| CACNA1G | chr17:50594993-50595061 | 8 | 9 | 3 | 13 | 11 | 15 | -1 | 7 | 49 | 66 | 39 | 17 | 16 | 81 | 82 | 54 | 82 | 41 | 32 | 65 | 52 | 9 | 7 | 13 | 11 | 9 | 12 | 16 | 11 | 82 | 53 | -1 | 0 | 13 | -1 | -1 | 10 | 4 | -1 | 30 | 4 | 9 | 11 | -1 | -1 | 9 | 7 | 14 | 8 | 7 | -1 |
| PROM1 | chr4:15980277-15980301 | 3 | 0 | 0 | 4 | 10 | 5 | 3 | 0 | 7 | 7 | 7 | 3 | 5 | 5 | 4 | 5 | 5 | 5 | 4 | 5 | 5 | -1 | 0 | 1 | 1 | 1 | 2 | 1 | 0 | 25 | 16 | 0 | 2 | 1 | 0 | 3 | 8 | 0 | 0 | 1 | 2 | 0 | 0 | 1 | 8 | 3 | 0 | 1 | 0 | -1 | 0 |
| CPNE5 | chr6:36838696-36839181\|36838782 | 8 | 6 | 7 | 9 | 16 | 7 | 11 | 7 | 2 | 1 | 1 | 4 | 4 | 1 | 1 | 2 | 2 | 1 | 2 | 4 | 2 | -1 | -1 | 6 | 7 | 7 | 9 | 10 | 6 | 62 | 42 | 7 | 9 | 7 | 8 | 37 | 7 | 5 | 9 | 11 | 6 | 8 | 7 | 6 | 5 | 4 | 15 | 5 | 5 | 6 | 13 |
| LMO7 | chr13:75838347-75838388 | 4 | 1 | 1 | 5 | 11 | 4 | 0 | 1 | 4 | 2 | 1 | 1 | 2 | 2 | 3 | 3 | 5 | 1 | 2 | 25 | 10 | 0 | 1 | 6 | 1 | 18 | 0 | 30 | 4 | 88 | 84 | 0 | 0 | 0 | 0 | 29 | 7 | 1 | 0 | 0 | 3 | 0 | 0 | 0 | 1 | 1 | 2 | 0 | 3 | 0 | 3 |
| NAV1 | chr1:201804489-201804497 | 6 | 6 | 3 | 6 | 6 | 6 | 8 | 6 | 35 | 32 | 36 | 24 | 16 | 34 | 32 | 36 | 26 | 27 | 36 | 39 | 30 | 7 | 2 | 6 | 6 | 5 | 3 | 6 | 12 | 6 | 47 | 4 | 3 | 5 | 4 | 4 | 7 | 6 | 4 | 3 | 4 | 4 | 4 | 7 | 3 | 4 | 3 | 4 | 6 | 3 | 4 |
| SLC25A36 | chr3:140954988-140955029 | 0 | 0 | 0 | 0 | 0 | 0 | 0 | 0 | 0 | 0 | 0 | 0 | 0 | 0 | 0 | 0 | 0 | 0 | 0 | 0 | 0 | 0 | 0 | 0 | 0 | 0 | 0 | 0 | 0 | 18 | 39 | 0 | 0 | 0 | 0 | 9 | 0 | 0 | 0 | 0 | 0 | 0 | 0 | 0 | 0 | 0 | 0 | 0 | 0 | 0 | 0 |
| CALCA | chr11:14971948-14972143 | 2 | 2 | 2 | 4 | 3 | 2 | 0 | 2 | -1 | -1 | 3 | 0 | 0 | 2 | 4 | -1 | 3 | 5 | -1 | 4 | 0 | 0 | 0 | 2 | 4 | 6 | 2 | 3 | 0 | 3 | 23 | 3 | 4 | 2 | 0 | 8 | 3 | -1 | 4 | 3 | 2 | 3 | 1 | 0 | 3 | 3 | 6 | 1 | 0 | 1 | 1 |
| CHN2 | chr7:29292925-29293608 | 1 | 2 | 0 | 1 | 3 | 2 | 0 | 1 | 0 | 1 | 1 | 5 | 5 | 2 | 1 | 0 | 2 | 2 | 1 | 1 | 0 | 0 | 0 | 1 | 0 | 1 | 0 | 1 | -1 | 6 | 41 | 0 | 0 | 1 | 0 | 23 | 2 | 0 | 0 | 0 | 2 | 1 | 1 | 0 | 1 | 0 | 0 | 1 | 1 | 3 | 0 |
| GGT7 | chr20:34850782\|34850132-34850917 | 3 | 2 | 3 | 1 | 2 | 1 | 1 | 2 | 2 | 1 | 2 | 0 | 0 | 1 | 1 | 2 | 1 | 1 | 2 | 3 | 3 | 1 | 1 | 2 | 2 | 2 | 2 | 2 | 2 | 25 | 37 | 2 | 2 | 2 | 2 | 8 | 4 | 1 | 1 | 2 | 2 | 2 | 2 | 3 | 3 | 2 | 2 | 1 | 1 | 1 | 3 |
| GGT7 | chr20:34850782-34850917\|34850852 | 0 | 1 | 1 | 0 | 1 | 0 | 0 | 0 | 2 | 1 | 2 | 0 | 0 | 1 | 0 | 2 | 1 | 1 | 2 | 3 | 2 | 0 | 0 | 1 | 1 | 1 | 0 | 1 | 0 | 29 | 43 | 1 | 0 | 1 | 1 | 9 | 3 | 0 | 0 | 0 | 1 | 0 | 0 | 0 | 1 | 1 | 1 | 0 | 0 | 0 | 1 |
| DENND5B | chr12:31432043-31432153 | 0 | 0 | 0 | 0 | 2 | 0 | 0 | 0 | 1 | 0 | 0 | 0 | 0 | 0 | 0 | 0 | 0 | 0 | 0 | 0 | 0 | 0 | 0 | 0 | 0 | 0 | 0 | 0 | 0 | 48 | 64 | 0 | 0 | 0 | 0 | 1 | 0 | 0 | 0 | 0 | 0 | 1 | 0 | 0 | 0 | 0 | 0 | 0 | 0 | 0 | 1 |
| LAMA2 | chr6:129292803-129293066\|129294134 | 3 | 2 | 2 | 4 | 5 | 4 | 1 | 2 | 3 | 11 | 1 | 4 | 8 | 24 | 22 | 3 | 3 | 4 | 3 | 2 | 3 | 1 | 2 | 3 | 2 | 4 | 3 | 3 | 2 | 28 | 61 | 2 | 3 | 5 | 4 | 37 | 6 | 2 | 2 | 10 | 3 | 3 | 4 | 2 | 1 | 3 | 4 | 3 | 3 | 3 | 11 |
| PHLDB2 | chr3:111949702-111949890 | 2 | 1 | 0 | 1 | 4 | 3 | 3 | 1 | 1 | 13 | 2 | 10 | 1 | 14 | 23 | 5 | 2 | 2 | 1 | 1 | 7 | 0 | 1 | 1 | 1 | 1 | 0 | 2 | 1 | 40 | 92 | 0 | 0 | 1 | 2 | 58 | 4 | 1 | 0 | 1 | 2 | 1 | 2 | 1 | 0 | 1 | 10 | 1 | 1 | 1 | 2 |
| ARHGEF10 | chr8:1858757-1858816 | 1 | 1 | 2 | 2 | 3 | 3 | 7 | 1 | 0 | 1 | 0 | 2 | 2 | 1 | 1 | 0 | 1 | 0 | 0 | 0 | 0 | 0 | 0 | 6 | 3 | 4 | 2 | 6 | 1 | 35 | 47 | 1 | 2 | 1 | 4 | 8 | 1 | 1 | 1 | 0 | 4 | 1 | 1 | 2 | 2 | 1 | 0 | 1 | 2 | 1 | 1 |
| THAP9 | chr4:82905861-82905926 | 4 | 5 | 2 | 3 | 6 | 4 | 0 | 2 | 44 | 38 | 39 | 7 | 9 | 31 | 24 | 50 | 27 | 34 | 40 | 52 | 49 | -1 | 0 | 3 | 2 | 2 | 2 | 3 | 0 | 14 | 24 | 6 | 1 | 4 | 1 | 1 | 9 | 3 | 1 | 1 | 1 | 1 | 0 | 2 | 3 | 2 | 3 | 3 | 2 | 2 | 2 |
| ARHGEF28 | chr5:73923087-73923164 | 15 | 18 | 19 | 14 | 17 | 13 | 16 | 17 | 12 | 12 | 13 | 18 | 16 | 14 | 13 | 12 | 10 | 11 | 12 | 12 | 10 | 12 | 16 | 14 | 18 | 15 | 15 | 15 | 20 | 20 | 33 | 19 | 20 | 18 | 16 | 18 | 13 | 18 | 22 | 15 | 16 | 14 | 14 | 18 | 18 | 18 | 15 | 14 | 16 | 15 | 14 |
| PPFIBP1 | chr12:27655143-27655235 | 0 | 0 | 1 | 0 | 2 | 0 | 0 | 0 | 14 | 52 | 8 | 11 | 4 | 52 | 58 | 24 | 28 | 12 | 5 | 2 | 9 | 0 | 1 | 0 | 0 | 0 | 0 | 0 | 1 | 48 | 83 | 4 | 1 | 1 | 3 | 52 | 1 | 0 | 2 | 5 | 2 | 0 | 0 | 1 | 0 | 1 | 0 | 3 | 0 | 0 | 6 |
| ANKRD36 | chr2:97199782-97199844 | 0 | 0 | 0 | 0 | 0 | 0 | 14 | 0 | 0 | 1 | 1 | 0 | 0 | 0 | 0 | 0 | 1 | 0 | 1 | 0 | 0 | 0 | 0 | 4 | 2 | 5 | 1 | 9 | 1 | 20 | 29 | 0 | 0 | 0 | 0 | 5 | 0 | 0 | 0 | 0 | 1 | 0 | 0 | 2 | 0 | 1 | 0 | 1 | 0 | 0 | 0 |
| DNAAF1 | chr16:84162190\|84162132-84162215 | 0 | 0 | 0 | -1 | 0 | 0 | -1 | 1 | 5 | 21 | 3 | 23 | 24 | 27 | 29 | 5 | 6 | 11 | 25 | 4 | 18 | 0 | 0 | 4 | 2 | -1 | 3 | 2 | 1 | 26 | 41 | 0 | -1 | 0 | 0 | 9 | 2 | 0 | 0 | 1 | 1 | 0 | 7 | -1 | 0 | 1 | 1 | 0 | 0 | -1 | 0 |
| SORBS2 | chr4:185676997-185677638 | 2 | 1 | 0 | 1 | 3 | 1 | 1 | 1 | 7 | 5 | 4 | 1 | 3 | 3 | 3 | 8 | 10 | 3 | 3 | 9 | 7 | 2 | 2 | 6 | 5 | 4 | 1 | 6 | 1 | 47 | 53 | 1 | 0 | 1 | 0 | 8 | 7 | 5 | 0 | 2 | 2 | 2 | 2 | 7 | 1 | 2 | 3 | 0 | 2 | 2 | 10 |
| GANC | chr15:42278025-42278056\|42278088 | 2 | 2 | 2 | 2 | 2 | 2 | 0 | 2 | 49 | 48 | 51 | 12 | 10 | 37 | 38 | 43 | 36 | 49 | 51 | 49 | 44 | 3 | 1 | 2 | 2 | 1 | 1 | 2 | 2 | 18 | 25 | 5 | 2 | 3 | 1 | 6 | 11 | 1 | 1 | 2 | 2 | 1 | 1 | 2 | 1 | 2 | 7 | 1 | 1 | 1 | 1 |
| EIF4E2 | chr2:232558822-232563808\|232558900 | 0 | 0 | 0 | 1 | 1 | 1 | 1 | 0 | 0 | 0 | 0 | 0 | 0 | 0 | 0 | 0 | 0 | 0 | 0 | 0 | 0 | 0 | 0 | 1 | 1 | 1 | 0 | 1 | 0 | 19 | 22 | 0 | 0 | 0 | 0 | 8 | 0 | 0 | 0 | 0 | 0 | 0 | 0 | 0 | 0 | 0 | 0 | 0 | 0 | 0 | 0 |
| CTIF | chr18:48625189-48625236 | 8 | 7 | 2 | 7 | 7 | 9 | 8 | 7 | 6 | 9 | 6 | 9 | 9 | 10 | 12 | 7 | 7 | 5 | 8 | 6 | 9 | 3 | 5 | 10 | 9 | 8 | 5 | 9 | 4 | 23 | 36 | 4 | 9 | 3 | 4 | 26 | 5 | 2 | 3 | 2 | 5 | 4 | 5 | 6 | 3 | 6 | 4 | 3 | 4 | 5 | 4 |
| SLC12A7 | chr5:1056582-1056596 | 4 | 2 | 1 | 12 | 12 | 24 | 15 | 3 | 2 | 3 | 1 | 2 | 2 | 4 | 4 | 1 | 6 | 1 | 1 | 3 | 15 | 7 | 3 | 35 | 9 | 20 | 4 | 25 | 6 | 62 | 83 | 0 | 0 | 3 | 3 | 35 | 4 | 4 | 0 | 10 | 9 | 3 | 4 | 3 | 0 | 4 | 1 | 2 | 9 | 7 | 3 |
| RAD17 | chr5:69371454-69371557 | 3 | 7 | 3 | 10 | 8 | 7 | 3 | 5 | 10 | 5 | 8 | 4 | 5 | 5 | 4 | 9 | 5 | 6 | 9 | 8 | 7 | 7 | 6 | 9 | 8 | 9 | 5 | 9 | 9 | 14 | 22 | 6 | 8 | 6 | 5 | 11 | 6 | 8 | 8 | 8 | 7 | 2 | 2 | 5 | 6 | 8 | 12 | 5 | 8 | 6 | 6 |
| HFM1 | chr1:91374905-91374965 | 1 | 1 | 0 | 0 | 3 | 1 | 0 | 1 | 8 | 0 | 3 | 6 | 5 | 2 | 4 | 0 | 6 | 6 | -1 | -1 | -1 | 0 | 0 | 1 | 3 | 1 | 4 | 5 | 0 | 28 | 30 | 0 | 0 | 2 | -1 | 12 | 1 | 1 | 1 | 0 | 4 | 0 | 2 | 2 | 0 | 1 | 1 | 0 | 0 | 0 | 0 |
| FHOD3 | chr18:36759118-36759141 | 0 | 0 | 0 | 0 | 1 | 0 | 0 | 0 | 0 | 0 | 0 | 0 | 0 | 0 | 0 | 0 | 0 | 0 | 0 | 0 | 0 | 0 | 0 | 0 | 0 | 0 | 0 | 0 | 0 | 35 | 50 | 0 | 0 | 0 | 0 | 38 | 0 | 0 | 0 | 0 | 0 | 0 | 0 | 0 | 0 | 0 | 0 | 0 | 0 | 0 | 10 |
| CLDN12 | chr7:90412007-90412049\|90412061 | 10 | 11 | 5 | 19 | 22 | 21 | 16 | 7 | 8 | 19 | 18 | 15 | 14 | 21 | 21 | 15 | 16 | 29 | 19 | 9 | 11 | 13 | 12 | 23 | 12 | 24 | 7 | 27 | 12 | 44 | 68 | 5 | 9 | 8 | 4 | 46 | 8 | 10 | 6 | 7 | 13 | 5 | 4 | 8 | 10 | 12 | 9 | 7 | 15 | 7 | 6 |
| NF1 | chr17:31367225\|31367249-31367278 | 2 | 2 | 1 | 4 | 4 | 6 | 8 | 1 | 3 | 6 | 4 | 16 | 14 | 7 | 8 | 4 | 3 | 4 | 5 | 3 | 4 | 0 | 4 | 9 | 5 | 10 | 2 | 13 | 4 | 41 | 56 | 1 | 0 | 1 | 1 | 42 | 2 | 2 | 1 | 1 | 5 | 1 | 1 | 3 | 1 | 4 | 3 | 1 | 3 | 2 | 1 |
| GRAMD3 | chr5:126466294-126466336 | 5 | 4 | 2 | 7 | 10 | 15 | 12 | 3 | 7 | 8 | 6 | 25 | 14 | 8 | 9 | 6 | 6 | 5 | 6 | 6 | 5 | 9 | 6 | 15 | 6 | 22 | 6 | 27 | 11 | 40 | 51 | 2 | 2 | 2 | 3 | 6 | 2 | 4 | 2 | 5 | 8 | 2 | 2 | 5 | 4 | 8 | 5 | 2 | 7 | 7 | 2 |
| PLPP5 | chr8:38267606-38267693\|38267732 | 7 | 5 | 6 | 6 | 6 | 5 | 4 | 5 | 11 | 10 | 10 | 11 | 12 | 12 | 12 | 8 | 8 | 9 | 11 | 7 | 10 | 3 | 4 | 8 | 5 | 9 | 4 | 9 | 3 | 12 | 22 | 4 | 6 | 5 | 4 | 17 | 7 | 2 | 4 | 7 | 5 | 4 | 5 | 4 | 4 | 4 | 4 | 3 | 4 | 4 | 4 |
| NRCAM | chr7:108150695-108150753 | 1 | 5 | 19 | 0 | 0 | 0 | 0 | 2 | 1 | 1 | 1 | 0 | 0 | 0 | 0 | 2 | 1 | 1 | 1 | 3 | 2 | 0 | 0 | 0 | 0 | 1 | 3 | 0 | 0 | 1 | 1 | 47 | 15 | 0 | 1 | 0 | 0 | 1 | 0 | 1 | 2 | 0 | 0 | 1 | 7 | 2 | 0 | 26 | 0 | 0 | 2 |
| CHRD | chr3:184381076-184381130 | 5 | 6 | 7 | 3 | 3 | 3 | 6 | 6 | 3 | 2 | 2 | 1 | 1 | 2 | 2 | 2 | 3 | 2 | 3 | 3 | 4 | 7 | 3 | 4 | 4 | 4 | 5 | 4 | 3 | 3 | 5 | 20 | 55 | 4 | 7 | 11 | 4 | 2 | 5 | 5 | 6 | 4 | 3 | 5 | 10 | 5 | 13 | 5 | 3 | 3 | 6 |
| GBAS | chr7:55991828-55991884 | 5 | 3 | 3 | 2 | 3 | 4 | 4 | 5 | 1 | 1 | 1 | 3 | 4 | 1 | 1 | 2 | 1 | 1 | 1 | 2 | 1 | 5 | 5 | 3 | 4 | 3 | 2 | 3 | 7 | 1 | 1 | 5 | 40 | 4 | 3 | 1 | 7 | 7 | 6 | 4 | 5 | 5 | 6 | 8 | 6 | 3 | 15 | 7 | 5 | 4 | 3 |
| TLE1 | chr9:81612292-81612393 | 4 | 7 | 7 | 4 | 4 | 4 | 7 | 5 | 2 | 2 | 2 | 1 | 1 | 1 | 1 | 2 | 2 | 1 | 1 | 2 | 3 | 3 | 3 | 5 | 21 | 3 | 6 | 2 | 3 | 4 | 4 | 8 | 82 | 5 | 7 | 14 | 2 | 2 | 27 | 5 | 10 | 4 | 4 | 9 | 12 | 14 | 4 | 7 | 4 | 4 | 12 |
| SIDT2 | chr11:117187892-117187954 | 1 | 1 | 1 | 1 | 1 | 1 | 1 | 1 | 0 | 1 | 2 | 1 | 1 | 2 | 3 | 1 | 1 | 3 | 2 | 0 | 0 | 0 | 1 | 2 | 8 | 1 | 1 | 1 | 1 | 0 | 0 | 6 | 49 | 1 | 4 | 2 | 1 | 0 | 11 | 1 | 4 | 1 | 1 | 22 | 0 | 5 | 0 | 5 | 1 | 1 | 0 |
| TMEM68 | chr8:55763422\|55763637-55763701 | 1 | 1 | 2 | 2 | 2 | 1 | 0 | 1 | 2 | 1 | 1 | 1 | 1 | 1 | 1 | 2 | 2 | 2 | 2 | 1 | 2 | 1 | 0 | 2 | 4 | 1 | 1 | 1 | 2 | 1 | 1 | 15 | 29 | 1 | 2 | 1 | 2 | 2 | 1 | 2 | 1 | 1 | 1 | 6 | 2 | 1 | 2 | 3 | 2 | 2 | 1 |
| TPTE2P5 | chr13:40877666\|40877713-40877861 | 6 | 10 | 12 | 9 | 10 | 6 | 0 | 7 | -1 | 0 | 5 | 5 | 3 | 0 | 6 | 7 | 14 | 4 | 3 | 9 | 6 | 0 | -1 | 8 | 7 | 6 | 5 | 7 | 0 | 6 | 7 | 13 | 38 | 11 | 10 | 8 | 5 | 1 | 7 | 7 | 5 | 8 | 5 | 9 | 7 | 8 | 2 | 7 | 7 | 4 | 7 |
| DIP2B | chr12:50686213-50686353 | 2 | 2 | 1 | 2 | 2 | 2 | 2 | 3 | 2 | 1 | 2 | 8 | 11 | 2 | 1 | 1 | 1 | 2 | 1 | 1 | 1 | 3 | -1 | 5 | 4 | 3 | 1 | 2 | -1 | 1 | 1 | 4 | 24 | 2 | 1 | 4 | 4 | 6 | 7 | 2 | 4 | 2 | 1 | 6 | 2 | 4 | 1 | 4 | 5 | 2 | 2 |
| PAPSS2 | chr10:87721756\|87721464-87721770 | 0 | 0 | 3 | 0 | 0 | 0 | 0 | 1 | 1 | 1 | 0 | 0 | 1 | 2 | 2 | 0 | 1 | 1 | 1 | 0 | 1 | 1 | 0 | 2 | 7 | 0 | 2 | 0 | 0 | 0 | 0 | 5 | 22 | 1 | 2 | 0 | 0 | 0 | 2 | 0 | 1 | 1 | 0 | 14 | 0 | 1 | 1 | 2 | 0 | 1 | 1 |
| PLCH2 | chr1:2479195-2479383 | 2 | 7 | 6 | 5 | 4 | 3 | 4 | 2 | 0 | 0 | 1 | 0 | 1 | 0 | 0 | 1 | 1 | 0 | 0 | 2 | 2 | 0 | -1 | 3 | 5 | 2 | 2 | 5 | 5 | 2 | 6 | 10 | 28 | 5 | 2 | 1 | 4 | 1 | 2 | 1 | 3 | 2 | 1 | 8 | 8 | 5 | 13 | 12 | 4 | 2 | 6 |
| PLCD4 | chr2:218622165\|218622186-218622228 | 12 | 15 | 7 | 11 | 14 | 11 | 23 | 12 | 2 | 2 | 2 | 7 | 8 | 3 | 1 | 3 | 4 | 2 | 3 | 8 | 2 | 0 | -1 | 25 | 24 | 20 | 22 | 18 | -1 | 16 | 10 | 21 | 66 | 11 | 17 | 5 | 17 | 13 | 15 | 8 | 21 | 16 | 17 | 21 | 15 | 21 | 5 | 14 | 22 | 13 | 4 |
| MALT1 | chr18:58733882-58733980 | 2 | 3 | 10 | 5 | 4 | 3 | 4 | 4 | 5 | 8 | 8 | 6 | 6 | 6 | 5 | 5 | 4 | 9 | 6 | 4 | 5 | 4 | 6 | 6 | 12 | 4 | 5 | 4 | 10 | 3 | 5 | 25 | 51 | 3 | 5 | 7 | 4 | 8 | 16 | 10 | 24 | 3 | 2 | 17 | 5 | 11 | 2 | 20 | 5 | 4 | 4 |
| PAPSS2 | chr10:87721756-87721770\|87726831 | 0 | 1 | 3 | 0 | 0 | 0 | 0 | 1 | 1 | 1 | 0 | 0 | 0 | 2 | 2 | 0 | 0 | 0 | 0 | 0 | 0 | 1 | 0 | 2 | 7 | 0 | 2 | 0 | 1 | 0 | 0 | 5 | 23 | 1 | 2 | 0 | 0 | 0 | 2 | 0 | 1 | 1 | 0 | 17 | 0 | 2 | 1 | 2 | 0 | 1 | 1 |
| UTRN | chr6:144287966-144288111 | 7 | 10 | 10 | 9 | 8 | 8 | 13 | 8 | 3 | 5 | 7 | 14 | 9 | 8 | 8 | 6 | 7 | 7 | 8 | 7 | 7 | 11 | 3 | 10 | 10 | 11 | 10 | 9 | 5 | 7 | 6 | 11 | 33 | 10 | 8 | 6 | 5 | 6 | 17 | 10 | 17 | 7 | 6 | 12 | 18 | 16 | 8 | 8 | 8 | 11 | 13 |
| CLUHP3 | chr16:31706421-31706454\|31706501\|31706740 | 12 | 11 | 10 | 11 | 11 | 12 | 10 | 11 | 7 | 8 | 11 | 10 | 9 | 7 | 8 | 10 | 8 | 8 | 9 | 12 | 9 | 11 | 5 | 12 | 13 | 11 | 14 | 10 | 10 | 10 | 12 | 22 | 30 | 14 | 14 | 9 | 14 | 9 | 16 | 10 | 12 | 13 | 13 | 14 | 15 | 13 | 12 | 14 | 12 | 13 | 16 |
| SLC39A14 | chr8:22409946-22410115 | 0 | 1 | 0 | 0 | 0 | 0 | 0 | 1 | 0 | 0 | 0 | 0 | 0 | 0 | 0 | 0 | 0 | 0 | 0 | 0 | 0 | 0 | 0 | 5 | 34 | 0 | 2 | 0 | 2 | 0 | 0 | 22 | 61 | 2 | 4 | 0 | 1 | 0 | 9 | 1 | 3 | 1 | 1 | 40 | 14 | 10 | 0 | 11 | 1 | 1 | 2 |
| SMC6 | chr2:17713385-17713580 | 7 | 11 | 8 | 7 | 7 | 7 | 7 | 9 | 7 | 7 | 6 | 11 | 11 | 6 | 6 | 7 | 6 | 7 | 5 | 7 | 7 | 5 | 13 | 9 | 8 | 9 | 10 | 8 | 9 | 6 | 7 | 10 | 21 | 9 | 8 | 7 | 7 | 9 | 10 | 14 | 10 | 10 | 9 | 13 | 12 | 9 | 7 | 11 | 11 | 10 | 8 |

SFig5, pg7

| Gene Symbol | Exon Location (hg38, human) | Adipose - Subcutaneous | Adipose - Visceral (Omentum) | Adrenal Gland | Artery - Aorta | Artery - Coronary | Artery - Tibial | Bladder | Breast - Mammary Tissue | Brain - Amygdala | Brain - Anterior cingulate cortex (BA24) | Brain - Caudate (basal ganglia) | Brain - Cerebellar Hemisphere | Brain - Cerebellum | Brain - Cortex | Brain - Frontal Cortex (BA9) | Brain - Hippocampus | Brain - Hypothalamus | Brain - Nucleus accumbens (basal ganglia) | Brain - Putamen (basal ganglia) | Brain - Spinal cord (cervical c-1) | Brain - Substantia nigra | Cervix - Ectocervix | Cervix - Endocervix | Colon - Sigmoid | Colon - Transverse | Esophagus - Gastroesophageal Junction | Esophagus - Mucosa | Esophagus - Muscularis | Fallopian Tube | Heart - Atrial Appendage | Heart - Left Ventricle | Kidney - Cortex | Liver | Lung | Minor Salivary Gland | Muscle - Skeletal | Nerve - Tibial | Ovary | Pancreas | Pituitary | Prostate | Skin - Not Sun Exposed (Suprapubic) | Skin - Sun Exposed (Lower leg) | Small Intestine - Terminal Ileum | Spleen | Stomach | Testis | Thyroid | Uterus | Vagina | Whole Blood |
|---|---|---|---|---|---|---|---|---|---|---|---|---|---|---|---|---|---|---|---|---|---|---|---|---|---|---|---|---|---|---|---|---|---|---|---|---|---|---|---|---|---|---|---|---|---|---|---|---|---|---|---|---|
| RPL32P3 | chr3:129385296-129390096 | 6 | 10 | 16 | 10 | 11 | 8 | 9 | 8 | 5 | 3 | 9 | 3 | 4 | 4 | 4 | 7 | 10 | 6 | 6 | 5 | 6 | 8 | 6 | 9 | 12 | 9 | 10 | 8 | 9 | 7 | 10 | 9 | 29 | 11 | 11 | 5 | 6 | 5 | 18 | 11 | 15 | 7 | 5 | 16 | 15 | 13 | 12 | 13 | 8 | 8 | 14 |
| RPL32P3 | chr3:129385296-129390096 | 6 | 10 | 16 | 10 | 11 | 8 | 9 | 8 | 5 | 3 | 9 | 3 | 4 | 4 | 4 | 7 | 10 | 6 | 6 | 5 | 6 | 8 | 6 | 9 | 12 | 9 | 10 | 8 | 9 | 7 | 10 | 9 | 29 | 11 | 11 | 5 | 6 | 5 | 18 | 11 | 15 | 7 | 5 | 16 | 15 | 13 | 12 | 13 | 8 | 8 | 14 |
| ZBTB33 | chrX:120252669-120252770 | 7 | 6 | 7 | 6 | 6 | 4 | -1 | 7 | 8 | 9 | 10 | 15 | 8 | 9 | 12 | 9 | 8 | 10 | 8 | 12 | 11 | 0 | -1 | 6 | 4 | 7 | 7 | 5 | -1 | 6 | 6 | 6 | 20 | 5 | 8 | 8 | 4 | 8 | 10 | 7 | 7 | 8 | 8 | 8 | 4 | 8 | 5 | 5 | 5 | 7 | 7 |
| CSAD | chr12:53164382-53166327 | 6 | 13 | 8 | 8 | 7 | 5 | 7 | 8 | 2 | 1 | 2 | 2 | 2 | 2 | 1 | 1 | 1 | 1 | 2 | 1 | 2 | 4 | 4 | 7 | 8 | 8 | 6 | 7 | 5 | 4 | 5 | 7 | 23 | 9 | 7 | 4 | 3 | 2 | 11 | 2 | 8 | 4 | 3 | 11 | 9 | 12 | 1 | 8 | 4 | 5 | 12 |
| GPR75-ASB3 | chr2:53756758-53763606 | 5 | 9 | 10 | 5 | 6 | 3 | 6 | 7 | 3 | 3 | 3 | 4 | 5 | 4 | 2 | 4 | 3 | 4 | 2 | 3 | 2 | 3 | 6 | 6 | 10 | 7 | 7 | 7 | 8 | 4 | 5 | 13 | 30 | 13 | 9 | 3 | 5 | 8 | 15 | 9 | 11 | 6 | 5 | 15 | 21 | 9 | 5 | 10 | 6 | 7 | 15 |
| CAPN3 | chr15:42362703-42362910 | 8 | 8 | 7 | 4 | 5 | 5 | 4 | 7 | 1 | 2 | 1 | 2 | 1 | 2 | 1 | 1 | 2 | 2 | 3 | 1 | 2 | 8 | 8 | 5 | 8 | 4 | 9 | 3 | 6 | 2 | 4 | 6 | 22 | 8 | 7 | 0 | 4 | 5 | 10 | 3 | 7 | 9 | 8 | 10 | 11 | 6 | 15 | 6 | 5 | 7 | 7 |
| GNPDA2 | chr4:44714596-44714659 | 8 | 11 | 9 | 8 | 9 | 6 | 9 | 10 | 10 | 8 | 9 | 13 | 18 | 8 | 5 | 10 | 10 | 8 | 9 | 12 | 9 | 7 | 9 | 13 | 12 | 9 | 11 | 9 | 8 | 11 | 14 | 14 | 31 | 10 | 12 | 18 | 15 | 8 | 21 | 11 | 15 | 7 | 7 | 12 | 18 | 12 | 12 | 15 | 12 | 13 | 8 |
| KEL | chr7:142946640-142946816 | 10 | 8 | -1 | 6 | 4 | 7 | 0 | 12 | 2 | 5 | 7 | 3 | 4 | 4 | 5 | 6 | 4 | 5 | 4 | 5 | 4 | -1 | 0 | 3 | 10 | 10 | 7 | 10 | -1 | 8 | 10 | 3 | 12 | 28 | 7 | 6 | 7 | 6 | 7 | 4 | 7 | 9 | 14 | 9 | 11 | 4 | 7 | 6 | 7 | 15 | 7 |
| SLC36A4 | chr11:93166265-93166310 | 0 | 0 | 0 | 0 | 0 | 0 | 0 | 0 | 0 | 0 | 0 | 0 | 0 | 0 | 0 | 0 | 0 | 0 | 0 | 0 | 0 | 0 | 0 | 0 | 0 | 0 | 0 | 0 | 1 | 0 | 1 | 0 | 0 | 0 | 0 | 21 | 0 | 0 | 0 | 0 | 0 | 0 | 0 | 0 | 0 | 0 | 0 | 0 | 0 | 0 | 0 |
| VPS8 | chr3:184911715\|184911807-184911864 | 0 | 0 | 0 | 0 | 0 | 0 | 0 | 0 | 1 | 0 | 0 | 0 | 0 | 0 | 0 | 0 | 0 | 0 | 0 | 1 | 1 | 0 | 0 | 0 | 0 | 0 | 0 | 0 | 0 | 0 | 1 | 0 | 0 | 0 | 1 | 29 | 1 | 0 | 0 | 0 | 0 | 0 | 0 | 0 | 0 | 0 | 0 | 0 | 0 | 0 | 0 |
| ACACA | chr17:37274622-37274690 | 0 | 0 | 0 | 0 | 0 | 0 | 0 | 0 | 1 | 1 | 2 | 1 | 1 | 1 | 1 | 1 | 1 | 4 | 1 | 0 | 0 | 0 | 0 | 0 | 0 | 0 | 0 | 0 | 0 | 2 | 1 | 0 | 0 | 0 | 1 | 43 | 0 | 0 | 0 | 0 | 0 | 0 | 0 | 1 | 0 | 0 | 1 | 0 | 0 | 0 | 0 |
| PRRC2B | chr9:131488752-131490769 | 2 | 1 | 0 | 2 | 1 | 2 | 2 | 1 | 0 | 0 | 0 | 1 | 1 | 0 | 0 | 0 | 0 | 0 | 1 | 0 | 0 | 1 | 0 | 2 | 2 | 2 | 0 | 2 | 0 | 1 | 2 | 0 | 0 | 1 | 2 | 60 | 2 | 1 | 0 | 1 | 2 | 1 | 1 | 1 | 0 | 1 | 1 | 0 | 2 | 1 | 1 |
| FUBP3 | chr9:130619327-130619388\|130619792\|130619641 | 0 | 0 | 0 | 0 | 0 | 0 | 0 | 0 | 0 | 0 | 0 | 1 | 1 | 0 | 0 | 0 | 0 | 0 | 0 | 0 | 0 | 0 | 0 | 0 | 0 | 0 | 0 | 0 | 0 | 1 | 1 | 0 | 0 | 0 | 0 | 30 | 0 | 0 | 0 | 0 | 0 | 0 | 0 | 0 | 0 | 0 | 1 | 0 | 0 | 0 | 0 |
| KIF13A | chr6:17821786-17821905 | 0 | 0 | 0 | 0 | 0 | 0 | 0 | 0 | 0 | 0 | 0 | 0 | 0 | 0 | 0 | 0 | 0 | 0 | 0 | 0 | 0 | 0 | 0 | 0 | 0 | 0 | 0 | 0 | 0 | 2 | 3 | 0 | 0 | 0 | 4 | 72 | 0 | 0 | 0 | 0 | 2 | 0 | 0 | 0 | 0 | 0 | 0 | 0 | 0 | 0 | 0 |
| SPAG9 | chr17:50999476-50999487 | 0 | 0 | 0 | 0 | 0 | 0 | 0 | 0 | 0 | 0 | 0 | 0 | 0 | 0 | 0 | 0 | 0 | 0 | 0 | 0 | 0 | 0 | 0 | 0 | 0 | 0 | 0 | 0 | 0 | 3 | 3 | 0 | 0 | 0 | 2 | 63 | 0 | 0 | 0 | 0 | 1 | 0 | 0 | 0 | 0 | 0 | 0 | 0 | 0 | 0 | 1 |
| GLB1 | chr3:33006041-33006094 | 0 | 0 | 0 | 0 | 0 | 1 | 0 | 0 | 0 | 1 | 1 | 1 | 1 | 1 | 1 | 0 | 0 | 0 | 1 | 0 | 1 | 0 | 0 | 0 | 0 | 0 | 0 | 0 | 0 | 1 | 2 | 0 | 0 | 0 | 0 | 23 | 0 | 0 | 0 | 0 | 0 | 0 | 0 | 0 | 0 | 0 | 1 | 0 | 0 | 0 | 0 |
| DIAPH1 | chr5:141518944-141518996 | 0 | 0 | 0 | 1 | 1 | 2 | 0 | 0 | 2 | 5 | 9 | 3 | 3 | 7 | 8 | 3 | 2 | 6 | 8 | 3 | 4 | 0 | 0 | 1 | 0 | 1 | 2 | 2 | 1 | 1 | 4 | 0 | 0 | 0 | 3 | 55 | 0 | 0 | 0 | 1 | 3 | 1 | 1 | 0 | 0 | 0 | 0 | 0 | 1 | 1 | 0 |
| TNS1 | chr2:217841199-217841285 | 3 | 2 | 0 | 5 | 4 | 5 | 3 | 2 | 1 | 1 | 1 | 2 | 2 | 1 | 1 | 1 | 1 | 1 | 1 | 0 | 1 | 2 | 1 | 4 | 4 | 2 | 2 | 2 | 3 | 2 | 3 | 0 | 1 | 2 | 6 | 73 | 3 | 1 | 1 | 0 | 2 | 2 | 3 | 3 | 0 | 3 | 1 | 1 | 3 | 2 | 4 |
| DAPK2 | chr15:63923036\|63922701-63923343 | 0 | 0 | 0 | 4 | 2 | 3 | 0 | 0 | 0 | 1 | 0 | 3 | 2 | 1 | 1 | 0 | 0 | 0 | 1 | 0 | 0 | 2 | 2 | 1 | 0 | 2 | 0 | 2 | 1 | 3 | 3 | 1 | 2 | 0 | 4 | 57 | 1 | 1 | 1 | 0 | 3 | 0 | 0 | 1 | 0 | 0 | 1 | 1 | 2 | 1 | 0 |
| ANK3 | chr10:60044074-60053727 | 1 | 1 | 1 | 1 | 1 | 2 | 0 | 1 | 0 | 0 | 0 | 0 | 0 | 0 | 0 | 0 | 0 | 0 | 1 | 1 | 0 | 2 | 0 | 1 | 1 | 1 | 1 | 1 | 0 | 6 | 9 | 1 | 1 | 1 | 4 | 82 | 1 | 1 | 1 | 0 | 5 | 1 | 1 | 1 | 1 | 1 | 1 | 0 | 0 | 1 | 2 |
| ZNF516 | chr18:76422935-76426449 | 0 | 0 | 0 | 0 | 0 | 0 | 0 | 0 | 0 | 0 | 0 | 0 | 0 | 0 | 0 | 0 | 0 | 0 | 0 | 0 | 0 | 0 | 0 | 0 | 0 | 0 | 0 | 0 | 0 | 2 | 3 | 0 | 0 | 0 | 0 | 22 | 0 | 0 | 0 | 0 | 0 | 0 | 0 | 0 | 0 | 0 | 0 | 0 | 0 | 0 | 0 |
| PPP1R12A | chr12:79777483-79777652 | 0 | 0 | 0 | 0 | 0 | 0 | 0 | 0 | 0 | 0 | 0 | 0 | 0 | 0 | 0 | 0 | 0 | 0 | 0 | 0 | 0 | 0 | 0 | 0 | 0 | 0 | 0 | 0 | 0 | 6 | 5 | 0 | 0 | 0 | 1 | 52 | 0 | 0 | 0 | 0 | 0 | 0 | 0 | 0 | 0 | 0 | 0 | 0 | 0 | 0 | 0 |
| B4GALT7 | chr5:177602032\|177601426-177602193 | 1 | 1 | 1 | 5 | 4 | 5 | 2 | 1 | 0 | 0 | 0 | 1 | 0 | 0 | 0 | 0 | 0 | 0 | 0 | 0 | 0 | 1 | 5 | 5 | 3 | 2 | 1 | 2 | 2 | 1 | 2 | 1 | 1 | 2 | 2 | 59 | 2 | 1 | 1 | 0 | 3 | 1 | 1 | 2 | 1 | 1 | 2 | 1 | 5 | 2 | 1 |
| FMR1 | chrX:147924919-147925055 | 4 | 3 | 2 | 5 | 4 | 4 | 2 | 3 | 4 | 2 | 2 | 1 | 1 | 1 | 1 | 1 | 3 | 3 | 2 | 4 | 3 | 4 | 2 | 2 | 5 | 3 | 3 | 3 | 4 | 1 | 4 | 6 | 2 | 5 | 4 | 53 | 4 | 2 | 2 | 2 | 2 | 4 | 4 | 4 | 3 | 4 | 1 | 2 | 3 | 3 | 4 |
| BICC1 | chr10:58648624-58648669 | 1 | 1 | 2 | 1 | 1 | 1 | -1 | 1 | -1 | 2 | 1 | 0 | 0 | 4 | 6 | 0 | 6 | 3 | 0 | 0 | -1 | 0 | 0 | 1 | 1 | 2 | 1 | 2 | 0 | 4 | 1 | 2 | 3 | 1 | 1 | 38 | 1 | 0 | 2 | 3 | 1 | 1 | 1 | 2 | 3 | 1 | 1 | 1 | 0 | 1 | 0 |
| USP36 | chr17:78836897-78836976 | 0 | 0 | 0 | 0 | 0 | 1 | 1 | 0 | 0 | 1 | 0 | 1 | 1 | 1 | 1 | 0 | 0 | 1 | 0 | 0 | 0 | 0 | 0 | 1 | 1 | 2 | 0 | 2 | 0 | 2 | 3 | 0 | 0 | 0 | 0 | 28 | 0 | 0 | 0 | 0 | 0 | 0 | 0 | 1 | 0 | 1 | 0 | 0 | 0 | 0 | 0 |
| CCPG1 | chr15:55374155\|55373472-55374250 | 1 | 1 | 0 | 2 | 1 | 2 | 0 | 1 | 3 | 7 | 5 | 8 | 10 | 11 | 8 | 4 | 3 | 12 | 7 | 1 | 1 | 1 | 0 | 2 | 1 | 1 | 1 | 1 | 2 | 2 | 4 | 0 | 0 | 1 | 2 | 62 | 1 | 1 | 0 | 1 | 1 | 1 | 1 | 1 | 0 | 1 | 0 | 0 | 1 | 1 | 1 |
| RPP40 | chr6:5001094-5001130 | 0 | 0 | 0 | 0 | 0 | 0 | 0 | 0 | 2 | 3 | 5 | 4 | 3 | 3 | 5 | 4 | 1 | 5 | 5 | 0 | 1 | 0 | 0 | 0 | 0 | 0 | 0 | 0 | 0 | 2 | 1 | 0 | 0 | 0 | 0 | 21 | 0 | 0 | 0 | 0 | 0 | 0 | 0 | 0 | 0 | 0 | 0 | 0 | 0 | 0 | 0 |
| B4GALT7 | chr5:177602032-177602193\|177603364\|177603900 | 4 | 3 | 2 | 7 | 6 | 7 | 5 | 3 | 1 | 1 | 1 | 3 | 3 | 1 | 1 | 1 | 1 | 1 | 1 | 1 | 1 | 3 | 7 | 7 | 4 | 3 | 3 | 3 | 5 | 2 | 3 | 3 | 2 | 5 | 4 | 58 | 4 | 3 | 2 | 1 | 5 | 3 | 3 | 5 | 4 | 3 | 5 | 3 | 7 | 4 | 2 |
| REEP1 | chr2:86239922-86240071 | 0 | 0 | 0 | 0 | 0 | 0 | 0 | 0 | 0 | 0 | 0 | 0 | 0 | 0 | 0 | 0 | 0 | 0 | 0 | 0 | 0 | 0 | 0 | 0 | 0 | 0 | 0 | 0 | 0 | 7 | 4 | 0 | 1 | 0 | 4 | 51 | 0 | 0 | 0 | 0 | 1 | 0 | 0 | 0 | 0 | 0 | 0 | 0 | 0 | 0 | -1 |
| FN3KRP | chr17:82723060\|82723027-82723111 | 2 | 1 | 0 | 1 | 1 | 1 | 1 | 1 | 0 | 0 | 0 | 1 | 2 | 0 | 0 | 0 | 0 | 0 | 0 | 0 | 0 | 0 | 1 | 1 | 1 | 1 | 1 | 1 | 1 | 2 | 2 | 1 | 1 | 2 | 1 | 23 | 1 | 1 | 1 | 1 | 1 | 2 | 3 | 3 | 3 | 1 | 2 | 1 | 1 | 1 | 2 |
| KDM4C | chr9:7018631-7018679 | 0 | 0 | 1 | 0 | 0 | 0 | 0 | 0 | 0 | 0 | 0 | 0 | 1 | 0 | 0 | 0 | 0 | 0 | 0 | 0 | 0 | 1 | 1 | 0 | 2 | 0 | 0 | 0 | 0 | 1 | 2 | 1 | 3 | 0 | 0 | 24 | 0 | 0 | 0 | 0 | 0 | 0 | 0 | 2 | 0 | 0 | 0 | 1 | 0 | 0 | 0 |
| ITGA7 | chr12:55686202-55686314 | 0 | 0 | 0 | 0 | 0 | 0 | 0 | 0 | 0 | 0 | 0 | 0 | 0 | 0 | 0 | 0 | 0 | 0 | 0 | 0 | 0 | 0 | 0 | 0 | 0 | 0 | 0 | 0 | 0 | 0 | 0 | 0 | 0 | 0 | 8 | 44 | 0 | 0 | 0 | 0 | 0 | 0 | 0 | 0 | 0 | 0 | 0 | 0 | 0 | 0 | 1 |
| CCDC39 | chr3:180641383-180641412 | 1 | 0 | -1 | 1 | 0 | 1 | 0 | 1 | 3 | 3 | 1 | 0 | 0 | 2 | 1 | 0 | 1 | 1 | 1 | 0 | 5 | 0 | 0 | 2 | 1 | 0 | 0 | 1 | 0 | 1 | 2 | 0 | 0 | 0 | 0 | 23 | 1 | 0 | 0 | 1 | 3 | 0 | 0 | 0 | 0 | 1 | 0 | 0 | 0 | 1 | 0 |
| PLB1 | chr2:28611432-28611580 | 0 | 0 | 0 | 1 | 1 | 1 | 0 | 0 | 0 | 1 | 1 | 0 | 0 | 1 | 0 | 0 | 1 | 1 | 1 | 0 | 0 | -1 | 0 | 0 | 1 | 1 | 1 | 1 | 1 | 1 | 2 | 1 | 0 | 0 | 1 | 22 | 0 | 1 | 0 | 0 | 1 | 0 | 0 | 0 | 0 | 1 | 3 | 0 | 1 | 1 | 0 |
| ZNF141 | chr4:342292-342426 | 1 | 2 | 1 | 1 | 0 | 0 | 0 | 1 | 1 | 1 | 0 | 0 | 0 | 0 | 1 | 5 | 0 | 0 | 1 | 1 | 0 | 0 | 0 | 0 | 1 | 1 | 0 | 1 | 0 | 2 | 3 | 0 | 1 | 1 | 0 | 20 | 0 | 1 | 1 | 2 | 0 | 0 | 1 | 0 | 2 | 1 | 2 | 1 | 0 | 1 | 2 |
| SNX9 | chr6:157888571-157888660 | 0 | 0 | 0 | 0 | 0 | 0 | 0 | 0 | 1 | 3 | 4 | 3 | 3 | 5 | 3 | 3 | 2 | 7 | 3 | 1 | 1 | 0 | 0 | 0 | 0 | 0 | 0 | 0 | 1 | 5 | 12 | 0 | 1 | 0 | 1 | 63 | 0 | 0 | 0 | 1 | 1 | 0 | 0 | 0 | 0 | 0 | 0 | 0 | 0 | 0 | 0 |
| ARHGAP11B | chr15:30735540-30735779 | 0 | 0 | 0 | 0 | 0 | 0 | 0 | -1 | 0 | 0 | 0 | 0 | 0 | 0 | 0 | 0 | 0 | 0 | 0 | 0 | 0 | -1 | -1 | 0 | 0 | 0 | 0 | 0 | -1 | 0 | 0 | -1 | 0 | 1 | 0 | 34 | 0 | 0 | 0 | 0 | 0 | 0 | 2 | 0 | -1 | 0 | 6 | 0 | 0 | 0 | 0 |
| GLIS3 | chr9:3932795-3932918 | 0 | 0 | 1 | 0 | 0 | 0 | 2 | 0 | 4 | 10 | 3 | 19 | 9 | 21 | 24 | 5 | 22 | 5 | 4 | 6 | 16 | 0 | 0 | 0 | 1 | 1 | 1 | 1 | 0 | 2 | 5 | 0 | 1 | 0 | 1 | 68 | 0 | 0 | 1 | 4 | 2 | 0 | 0 | 0 | 0 | 0 | 11 | 0 | 0 | 1 | 2 |
| AGPAT5 | chr8:6742533-6742577 | 0 | 0 | 0 | 0 | 0 | 0 | 0 | 0 | 0 | 0 | 0 | 0 | 0 | 0 | 0 | 0 | 0 | 0 | 0 | 0 | 0 | 0 | 0 | 0 | 0 | 0 | 0 | 0 | 0 | 4 | 5 | 0 | 0 | 0 | 1 | 29 | 0 | 0 | 0 | 0 | 1 | 0 | 0 | 0 | 0 | 0 | 0 | 0 | 0 | 0 | 0 |
| PLCXD1 | chrX:288457-288575 | 1 | 1 | 1 | 3 | 2 | 2 | 1 | 1 | 1 | 1 | 0 | 3 | 3 | 2 | 1 | 1 | 1 | 1 | 1 | 2 | 1 | 1 | 1 | 1 | 1 | 1 | 1 | 1 | 0 | 1 | 3 | 1 | 3 | 1 | 1 | 20 | 1 | 0 | 1 | 1 | 1 | 0 | 0 | 1 | 1 | 0 | 1 | 1 | 1 | 1 | 1 |
| TRIM36 | chr5:115159641-115159670 | 0 | 1 | 0 | 0 | 0 | 0 | 0 | 0 | 6 | 8 | 10 | 3 | 6 | 12 | 14 | 13 | 2 | 6 | 6 | 0 | 2 | 0 | 0 | 2 | 2 | 1 | 0 | 1 | 0 | -1 | -1 | -1 | 1 | 0 | 0 | 33 | 0 | 0 | 1 | 2 | 1 | 0 | 0 | 2 | 0 | 1 | 0 | 0 | 0 | 2 | 0 |
| ROCK2 | chr2:11196687\|11196693-11196863 | 0 | 0 | 0 | 0 | 0 | 0 | 0 | 0 | 1 | 0 | 1 | 0 | 0 | 0 | 0 | 0 | 0 | 0 | 0 | 0 | 0 | 0 | 0 | 0 | 0 | 0 | 0 | 0 | 0 | 8 | 16 | 0 | 0 | 0 | 2 | 79 | 0 | 0 | 0 | 0 | 3 | 0 | 0 | 0 | 0 | 0 | 0 | 0 | 0 | 0 | 1 |
| LINC01094 | chr4:78648393-78648475 | 2 | 3 | 3 | 3 | 3 | 2 | -1 | 1 | 4 | 5 | 4 | 4 | 4 | 3 | 3 | 4 | 3 | 3 | 4 | 4 | 3 | 0 | 0 | 4 | 3 | 2 | 1 | 3 | -1 | 3 | 5 | 6 | 0 | 3 | 3 | 49 | 4 | 1 | 2 | 2 | 6 | 1 | 1 | 4 | 7 | 1 | 5 | 1 | 3 | 1 | 6 |
| IFT88 | chr13:20632279-20633028 | 3 | 1 | 0 | 1 | 1 | 1 | 0 | 1 | 2 | 2 | 2 | 1 | 1 | 2 | 2 | 2 | 1 | 2 | 1 | 2 | 1 | 2 | 1 | 1 | 1 | 1 | 0 | 1 | 1 | 4 | 5 | 0 | 0 | 1 | 1 | 33 | 7 | 2 | 0 | 0 | 1 | 1 | 2 | 1 | 0 | 0 | 1 | 0 | 1 | 1 | 0 |
| RANBP2 | chr2:108769260-108769337\|108769867 | 1 | 2 | 1 | 1 | 1 | 1 | 0 | 1 | 1 | 1 | 1 | 1 | 1 | 1 | 1 | 1 | 1 | 1 | 2 | 2 | 1 | 1 | 0 | 1 | 2 | 2 | 1 | 1 | 0 | 7 | 11 | 2 | 2 | 2 | 2 | 45 | 1 | 1 | 1 | 1 | 2 | 1 | 0 | 1 | 1 | 1 | 1 | 1 | 1 | 1 | 2 |
| PHTF2 | chr7:77893982\|77893940\|77893903-77893993 | 0 | 0 | 0 | 0 | 0 | 0 | 0 | 0 | 0 | 1 | 0 | 0 | 0 | 0 | 0 | 0 | 0 | 0 | 1 | 0 | 0 | 0 | 0 | 0 | 0 | 0 | 0 | 0 | 0 | 7 | 14 | 0 | 0 | 0 | 8 | 64 | 1 | 0 | 1 | 0 | 2 | 0 | 0 | 0 | 0 | 0 | 0 | 0 | 0 | 0 | 0 |
| KSR1 | chr17:27621193-27621273\|27622343\|27622143 | 0 | 0 | 1 | 0 | 0 | 0 | 0 | 0 | 17 | 11 | 15 | 14 | 12 | 14 | 10 | 23 | 17 | 10 | 18 | 39 | 24 | 0 | 0 | 0 | 0 | 0 | 0 | 1 | 0 | 1 | 1 | 0 | 0 | 0 | 5 | 79 | 6 | 0 | 0 | 2 | 1 | 0 | 0 | 0 | 0 | 0 | 16 | 1 | 0 | 0 | 0 |
| RTTN | chr18:70128054-70128098 | 3 | 1 | 1 | 2 | 1 | 3 | 0 | 2 | 1 | 1 | 1 | 0 | 1 | 2 | 1 | 0 | 1 | 1 | 1 | 0 | 1 | -1 | 0 | 2 | 1 | 1 | 1 | 1 | -1 | 3 | 3 | 1 | 2 | 2 | 1 | 27 | 1 | 1 | 2 | 1 | 0 | 2 | 5 | 2 | 0 | 2 | 2 | 0 | 0 | 0 | 0 |
| COL4A5 | chrX:108570649-108570795 | 2 | 2 | 2 | 2 | 3 | 2 | 4 | 3 | 1 | 1 | 2 | 2 | 6 | 3 | 4 | 4 | 3 | 2 | 1 | 6 | 2 | 4 | 3 | 4 | 3 | 3 | 4 | 4 | 2 | 2 | 3 | 2 | 4 | 3 | 3 | 30 | 4 | 2 | 5 | 7 | 4 | 4 | 2 | 4 | 2 | 4 | 5 | 3 | 3 | 4 | 0 |
| NOTCH2 | chr1:119950080-119950138 | 1 | 0 | 0 | 0 | 0 | 0 | 1 | 1 | 2 | 0 | 0 | 0 | 0 | 0 | 1 | 1 | 1 | 1 | 0 | 0 | 0 | 0 | 0 | 0 | 0 | 0 | 0 | 0 | 0 | 0 | 1 | 0 | 1 | 1 | 0 | 23 | 1 | 0 | 0 | 0 | 0 | 1 | 1 | 1 | 1 | 0 | 1 | 0 | 0 | 0 | 7 |
| ATL3 | chr11:63635847-63635993 | 3 | 2 | 0 | 3 | 2 | 3 | 2 | 3 | 2 | 3 | 2 | 3 | 3 | 3 | 1 | 3 | 3 | 4 | 2 | 2 | 2 | 2 | 2 | 3 | 2 | 2 | 1 | 3 | 1 | 9 | 8 | 1 | 1 | 5 | 1 | 45 | 7 | 3 | 0 | 1 | 2 | 3 | 3 | 1 | 0 | 1 | 6 | 2 | 3 | 3 | 1 |
| IDH3B | chr20:2659109-2659312 | 0 | 0 | 0 | 0 | 0 | 0 | 0 | 0 | 1 | 1 | 1 | 3 | 3 | 2 | 2 | 2 | 1 | 2 | 1 | 1 | 1 | 0 | 0 | 0 | 0 | 1 | 0 | 1 | 0 | 4 | 5 | 0 | 0 | 0 | 1 | 25 | 0 | 0 | 0 | 0 | 0 | 0 | 0 | 0 | 0 | 0 | 0 | 0 | 0 | 0 | 0 |
| SGMS1 | chr10:50340387-50340464 | 4 | 2 | 2 | 1 | 2 | 2 | 2 | 3 | 1 | 2 | 1 | 5 | 5 | 3 | 2 | 1 | 1 | 2 | 1 | 1 | 1 | 0 | 2 | 2 | 2 | 2 | 1 | 2 | 3 | 3 | 4 | 1 | 1 | 2 | 2 | 24 | 2 | 3 | 3 | 2 | 3 | 4 | 5 | 4 | 3 | 3 | 3 | 1 | 2 | 1 | 4 |
| PRKD3 | chr2:37291523-37292976 | 2 | 2 | 3 | 3 | 2 | 2 | 2 | 2 | 6 | 3 | 4 | 3 | 3 | 2 | 2 | 5 | 3 | 3 | 2 | 5 | 4 | 5 | 1 | 3 | 2 | 3 | 3 | 3 | 2 | 5 | 6 | 3 | 6 | 4 | 4 | 28 | 3 | 3 | 5 | 4 | 4 | 3 | 2 | 5 | 4 | 3 | 2 | 3 | 2 | 4 | 6 |
| TTTY15 | chrY:12681671-12681709 | 0 | 0 | 0 | 0 | 0 | 0 | 0 | 0 | 0 | 0 | 0 | 0 | 0 | 0 | 0 | 0 | 0 | 0 | 0 | 0 | 0 | -1 | 0 | 0 | 0 | 0 | 0 | 0 | -1 | 6 | 11 | 0 | 0 | 0 | 0 | 41 | 0 | 0 | 0 | 0 | 0 | 0 | 0 | 0 | 0 | 0 | 0 | 0 | 0 | -1 | 0 |
| PPP3CC | chr8:22531302-22531328 | 0 | 0 | 0 | 0 | 0 | 0 | 0 | 0 | 1 | 0 | 0 | 0 | 0 | 0 | 0 | 0 | 1 | 0 | 1 | 1 | 0 | 0 | 0 | 0 | 0 | 0 | 0 | 1 | 0 | 13 | 23 | 0 | 0 | 0 | 7 | 90 | 0 | 0 | 0 | 0 | 3 | 0 | 0 | 0 | 0 | 0 | 0 | 0 | 0 | 0 | 0 |
| ANK2 | chr4:113304224-113304277 | 0 | 0 | 0 | 0 | 1 | 0 | 0 | 0 | 0 | 0 | 0 | 0 | 0 | 0 | 0 | 0 | 0 | 0 | 0 | 0 | 0 | 0 | 0 | 0 | 0 | 0 | 0 | 0 | 0 | 16 | 23 | 0 | 1 | 0 | 5 | 93 | 0 | 0 | 1 | 0 | 2 | 0 | 0 | 0 | 0 | 0 | 0 | 0 | 0 | 0 | 8 |
| BTBD10 | chr11:13403911\|13403900-13404015 | 3 | 3 | 1 | 2 | 2 | 1 | 0 | 2 | 2 | 1 | 2 | 1 | 1 | 1 | 1 | 2 | 1 | 1 | 1 | 4 | 2 | 1 | 1 | 1 | 1 | 1 | 1 | 1 | 1 | 8 | 15 | 1 | 1 | 4 | 2 | 59 | 9 | 1 | 0 | 1 | 2 | 2 | 1 | 1 | 1 | 1 | 0 | 2 | 1 | 1 | 1 |
| SNHG14 | chr15:25077219-25077316 | 1 | 1 | 1 | 3 | 2 | 1 | 2 | 1 | 2 | 4 | 3 | 14 | 17 | 6 | 7 | 3 | 4 | 3 | 4 | 1 | 4 | 1 | 1 | 2 | 2 | 3 | 1 | 2 | 1 | 6 | 8 | 0 | 1 | 1 | 4 | 59 | 0 | 1 | 0 | 5 | 2 | 1 | 2 | 1 | 0 | 2 | 0 | 0 | 1 | 2 | 3 |
| CCDC85C | chr14:99528218-99528256 | 2 | 2 | 1 | 3 | 5 | 4 | 7 | 2 | 6 | 7 | 8 | 6 | 6 | 6 | 7 | 8 | 5 | 10 | 9 | 4 | 5 | 3 | 3 | 6 | 4 | 5 | 4 | 6 | 1 | 14 | 17 | 1 | 3 | 3 | 2 | 65 | 4 | 2 | 3 | 3 | 5 | 2 | 2 | 4 | 3 | 3 | 8 | 2 | 3 | 5 | 4 |
| SLC35B3 | chr6:8428975-8428994 | 0 | 0 | 0 | 1 | 0 | 1 | 0 | 1 | 9 | 5 | 7 | 1 | 1 | 5 | 2 | 5 | 3 | 6 | 5 | 13 | 8 | 0 | 0 | 0 | 0 | 0 | 0 | 0 | 0 | 6 | 15 | 0 | 0 | 1 | 0 | 49 | 2 | 0 | 0 | 0 | 0 | 1 | 0 | 0 | 0 | 0 | 2 | 0 | 0 | 0 | 0 |
| RGS12 | chr4:3386525-3386573 | 1 | 1 | 1 | 1 | 1 | 1 | 0 | 1 | 3 | 2 | 11 | 3 | 3 | 1 | 3 | 1 | 3 | 15 | 9 | 1 | 2 | 1 | 0 | 2 | 1 | 2 | 0 | 1 | 0 | 2 | 1 | 1 | 1 | 1 | 1 | 23 | 1 | 1 | 0 | 4 | 1 | 0 | 0 | 1 | 1 | 2 | 0 | 1 | 1 | 0 | 6 |
| USP47 | chr11:11873800-11873859\|11873915 | 7 | 6 | 3 | 5 | 5 | 6 | 5 | 6 | 5 | 5 | 3 | 6 | 6 | 5 | 6 | 4 | 4 | 3 | 4 | 4 | 3 | 5 | 2 | 7 | 6 | 5 | 5 | 5 | 2 | 14 | 26 | 4 | 5 | 5 | 7 | 79 | 5 | 2 | 4 | 2 | 5 | 4 | 5 | 7 | 5 | 6 | 5 | 4 | 4 | 4 | 6 |
| PPP6R1 | chr19:55252094-55252129 | 2 | 1 | 1 | 4 | 4 | 6 | 6 | 2 | 16 | 13 | 24 | 5 | 5 | 13 | 14 | 16 | 5 | 24 | 23 | 2 | 6 | 0 | 3 | 8 | 5 | 7 | 4 | 6 | 4 | 5 | 6 | 1 | 1 | 3 | 3 | 39 | 2 | 1 | 3 | 6 | 4 | 4 | 4 | 3 | 1 | 4 | 2 | 2 | 3 | 4 | 1 |
| ROCK2 | chr2:11184666-11184769\|11186923 | 3 | 4 | 1 | 2 | 2 | 2 | 1 | 3 | 2 | 3 | 3 | 5 | 5 | 3 | 3 | 3 | 3 | 2 | 2 | 1 | 1 | 2 | 2 | 2 | 1 | 1 | 1 | 2 | 4 | 16 | 23 | 1 | 2 | 2 | 4 | 77 | 2 | 1 | 1 | 3 | 4 | 2 | 2 | 2 | 1 | 2 | 4 | 1 | 1 | 1 | 2 |
| PAPD4 | chr5:79680919-79680984 | 0 | 0 | 0 | 0 | 0 | 0 | 0 | 0 | 0 | 0 | 0 | 0 | 0 | 0 | 0 | 0 | 0 | 0 | 0 | 0 | 0 | 0 | 0 | 0 | 0 | 0 | 0 | 0 | 0 | 2 | 8 | 0 | 0 | 0 | 0 | 25 | 0 | 0 | 0 | 0 | 0 | 0 | 0 | 0 | 0 | 0 | 0 | 0 | 0 | 0 | 0 |
| RHBDF1 | chr16:64333-64440\|64437 | 0 | 0 | 0 | 0 | 0 | 0 | 0 | 0 | 1 | 3 | 6 | 25 | 21 | 6 | 6 | 2 | 1 | 8 | 4 | 1 | 1 | 0 | 0 | 0 | 0 | 0 | 0 | 0 | 0 | 5 | 12 | 0 | 0 | 0 | 1 | 71 | 0 | 0 | 0 | 0 | 1 | 0 | 0 | 0 | 0 | 0 | 0 | 0 | 0 | 0 | 4 |
| COPS5 | chr8:67060497-67060545\|67061422 | 2 | 1 | 0 | 1 | 1 | 1 | 1 | 2 | 2 | 1 | 1 | 1 | 1 | 1 | 1 | 2 | 1 | 1 | 1 | 2 | 2 | 1 | 1 | 1 | 0 | 1 | 0 | 1 | 2 | 0 | 0 | 1 | 0 | 1 | 1 | 24 | 3 | 2 | 1 | 1 | 1 | 2 | 2 | 1 | 1 | 1 | 8 | 1 | 1 | 1 | 1 |
| LIMCH1 | chr4:41620424-41633808 | 0 | 0 | 0 | 1 | 1 | 1 | 0 | 0 | 0 | 0 | 0 | 0 | 0 | 0 | 0 | 0 | 0 | 0 | 0 | 0 | 0 | 0 | 0 | 1 | 1 | 1 | 1 | 1 | 0 | 2 | 3 | 0 | 3 | 1 | 21 | 85 | 0 | 0 | 1 | 1 | 5 | 0 | 1 | 1 | 0 | 1 | 2 | 1 | 0 | 1 | 23 |
| COL28A1 | chr7:7417301-7417459 | 1 | 1 | 1 | 1 | 0 | 0 | 1 | 1 | 3 | 4 | 1 | 9 | 10 | 9 | 5 | 2 | 3 | 2 | 2 | 2 | 0 | 0 | 0 | 0 | 0 | 0 | 1 | 0 | 1 | 1 | 3 | 1 | 1 | 1 | 1 | 29 | 1 | 1 | 0 | 2 | 1 | 1 | 1 | 0 | -1 | 0 | 0 | 1 | 1 | 1 | 0 |
| PCNT | chr21:46394504-46394536 | 1 | 0 | 0 | 1 | 1 | 1 | 0 | 1 | 1 | 0 | 1 | 0 | 0 | 1 | 0 | 0 | 1 | 1 | 1 | 1 | 2 | 0 | 0 | 0 | 1 | 0 | 0 | 4 | 0 | 1 | 1 | 0 | 2 | 1 | 31 | 86 | 2 | 1 | 1 | 0 | 4 | 1 | 1 | 1 | 1 | 0 | 0 | 2 | 0 | 0 | 3 |
| C2CD5 | chr12:22512647-22512679 | 0 | 0 | 0 | 0 | 0 | 0 | 0 | 0 | 13 | 29 | 16 | 16 | 15 | 26 | 34 | 16 | 9 | 15 | 15 | 2 | 9 | 1 | 1 | 0 | 0 | 0 | 0 | 0 | 0 | 5 | 6 | 0 | 0 | 0 | 1 | 44 | 0 | 0 | 0 | 1 | 1 | 0 | 0 | 0 | 0 | 0 | 0 | 0 | 0 | 0 | 0 |
| PPFIBP2 | chr11:7617112-7617294\|7617311 | 0 | 0 | 0 | 0 | 0 | 0 | 0 | 0 | 0 | 1 | 0 | 1 | 1 | 0 | 0 | 0 | 0 | 0 | 0 | 0 | 0 | 0 | 0 | 0 | 0 | 0 | 0 | 0 | 0 | 17 | 20 | 0 | 0 | 0 | 1 | 70 | 0 | 0 | 0 | 0 | 0 | 0 | 0 | 0 | 0 | 0 | 0 | 0 | 0 | 0 | 0 |
| AP3B1 | chr5:78003459-78003494 | 6 | 4 | 2 | 5 | 4 | 6 | 5 | 6 | 9 | 10 | 7 | 28 | 23 | 15 | 15 | 14 | 9 | 7 | 10 | 16 | 12 | 8 | 6 | 7 | 6 | 5 | 3 | 4 | 6 | 7 | 11 | 3 | 6 | 4 | 4 | 64 | 6 | 4 | 2 | 1 | 4 | 5 | 5 | 7 | 3 | 5 | 2 | 2 | 8 | 4 | 3 |
| PRKCA | chr17:66796448-66796495 | 0 | 1 | 0 | 1 | 2 | 2 | 4 | 1 | 1 | 1 | 1 | 1 | 1 | 1 | 1 | 1 | 1 | 1 | 1 | 3 | 2 | 0 | 1 | 4 | 2 | 3 | 1 | 3 | 1 | 13 | 29 | 0 | 3 | 1 | 2 | 83 | 3 | 0 | 1 | 1 | 2 | 1 | 1 | 1 | 0 | 2 | 0 | 1 | 2 | 1 | 0 |
| RASA2 | chr3:141559338-141559350 | 0 | 0 | 0 | 0 | 0 | 0 | 1 | 0 | 0 | 0 | 0 | 0 | 0 | 0 | 0 | 0 | 0 | 1 | 0 | 0 | 0 | 0 | 0 | 0 | 0 | 0 | 0 | 0 | 0 | 11 | 23 | 0 | 0 | 0 | 0 | 67 | 0 | 0 | 0 | 0 | 0 | 0 | 0 | 0 | 0 | 0 | 0 | 0 | 0 | 0 | 0 |
| SEMA6C | chr1:151133960-151134055 | 2 | 1 | 2 | 3 | 2 | 3 | 3 | 2 | 1 | 1 | 2 | 3 | 2 | 2 | 2 | 2 | 1 | 2 | 2 | 2 | 2 | 2 | 4 | 2 | 2 | 2 | 2 | 2 | 2 | 3 | 5 | 3 | 4 | 2 | 22 | 65 | 2 | 1 | 3 | 1 | 10 | 2 | 2 | 3 | 1 | 2 | 1 | 2 | 2 | 2 | 14 |
| ADIPOR1 | chr1:202954241-202954307 | 8 | 4 | 4 | 7 | 5 | 9 | 7 | 7 | 8 | 15 | 9 | 21 | 21 | 20 | 22 | 10 | 8 | 8 | 9 | 5 | 6 | 7 | 5 | 8 | 6 | 7 | 3 | 6 | 9 | 13 | 22 | 5 | 4 | 5 | 4 | 72 | 7 | 4 | 3 | 3 | 6 | 7 | 7 | 7 | 3 | 6 | 2 | 3 | 7 | 4 | 2 |
| IQGAP1 | chr15:90431221-90431252 | 0 | 0 | 0 | 0 | 0 | 0 | 0 | 0 | 14 | 17 | 17 | 1 | 4 | 15 | 13 | 13 | 9 | 14 | 12 | 6 | 13 | 0 | 0 | 1 | 1 | 1 | 0 | 1 | 1 | 2 | 6 | 0 | 1 | 0 | 0 | 25 | 7 | 0 | 0 | 0 | 0 | 0 | 0 | 1 | 0 | 1 | 0 | 0 | 0 | 0 | 0 |
| EIF2AK3 | chr2:88612285-88612336 | 0 | 0 | 0 | 0 | 0 | 0 | 0 | 0 | 0 | 0 | 0 | 0 | 0 | 0 | 0 | 1 | 0 | 0 | 0 | 0 | 0 | 0 | 0 | 0 | 0 | 0 | 0 | 0 | 0 | 9 | 16 | 0 | 0 | 0 | 0 | 46 | 0 | 0 | 0 | 0 | 1 | 0 | 0 | 1 | 0 | 0 | 0 | 0 | 0 | 0 | 0 |
| PMS1 | chr2:189785346-189785471 | 6 | 5 | 3 | 7 | 5 | 7 | 6 | 5 | 14 | 16 | 9 | 11 | 10 | 12 | 19 | 10 | 13 | 13 | 9 | 11 | 12 | 6 | 2 | 7 | 5 | 4 | 3 | 6 | 5 | 9 | 13 | 3 | 3 | 5 | 3 | 37 | 7 | 3 | 3 | 3 | 5 | 5 | 5 | 5 | 4 | 4 | 4 | 3 | 5 | 5 | 6 |
| FMNL2 | chr2:152620962-152621091\|152621158\|152625000 | 2 | 3 | 1 | 4 | 5 | 10 | 14 | 2 | 0 | 0 | 0 | 0 | 0 | 0 | 0 | 0 | 0 | 0 | 0 | 0 | 0 | 1 | 2 | 12 | 7 | 9 | 3 | 12 | 3 | 11 | 21 | 2 | 4 | 5 | 5 | 73 | 1 | 1 | 1 | 1 | 6 | 2 | 1 | 4 | 4 | 5 | 1 | 2 | 2 | 3 | 5 |
| FAM91A1 | chr8:123788201-123788247 | 10 | 8 | 5 | 8 | 7 | 10 | 10 | 10 | 9 | 6 | 7 | 3 | 4 | 7 | 5 | 12 | 8 | 8 | 9 | 18 | 14 | 4 | 4 | 9 | 8 | 9 | 4 | 11 | 6 | 17 | 21 | 6 | 7 | 12 | 8 | 63 | 14 | 5 | 7 | 7 | 6 | 13 | 12 | 10 | 9 | 10 | 7 | 5 | 7 | 7 | 12 |
| CCDC93 | chr2:117979080\|117985389-117985406 | 1 | 1 | 0 | 2 | 3 | 4 | 9 | 1 | 2 | 2 | 2 | 3 | 4 | 2 | 2 | 1 | 1 | 3 | 2 | 1 | 2 | 6 | 1 | 5 | 2 | 4 | 1 | 4 | 2 | 18 | 23 | 1 | 1 | 1 | 2 | 70 | 1 | 2 | 1 | 1 | 4 | 1 | 1 | 1 | 1 | 1 | 2 | 1 | 4 | 2 | 1 |
| DNAH11 | chr7:21807539-21807808 | 2 | 1 | 1 | 2 | 0 | 2 | 0 | 0 | 2 | -1 | 1 | 0 | 0 | 0 | 0 | 3 | 1 | 6 | 0 | 3 | -1 | 0 | 0 | 0 | 1 | 1 | 3 | 1 | 3 | 6 | 2 | -1 | 1 | 2 | -1 | 21 | 1 | 3 | 3 | 1 | 0 | 3 | 3 | 2 | 0 | 1 | 0 | 1 | 2 | 0 | 0 |
| HKR1 | chr19:37322438-37322479 | 4 | 3 | 1 | 4 | 4 | 5 | 4 | 4 | 3 | 2 | 5 | 4 | 4 | 3 | 2 | 3 | 2 | 6 | 5 | 2 | 2 | 2 | 1 | 7 | 5 | 5 | 4 | 6 | 5 | 7 | 11 | 3 | 4 | 4 | 3 | 34 | 3 | 2 | 3 | 2 | 3 | 6 | 5 | 8 | 8 | 6 | 4 | 3 | 4 | 4 | 7 |
| ADSS | chr1:244416800-244416916 | 0 | 0 | 0 | 0 | 0 | 0 | 0 | 0 | 0 | 0 | 0 | 0 | 0 | 0 | 0 | 0 | 0 | 0 | 0 | 0 | 0 | 0 | 0 | 0 | 0 | 0 | 0 | 0 | 0 | 7 | 10 | 0 | 0 | 0 | 0 | 29 | 0 | 0 | 0 | 0 | 0 | 0 | 0 | 0 | 0 | 0 | 0 | 0 | 0 | 0 | 0 |
| MEF2D | chr1:156480643-156480777 | 1 | 1 | 0 | 1 | 1 | 1 | 2 | 1 | 0 | 0 | 1 | 0 | 0 | 0 | 0 | 0 | 0 | 0 | 0 | 1 | 0 | 0 | 1 | 2 | 2 | 1 | 1 | 1 | 0 | 18 | 29 | 1 | 1 | 1 | 14 | 86 | 1 | 0 | 1 | 0 | 3 | 0 | 0 | 1 | 1 | 1 | 0 | 1 | 1 | 0 | 1 |
| R3HDM2 | chr12:57288857-57288910 | 2 | 1 | 1 | 1 | 1 | 1 | 1 | 1 | 26 | 49 | 22 | 28 | 26 | 52 | 56 | 22 | 31 | 39 | 21 | 5 | 11 | 0 | 1 | 2 | 1 | 1 | 1 | 1 | 1 | 13 | 13 | 1 | 1 | 1 | 3 | 72 | 2 | 1 | 1 | 3 | 1 | 1 | 1 | 1 | 0 | 1 | 2 | 1 | 1 | 1 | 1 |
| TMEFF1 | chr9:100499272\|100499223-100499303 | 0 | 0 | 0 | 0 | 0 | 0 | 0 | 0 | 2 | 2 | 2 | 3 | 6 | 5 | 2 | 1 | 1 | 1 | 2 | 1 | 2 | 0 | 0 | 0 | 0 | 0 | 0 | 0 | 0 | 5 | 11 | 0 | 0 | 0 | 1 | 35 | 0 | 0 | 0 | 0 | 1 | 0 | 1 | 0 | 0 | 0 | 7 | 0 | 0 | 0 | 0 |
| DNM2 | chr19:10803639-10803668 | 0 | 0 | 0 | 0 | 0 | 0 | 0 | 0 | 1 | 2 | 3 | 11 | 11 | 3 | 5 | 1 | 3 | 3 | 3 | 1 | 1 | 0 | 0 | 0 | 0 | 0 | 0 | 0 | 0 | 11 | 17 | 0 | 0 | 0 | 1 | 52 | 0 | 0 | 0 | 0 | 0 | 0 | 0 | 0 | 0 | 0 | 0 | 0 | 0 | 0 | 0 |
| SPTAN1 | chr9:128555349-128555399 | 0 | 0 | 0 | 0 | 0 | 0 | 0 | 0 | 1 | 4 | 1 | 2 | 1 | 4 | 5 | 1 | 1 | 1 | 2 | 1 | 3 | 0 | 0 | 0 | 0 | 0 | 0 | 0 | 0 | 11 | 11 | 0 | 0 | 0 | 1 | 37 | 0 | 0 | 0 | 0 | 0 | 0 | 0 | 0 | 0 | 0 | 0 | 0 | 0 | 0 | 0 |
| EML4 | chr2:42264919-42264951 | 5 | 4 | 10 | 7 | 7 | 8 | 8 | 5 | 4 | 8 | 4 | 4 | 4 | 7 | 13 | 9 | 2 | 4 | 4 | 2 | 7 | 1 | 7 | 7 | 8 | 7 | 4 | 6 | 5 | 10 | 10 | 12 | 6 | 5 | 9 | 45 | 4 | 10 | 6 | 8 | 8 | 5 | 4 | 10 | 8 | 7 | 6 | 15 | 8 | 7 | 4 |
| RAD54B | chr8:94373012-94373082 | 0 | 0 | 0 | 0 | 0 | 1 | 0 | 0 | 3 | 8 | 8 | 10 | 10 | 9 | 9 | 7 | 3 | 8 | 10 | 2 | 3 | 0 | 0 | 0 | 0 | 0 | 0 | 0 | 0 | 4 | 6 | 0 | 0 | 0 | 0 | 28 | 1 | 0 | 0 | 0 | 0 | 0 | 0 | 0 | 0 | 0 | 3 | 0 | 0 | 0 | 0 |
| RYR3 | chr15:33843488-33843574 | 1 | 3 | 2 | 1 | 2 | 2 | 6 | 1 | 11 | 21 | 66 | 10 | 11 | 20 | 21 | 36 | 39 | 50 | 64 | 17 | 21 | 0 | 1 | 3 | 2 | 3 | 2 | 4 | 1 | 6 | 9 | 1 | -1 | 2 | 8 | 95 | 10 | 1 | -1 | 3 | 3 | 1 | 1 | 4 | 2 | 2 | 41 | 3 | 1 | 2 | -1 |
| AKAP13 | chr15:85676976-85677098 | 0 | 1 | 0 | 1 | 1 | 0 | 1 | 0 | 5 | 5 | 4 | 4 | 1 | 5 | 5 | 3 | 3 | 12 | 4 | 1 | 2 | 0 | 0 | 1 | 0 | 1 | 0 | 1 | 0 | 11 | 22 | 0 | 1 | 0 | 1 | 52 | 0 | 0 | 0 | 1 | 1 | 0 | 0 | 0 | 0 | 0 | 0 | 0 | 0 | 0 | 1 |
| CIZ1 | chr9:128169623-128169700\|128169771 | 1 | 2 | 1 | 1 | 1 | 1 | 1 | 1 | 2 | 1 | 2 | 1 | 1 | 2 | 1 | 2 | 2 | 1 | 2 | 3 | 3 | 1 | 0 | 1 | 1 | 1 | 1 | 1 | 0 | 7 | 11 | 1 | 4 | 1 | 1 | 27 | 2 | 0 | 1 | 1 | 1 | 1 | 1 | 1 | 2 | 1 | 1 | 1 | 1 | 1 | 2 |
| USP54 | chr10:73518189-73518221 | 0 | 0 | 0 | 0 | 0 | 0 | 0 | 0 | 0 | 0 | 0 | 0 | 0 | 0 | 0 | 0 | 0 | 0 | 0 | 0 | 0 | 0 | 0 | 0 | 0 | 0 | 0 | 0 | 0 | 11 | 20 | 0 | 1 | 0 | 1 | 50 | 0 | 0 | 0 | 0 | 0 | 0 | 0 | 0 | 0 | 0 | 0 | 0 | 0 | 0 | 2 |
| CIZ1 | chr9:128169623\|128169604-128169700 | 1 | 2 | 1 | 1 | 1 | 1 | 1 | 1 | 2 | 1 | 2 | 1 | 1 | 1 | 1 | 2 | 2 | 1 | 2 | 3 | 3 | 1 | 0 | 1 | 1 | 1 | 1 | 1 | 0 | 7 | 10 | 1 | 4 | 1 | 1 | 26 | 2 | 0 | 1 | 1 | 1 | 1 | 1 | 1 | 1 | 1 | 1 | 1 | 1 | 1 | 2 |
| RAPH1 | chr2:203441823\|203442598-203442673 | 1 | 1 | 0 | 2 | 2 | 2 | 1 | 1 | 3 | 4 | 1 | 1 | 2 | 0 | 1 | 2 | 0 | 3 | 2 | 3 | 0 | -1 | 0 | 4 | 1 | 2 | 1 | 2 | -1 | 10 | 19 | 1 | 2 | 1 | 1 | 56 | 2 | 3 | 0 | 1 | 3 | 1 | 1 | 1 | 2 | 1 | 17 | 1 | 0 | 1 | 1 |
| ASH2L | chr8:38117452-38117469 | 0 | 0 | 0 | 0 | 0 | 0 | 0 | 0 | 12 | 12 | 12 | 3 | 4 | 11 | 11 | 9 | 8 | 11 | 11 | 4 | 7 | 0 | 0 | 0 | 1 | 0 | 0 | 0 | 1 | 7 | 16 | 0 | 0 | 0 | 1 | 48 | 0 | 0 | 0 | 15 | 0 | 0 | 0 | 1 | 2 | 0 | 2 | 0 | 0 | 0 | 1 |
| MOB1B | chr4:70980999\|70980994-70981070 | 3 | 4 | 2 | 7 | 7 | 5 | 7 | 4 | 5 | 10 | 6 | 8 | 11 | 10 | 12 | 7 | 5 | 9 | 7 | 4 | 4 | -1 | 0 | 6 | 6 | 6 | 4 | 6 | -1 | 9 | 16 | 2 | 4 | 3 | 4 | 46 | 4 | 5 | 4 | 3 | 5 | 2 | 2 | 4 | 3 | 4 | 9 | 2 | 6 | 3 | 2 |
| STRN4 | chr19:46733816-46733875 | 3 | 5 | 2 | 4 | 5 | 3 | 5 | 3 | 12 | 8 | 11 | 5 | 6 | 9 | 7 | 11 | 8 | 12 | 8 | 8 | 8 | 3 | 4 | 7 | 4 | 4 | 3 | 5 | 3 | 7 | 6 | 2 | 5 | 6 | 3 | 39 | 6 | 2 | 3 | 10 | 3 | 4 | 3 | 5 | 9 | 3 | 1 | 3 | 4 | 4 | 17 |
| TAF12 | chr1:28623946\|28623942-28624030 | 0 | 0 | 0 | 0 | 1 | 0 | 0 | 0 | 0 | 0 | 0 | 0 | 0 | 0 | 0 | 0 | 0 | 0 | 0 | 0 | 0 | 0 | 0 | 0 | 0 | 0 | 0 | 0 | 0 | 11 | 33 | 0 | 0 | 0 | 3 | 72 | 0 | 0 | 0 | 0 | 2 | 0 | 0 | 0 | 0 | 0 | 0 | 0 | 0 | 0 | 0 |
| COL23A1 | chr5:178265655-178265726 | 0 | 0 | 0 | 0 | 1 | 0 | 0 | 0 | 0 | 1 | 0 | 0 | 0 | 0 | 0 | 0 | 0 | 0 | 0 | 0 | 0 | 0 | 0 | 0 | 0 | 0 | 0 | 0 | -1 | 6 | 11 | 0 | 0 | 0 | 0 | 26 | 0 | 0 | 0 | 0 | 1 | 0 | 0 | 0 | 0 | 0 | 1 | 0 | 0 | 0 | 0 |
| MEF2C | chr5:88760988-88761125 | 6 | 8 | 5 | 6 | 7 | 5 | 5 | 7 | 2 | 1 | 3 | 2 | 2 | 1 | 1 | 2 | 4 | 4 | 5 | 8 | 6 | 4 | 5 | 6 | 8 | 6 | 9 | 8 | 5 | 11 | 21 | 7 | 7 | 8 | 24 | 68 | 7 | 2 | 8 | 4 | 23 | 7 | 8 | 12 | 11 | 8 | 2 | 7 | 4 | 5 | 10 |

SFig5, pg8

| Gene Symbol | Exon Location (hg38, human) | Adipose - Subcutaneous | Adipose - Visceral (Omentum) | Adrenal Gland | Artery - Aorta | Artery - Coronary | Artery - Tibial | Bladder | Breast - Mammary Tissue | Brain - Amygdala | Brain - Anterior cingulate cortex (BA24) | Brain - Caudate (basal ganglia) | Brain - Cerebellar Hemisphere | Brain - Cerebellum | Brain - Cortex | Brain - Frontal Cortex (BA9) | Brain - Hippocampus | Brain - Hypothalamus | Brain - Nucleus accumbens (basal ganglia) | Brain - Putamen (basal ganglia) | Brain - Spinal cord (cervical c-1) | Brain - Substantia nigra | Cervix - Ectocervix | Cervix - Endocervix | Colon - Sigmoid | Colon - Transverse | Esophagus - Gastroesophageal Junction | Esophagus - Mucosa | Esophagus - Muscularis | Fallopian Tube | Heart - Atrial Appendage | Heart - Left Ventricle | Kidney - Cortex | Liver | Lung | Minor Salivary Gland | Muscle - Skeletal | Nerve - Tibial | Ovary | Pancreas | Pituitary | Prostate | Skin - Not Sun Exposed (Suprapubic) | Skin - Sun Exposed (Lower leg) | Small Intestine - Terminal Ileum | Spleen | Stomach | Testis | Thyroid | Uterus | Vagina | Whole Blood |
|---|---|---|---|---|---|---|---|---|---|---|---|---|---|---|---|---|---|---|---|---|---|---|---|---|---|---|---|---|---|---|---|---|---|---|---|---|---|---|---|---|---|---|---|---|---|---|---|---|---|---|---|---|
| COL4A5 | chrX:108536002-108536053 | 2 | 2 | 2 | 2 | 2 | 2 | 3 | 3 | 3 | 1 | 2 | 3 | 5 | 1 | 2 | 3 | 4 | 2 | 1 | 2 | 8 | 3 | 2 | 2 | 4 | 4 | 4 | 2 | 2 | 14 | 19 | 2 | 2 | 4 | 3 | 44 | 2 | 1 | 1 | 4 | 3 | 3 | 2 | 3 | 2 | 3 | 2 | 2 | 3 | 3 | -1 |
| RP11-176H8.1 | chr14:31340450-31340476 | 1 | 2 | 1 | 2 | 2 | 2 | 2 | 1 | 2 | 3 | 2 | 8 | 7 | 3 | 5 | 1 | 5 | 1 | 1 | 1 | 5 | 5 | 0 | 4 | 2 | 2 | 0 | 2 | 2 | 25 | 30 | 0 | 0 | 1 | 1 | 74 | 1 | 2 | 1 | 2 | 1 | 2 | 1 | 3 | 1 | 2 | 1 | 1 | 1 | 2 | 1 |
| USP6 | chr17:5119968-5120139 | 1 | 2 | 0 | 2 | 4 | 2 | -1 | 2 | -1 | 2 | 0 | 0 | -1 | -1 | -1 | 0 | 2 | 0 | 0 | 0 | 0 | -1 | -1 | 3 | 2 | 1 | 2 | 3 | 0 | 6 | 8 | 2 | 0 | 1 | 2 | 25 | 2 | 2 | 1 | 0 | 1 | 1 | 1 | 1 | 4 | 1 | 3 | 0 | 4 | 5 | 0 |
| CDC14A | chr1:100383466-100383510\|100383640 | 2 | 2 | 1 | 7 | 6 | 12 | 4 | 2 | 2 | 3 | 2 | 15 | 3 | 1 | 3 | 2 | 2 | 2 | 2 | 5 | 1 | -1 | -1 | 19 | 8 | 8 | 2 | 9 | 2 | 23 | 17 | 3 | 3 | 3 | 2 | 66 | 4 | 3 | 1 | 1 | 3 | 2 | 1 | 3 | 1 | 3 | 2 | 1 | 8 | 4 | 2 |
| FBXW2 | chr9:120792887-120793015 | 3 | 2 | 2 | 2 | 2 | 2 | 1 | 2 | 3 | 2 | 4 | 2 | 2 | 1 | 3 | 4 | 2 | 3 | 4 | 4 | 4 | 2 | 1 | 2 | 2 | 2 | 2 | 2 | 2 | 8 | 13 | 2 | 1 | 2 | 2 | 27 | 3 | 1 | 2 | 3 | 2 | 2 | 2 | 2 | 2 | 2 | 4 | 2 | 2 | 3 | 1 |
| ZNRF2 | chr7:30350761-30350837 | 0 | 0 | 0 | 0 | 0 | 0 | 0 | 0 | 4 | 8 | 2 | 24 | 25 | 13 | 14 | 4 | 2 | 3 | 2 | 1 | 1 | 1 | 0 | 0 | 0 | 0 | 0 | 0 | 0 | 1 | 2 | 0 | 0 | 0 | 1 | 47 | 0 | 0 | 0 | 0 | 1 | 0 | 0 | 0 | 0 | 0 | 0 | 0 | 1 | 0 | 0 |
| SEPN1 | chr1:25802016-25802117 | 10 | 11 | 10 | 10 | 11 | 12 | 13 | 10 | 17 | 17 | 16 | 10 | 12 | 16 | 18 | 19 | 16 | 17 | 15 | 18 | 19 | 11 | 10 | 15 | 13 | 12 | 12 | 13 | 11 | 12 | 14 | 10 | 10 | 10 | 10 | 31 | 11 | 11 | 17 | 10 | 12 | 10 | 10 | 12 | 9 | 13 | 13 | 12 | 11 | 11 | 12 |
| DENND1A | chr9:123411776-123411829 | 14 | 8 | 2 | 10 | 11 | 10 | 6 | 10 | 2 | 2 | 4 | 2 | 1 | 2 | 2 | 3 | 1 | 4 | 4 | 2 | 2 | 10 | 7 | 8 | 4 | 8 | 10 | 7 | 7 | 25 | 32 | 3 | 5 | 9 | 11 | 82 | 17 | 6 | 1 | 3 | 5 | 15 | 19 | 4 | 6 | 5 | 2 | 3 | 12 | 8 | 9 |
| YWHAE | chr17:1347962-1348008 | 0 | 0 | 0 | 0 | 1 | 1 | 0 | 0 | 1 | 2 | 1 | 1 | 1 | 3 | 3 | 2 | 1 | 1 | 1 | 1 | 1 | 0 | 0 | 0 | 0 | 0 | 0 | 1 | 0 | 26 | 40 | 0 | 0 | 0 | 8 | 90 | 1 | 0 | 0 | 0 | 3 | 0 | 0 | 0 | 0 | 0 | 0 | 0 | 0 | 0 | 1 |
| PRPF18 | chr10:13597645-13597671 | 0 | 0 | 0 | 0 | 0 | 0 | 0 | 0 | 23 | 35 | 28 | 41 | 41 | 42 | 52 | 25 | 6 | 38 | 26 | 4 | 6 | 0 | 0 | 0 | 0 | 0 | 0 | 0 | 0 | 9 | 13 | 0 | 0 | 0 | 2 | 81 | 0 | 0 | 0 | 0 | 1 | 0 | 0 | 0 | 0 | 0 | 0 | 0 | 0 | 0 | 0 |
| SDHAP3 | chr5:1571328-1571440 | 1 | 2 | 2 | 1 | 1 | 2 | 0 | 2 | 6 | 6 | 19 | 4 | 5 | 8 | 5 | 10 | 10 | 27 | 19 | 2 | 9 | -1 | 0 | 4 | 2 | 3 | 1 | 3 | 0 | 4 | 7 | 0 | 1 | 3 | 1 | 25 | 1 | 3 | 3 | 1 | 2 | 1 | 1 | 3 | 10 | 2 | 2 | 1 | 1 | 3 | 4 |
| OSBPL2 | chr20:62280009-62280122 | 6 | 3 | 1 | 9 | 7 | 8 | 5 | 5 | 0 | 0 | 1 | 3 | 4 | 1 | 0 | 1 | 0 | 0 | 0 | 0 | 0 | 4 | 4 | 10 | 6 | 5 | 3 | 5 | 5 | 14 | 19 | 2 | 4 | 5 | 4 | 42 | 4 | 2 | 7 | 2 | 4 | 7 | 8 | 5 | 4 | 4 | 3 | 3 | 6 | 4 | 3 |
| PRDM2 | chr1:13769118\|13769039-13769173 | 7 | 6 | 13 | 9 | 10 | 12 | 12 | 6 | 17 | 16 | 15 | 14 | 13 | 17 | 17 | 22 | 19 | 12 | 16 | 25 | 22 | 5 | 4 | 10 | 8 | 9 | 11 | 10 | 6 | 18 | 29 | 10 | 11 | 6 | 10 | 59 | 13 | 4 | 8 | 4 | 8 | 8 | 10 | 8 | 7 | 11 | 14 | 8 | 5 | 5 | 4 |
| FSD1L | chr9:105513599-105513631 | 13 | 8 | 7 | 7 | 9 | 9 | 0 | 9 | 46 | 42 | 54 | 17 | 19 | 41 | 41 | 42 | 36 | 46 | 36 | 31 | 28 | 0 | 0 | 8 | 2 | 7 | 5 | 11 | 5 | 23 | 32 | 3 | 6 | 13 | 14 | 95 | 40 | 9 | 4 | 8 | 11 | 11 | 12 | 7 | 9 | 5 | 10 | 5 | 3 | 10 | 13 |
| DSCR3 | chr21:37266792-37266847 | 0 | 0 | 0 | 0 | 0 | 0 | 0 | 0 | 2 | 3 | 2 | 0 | 0 | 1 | 1 | 3 | 2 | 2 | 2 | 3 | 2 | 1 | 0 | 0 | 0 | 0 | 0 | 0 | 0 | 6 | 10 | 0 | 0 | 0 | 0 | 21 | 0 | 0 | 0 | 0 | 0 | 0 | 0 | 0 | 0 | 0 | 1 | 0 | 0 | 0 | 0 |
| PMS2P3 | chr7:75524054-75524256 | 2 | 3 | 2 | 3 | 3 | 3 | 3 | 2 | 5 | 5 | 4 | 9 | 10 | 6 | 5 | 5 | 5 | 5 | 5 | 6 | 4 | 1 | 3 | 2 | 2 | 3 | 1 | 3 | 10 | 6 | 7 | 2 | 2 | 3 | 2 | 26 | 5 | 2 | 2 | 4 | 3 | 1 | 1 | 1 | 1 | 3 | 3 | 2 | 3 | 2 | 1 |
| MYO5A | chr15:52415116-52415202\|52415179 | 1 | 1 | 0 | 0 | 0 | 1 | 0 | 1 | 0 | 0 | 0 | 0 | 0 | 0 | 0 | 0 | 0 | 0 | 0 | 0 | 0 | -1 | 0 | 1 | 1 | 0 | 0 | 1 | 0 | 29 | 35 | 0 | 2 | 1 | 3 | 79 | 1 | 0 | 1 | 0 | 0 | 0 | 0 | 1 | 1 | 1 | 1 | 1 | 1 | 0 | 0 |
| ITPR1 | chr3:4709692-4709729 | 0 | 0 | 0 | 0 | 0 | 0 | 0 | 0 | 0 | 0 | 0 | 0 | 0 | 0 | 0 | 0 | 1 | 0 | 0 | 1 | 1 | 1 | 0 | 0 | 0 | 0 | 0 | 0 | 0 | 9 | 7 | 0 | 0 | 0 | 0 | 20 | 0 | 0 | 0 | 0 | 0 | 0 | 0 | 0 | 0 | 0 | 0 | 0 | 0 | 0 | 0 |
| ABCC9 | chr12:21805136-21805311 | 0 | 0 | 0 | 0 | 1 | 0 | 0 | 0 | 1 | 1 | 0 | 0 | 2 | 2 | 3 | 0 | 1 | 1 | 0 | 1 | 2 | 0 | 1 | 0 | 0 | 0 | 0 | 0 | 0 | 21 | 36 | 0 | 0 | 0 | 5 | 82 | 0 | 0 | 0 | 0 | 1 | 0 | 0 | 0 | 1 | 0 | 1 | 0 | 0 | 0 | 24 |
| SSRP1 | chr11:57330733-57330741 | 1 | 0 | 0 | 3 | 3 | 4 | 8 | 0 | 1 | 3 | 4 | 2 | 2 | 5 | 5 | 2 | 1 | 3 | 4 | 0 | 0 | 1 | 1 | 12 | 5 | 7 | 1 | 9 | 2 | 26 | 41 | 0 | 0 | 0 | 2 | 85 | 0 | 1 | 0 | 0 | 3 | 0 | 0 | 2 | 0 | 2 | 0 | 0 | 2 | 1 | 0 |
| KCNQ5 | chr6:73129793-73129849 | 0 | -1 | 0 | 0 | 0 | 1 | 0 | -1 | 0 | 0 | 0 | 0 | -1 | 0 | 0 | 0 | 0 | 0 | 0 | 0 | 0 | 0 | 0 | 0 | -1 | 0 | 0 | 1 | 0 | -1 | 0 | 0 | 0 | 1 | 20 | 36 | 3 | -1 | -1 | 0 | 0 | -1 | 3 | 0 | 0 | 1 | 1 | 0 | 0 | 0 | 0 |
| FAM179A | chr2:29046647-29046718 | 1 | 1 | 1 | 1 | 1 | 2 | 1 | 1 | 1 | 1 | 1 | 1 | 0 | 3 | 3 | 1 | 1 | 2 | 2 | 0 | 0 | 0 | 0 | 2 | 1 | 2 | 1 | 2 | 0 | 3 | 4 | 0 | 1 | 0 | 25 | 44 | 2 | 1 | 1 | 1 | 2 | 1 | 0 | 1 | 1 | 2 | 0 | 2 | 1 | 1 | 0 |
| ZNF267 | chr16:31890207-31890250 | 4 | 7 | 7 | 6 | 6 | 5 | 6 | 6 | 18 | 17 | 16 | 3 | 4 | 13 | 10 | 11 | 14 | 14 | 16 | 11 | 15 | 5 | -1 | 5 | 3 | 7 | 12 | 9 | 7 | 22 | 34 | 6 | 5 | 4 | 8 | 70 | 6 | 18 | 5 | 6 | 7 | 10 | 9 | 3 | 2 | 5 | 12 | 9 | 6 | 7 | 0 |
| SCN3B | chr11:123641342-123641817\|123641407 | 8 | 5 | 2 | 2 | 1 | 5 | -1 | 4 | 1 | 0 | 0 | 6 | 9 | 0 | 0 | 0 | 0 | 0 | 1 | 0 | 0 | -1 | -1 | 3 | 3 | 2 | 3 | 3 | -1 | 4 | 2 | -1 | 0 | 4 | 7 | 34 | 6 | 6 | 2 | 2 | 3 | 9 | 8 | 2 | -1 | 7 | 8 | 4 | 4 | 2 | 0 |
| CEP250 | chr20:35469566\|35469495-35469665 | 5 | 3 | 1 | 10 | 10 | 11 | 8 | 6 | 2 | 2 | 3 | 9 | 8 | 3 | 4 | 2 | 4 | 5 | 2 | 1 | 1 | 8 | -1 | 18 | 9 | 12 | 2 | 13 | 6 | 16 | 18 | 1 | 1 | 2 | 3 | 56 | 4 | 4 | 1 | 8 | 6 | 2 | 2 | 5 | 2 | 5 | 3 | 2 | 8 | 3 | 1 |
| IMPDH2 | chr3:49025995-49026066 | 4 | 3 | 2 | 8 | 6 | 7 | 8 | 4 | 3 | 3 | 4 | 3 | 3 | 3 | 3 | 2 | 2 | 3 | 4 | 2 | 3 | 5 | 5 | 10 | 5 | 8 | 1 | 9 | 5 | 18 | 36 | 4 | 2 | 3 | 6 | 67 | 3 | 6 | 2 | 2 | 5 | 1 | 2 | 4 | 2 | 5 | 2 | 3 | 7 | 3 | 1 |
| LATS1 | chr6:149697092-149697186 | 1 | 1 | 1 | 2 | 2 | 1 | 1 | 0 | 1 | 4 | 4 | 9 | 8 | 7 | 8 | 4 | 1 | 3 | 3 | 1 | 1 | 0 | 1 | 2 | 3 | 1 | 3 | 1 | 2 | 4 | 10 | 2 | 6 | 1 | 2 | 27 | 0 | 0 | 2 | 2 | 1 | 1 | 1 | 3 | 1 | 2 | 10 | 1 | 0 | 1 | 1 |
| CA5BP1 | chrX:15681001-15681047 | 7 | 10 | 9 | 9 | 9 | 9 | 7 | 9 | 9 | 6 | 9 | 9 | 9 | 6 | 6 | 9 | 9 | 7 | 9 | 13 | 14 | 13 | 9 | 7 | 8 | 7 | 6 | 7 | 9 | 7 | 10 | 9 | 10 | 10 | 7 | 24 | 10 | 8 | 8 | 7 | 8 | 6 | 6 | 8 | 8 | 8 | 10 | 8 | 10 | 7 | 11 |
| UBR3 | chr2:169947905-169947937 | 2 | 1 | 0 | 2 | 2 | 2 | 1 | 2 | 3 | 1 | 3 | 1 | 1 | 2 | 1 | 5 | 3 | 1 | 5 | 5 | 5 | 1 | 0 | 1 | 1 | 0 | 1 | 1 | 0 | 28 | 53 | 1 | 3 | 3 | 11 | 98 | 3 | 0 | 1 | 0 | 3 | 3 | 2 | 1 | 1 | 1 | 1 | 0 | 1 | 1 | 2 |
| LATS1 | chr6:149681461-149681523 | 1 | 2 | 3 | 4 | 4 | 3 | 8 | 1 | 1 | 1 | 1 | 2 | 2 | 2 | 2 | 2 | 1 | 1 | 3 | 1 | 2 | 7 | 3 | 7 | 4 | 4 | 1 | 5 | 4 | 23 | 35 | 2 | 1 | 1 | 1 | 68 | 2 | 4 | 0 | 1 | 2 | 1 | 0 | 2 | 1 | 3 | 0 | 1 | 4 | 3 | 0 |
| MAP4 | chr3:47909038-47912421 | 0 | 1 | 0 | 2 | 2 | 2 | 4 | 0 | 0 | 0 | 0 | 0 | 1 | 0 | 0 | 0 | 0 | 1 | 0 | 0 | 0 | 1 | 1 | 3 | 3 | 1 | 1 | 1 | 1 | 29 | 48 | 0 | 1 | 1 | 8 | 90 | 0 | 0 | 1 | 0 | 3 | 0 | 0 | 2 | 1 | 1 | 7 | 1 | 2 | 1 | 3 |
| TKT | chr3:53243583-53243622 | 1 | 1 | 0 | 4 | 3 | 3 | 3 | 1 | 0 | 0 | 0 | 0 | 0 | 0 | 0 | 0 | 0 | 0 | 0 | 0 | 0 | 1 | 2 | 6 | 2 | 3 | 0 | 3 | 2 | 15 | 25 | 0 | 0 | 1 | 0 | 47 | 3 | 1 | 0 | 0 | 3 | 1 | 1 | 1 | 0 | 1 | 1 | 0 | 3 | 1 | 0 |
| EXOC1 | chr4:55872755\|55872787-55872808 | 12 | 11 | 14 | 6 | 9 | 4 | 3 | 8 | 50 | 26 | 46 | 9 | 11 | 28 | 18 | 47 | 26 | 40 | 52 | 78 | 53 | 1 | 4 | 7 | 4 | 5 | 1 | 5 | 2 | 17 | 32 | 4 | 4 | 14 | 4 | 82 | 21 | 2 | 3 | 6 | 5 | 2 | 1 | 5 | 21 | 4 | 8 | 2 | 2 | 2 | 29 |
| SNX6 | chr14:34604135-34604268 | 7 | 5 | 4 | 6 | 5 | 8 | 7 | 7 | 17 | 15 | 14 | 13 | 14 | 15 | 11 | 19 | 15 | 13 | 16 | 16 | 19 | 6 | 7 | 7 | 4 | 6 | 3 | 7 | 8 | 16 | 22 | 6 | 5 | 6 | 5 | 43 | 11 | 11 | 3 | 9 | 10 | 5 | 6 | 6 | 6 | 5 | 7 | 5 | 8 | 7 | 4 |
| TCF7L1 | chr2:85308611-85308673 | 0 | 0 | 0 | 0 | 0 | 0 | 0 | 0 | 2 | 2 | 3 | 19 | 13 | 3 | 3 | 3 | 3 | 1 | 3 | 1 | 2 | 0 | 0 | 0 | 0 | 0 | 0 | 0 | 0 | 4 | 3 | 0 | 0 | 0 | 1 | 22 | 2 | 0 | 0 | 0 | 0 | 0 | 1 | 0 | 0 | 0 | 2 | 0 | 0 | 0 | 1 |
| LRIG3 | chr12:58880369-58880478 | 3 | 1 | 1 | 3 | 3 | 5 | 9 | 2 | 13 | 24 | 11 | 10 | 14 | 22 | 24 | 15 | 17 | 13 | 15 | 13 | 15 | 2 | 2 | 5 | 2 | 7 | 1 | 9 | 1 | 8 | 30 | 0 | 1 | 2 | 1 | 56 | 5 | 1 | 1 | 5 | 4 | 1 | 1 | 2 | 0 | 2 | 8 | 0 | 1 | 2 | 2 |
| KIAA1217 | chr10:24367119-24367199\|24367168 | 1 | 1 | 1 | 1 | 2 | 2 | 0 | 1 | 4 | 1 | 3 | 1 | 1 | 1 | 1 | 1 | 2 | 1 | 1 | 5 | 2 | -1 | 0 | 3 | 3 | 2 | 1 | 3 | 3 | 26 | 22 | 1 | 2 | 1 | 2 | 54 | 3 | 1 | 1 | 0 | 2 | 1 | 0 | 1 | 1 | 2 | 2 | 1 | 1 | 1 | -1 |
| USP40 | chr2:233488781-233488903 | 1 | 1 | 0 | 1 | 1 | 2 | 0 | 1 | 4 | 9 | 4 | 19 | 17 | 10 | 10 | 6 | 7 | 7 | 6 | 4 | 4 | 0 | 0 | 1 | 1 | 1 | 0 | 1 | 1 | 9 | 12 | 0 | 0 | 1 | 1 | 35 | 2 | 0 | 0 | 1 | 1 | 1 | 1 | 1 | 0 | 1 | 1 | 0 | 1 | 1 | 1 |
| WDR11 | chr10:120887210-120887268\|120887271 | 1 | 1 | 0 | 1 | 2 | 1 | 1 | 1 | 17 | 10 | 14 | 2 | 4 | 11 | 7 | 16 | 8 | 12 | 17 | 18 | 16 | 0 | 0 | 2 | 1 | 2 | 0 | 2 | 1 | 15 | 22 | 0 | 1 | 1 | 1 | 43 | 9 | 1 | 0 | 1 | 1 | 0 | 0 | 1 | 1 | 1 | 1 | 0 | 0 | 1 | 0 |
| DMXL1 | chr5:119178575-119178688 | 2 | 3 | 1 | 3 | 3 | 3 | 2 | 2 | 14 | 19 | 12 | 12 | 8 | 21 | 29 | 14 | 17 | 13 | 13 | 18 | 20 | 2 | 1 | 3 | 4 | 3 | 2 | 3 | 4 | 17 | 19 | 2 | 1 | 3 | 2 | 47 | 6 | 4 | 2 | 7 | 2 | 2 | 2 | 4 | 3 | 3 | 20 | 2 | 3 | 2 | 4 |
| ADARB1 | chr21:45205660-45205752\|45207388 | 4 | 4 | 4 | 3 | 4 | 4 | 6 | 4 | 1 | 1 | 1 | 2 | 2 | 1 | 1 | 1 | 1 | 1 | 1 | 4 | 2 | 4 | 4 | 8 | 7 | 6 | 5 | 7 | 5 | 30 | 26 | 3 | 5 | 3 | 7 | 61 | 5 | 4 | 5 | 3 | 7 | 4 | 3 | 5 | 4 | 5 | 17 | 4 | 3 | 5 | 4 |
| PHKA1 | chrX:72601991-72602029 | 15 | 8 | 2 | 13 | 10 | 11 | 6 | 10 | 6 | 8 | 6 | 3 | 4 | 6 | 9 | 8 | 5 | 6 | 6 | 6 | 6 | 3 | 6 | 8 | 7 | 6 | 4 | 6 | 6 | 17 | 38 | 3 | 11 | 5 | 18 | 82 | 13 | 2 | 3 | 1 | 6 | 7 | 12 | 11 | 8 | 5 | 4 | 3 | 5 | 3 | 34 |
| ACO1 | chr9:32386371-32386438 | 5 | 6 | 7 | 9 | 8 | 11 | 9 | 6 | 9 | 9 | 9 | 17 | 17 | 9 | 9 | 9 | 10 | 15 | 9 | 5 | 5 | 7 | 12 | 8 | 7 | 8 | 5 | 6 | 10 | 6 | 7 | 9 | 4 | 9 | 7 | 34 | 9 | 10 | 9 | 11 | 9 | 7 | 7 | 8 | 6 | 6 | 16 | 8 | 11 | 9 | 3 |
| ZNF789 | chr7:99485208-99485368 | 7 | 5 | 7 | 6 | 6 | 8 | 3 | 5 | 18 | 19 | 18 | 16 | 15 | 16 | 20 | 20 | 21 | 21 | 21 | 30 | 18 | 3 | 2 | 4 | 3 | 5 | 4 | 6 | 2 | 16 | 16 | 5 | 3 | 4 | 5 | 40 | 9 | 3 | 2 | 5 | 3 | 4 | 5 | 3 | 2 | 5 | 12 | 5 | 3 | 3 | 3 |
| FBXW2 | chr9:120776931-120777021 | 6 | 5 | 2 | 4 | 3 | 5 | 9 | 5 | 3 | 3 | 2 | 7 | 8 | 3 | 3 | 3 | 2 | 2 | 3 | 5 | 3 | 6 | 2 | 7 | 5 | 5 | 2 | 6 | 4 | 3 | 3 | 2 | 2 | 5 | 3 | 21 | 5 | 3 | 2 | 1 | 4 | 5 | 4 | 6 | 4 | 5 | 4 | 2 | 2 | 3 | 4 |
| CDC42BPB | chr14:102956240-102956290 | 1 | 1 | 2 | 0 | 1 | 1 | 1 | 1 | 23 | 14 | 27 | 8 | 8 | 13 | 11 | 27 | 26 | 22 | 32 | 47 | 35 | 1 | 1 | 1 | 1 | 1 | 0 | 1 | 1 | 11 | 17 | 2 | 1 | 1 | 1 | 33 | 6 | 1 | 2 | 5 | 1 | 0 | 0 | 1 | 1 | 1 | 2 | 1 | 1 | 0 | 2 |
| KIF13B | chr8:29141765-29141937 | 0 | 0 | 0 | 0 | 0 | 0 | 0 | 0 | 1 | 2 | 2 | 13 | 20 | 5 | 5 | 2 | 2 | 1 | 1 | 3 | 2 | 0 | 0 | 0 | 0 | 0 | 0 | 0 | 0 | 8 | 19 | 0 | 0 | 0 | 0 | 44 | 0 | 0 | 0 | 1 | 0 | 0 | 0 | 0 | 0 | 0 | 0 | 0 | 0 | 0 | 0 |
| NSMCE4A | chr10:121973523-121973599 | 4 | 2 | 2 | 3 | 3 | 3 | 3 | 3 | 8 | 9 | 7 | 15 | 14 | 10 | 9 | 6 | 7 | 6 | 6 | 6 | 7 | 3 | 2 | 3 | 2 | 3 | 2 | 2 | 2 | 4 | 6 | 2 | 3 | 3 | 3 | 23 | 4 | 2 | 3 | 3 | 3 | 4 | 4 | 3 | 1 | 3 | 4 | 3 | 2 | 2 | 1 |
| EXOC1 | chr4:55872755-55872808\|55877757 | 14 | 14 | 14 | 8 | 10 | 5 | 3 | 10 | 51 | 27 | 48 | 10 | 13 | 30 | 19 | 47 | 26 | 41 | 53 | 79 | 53 | 3 | 4 | 8 | 5 | 6 | 2 | 7 | 2 | 19 | 33 | 5 | 8 | 18 | 4 | 83 | 25 | 4 | 5 | 8 | 7 | 3 | 3 | 6 | 23 | 5 | 10 | 4 | 3 | 3 | 32 |
| INPP5B | chr1:37883661-37883732\|37883867 | 2 | 6 | 1 | 3 | 3 | 3 | 3 | 2 | 8 | 6 | 7 | 9 | 11 | 6 | 6 | 4 | 5 | 6 | 6 | 3 | 4 | 4 | 8 | 7 | 4 | 6 | 3 | 6 | 1 | 15 | 10 | 2 | 8 | 3 | 2 | 34 | 7 | 8 | 2 | 6 | 6 | 1 | 1 | 2 | 2 | 2 | 3 | 4 | 6 | 5 | 2 |
| MGME1 | chr20:17972707-17972751 | 5 | 8 | 2 | 10 | 10 | 8 | 9 | 7 | 1 | 1 | 1 | 1 | 1 | 2 | 1 | 1 | 1 | 0 | 1 | 0 | 1 | 10 | 6 | 6 | 3 | 8 | 16 | 8 | 6 | 23 | 31 | 4 | 9 | 10 | 12 | 63 | 4 | 5 | 2 | 1 | 9 | 13 | 12 | 4 | 2 | 6 | 1 | 8 | 6 | 12 | 1 |
| PPP1R12B | chr1:202575002-202575182 | 0 | 0 | 0 | 0 | 0 | 0 | 0 | 0 | 3 | 17 | 5 | 21 | 27 | 17 | 26 | 5 | 9 | 4 | 5 | 10 | 10 | 0 | 0 | 0 | 0 | 0 | 0 | 0 | 0 | 10 | 16 | 0 | 0 | 0 | 2 | 48 | 0 | 0 | 0 | 0 | 1 | 1 | 0 | 0 | 0 | 0 | 3 | 0 | 0 | 0 | 1 |
| ABCD3 | chr1:94472173-94472242 | 0 | 0 | 0 | 0 | 0 | 0 | 0 | 0 | 0 | 0 | 0 | 0 | 0 | 0 | 0 | 0 | 0 | 0 | 0 | 0 | 0 | 0 | 0 | 0 | 0 | 0 | 0 | 0 | 0 | 11 | 20 | 0 | 0 | 0 | 1 | 34 | 0 | 0 | 0 | 0 | 0 | 0 | 0 | 0 | 0 | 0 | 0 | 0 | 0 | 0 | 0 |
| ACTR2 | chr2:65242049-65242063\|65246307 | 0 | 1 | 0 | 0 | 0 | 0 | 1 | 0 | 2 | 7 | 2 | 8 | 17 | 10 | 6 | 3 | 3 | 1 | 2 | 1 | 3 | 0 | 0 | 0 | 0 | 0 | 0 | 0 | 1 | 5 | 5 | 0 | 1 | 1 | 0 | 27 | 0 | 0 | 0 | 0 | 1 | 1 | 0 | 1 | 0 | 0 | 1 | 0 | 0 | 0 | 0 |
| PCNT | chr21:46430759-46431329\|46430973\|46430809 | 4 | 3 | 2 | 5 | 4 | 4 | 6 | 4 | 2 | 1 | 2 | 3 | 3 | 1 | 1 | 2 | 1 | 1 | 2 | 3 | 2 | 6 | 3 | 7 | 7 | 4 | 7 | 4 | 3 | 4 | 3 | 2 | 3 | 5 | 17 | 34 | 3 | 2 | 2 | 2 | 4 | 12 | 14 | 6 | 4 | 5 | 5 | 3 | 4 | 8 | 7 |
| ZIK1 | chr19:57589457-57589706 | 0 | 0 | 0 | 0 | 0 | 0 | 0 | 0 | 0 | 0 | 0 | 1 | 1 | 0 | 1 | 0 | 0 | 0 | 0 | 0 | 0 | 0 | 0 | 0 | 0 | 0 | 0 | 0 | 0 | 14 | 24 | 0 | 0 | 0 | 0 | 42 | 0 | 0 | 0 | 0 | 0 | 0 | 0 | 0 | 0 | 0 | 6 | 0 | 0 | 0 | 0 |
| RNF115 | chr1:145787012-145787063 | 7 | 5 | 3 | 7 | 8 | 8 | 4 | 6 | 6 | 5 | 5 | 2 | 3 | 6 | 7 | 5 | 4 | 4 | 5 | 5 | 7 | 6 | 4 | 8 | 7 | 6 | 6 | 6 | 4 | 33 | 47 | 6 | 21 | 8 | 9 | 83 | 10 | 3 | 5 | 4 | 6 | 8 | 8 | 6 | 4 | 6 | 7 | 5 | 5 | 6 | 5 |
| BIN1 | chr2:127060596-127060640 | 4 | 2 | 0 | 9 | 5 | 13 | 6 | 4 | 0 | 1 | 1 | 1 | 1 | 1 | 1 | 0 | 0 | 1 | 1 | 0 | 0 | 6 | 5 | 6 | 4 | 4 | 4 | 7 | 3 | 6 | 16 | 0 | 2 | 3 | 63 | 99 | 3 | 1 | 2 | 1 | 21 | 6 | 8 | 2 | 1 | 3 | 3 | 4 | 6 | 5 | 6 |
| LGR5 | chr12:71462532-71462654 | 6 | 6 | 2 | 3 | 2 | 4 | 0 | 5 | 8 | 10 | 10 | 15 | 5 | 10 | 7 | 7 | 8 | 12 | 6 | 8 | 6 | 0 | 0 | 3 | 3 | 8 | 3 | 3 | 8 | 5 | -1 | 0 | 5 | 4 | 4 | 26 | 8 | 5 | -1 | 5 | 4 | 4 | 5 | 3 | 0 | 2 | -1 | 3 | 6 | 5 | -1 |
| PRKD3 | chr2:37275443-37275524 | 2 | 1 | 1 | 1 | 2 | 2 | 2 | 2 | 1 | 1 | 2 | 1 | 4 | 2 | 1 | 2 | 2 | 1 | 3 | 2 | 3 | 0 | 2 | 2 | 2 | 2 | 1 | 2 | 1 | 4 | 5 | 2 | 1 | 1 | 1 | 22 | 4 | 3 | 1 | 1 | 2 | 2 | 2 | 3 | 1 | 2 | 14 | 1 | 2 | 2 | 0 |
| ARHGAP19 | chr10:97265242-97265334 | 2 | 1 | 0 | 1 | 4 | 1 | 2 | 1 | 10 | 11 | 11 | 2 | 2 | 10 | 9 | 9 | 6 | 9 | 12 | 8 | 7 | 0 | 0 | 5 | 2 | 2 | 0 | 3 | 0 | 5 | 7 | 2 | 1 | 1 | 4 | 28 | 17 | 1 | 0 | 1 | 1 | 1 | 1 | 0 | 1 | 1 | 0 | 1 | 1 | 1 | 1 |
| BSDC1 | chr1:32369230-32369338 | 3 | 4 | 1 | 3 | 3 | 3 | 3 | 3 | 5 | 3 | 5 | 3 | 3 | 4 | 3 | 5 | 4 | 4 | 6 | 7 | 6 | 4 | 4 | 4 | 3 | 3 | 2 | 4 | 4 | 10 | 12 | 2 | 2 | 3 | 1 | 22 | 6 | 4 | 1 | 2 | 2 | 1 | 1 | 2 | 2 | 2 | 3 | 2 | 4 | 3 | 2 |
| RAP1GDS1 | chr4:98387436-98387503\|98387507 | 6 | 5 | 3 | 14 | 10 | 13 | 10 | 5 | 3 | 3 | 3 | 7 | 7 | 3 | 3 | 2 | 3 | 3 | 3 | 1 | 2 | 13 | 15 | 16 | 8 | 13 | 3 | 12 | 11 | 11 | 17 | 4 | 5 | 4 | 4 | 48 | 4 | 13 | 3 | 6 | 10 | 3 | 3 | 7 | 4 | 7 | 3 | 5 | 17 | 8 | 1 |
| COL4A4 | chr2:227144653-227144833 | 3 | 3 | 1 | 2 | 2 | 2 | -1 | 2 | 7 | 7 | 2 | 6 | 8 | 8 | 5 | 8 | 14 | 3 | 5 | 4 | 3 | 0 | 0 | 1 | 1 | 2 | 1 | 1 | -1 | 3 | 2 | 1 | 1 | 2 | 6 | 21 | 3 | 3 | 0 | 5 | 3 | 3 | 2 | 2 | 3 | 2 | 7 | 1 | 1 | 5 | 7 |
| ACAP2 | chr3:195312947-195312967 | 6 | 7 | 2 | 4 | 6 | 3 | 6 | 9 | 33 | 32 | 45 | 3 | 7 | 29 | 23 | 23 | 25 | 38 | 37 | 15 | 25 | 7 | 4 | 7 | 3 | 5 | 1 | 5 | 2 | 22 | 27 | 5 | 1 | 5 | 2 | 57 | 24 | 6 | 1 | 5 | 5 | 2 | 2 | 3 | 3 | 3 | 6 | 4 | 4 | 5 | 1 |
| L3MBTL2 | chr22:41227519-41227679\|41227574 | 5 | 5 | 3 | 7 | 6 | 6 | 4 | 5 | 5 | 4 | 4 | 9 | 10 | 4 | 3 | 4 | 5 | 4 | 4 | 7 | 6 | 8 | 12 | 5 | 4 | 4 | 3 | 4 | 5 | 8 | 6 | 3 | 5 | 7 | 4 | 24 | 9 | 7 | 5 | 7 | 5 | 5 | 6 | 4 | 6 | 3 | 7 | 5 | 6 | 7 | 7 |
| RALGAPB | chr20:38520240-38520275 | 0 | 0 | 0 | 0 | 0 | 0 | 0 | 0 | 12 | 21 | 10 | 34 | 34 | 27 | 36 | 10 | 18 | 28 | 14 | 3 | 9 | 0 | 0 | 0 | 0 | 0 | 0 | 0 | 0 | 2 | 3 | 0 | 0 | 0 | 1 | 48 | 0 | 0 | 0 | 0 | 0 | 0 | 0 | 0 | 0 | 0 | 0 | 0 | 0 | 0 | 0 |
| DIP2B | chr12:50707914-50708048 | 0 | 0 | 0 | 0 | 0 | 0 | 0 | 0 | 1 | 2 | 1 | 3 | 3 | 2 | 2 | 1 | 1 | 1 | 1 | 0 | 1 | 0 | 0 | 0 | 0 | 0 | 0 | 0 | 0 | 7 | 12 | 0 | 0 | 0 | 0 | 20 | 0 | 0 | 0 | 0 | 0 | 0 | 0 | 0 | 0 | 0 | 0 | 0 | 0 | 0 | 0 |
| MAP4 | chr3:47987383-47987433 | 2 | 2 | 1 | 2 | 3 | 3 | 3 | 2 | 7 | 9 | 7 | 10 | 10 | 9 | 12 | 7 | 7 | 7 | 7 | 7 | 7 | 1 | 1 | 3 | 3 | 3 | 1 | 3 | 2 | 17 | 28 | 1 | 2 | 2 | 4 | 46 | 2 | 1 | 1 | 2 | 2 | 1 | 1 | 2 | 1 | 3 | 1 | 1 | 2 | 2 | 3 |
| MFF | chr2:227342745-227342819 | 27 | 23 | 17 | 20 | 22 | 20 | 18 | 24 | 13 | 12 | 13 | 7 | 8 | 12 | 10 | 13 | 11 | 11 | 14 | 14 | 14 | 24 | 18 | 18 | 18 | 18 | 18 | 17 | 21 | 49 | 45 | 17 | 17 | 21 | 23 | 82 | 25 | 18 | 23 | 15 | 20 | 20 | 21 | 20 | 24 | 21 | 17 | 16 | 20 | 20 | 22 |
| AKAP2 | chr9:110162094-110162132 | 0 | 0 | 0 | 0 | 0 | 0 | 0 | 0 | 7 | 31 | 7 | 20 | 5 | 28 | 45 | 10 | 22 | 15 | 5 | 2 | 13 | 0 | 0 | 0 | 0 | 0 | 0 | 0 | 0 | 8 | 6 | 0 | 0 | 0 | 0 | 16 | 0 | 0 | 0 | 0 | 0 | 0 | 0 | 0 | 0 | 0 | 0 | 0 | 0 | 0 | 0 |
| PLPP1 | chr5:55490960-55491114 | 9 | 9 | 7 | 14 | 12 | 12 | 10 | 9 | 13 | 15 | 15 | 18 | 18 | 15 | 17 | 13 | 13 | 15 | 14 | 10 | 13 | 9 | 11 | 10 | 9 | 9 | 9 | 9 | 12 | 18 | 35 | 7 | 8 | 12 | 11 | 54 | 12 | 10 | 10 | 12 | 9 | 11 | 11 | 9 | 7 | 9 | 21 | 9 | 11 | 10 | 9 |
| SLC30A7 | chr1:100910370-100910412 | 1 | 1 | 1 | 0 | 1 | 0 | 0 | 1 | 17 | 9 | 11 | 2 | 2 | 11 | 6 | 19 | 9 | 15 | 14 | 19 | 15 | 0 | 0 | 1 | 0 | 1 | 0 | 1 | 2 | 10 | 19 | 0 | 0 | 0 | 1 | 30 | 2 | 0 | 1 | 1 | 1 | 0 | 0 | 0 | 0 | 0 | 5 | 0 | 1 | 1 | 0 |
| COPS5 | chr8:67060497\|67060321-67060545 | 2 | 1 | 0 | 1 | 1 | 1 | 1 | 1 | 3 | 1 | 1 | 1 | 1 | 1 | 1 | 1 | 2 | 1 | 2 | 2 | 3 | 2 | 2 | 1 | 1 | 0 | 0 | 0 | 2 | 0 | 0 | 1 | 0 | 1 | 1 | 24 | 3 | 2 | 0 | 1 | 1 | 1 | 1 | 1 | 0 | 0 | 17 | 1 | 1 | 1 | 1 |
| PKP4 | chr2:158533438-158533534 | 1 | 1 | 1 | 2 | 2 | 2 | 2 | 1 | 5 | 19 | 11 | 6 | 7 | 19 | 19 | 7 | 4 | 18 | 6 | 1 | 1 | 1 | 1 | 2 | 1 | 2 | 1 | 2 | 2 | 15 | 18 | 1 | 1 | 3 | 1 | 33 | 6 | 2 | 1 | 3 | 1 | 1 | 1 | 2 | 2 | 1 | 3 | 1 | 1 | 1 | 6 |
| GAS2 | chr11:22731331-22731425\|22731396 | 0 | 0 | 1 | 1 | 1 | 0 | 0 | 0 | 14 | 41 | 31 | 14 | 21 | 45 | 42 | 17 | 41 | 39 | 17 | 1 | 2 | 0 | 0 | 6 | 2 | 0 | 0 | 1 | 0 | 40 | 35 | 0 | 0 | 1 | 4 | 79 | 0 | 0 | 0 | 13 | 1 | 0 | 0 | 1 | 1 | 1 | 6 | 1 | 1 | 0 | 4 |
| IWS1 | chr2:127485874-127486033 | 0 | 0 | 0 | 1 | 2 | 2 | 2 | 0 | 1 | 1 | 2 | 2 | 2 | 1 | 1 | 1 | 0 | 3 | 2 | 0 | 0 | 0 | 0 | 2 | 1 | 1 | 0 | 2 | 1 | 32 | 47 | 0 | 0 | 0 | 2 | 77 | 0 | 0 | 0 | 0 | 1 | 0 | 0 | 1 | 0 | 1 | 1 | 0 | 1 | 1 | 0 |
| MAX | chr14:65077328\|65077344-65077428 | 8 | 7 | 3 | 8 | 6 | 9 | 10 | 8 | 12 | 12 | 11 | 11 | 12 | 15 | 14 | 12 | 9 | 9 | 10 | 10 | 14 | 6 | 8 | 9 | 7 | 8 | 7 | 8 | 6 | 6 | 7 | 6 | 10 | 13 | 5 | 39 | 8 | 6 | 14 | 3 | 7 | 7 | 7 | 9 | 11 | 9 | 23 | 5 | 9 | 8 | 19 |
| PAAF1 | chr11:73889172\|73888435\|73889122\|73889087-73889245 | 0 | 0 | 0 | 0 | 0 | 0 | 0 | 0 | 12 | 15 | 15 | 4 | 6 | 12 | 10 | 9 | 7 | 13 | 12 | 5 | 6 | 1 | 0 | 1 | 0 | 0 | 0 | 0 | 0 | 22 | 39 | 0 | 0 | 0 | 2 | 61 | 4 | 0 | 0 | 1 | 0 | 0 | 0 | 0 | 0 | 0 | 0 | 0 | 0 | 0 | 0 |
| PLCE1 | chr10:94282791\|94282794-94282832 | 3 | 1 | 3 | 2 | 2 | 4 | 2 | 3 | 13 | 32 | 10 | 9 | 8 | 19 | 30 | 13 | 16 | 10 | 14 | 7 | 9 | 2 | 0 | 4 | 6 | 3 | 1 | 3 | 3 | 19 | 10 | 1 | 1 | 1 | 3 | 32 | 2 | 2 | 2 | 2 | 3 | 2 | 5 | 7 | 1 | 5 | 6 | 10 | 2 | 3 | -1 |
| CRYZL1 | chr21:33641179-33641305 | 1 | 1 | 1 | 1 | 1 | 1 | 0 | 1 | 4 | 8 | 6 | 20 | 18 | 11 | 16 | 6 | 8 | 7 | 5 | 5 | 5 | 0 | 0 | 1 | 1 | 1 | 1 | 1 | 1 | 11 | 21 | 1 | 1 | 1 | 2 | 42 | 2 | 1 | 1 | 2 | 1 | 1 | 1 | 1 | 1 | 1 | 13 | 1 | 1 | 1 | 2 |
| PLCE1 | chr10:94282791-94282832\|94282940 | 3 | 1 | 2 | 1 | 1 | 3 | 2 | 2 | 12 | 27 | 8 | 8 | 6 | 17 | 24 | 10 | 13 | 7 | 11 | 6 | 7 | 1 | 0 | 3 | 5 | 2 | 2 | 2 | 1 | 16 | 9 | 1 | 1 | 1 | 3 | 27 | 2 | 1 | 2 | 2 | 3 | 2 | 4 | 6 | 1 | 4 | 6 | 9 | 2 | 2 | -1 |
| RP11-400K9.4 | chr8:53521209-53521379 | 3 | 4 | 7 | 6 | 6 | 5 | 2 | 4 | 2 | 3 | 5 | 7 | 3 | 4 | 4 | 3 | 4 | 7 | 6 | 4 | 3 | -1 | -1 | 5 | 3 | 3 | 3 | 4 | 0 | 5 | 18 | -1 | -1 | 3 | 6 | 32 | 4 | 4 | 5 | 10 | 5 | 4 | 4 | 1 | 10 | 4 | 5 | 4 | 2 | 3 | 0 |
| KCNMA1 | chr10:76974487-76974573 | 13 | 10 | 7 | 9 | 10 | 9 | 12 | 12 | 12 | 14 | 13 | 10 | 13 | 12 | 12 | 13 | 11 | 14 | 12 | 12 | 14 | 12 | 22 | 12 | 13 | 9 | 9 | 8 | 17 | 14 | 15 | 5 | 10 | 11 | 8 | 84 | 15 | 14 | 10 | 12 | 11 | 11 | 11 | 11 | 8 | 9 | 63 | 14 | 20 | 13 | 10 |
| MYO6 | chr6:75867431-75867523 | 0 | 0 | 0 | 0 | 1 | 1 | 0 | 0 | 4 | 6 | 5 | 24 | 30 | 13 | 10 | 6 | 10 | 9 | 5 | 5 | 7 | 0 | 0 | 2 | 1 | 1 | 0 | 1 | 0 | 10 | 13 | 0 | 0 | 0 | 0 | 44 | 1 | 0 | 0 | 1 | 1 | 0 | 0 | 0 | 0 | 0 | 3 | 0 | 0 | 0 | 0 |
| NAP1L1 | chr12:76057744-76057776 | 0 | 0 | 0 | 0 | 0 | 0 | 0 | 0 | 4 | 8 | 5 | 5 | 6 | 9 | 8 | 4 | 3 | 4 | 4 | 1 | 2 | 0 | 0 | 0 | 0 | 0 | 0 | 0 | 0 | 7 | 13 | 0 | 0 | 0 | 1 | 20 | 0 | 0 | 0 | 0 | 1 | 0 | 0 | 0 | 0 | 0 | 0 | 0 | 0 | 0 | 0 |
| ATP11B | chr3:182844514-182844593 | 12 | 9 | 4 | 4 | 6 | 2 | 1 | 9 | 40 | 16 | 34 | 4 | 5 | 26 | 15 | 40 | 27 | 31 | 31 | 46 | 51 | 3 | 3 | 6 | 2 | 5 | 0 | 4 | 3 | 8 | 9 | 4 | 5 | 12 | 2 | 48 | 33 | 1 | 0 | 2 | 2 | 2 | 1 | 4 | 7 | 1 | 6 | 3 | 2 | 2 | 7 |
| OSBPL9 | chr1:51740143-51740211\|51742030 | 0 | 0 | 0 | 0 | 0 | 0 | 0 | 0 | 7 | 16 | 7 | 45 | 38 | 20 | 26 | 12 | 12 | 8 | 9 | 15 | 12 | 0 | 0 | 0 | 0 | 0 | 0 | 0 | 0 | 14 | 29 | 0 | 0 | 0 | 2 | 63 | 0 | 0 | 0 | 1 | 1 | 0 | 0 | 0 | 0 | 0 | 4 | 0 | 0 | 0 | 1 |
| HSPG2 | chr1:21893589-21893663 | 3 | 1 | 1 | 2 | 2 | 3 | 2 | 2 | 2 | 8 | 6 | 4 | 3 | 3 | 2 | 4 | 6 | 5 | 4 | 4 | 4 | 3 | 2 | 3 | 2 | 3 | 2 | 3 | 2 | 19 | 30 | 2 | 1 | 1 | 2 | 44 | 6 | 1 | 1 | 1 | 2 | 1 | 2 | 2 | 1 | 2 | 1 | 1 | 1 | 2 | 5 |
| PRR11 | chr17:59163599-59163664 | 4 | 4 | 4 | 3 | 3 | 2 | -1 | 5 | 5 | 6 | 7 | -1 | 1 | 4 | 13 | 7 | 4 | 1 | 7 | 11 | 3 | 0 | 0 | 5 | 3 | 9 | 3 | 5 | -1 | 5 | 11 | -1 | 2 | 3 | 2 | 23 | 5 | 5 | 3 | -1 | 5 | 2 | 3 | 5 | 4 | 6 | 6 | 4 | 1 | 3 | 4 |
| MAX | chr14:65077344-65077428\|65077643 | 7 | 6 | 3 | 7 | 5 | 8 | 8 | 7 | 10 | 10 | 9 | 8 | 9 | 11 | 11 | 10 | 7 | 7 | 9 | 9 | 12 | 5 | 7 | 8 | 6 | 7 | 6 | 7 | 5 | 5 | 6 | 6 | 9 | 12 | 5 | 32 | 7 | 5 | 12 | 3 | 6 | 6 | 6 | 8 | 10 | 8 | 20 | 4 | 7 | 7 | 18 |
| EXOG | chr3:38513750\|38513773-38513868 | 0 | 1 | 0 | 0 | 1 | 0 | 0 | 1 | 2 | 2 | 1 | 13 | 12 | 3 | 3 | 2 | 3 | 2 | 2 | 4 | 3 | 1 | 0 | 1 | 0 | 1 | 0 | 1 | 0 | 9 | 14 | 0 | 1 | 0 | 1 | 31 | 1 | 0 | 1 | 0 | 1 | 0 | 1 | 1 | 1 | 1 | 16 | 0 | 0 | 1 | 0 |
| GPATCH8 | chr17:44407217-44407266 | 1 | 1 | 1 | 0 | 0 | 0 | 1 | 1 | 13 | 19 | 11 | 18 | 19 | 18 | 16 | 9 | 9 | 12 | 11 | 7 | 7 | 0 | 0 | 1 | 0 | 0 | 1 | 0 | 1 | 8 | 7 | 0 | 1 | 1 | 1 | 27 | 3 | 0 | 1 | 2 | 1 | 1 | 1 | 1 | 0 | 1 | 6 | 1 | 0 | 1 | 1 |
| TMED2 | chr12:123587611-123587631 | 0 | 0 | 0 | 0 | 0 | 0 | 0 | 0 | 1 | 0 | 0 | 0 | 0 | 0 | 0 | 0 | 0 | 0 | 1 | 0 | 0 | 0 | 0 | 0 | 0 | 0 | 0 | 0 | 0 | 38 | 58 | 0 | 0 | 0 | 1 | 85 | 0 | 0 | 0 | 0 | 1 | 0 | 0 | 0 | 0 | 0 | 0 | 0 | 0 | 0 | 0 |
| USP25 | chr21:15847663-15849872 | 7 | 6 | 5 | 8 | 8 | 10 | 8 | 7 | 5 | 5 | 4 | 4 | 4 | 5 | 4 | 6 | 4 | 5 | 5 | 6 | 5 | 6 | 5 | 7 | 7 | 7 | 5 | 8 | 9 | 33 | 41 | 5 | 5 | 5 | 7 | 61 | 8 | 6 | 5 | 4 | 7 | 5 | 6 | 7 | 5 | 7 | 1 | 5 | 6 | 5 | 3 |
| PTBP3 | chr9:112330441-112330474 | 2 | 3 | 4 | 3 | 3 | 3 | 3 | 2 | 15 | 17 | 17 | 21 | 22 | 17 | 22 | 10 | 11 | 26 | 23 | 5 | 11 | 4 | 7 | 4 | 2 | 3 | 3 | 4 | 5 | 14 | 24 | 3 | 3 | 2 | 2 | 46 | 3 | 1 | 2 | 4 | 4 | 2 | 2 | 2 | 3 | 3 | 22 | 3 | 3 | 2 | 2 |
| MYO5A | chr15:52331699-52331836 | 3 | 5 | 2 | 4 | 5 | 3 | 1 | 4 | 11 | 5 | 15 | 2 | 2 | 6 | 3 | 16 | 7 | 8 | 17 | 33 | 17 | 6 | 2 | 13 | 7 | 7 | 3 | 6 | 2 | 15 | 20 | 4 | 6 | 5 | 3 | 34 | 14 | 8 | 2 | 2 | 6 | 2 | 2 | 5 | 4 | 4 | 2 | 2 | 5 | 6 | 6 |
| FRMD4B | chr3:69173042-69173133 | 1 | 1 | 2 | 1 | 1 | 0 | 1 | 0 | 2 | 1 | 1 | 1 | 2 | 2 | 1 | 2 | 2 | 1 | 2 | 2 | 2 | 0 | 0 | 1 | 0 | 1 | 0 | 1 | 1 | 29 | 27 | 1 | 0 | 0 | 0 | 47 | 1 | 0 | 0 | 0 | 0 | 0 | 0 | 1 | 2 | 1 | 1 | 0 | 0 | 0 | 0 |
| ARHGAP32 | chr11:128979780-128979855 | 1 | 2 | 0 | 4 | 3 | 2 | 1 | 1 | 8 | 4 | 8 | 4 | 6 | 4 | 3 | 7 | 8 | 5 | 9 | 10 | 10 | 3 | -1 | 4 | 1 | 3 | 1 | 3 | 1 | 27 | 36 | 0 | 0 | 1 | 1 | 54 | 8 | 6 | 1 | 2 | 1 | 1 | 0 | 0 | 1 | 2 | 3 | 1 | 3 | 1 | 1 |
| INTU | chr4:127664741-127664824 | 10 | 4 | 2 | 6 | 7 | 9 | 2 | 6 | 6 | 8 | 6 | 21 | 8 | 9 | 7 | 18 | 13 | 6 | 8 | 46 | 42 | -1 | 5 | 3 | 4 | 3 | 2 | 3 | 0 | 8 | 10 | 5 | 3 | 4 | 3 | 24 | 6 | 1 | 2 | 2 | 2 | 7 | 10 | 2 | 6 | 3 | 4 | 2 | 3 | 4 | 0 |
| AGFG1 | chr2:227550059-227550106 | 4 | 5 | 3 | 4 | 4 | 3 | 5 | 4 | 24 | 50 | 25 | 63 | 61 | 53 | 61 | 27 | 34 | 42 | 25 | 4 | 14 | 2 | 2 | 6 | 4 | 5 | 1 | 5 | 5 | 16 | 26 | 2 | 2 | 6 | 4 | 80 | 9 | 4 | 1 | 8 | 3 | 2 | 2 | 5 | 9 | 3 | 0 | 2 | 4 | 3 | 11 |
| AP1S2 | chrX:15840497-15840591 | 0 | 0 | 0 | 0 | 2 | 0 | 0 | 0 | 2 | 3 | 1 | 1 | 1 | 2 | 2 | 2 | 0 | 1 | 1 | 1 | 1 | 0 | 0 | 0 | 0 | 0 | 0 | 0 | 0 | 44 | 64 | 0 | 1 | 0 | 14 | 93 | 0 | 0 | 1 | 0 | 8 | 0 | 0 | 0 | 0 | 0 | 0 | 1 | 0 | 0 | 0 |
| MIA3 | chr1:222662458-222662529 | 2 | 3 | 3 | 3 | 3 | 2 | 2 | 3 | 6 | 6 | 7 | 7 | 7 | 7 | 7 | 6 | 5 | 7 | 6 | 4 | 6 | 4 | 2 | 3 | 5 | 3 | 16 | 2 | 3 | 36 | 63 | 4 | 14 | 8 | 6 | 88 | 4 | 2 | 2 | 3 | 4 | 16 | 13 | 4 | 2 | 6 | 1 | 3 | 2 | 6 | 1 |
| PBRM1 | chr3:52624897-52624941 | 7 | 6 | 7 | 9 | 7 | 8 | 7 | 7 | 20 | 30 | 13 | 57 | 46 | 30 | 39 | 19 | 27 | 21 | 13 | 14 | 19 | 5 | 8 | 10 | 10 | 6 | 4 | 6 | 9 | 23 | 30 | 6 | 6 | 4 | 10 | 65 | 9 | 6 | 9 | 4 | 8 | 7 | 6 | 9 | 4 | 11 | 5 | 9 | 9 | 7 | 3 |
| TXNL1 | chr18:56604436-56604478\|56605384 | 0 | 0 | 0 | 0 | 1 | 0 | 0 | 0 | 0 | 0 | 0 | 0 | 0 | 0 | 0 | 0 | 0 | 0 | 1 | 0 | 0 | 0 | 0 | 0 | 0 | 0 | 0 | 0 | 0 | 38 | 54 | 0 | 0 | 0 | 3 | 78 | 1 | 0 | 0 | 0 | 1 | 0 | 0 | 0 | 0 | 0 | 0 | 0 | 0 | 0 | 0 |
| YWHAB | chr20:44887648-44887742 | 10 | 14 | 11 | 13 | 14 | 12 | 13 | 12 | 8 | 8 | 9 | 13 | 13 | 9 | 9 | 9 | 7 | 7 | 9 | 9 | 9 | 15 | 16 | 14 | 14 | 13 | 12 | 13 | 10 | 14 | 16 | 13 | 16 | 13 | 11 | 26 | 11 | 13 | 12 | 11 | 13 | 10 | 8 | 15 | 13 | 14 | 15 | 13 | 14 | 13 | 9 |
| UBFD1 | chr16:23564039-23564109\|23564732\|23564089\|23565244 | 29 | 15 | 10 | 22 | 19 | 23 | 15 | 24 | 7 | 5 | 8 | 17 | 20 | 9 | 7 | 8 | 4 | 7 | 9 | 7 | 5 | 21 | 13 | 16 | 19 | 15 | 11 | 15 | 19 | 19 | 20 | 14 | 21 | 21 | 20 | 52 | 23 | 12 | 19 | 7 | 17 | 22 | 26 | 26 | 23 | 18 | 12 | 13 | 15 | 15 | 16 |

SFig5, pg9

| Gene Symbol | Exon Location (hg38, human) | Adipose - Subcutaneous | Adipose - Visceral (Omentum) | Adrenal Gland | Artery - Aorta | Artery - Coronary | Artery - Tibial | Bladder | Breast - Mammary Tissue | Brain - Amygdala | Brain - Anterior cingulate cortex (BA24) | Brain - Caudate (basal ganglia) | Brain - Cerebellar Hemisphere | Brain - Cerebellum | Brain - Cortex | Brain - Frontal Cortex (BA9) | Brain - Hippocampus | Brain - Hypothalamus | Brain - Nucleus accumbens (basal ganglia) | Brain - Putamen (basal ganglia) | Brain - Spinal cord (cervical c-1) | Brain - Substantia nigra | Cervix - Ectocervix | Cervix - Endocervix | Colon - Sigmoid | Colon - Transverse | Esophagus - Gastroesophageal Junction | Esophagus - Mucosa | Esophagus - Muscularis | Fallopian Tube | Heart - Atrial Appendage | Heart - Left Ventricle | Kidney - Cortex | Liver | Lung | Minor Salivary Gland | Muscle - Skeletal | Nerve - Tibial | Ovary | Pancreas | Pituitary | Prostate | Skin - Not Sun Exposed (Suprapubic) | Skin - Sun Exposed (Lower leg) | Small Intestine - Terminal Ileum | Spleen | Stomach | Testis | Thyroid | Uterus | Vagina | Whole Blood |
|---|---|---|---|---|---|---|---|---|---|---|---|---|---|---|---|---|---|---|---|---|---|---|---|---|---|---|---|---|---|---|---|---|---|---|---|---|---|---|---|---|---|---|---|---|---|---|---|---|---|---|---|---|
| DIP2C | chr10:311537\|311335-311566 | 23 | 16 | 4 | 37 | 29 | 33 | 14 | 21 | 6 | 5 | 8 | 4 | 3 | 5 | 6 | 7 | 6 | 6 | 8 | 6 | 8 | 31 | 38 | 16 | 15 | 13 | 15 | 11 | 23 | 10 | 5 | 8 | 13 | 22 | 19 | 92 | 39 | 23 | 9 | 12 | 17 | 31 | 36 | 10 | 10 | 12 | 5 | 13 | 33 | 27 | 11 |
| GAK | chr4:859362-859493 | 4 | 4 | 2 | 6 | 5 | 4 | 4 | 4 | 5 | 7 | 5 | 27 | 26 | 8 | 9 | 6 | 4 | 5 | 5 | 3 | 3 | 5 | 3 | 4 | 3 | 3 | 3 | 3 | 5 | 4 | 4 | 3 | 3 | 6 | 3 | 31 | 5 | 4 | 2 | 3 | 3 | 4 | 3 | 4 | 3 | 2 | 3 | 3 | 7 | 4 | 4 |
| PPP1R12A | chr12:79810395-79810511 | 0 | 0 | 0 | 0 | 0 | 0 | 0 | 0 | 1 | 0 | 1 | 0 | 0 | 0 | 0 | 0 | 0 | 1 | 0 | 0 | 0 | 0 | 0 | 0 | 0 | 0 | 0 | 0 | 0 | 34 | 36 | 0 | 0 | 0 | 1 | 57 | 0 | 0 | 0 | 0 | 0 | 0 | 0 | 0 | 0 | 0 | 0 | 0 | 0 | 0 | 0 |
| ZSWIM6 | chr5:61525016-61525147 | 12 | 10 | 7 | 10 | 11 | 10 | 14 | 10 | 8 | 8 | 9 | 23 | 23 | 9 | 9 | 6 | 7 | 9 | 12 | 7 | 9 | 15 | 2 | 13 | 11 | 12 | 9 | 11 | 7 | 8 | 10 | 7 | 13 | 8 | 11 | 34 | 15 | 11 | 10 | 10 | 12 | 10 | 11 | 13 | 9 | 11 | 5 | 10 | 10 | 9 | 6 |
| UBE2D3 | chr4:102798928-102798977 | 0 | 0 | 0 | 1 | 2 | 1 | 0 | 0 | 0 | 0 | 0 | 0 | 0 | 0 | 0 | 0 | 0 | 0 | 0 | 0 | 0 | 0 | 0 | 1 | 1 | 1 | 0 | 1 | 1 | 49 | 50 | 0 | 0 | 0 | 11 | 81 | 1 | 0 | 0 | 0 | 2 | 0 | 0 | 0 | 0 | 0 | 0 | 0 | 0 | 0 | 0 |
| SORBS1 | chr10:95371326-95372049\|95372928 | 1 | 2 | 4 | 3 | 4 | 3 | 4 | 2 | 6 | 16 | 17 | 43 | 35 | 17 | 24 | 7 | 11 | 24 | 18 | 4 | 7 | 3 | 5 | 3 | 3 | 3 | 4 | 3 | 4 | 44 | 57 | 2 | 4 | 3 | 14 | 94 | 1 | 3 | 4 | 18 | 5 | 3 | 2 | 3 | 5 | 3 | 14 | 4 | 3 | 4 | 27 |
| HSPG2 | chr1:21868885-21870872 | 0 | 1 | 1 | 1 | 1 | 1 | 1 | 1 | 3 | 2 | 4 | 2 | 3 | 4 | 3 | 2 | 6 | 2 | 3 | 3 | 4 | 2 | 1 | 1 | 1 | 1 | 1 | 1 | 0 | 21 | 30 | 0 | 0 | 1 | 1 | 41 | 3 | 0 | 1 | 1 | 1 | 0 | 0 | 1 | 0 | 1 | 1 | 0 | 0 | 1 | 4 |
| FRS2 | chr12:69480299-69480359 | 6 | 5 | 3 | 4 | 6 | 5 | 5 | 6 | 14 | 12 | 10 | 12 | 14 | 11 | 14 | 11 | 11 | 7 | 10 | 13 | 9 | 0 | -1 | 5 | 6 | 4 | 3 | 6 | 4 | 17 | 25 | 2 | 5 | 5 | 7 | 36 | 8 | 4 | 3 | 3 | 4 | 6 | 6 | 5 | 11 | 7 | 3 | 3 | 3 | 4 | 6 |
| AKAP9 | chr7:92063462-92063500 | 0 | 0 | 0 | 0 | 1 | 0 | 0 | 0 | 2 | 1 | 1 | 0 | 1 | 1 | 1 | 1 | 1 | 1 | 2 | 3 | 1 | 0 | 0 | 1 | 0 | 0 | 2 | 1 | 0 | 44 | 66 | 4 | 1 | 1 | 2 | 90 | 0 | 0 | 0 | 0 | 1 | 0 | 0 | 0 | 0 | 0 | 7 | 0 | 0 | 1 | 1 |
| EHBP1L1 | chr11:65580928-65580963 | 1 | 1 | 1 | 1 | 1 | 1 | 2 | 1 | 3 | 5 | 3 | 47 | 55 | 7 | 6 | 4 | 4 | 4 | 3 | 3 | 1 | 0 | 1 | 2 | 1 | 2 | 1 | 3 | 1 | 34 | 49 | 1 | 1 | 0 | 8 | 89 | 1 | 0 | 1 | 1 | 3 | 0 | 1 | 1 | 0 | 2 | 4 | 1 | 1 | 1 | 0 |
| QTRT2 | chr3:114077476-114077585 | 5 | 4 | 2 | 9 | 8 | 10 | 9 | 6 | 3 | 7 | 5 | 12 | 12 | 9 | 10 | 5 | 6 | 9 | 5 | 2 | 3 | 8 | 7 | 12 | 7 | 10 | 3 | 10 | 6 | 12 | 15 | 4 | 4 | 4 | 5 | 28 | 4 | 6 | 3 | 9 | 7 | 4 | 3 | 7 | 3 | 5 | 4 | 3 | 10 | 5 | 2 |
| PDPN | chr1:13609604-13609663 | 1 | 0 | 0 | 0 | 0 | 0 | 0 | 1 | 1 | 9 | 0 | 0 | 0 | 18 | 25 | 1 | 0 | 0 | 0 | 0 | 1 | 0 | 0 | 0 | 0 | 0 | 0 | 0 | 0 | 0 | 0 | 0 | 0 | 0 | 0 | 1 | 0 | 0 | 0 | 0 | 0 | 0 | 0 | 0 | 0 | 0 | 0 | 0 | 0 | 0 | 0 |
| RBM24 | chr6:17285053\|17284876-17285165 | 1 | 1 | 5 | 1 | 1 | 1 | 0 | 2 | 3 | 2 | 1 | 2 | 2 | 2 | 1 | 2 | 4 | 1 | 2 | 1 | 1 | 0 | 0 | 1 | 0 | 1 | 3 | 0 | 0 | 8 | 6 | 6 | 5 | 3 | 28 | 50 | 2 | 1 | 15 | 1 | 10 | 3 | 4 | 1 | 5 | 1 | 1 | 4 | 0 | 1 | 31 |
| MEF2A | chr15:99703362-99703385 | 2 | 1 | 1 | 1 | 2 | 1 | 2 | 2 | 9 | 30 | 12 | 19 | 25 | 29 | 30 | 11 | 6 | 20 | 13 | 3 | 5 | 1 | 0 | 3 | 2 | 2 | 1 | 2 | 3 | 33 | 37 | 0 | 1 | 2 | 2 | 60 | 2 | 1 | 1 | 1 | 2 | 2 | 2 | 2 | 2 | 2 | 1 | 1 | 2 | 1 | 1 |
| CSRNP2 | chr12:51065065-51065107 | 1 | 2 | 2 | 4 | 4 | 2 | 7 | 2 | 1 | 1 | 1 | 0 | 0 | 0 | 0 | 0 | 0 | 1 | 2 | 0 | 1 | 1 | 0 | 4 | 4 | 4 | 2 | 4 | 1 | 25 | 43 | 2 | 5 | 1 | 3 | 54 | 1 | 1 | 1 | 1 | 3 | 1 | 0 | 3 | 2 | 3 | 0 | 2 | 2 | 1 | 1 |
| C2CD5 | chr12:22514959-22515150 | 0 | 0 | 0 | 0 | 0 | 0 | 0 | 0 | 2 | 1 | 2 | 2 | 3 | 1 | 1 | 3 | 2 | 1 | 4 | 8 | 4 | 0 | 0 | 0 | 0 | 0 | 0 | 0 | 0 | 33 | 50 | 0 | 0 | 0 | 1 | 66 | 0 | 0 | 0 | 0 | 0 | 0 | 0 | 0 | 0 | 0 | 1 | 0 | 0 | 0 | 0 |
| LRRFIP2 | chr3:37109527-37109571 | 20 | 14 | 12 | 19 | 21 | 19 | 19 | 21 | 13 | 26 | 22 | 17 | 24 | 21 | 17 | 19 | 15 | 24 | 14 | 10 | 21 | 0 | -1 | 23 | 17 | 19 | 18 | 17 | -1 | 14 | 24 | -1 | 35 | 22 | 67 | 98 | 27 | 14 | 15 | -1 | 31 | 17 | 19 | 23 | 30 | 19 | 6 | 18 | 14 | 13 | 32 |
| GLT1D1 | chr12:128906907-128906954 | 3 | 1 | -1 | 36 | 11 | 24 | 0 | 15 | 3 | 3 | 10 | 5 | 4 | 4 | 5 | 3 | 2 | 8 | 10 | 5 | 5 | -1 | -1 | 17 | 12 | 8 | 6 | 18 | -1 | 10 | 51 | 0 | 0 | 1 | -1 | 96 | 19 | -1 | 1 | 5 | 34 | 2 | 3 | 7 | 0 | 8 | 0 | 1 | 42 | 26 | 0 |
| EPS15L1 | chr19:16373915-16373962 | 5 | 4 | 1 | 5 | 8 | 7 | 7 | 4 | 3 | 1 | 3 | 0 | 0 | 1 | 1 | 1 | 2 | 2 | 4 | 2 | 2 | 3 | 1 | 10 | 6 | 7 | 1 | 9 | 4 | 50 | 54 | 1 | 0 | 3 | 9 | 79 | 9 | 2 | 1 | 0 | 4 | 2 | 2 | 4 | 2 | 4 | 2 | 2 | 4 | 3 | 4 |
| SPATS2 | chr12:49512862-49512905 | 17 | 16 | 9 | 15 | 19 | 14 | 12 | 15 | 11 | 5 | 13 | 2 | 2 | 4 | 3 | 8 | 7 | 9 | 12 | 15 | 11 | 7 | 7 | 13 | 12 | 13 | 9 | 13 | 18 | 25 | 32 | 12 | 18 | 13 | 6 | 54 | 25 | 8 | 8 | 5 | 7 | 7 | 6 | 16 | 20 | 9 | 2 | 10 | 11 | 9 | 16 |
| CNNM2 | chr10:103054663-103054727 | 10 | 5 | 2 | 12 | 13 | 18 | 13 | 8 | 6 | 6 | 5 | 11 | 12 | 7 | 6 | 6 | 4 | 6 | 6 | 2 | 3 | 16 | 12 | 13 | 5 | 11 | 5 | 12 | 4 | 15 | 18 | 3 | 3 | 3 | 2 | 39 | 7 | 5 | 2 | 4 | 8 | 4 | 5 | 7 | 3 | 5 | 9 | 4 | 13 | 9 | 3 |
| COG1 | chr17:73207702-73207739 | 3 | 1 | 1 | 2 | 3 | 4 | 3 | 2 | 38 | 27 | 44 | 5 | 5 | 19 | 17 | 29 | 22 | 57 | 39 | 4 | 8 | 3 | 2 | 6 | 2 | 4 | 1 | 5 | 2 | 49 | 67 | 1 | 1 | 2 | 4 | 88 | 3 | 2 | 1 | 6 | 2 | 1 | 2 | 3 | 1 | 2 | 1 | 1 | 1 | 2 | 3 |
| BMP1 | chr8:22192235-22192335 | 12 | 8 | 4 | 10 | 8 | 12 | 9 | 11 | 9 | 8 | 8 | 11 | 9 | 10 | 8 | 12 | 9 | 7 | 9 | 9 | 10 | 9 | 9 | 10 | 7 | 8 | 7 | 8 | 9 | 8 | 9 | 9 | 6 | 10 | 9 | 23 | 14 | 11 | 6 | 8 | 9 | 13 | 14 | 8 | 7 | 8 | 11 | 9 | 11 | 12 | 9 |
| RP11-463D19.2 | chr8:73709032-73709162 | 7 | 9 | 7 | 10 | 9 | 10 | 8 | 6 | 6 | 6 | 6 | 10 | 7 | 7 | 10 | 7 | 5 | 5 | 7 | 9 | 7 | 9 | 4 | 9 | 9 | 8 | 10 | 9 | 6 | 11 | 20 | 6 | 10 | 9 | 8 | 31 | 7 | 5 | 13 | 5 | 9 | 5 | 5 | 8 | 8 | 9 | 21 | 6 | 6 | 10 | 6 |
| E2F6 | chr2:11458281-11458351 | 16 | 19 | 13 | 15 | 15 | 12 | 20 | 19 | 21 | 32 | 23 | 27 | 24 | 27 | 24 | 20 | 32 | 30 | 22 | 23 | 25 | 12 | -1 | 17 | 16 | 16 | 11 | 19 | 20 | 27 | 35 | 14 | 26 | 10 | 17 | 58 | 16 | 8 | 13 | 16 | 15 | 12 | 8 | 17 | 15 | 16 | 34 | 9 | 11 | 11 | 15 |
| PIGX | chr3:196728702-196728755 | 2 | 4 | 5 | 2 | 4 | 2 | 4 | 2 | 16 | 21 | 16 | 43 | 41 | 20 | 22 | 15 | 40 | 13 | 15 | 14 | 22 | 3 | 3 | 5 | 4 | 4 | 3 | 5 | 1 | 29 | 54 | 3 | 4 | 3 | 4 | 76 | 2 | 4 | 2 | 19 | 3 | 2 | 1 | 4 | 3 | 4 | 4 | 2 | 3 | 3 | 3 |
| MFN2 | chr1:11982523\|11982506-11982590 | 4 | 4 | 3 | 3 | 5 | 3 | 3 | 4 | 12 | 12 | 10 | 14 | 13 | 12 | 13 | 13 | 11 | 12 | 11 | 11 | 12 | 3 | 1 | 4 | 4 | 3 | 5 | 4 | 2 | 28 | 34 | 3 | 8 | 4 | 8 | 46 | 5 | 1 | 4 | 2 | 4 | 4 | 4 | 4 | 3 | 4 | 4 | 2 | 2 | 3 | 4 |
| SLC29A3 | chr10:71352567-71352818 | 5 | 4 | 1 | 3 | 4 | 6 | 2 | 7 | 7 | 8 | 5 | 27 | 24 | 14 | 13 | 5 | 10 | 6 | 5 | 9 | 7 | 6 | 4 | 4 | 2 | 3 | 3 | 3 | 0 | 10 | 9 | 3 | 5 | 4 | 4 | 29 | 8 | 4 | 1 | 3 | 4 | 5 | 7 | 2 | 2 | 1 | 6 | 5 | 4 | 5 | 6 |
| CARS2 | chr13:110697918-110697967 | 6 | 4 | 2 | 7 | 6 | 7 | 6 | 5 | 7 | 12 | 8 | 15 | 15 | 13 | 14 | 7 | 8 | 9 | 9 | 5 | 6 | 5 | 5 | 6 | 5 | 5 | 3 | 6 | 5 | 8 | 7 | 2 | 2 | 4 | 4 | 20 | 6 | 6 | 2 | 7 | 6 | 4 | 5 | 4 | 3 | 4 | 3 | 4 | 7 | 4 | 2 |
| DPYSL4 | chr10:132192369\|132192283-132192520 | 3 | 5 | 8 | 6 | 7 | 5 | -1 | 3 | 23 | 18 | 15 | 20 | 24 | 20 | 15 | 30 | 20 | 15 | 15 | 47 | 40 | -1 | -1 | 16 | 10 | 12 | 5 | 12 | -1 | 15 | 10 | 10 | 6 | 6 | 10 | 49 | 5 | 4 | 11 | 27 | 7 | 4 | 3 | 10 | 6 | 10 | 19 | 5 | 5 | 6 | 20 |
| KMT2C | chr7:152195522-152195569 | 1 | 1 | 7 | 3 | 3 | 1 | 5 | 1 | 47 | 46 | 51 | 9 | 14 | 35 | 37 | 28 | 33 | 52 | 37 | 20 | 32 | 0 | -1 | 3 | 3 | 2 | 1 | 3 | -1 | 48 | 64 | 4 | 2 | 1 | 4 | 91 | 2 | 22 | 1 | 35 | 3 | 0 | 1 | 2 | 2 | 2 | 8 | 1 | 3 | 1 | 1 |
| VSIG8 | chr1:159842227-159842314 | 10 | 5 | 4 | 9 | 8 | 10 | 10 | 5 | 10 | 11 | 9 | 37 | 35 | 13 | 13 | 14 | 9 | 13 | 10 | 11 | 10 | 0 | -1 | 9 | 7 | 10 | 5 | 11 | 7 | 24 | 31 | 3 | -1 | 5 | 6 | 54 | 16 | 7 | 3 | 9 | 8 | 6 | 6 | 6 | 4 | 7 | 5 | 5 | 6 | 7 | 5 |
| SLC17A5 | chr6:73642846-73642923 | 0 | 0 | 0 | 0 | 1 | 0 | 1 | 0 | 0 | 0 | 0 | 0 | 0 | 0 | 0 | 0 | 0 | 0 | 0 | 0 | 0 | 1 | 0 | 1 | 0 | 1 | 0 | 1 | 0 | 14 | 20 | 0 | 0 | 0 | 0 | 25 | 0 | 0 | 0 | 0 | 1 | 0 | 0 | 0 | 0 | 1 | 0 | 0 | 0 | 0 | 0 |
| ZSCAN5A | chr19:56314250-56314343 | 10 | 9 | 4 | 10 | 10 | 8 | 8 | 8 | 8 | 12 | 8 | 19 | 22 | 11 | 12 | 10 | 12 | 6 | 8 | 10 | 9 | 6 | 6 | 10 | 8 | 9 | 12 | 11 | 4 | 7 | 8 | 6 | 12 | 5 | 8 | 32 | 8 | 8 | 5 | 7 | 9 | 8 | 7 | 8 | 6 | 8 | 22 | 5 | 8 | 9 | 4 |
| MALT1 | chr18:58720380-58720422 | 1 | 1 | 1 | 1 | 1 | 1 | 0 | 1 | 9 | 27 | 12 | 46 | 58 | 39 | 39 | 14 | 27 | 20 | 17 | 12 | 17 | 0 | 1 | 1 | 1 | 1 | 0 | 1 | 2 | 26 | 28 | 1 | 2 | 1 | 1 | 70 | 1 | 0 | 1 | 1 | 1 | 1 | 0 | 0 | 1 | 1 | 2 | 1 | 1 | 1 | 0 |
| ERI1 | chr8:9004074-9004296 | 2 | 3 | 1 | 3 | 3 | 4 | 4 | 2 | 24 | 15 | 17 | 14 | 15 | 16 | 16 | 20 | 16 | 17 | 19 | 16 | 17 | 3 | 4 | 4 | 3 | 3 | 2 | 3 | 5 | 5 | 13 | 4 | 5 | 5 | 3 | 20 | 5 | 4 | 5 | 3 | 4 | 2 | 1 | 3 | 5 | 3 | 1 | 4 | 4 | 3 | 2 |
| RPS24 | chr10:78037965-78037982 | 1 | 0 | 0 | 1 | 4 | 6 | 26 | 0 | 7 | 13 | 20 | 27 | 28 | 23 | 24 | 12 | 3 | 12 | 21 | 2 | 7 | 2 | 1 | 32 | 17 | 20 | 3 | 30 | 4 | 37 | 55 | 0 | 0 | 0 | 6 | 87 | 0 | 0 | 0 | 0 | 9 | 0 | 0 | 9 | 0 | 9 | 0 | 1 | 2 | 2 | 1 |
| REEP3 | chr10:63599670-63599712 | 0 | 0 | 0 | 0 | 0 | 0 | 0 | 0 | 5 | 13 | 4 | 59 | 66 | 17 | 17 | 6 | 8 | 4 | 2 | 1 | 4 | 1 | 0 | 0 | 0 | 0 | 0 | 0 | 1 | 20 | 18 | 0 | 0 | 0 | 0 | 72 | 0 | 0 | 0 | 1 | 0 | 0 | 0 | 0 | 0 | 0 | 0 | 0 | 0 | 0 | 0 |
| ANKK1 | chr11:113395585-113395614 | 25 | 21 | 7 | 31 | 21 | 19 | -1 | 21 | 16 | 24 | 22 | 21 | 18 | 29 | 43 | 25 | 15 | 37 | 44 | 38 | 37 | -1 | -1 | 30 | 13 | 27 | 15 | 18 | 16 | 26 | 29 | 13 | 15 | 22 | 14 | 69 | 36 | 18 | 13 | 15 | 15 | 26 | 23 | 19 | 15 | 25 | 15 | 12 | 24 | 22 | 12 |
| PRRC2B | chr9:131486481-131486541 | 14 | 8 | 3 | 4 | 5 | 6 | 8 | 12 | 1 | 1 | 1 | 1 | 1 | 1 | 1 | 1 | 1 | 1 | 1 | 3 | 2 | 6 | 5 | 6 | 8 | 5 | 7 | 6 | 7 | 6 | 6 | 6 | 10 | 9 | 7 | 36 | 9 | 5 | 7 | 2 | 6 | 13 | 15 | 15 | 16 | 8 | 26 | 6 | 5 | 7 | 12 |
| GOLGA4 | chr3:37266842-37266940\|37267255 | 18 | 17 | 10 | 13 | 14 | 8 | 4 | 15 | 47 | 47 | 52 | 10 | 14 | 47 | 47 | 51 | 40 | 44 | 56 | 56 | 52 | 12 | 11 | 10 | 6 | 10 | 2 | 11 | 9 | 45 | 36 | 5 | 14 | 9 | 7 | 66 | 31 | 14 | 3 | 8 | 9 | 3 | 2 | 4 | 9 | 6 | 13 | 5 | 7 | 3 | 2 |
| EIF4G3 | chr1:20997601-20997633 | 7 | 8 | 13 | 8 | 8 | 8 | 8 | 9 | 32 | 25 | 31 | 22 | 23 | 27 | 26 | 31 | 24 | 23 | 30 | 32 | 36 | 11 | 4 | 7 | 4 | 6 | 4 | 7 | 7 | 27 | 37 | 10 | 7 | 5 | 7 | 51 | 9 | 8 | 7 | 9 | 7 | 6 | 6 | 4 | 5 | 7 | 25 | 8 | 7 | 6 | 1 |
| ABLIM2 | chr4:8019618\|8019423-8027857 | 2 | 5 | 3 | 4 | 5 | 8 | 6 | 4 | 62 | 63 | 58 | 39 | 41 | 59 | 58 | 56 | 53 | 65 | 55 | 36 | 42 | 4 | -1 | 21 | 11 | 7 | 4 | 13 | 0 | 33 | 50 | 2 | 8 | 6 | 39 | 87 | 7 | 4 | 3 | 15 | 27 | 1 | 2 | 9 | 2 | 7 | 17 | 4 | 6 | 4 | 37 |
| COBL | chr7:51187914-51187958 | 9 | 3 | 11 | 19 | 16 | 33 | -1 | 2 | 0 | 1 | 1 | 0 | 1 | 1 | 1 | 0 | 0 | 1 | 1 | 0 | 0 | 0 | 0 | 4 | 1 | 8 | 2 | 11 | 0 | 47 | 47 | 2 | 2 | 2 | 23 | 99 | 2 | 8 | 1 | 1 | 10 | 0 | 1 | 1 | -1 | 2 | 2 | 4 | 8 | 1 | 65 |
| GFPT1 | chr2:69354259-69354312 | 4 | 3 | 1 | 8 | 10 | 14 | 6 | 4 | 3 | 7 | 2 | 4 | 3 | 5 | 7 | 3 | 4 | 3 | 1 | 1 | 2 | 1 | 3 | 7 | 2 | 9 | 1 | 9 | 8 | 41 | 69 | 1 | 1 | 2 | 1 | 81 | 3 | 2 | 0 | 1 | 3 | 2 | 2 | 2 | 1 | 2 | 0 | 1 | 6 | 3 | 0 |
| CDC42BPA | chr1:227010878\|227010904-227011063 | 6 | 5 | 1 | 3 | 5 | 6 | 15 | 4 | 55 | 29 | 47 | 6 | 8 | 34 | 23 | 56 | 36 | 39 | 54 | 70 | 57 | 5 | 1 | 12 | 6 | 10 | 4 | 14 | 2 | 39 | 45 | 0 | 0 | 1 | 3 | 67 | 32 | 2 | 0 | 1 | 10 | 2 | 2 | 4 | 1 | 4 | 3 | 1 | 2 | 3 | 7 |
| WDR20 | chr14:102193463-102193514 | 8 | 7 | 9 | 7 | 7 | 6 | 8 | 7 | 9 | 11 | 10 | 11 | 10 | 10 | 13 | 8 | 9 | 11 | 10 | 5 | 9 | 6 | 6 | 10 | 8 | 8 | 5 | 9 | 8 | 16 | 20 | 11 | 8 | 6 | 8 | 25 | 6 | 5 | 7 | 7 | 8 | 5 | 5 | 9 | 7 | 8 | 6 | 7 | 7 | 4 | 3 |
| UBR4 | chr1:19149763-19149795 | 3 | 2 | 1 | 2 | 2 | 3 | 4 | 2 | 16 | 36 | 20 | 77 | 76 | 42 | 47 | 23 | 34 | 36 | 22 | 7 | 13 | 1 | 5 | 4 | 3 | 4 | 2 | 6 | 2 | 26 | 36 | 0 | 0 | 2 | 5 | 87 | 6 | 2 | 1 | 25 | 2 | 1 | 2 | 2 | 1 | 2 | 1 | 1 | 2 | 2 | 1 |
| PPP3CA | chr4:101029166-101029195 | 3 | 3 | 1 | 2 | 4 | 2 | 1 | 2 | 70 | 85 | 89 | 69 | 65 | 85 | 87 | 73 | 60 | 76 | 89 | 43 | 71 | 1 | 1 | 6 | 4 | 5 | 2 | 8 | 5 | 18 | 52 | 1 | 0 | 4 | 3 | 86 | 19 | 1 | 1 | 24 | 2 | 2 | 2 | 4 | 4 | 2 | 22 | 2 | 1 | 2 | 2 |
| ZNF347 | chr19:53149936-53149984 | 6 | 3 | 1 | 12 | 10 | 13 | 7 | 5 | 2 | 2 | 1 | 3 | 6 | 3 | 3 | 1 | 1 | 2 | 1 | 3 | 2 | -1 | -1 | 14 | 8 | 12 | 7 | 12 | -1 | 4 | 7 | 1 | 2 | 4 | 3 | 28 | 3 | 3 | 2 | 0 | 4 | 7 | 6 | 5 | 3 | 6 | 5 | 2 | 6 | 5 | 0 |
| HM13 | chr20:31568078-31568280\|31568224 | 10 | 7 | 5 | 9 | 7 | 11 | 7 | 8 | 32 | 50 | 36 | 58 | 57 | 59 | 60 | 43 | 42 | 41 | 38 | 30 | 35 | 6 | 5 | 8 | 5 | 7 | 8 | 7 | 5 | 26 | 44 | 5 | 10 | 6 | 5 | 71 | 14 | 4 | 5 | 5 | 5 | 6 | 7 | 5 | 5 | 5 | 7 | 4 | 6 | 6 | 7 |
| ZNF189 | chr9:101407412-101407532 | 1 | 1 | 2 | 1 | 1 | 0 | 0 | 1 | 10 | 9 | 7 | 4 | 3 | 7 | 8 | 14 | 12 | 8 | 12 | 26 | 20 | 0 | 0 | 1 | 1 | 1 | 1 | 1 | 1 | 2 | 2 | 2 | 3 | 1 | 1 | 5 | 1 | 0 | 2 | 2 | 2 | 1 | 1 | 1 | 1 | 1 | 1 | 1 | 1 | 0 | 1 |
| ASB8 | chr12:48150087-48150140 | 5 | 6 | 2 | 3 | 3 | 3 | 4 | 5 | 12 | 14 | 11 | 9 | 9 | 15 | 16 | 15 | 9 | 13 | 12 | 12 | 10 | 2 | 3 | 4 | 3 | 4 | 3 | 4 | 3 | 20 | 24 | 4 | 5 | 5 | 6 | 31 | 7 | 2 | 4 | 3 | 5 | 2 | 3 | 3 | 5 | 3 | 11 | 3 | 3 | 3 | 4 |
| TRAK1 | chr3:42220499-42220591 | 2 | 1 | 0 | 2 | 3 | 1 | 1 | 1 | 3 | 2 | 3 | 0 | 0 | 2 | 1 | 3 | 2 | 2 | 3 | 8 | 4 | 2 | 0 | 2 | 1 | 2 | 0 | 2 | 1 | 44 | 45 | 0 | 0 | 2 | 2 | 62 | 21 | 0 | 0 | 0 | 2 | 0 | 0 | 0 | 1 | 0 | 1 | 0 | 1 | 1 | 1 |
| MYL6 | chr12:56158902-56158920 | 1 | 1 | 0 | 1 | 1 | 1 | 2 | 1 | 9 | 10 | 7 | 13 | 13 | 13 | 16 | 12 | 7 | 7 | 11 | 9 | 9 | 1 | 1 | 2 | 1 | 2 | 1 | 2 | 1 | 22 | 29 | 0 | 1 | 1 | 1 | 36 | 3 | 0 | 0 | 1 | 1 | 1 | 1 | 1 | 1 | 1 | 0 | 0 | 1 | 1 | 0 |
| TNK2 | chr3:195869497\|195869428-195869541 | 17 | 12 | 10 | 4 | 7 | 5 | 8 | 13 | 66 | 73 | 67 | 62 | 63 | 73 | 73 | 66 | 61 | 70 | 66 | 45 | 55 | 7 | 6 | 8 | 10 | 7 | 13 | 9 | 10 | 48 | 53 | 13 | 18 | 12 | 16 | 89 | 24 | 6 | 9 | 13 | 9 | 16 | 18 | 13 | 15 | 12 | 30 | 15 | 4 | 9 | 20 |
| GLE1 | chr9:128532665-128532691 | 3 | 4 | 3 | 2 | 2 | 2 | 3 | 3 | 7 | 5 | 4 | 8 | 9 | 6 | 5 | 4 | 3 | 4 | 4 | 6 | 4 | 2 | 4 | 2 | 2 | 3 | 2 | 3 | 1 | 15 | 26 | 3 | 6 | 3 | 2 | 29 | 3 | 1 | 3 | 2 | 2 | 2 | 3 | 4 | 3 | 3 | 2 | 2 | 1 | 2 | 4 |
| RTN3 | chr11:63704851-63704907 | 2 | 2 | 2 | 2 | 2 | 3 | 2 | 2 | 13 | 19 | 15 | 25 | 24 | 19 | 19 | 15 | 15 | 18 | 14 | 10 | 11 | 1 | 2 | 2 | 1 | 2 | 1 | 2 | 2 | 25 | 31 | 1 | 1 | 1 | 1 | 43 | 3 | 2 | 1 | 3 | 3 | 1 | 1 | 2 | 1 | 1 | 3 | 1 | 2 | 1 | 1 |
| GULP1 | chr2:188582341-188582565\|188582413 | 14 | 9 | 6 | 19 | 17 | 23 | 16 | 13 | 53 | 73 | 62 | 72 | 52 | 76 | 84 | 43 | 61 | 64 | 55 | 26 | 46 | 13 | 12 | 21 | 16 | 18 | 8 | 19 | 11 | 17 | 9 | 6 | 7 | 13 | 7 | 62 | 32 | 9 | 6 | 10 | 12 | 14 | 17 | 10 | 5 | 11 | 23 | 7 | 14 | 12 | 21 |
| ACSS3 | chr12:81199660-81199711 | 6 | 4 | 1 | 9 | 7 | 10 | 5 | 5 | 7 | 8 | 7 | 7 | 9 | 5 | 5 | 7 | 8 | 8 | 6 | 4 | 6 | 12 | 11 | 6 | 7 | 4 | 4 | 6 | 7 | 25 | 17 | 2 | 3 | 6 | 3 | 35 | 22 | 6 | 3 | 11 | 7 | 6 | 7 | 6 | 4 | 7 | 10 | 4 | 10 | 9 | 3 |
| CSDE1 | chr1:114741527-114741673\|114741615 | 1 | 1 | 1 | 3 | 4 | 4 | 5 | 1 | 5 | 4 | 4 | 2 | 2 | 4 | 4 | 6 | 5 | 3 | 5 | 15 | 9 | 1 | 1 | 8 | 5 | 5 | 2 | 8 | 2 | 61 | 79 | 1 | 1 | 1 | 9 | 93 | 2 | 1 | 0 | 1 | 7 | 1 | 1 | 3 | 1 | 4 | 0 | 1 | 2 | 2 | 1 |
| CEP350 | chr1:180073786-180073908 | 0 | 0 | 0 | 0 | 0 | 0 | 0 | 0 | 7 | 23 | 8 | 81 | 74 | 25 | 32 | 11 | 18 | 13 | 9 | 14 | 8 | 0 | 0 | 0 | 0 | 0 | 0 | 0 | 0 | 35 | 57 | 0 | 0 | 0 | 4 | 91 | 0 | 0 | 0 | 1 | 2 | 0 | 0 | 0 | 0 | 0 | 0 | 0 | 0 | 0 | 0 |
| AC084219.2 | chr19:44026026-44026086 | 24 | 19 | 16 | 20 | 20 | 26 | 34 | 24 | 22 | 29 | 16 | 18 | 21 | 27 | 41 | 30 | 29 | 28 | 26 | 23 | 32 | -1 | -1 | 20 | 19 | 16 | 16 | 19 | -1 | 20 | 25 | 14 | 10 | 19 | 14 | 58 | 22 | 18 | 26 | 12 | 14 | 22 | 18 | 22 | 17 | 23 | 21 | 11 | 17 | 13 | 10 |
| LRRFIP2 | chr3:37102924-37127680 | 1 | 1 | 0 | 3 | 5 | 4 | 6 | 1 | 2 | 2 | 2 | 1 | 0 | 2 | 1 | 1 | 1 | 1 | 2 | 1 | 1 | 1 | 1 | 7 | 3 | 6 | 1 | 7 | 3 | 68 | 74 | 0 | 1 | 1 | 3 | 92 | 4 | 0 | 0 | 0 | 5 | 0 | 0 | 3 | 0 | 3 | 0 | 1 | 1 | 1 | 0 |
| HM13 | chr20:31568078\|31567640-31568280 | 8 | 6 | 4 | 7 | 6 | 8 | 5 | 7 | 27 | 42 | 31 | 49 | 49 | 52 | 52 | 37 | 36 | 35 | 32 | 26 | 30 | 5 | 5 | 7 | 5 | 6 | 7 | 6 | 4 | 22 | 37 | 4 | 8 | 5 | 4 | 57 | 12 | 4 | 4 | 4 | 5 | 5 | 6 | 4 | 5 | 4 | 6 | 3 | 5 | 5 | 6 |
| ZBED8 | chr5:160397442-160397515 | 9 | 6 | 7 | 8 | 7 | 6 | -1 | 7 | 6 | 7 | 7 | 8 | 12 | 7 | 7 | 8 | 7 | 6 | 5 | 5 | 3 | -1 | -1 | 8 | 9 | 7 | 4 | 8 | 4 | 9 | 8 | 1 | 11 | 7 | 3 | 22 | 6 | 5 | 5 | 7 | 5 | 5 | 5 | 6 | 6 | 8 | 3 | 4 | 7 | 4 | 6 |
| RPN2 | chr20:37238402-37238449 | 5 | 5 | 2 | 6 | 6 | 8 | 8 | 4 | 27 | 42 | 29 | 52 | 48 | 46 | 49 | 33 | 35 | 34 | 31 | 25 | 28 | 3 | 2 | 9 | 4 | 7 | 1 | 8 | 3 | 34 | 53 | 2 | 5 | 2 | 2 | 73 | 7 | 2 | 1 | 2 | 3 | 1 | 2 | 3 | 2 | 3 | 24 | 1 | 3 | 2 | 2 |
| OTOF | chr2:26460008-26460205 | 2 | 3 | 4 | 4 | 2 | 4 | 0 | 2 | 76 | 77 | 81 | 58 | 57 | 82 | 83 | 77 | 72 | 76 | 84 | 55 | 66 | 0 | -1 | 8 | 11 | 13 | 3 | 15 | 0 | 7 | 21 | -1 | 0 | 2 | -1 | 59 | 3 | 2 | 1 | 4 | 3 | 3 | 1 | 6 | 2 | 5 | 2 | 3 | -1 | 1 | 1 |
| SLC16A9 | chr10:59707106-59707188 | 1 | 2 | 2 | 2 | 2 | 2 | 0 | 1 | 8 | 9 | 8 | 67 | 63 | 9 | 8 | 8 | 13 | 8 | 9 | 8 | 8 | 0 | 2 | 2 | 1 | 1 | 1 | 2 | 2 | 20 | 57 | 2 | 0 | 3 | 2 | 79 | 2 | 1 | 3 | 4 | 2 | 0 | 5 | 0 | 3 | 2 | 1 | 2 | 2 | 2 | -1 |
| CDC42BPA | chr1:227059373-227059411 | 0 | 1 | 1 | 0 | 0 | 0 | 0 | 0 | 8 | 15 | 7 | 34 | 30 | 16 | 18 | 10 | 26 | 7 | 9 | 15 | 19 | 0 | 1 | 1 | 1 | 0 | 0 | 1 | 1 | 18 | 20 | 1 | 0 | 0 | 2 | 36 | 1 | 0 | 0 | 2 | 1 | 0 | 1 | 2 | 0 | 1 | 1 | 1 | 0 | 1 | 3 |
| SCAF8 | chr6:154741956-154742018 | 4 | 4 | 5 | 5 | 6 | 6 | 6 | 4 | 21 | 20 | 18 | 11 | 10 | 18 | 17 | 15 | 13 | 19 | 14 | 11 | 18 | 7 | 4 | 3 | 4 | 4 | 4 | 4 | 8 | 7 | 13 | 8 | 5 | 6 | 3 | 20 | 9 | 5 | 4 | 6 | 3 | 4 | 3 | 4 | 4 | 5 | 15 | 5 | 4 | 3 | 3 |
| GMPPA | chr2:219504854-219504868 | 0 | 0 | 0 | 0 | 1 | 0 | 1 | 0 | 13 | 28 | 10 | 51 | 48 | 40 | 44 | 23 | 20 | 6 | 11 | 35 | 25 | 1 | 0 | 1 | 0 | 0 | 0 | 1 | 0 | 53 | 64 | 0 | 0 | 0 | 1 | 86 | 1 | 0 | 0 | 1 | 1 | 0 | 0 | 0 | 0 | 0 | 1 | 0 | 0 | 0 | 1 |
| CDC42BPA | chr1:227010878-227011063\|227013126 | 6 | 5 | 1 | 3 | 5 | 6 | 15 | 4 | 54 | 28 | 47 | 7 | 8 | 33 | 23 | 55 | 35 | 39 | 54 | 67 | 55 | 5 | 1 | 12 | 6 | 10 | 4 | 14 | 2 | 41 | 47 | 0 | 0 | 1 | 3 | 63 | 29 | 2 | 0 | 2 | 10 | 2 | 2 | 4 | 1 | 4 | 3 | 1 | 2 | 3 | 7 |
| SMARCA2 | chr9:2170419-2170472 | 7 | 9 | 11 | 9 | 9 | 7 | 10 | 9 | 8 | 8 | 8 | 9 | 9 | 8 | 9 | 8 | 7 | 7 | 9 | 9 | 9 | 7 | 8 | 8 | 13 | 8 | 17 | 8 | 9 | 10 | 11 | 11 | 20 | 14 | 13 | 30 | 8 | 7 | 15 | 10 | 10 | 16 | 14 | 15 | 16 | 12 | 11 | 11 | 7 | 10 | 20 |
| AGFG1 | chr2:227532138-227532209 | 1 | 1 | 1 | 1 | 1 | 1 | 0 | 1 | 3 | 1 | 2 | 1 | 1 | 2 | 1 | 3 | 1 | 2 | 3 | 4 | 2 | 0 | 2 | 1 | 0 | 1 | 0 | 1 | 1 | 26 | 44 | 0 | 1 | 0 | 1 | 49 | 2 | 0 | 0 | 1 | 1 | 0 | 0 | 0 | 0 | 1 | 20 | 0 | 1 | 0 | 0 |
| ACTB | chr7:5529019-5529059 | 0 | 0 | 0 | 0 | 0 | 0 | 2 | 0 | 0 | 0 | 0 | 1 | 1 | 0 | 0 | 0 | 0 | 0 | 0 | 0 | 0 | 0 | 0 | 1 | 1 | 1 | 0 | 1 | 0 | 50 | 30 | 0 | 0 | 0 | 1 | 53 | 0 | 0 | 0 | 1 | 0 | 0 | 0 | 1 | 0 | 1 | 0 | 0 | 0 | 0 | 0 |
| PPP1R3F | chrX:49283680\|49283700-49283734 | 0 | 0 | 0 | 0 | 0 | 0 | 0 | 0 | 3 | 1 | 4 | 1 | 1 | 1 | 1 | 4 | 3 | 2 | 5 | 30 | 13 | 0 | 0 | 0 | 0 | 0 | 0 | 0 | 0 | 0 | 0 | 0 | 0 | 0 | 0 | 0 | 5 | 0 | 0 | 0 | 0 | 0 | 0 | 0 | 0 | 0 | 0 | 0 | 0 | 0 | 0 |
| LAMA3 | chr18:23858260-23858390 | 1 | 1 | 0 | 1 | 1 | 0 | 4 | 1 | 2 | 3 | 4 | 1 | 2 | 2 | 2 | 1 | 2 | 4 | 1 | 7 | 3 | 1 | 0 | 3 | 1 | 1 | 0 | 2 | 0 | 1 | 1 | 0 | 2 | 3 | 0 | 2 | 22 | 0 | 0 | 0 | 1 | 1 | 0 | 0 | 0 | 1 | 1 | 0 | 0 | 0 | -1 |
| ATM | chr11:108320687\|108320710-108320780 | 0 | 0 | 0 | 0 | 0 | 0 | 0 | 0 | 22 | 14 | 26 | 1 | 2 | 22 | 16 | 28 | 13 | 15 | 29 | 29 | 36 | 0 | 0 | 1 | 0 | 0 | 0 | 0 | 1 | 0 | 0 | 0 | 0 | 0 | 1 | 6 | 0 | 0 | 0 | 0 | 0 | 0 | 0 | 0 | 0 | 0 | 0 | 0 | 0 | 0 | 0 |
| DOCK9 | chr13:98867064-98867105 | 0 | 0 | 0 | 0 | 1 | 0 | 1 | 0 | 12 | 8 | 13 | 5 | 8 | 13 | 11 | 10 | 9 | 12 | 13 | 9 | 11 | 0 | 0 | 0 | 1 | 1 | 0 | 1 | 0 | 1 | 1 | 0 | 0 | 0 | 0 | 13 | 20 | 0 | 0 | 2 | 0 | 1 | 1 | 1 | 1 | 0 | 0 | 0 | 0 | 0 | 0 |
| LAMA3 | chr18:23925827-23925881 | 3 | 2 | 1 | 2 | 2 | 2 | 1 | 2 | 5 | 4 | 5 | 8 | 5 | 4 | 3 | 3 | 4 | 7 | 3 | 5 | 2 | 2 | 1 | 5 | 2 | 3 | 1 | 4 | 2 | 5 | 19 | 1 | 1 | 1 | 1 | 15 | 32 | 1 | 1 | 0 | 2 | 1 | 1 | 1 | 1 | 1 | 2 | 2 | 1 | 1 | 3 |
| IGSF1 | chrX:131280875-131281054 | 13 | 3 | 3 | 1 | 1 | 1 | 0 | 5 | 4 | 3 | 3 | 0 | 3 | 4 | 2 | 6 | 3 | 4 | 7 | 3 | 4 | 0 | 0 | 4 | 5 | 2 | 4 | 5 | 0 | 4 | 3 | -1 | 0 | 3 | 5 | 13 | 26 | 2 | 2 | 6 | 4 | 7 | 6 | 5 | 0 | 3 | 3 | 3 | 4 | 2 | -1 |
| NFASC | chr1:204984027-204986114 | 14 | 23 | 31 | 17 | 18 | 12 | 18 | 14 | 74 | 39 | 72 | 15 | 16 | 54 | 32 | 76 | 49 | 61 | 77 | 89 | 74 | 17 | 15 | 19 | 20 | 14 | 10 | 14 | 12 | 68 | 28 | 24 | 21 | 15 | 15 | 18 | 90 | 11 | 19 | 7 | 18 | 8 | 7 | 20 | 23 | 22 | 21 | 23 | 15 | 14 | 28 |
| ETFDH | chr4:158700875-158700919 | 5 | 1 | 0 | 2 | 1 | 2 | 2 | 3 | 15 | 8 | 10 | 3 | 4 | 9 | 7 | 16 | 11 | 10 | 12 | 42 | 17 | 2 | 2 | 2 | 1 | 1 | 1 | 1 | 3 | 0 | 0 | 1 | 0 | 2 | 1 | 0 | 11 | 3 | 1 | 1 | 2 | 3 | 5 | 1 | 3 | 1 | 6 | 2 | 2 | 1 | 1 |
| TUBG2 | chr17:42661201\|42660880-42661227 | 16 | 15 | 8 | 10 | 10 | 7 | 7 | 12 | 29 | 16 | 29 | 11 | 11 | 13 | 11 | 25 | 27 | 24 | 32 | 73 | 48 | 6 | 6 | 7 | 8 | 7 | 9 | 6 | 10 | 14 | 21 | 8 | 13 | 12 | 7 | 16 | 31 | 8 | 7 | 6 | 7 | 10 | 9 | 11 | 11 | 9 | 18 | 7 | 7 | 8 | 11 |
| SERPINB9 | chr6:2897926-2898069\|2898048\|2898073 | 8 | 9 | 4 | 10 | 10 | 10 | 6 | 9 | 13 | 11 | 17 | 5 | 5 | 11 | 10 | 12 | 11 | 13 | 11 | 18 | 15 | 7 | 8 | 12 | 8 | 10 | 6 | 9 | 6 | 12 | 9 | 7 | 9 | 16 | 8 | 13 | 24 | 9 | 10 | 9 | 8 | 6 | 8 | 6 | 5 | 5 | 4 | 11 | 11 | 14 | 6 |
| PHLDB1 | chr11:118655860-118655892 | 1 | 1 | 0 | 0 | 1 | 0 | 1 | 0 | 48 | 42 | 48 | 10 | 8 | 50 | 42 | 56 | 43 | 43 | 52 | 62 | 57 | 0 | 1 | 2 | 1 | 2 | 1 | 3 | 1 | 1 | 2 | 0 | 0 | 0 | 1 | 1 | 9 | 0 | 0 | 1 | 1 | 1 | 0 | 1 | 0 | 1 | 0 | 0 | 0 | 1 | 4 |
| CLCN5 | chrX:50088413-50088484 | 1 | 1 | 12 | 11 | 3 | 4 | 0 | 1 | 7 | 9 | 16 | 4 | 7 | 12 | 5 | 7 | 8 | 17 | 21 | 9 | 7 | -1 | 0 | 3 | 2 | 6 | 1 | 4 | -1 | 6 | 13 | 4 | 1 | 1 | 2 | 12 | 1 | 37 | 1 | 16 | 8 | 0 | 0 | 1 | 2 | 2 | 2 | 3 | 12 | 1 | 1 |
| ASTN1 | chr1:176931526-176931684 | 7 | 5 | 5 | 7 | -1 | 8 | 0 | 5 | 2 | 1 | 1 | 3 | 4 | 2 | 1 | 1 | 1 | 1 | 1 | 2 | 2 | 0 | 0 | 0 | 1 | 3 | 4 | 3 | -1 | 7 | 4 | 0 | -1 | 2 | 5 | 5 | 6 | 4 | 29 | 11 | 5 | 3 | 7 | 1 | 4 | 5 | 11 | 6 | 5 | 2 | 0 |
| DCT | chr13:94444363\|94445689-94445761 | 13 | 18 | 20 | 11 | 8 | 12 | 0 | 18 | 6 | 2 | 3 | 3 | 1 | 2 | 1 | 2 | 2 | 3 | 3 | 2 | 2 | 0 | 0 | 7 | 10 | 10 | 7 | 8 | -1 | 8 | 7 | -1 | 22 | 18 | 6 | 6 | 9 | 6 | 48 | 12 | 12 | 7 | 9 | 16 | -1 | 12 | 21 | 21 | 7 | 5 | 14 |
| TMCO4 | chr1:19747478-19747491 | 0 | 0 | 0 | 0 | 0 | 0 | 0 | 0 | 0 | 4 | 5 | 7 | 1 | 3 | 2 | 1 | 24 | 4 | 7 | 1 | 7 | 0 | 1 | 0 | 0 | 0 | 0 | 0 | 0 | 0 | 0 | 0 | 0 | 0 | 0 | 0 | 0 | 0 | 12 | 0 | 0 | 0 | 0 | 0 | 0 | 0 | 0 | 0 | 0 | 0 | 0 |
| D2HGDH | chr2:241745337-241747703 | 8 | 10 | 4 | 14 | 14 | 12 | 13 | 7 | 4 | 2 | 4 | 5 | 4 | 4 | 4 | 4 | 4 | 6 | 3 | 6 | 4 | 9 | 8 | 13 | 9 | 16 | 8 | 15 | 10 | 10 | 9 | 4 | 5 | 13 | 10 | 4 | 8 | 12 | 10 | 39 | 9 | 8 | 7 | 10 | 7 | 9 | 11 | 8 | 10 | 11 | 9 |
| PIEZO2 | chr18:10735232-10735330 | 0 | 1 | 4 | 0 | 0 | 0 | 5 | 7 | 0 | 3 | 0 | 0 | 0 | 4 | 6 | 1 | 2 | 0 | 0 | 0 | 0 | 0 | 0 | 3 | 5 | 1 | 2 | 5 | -1 | 0 | 0 | 0 | 3 | 1 | 3 | 0 | 0 | 1 | 17 | 35 | 5 | 2 | 0 | 2 | 0 | 6 | 11 | 1 | 0 | 0 | -1 |
| R3HDM1 | chr2:135695528\|135695767-135695906 | 6 | 7 | 4 | 8 | 7 | 7 | 6 | 7 | 5 | 8 | 6 | 13 | 14 | 12 | 9 | 6 | 6 | 6 | 6 | 6 | 6 | 10 | 6 | 7 | 6 | 7 | 4 | 5 | 9 | 7 | 8 | 6 | 7 | 7 | 7 | 11 | 8 | 9 | 9 | 23 | 7 | 8 | 8 | 8 | 8 | 6 | 6 | 7 | 10 | 8 | 7 |
| TAB3 | chrX:30855817-30855932 | 0 | 0 | 6 | 0 | 0 | 0 | 0 | 0 | 25 | 15 | 21 | 1 | 3 | 14 | 9 | 16 | 9 | 23 | 15 | 7 | 10 | 0 | 0 | 0 | 0 | 0 | 0 | 0 | 0 | 3 | 3 | 4 | 0 | 0 | 0 | 1 | 0 | 9 | 1 | 17 | 2 | 0 | 0 | 0 | 1 | 1 | 8 | 2 | 0 | 0 | 0 |
| PXDN | chr2:1653916-1653980 | 0 | 0 | 1 | 1 | 0 | 0 | 1 | 0 | 12 | 27 | 11 | 11 | 25 | 33 | 27 | 12 | 29 | 19 | 12 | 5 | 19 | 0 | 1 | 1 | 0 | 0 | 0 | 0 | 1 | 1 | 0 | 1 | 0 | 0 | 0 | 0 | 0 | 6 | 0 | 27 | 1 | 0 | 0 | 0 | 0 | 0 | 0 | 0 | 0 | 1 | 0 |
| PHF21B | chr22:44889760-44889782 | 1 | 1 | 16 | 1 | 3 | 4 | 0 | 1 | 5 | 7 | 4 | 15 | 15 | 7 | 7 | 10 | 19 | 9 | 6 | 1 | 5 | 0 | 0 | 2 | 4 | 1 | 0 | 0 | 0 | 1 | 4 | 1 | 0 | 3 | 0 | 1 | 0 | 1 | 49 | 57 | 0 | 0 | 2 | 0 | 0 | 13 | 2 | 0 | 1 | 0 | -1 |
| SEMA4D | chr9:89367123\|89365682-89367158 | 3 | 3 | 7 | 3 | 3 | 1 | 3 | 2 | 4 | 15 | 3 | 31 | 42 | 18 | 21 | 3 | 7 | 6 | 2 | 0 | 1 | -1 | 0 | 12 | 7 | 5 | 2 | 5 | 0 | 0 | 0 | 2 | 2 | 4 | 2 | 0 | 1 | 2 | 10 | 45 | 4 | 2 | 4 | 6 | 5 | 6 | 7 | 4 | 1 | 4 | 3 |
| DOCK5 | chr8:25376543-25376665 | 2 | 4 | 2 | 4 | 5 | 4 | 0 | 2 | 3 | 2 | 3 | 3 | 7 | 2 | 4 | 3 | 5 | 4 | 5 | 3 | 2 | -1 | 0 | 13 | 4 | 9 | 2 | 12 | 5 | 5 | 3 | 2 | 2 | 2 | 2 | 8 | 8 | 3 | 2 | 22 | 4 | 2 | 1 | 3 | 2 | 4 | 3 | 1 | 3 | 4 | 1 |
| SEMA4D | chr9:89367123-89367158\|89372961 | 3 | 2 | 7 | 3 | 2 | 1 | 1 | 3 | 4 | 16 | 4 | 32 | 43 | 19 | 21 | 4 | 8 | 7 | 3 | 1 | 2 | -1 | 0 | 11 | 6 | 5 | 1 | 5 | -1 | 1 | 1 | 1 | 1 | 3 | 2 | 1 | 1 | 2 | 10 | 46 | 2 | 3 | 4 | 5 | 4 | 6 | 9 | 5 | 1 | 1 | 2 |
| RGS6 | chr14:72553290-72553356 | 1 | 1 | 0 | 1 | 1 | 1 | 0 | 1 | 33 | 39 | 14 | 21 | 13 | 41 | 40 | 27 | 29 | 23 | 15 | 4 | 26 | 0 | 0 | 0 | 0 | 1 | 1 | 0 | 0 | 1 | 0 | 0 | 0 | 1 | 0 | 2 | 5 | 0 | -1 | 14 | 3 | 3 | 3 | 1 | 1 | 2 | 0 | 0 | 1 | 2 | 1 |
| NBEAP1 | chr15:20669973-20670095\|20670070 | 7 | 7 | 3 | 6 | 7 | 6 | -1 | 8 | 10 | 9 | 10 | 8 | 11 | 9 | 7 | 11 | 12 | 8 | 11 | 14 | 15 | 0 | -1 | 9 | 7 | 6 | 3 | 8 | -1 | 10 | 10 | -1 | 5 | 8 | 9 | 9 | 10 | 16 | 11 | 6 | 38 | 10 | 10 | 6 | 9 | 13 | 22 | 6 | 7 | 12 | 21 |
| CALCOCO1 | chr12:53712775-53712846 | 8 | 5 | 2 | 3 | 3 | 4 | 6 | 6 | 2 | 1 | 1 | 1 | 1 | 1 | 1 | 2 | 2 | 1 | 1 | 2 | 2 | 3 | 2 | 3 | 6 | 2 | 10 | 2 | 3 | 1 | 1 | 5 | 5 | 7 | 7 | 3 | 6 | 3 | 10 | 2 | 4 | 25 | 28 | 6 | 5 | 5 | 2 | 4 | 2 | 5 | 7 |
| CTAGE5 | chr14:39292680-39292706 | 1 | 2 | 2 | 2 | 2 | 1 | 0 | 1 | 3 | 1 | 4 | 2 | 5 | 2 | 2 | 6 | 3 | 5 | 4 | 14 | 10 | 4 | 1 | 6 | 20 | 1 | 2 | 1 | 5 | 2 | 2 | 12 | 12 | 2 | 3 | 2 | 2 | 3 | 3 | 1 | 5 | 1 | 1 | 51 | 4 | 6 | 3 | 2 | 2 | 2 | 3 |
| SLC6A13 | chr12:223746-223845 | 5 | 6 | 8 | 2 | 2 | 2 | 0 | 6 | 3 | 2 | 2 | 2 | 2 | 2 | 1 | 2 | 1 | 1 | 2 | 2 | 2 | -1 | -1 | -1 | -1 | 1 | -1 | 4 | 2 | 1 | 3 | 16 | 25 | 6 | 0 | 2 | 2 | -1 | 0 | 2 | 3 | 4 | 0 | 39 | 13 | 7 | 2 | 4 | 0 | 0 | 8 |
| CACNB2 | chr10:18519717-18522238\|18527448 | 16 | 11 | 4 | 15 | 13 | 18 | 17 | 13 | 1 | 1 | 2 | 2 | 2 | 1 | 1 | 0 | 1 | 2 | 2 | 5 | 1 | 12 | -1 | 13 | 26 | 12 | 13 | 10 | 7 | 7 | 4 | 7 | 18 | 12 | 19 | 22 | 11 | 10 | 21 | 4 | 15 | 18 | 19 | 49 | 8 | 11 | 7 | 10 | 14 | 20 | -1 |
| FCGR2B | chr1:161675257-161675313 | 5 | 8 | 9 | 10 | 11 | 11 | 13 | 9 | 11 | 10 | 12 | 7 | 13 | 13 | 13 | 12 | 9 | 11 | 13 | 10 | 9 | -1 | 24 | 12 | 18 | 7 | 24 | 7 | 12 | 8 | 10 | 22 | 9 | 15 | 18 | 8 | 8 | 11 | 16 | 17 | 26 | 12 | 11 | 56 | 74 | 22 | 20 | 18 | 16 | 17 | 29 |

SFig5, pg10

| Gene Symbol | Exon Location (hg38, human) | Adipose - Subcutaneous | Adipose - Visceral (Omentum) | Adrenal Gland | Artery - Aorta | Artery - Coronary | Artery - Tibial | Bladder | Breast - Mammary Tissue | Brain - Amygdala | Brain - Anterior cingulate cortex (BA24) | Brain - Caudate (basal ganglia) | Brain - Cerebellar Hemisphere | Brain - Cerebellum | Brain - Cortex | Brain - Frontal Cortex (BA9) | Brain - Hippocampus | Brain - Hypothalamus | Brain - Nucleus accumbens (basal ganglia) | Brain - Putamen (basal ganglia) | Brain - Spinal cord (cervical c-1) | Brain - Substantia nigra | Cervix - Ectocervix | Cervix - Endocervix | Colon - Sigmoid | Colon - Transverse | Esophagus - Gastroesophageal Junction | Esophagus - Mucosa | Esophagus - Muscularis | Fallopian Tube | Heart - Atrial Appendage | Heart - Left Ventricle | Kidney - Cortex | Liver | Lung | Minor Salivary Gland | Muscle - Skeletal | Nerve - Tibial | Ovary | Pancreas | Pituitary | Prostate | Skin - Not Sun Exposed (Suprapubic) | Skin - Sun Exposed (Lower leg) | Small Intestine - Terminal Ileum | Spleen | Stomach | Testis | Thyroid | Uterus | Vagina | Whole Blood |
|---|---|---|---|---|---|---|---|---|---|---|---|---|---|---|---|---|---|---|---|---|---|---|---|---|---|---|---|---|---|---|---|---|---|---|---|---|---|---|---|---|---|---|---|---|---|---|---|---|---|---|---|---|
| RUNDC3A | chr17:44317037\|44317040\|44316844-44317123 | 5 | 6 | 7 | 6 | 8 | 6 | 0 | 7 | 4 | 4 | 4 | 3 | 3 | 3 | 4 | 4 | 4 | 5 | 4 | 5 | 4 | -1 | 0 | 4 | 8 | 5 | 5 | 5 | 0 | 5 | 5 | 7 | -1 | 7 | 7 | 6 | 8 | 7 | 4 | 6 | 5 | 6 | 5 | 6 | 22 | 6 | 11 | 7 | -1 | 5 | 12 |
| PLEKHA1 | chr10:122428082-122428146 | 0 | 0 | 0 | 0 | 0 | 0 | 0 | 0 | 0 | 0 | 0 | 0 | 0 | 0 | 0 | 0 | 0 | 0 | 0 | 0 | 0 | 0 | 0 | 0 | 0 | 0 | 0 | 0 | 0 | 0 | 0 | 0 | 0 | 0 | 0 | 0 | 0 | 0 | 0 | 0 | 0 | 0 | 0 | 0 | 0 | 0 | 27 | 0 | 0 | 0 | 0 |
| YWHAZ | chr8:100923219\|100923211-100923307 | 0 | 0 | 0 | 0 | 0 | 0 | 1 | 0 | 0 | 0 | 0 | 0 | 0 | 0 | 0 | 0 | 0 | 0 | 0 | 0 | 0 | 0 | 0 | 0 | 0 | 0 | 0 | 0 | 0 | 0 | 0 | 0 | 0 | 0 | 0 | 0 | 0 | 0 | 0 | 0 | 0 | 0 | 0 | 0 | 0 | 0 | 28 | 0 | 0 | 0 | 0 |
| CD37 | chr19:49337626-49337808 | 0 | 0 | 0 | 0 | 0 | 0 | 0 | 0 | 0 | 0 | 0 | 0 | 0 | 0 | 0 | 0 | 0 | 0 | 0 | 0 | 0 | 0 | 0 | 0 | 0 | 0 | 0 | 0 | 0 | 0 | 0 | 0 | 0 | 0 | 0 | 0 | 0 | 0 | 0 | 0 | 0 | 0 | 0 | 0 | 0 | 0 | 23 | 0 | 0 | 0 | 0 |
| TNFRSF21 | chr6:47267820-47267954 | 0 | 0 | 0 | 0 | 0 | 0 | 0 | 0 | 0 | 0 | 0 | 0 | 0 | 0 | 0 | 0 | 0 | 0 | 0 | 0 | 0 | 0 | 0 | 0 | 0 | 0 | 0 | 0 | 1 | 0 | 0 | 0 | 0 | 0 | 0 | 0 | 0 | 0 | 0 | 0 | 0 | 0 | 0 | 0 | 0 | 0 | 32 | 0 | 0 | 0 | 0 |
| FGD4 | chr12:32600410-32600509 | 0 | 1 | 0 | 0 | 0 | 0 | 0 | 0 | 0 | 1 | 0 | 0 | 0 | 0 | 0 | 0 | 0 | 0 | 0 | 0 | 0 | 0 | 0 | 0 | 1 | 0 | 0 | 1 | 1 | 0 | 0 | 0 | 1 | 0 | 0 | 0 | 0 | 0 | 1 | 0 | 1 | 0 | 0 | 1 | 0 | 0 | 55 | 0 | 0 | 0 | 0 |
| ADORA3 | chr1:111501195-111501359 | 0 | 0 | 0 | 0 | 0 | 0 | 0 | 0 | 0 | 0 | 0 | 0 | 0 | 0 | 0 | 0 | 0 | 0 | 0 | 0 | 0 | 0 | 0 | 0 | 0 | 0 | 0 | 0 | 0 | 0 | 0 | 0 | 0 | 0 | 0 | 0 | 0 | 0 | 0 | 0 | 0 | 0 | 0 | 0 | 0 | 0 | 22 | 0 | 0 | 0 | 0 |
| MAOA | chrX:43740826-43741022 | 0 | 0 | 0 | 0 | 0 | 0 | 0 | 0 | 0 | 0 | 0 | 0 | 0 | 0 | 0 | 0 | 0 | 0 | 0 | 1 | 0 | 0 | 0 | 0 | 0 | 0 | 0 | 0 | 0 | 0 | 0 | 0 | 0 | 0 | 0 | 1 | 0 | 0 | 0 | 0 | 0 | 0 | 0 | 0 | 0 | 0 | 28 | 0 | 0 | 0 | 0 |
| FAP | chr2:162214841-162215028 | 0 | 0 | 0 | 0 | 0 | 0 | 0 | 0 | 0 | 0 | 0 | 0 | 0 | 0 | 0 | 0 | 0 | 0 | 0 | 0 | 0 | 0 | 0 | 0 | 0 | 0 | 0 | 0 | 0 | 0 | 0 | 0 | 0 | 0 | 0 | 0 | 0 | 0 | 0 | 0 | 0 | 0 | 0 | 0 | 0 | 0 | 24 | 0 | 0 | 0 | 0 |
| TG | chr8:132932993-132933125 | 0 | 0 | 0 | 0 | 0 | 0 | 0 | 0 | 0 | 0 | 0 | 0 | 0 | 0 | 0 | 0 | 0 | 0 | 0 | 0 | 0 | 0 | 0 | 0 | 0 | 0 | 0 | 0 | 0 | 0 | 0 | 0 | 0 | 0 | 0 | 0 | 0 | 0 | 1 | 0 | 0 | 0 | 0 | 0 | 0 | 0 | 32 | 0 | 0 | 0 | 1 |
| CASQ2 | chr1:115726456-115726597 | 0 | 0 | 0 | 0 | 0 | 0 | 0 | 0 | 0 | 0 | 0 | 0 | 0 | 1 | 0 | 0 | 0 | 0 | 0 | 0 | 0 | 1 | 1 | 0 | 0 | 0 | 0 | 0 | 0 | 0 | 0 | 0 | 0 | 0 | 0 | 0 | 0 | 0 | 0 | 0 | 0 | 0 | 0 | 0 | 0 | 0 | 38 | 0 | 0 | 0 | 0 |
| CHCHD3 | chr7:133022406-133022495 | 0 | 0 | 0 | 0 | 0 | 0 | 0 | 0 | 0 | 0 | 0 | 0 | 0 | 0 | 0 | 0 | 0 | 0 | 0 | 0 | 0 | 0 | 1 | 0 | 0 | 0 | 0 | 0 | 0 | 0 | 0 | 0 | 0 | 0 | 0 | 0 | 0 | 0 | 0 | 0 | 0 | 0 | 0 | 0 | 0 | 0 | 23 | 0 | 0 | 0 | 0 |
| AL365273.1 | chr10:95849411-95849616 | 0 | 0 | 0 | 0 | 0 | 0 | 0 | 0 | 0 | 0 | 0 | 0 | 1 | 0 | 0 | 0 | 0 | 0 | 0 | 0 | 0 | 0 | 0 | 0 | 0 | 0 | 0 | 0 | 0 | 0 | 0 | 0 | 0 | 0 | 0 | 0 | 1 | 0 | 0 | 1 | 0 | 0 | 0 | 0 | 0 | 0 | 35 | 0 | 0 | 0 | 0 |
| C1orf106 | chr1:200908116-200908418 | 0 | 0 | 0 | 0 | 0 | 0 | 0 | 0 | 0 | 0 | 0 | 0 | 0 | 1 | 0 | 0 | 0 | 0 | 0 | 0 | 0 | 0 | 0 | 0 | 0 | 0 | 0 | 1 | 0 | 0 | 0 | 0 | 0 | 0 | 0 | 0 | 0 | 0 | 0 | 0 | 0 | 0 | 0 | 0 | 0 | 0 | 35 | 0 | 0 | 0 | 0 |
| DENND1A | chr9:123452826-123452908 | 1 | 1 | 1 | 1 | 1 | 1 | 1 | 2 | 2 | 2 | 2 | 2 | 2 | 1 | 1 | 2 | 2 | 1 | 1 | 2 | 1 | 2 | 1 | 1 | 2 | 1 | 1 | 1 | 2 | 2 | 1 | 1 | 1 | 1 | 1 | 1 | 1 | 2 | 1 | 1 | 1 | 2 | 2 | 2 | 1 | 1 | 70 | 2 | 2 | 1 | 2 |
| ANKAR | chr2:189743275-189743474 | 0 | 0 | 0 | 0 | 0 | 0 | 0 | 1 | 1 | 0 | 0 | 0 | 1 | 0 | 0 | 0 | 1 | 1 | 0 | 1 | 1 | 0 | 0 | 0 | 0 | 0 | 1 | 0 | 0 | 0 | 0 | 1 | 1 | 0 | 1 | 0 | 0 | 1 | 1 | 2 | 1 | 0 | 1 | 1 | 1 | 0 | 64 | 1 | 0 | 0 | 1 |
| ANKFN1 | chr17:56451931-56452053 | 0 | 0 | 0 | 0 | 2 | 0 | 0 | 0 | 1 | 0 | 0 | 0 | -1 | 0 | 0 | 0 | 0 | 0 | 0 | 0 | 1 | 0 | 0 | 0 | 0 | 0 | 0 | 0 | 0 | 0 | 0 | -1 | 0 | 0 | 0 | 0 | 0 | 0 | 0 | 1 | 0 | 0 | 0 | 0 | 0 | 0 | 65 | 0 | 0 | 0 | 0 |
| MTMR6 | chr13:25260574-25260687 | 1 | 0 | 0 | 0 | 0 | 0 | 0 | 0 | 0 | 0 | 0 | 0 | 0 | 0 | 0 | 0 | 0 | 0 | 0 | 0 | 0 | 0 | 0 | 0 | 0 | 0 | 0 | 0 | 0 | 0 | 0 | 0 | 0 | 0 | 0 | 0 | 0 | 0 | 0 | 0 | 0 | 0 | 1 | 0 | 0 | 0 | 22 | 0 | 0 | 0 | 0 |
| TRAF3IP3 | chr1:209773456-209773606 | 0 | 0 | 0 | 0 | 1 | 0 | 1 | 0 | 0 | 0 | 0 | 0 | 0 | 0 | 0 | 0 | 0 | 1 | 1 | 0 | 0 | 0 | 0 | 0 | 0 | 1 | 0 | 0 | 0 | 0 | 0 | 0 | 1 | 0 | 0 | 2 | 0 | 0 | 0 | 0 | 0 | 1 | 1 | 0 | 0 | 0 | 38 | 0 | 0 | 0 | 0 |
| MCC | chr5:113102161-113102316 | 0 | 0 | 0 | 0 | 0 | 0 | 0 | 0 | 0 | 0 | 0 | 0 | 0 | 0 | 0 | 0 | 0 | 0 | 0 | 0 | 0 | 0 | 0 | 0 | 0 | 0 | 0 | 0 | 0 | 1 | 0 | 0 | 0 | 0 | 0 | 0 | 0 | 0 | 0 | 0 | 0 | 0 | 0 | 0 | 0 | 0 | 21 | 0 | 0 | 0 | 0 |
| RPH3A | chr12:112892279-112892385 | 0 | 0 | 1 | 0 | 0 | 0 | 0 | -1 | 0 | 0 | 0 | 0 | 1 | 0 | 0 | 0 | 0 | 0 | 0 | 0 | 0 | -1 | -1 | 0 | 0 | 0 | 0 | -1 | 0 | 0 | 0 | 0 | 0 | 0 | 0 | 2 | 0 | 0 | 0 | 0 | 0 | -1 | 1 | 0 | 0 | 0 | 72 | 0 | 0 | 0 | 0 |
| AMOTL1 | chr11:94857664-94859414 | 0 | 0 | 0 | 0 | 0 | 0 | 0 | 0 | 1 | 0 | 0 | 0 | 0 | 0 | 0 | 0 | 0 | 1 | 0 | 1 | 0 | 0 | 0 | 0 | 0 | 0 | 0 | 0 | 1 | 1 | 0 | 0 | 1 | 0 | 1 | 2 | 1 | 0 | 0 | 0 | 1 | 0 | 0 | 0 | 0 | 0 | 39 | 0 | 0 | 0 | 1 |
| SEMA5A | chr5:9193225-9193294 | 1 | 1 | 1 | 1 | 0 | 0 | 0 | 0 | 0 | 0 | 0 | 1 | 1 | 0 | 0 | 0 | 0 | 1 | 0 | 1 | 0 | 0 | 0 | 1 | 2 | 0 | 1 | 1 | 0 | 0 | 0 | 0 | 0 | 1 | 0 | 1 | 0 | 0 | 1 | 1 | 1 | 0 | 1 | 1 | 0 | 1 | 59 | 1 | 2 | 1 | -1 |
| SMAD2 | chr18:47868052-47868097 | 1 | 1 | 0 | 1 | 0 | 1 | 1 | 1 | 0 | 0 | 0 | 0 | 0 | 0 | 0 | 0 | 0 | 0 | 0 | 0 | 1 | 1 | 1 | 1 | 1 | 0 | 1 | 1 | 1 | 0 | 0 | 1 | 1 | 1 | 2 | 1 | 1 | 1 | 0 | 1 | 1 | 1 | 1 | 1 | 1 | 1 | 34 | 1 | 1 | 1 | 1 |
| UNC5D | chr8:35747568-35747752 | 0 | 0 | 0 | -1 | -1 | -1 | -1 | 0 | 0 | 0 | 0 | 0 | 0 | 0 | 0 | 0 | 0 | 1 | 0 | 0 | 0 | 0 | 0 | 0 | 0 | 0 | 0 | 0 | -1 | 0 | 0 | 0 | 0 | 0 | 0 | 0 | 0 | 0 | 0 | 0 | 0 | 0 | 0 | 0 | 0 | 0 | 42 | 0 | 0 | 0 | 0 |
| ATPAF2 | chr17:18025251-18025425 | 0 | 0 | 0 | 0 | 0 | 0 | 0 | 0 | 0 | 0 | 0 | 1 | 1 | 0 | 0 | 0 | 0 | 0 | 0 | 1 | 0 | 0 | 0 | 0 | 0 | 0 | 0 | 0 | 0 | 0 | 0 | 0 | 0 | 0 | 0 | 0 | 0 | 1 | 0 | 0 | 1 | 0 | 0 | 0 | 1 | 0 | 22 | 0 | 1 | 1 | 0 |
| ADAMTSL3 | chr15:83732825-83732905 | 0 | 0 | 0 | 0 | 0 | 0 | 0 | 0 | 0 | 0 | 0 | 0 | 1 | 1 | 0 | 1 | 0 | 1 | 1 | 1 | 0 | 0 | 0 | 0 | 0 | 0 | 0 | 0 | 0 | 0 | 0 | 0 | 0 | 0 | 0 | 0 | 0 | 1 | 0 | 0 | 0 | 0 | 0 | 0 | 0 | 0 | 22 | 0 | 0 | 0 | 0 |
| STAB2 | chr12:103701737-103701807 | 0 | 0 | 0 | 0 | 0 | 0 | 0 | 0 | 0 | 0 | 0 | -1 | 0 | 0 | 0 | -1 | 0 | 0 | -1 | -1 | -1 | 0 | -1 | 0 | 0 | 0 | 0 | 0 | 0 | 1 | 0 | 0 | 0 | 0 | 0 | 0 | 0 | 0 | 0 | 0 | 0 | 0 | 0 | 0 | 0 | 0 | 24 | 0 | 0 | 0 | 0 |
| PHGDH | chr1:119724319-119724875 | 0 | 0 | 0 | 0 | 0 | 0 | 0 | 0 | 0 | 0 | 0 | 0 | 0 | 0 | 0 | 0 | 0 | 0 | 0 | 0 | 0 | 0 | 0 | 0 | 0 | 0 | 0 | 0 | 0 | 0 | 0 | 0 | 1 | 0 | 1 | 0 | 1 | 0 | 1 | 0 | 1 | 0 | 0 | 0 | 1 | 0 | 24 | 0 | 0 | 0 | 1 |
| PFKP | chr10:3125097-3125249\|3128287 | 0 | 0 | 1 | 0 | 0 | 0 | 1 | 0 | 0 | 0 | 0 | 1 | 1 | 0 | 0 | 0 | 0 | 0 | 0 | 0 | 0 | 0 | 0 | 1 | 3 | 0 | 0 | 0 | 0 | 0 | 0 | 1 | 1 | 0 | 0 | 1 | 0 | 0 | 1 | 2 | 1 | 0 | 0 | 3 | 1 | 1 | 62 | 1 | 0 | 0 | 1 |
| MAGI2 | chr7:78367211-78367299 | 1 | 0 | 0 | 0 | 1 | 1 | 1 | 1 | 1 | 0 | 0 | 0 | 0 | 1 | 0 | 0 | 1 | 0 | 0 | 0 | 0 | 0 | 0 | 0 | 0 | 0 | 0 | 0 | 0 | 1 | 2 | 0 | 0 | 1 | 0 | 1 | 0 | 0 | 2 | 1 | 2 | 1 | 2 | 0 | 0 | 0 | 60 | 1 | 0 | 0 | 0 |
| PCDHAC2 | chr5:140979674-140979926 | 0 | 0 | 0 | 0 | 0 | 0 | 0 | 0 | 0 | 0 | 0 | 0 | 0 | 0 | 0 | 0 | 0 | 0 | 0 | 0 | 0 | 0 | 1 | 0 | 0 | 0 | 0 | 0 | 0 | 0 | 1 | 0 | 0 | 0 | 0 | 0 | 0 | 0 | 0 | 0 | 0 | 0 | 0 | 0 | 0 | 0 | 29 | 0 | 0 | 0 | 1 |
| AC008074.1 | chr2:64583759-64583892 | 0 | 0 | 0 | 0 | 0 | 0 | 0 | 0 | 0 | 0 | 0 | 0 | 0 | 0 | 0 | 0 | 0 | 0 | 0 | 0 | 0 | -1 | 0 | 0 | -1 | 0 | 0 | 0 | -1 | 0 | 0 | 0 | -1 | 0 | 0 | -1 | 0 | -1 | 0 | 0 | 0 | 1 | 0 | 0 | 0 | 0 | 44 | 0 | 0 | 0 | -1 |
| ATP2C2 | chr16:84438184-84438652 | 0 | 0 | 0 | -1 | 0 | 0 | 0 | 0 | 0 | 0 | 0 | 0 | 0 | 1 | 1 | 0 | 0 | 0 | 1 | 1 | 0 | 0 | 0 | 0 | 0 | 1 | 0 | 0 | 1 | 0 | 0 | 0 | 1 | 0 | 0 | 0 | 1 | 0 | 0 | 1 | 0 | 0 | 0 | 0 | 1 | 0 | 46 | 0 | 0 | 1 | 1 |
| SEC24D | chr4:118816744-118816850 | 1 | 0 | 0 | 1 | 0 | 1 | 1 | 1 | 0 | 0 | 0 | 0 | 1 | 0 | 1 | 0 | 1 | 1 | 1 | 0 | 1 | 0 | 0 | 1 | 0 | 1 | 0 | 1 | 0 | 1 | 0 | 0 | 0 | 1 | 0 | 1 | 1 | 1 | 0 | 1 | 1 | 1 | 1 | 0 | 0 | 1 | 25 | 1 | 1 | 0 | 0 |
| AL109923.1 | chr20:34430188-34430337 | 1 | 1 | 0 | 0 | 0 | 1 | 1 | 1 | 0 | 0 | 1 | 1 | 2 | 0 | 0 | 0 | 0 | 0 | 0 | 0 | 0 | 0 | 0 | 1 | 1 | 0 | 0 | 1 | 1 | 1 | 1 | 1 | 1 | 1 | 1 | 0 | 1 | 1 | 1 | 1 | 1 | 1 | 1 | 1 | 1 | 1 | 28 | 1 | 1 | 1 | 0 |
| SNX13 | chr7:17831074-17831247\|17832503 | 1 | 1 | 1 | 0 | 1 | 1 | 0 | 1 | 1 | 1 | 2 | 2 | 1 | 1 | 2 | 4 | 1 | 2 | 1 | 1 | 1 | 0 | 1 | 1 | 1 | 0 | 1 | 0 | 1 | 1 | 0 | 1 | 2 | 1 | 1 | 1 | 1 | 1 | 1 | 1 | 1 | 1 | 0 | 1 | 2 | 1 | 31 | 0 | 1 | 1 | 1 |
| CD47 | chr3:108055520\|108054253-108055566 | 2 | 1 | 1 | 1 | 1 | 2 | 2 | 2 | 4 | 2 | 3 | 1 | 1 | 2 | 2 | 2 | 2 | 3 | 4 | 1 | 2 | 2 | 1 | 1 | 1 | 1 | 1 | 1 | 2 | 3 | 3 | 1 | 1 | 2 | 1 | 3 | 2 | 2 | 1 | 2 | 2 | 3 | 3 | 3 | 2 | 2 | 61 | 1 | 1 | 1 | 2 |
| RP5-1052I5.2 | chr1:87099290-87099372 | 0 | 0 | 0 | 0 | 0 | 0 | 1 | 0 | 0 | 0 | 0 | 0 | 0 | 0 | 0 | 0 | 0 | 0 | 0 | 0 | 0 | 0 | 0 | 0 | 1 | 0 | 0 | 0 | 0 | 0 | 0 | 1 | 0 | 0 | 0 | 0 | 0 | 0 | 1 | 0 | 0 | 0 | 0 | 1 | 1 | 0 | 24 | 1 | 0 | 0 | 1 |
| PDGFC | chr4:156835915-156836007 | 0 | 1 | 0 | 0 | 0 | 0 | 0 | 0 | 1 | 0 | 1 | 0 | 0 | 1 | 1 | 0 | 1 | 0 | 0 | 0 | 0 | 0 | 0 | 0 | 1 | 0 | 1 | 0 | 0 | 1 | 1 | 0 | 1 | 1 | 1 | 0 | 0 | 0 | 1 | 1 | 0 | 0 | 0 | 0 | 1 | 1 | 24 | 1 | 0 | 0 | 1 |
| SLC44A3 | chr1:94844366-94844765 | 0 | 0 | 0 | 1 | 0 | 1 | 0 | 0 | 0 | 0 | 0 | 0 | 0 | 0 | 1 | 0 | 0 | 0 | 0 | 0 | 0 | 0 | -1 | 0 | 0 | 1 | 0 | 0 | 0 | 0 | 1 | 0 | 0 | 0 | 0 | 1 | 1 | 1 | 0 | 0 | 0 | 0 | 0 | 0 | 1 | 0 | 39 | 0 | 1 | 1 | 0 |
| HIBCH | chr2:190245088-190245370 | 0 | 0 | 0 | 1 | 0 | 0 | 0 | 0 | 0 | 0 | 0 | 0 | 0 | 0 | 0 | 0 | 0 | 0 | 0 | 0 | 0 | 0 | 0 | 0 | 0 | 0 | 0 | 0 | 0 | 0 | 0 | 0 | 0 | 1 | 0 | 0 | 0 | 1 | 0 | 0 | 0 | 0 | 0 | 0 | 0 | 0 | 21 | 0 | 0 | 1 | 1 |
| MAST4 | chr5:66902649-66902736 | 0 | 0 | 0 | 0 | 0 | 0 | 0 | 0 | 0 | 0 | 0 | 0 | 2 | 0 | 0 | 0 | 0 | 0 | 0 | 0 | 0 | 0 | 0 | 0 | 0 | 0 | 0 | 0 | 0 | 0 | 0 | 0 | 0 | 0 | 1 | 0 | 0 | 0 | 0 | 0 | 0 | 0 | 0 | 0 | 0 | 0 | 29 | 0 | 0 | 0 | 1 |
| PDE1A | chr2:182195613-182195758 | 1 | 0 | 0 | 0 | 0 | 0 | 0 | 1 | 1 | 0 | 1 | 0 | 0 | 0 | 0 | 0 | 1 | 1 | 1 | 1 | 1 | 1 | 0 | 0 | 0 | 0 | 1 | 0 | 0 | 0 | 0 | 0 | 1 | 0 | 0 | 1 | 1 | 0 | 0 | 1 | 0 | 1 | 1 | 1 | 1 | 0 | 22 | 0 | 0 | 0 | 1 |
| NEK11 | chr3:131133246-131133355 | 0 | 0 | 0 | 0 | 0 | 0 | 0 | 0 | 0 | 0 | 0 | 0 | 1 | 0 | 1 | 0 | 0 | 0 | 0 | 0 | 0 | 0 | 0 | 0 | 0 | 1 | 0 | 0 | 2 | 0 | 0 | 0 | 1 | 0 | 0 | 0 | 1 | 0 | 0 | 0 | 1 | 0 | 0 | 1 | 0 | 0 | 30 | 0 | 0 | 0 | 1 |
| ATP11B | chr3:182829923-182830002 | 0 | 1 | 0 | 0 | 0 | 0 | 0 | 0 | 0 | 0 | 0 | 0 | 0 | 0 | 0 | 0 | 0 | 0 | 0 | 0 | 0 | 0 | 0 | 0 | 0 | 0 | 0 | 0 | 0 | 0 | 0 | 0 | 0 | 1 | 0 | 0 | 1 | 0 | 0 | 0 | 0 | 0 | 0 | 1 | 1 | 1 | 21 | 0 | 0 | 0 | 1 |
| ATPAF1 | chr1:46663833-46663923 | 1 | 1 | 0 | 1 | 1 | 1 | 1 | 1 | 0 | 0 | 0 | 0 | 0 | 0 | 0 | 0 | 0 | 0 | 0 | 0 | 0 | 1 | 0 | 0 | 1 | 0 | 1 | 0 | 0 | 0 | 0 | 0 | 0 | 1 | 1 | 0 | 1 | 1 | 0 | 0 | 1 | 2 | 3 | 1 | 1 | 0 | 42 | 1 | 1 | 1 | 1 |
| TBC1D32 | chr6:121224363-121224451 | 0 | 1 | 0 | 0 | 0 | 1 | 0 | 1 | 0 | 0 | 0 | 1 | 1 | 0 | 0 | 0 | 0 | 0 | 1 | 0 | 0 | 0 | 0 | 0 | 0 | 0 | 1 | 0 | 0 | 1 | 0 | 0 | 1 | 0 | 0 | 0 | 0 | 0 | 0 | 0 | 0 | 0 | 0 | 1 | 0 | 0 | 22 | 1 | 0 | 0 | 0 |
| SMARCC1 | chr3:47765615-47765658 | 1 | 1 | 0 | 1 | 1 | 0 | 0 | 1 | 0 | 1 | 0 | 1 | 1 | 1 | 0 | 0 | 0 | 0 | 1 | 0 | 0 | 1 | 0 | 0 | 0 | 1 | 0 | 0 | 0 | 1 | 1 | 0 | 1 | 1 | 1 | 1 | 1 | 1 | 1 | 1 | 1 | 1 | 1 | 1 | 1 | 0 | 24 | 1 | 0 | 0 | 1 |
| PLEKHA3 | chr2:178502336-178502483 | 0 | 0 | 0 | 0 | 0 | 0 | 0 | 0 | 4 | 2 | 2 | 1 | 1 | 0 | 1 | 0 | 1 | 2 | 2 | 0 | 0 | 0 | 0 | 0 | 0 | 0 | 0 | 0 | 0 | 1 | 0 | 0 | 0 | 0 | 0 | 0 | 0 | 0 | 0 | 0 | 0 | 0 | 0 | 0 | 0 | 0 | 22 | 0 | 1 | 0 | 0 |
| MYO3B | chr2:170371573-170371716 | 0 | 0 | 0 | 0 | 0 | 0 | 0 | 0 | 0 | 0 | 0 | 0 | -1 | 0 | -1 | 0 | 0 | 0 | 0 | 0 | 0 | -1 | 0 | 0 | 0 | 0 | 0 | 0 | -1 | 0 | 0 | 0 | 0 | 0 | 0 | 0 | 0 | 0 | 0 | 0 | 0 | 0 | 0 | 0 | 0 | 0 | 26 | 0 | 0 | -1 | 1 |
| SDK1 | chr7:4219030-4219627 | 1 | 1 | 0 | 1 | 1 | 1 | 0 | 1 | 1 | 0 | 1 | 0 | 0 | 1 | 0 | 0 | 0 | 1 | 1 | 0 | 0 | 0 | 1 | 1 | 1 | 0 | 0 | 1 | 1 | 0 | 0 | 1 | 0 | 0 | 0 | 0 | 1 | 1 | 0 | 0 | 1 | 0 | 0 | 1 | 1 | 1 | 26 | 1 | 0 | 0 | 2 |
| HPSE2 | chr10:98466657-98472691 | 0 | 1 | 0 | 0 | 0 | 0 | 0 | 0 | 0 | 0 | 0 | 0 | 0 | 0 | 0 | 0 | 0 | 0 | 0 | 0 | 0 | 0 | 0 | 0 | 0 | 0 | 0 | 0 | 0 | 0 | 0 | 0 | 0 | 0 | 0 | 0 | 0 | 1 | 0 | 0 | 0 | 0 | 0 | 0 | 0 | 0 | 21 | 0 | 0 | 0 | -1 |
| MPP7 | chr10:28081049-28081119 | 0 | 0 | 0 | 0 | 0 | 0 | 0 | 1 | 0 | 1 | 0 | 1 | 1 | 0 | 1 | 0 | 0 | 1 | 3 | 1 | 0 | 0 | 0 | 0 | 0 | 0 | 0 | 0 | 1 | 1 | 1 | 1 | 1 | 0 | 0 | 1 | 1 | 0 | 1 | 1 | 1 | 0 | 0 | 0 | 1 | 0 | 24 | 0 | 0 | 0 | 0 |
| C7orf55-LUC7L2 | chr7:139369179\|139369127-139369251 | 1 | 1 | 1 | 1 | 0 | 1 | -1 | 1 | 1 | 2 | 1 | -1 | 1 | 3 | 3 | 1 | 2 | 0 | 1 | 1 | 1 | 1 | 0 | 1 | 1 | 1 | 1 | 1 | 1 | 0 | 1 | 2 | 1 | 1 | 1 | 1 | 0 | 1 | 1 | 1 | 1 | 1 | 1 | 1 | 1 | 2 | 41 | 1 | 2 | 1 | 1 |
| NBEA | chr13:35589486-35589815 | 0 | 2 | 0 | 1 | 1 | 0 | 0 | 1 | 1 | 1 | 0 | 0 | 1 | 0 | 0 | 1 | 0 | 0 | 0 | 4 | 0 | 0 | 0 | 1 | 0 | 0 | 1 | 0 | 0 | 1 | 0 | 1 | 3 | 2 | 0 | 1 | 2 | 2 | 1 | 2 | 1 | 1 | 1 | 0 | 0 | 0 | 51 | 1 | 1 | 1 | 1 |
| ADD2 | chr2:70665680-70665790 | 0 | 0 | 0 | 1 | 0 | 0 | 0 | 0 | 0 | 0 | 0 | 0 | 0 | 0 | 0 | 0 | 0 | 0 | 0 | 0 | 0 | 0 | 0 | 0 | 0 | 1 | 0 | 0 | 0 | 0 | 0 | 0 | 0 | 0 | 0 | 1 | 0 | 0 | 0 | 0 | 1 | 0 | 0 | 0 | 1 | 0 | 20 | 0 | 0 | 0 | 0 |
| CENPJ | chr13:24884703-24884793 | 2 | 1 | 1 | 1 | 1 | 1 | 1 | 1 | 1 | 0 | 0 | 1 | 0 | 0 | 0 | 0 | 0 | 0 | 1 | 1 | 1 | 2 | 1 | 1 | 2 | 1 | 2 | 1 | 1 | 1 | 1 | 1 | 1 | 1 | 2 | 1 | 2 | 1 | 1 | 1 | 1 | 1 | 1 | 2 | 2 | 1 | 28 | 1 | 1 | 2 | 1 |
| IQSEC2 | chrX:53236957-53237516\|53237501 | 1 | 1 | 0 | 1 | 1 | 1 | 1 | 1 | 0 | 0 | 0 | 1 | 1 | 0 | 0 | 0 | 0 | 0 | 0 | 0 | 1 | 0 | 2 | 1 | 1 | 1 | 1 | 1 | 1 | 1 | 1 | 1 | 1 | 1 | 1 | 1 | 1 | 0 | 1 | 1 | 1 | 1 | 0 | 1 | 1 | 0 | 24 | 1 | 1 | 1 | 1 |
| STXBP5 | chr6:147308502-147308592 | 1 | 0 | 0 | 0 | 0 | 0 | 0 | 1 | 0 | 0 | 0 | 0 | 0 | 0 | 0 | 0 | 0 | 0 | 0 | 0 | 0 | 0 | 0 | 0 | 0 | 0 | 0 | 0 | 0 | 0 | 0 | 0 | 0 | 1 | 1 | 0 | 0 | 0 | 1 | 0 | 0 | 1 | 1 | 1 | 1 | 1 | 22 | 1 | 1 | 0 | 1 |
| TPM3 | chr1:154191629-154191733 | 0 | 1 | 1 | 0 | 0 | 0 | 0 | 0 | 0 | 0 | 0 | 0 | 0 | 0 | 0 | 1 | 0 | 0 | 0 | 0 | 0 | -1 | -1 | 0 | 0 | 0 | 0 | 0 | 6 | 0 | 0 | 0 | 0 | 1 | 0 | 0 | 0 | 1 | 0 | 1 | 0 | 0 | 0 | 1 | 2 | 1 | 73 | 0 | 1 | 1 | 0 |
| RP11-426C22.7 | chr16:29227488-29227749 | 0 | 0 | 0 | -1 | -1 | 0 | -1 | 0 | 0 | 0 | 0 | 0 | 0 | 0 | 0 | 0 | 0 | 0 | 0 | 0 | 0 | 0 | -1 | 0 | 0 | 0 | 0 | 0 | -1 | 0 | 0 | 0 | 0 | 0 | -1 | 0 | 0 | 0 | 0 | 0 | 0 | 0 | -1 | 0 | 0 | 0 | 28 | 0 | 0 | -1 | 0 |
| SESTD1 | chr2:179141595-179142101 | 0 | 0 | 0 | 0 | 0 | 0 | 0 | 0 | 0 | 0 | 0 | 1 | 1 | 1 | 0 | 0 | 0 | 0 | 0 | 0 | 0 | 0 | 0 | 0 | 0 | 0 | 0 | 0 | 0 | 1 | 1 | 0 | 2 | 1 | 0 | 1 | 0 | 1 | 1 | 1 | 0 | 0 | 0 | 0 | 1 | 1 | 25 | 0 | 0 | 0 | 1 |
| DUSP4 | chr8:29345370-29346074 | 0 | 0 | 0 | 1 | 0 | 1 | 0 | 0 | 3 | 1 | 1 | 2 | 2 | 1 | 1 | 1 | 2 | 1 | 2 | 0 | 0 | 0 | 0 | 0 | 0 | 0 | 0 | 0 | 0 | 0 | 0 | 0 | 0 | 0 | 0 | 1 | 1 | 1 | 1 | 0 | 0 | 0 | 0 | 1 | 0 | 0 | 24 | 1 | 1 | 0 | 1 |
| SNX13 | chr7:17888560-17888788 | 0 | 1 | 0 | 0 | 0 | 0 | 1 | 0 | 0 | 0 | 0 | 0 | 0 | 0 | 0 | 1 | 0 | 0 | 0 | 0 | 0 | 0 | 1 | 0 | 0 | 0 | 0 | 0 | 1 | 0 | 0 | 0 | 0 | 1 | 0 | 0 | 0 | 0 | 0 | 1 | 1 | 1 | 0 | 1 | 0 | 1 | 30 | 0 | 0 | 1 | 3 |
| C15orf65 | chr15:55417698-55417759 | 0 | 0 | 0 | 0 | 0 | 0 | 0 | 1 | 1 | 0 | 1 | 0 | 1 | 0 | 1 | 1 | 0 | 1 | 0 | 1 | 1 | 0 | 0 | 0 | 0 | 0 | 0 | 0 | 0 | 0 | 1 | 0 | 1 | 1 | 0 | 0 | 1 | 1 | 0 | 0 | 0 | 0 | 0 | 0 | 0 | 1 | 21 | 0 | 0 | 0 | 1 |
| LRR1 | chr14:49612472-49612624 | 0 | 1 | 1 | 0 | 0 | 0 | 0 | 1 | 1 | 1 | 1 | 0 | 0 | 0 | 0 | 0 | 0 | 0 | 1 | 0 | 0 | 0 | 0 | 0 | 1 | 0 | 0 | 0 | 1 | 0 | 0 | 0 | 1 | 1 | 1 | 0 | 0 | 0 | 1 | 1 | 1 | 0 | 0 | 1 | 1 | 1 | 26 | 0 | 0 | 0 | 1 |
| AC005562.1 | chr17:30615896-30615980 | 4 | 2 | 1 | 2 | 2 | 2 | 2 | 4 | 0 | 1 | 1 | 5 | 4 | 1 | 1 | 1 | 1 | 0 | 1 | 1 | 0 | 2 | 0 | 2 | 2 | 2 | 2 | 2 | 4 | 2 | 1 | 2 | 1 | 2 | 1 | 2 | 2 | 2 | 2 | 1 | 2 | 4 | 4 | 2 | 3 | 2 | 60 | 1 | 2 | 2 | 2 |
| NPAS2 | chr2:100994424\|100994372-100994580 | 0 | 1 | 1 | 1 | 1 | 1 | 2 | 1 | 1 | 0 | 0 | 1 | 1 | 0 | 0 | 1 | 1 | 1 | 0 | 1 | 1 | 1 | 1 | 1 | 1 | 1 | 0 | 1 | 1 | 1 | 1 | 2 | 1 | 1 | 1 | 1 | 1 | 2 | 2 | 1 | 1 | 1 | 0 | 1 | 2 | 1 | 27 | 2 | 1 | 1 | 1 |
| CAPN3 | chr15:42385679-42385956 | 0 | 1 | 0 | 0 | 0 | 0 | 0 | 0 | 1 | 1 | 2 | 1 | 1 | 1 | 1 | 2 | 1 | 1 | 2 | 0 | 1 | 0 | 2 | 0 | 0 | 0 | 0 | 0 | 1 | 0 | 0 | 1 | 0 | 0 | 0 | 0 | 1 | 1 | 1 | 0 | 0 | 0 | 0 | 0 | 0 | 0 | 20 | 0 | 0 | 0 | 1 |
| DHX57 | chr2:38816095-38816185 | 1 | 1 | 1 | 1 | 1 | 1 | 0 | 1 | 0 | 0 | 1 | 1 | 1 | 0 | 0 | 1 | 0 | 1 | 1 | 1 | 0 | 0 | 0 | 1 | 1 | 1 | 1 | 1 | 0 | 1 | 1 | 1 | 1 | 1 | 1 | 1 | 1 | 1 | 0 | 1 | 1 | 1 | 1 | 1 | 2 | 1 | 23 | 1 | 1 | 1 | 1 |
| HOXB-AS1 | chr17:48548925\|48548912-48549057 | 1 | 1 | 2 | 1 | 1 | 1 | 0 | 1 | 0 | -1 | -1 | 0 | 0 | 0 | 0 | 0 | 0 | 0 | -1 | 0 | 0 | 0 | 3 | 1 | 0 | 0 | 1 | 0 | 0 | 1 | 1 | 2 | 1 | 2 | -1 | 1 | 1 | 4 | 1 | 1 | 1 | 2 | 3 | 2 | 2 | 1 | 70 | 1 | 1 | 1 | 5 |
| MYO3B | chr2:170527095-170527202 | 0 | 0 | -1 | 0 | 0 | 0 | 0 | 1 | 0 | 0 | 0 | 0 | 0 | 0 | -1 | 0 | 0 | 0 | 0 | 0 | 0 | 0 | 0 | -1 | 0 | -1 | 0 | 0 | 0 | 0 | 0 | 0 | 0 | 0 | 0 | 1 | 1 | 0 | 1 | 0 | 0 | 0 | 1 | 0 | -1 | 0 | 30 | 0 | 0 | 0 | 1 |
| COL11A1 | chr1:102925501-102925618 | 1 | 0 | 0 | 0 | 1 | 1 | 0 | 1 | 0 | 0 | 0 | 0 | 0 | 0 | 0 | 0 | 0 | 0 | 0 | 0 | 0 | 0 | 0 | 0 | 0 | 0 | -1 | 0 | 0 | 0 | 0 | 0 | -1 | 1 | 0 | 1 | 1 | 0 | 0 | 0 | 0 | 0 | 0 | 0 | 1 | 0 | 25 | 0 | 0 | 0 | 0 |
| PDZD8 | chr10:117285694-117285799 | 1 | 2 | 2 | 2 | 2 | 2 | 2 | 2 | 1 | 2 | 2 | 2 | 3 | 2 | 2 | 3 | 3 | 2 | 3 | 2 | 3 | 4 | 1 | 3 | 4 | 2 | 1 | 3 | 2 | 3 | 3 | 3 | 3 | 2 | 1 | 3 | 2 | 3 | 3 | 2 | 3 | 1 | 2 | 5 | 4 | 3 | 50 | 2 | 2 | 3 | 3 |
| PLCD4 | chr2:218633386-218633481 | 1 | 2 | 2 | 1 | 2 | 1 | 1 | 2 | 1 | 3 | 2 | 1 | 1 | 0 | 1 | 1 | 1 | 0 | 1 | 5 | 2 | 0 | 0 | 2 | 2 | 2 | 1 | 2 | 0 | 2 | 1 | 3 | 4 | 1 | 1 | 1 | 2 | 2 | 3 | 1 | 2 | 2 | 2 | 2 | 1 | 2 | 46 | 1 | 1 | 2 | 1 |
| BNIP1 | chr5:173162073-173162145 | 0 | 1 | 0 | 1 | 0 | 0 | 0 | 0 | 0 | 0 | 0 | 0 | 0 | 0 | 0 | 0 | 0 | 0 | 0 | 0 | 0 | 0 | 0 | 0 | 0 | 0 | 0 | 0 | 2 | 1 | 1 | 0 | 1 | 1 | 0 | 0 | 0 | 0 | 1 | 0 | 0 | 1 | 0 | 0 | 1 | 0 | 21 | 0 | 0 | 0 | 0 |
| CUBN | chr10:16875534-16875597 | 1 | 1 | 1 | 1 | 0 | 1 | 0 | 1 | 0 | 2 | 0 | 0 | 0 | 0 | 0 | 0 | 0 | 0 | 0 | 0 | 0 | 0 | 0 | 0 | 0 | 0 | 2 | 1 | 0 | 0 | 0 | 0 | 0 | 1 | 0 | 0 | 0 | 0 | 0 | 0 | 1 | 1 | 1 | 0 | 1 | 0 | 22 | 0 | 0 | 0 | 0 |
| ATL3 | chr11:63647977-63648113 | 1 | 1 | 1 | 1 | 1 | 1 | 2 | 2 | 1 | 1 | 0 | 3 | 3 | 1 | 0 | 1 | 1 | 0 | 2 | 0 | 0 | 0 | 2 | 1 | 1 | 1 | 1 | 1 | 1 | 1 | 1 | 2 | 1 | 1 | 1 | 3 | 2 | 2 | 2 | 1 | 2 | 1 | 1 | 1 | 1 | 1 | 33 | 2 | 1 | 1 | 0 |
| CHIC1 | chrX:73586633-73586810 | 1 | 1 | 1 | 1 | 0 | 1 | 0 | 1 | 0 | 0 | 0 | 0 | 1 | 0 | 0 | 0 | 0 | 1 | 0 | 0 | 0 | 0 | 0 | 0 | 1 | 0 | 1 | 1 | 0 | 1 | 1 | 3 | 0 | 1 | 1 | 1 | 0 | 0 | 0 | 0 | 1 | 1 | 1 | 1 | 1 | 1 | 30 | 1 | 0 | 1 | 2 |
| TRAF3IP1 | chr2:238338362-238344598 | 2 | 3 | 4 | 2 | 3 | 2 | 4 | 2 | 2 | 1 | 3 | 3 | 2 | 2 | 1 | 1 | 2 | 3 | 2 | 3 | 1 | -1 | 0 | 2 | 2 | 2 | 3 | 1 | 6 | 3 | 3 | 2 | 5 | 4 | 2 | 1 | 2 | 2 | 4 | 4 | 2 | 3 | 1 | 3 | 3 | 3 | 71 | 4 | 2 | 3 | 5 |
| CASD1 | chr7:94542953-94543098 | 1 | 1 | 1 | 1 | 0 | 1 | 0 | 1 | 0 | 0 | 0 | 0 | 1 | 0 | 0 | 0 | 0 | 0 | 0 | 1 | 0 | 1 | 2 | 1 | 1 | 0 | 1 | 1 | 1 | 1 | 0 | 0 | 1 | 1 | 1 | 1 | 1 | 2 | 2 | 1 | 1 | 1 | 1 | 1 | 1 | 1 | 25 | 1 | 1 | 0 | 1 |
| TPP2 | chr13:102667758-102667883 | 2 | 4 | 4 | 3 | 3 | 2 | 4 | 3 | 1 | 2 | 2 | 2 | 2 | 2 | 1 | 2 | 2 | 2 | 2 | 2 | 1 | 2 | 3 | 3 | 3 | 3 | 4 | 3 | 3 | 3 | 4 | 3 | 4 | 2 | 2 | 3 | 2 | 3 | 3 | 3 | 3 | 2 | 2 | 3 | 4 | 4 | 36 | 2 | 2 | 3 | 3 |
| SEH1L | chr18:12964436-12964495 | 2 | 2 | 1 | 1 | 1 | 0 | 1 | 1 | 0 | 0 | 0 | 0 | 0 | 0 | 0 | 0 | 0 | 0 | 0 | 0 | 0 | 1 | 0 | 1 | 1 | 1 | 1 | 1 | 1 | 1 | 1 | 1 | 1 | 1 | 1 | 1 | 1 | 1 | 1 | 1 | 1 | 2 | 2 | 2 | 2 | 1 | 31 | 1 | 1 | 1 | 2 |
| NAPEPLD | chr7:103124553-103124607 | 0 | 0 | 0 | 0 | 0 | 0 | 1 | 1 | 0 | 0 | 0 | 0 | 1 | 1 | 0 | 0 | 0 | 0 | 0 | 0 | 0 | 0 | 0 | 1 | 1 | 0 | 1 | 0 | 2 | 0 | 0 | 1 | 2 | 0 | 1 | 0 | 0 | 0 | 1 | 1 | 2 | 1 | 1 | 1 | 1 | 1 | 24 | 0 | 1 | 0 | 0 |
| PSME4 | chr2:53942159-53942224 | 1 | 1 | 0 | 1 | 1 | 1 | 0 | 1 | 1 | 0 | 0 | 0 | 0 | 1 | 1 | 1 | 0 | 0 | 0 | 2 | 0 | 1 | 0 | 0 | 1 | 0 | 1 | 1 | 1 | 0 | 1 | 0 | 0 | 1 | 0 | 0 | 1 | 1 | 0 | 0 | 1 | 1 | 1 | 1 | 1 | 1 | 21 | 1 | 0 | 1 | 2 |
| PCDHAC2 | chr5:140984066-140984149\|140984249 | 2 | 1 | 1 | 1 | 1 | 1 | 0 | 2 | 0 | 0 | 1 | 1 | 1 | 1 | 0 | 0 | 0 | 0 | 1 | 1 | 1 | 0 | -1 | 0 | 1 | 0 | 1 | 1 | -1 | 1 | 0 | 0 | 1 | 2 | 1 | 1 | 1 | 1 | 1 | 1 | 0 | 2 | 2 | 0 | 2 | 1 | 40 | 1 | 1 | 2 | 1 |
| COL25A1 | chr4:108823068-108824035\|108823139 | 0 | 0 | -1 | 0 | 0 | 0 | 0 | 0 | 0 | 0 | 1 | 0 | -1 | 0 | 0 | 0 | 0 | 0 | 0 | -1 | 0 | 0 | 0 | 0 | 1 | 0 | 0 | 0 | 0 | 1 | 0 | 0 | 0 | 0 | -1 | 0 | 0 | 0 | 0 | 0 | 2 | 0 | 0 | -1 | -1 | 1 | 28 | 0 | 0 | 0 | -1 |
| IFT122 | chr3:129466890-129467066 | 5 | 4 | 2 | 4 | 4 | 5 | 6 | 5 | 4 | 2 | 4 | 4 | 4 | 2 | 3 | 3 | 3 | 2 | 3 | 4 | 3 | 4 | 6 | 3 | 3 | 3 | 3 | 3 | 8 | 4 | 3 | 4 | 3 | 6 | 2 | 6 | 5 | 7 | 4 | 3 | 4 | 5 | 4 | 3 | 4 | 3 | 72 | 4 | 4 | 5 | 3 |
| RP11-697E2.6 | chr15:90262492-90262669 | 3 | 1 | 2 | 0 | 0 | 2 | 0 | 2 | 0 | 1 | 1 | 1 | 1 | 1 | 0 | 1 | 0 | 1 | 0 | 0 | 1 | 0 | 0 | 1 | 4 | 0 | 2 | 0 | 0 | 1 | 1 | 2 | 1 | 2 | 2 | 2 | 0 | 0 | 1 | 1 | 1 | 2 | 2 | 2 | 0 | 1 | 51 | 1 | 0 | 2 | 2 |
| SLC5A1 | chr22:32099990\|32099987-32100203 | 1 | 0 | 0 | 0 | 0 | 0 | 0 | 1 | 0 | 0 | 0 | 0 | 0 | -1 | 0 | -1 | -1 | 0 | 0 | 0 | 0 | 3 | -1 | 0 | 0 | 0 | 0 | 1 | 0 | 0 | 0 | 1 | 2 | 0 | 0 | 0 | 0 | 0 | 0 | 1 | 2 | 1 | 0 | 0 | 0 | 1 | 39 | 3 | 0 | 0 | 0 |
| ELMO2 | chr20:46382206-46382241 | 0 | 0 | 0 | 0 | 0 | 0 | 0 | 0 | 3 | 2 | 1 | 2 | 1 | 2 | 1 | 5 | 4 | 1 | 2 | 30 | 11 | 0 | 0 | 0 | 0 | 0 | 0 | 0 | 0 | 0 | 0 | 0 | 0 | 0 | 0 | 0 | 0 | 0 | 0 | 0 | 0 | 0 | 0 | 0 | 0 | 0 | 9 | 0 | 0 | 0 | 0 |
| LINC01490 | chr12:80779901-80780456 | 0 | 1 | 0 | 0 | 0 | 0 | 0 | 0 | -1 | 0 | 0 | -1 | 0 | 0 | 0 | 0 | -1 | 0 | 0 | 0 | 0 | -1 | 0 | 0 | 0 | 0 | 3 | 0 | 0 | 0 | 2 | -1 | 4 | 0 | 0 | -1 | 0 | 0 | 0 | 0 | 0 | 2 | 0 | 0 | 0 | 0 | 47 | 0 | 0 | 0 | 0 |
| TNNI3K | chr1:74365972-74366091 | 2 | 1 | 0 | 1 | 2 | 2 | 1 | 1 | 1 | 0 | 0 | 1 | 1 | 1 | 1 | 2 | 1 | 0 | 0 | 0 | 1 | 3 | 3 | 1 | 1 | 1 | 1 | 1 | 1 | 1 | 1 | 1 | 6 | 2 | 1 | 2 | 2 | 1 | 1 | 1 | 1 | 2 | 2 | 1 | 1 | 1 | 49 | 1 | 1 | 3 | 1 |
| YPEL3 | chr16:30094385-30094475 | 2 | 1 | 1 | 1 | 1 | 1 | 2 | 2 | 0 | 0 | 0 | 1 | 1 | 0 | 0 | 1 | 0 | 0 | 1 | 1 | 1 | 1 | 1 | 2 | 2 | 2 | 1 | 2 | 1 | 2 | 2 | 1 | 1 | 1 | 2 | 1 | 1 | 1 | 1 | 1 | 2 | 2 | 2 | 2 | 1 | 2 | 20 | 1 | 1 | 1 | 1 |
| VEPH1 | chr3:157468891-157468991 | 0 | 1 | 0 | 0 | 1 | 2 | 0 | 0 | -1 | 0 | 3 | 0 | 0 | -1 | 2 | 0 | 0 | 1 | 0 | 0 | 0 | 0 | 0 | 0 | 1 | 0 | 2 | 0 | 0 | 0 | 1 | 0 | 2 | 1 | 0 | 0 | 0 | 0 | 0 | 0 | 0 | 0 | 0 | 0 | 0 | 0 | 30 | 0 | -1 | 0 | 0 |
| SLC44A5 | chr1:75311615-75311705 | 2 | 0 | 0 | 0 | 0 | 0 | 0 | 0 | -1 | 0 | 0 | 0 | 0 | 0 | 1 | 0 | 2 | 0 | 0 | 0 | 0 | 0 | -1 | 0 | 1 | 0 | 0 | 1 | 0 | 0 | 0 | 0 | 0 | 0 | 0 | 0 | 0 | 0 | 0 | 0 | 0 | 0 | 0 | 0 | 0 | 0 | 20 | 0 | 0 | 0 | 0 |
| MYRFL | chr12:69925889-69925948 | 0 | 0 | 1 | 0 | 0 | -1 | -1 | 1 | 0 | -1 | 0 | 0 | 0 | 0 | 0 | 0 | 0 | 0 | 0 | 0 | 0 | -1 | 0 | 0 | 0 | 0 | 0 | 0 | 0 | 0 | 0 | 1 | 0 | 0 | 0 | -1 | 0 | 0 | 0 | 0 | 0 | 0 | 0 | 0 | 0 | 0 | 23 | 0 | 0 | 0 | -1 |
| MAGI2 | chr7:78337726-78337825 | 1 | 1 | 1 | 0 | 1 | 1 | 2 | 0 | 0 | 0 | 1 | 0 | 0 | 1 | 0 | 0 | 0 | 1 | 0 | 0 | 0 | -1 | 0 | 1 | 0 | 0 | 1 | 1 | 0 | 0 | 0 | 0 | 0 | 1 | 0 | 0 | 0 | 1 | 1 | 0 | 1 | 0 | 0 | 1 | 1 | 0 | 24 | 1 | 0 | 0 | 0 |
| ATOH8 | chr2:85772862\|85780373\|85780965-85784660 | 3 | 3 | 2 | 2 | 2 | 2 | 3 | 3 | 3 | 4 | 4 | 3 | 3 | 4 | 4 | 4 | 4 | 4 | 3 | 4 | 4 | 2 | 3 | 3 | 2 | 2 | 3 | 2 | 1 | 3 | 3 | 2 | 3 | 2 | 3 | 4 | 2 | 3 | 3 | 4 | 4 | 2 | 2 | 3 | 2 | 2 | 32 | 3 | 3 | 2 | 3 |
| C6orf106 | chr6:34649709-34649830 | 3 | 3 | 2 | 2 | 2 | 2 | 2 | 3 | 4 | 5 | 3 | 4 | 5 | 5 | 5 | 4 | 3 | 3 | 3 | 2 | 3 | 3 | 4 | 3 | 4 | 2 | 3 | 2 | 3 | 1 | 1 | 3 | 2 | 3 | 4 | 2 | 2 | 4 | 4 | 3 | 4 | 5 | 5 | 4 | 6 | 4 | 54 | 3 | 3 | 3 | 3 |
| RP11-145E5.5 | chr9:22031620-22031680 | 0 | 2 | 2 | 0 | 0 | 0 | 0 | 2 | 0 | 0 | 0 | 0 | 0 | 0 | 0 | 0 | 0 | 0 | 0 | 0 | 0 | -1 | -1 | 0 | 0 | 0 | 0 | 0 | -1 | 0 | 0 | -1 | 0 | 0 | 0 | 0 | 1 | 0 | 0 | 0 | 0 | 0 | 0 | 0 | 0 | 0 | 32 | 0 | 0 | 0 | 0 |
| DCC | chr18:53500204-53500268 | 0 | 0 | 0 | 0 | -1 | -1 | 0 | 1 | 0 | 0 | 0 | 0 | 0 | 0 | 0 | 0 | 0 | 0 | 0 | 0 | 0 | 0 | 0 | 0 | 1 | 0 | 0 | -1 | 0 | -1 | 0 | 0 | 0 | 0 | -1 | -1 | 0 | 0 | 0 | 0 | 0 | 0 | 0 | 0 | 0 | 0 | 24 | -1 | 0 | -1 | 0 |
| SYNJ1 | chr21:32691285-32691414 | 1 | 1 | 1 | 1 | 1 | 1 | 1 | 1 | 1 | 0 | 1 | 1 | 2 | 0 | 0 | 1 | 0 | 0 | 1 | 1 | 1 | 0 | 0 | 1 | 1 | 1 | 1 | 1 | -1 | 1 | 1 | 1 | 3 | 1 | 2 | 1 | 2 | 2 | 1 | 1 | 1 | 2 | 1 | 1 | 1 | 2 | 33 | 1 | 1 | 1 | 2 |
| PARD3B | chr2:205476127-205476283 | 1 | 1 | 1 | 1 | 1 | 0 | 0 | 1 | 1 | 1 | 0 | 0 | 1 | 0 | 1 | 1 | 1 | 0 | 0 | 1 | 1 | 2 | 0 | 0 | 1 | 0 | 1 | 0 | 0 | 2 | 1 | 1 | 1 | 1 | 1 | 1 | 1 | 1 | 1 | 1 | 0 | 1 | 1 | 2 | 1 | 1 | 30 | 1 | 1 | 1 | 4 |
| NOXA1 | chr9:137428441-137428616 | 1 | 1 | 0 | 1 | 1 | 1 | 0 | 1 | 2 | 2 | 2 | 1 | 1 | 2 | 1 | 2 | 2 | 2 | 2 | 3 | 1 | 1 | 1 | 1 | 1 | 1 | 1 | 1 | 3 | 2 | 1 | 1 | 1 | 1 | 1 | 1 | 4 | 2 | 1 | 1 | 1 | 1 | 2 | 1 | 2 | 1 | 29 | 1 | 1 | 1 | 2 |
| RP11-564D11.3 | chr10:122883725-122883931 | 0 | 0 | 0 | 0 | 0 | 0 | 0 | 0 | 0 | 0 | 0 | 0 | 0 | 1 | 0 | 0 | 2 | 0 | 1 | 1 | 0 | -1 | 0 | 0 | 0 | 0 | 0 | 0 | 0 | 0 | 0 | 0 | 0 | 1 | -1 | -1 | 0 | -1 | 0 | 0 | 0 | 0 | 0 | 0 | 0 | 0 | 25 | 2 | 0 | -1 | 0 |

SFig5, pg11

| Gene Symbol | Exon Location (hg38, human) | Adipose - Subcutaneous | Adipose - Visceral (Omentum) | Adrenal Gland | Artery - Aorta | Artery - Coronary | Artery - Tibial | Bladder | Breast - Mammary Tissue | Brain - Amygdala | Brain - Anterior cingulate cortex (BA24) | Brain - Caudate (basal ganglia) | Brain - Cerebellar Hemisphere | Brain - Cerebellum | Brain - Cortex | Brain - Frontal Cortex (BA9) | Brain - Hippocampus | Brain - Hypothalamus | Brain - Nucleus accumbens (basal ganglia) | Brain - Putamen (basal ganglia) | Brain - Spinal cord (cervical c-1) | Brain - Substantia nigra | Cervix - Ectocervix | Cervix - Endocervix | Colon - Sigmoid | Colon - Transverse | Esophagus - Gastroesophageal Junction | Esophagus - Mucosa | Esophagus - Muscularis | Fallopian Tube | Heart - Atrial Appendage | Heart - Left Ventricle | Kidney - Cortex | Liver | Lung | Minor Salivary Gland | Muscle - Skeletal | Nerve - Tibial | Ovary | Pancreas | Pituitary | Prostate | Skin - Not Sun Exposed (Suprapubic) | Skin - Sun Exposed (Lower leg) | Small Intestine - Terminal Ileum | Spleen | Stomach | Testis | Thyroid | Uterus | Vagina | Whole Blood |
|---|---|---|---|---|---|---|---|---|---|---|---|---|---|---|---|---|---|---|---|---|---|---|---|---|---|---|---|---|---|---|---|---|---|---|---|---|---|---|---|---|---|---|---|---|---|---|---|---|---|---|---|---|
| RP11-696N14.1 | chr4:99291326-99292444 | 1 | 1 | 1 | 2 | 1 | 1 | 1 | 1 | 0 | 1 | 0 | 0 | 0 | 3 | 0 | 2 | 0 | 2 | 1 | 0 | 0 | 0 | 0 | 0 | 0 | 0 | 1 | 1 | 0 | 0 | 0 | 1 | 0 | 1 | 0 | 1 | 1 | 0 | 1 | 0 | 0 | 1 | 1 | 1 | 1 | 1 | 21 | 1 | 1 | 0 | 0 |
| CPA2 | chr7:130273706-130273811 | 1 | 2 | 2 | 1 | 2 | 2 | 2 | 2 | 24 | 15 | 16 | 7 | 3 | 14 | 12 | 19 | 27 | 23 | 14 | 61 | 40 | 0 | -1 | 1 | 2 | 1 | 1 | 1 | 0 | 1 | 2 | 2 | 1 | 2 | 1 | 2 | 1 | 3 | 1 | 6 | 1 | 1 | 1 | 1 | 1 | 1 | 55 | 2 | 4 | 1 | 2 |
| CHD7 | chr8:60809333-60809384 | 1 | 1 | 1 | 1 | 0 | 0 | 0 | 1 | 0 | 1 | 1 | 1 | 1 | 0 | 2 | 1 | 0 | 2 | 1 | 1 | 0 | 0 | -1 | 1 | 1 | 1 | 1 | 1 | -1 | 0 | 1 | 1 | 3 | 1 | 1 | 0 | 1 | 0 | 2 | 0 | 0 | 0 | 1 | 2 | 1 | 1 | 34 | 1 | 0 | 1 | 1 |
| DNAH9 | chr17:11940618-11940662 | 2 | 0 | 0 | -1 | 0 | 0 | 0 | 1 | 0 | 0 | 0 | 0 | 1 | 0 | 0 | 0 | 0 | 0 | 1 | 1 | 0 | 0 | 0 | 0 | 0 | 0 | 0 | 0 | 0 | 0 | 0 | 0 | -1 | 0 | 0 | 0 | 0 | 1 | 1 | 0 | 0 | 0 | 0 | 0 | 0 | 0 | 21 | 0 | 0 | 0 | 0 |
| MAATS1 | chr3:119752301-119752376 | 3 | 2 | 1 | 2 | 2 | 2 | 0 | 2 | 2 | 1 | 2 | 2 | 1 | 2 | 2 | 2 | 2 | 1 | 2 | 2 | 1 | 2 | 0 | 2 | 2 | 2 | 1 | 2 | 3 | 3 | 2 | 2 | 1 | 3 | 3 | 5 | 3 | 3 | 2 | 2 | 1 | 3 | 2 | 3 | 2 | 2 | 39 | 2 | 2 | 2 | 2 |
| SNX13 | chr7:17834998-17835141 | 0 | 0 | 0 | 0 | 0 | 0 | 0 | 0 | 0 | 0 | 0 | 1 | 0 | 0 | 0 | 0 | 0 | 0 | 0 | 0 | 0 | 0 | 0 | 0 | 0 | 0 | 0 | 0 | 4 | 0 | 0 | 0 | 0 | 0 | 0 | 0 | 0 | 0 | 0 | 0 | 0 | 0 | 0 | 0 | 0 | 0 | 29 | 0 | 0 | 0 | 0 |
| KIF5C | chr2:149020176-149020382 | 2 | 1 | 0 | 0 | 1 | 0 | 1 | 0 | 0 | 0 | 0 | 0 | 0 | 0 | 0 | 0 | 0 | 0 | 0 | 0 | 0 | -1 | 0 | 1 | 0 | 0 | 1 | 0 | 0 | 0 | 0 | 1 | 0 | 1 | 0 | 0 | 0 | 0 | 0 | 0 | 0 | 0 | 0 | 1 | 0 | 0 | 22 | 0 | 0 | 0 | 2 |
| MYO19 | chr17:36530976-36531869\|36531067 | 6 | 4 | 5 | 6 | 5 | 5 | 6 | 5 | 3 | 3 | 6 | 6 | 5 | 4 | 4 | 5 | 4 | 3 | 4 | 6 | 4 | 3 | 3 | 5 | 4 | 5 | 5 | 5 | 2 | 5 | 5 | 4 | 5 | 5 | 5 | 4 | 5 | 4 | 4 | 4 | 5 | 4 | 5 | 4 | 5 | 4 | 38 | 5 | 4 | 4 | 4 |
| POU2F1 | chr1:167376553-167376587 | 1 | 1 | 0 | 1 | 0 | 1 | 0 | 1 | 0 | 0 | 0 | 0 | 0 | 0 | 0 | 0 | 0 | 0 | 0 | 0 | 0 | 0 | 0 | 0 | 0 | 0 | 0 | 0 | 0 | 0 | 1 | 1 | 0 | 1 | 0 | 1 | 1 | 1 | 0 | 1 | 0 | 2 | 2 | 1 | 0 | 1 | 23 | 0 | 0 | 0 | 1 |
| TTC28 | chr22:28017711-28029116 | 1 | 1 | 0 | 1 | 1 | 1 | 1 | 0 | 1 | 1 | 0 | 0 | 0 | 0 | 0 | 0 | 1 | 0 | 0 | 0 | 0 | 0 | 2 | 1 | 1 | 1 | 1 | 0 | 1 | 0 | 0 | 0 | 1 | 0 | 0 | 0 | 1 | 1 | 0 | 1 | 1 | 1 | 1 | 0 | 0 | 1 | 20 | 0 | 1 | 1 | 0 |
| PAK1 | chr11:77356507-77357707\|77356561 | 3 | 2 | 1 | 1 | 1 | 1 | 1 | 2 | 0 | 0 | 1 | 0 | 0 | 0 | 0 | 0 | 0 | 0 | 0 | 1 | 0 | 1 | 1 | 1 | 2 | 1 | 1 | 1 | 3 | 1 | 2 | 2 | 3 | 2 | 1 | 1 | 5 | 2 | 2 | 1 | 2 | 2 | 2 | 3 | 4 | 2 | 42 | 1 | 1 | 1 | 1 |
| NHLRC2 | chr10:113907226-113907359 | 2 | 1 | 0 | 1 | 1 | 1 | 2 | 1 | 0 | 0 | 0 | 1 | 1 | 1 | 0 | 1 | 1 | 1 | 0 | 0 | 1 | 0 | 1 | 1 | 1 | 1 | 0 | 1 | 5 | 1 | 0 | 1 | 0 | 1 | 2 | 0 | 1 | 1 | 1 | 1 | 1 | 1 | 1 | 2 | 1 | 1 | 31 | 1 | 1 | 1 | 1 |
| ANO3 | chr11:26643887-26644051 | 1 | 2 | 0 | 1 | 0 | 1 | 0 | 1 | 0 | 0 | 0 | 0 | 0 | 0 | 0 | 0 | 0 | 0 | 0 | -1 | 0 | 0 | 0 | 0 | 0 | 0 | 0 | 0 | 0 | 1 | -1 | 0 | 1 | 0 | 0 | 0 | 1 | 1 | 1 | 1 | 0 | 1 | 0 | 0 | 0 | 0 | 23 | 1 | 1 | 0 | 0 |
| ZDHHC20 | chr13:21404225-21404342 | 4 | 2 | 0 | 0 | 0 | 1 | 0 | 5 | 2 | 1 | 3 | 1 | 1 | 3 | 1 | 2 | 1 | 0 | 4 | 3 | 3 | 0 | 0 | 1 | 1 | 1 | 1 | 1 | 0 | 2 | 2 | 1 | 1 | 4 | 1 | 3 | 3 | 2 | 2 | 3 | 2 | 4 | 5 | 5 | 2 | 2 | 65 | 2 | 1 | 1 | 5 |
| CFTR | chr7:117642868-117642983 | 1 | 0 | 0 | -1 | -1 | 0 | 0 | 1 | 0 | 0 | 1 | 0 | -1 | 3 | 0 | 1 | 0 | 4 | 2 | 0 | 0 | 0 | 0 | 0 | 0 | 0 | 0 | -1 | 0 | -1 | 2 | 1 | 0 | 0 | 0 | 0 | 0 | 0 | 1 | 0 | 2 | 0 | 0 | 1 | 0 | 1 | 29 | -1 | 0 | 0 | 0 |
| DOCK8 | chr9:365591-366552 | 0 | 1 | 0 | 1 | 1 | 1 | 0 | 0 | 1 | 1 | 1 | 0 | 0 | 1 | 1 | 1 | 1 | 1 | 0 | 0 | 1 | 0 | 0 | 1 | 1 | 1 | 1 | 0 | -1 | 2 | 1 | 1 | 4 | 2 | 1 | 2 | 1 | 0 | 0 | 1 | 3 | 1 | 0 | 1 | 1 | 1 | 44 | 1 | 1 | 0 | 5 |
| AKAP1 | chr17:57107946\|57107704-57108048 | 2 | 2 | 2 | 3 | 3 | 3 | 3 | 3 | 3 | 2 | 3 | 4 | 5 | 2 | 2 | 2 | 3 | 3 | 2 | 2 | 2 | 4 | 2 | 3 | 3 | 3 | 2 | 3 | 2 | 2 | 2 | 3 | 3 | 3 | 3 | 2 | 3 | 4 | 3 | 6 | 5 | 2 | 2 | 4 | 4 | 3 | 41 | 4 | 3 | 3 | 4 |
| TMEM219 | chr16:29972042-29972140 | 1 | 2 | 2 | 2 | 2 | 2 | 2 | 2 | 1 | 1 | 1 | 3 | 3 | 2 | 2 | 1 | 1 | 1 | 1 | 1 | 1 | 2 | 1 | 1 | 3 | 1 | 2 | 1 | 2 | 1 | 1 | 1 | 2 | 3 | 2 | 2 | 1 | 1 | 2 | 1 | 2 | 2 | 1 | 3 | 3 | 2 | 26 | 1 | 1 | 2 | 2 |
| INPP4B | chr4:142425207-142425263 | 2 | 2 | 0 | 1 | 1 | 1 | 0 | 3 | 2 | 1 | 1 | 0 | 1 | 0 | 0 | 0 | 1 | 0 | 0 | 2 | 0 | 0 | 0 | 1 | 1 | 1 | 1 | 1 | 0 | 0 | 1 | -1 | -1 | 1 | 2 | 3 | 4 | 1 | 0 | 1 | 2 | 1 | 3 | 1 | 2 | 0 | 47 | 3 | 1 | 1 | 1 |
| CD24 | chr6:106971942-106974168 | 1 | 1 | 1 | 1 | 0 | 0 | 1 | 1 | 2 | 2 | 1 | 0 | -1 | 4 | 1 | 3 | 0 | 1 | -1 | 3 | 5 | 1 | 2 | 0 | 0 | 0 | 0 | 1 | -1 | 0 | 1 | 1 | 2 | 1 | 1 | 1 | 4 | 0 | 0 | 2 | 2 | 1 | 1 | 1 | 2 | 1 | 37 | 1 | 1 | 0 | 1 |
| MAGI2 | chr7:78198140-78198235 | 0 | 0 | 1 | 0 | 0 | 0 | 0 | 0 | 0 | 0 | 0 | 0 | 0 | 0 | 0 | 0 | 0 | 0 | 0 | 0 | 0 | 0 | -1 | 0 | 0 | 0 | 1 | 0 | 0 | 1 | 1 | 1 | 0 | 1 | 2 | 0 | 1 | 0 | 2 | 1 | 1 | 2 | 1 | 0 | 0 | 1 | 27 | 1 | 0 | 2 | 0 |
| SORCS3 | chr10:105258774-105258881 | 0 | 0 | 1 | 0 | 1 | 0 | 0 | -1 | 1 | 0 | 0 | 0 | 0 | 0 | 0 | 0 | 0 | 0 | 0 | 0 | 0 | 0 | 0 | 0 | 0 | 1 | 0 | 0 | 0 | 0 | 0 | -1 | 0 | 0 | 0 | -1 | 1 | 1 | 0 | 1 | 0 | 3 | 1 | 2 | -1 | 0 | 30 | 0 | 0 | -1 | 0 |
| VAV3 | chr1:107777608-107777770 | 1 | 0 | 0 | 0 | 0 | 0 | 0 | 0 | 1 | 0 | 2 | 0 | 0 | 0 | 0 | 0 | 0 | 0 | 0 | 0 | 0 | 0 | -1 | 0 | 0 | 0 | 1 | 1 | 0 | 0 | 0 | -1 | 0 | 1 | 2 | 0 | 0 | 0 | 0 | 1 | 0 | 1 | 1 | 1 | 0 | 0 | 23 | 0 | 1 | 0 | 0 |
| FFAR4 | chr10:93579018-93579206 | 0 | 0 | 0 | 1 | 0 | 1 | 0 | 0 | 0 | 0 | 1 | 0 | 0 | 0 | 0 | 0 | 0 | 1 | -1 | 0 | 0 | 0 | 0 | 0 | 0 | 1 | 2 | 0 | 0 | 0 | -1 | 0 | 0 | 1 | 0 | 0 | 0 | 0 | 0 | 0 | 0 | 1 | 1 | 1 | 0 | 0 | 21 | 1 | 0 | 0 | 0 |
| SEPT10 | chr2:109545385\|109545460\|109544955-109545542 | 2 | 4 | 3 | 3 | 4 | 2 | 4 | 3 | 1 | 3 | 3 | 1 | 10 | 1 | 2 | 2 | 2 | 1 | 0 | 2 | 2 | 4 | 5 | 5 | 5 | 5 | 2 | 5 | 10 | 5 | 5 | 3 | 4 | 2 | 2 | 8 | 2 | 2 | 2 | 2 | 3 | 2 | 1 | 5 | 3 | 4 | 83 | 2 | 5 | 3 | 5 |
| TTC39B | chr9:15194545-15194735 | 1 | 1 | 0 | 0 | 1 | 0 | -1 | 1 | 0 | 0 | 0 | 1 | 2 | 0 | 0 | 1 | 0 | 0 | 0 | 3 | 2 | 0 | 0 | 1 | 2 | 1 | 1 | 0 | 0 | 1 | 1 | 1 | 2 | 1 | 0 | 0 | 1 | 0 | 1 | 1 | 2 | 0 | 0 | 2 | 2 | 1 | 25 | 1 | 1 | 1 | 2 |
| MTRR | chr5:7883724-7883863 | 2 | 1 | 0 | 1 | 1 | 1 | 2 | 2 | 1 | 2 | 1 | 2 | 3 | 1 | 2 | 1 | 1 | 2 | 1 | 1 | 1 | 3 | 0 | 1 | 1 | 1 | 1 | 1 | 0 | 2 | 1 | 0 | 0 | 2 | 1 | 2 | 2 | 2 | 1 | 1 | 1 | 2 | 2 | 1 | 1 | 1 | 27 | 1 | 1 | 1 | 1 |
| HERC4 | chr10:68038431\|68039417-68039488 | 1 | 2 | 1 | 1 | 1 | 1 | 1 | 1 | 1 | 0 | 0 | 1 | 1 | 0 | 1 | 0 | 1 | 0 | 1 | 0 | 1 | 2 | 0 | 1 | 1 | 1 | 1 | 1 | 2 | 1 | 1 | 1 | 4 | 2 | 1 | 0 | 2 | 1 | 2 | 1 | 1 | 1 | 1 | 2 | 3 | 1 | 29 | 1 | 1 | 1 | 1 |
| DGKI | chr7:137594673-137594764 | 0 | 0 | 0 | 1 | 0 | 2 | 0 | 2 | 1 | 0 | 1 | 0 | 1 | 1 | 1 | 0 | 0 | 1 | 1 | 0 | 0 | 0 | -1 | 0 | 0 | 0 | 0 | 0 | -1 | 6 | 0 | 0 | 0 | 1 | 1 | 0 | 1 | 0 | 0 | 1 | -1 | 4 | 3 | 0 | -1 | -1 | 50 | 1 | 0 | 0 | 0 |
| SCAMP2 | chr15:74853368-74853496 | 0 | 0 | 0 | 0 | 0 | 0 | 0 | 0 | 0 | 0 | 0 | 1 | 0 | 0 | 0 | 0 | 0 | 0 | 0 | 0 | 0 | 0 | 0 | 0 | 0 | 0 | 0 | 0 | 0 | 1 | 2 | 1 | 1 | 0 | 0 | 7 | 0 | 0 | 0 | 0 | 0 | 0 | 0 | 1 | 0 | 0 | 43 | 0 | 0 | 0 | 0 |
| IGBP1 | chrX:70162346\|70162285-70162464 | 10 | 8 | 8 | 8 | 8 | 9 | 11 | 10 | 7 | 6 | 6 | 8 | 8 | 6 | 6 | 6 | 6 | 6 | 6 | 8 | 7 | 10 | 8 | 11 | 9 | 9 | 7 | 10 | 11 | 9 | 7 | 7 | 10 | 9 | 9 | 7 | 10 | 7 | 12 | 7 | 11 | 9 | 9 | 10 | 10 | 11 | 64 | 8 | 8 | 8 | 5 |
| FAM92A1 | chr8:93727131-93727244 | 1 | 1 | 1 | 1 | 1 | 1 | 1 | 1 | 1 | 1 | 1 | 1 | 1 | 1 | 1 | 0 | 1 | 1 | 0 | 1 | 1 | 0 | 1 | 1 | 1 | 1 | 1 | 1 | 5 | 0 | 0 | 2 | 2 | 1 | 1 | 1 | 2 | 1 | 1 | 1 | 1 | 2 | 1 | 1 | 1 | 1 | 27 | 1 | 1 | 1 | 2 |
| CHP1 | chr15:41264165-41264235 | 5 | 4 | 2 | 4 | 4 | 5 | 4 | 5 | 2 | 1 | 2 | 3 | 3 | 2 | 2 | 2 | 2 | 2 | 3 | 3 | 2 | 2 | 3 | 4 | 4 | 4 | 3 | 4 | 3 | 3 | 3 | 3 | 6 | 4 | 4 | 7 | 6 | 5 | 7 | 4 | 6 | 6 | 5 | 7 | 5 | 5 | 49 | 4 | 4 | 4 | 3 |
| ARHGAP27 | chr17:45401580-45401668 | 1 | 1 | 1 | 1 | 1 | 1 | 0 | 1 | 1 | 1 | 1 | 3 | 4 | 1 | 0 | 1 | 1 | 1 | 1 | 1 | 0 | 1 | 1 | 1 | 1 | 1 | 1 | 1 | 1 | 0 | 1 | 1 | 1 | 0 | 1 | 1 | 1 | 1 | 1 | 1 | 1 | 1 | 1 | 1 | 1 | 1 | 21 | 1 | 1 | 1 | 1 |
| SLC35A1 | chr6:87506881-87507061 | 1 | 1 | 1 | 1 | 1 | 1 | 2 | 1 | 2 | 2 | 1 | 6 | 6 | 2 | 1 | 1 | 1 | 1 | 2 | 1 | 1 | 1 | 0 | 1 | 1 | 1 | 1 | 1 | 1 | 1 | 1 | 1 | 1 | 1 | 1 | 1 | 1 | 1 | 1 | 1 | 1 | 2 | 2 | 2 | 2 | 1 | 30 | 1 | 1 | 1 | 1 |
| FAP | chr2:162211436-162211492 | 1 | 2 | 3 | 1 | 1 | 1 | -1 | 2 | 48 | 38 | 30 | -1 | -1 | 54 | 49 | -1 | 31 | 39 | -1 | -1 | 22 | 3 | 2 | 2 | 2 | 4 | 1 | 4 | 2 | 2 | 2 | 0 | 1 | 2 | 1 | 2 | 2 | 2 | 4 | 10 | 3 | 2 | 1 | 2 | 2 | 4 | 64 | 1 | 2 | 2 | 1 |
| SFXN5 | chr2:72956980-72957103 | 3 | 4 | 3 | 3 | 3 | 3 | 2 | 3 | 5 | 5 | 5 | 5 | 4 | 5 | 5 | 5 | 4 | 6 | 5 | 4 | 4 | 9 | 4 | 3 | 3 | 2 | 3 | 3 | 4 | 3 | 3 | 4 | 5 | 3 | 3 | 3 | 4 | 4 | 4 | 3 | 3 | 4 | 4 | 3 | 3 | 3 | 42 | 4 | 4 | 3 | 3 |
| UTP18 | chr17:51286483-51286552 | 2 | 2 | 1 | 2 | 2 | 2 | 2 | 2 | 2 | 3 | 2 | 4 | 5 | 3 | 3 | 2 | 2 | 2 | 2 | 3 | 3 | 2 | 2 | 2 | 2 | 2 | 2 | 2 | 3 | 2 | 2 | 3 | 4 | 2 | 2 | 2 | 3 | 1 | 3 | 1 | 3 | 2 | 2 | 2 | 2 | 2 | 26 | 2 | 2 | 2 | 2 |
| FAM200B | chr4:15684850-15684959 | 5 | 3 | 3 | 1 | 2 | 1 | 2 | 4 | 1 | 1 | 1 | 1 | 1 | 1 | 1 | 1 | 1 | 1 | 1 | 1 | 1 | 3 | 1 | 2 | 2 | 1 | 3 | 1 | 1 | 3 | 3 | 3 | 5 | 3 | 3 | 2 | 2 | 2 | 6 | 1 | 2 | 3 | 3 | 3 | 3 | 4 | 49 | 2 | 1 | 2 | 6 |
| NTRK3 | chr15:87931026-87931236 | 0 | 1 | 1 | 0 | 0 | 0 | 0 | 0 | 1 | 0 | 0 | 1 | 1 | 0 | 0 | 0 | 0 | 0 | 0 | 3 | 0 | -1 | 0 | 0 | 0 | 1 | 0 | 0 | 0 | 2 | 1 | 0 | -1 | 1 | 1 | 0 | 0 | 0 | -1 | 3 | 1 | 0 | 0 | 0 | 0 | 0 | 25 | 1 | 0 | 1 | 0 |
| RP11-755F10.3 | chr11:66261264-66261416\|66261355 | 2 | 2 | 1 | 1 | 1 | 2 | 1 | 2 | 1 | 3 | 2 | 3 | 4 | 2 | 2 | 2 | 1 | 1 | 1 | 1 | 4 | 1 | 0 | 1 | 1 | 1 | 2 | 1 | 2 | 2 | 3 | 1 | 1 | 2 | 1 | 1 | 3 | 4 | 2 | 3 | 3 | 2 | 2 | 3 | 2 | 2 | 32 | 2 | 1 | 1 | 3 |
| CES1 | chr16:55812336-55812430 | 0 | 1 | 1 | 1 | 1 | 1 | 2 | 1 | 0 | 1 | 1 | 0 | 1 | 1 | 0 | 2 | 0 | 1 | 1 | 1 | 0 | 2 | 4 | 2 | 2 | 1 | 1 | 1 | 1 | 0 | 0 | 1 | 1 | 1 | 1 | 1 | 1 | 1 | 1 | 1 | 2 | 1 | 1 | 1 | 2 | 2 | 22 | 1 | 1 | 2 | 1 |
| CLPB | chr11:72372925-72373014 | 4 | 2 | 2 | 2 | 2 | 2 | 2 | 4 | 2 | 2 | 2 | 1 | 2 | 3 | 1 | 2 | 1 | 2 | 2 | 2 | 2 | 0 | 3 | 2 | 2 | 2 | 2 | 1 | 1 | 2 | 1 | 3 | 2 | 2 | 4 | 2 | 3 | 2 | 4 | 2 | 3 | 4 | 4 | 3 | 3 | 3 | 37 | 4 | 1 | 2 | 3 |
| PC | chr11:66855027-66860669 | 1 | 4 | 1 | 1 | 2 | 0 | 1 | 3 | 0 | 0 | 0 | 0 | 0 | 0 | 0 | 1 | 1 | 0 | 0 | 0 | 0 | 1 | 0 | 1 | 1 | 3 | 1 | 1 | 3 | 1 | 1 | 1 | 0 | 1 | 1 | 0 | 1 | 0 | 1 | 1 | 0 | 1 | 0 | 2 | 1 | 1 | 28 | 1 | 0 | 1 | 1 |
| TRDN | chr6:123463850-123464061 | 0 | 0 | 1 | 0 | 1 | 0 | -1 | 0 | 1 | 0 | 1 | 0 | 1 | 1 | 0 | 0 | 1 | 1 | 0 | 0 | 0 | -1 | 0 | 0 | 1 | 1 | 0 | 1 | 0 | 1 | 4 | 0 | 2 | 0 | 0 | 0 | 0 | 0 | 0 | 0 | 0 | 1 | 0 | 0 | 1 | 0 | 27 | 0 | 1 | 1 | 0 |
| MAST4 | chr5:66870778-66870852 | 2 | 2 | 2 | 3 | 2 | 3 | 0 | 2 | 7 | 26 | 3 | 5 | -1 | 29 | 34 | 11 | 24 | 4 | 4 | -1 | 8 | 0 | 0 | 4 | 3 | 2 | 2 | 3 | 0 | 1 | 1 | 3 | 1 | 3 | 3 | 5 | 5 | 2 | 1 | 2 | 1 | 2 | 3 | 2 | 2 | 2 | 46 | 2 | 2 | 1 | 3 |
| ANKAR | chr2:189676456-189677091 | 1 | 1 | 0 | 2 | 3 | 3 | 3 | 2 | 4 | 1 | 1 | 1 | 1 | 0 | 2 | 0 | 0 | 2 | 1 | 0 | 0 | -1 | 0 | 2 | 1 | 1 | 1 | 2 | -1 | 1 | 1 | 0 | 0 | 1 | 0 | 1 | 2 | 3 | 1 | 1 | 1 | 1 | 1 | 1 | 1 | 1 | 31 | 1 | 2 | 2 | 1 |
| SAV1 | chr14:50649307-50649389 | 1 | 2 | 0 | 1 | 1 | 1 | 1 | 2 | 0 | 1 | 0 | 1 | 1 | 0 | 0 | 0 | 0 | 0 | 0 | 0 | 0 | 2 | 0 | 1 | 1 | 1 | 1 | 0 | 2 | 0 | 0 | 0 | 1 | 2 | 1 | 0 | 1 | 1 | 1 | 0 | 1 | 2 | 1 | 2 | 1 | 2 | 22 | 0 | 1 | 0 | 2 |
| HECW1 | chr7:43484144-43484310 | 0 | 0 | 0 | 1 | 0 | 0 | 0 | 0 | 0 | 0 | 0 | 1 | 1 | 0 | 0 | 0 | 0 | 0 | 0 | 1 | 0 | -1 | -1 | 0 | 0 | 0 | 0 | 0 | -1 | 0 | 0 | 0 | 0 | 2 | 0 | 0 | 0 | 0 | 0 | 0 | 1 | 0 | 4 | 0 | 0 | 0 | 27 | 0 | 0 | 0 | 0 |
| CLVS2 | chr6:123050837-123050958 | -1 | -1 | 0 | 0 | 0 | 1 | -1 | 3 | 0 | 0 | 0 | 0 | 0 | 0 | 0 | 0 | 1 | 0 | 0 | 0 | 0 | 0 | 0 | 1 | 0 | 0 | 0 | -1 | 0 | 0 | 0 | -1 | -1 | 0 | 0 | 1 | -1 | 0 | 0 | 1 | 2 | 0 | 9 | 0 | -1 | 0 | 59 | 0 | 0 | 2 | 0 |
| RP11-33B1.1 | chr4:119457971-119458170\|119458077 | 3 | 5 | 5 | 4 | 5 | 3 | 2 | 3 | 8 | 8 | 7 | 4 | 4 | 6 | 7 | 6 | 8 | 9 | 6 | 3 | 3 | 3 | 6 | 5 | 5 | 3 | 6 | 3 | 5 | 4 | 5 | 5 | 6 | 6 | 4 | 2 | 4 | 5 | 6 | 7 | 5 | 3 | 2 | 4 | 6 | 5 | 45 | 5 | 4 | 5 | 5 |
| WDR27 | chr6:169636371-169636504 | 11 | 6 | 7 | 9 | 9 | 10 | 8 | 9 | 11 | 5 | 9 | 6 | 6 | 7 | 5 | 5 | 11 | 8 | 7 | 9 | 7 | 3 | 5 | 7 | 8 | 7 | 6 | 8 | 10 | 7 | 7 | 5 | 4 | 11 | 9 | 10 | 10 | 9 | 10 | 12 | 8 | 7 | 9 | 9 | 10 | 7 | 73 | 10 | 7 | 6 | 10 |
| ZNF806 | chr2:132312585-132312746 | 1 | 3 | 0 | 0 | 0 | 0 | 0 | 0 | 4 | 3 | 2 | 4 | 1 | 6 | 4 | 7 | 3 | 2 | 3 | 1 | 5 | 0 | -1 | 1 | 0 | 0 | 2 | 1 | 0 | 0 | 0 | 0 | 0 | 0 | -1 | 0 | 2 | 0 | 0 | 0 | 0 | 0 | 0 | 0 | 0 | 0 | 24 | 0 | 0 | 0 | 0 |
| SPATA5 | chr4:123122237-123122283 | 1 | 1 | 1 | 2 | 1 | 1 | -1 | 1 | 4 | 0 | 1 | 5 | 2 | 3 | 0 | 1 | 0 | 1 | 3 | 1 | 3 | -1 | 0 | 2 | 1 | 1 | 1 | 1 | 0 | 1 | 1 | 0 | 1 | 1 | 0 | 1 | 2 | 2 | 2 | 1 | 2 | 2 | 3 | 3 | 3 | 2 | 30 | 1 | 2 | 2 | 3 |
| TPK1 | chr7:144606929-144607050 | 3 | 2 | 1 | 2 | 2 | 2 | 2 | 3 | 0 | 1 | 1 | 1 | 2 | 2 | 1 | 1 | 1 | 0 | 1 | 1 | 1 | -1 | 0 | 2 | 2 | 2 | 3 | 2 | -1 | 1 | 1 | 2 | 3 | 2 | 3 | 2 | 3 | 2 | 2 | 2 | 2 | 5 | 3 | 4 | 2 | 1 | 38 | 1 | 2 | 1 | 1 |
| SCMH1 | chr1:41162779-41162979\|41163017 | 2 | 2 | 1 | 1 | 1 | 1 | 1 | 2 | 1 | 1 | 1 | 1 | 1 | 1 | 1 | 1 | 1 | 1 | 1 | 1 | 0 | 3 | 4 | 1 | 2 | 1 | 2 | 1 | 2 | 1 | 1 | 2 | 3 | 3 | 1 | 1 | 3 | 4 | 1 | 2 | 2 | 2 | 2 | 1 | 3 | 1 | 27 | 2 | 3 | 2 | 3 |
| VRK1 | chr14:96877502-96877573 | 3 | 2 | 1 | 2 | 1 | 2 | 4 | 3 | 3 | 5 | 3 | 5 | 4 | 4 | 5 | 4 | 3 | 3 | 3 | 3 | 2 | 3 | 0 | 2 | 1 | 1 | 1 | 1 | 2 | 3 | 2 | 3 | 2 | 2 | 2 | 6 | 3 | 1 | 2 | 1 | 1 | 3 | 3 | 2 | 2 | 2 | 36 | 1 | 1 | 2 | 2 |
| SUCLA2 | chr13:47939267-47939409 | 4 | 5 | 1 | 3 | 3 | 2 | 0 | 2 | -1 | 0 | 0 | 3 | 3 | 1 | 0 | -1 | 0 | 0 | 0 | 0 | 0 | 0 | -1 | 3 | 3 | 1 | 1 | 1 | 0 | 2 | 1 | 0 | 5 | 4 | 2 | 3 | 6 | 0 | 1 | 2 | 3 | 3 | 5 | 4 | 3 | 5 | 57 | 3 | 1 | 2 | 3 |
| ZFP28 | chr19:56548378-56548478 | 1 | 1 | 1 | 1 | 2 | 1 | 1 | 1 | 1 | 0 | 2 | 1 | 1 | 1 | 1 | 1 | 1 | 1 | 1 | 1 | 2 | 2 | 4 | 2 | 2 | 1 | 2 | 1 | 5 | 1 | 1 | 3 | 1 | 1 | 2 | 0 | 2 | 3 | 2 | 1 | 2 | 1 | 1 | 2 | 3 | 2 | 29 | 1 | 2 | 2 | 2 |
| MFSD14A | chr1:100043073-100043229 | 6 | 6 | 3 | 4 | 3 | 4 | 3 | 5 | 5 | 5 | 4 | 7 | 6 | 5 | 5 | 4 | 5 | 5 | 4 | 2 | 4 | 5 | 5 | 3 | 5 | 3 | 4 | 3 | 4 | 5 | 4 | 4 | 4 | 7 | 4 | 6 | 5 | 4 | 7 | 6 | 5 | 6 | 6 | 6 | 6 | 5 | 39 | 5 | 3 | 5 | 6 |
| HMCN2 | chr9:130419208-130419587 | 2 | 1 | 1 | 1 | 0 | 1 | 1 | 1 | 0 | 0 | 5 | 1 | 0 | 0 | 0 | 3 | 1 | 3 | 0 | 0 | 0 | 0 | 0 | 0 | 0 | 0 | 0 | 0 | 1 | 1 | 0 | 1 | 0 | 1 | 0 | 1 | 0 | 0 | 0 | 4 | 1 | 2 | 2 | 0 | 1 | 1 | 23 | 0 | 0 | 0 | 1 |
| SLC6A15 | chr12:84864850-84864985 | 5 | 0 | -1 | 0 | 0 | 6 | -1 | 6 | 0 | 0 | 1 | 2 | 1 | 1 | 1 | 0 | 1 | 2 | 1 | 1 | 1 | 0 | 0 | 3 | 4 | 1 | 1 | 1 | -1 | 0 | 0 | 0 | 0 | 3 | 3 | 0 | 6 | -1 | 0 | 2 | -1 | 3 | 3 | -1 | -1 | 0 | 76 | 5 | 8 | 2 | 0 |
| TRDN | chr6:123251056-123251205 | 1 | 1 | 1 | 0 | 0 | 0 | -1 | 0 | 0 | 0 | 0 | 0 | 0 | 0 | 0 | 0 | 1 | 3 | 0 | 0 | 0 | -1 | -1 | 2 | 2 | 0 | 0 | 0 | 0 | 0 | 0 | 0 | -1 | 0 | 0 | 0 | 0 | 0 | 0 | 0 | 0 | 0 | 1 | 1 | 0 | 0 | 20 | 0 | 0 | 0 | 0 |
| ZDHHC13 | chr11:19168825-19168968 | 2 | 2 | 0 | 2 | 1 | 2 | 3 | 3 | 0 | 0 | 0 | 0 | 0 | 0 | 0 | 0 | 0 | 0 | 0 | 0 | 0 | 1 | 0 | 1 | 1 | 1 | 0 | 1 | 1 | 1 | 0 | 0 | 1 | 1 | 1 | 1 | 2 | 2 | 1 | 0 | 2 | 2 | 2 | 2 | 2 | 1 | 22 | 1 | 1 | 1 | 2 |
| IFT43 | chr14:75906555-75906677 | 3 | 5 | 4 | 4 | 5 | 4 | 5 | 4 | 4 | 5 | 5 | 5 | 6 | 6 | 5 | 5 | 5 | 5 | 4 | 3 | 3 | 7 | 4 | 5 | 6 | 4 | 6 | 4 | 4 | 4 | 4 | 6 | 4 | 5 | 4 | 3 | 3 | 5 | 5 | 7 | 6 | 3 | 2 | 4 | 5 | 5 | 39 | 5 | 4 | 6 | 4 |
| TMEM237 | chr2:201634804\|201633660-201634893 | 6 | 3 | 1 | 3 | 3 | 4 | 4 | 5 | 3 | 2 | 2 | 2 | 2 | 2 | 1 | 2 | 1 | 1 | 2 | 3 | 3 | 2 | 5 | 4 | 4 | 4 | 3 | 4 | 2 | 3 | 3 | 5 | 3 | 3 | 4 | 5 | 6 | 3 | 5 | 1 | 4 | 5 | 6 | 5 | 4 | 4 | 40 | 3 | 3 | 4 | 3 |
| CADPS2 | chr7:122617465-122617530 | 3 | 2 | 1 | 1 | 1 | 2 | 5 | 3 | 1 | 1 | 1 | 3 | 3 | 2 | 0 | 1 | 1 | 1 | 1 | 1 | 1 | 3 | 2 | 2 | 1 | 1 | 2 | 1 | 1 | 3 | 1 | 1 | 2 | 2 | 3 | 4 | 2 | 2 | 3 | 3 | 3 | 3 | 3 | 2 | 1 | 2 | 30 | 2 | 3 | 3 | 3 |
| SYNJ2 | chr6:158093888-158094037 | 0 | 1 | 0 | 0 | 0 | 0 | 1 | 0 | 6 | 17 | 2 | 5 | 5 | 16 | 24 | 5 | 3 | 2 | 1 | 1 | 1 | 2 | 0 | 1 | 0 | 1 | 2 | 2 | 0 | 6 | 2 | 0 | 1 | 0 | 0 | 0 | 0 | 0 | 0 | 1 | 1 | 0 | 0 | 0 | 0 | 1 | 37 | 1 | 1 | 1 | 0 |
| RP11-248J23.7 | chr10:95865273-95865411 | 2 | 3 | 3 | 3 | 3 | 3 | 4 | 3 | 3 | 2 | 1 | 1 | 2 | 2 | 2 | 3 | 3 | 1 | 3 | 2 | 3 | 4 | 2 | 3 | 2 | 2 | 2 | 2 | 1 | 2 | 3 | 2 | 2 | 3 | 3 | 3 | 4 | 3 | 2 | 2 | 3 | 2 | 2 | 2 | 3 | 2 | 22 | 2 | 2 | 3 | 2 |
| KIF9 | chr3:47282140-47282385 | 3 | 5 | 4 | 3 | 3 | 3 | 4 | 5 | 4 | 5 | 4 | 6 | 7 | 7 | 7 | 4 | 5 | 4 | 4 | 3 | 3 | 7 | 1 | 4 | 4 | 4 | 4 | 3 | 3 | 4 | 4 | 3 | 4 | 5 | 3 | 3 | 4 | 4 | 3 | 5 | 5 | 5 | 5 | 4 | 3 | 5 | 36 | 5 | 5 | 4 | 4 |
| GALC | chr14:87986969-87987056 | 2 | 1 | 1 | 1 | 2 | 2 | 2 | 2 | 2 | 1 | 1 | 3 | 4 | 2 | 1 | 0 | 1 | 1 | 2 | 1 | 1 | 0 | 0 | 1 | 1 | 1 | 1 | 2 | 3 | 2 | 2 | 2 | 2 | 2 | 2 | 3 | 2 | 2 | 1 | 1 | 1 | 3 | 3 | 2 | 2 | 1 | 26 | 1 | 1 | 2 | 1 |
| CTD-2341M24.1 | chr14:86104747-86104888 | 3 | 1 | 0 | 1 | 0 | 1 | 0 | 1 | 1 | 0 | 1 | 1 | 3 | 0 | 1 | 1 | 0 | 1 | 1 | 2 | 0 | 0 | 0 | 1 | 1 | 1 | 1 | 0 | 0 | 3 | 1 | 0 | 1 | 1 | 1 | 1 | 2 | 1 | 1 | 0 | 1 | 2 | 1 | 1 | 1 | 1 | 23 | 1 | 0 | 1 | 1 |
| MYRFL | chr12:69948400\|69948384\|69948364-69948488 | 0 | -1 | 4 | 0 | 2 | 2 | 0 | 1 | 0 | 0 | 0 | 0 | 1 | 0 | 3 | 1 | 0 | 0 | 0 | 0 | 0 | 0 | -1 | 0 | 1 | 0 | 1 | 0 | -1 | 3 | 0 | 2 | 0 | 1 | 3 | 0 | 1 | 0 | 2 | 2 | 2 | 0 | 0 | 0 | 1 | 0 | 42 | 5 | 0 | -1 | 4 |
| GPATCH2 | chr1:217447112\|217447108-217447176 | 2 | 2 | 2 | 1 | 1 | 1 | 2 | 2 | 1 | 1 | 1 | 2 | 0 | 1 | 0 | 1 | 1 | 1 | 1 | 1 | 2 | 1 | 5 | 2 | 1 | 2 | 1 | 2 | 1 | 4 | 1 | 1 | 2 | 3 | 2 | 1 | 1 | 2 | 2 | 1 | 2 | 2 | 1 | 3 | 2 | 2 | 25 | 2 | 2 | 2 | 3 |
| RASGRF1 | chr15:78978928-78979206\|78979202 | 2 | 5 | 7 | 0 | 2 | 2 | 0 | 2 | 5 | 2 | 5 | 1 | 1 | 1 | 1 | 9 | 13 | 9 | 7 | 73 | 45 | -1 | -1 | 8 | 4 | 3 | 2 | 3 | 0 | 0 | -1 | 5 | -1 | 2 | 4 | -1 | 3 | 9 | 2 | 6 | 5 | 6 | 3 | 2 | 4 | 3 | 75 | 5 | 6 | 2 | 1 |
| RAB36 | chr22:23155470-23155658 | 1 | 0 | 0 | 1 | 0 | 1 | 0 | 2 | 1 | 0 | 0 | 0 | 0 | 0 | 0 | 0 | 0 | 0 | 0 | 0 | 0 | 0 | 2 | 1 | 1 | 1 | 0 | 1 | 5 | 0 | 0 | 0 | 2 | 3 | 1 | 4 | 1 | 2 | 1 | 1 | 1 | 1 | 2 | 1 | 1 | 1 | 29 | 1 | 1 | 1 | 2 |
| C3orf67 | chr3:58992508-58992615 | 0 | 0 | 0 | 0 | 1 | 0 | 0 | 0 | 0 | 1 | 1 | 0 | 0 | 0 | 0 | 1 | 0 | 0 | 2 | 0 | 0 | -1 | 0 | 0 | 0 | 0 | 0 | 1 | 0 | 1 | 1 | 0 | 1 | 1 | 0 | -1 | 0 | 0 | 0 | 0 | 1 | 1 | 0 | 1 | 2 | 3 | 20 | 1 | 0 | 0 | 0 |
| BRD8 | chr5:138157642-138157894\|138157818 | 3 | 5 | 3 | 3 | 3 | 2 | 2 | 3 | 2 | 1 | 2 | 2 | 2 | 1 | 1 | 2 | 2 | 1 | 1 | 4 | 2 | 5 | 5 | 2 | 2 | 2 | 3 | 2 | 2 | 6 | 3 | 5 | 4 | 5 | 4 | 2 | 3 | 8 | 3 | 3 | 6 | 3 | 3 | 3 | 4 | 3 | 41 | 4 | 5 | 5 | 5 |
| PPFIBP2 | chr11:7548344-7548499 | 3 | 3 | 1 | 4 | 3 | 2 | 1 | 3 | 1 | 1 | 2 | 2 | 6 | 1 | 2 | 1 | 4 | 2 | 1 | 2 | 1 | 1 | 0 | 3 | 3 | 2 | 4 | 2 | -1 | 1 | 1 | 4 | 4 | 4 | 2 | 3 | 2 | 3 | 2 | 3 | 3 | 2 | 2 | 3 | 3 | 2 | 39 | 2 | 4 | 4 | 5 |
| PITPNM1 | chr11:67501103-67501224 | 3 | 2 | 1 | 1 | 1 | 1 | 5 | 2 | 0 | 0 | 1 | 2 | 2 | 0 | 0 | 1 | 0 | 0 | 1 | 1 | 0 | 0 | 2 | 2 | 2 | 1 | 1 | 1 | 1 | 1 | 1 | 2 | 2 | 2 | 2 | 1 | 2 | 2 | 3 | 2 | 2 | 2 | 2 | 3 | 3 | 2 | 24 | 3 | 2 | 2 | 2 |
| SSUH2 | chr3:8626914-8627085 | 2 | 3 | 4 | 1 | 3 | 2 | 0 | 2 | 1 | 1 | 2 | 0 | 0 | 0 | 0 | 0 | 2 | 1 | 0 | 0 | 1 | 1 | 0 | -1 | 4 | 4 | 1 | 2 | 1 | 0 | 3 | 3 | 1 | 2 | 4 | 2 | 2 | 2 | 3 | 3 | 2 | 2 | 1 | 4 | 2 | 1 | 39 | 4 | 2 | 3 | -1 |
| RASGRF2 | chr5:81183181-81183319 | 5 | 3 | 5 | 3 | 3 | 4 | -1 | 4 | 1 | 1 | 1 | 1 | 1 | 1 | 1 | 1 | 1 | 1 | 1 | 2 | 2 | 0 | 0 | 2 | 2 | 3 | 4 | 5 | -1 | 3 | 3 | 2 | 5 | 3 | 5 | 5 | 4 | 4 | 4 | 4 | 3 | 7 | 7 | 2 | 4 | 4 | 50 | 4 | 3 | 3 | 3 |
| CREM | chr10:35137782-35137879 | 5 | 7 | 5 | 5 | 6 | 6 | 6 | 5 | 2 | 3 | 3 | 4 | 3 | 3 | 3 | 3 | 3 | 2 | 3 | 3 | 2 | 4 | 3 | 5 | 7 | 5 | 5 | 5 | 5 | 4 | 4 | 6 | 4 | 5 | 5 | 2 | 3 | 4 | 6 | 7 | 5 | 4 | 3 | 7 | 8 | 7 | 42 | 7 | 5 | 5 | 4 |
| RFX1 | chr19:13978750-13979079 | 7 | 5 | 3 | 8 | 8 | 7 | 12 | 8 | 6 | 7 | 7 | 15 | 14 | 5 | 7 | 8 | 10 | 8 | 6 | 8 | 4 | 7 | 6 | 6 | 5 | 7 | 4 | 7 | 7 | 9 | 7 | 11 | 2 | 8 | 8 | 5 | 10 | 12 | 5 | 11 | 8 | 10 | 11 | 9 | 7 | 5 | 75 | 8 | 7 | 7 | 5 |
| FER | chr5:108820151\|108820350-108820524 | 3 | 3 | 2 | 2 | 1 | 3 | 1 | 4 | 1 | 3 | 2 | 2 | 1 | 2 | 3 | 2 | 2 | 2 | 3 | 2 | 3 | 4 | -1 | 2 | 2 | 2 | 3 | 2 | 2 | 3 | 1 | 3 | 3 | 4 | 4 | 2 | 3 | 2 | 3 | 4 | 3 | 4 | 5 | 2 | 1 | 2 | 32 | 3 | 2 | 3 | 2 |
| DIS3L2 | chr2:232111197-232111255 | 2 | 2 | 1 | 3 | 3 | 3 | 1 | 3 | 2 | 1 | 1 | 1 | 1 | 1 | 1 | 1 | 1 | 2 | 1 | 1 | 1 | 1 | 2 | 2 | 2 | 2 | 1 | 1 | 3 | 2 | 3 | 2 | 2 | 3 | 2 | 2 | 3 | 5 | 3 | 3 | 4 | 2 | 2 | 3 | 3 | 2 | 23 | 4 | 3 | 2 | 2 |
| PDE11A | chr2:177656532-177656675 | 1 | 2 | 2 | 2 | 0 | 1 | 0 | 1 | 6 | 5 | 3 | 1 | 0 | 2 | 1 | 2 | 4 | 4 | 4 | 3 | 3 | 0 | 0 | 1 | 1 | 0 | -1 | -1 | -1 | 8 | 4 | -1 | 1 | -1 | 1 | 3 | 3 | 0 | 0 | 3 | 1 | 2 | 0 | 1 | -1 | 0 | 46 | 0 | 0 | 0 | 0 |
| PHYKPL | chr5:178223320-178223945 | 3 | 4 | 3 | 3 | 3 | 3 | 5 | 4 | 4 | 2 | 3 | 7 | 8 | 4 | 2 | 3 | 4 | 3 | 3 | 3 | 3 | 3 | 6 | 4 | 5 | 4 | 4 | 4 | 7 | 4 | 4 | 5 | 3 | 5 | 4 | 2 | 4 | 6 | 5 | 4 | 6 | 4 | 4 | 6 | 6 | 4 | 39 | 5 | 5 | 5 | 3 |
| KLHL23 | chr2:169697893\|169697865-169697982 | 8 | 8 | 5 | 5 | 4 | 4 | 12 | 8 | 4 | 3 | 4 | 3 | 3 | 3 | 3 | 4 | 3 | 4 | 4 | 3 | 7 | 6 | 1 | 6 | 7 | 5 | 6 | 6 | 9 | 5 | 3 | 4 | 7 | 7 | 6 | 4 | 4 | 3 | 5 | 4 | 4 | 6 | 8 | 9 | 8 | 7 | 64 | 5 | 5 | 5 | 8 |
| SLCO1C1 | chr12:20714673-20714772 | 0 | 0 | 0 | 1 | 2 | 1 | 0 | 1 | 0 | 0 | 0 | 0 | 0 | 0 | 0 | 0 | 0 | 0 | 0 | 0 | 0 | -1 | 0 | 0 | 1 | 3 | 0 | 0 | 0 | 1 | 1 | 0 | 0 | 0 | 0 | 0 | 0 | 0 | 1 | 0 | 3 | 0 | 3 | 0 | 0 | 5 | 30 | 0 | 0 | -1 | 0 |
| BPIFB1 | chr20:33302615-33302748 | 1 | 1 | 0 | 2 | 0 | 1 | 0 | 1 | 0 | -1 | 0 | 0 | 0 | 0 | 0 | 0 | 0 | 0 | 0 | 0 | 0 | 0 | 0 | 0 | 0 | 0 | 0 | 3 | 0 | 0 | 0 | 0 | 0 | 0 | 0 | 1 | 0 | 0 | 0 | 0 | 0 | 1 | 5 | -1 | 0 | 0 | 26 | 1 | 0 | 0 | 0 |
| LMX1A | chr1:165207184-165207614 | 0 | 0 | 0 | 0 | 0 | 0 | 0 | 0 | 0 | 0 | -1 | 0 | 0 | 0 | -1 | 0 | 0 | 0 | -1 | 0 | 0 | 0 | 0 | 0 | 4 | 0 | 0 | 1 | 0 | 0 | 0 | -1 | 0 | 1 | 0 | 1 | 1 | 0 | -1 | 0 | 0 | 1 | 1 | 0 | 0 | 0 | 21 | 0 | 0 | 0 | 0 |
| MYPOP | chr19:45892170-45892279 | 1 | 2 | 2 | 2 | 2 | 1 | 3 | 1 | 0 | 1 | 1 | 1 | 1 | 1 | 1 | 1 | 1 | 1 | 1 | 0 | 1 | 4 | 2 | 2 | 4 | 1 | 2 | 1 | 1 | 1 | 1 | 2 | 3 | 3 | 3 | 2 | 1 | 1 | 1 | 2 | 1 | 1 | 1 | 4 | 3 | 3 | 24 | 2 | 1 | 1 | 2 |
| ANXA4 | chr2:69803393-69803515 | 4 | 1 | 1 | 2 | 1 | 2 | 1 | 3 | 1 | 1 | 1 | 2 | 1 | 1 | 1 | 1 | 1 | 1 | 1 | 2 | 1 | 2 | 2 | 1 | 1 | 1 | 1 | 1 | 2 | 1 | 1 | 1 | 0 | 2 | 2 | 3 | 2 | 1 | 1 | 1 | 2 | 3 | 4 | 1 | 2 | 1 | 25 | 1 | 2 | 2 | 1 |
| SUMF1 | chr3:3824871-3825381 | 3 | 3 | 3 | 3 | 0 | 3 | -1 | 3 | -1 | 6 | 8 | 6 | 8 | 4 | 9 | -1 | 8 | 7 | -1 | 0 | -1 | -1 | 0 | 0 | 4 | 2 | 3 | 1 | 0 | 2 | 1 | -1 | 3 | 2 | 4 | 1 | 1 | 3 | 4 | 7 | 5 | 4 | 4 | 5 | 4 | 4 | 55 | 5 | 0 | 2 | 3 |
| RUNX2 | chr6:45492881-45492962 | 3 | 3 | 6 | 5 | 4 | 3 | 0 | 3 | 6 | 1 | 8 | 0 | -1 | 2 | 4 | 1 | 3 | 3 | 8 | 3 | 3 | -1 | -1 | 2 | 3 | 4 | 3 | 3 | -1 | 6 | 2 | 2 | -1 | 3 | 3 | 2 | 4 | 6 | 3 | 2 | 2 | 3 | 3 | 4 | 2 | 5 | 51 | 2 | 3 | 2 | 2 |
| CA10 | chr17:51633274-51633378 | 0 | 0 | 1 | 0 | -1 | 0 | -1 | -1 | 0 | 0 | 0 | 0 | 0 | 0 | 0 | 0 | 0 | 0 | 0 | 0 | 0 | 0 | 0 | 1 | 2 | 0 | 3 | 0 | 0 | 0 | 0 | 0 | 0 | 2 | 3 | 1 | 0 | 0 | 0 | 0 | 0 | -1 | 0 | 1 | 0 | 0 | 30 | 5 | 0 | 2 | 0 |
| DSG2 | chr18:31509298-31509335 | 2 | 2 | 0 | 2 | 2 | 2 | 1 | 2 | -1 | -1 | -1 | 0 | -1 | 0 | -1 | 0 | 0 | 0 | 0 | 0 | 0 | 0 | 0 | 2 | 2 | 1 | 2 | 2 | 0 | 1 | 2 | 2 | 1 | 2 | 2 | 4 | 1 | 1 | 2 | 1 | 2 | 1 | 1 | 2 | 0 | 3 | 26 | 2 | 3 | 1 | 0 |
| INPP1 | chr2:190365113-190365202\|190365683 | 5 | 5 | 2 | 6 | 5 | 6 | 4 | 6 | 2 | 2 | 2 | 2 | 2 | 2 | 2 | 2 | 2 | 2 | 2 | 2 | 2 | 3 | 5 | 4 | 3 | 4 | 3 | 4 | 9 | 2 | 3 | 3 | 8 | 5 | 3 | 4 | 7 | 4 | 6 | 4 | 3 | 4 | 3 | 4 | 4 | 5 | 44 | 3 | 6 | 3 | 4 |
| PC | chr11:66856146-66860669 | 1 | 4 | 1 | 1 | 2 | 0 | 1 | 3 | 0 | 0 | 0 | 0 | 0 | 0 | 0 | 1 | 0 | 0 | 0 | 0 | 1 | 1 | 0 | 1 | 1 | 3 | 1 | 1 | 3 | 1 | 1 | 1 | 0 | 1 | 1 | 0 | 1 | 0 | 1 | 1 | 0 | 1 | 0 | 2 | 1 | 0 | 21 | 1 | 0 | 1 | 1 |
| GSTCD | chr4:105832935-105833041 | 2 | 2 | 1 | 1 | 2 | 2 | 4 | 3 | 1 | 0 | 0 | 1 | 2 | 1 | 0 | 1 | 0 | 1 | 0 | 2 | 0 | 0 | 0 | 1 | 1 | 1 | 1 | 1 | -1 | 1 | 1 | 2 | 1 | 1 | 1 | 2 | 2 | 3 | 2 | 2 | 1 | 2 | 4 | 1 | 0 | 2 | 26 | 2 | 1 | 2 | 1 |
| RB1CC1 | chr8:52685937-52686156 | 4 | 6 | 2 | 5 | 5 | 5 | 4 | 4 | 10 | 9 | 9 | 9 | 9 | 8 | 7 | 8 | 7 | 10 | 9 | 7 | 6 | 5 | 3 | 5 | 4 | 5 | 3 | 5 | 3 | 9 | 6 | 6 | 5 | 6 | 4 | 5 | 5 | 7 | 3 | 8 | 5 | 4 | 4 | 4 | 5 | 4 | 42 | 4 | 5 | 4 | 3 |
| TMEM231 | chr16:75542780-75542979 | 1 | 2 | 0 | 1 | 2 | 1 | 0 | 2 | 1 | 0 | 1 | 0 | 0 | 0 | 0 | 1 | 1 | 1 | 0 | 1 | 0 | 1 | 3 | 1 | 1 | 1 | 1 | 1 | 2 | 1 | 1 | 1 | 2 | 4 | 2 | 2 | 2 | 1 | 1 | 1 | 3 | 2 | 2 | 1 | 1 | 1 | 22 | 1 | 1 | 2 | 3 |
| NCAM2 | chr21:21385604-21385719 | 3 | 2 | 1 | 1 | 1 | 1 | 0 | 2 | 1 | 1 | 2 | 2 | 1 | 1 | 1 | 1 | 1 | 1 | 1 | 2 | 1 | -1 | 0 | 1 | 2 | 1 | 1 | 1 | 0 | 1 | 2 | 0 | 7 | 1 | 3 | 2 | 2 | 1 | 2 | 1 | 2 | 2 | 4 | 1 | 1 | 2 | 34 | 2 | 1 | 2 | -1 |
| INTS12 | chr4:105702886-105703023 | 5 | 4 | 3 | 6 | 5 | 5 | 5 | 6 | 2 | 2 | 3 | 3 | 4 | 2 | 2 | 2 | 2 | 4 | 4 | 2 | 3 | -1 | -1 | 4 | 3 | 4 | 5 | 4 | 7 | 4 | 4 | 3 | 4 | 5 | 5 | 5 | 5 | 4 | 5 | 5 | 5 | 6 | 6 | 7 | 5 | 4 | 44 | 3 | 3 | 3 | 3 |
| TLK2 | chr17:62574345-62574410 | 2 | 4 | 3 | 3 | 4 | 2 | 2 | 3 | 2 | 1 | 2 | 1 | 2 | 2 | 1 | 2 | 3 | 3 | 2 | 3 | 2 | 6 | 0 | 4 | 3 | 3 | 4 | 3 | -1 | 4 | 3 | 3 | 4 | 3 | 4 | 1 | 4 | 2 | 3 | 2 | 3 | 4 | 3 | 5 | 5 | 3 | 38 | 4 | 3 | 4 | 5 |
| RHOBTB1 | chr10:60936213-60936284 | 6 | 5 | 3 | 4 | 3 | 4 | 5 | 5 | 3 | 3 | 8 | 5 | 7 | 3 | 3 | 2 | 6 | 3 | 5 | 3 | 5 | 6 | 3 | 3 | 4 | 4 | 4 | 3 | 5 | 5 | 5 | 8 | 7 | 5 | 7 | 3 | 5 | 5 | 5 | 7 | 5 | 6 | 5 | 7 | 6 | 6 | 35 | 5 | 5 | 4 | 4 |

SFig5, pg12

| Gene Symbol | Exon Location (hg38, human) | Adipose - Subcutaneous | Adipose - Visceral (Omentum) | Adrenal Gland | Artery - Aorta | Artery - Coronary | Artery - Tibial | Bladder | Breast - Mammary Tissue | Brain - Amygdala | Brain - Anterior cingulate cortex (BA24) | Brain - Caudate (basal ganglia) | Brain - Cerebellar Hemisphere | Brain - Cerebellum | Brain - Cortex | Brain - Frontal Cortex (BA9) | Brain - Hippocampus | Brain - Hypothalamus | Brain - Nucleus accumbens (basal ganglia) | Brain - Putamen (basal ganglia) | Brain - Spinal cord (cervical c-1) | Brain - Substantia nigra | Cervix - Ectocervix | Cervix - Endocervix | Colon - Sigmoid | Colon - Transverse | Esophagus - Gastroesophageal Junction | Esophagus - Mucosa | Esophagus - Muscularis | Fallopian Tube | Heart - Atrial Appendage | Heart - Left Ventricle | Kidney - Cortex | Liver | Lung | Minor Salivary Gland | Muscle - Skeletal | Nerve - Tibial | Ovary | Pancreas | Pituitary | Prostate | Skin - Not Sun Exposed (Suprapubic) | Skin - Sun Exposed (Lower leg) | Small Intestine - Terminal Ileum | Spleen | Stomach | Testis | Thyroid | Uterus | Vagina | Whole Blood |
|---|---|---|---|---|---|---|---|---|---|---|---|---|---|---|---|---|---|---|---|---|---|---|---|---|---|---|---|---|---|---|---|---|---|---|---|---|---|---|---|---|---|---|---|---|---|---|---|---|---|---|---|---|
| FAM19A2 | chr12:61742911-61743013 | 1 | 2 | 3 | 1 | 0 | 1 | 3 | 1 | 0 | 1 | 1 | 1 | 1 | 1 | 1 | 1 | 1 | 1 | 2 | 1 | 1 | 0 | 0 | 0 | 2 | 2 | 1 | 2 | -1 | 1 | 0 | -1 | 0 | 2 | 1 | 2 | 2 | 0 | 0 | 0 | 1 | 0 | 1 | 1 | 1 | 2 | 23 | 2 | 1 | 1 | 1 |
| EVI2A | chr17:31320373-31320513 | 1 | 2 | 3 | 2 | 2 | 1 | 3 | 2 | 2 | 2 | 2 | 2 | 2 | 2 | 2 | 2 | 2 | 2 | 2 | 2 | 2 | 0 | 0 | 2 | 2 | 2 | 2 | 2 | 0 | 1 | 1 | 3 | 1 | 2 | 3 | 2 | 2 | 2 | 3 | 2 | 1 | 2 | 2 | 2 | 2 | 2 | 21 | 2 | 2 | 2 | 2 |
| THCAT158 | chr17:47305522-47305710 | 3 | 2 | 0 | 1 | 3 | 1 | 0 | 2 | -1 | -1 | 0 | 0 | 0 | 0 | 0 | -1 | 3 | 0 | 0 | 1 | 0 | -1 | 0 | 2 | 1 | 3 | 3 | 2 | -1 | 2 | 0 | -1 | 1 | 3 | 0 | 3 | 3 | 2 | 3 | -1 | 0 | 0 | 2 | 2 | 0 | 4 | 35 | 1 | 1 | -1 | -1 |
| EXOC6B | chr2:72438766-72438828 | 2 | 0 | 0 | 0 | 0 | 1 | 0 | 2 | 0 | 0 | 0 | 0 | 0 | 0 | 0 | 0 | 0 | 0 | 0 | 0 | 0 | 1 | 0 | 1 | 1 | 1 | 1 | 0 | 0 | 1 | 0 | 0 | 0 | 1 | 1 | 1 | 1 | 0 | 0 | 1 | 1 | 3 | 4 | 1 | 1 | 1 | 22 | 0 | 0 | 1 | 3 |
| DSTN | chr20:17591932-17592063 | 5 | 5 | 5 | 4 | 4 | 4 | 4 | 5 | 11 | 10 | 10 | 5 | 7 | 9 | 7 | 10 | 9 | 12 | 10 | 11 | 9 | 6 | 5 | 6 | 5 | 5 | 5 | 5 | 5 | 6 | 4 | 5 | 4 | 5 | 5 | 5 | 6 | 5 | 4 | 7 | 6 | 4 | 4 | 6 | 5 | 5 | 23 | 5 | 6 | 5 | 3 |
| TMTC3 | chr12:88172966-88173089 | 2 | 2 | 2 | 2 | 2 | 3 | 4 | 4 | 2 | 8 | 2 | 7 | 10 | 5 | 4 | 5 | 1 | 0 | 0 | 2 | 1 | 3 | 0 | 3 | 2 | 3 | 2 | 1 | -1 | 2 | 1 | 4 | 1 | 2 | 5 | 2 | 2 | 2 | 2 | 4 | 4 | 3 | 3 | 5 | 3 | 4 | 44 | 3 | 3 | 3 | 2 |
| RP11-164O23.5 | chr1:207147005-207147087 | 2 | 2 | 1 | 1 | 0 | 2 | 0 | 1 | 2 | 1 | 1 | 0 | 2 | 1 | 1 | 1 | 2 | 2 | 1 | 3 | 2 | -1 | 0 | 3 | 1 | 1 | 1 | 2 | 0 | 2 | 2 | 0 | -1 | 3 | 2 | 1 | 3 | 2 | 2 | 1 | 2 | 4 | 3 | 5 | 2 | 1 | 33 | 2 | 3 | 2 | -1 |
| IQGAP3 | chr1:156555951-156556148 | 3 | 1 | 1 | 1 | 1 | 4 | 0 | 2 | 0 | 0 | 0 | 0 | 0 | 0 | 0 | 0 | 0 | 0 | 0 | 0 | 0 | 0 | 0 | 1 | 1 | 1 | 1 | 2 | 0 | 1 | 1 | 0 | 2 | 2 | 0 | 0 | 0 | 0 | 0 | 0 | 3 | 2 | 1 | 1 | 0 | 1 | 24 | 1 | 0 | 2 | 1 |
| MICU1 | chr10:72532995\|72531460-72533162 | 1 | 1 | 1 | 1 | 1 | 1 | 1 | 1 | 1 | 2 | 1 | 7 | 8 | 2 | 2 | 1 | 2 | 2 | 2 | 1 | 1 | 1 | 2 | 1 | 1 | 1 | 2 | 1 | 2 | 2 | 2 | 1 | 1 | 1 | 2 | 4 | 1 | 2 | 1 | 3 | 1 | 1 | 1 | 1 | 1 | 1 | 32 | 1 | 1 | 1 | 1 |
| ALK | chr2:29294113-29294256 | 0 | 0 | 2 | 0 | 0 | -1 | -1 | 2 | 0 | 2 | 1 | 3 | 2 | 1 | 1 | 1 | 1 | 2 | 1 | 2 | 1 | -1 | -1 | 2 | 0 | 1 | 1 | 1 | -1 | 0 | -1 | 0 | 0 | 0 | 0 | 0 | 0 | 0 | 0 | 2 | -1 | -1 | -1 | 0 | 0 | 1 | 21 | 0 | 0 | 0 | 0 |
| KCNT2 | chr1:196228532-196228722 | 4 | 3 | 1 | 2 | 2 | 2 | 1 | 3 | 1 | 0 | 1 | 4 | 2 | 1 | 1 | 1 | 0 | 1 | 1 | 2 | 1 | 0 | 0 | 2 | 1 | 1 | 1 | 2 | -1 | 2 | 1 | 3 | 3 | 3 | 2 | 3 | 4 | 1 | 1 | 1 | 1 | 5 | 4 | 3 | 3 | 2 | 31 | 2 | 1 | 1 | -1 |
| PNPT1 | chr2:55654454-55654500 | 3 | 2 | 1 | 1 | 1 | 2 | 2 | 3 | 1 | 1 | 1 | 0 | 1 | 1 | 1 | 1 | 1 | 1 | 1 | 1 | 1 | 3 | 4 | 2 | 2 | 1 | 1 | 2 | 1 | 1 | 1 | 1 | 1 | 2 | 1 | 1 | 2 | 1 | 3 | 1 | 2 | 3 | 4 | 3 | 3 | 2 | 27 | 2 | 2 | 2 | 5 |
| DBNL | chr7:44067801\|44067784-44067984 | 6 | 5 | 6 | 6 | 5 | 6 | 5 | 5 | 4 | 1 | 4 | 3 | 4 | 2 | 3 | 3 | 3 | 3 | 3 | 5 | 4 | 3 | 5 | 4 | 5 | 4 | 3 | 5 | 9 | 5 | 5 | 5 | 4 | 7 | 7 | 7 | 6 | 7 | 5 | 6 | 6 | 6 | 6 | 6 | 7 | 5 | 36 | 5 | 5 | 5 | 10 |
| ST3GAL3 | chr1:43901522-43901678 | 3 | 1 | 2 | 1 | 2 | 1 | 0 | 5 | 1 | 1 | 2 | 2 | 1 | 1 | 1 | 0 | 0 | 1 | 2 | -1 | 1 | 0 | -1 | 1 | 1 | 2 | 7 | 1 | 0 | 3 | 2 | 0 | -1 | 4 | -1 | 2 | 2 | 3 | 0 | 5 | 2 | 3 | 2 | 3 | 2 | 3 | 44 | 3 | 0 | 6 | 2 |
| TMPRSS4 | chr11:118113923-118114052 | 4 | 3 | 3 | 1 | -1 | 2 | 4 | 8 | 0 | 0 | -1 | 0 | 0 | 0 | -1 | 0 | -1 | 0 | 0 | -1 | -1 | 3 | 4 | 3 | 2 | 2 | 2 | 1 | -1 | -1 | 5 | 4 | 3 | 11 | 2 | 2 | 7 | 1 | 2 | 4 | 5 | 6 | 7 | 3 | 6 | 4 | 61 | 3 | 1 | 4 | 5 |
| REV3L | chr6:111330186-111331056\|111330312 | 5 | 6 | 6 | 6 | 5 | 5 | 6 | 5 | 9 | 7 | 7 | 7 | 7 | 7 | 7 | 8 | 10 | 10 | 10 | 6 | 8 | 4 | 7 | 5 | 5 | 4 | 4 | 5 | 5 | 7 | 7 | 7 | 8 | 4 | 5 | 4 | 6 | 4 | 6 | 6 | 6 | 6 | 5 | 5 | 6 | 6 | 28 | 6 | 5 | 4 | 5 |
| LRRC49 | chr15:70898367-70898455 | 2 | 2 | 0 | 0 | 0 | 1 | 1 | 1 | 0 | 0 | 0 | 0 | 0 | 0 | 0 | 0 | 0 | 0 | 0 | 0 | 0 | 0 | 1 | 1 | 1 | 0 | 1 | 0 | 1 | 0 | 1 | 2 | 3 | 2 | 4 | 1 | 1 | 0 | 3 | 0 | 0 | 2 | 2 | 1 | 0 | 1 | 27 | 1 | 0 | 0 | 5 |
| FNBP1L | chr1:93522888-93523091 | 4 | 4 | 1 | 3 | 4 | 4 | 4 | 4 | 2 | 3 | 2 | 8 | 7 | 3 | 1 | 3 | 3 | 2 | 2 | 2 | 2 | 2 | 10 | 3 | 2 | 3 | 2 | 2 | 1 | 2 | 2 | 3 | 2 | 3 | 3 | 2 | 5 | 6 | 2 | 4 | 4 | 4 | 4 | 2 | 4 | 2 | 41 | 3 | 5 | 5 | 3 |
| NMNAT3 | chr3:139672603-139672712 | 1 | 2 | 2 | 1 | 1 | 1 | 1 | 2 | 1 | 0 | 1 | 1 | 1 | 0 | 1 | 1 | 1 | 1 | 0 | 0 | 2 | 6 | -1 | 1 | 1 | 1 | 2 | 1 | 4 | 1 | 1 | 1 | 2 | 1 | 2 | 2 | 1 | 2 | 2 | 2 | 1 | 2 | 2 | 2 | 2 | 1 | 27 | 1 | 1 | 1 | 2 |
| SP4 | chr7:21481502-21481682 | 1 | 2 | 1 | 3 | 1 | 2 | 1 | 1 | 2 | 2 | 1 | 1 | 2 | 3 | 1 | 1 | 1 | 1 | 1 | 5 | 1 | 0 | 0 | 2 | 3 | 2 | 2 | 1 | 1 | 2 | 1 | 7 | 3 | 3 | 4 | 2 | 2 | 3 | 4 | 3 | 3 | 3 | 2 | 3 | 4 | 2 | 31 | 4 | 3 | 3 | 3 |
| ADH4 | chr4:99124692-99124802 | 2 | 1 | 2 | 1 | 0 | 1 | 0 | 1 | 0 | 0 | 0 | 0 | 0 | 0 | 0 | 0 | -1 | 0 | -1 | 0 | -1 | -1 | 0 | 3 | 1 | 1 | 2 | 1 | 0 | 2 | 1 | 1 | 1 | 2 | 2 | 2 | 1 | 1 | 2 | 4 | 2 | 3 | 2 | 3 | 1 | 3 | 24 | 1 | 0 | 1 | 1 |
| MOBP | chr3:39517716-39517785 | 2 | 2 | 1 | 1 | 0 | 2 | -1 | 2 | 1 | 1 | 1 | 1 | 1 | 1 | 1 | 1 | 1 | 1 | 1 | 0 | 1 | -1 | -1 | 2 | 2 | 1 | 1 | 0 | -1 | 0 | 0 | 0 | 1 | 1 | 0 | 3 | 1 | 1 | 1 | 1 | -1 | 3 | 1 | 0 | 0 | 1 | 25 | 1 | -1 | 0 | 1 |
| POLK | chr5:75568677-75568790 | 6 | 4 | 3 | 4 | 4 | 5 | 5 | 6 | 5 | 4 | 5 | 7 | 8 | 5 | 4 | 5 | 4 | 5 | 6 | 5 | 5 | 5 | 11 | 5 | 5 | 4 | 4 | 4 | 7 | 3 | 3 | 6 | 7 | 5 | 6 | 2 | 8 | 6 | 6 | 7 | 6 | 7 | 7 | 7 | 7 | 6 | 45 | 7 | 5 | 5 | 3 |
| SGTB | chr5:65704687-65704796 | 2 | 3 | 2 | 2 | 2 | 2 | 1 | 2 | 3 | 3 | 4 | 3 | 3 | 3 | 2 | 3 | 2 | 5 | 4 | 3 | 2 | 3 | 0 | 2 | 3 | 2 | 2 | 2 | 0 | 3 | 2 | 2 | 3 | 2 | 2 | 2 | 3 | 3 | 1 | 2 | 2 | 2 | 2 | 3 | 2 | 3 | 22 | 2 | 3 | 4 | 1 |
| RP11-571M6.15 | chr12:57774629-57774767 | 13 | 11 | 11 | 10 | 10 | 10 | 12 | 12 | 5 | 2 | 6 | 6 | 4 | 2 | 4 | 6 | 5 | 4 | 2 | 6 | 3 | 7 | 10 | 11 | 12 | 11 | 12 | 11 | 9 | 8 | 10 | 12 | 11 | 12 | 11 | 9 | 10 | 8 | 14 | 4 | 13 | 10 | 11 | 11 | 8 | 11 | 59 | 9 | 9 | 8 | 13 |
| ZEB1 | chr10:31355147-31387798 | 3 | 3 | 2 | 2 | 2 | 2 | 2 | 2 | 1 | 2 | 2 | 2 | 2 | 2 | 2 | 2 | 2 | 2 | 1 | 1 | 2 | 3 | 3 | 2 | 2 | 2 | 3 | 2 | 3 | 2 | 2 | 4 | 3 | 3 | 3 | 1 | 3 | 3 | 7 | 4 | 3 | 3 | 2 | 3 | 6 | 2 | 28 | 4 | 1 | 3 | 2 |
| CDKL3 | chr5:134299656-134299750 | 1 | 2 | 1 | 1 | 1 | 1 | -1 | 0 | 0 | 0 | 0 | 0 | 0 | 0 | 0 | 0 | 0 | 0 | 0 | 0 | 0 | 0 | 0 | 1 | 1 | 1 | 1 | 0 | 0 | 0 | 0 | 2 | 1 | 1 | 0 | 0 | 0 | 1 | 1 | 0 | 0 | 2 | 1 | 1 | 3 | 2 | 21 | 0 | 0 | 1 | 4 |
| RP11-255H23.5 | chr19:23834267-23834404 | 2 | 3 | 3 | 1 | 3 | 2 | -1 | 2 | 1 | 1 | 2 | 1 | 0 | 3 | 0 | 1 | 0 | 0 | 1 | 1 | 0 | 0 | 0 | 5 | 3 | 1 | 6 | 2 | 6 | 2 | 2 | 2 | -1 | 4 | 3 | 2 | 2 | 1 | 3 | 1 | 5 | 2 | 2 | 0 | 3 | 7 | 43 | 2 | 1 | 1 | -1 |
| RECK | chr9:36113544\|36113679-36113871 | 2 | 1 | 2 | 1 | 1 | 1 | 0 | 1 | 1 | 2 | 2 | 4 | 4 | 2 | 1 | 1 | 1 | 2 | 2 | 4 | 1 | 1 | 1 | 2 | 1 | 1 | 1 | 1 | 1 | 1 | 2 | 3 | 3 | 2 | 1 | 2 | 2 | 1 | 2 | 4 | 2 | 2 | 2 | 2 | 3 | 1 | 21 | 2 | 1 | 1 | 3 |
| RP11-207C16.4 | chr9:5719875-5720020 | 7 | 3 | 4 | 3 | 3 | 4 | -1 | 6 | 4 | 5 | 3 | 5 | 3 | 5 | 4 | 4 | 4 | 3 | 4 | 4 | 2 | -1 | -1 | 3 | 5 | 3 | 4 | 4 | -1 | 3 | 3 | 3 | 1 | 3 | 2 | 13 | 7 | 3 | 5 | 6 | 3 | 8 | 9 | 5 | 3 | 4 | 64 | 4 | 4 | 3 | 3 |
| EFCAB13 | chr17:47328793-47329038 | 2 | 2 | 1 | 2 | 2 | 2 | 1 | 3 | 5 | 5 | 5 | 3 | 3 | 3 | 3 | 3 | 3 | 7 | 5 | 3 | 4 | 0 | 2 | 2 | 2 | 2 | 2 | 1 | 3 | 2 | 1 | 3 | 5 | 3 | 1 | 3 | 3 | 2 | 2 | 1 | 2 | 2 | 2 | 2 | 2 | 2 | 22 | 1 | 3 | 2 | 5 |
| CDCA3 | chr12:6847241-6847367 | 4 | 6 | 3 | 2 | 1 | 0 | 0 | 2 | 0 | 0 | 0 | 0 | 0 | 1 | 0 | 0 | 0 | -1 | 0 | 0 | 0 | 0 | 0 | 2 | 6 | 5 | 5 | 4 | 0 | 6 | 2 | -1 | 2 | 3 | 1 | 1 | 2 | -1 | 4 | 3 | 1 | 3 | 2 | 2 | 5 | 6 | 48 | 3 | 1 | 2 | 3 |
| WDFY3 | chr4:84816037-84816150 | 3 | 3 | 2 | 2 | 2 | 3 | 2 | 4 | 1 | 1 | 1 | 1 | 2 | 1 | 1 | 1 | 0 | 1 | 1 | 0 | 0 | 0 | 0 | 3 | 2 | 2 | 2 | 2 | 4 | 3 | 3 | 3 | 2 | 3 | 2 | 4 | 3 | 2 | 3 | 2 | 3 | 4 | 4 | 3 | 6 | 3 | 27 | 3 | 2 | 4 | 3 |
| FOXP1 | chr3:70972011-70972180 | 2 | 2 | 1 | 1 | 1 | 2 | 3 | 3 | 4 | 2 | 2 | 2 | 2 | 2 | 1 | 3 | 3 | 1 | 2 | 3 | 2 | 2 | 2 | 2 | 2 | 2 | 1 | 2 | 3 | 3 | 4 | 0 | 1 | 2 | 1 | 7 | 4 | 3 | 1 | 3 | 2 | 3 | 2 | 2 | 1 | 2 | 27 | 2 | 2 | 2 | 1 |
| CLC | chr19:39735697\|39735822-39735845 | 2 | 0 | 1 | 1 | 0 | 0 | -1 | 0 | 0 | 0 | 0 | 0 | 0 | 0 | 0 | 3 | 0 | 0 | 0 | 0 | 0 | -1 | 0 | 2 | 2 | 0 | 1 | 1 | -1 | 0 | 0 | 0 | 0 | 1 | 0 | 0 | 1 | 0 | 0 | 0 | 0 | 1 | 5 | 1 | 1 | 1 | 21 | 0 | 0 | 0 | 0 |
| SMC6 | chr2:17682043-17682099 | 2 | 2 | 1 | 2 | 1 | 3 | 3 | 2 | 1 | 2 | 1 | 1 | 0 | 1 | 1 | 1 | 1 | 1 | 1 | 2 | 1 | 2 | 0 | 1 | 1 | 1 | 0 | 1 | 5 | 0 | 0 | 2 | 0 | 1 | 1 | 1 | 2 | 2 | 1 | 1 | 1 | 1 | 1 | 1 | 1 | 2 | 22 | 1 | 1 | 0 | 1 |
| PLA2G4E | chr15:41996512-41996718 | 2 | 0 | 0 | 0 | 0 | 2 | 0 | 0 | 0 | 0 | 0 | 0 | 0 | 0 | 0 | 0 | -1 | -1 | 0 | 0 | 0 | 0 | -1 | -1 | 0 | 2 | 2 | -1 | 0 | 0 | 0 | -1 | 0 | 2 | 2 | 1 | 2 | 0 | -1 | 0 | 0 | 2 | 2 | 0 | 0 | 0 | 22 | -1 | 0 | 2 | 0 |
| PTPRA | chr20:3034901-3035086 | 3 | 3 | 1 | 1 | 2 | 2 | 1 | 3 | 0 | 0 | 0 | 3 | 3 | 1 | 0 | 1 | 1 | 0 | 0 | 1 | 0 | 3 | 4 | 1 | 1 | 1 | 2 | 1 | 2 | 4 | 3 | 2 | 3 | 3 | 2 | 7 | 2 | 3 | 2 | 2 | 2 | 2 | 2 | 2 | 3 | 1 | 25 | 2 | 3 | 2 | 2 |
| GRM2 | chr3:51713964-51714148\|51714121 | 3 | 5 | -1 | 9 | 9 | 6 | 0 | 3 | 2 | 0 | 2 | 4 | 4 | 1 | 0 | 1 | 1 | 2 | 4 | 1 | 2 | -1 | 0 | 6 | 8 | 7 | 1 | 4 | 0 | 3 | 9 | 0 | 0 | 4 | 2 | 9 | 10 | 6 | 3 | 1 | 8 | 2 | 1 | 2 | 5 | 2 | 71 | 3 | 1 | 7 | 5 |
| RP11-458F8.1 | chr7:66847861-66848012\|66847998\|66848296 | 2 | 3 | 2 | 3 | 3 | 4 | 6 | 3 | 6 | 3 | 4 | 2 | 1 | 6 | 3 | 7 | 6 | 4 | 4 | 6 | 5 | 0 | 0 | 3 | 2 | 3 | 3 | 4 | 0 | 3 | 3 | 2 | 4 | 4 | 3 | 5 | 4 | 2 | 6 | 3 | 2 | 3 | 4 | 4 | 3 | 3 | 31 | 3 | 3 | 2 | 5 |
| MTFR2 | chr6:136247461-136247560 | 4 | 3 | 4 | 0 | 2 | 2 | 0 | 3 | -1 | 1 | 4 | 0 | 2 | 0 | 1 | 4 | 2 | 4 | 1 | 0 | 0 | 0 | 0 | 2 | 2 | 1 | 2 | 1 | 0 | 1 | 3 | 3 | 4 | 2 | 2 | 3 | 5 | 3 | 3 | 1 | 6 | 3 | 3 | 4 | 3 | 1 | 32 | 5 | 1 | 1 | 2 |
| PLVAP | chr19:17375456-17375572 | 0 | 0 | 0 | 0 | 0 | 0 | 0 | 0 | 1 | 0 | 1 | 1 | 2 | 1 | 1 | 1 | 2 | 3 | 3 | 1 | 1 | 0 | 0 | 0 | 0 | 0 | 0 | 0 | 0 | 0 | 0 | 0 | 1 | 0 | 0 | 7 | 0 | 0 | 0 | 0 | 0 | 0 | 0 | 0 | 0 | 0 | 24 | 0 | 0 | 0 | 1 |
| LGR5 | chr12:71558684-71558815 | 3 | 3 | 2 | 2 | 2 | 3 | 0 | 2 | 1 | 0 | 3 | 3 | 0 | 2 | 1 | 2 | 1 | 3 | 1 | 1 | 2 | -1 | 0 | 2 | 0 | 2 | 1 | 2 | 0 | 0 | 0 | 0 | 1 | 1 | 2 | 5 | 2 | 0 | -1 | 2 | 3 | 3 | 4 | 1 | 0 | 1 | 30 | 1 | 1 | 1 | -1 |
| TIAM2 | chr6:155168845-155168916 | 6 | 2 | 5 | 3 | 3 | 3 | -1 | 3 | 2 | 2 | 1 | 3 | 3 | 3 | 1 | 5 | 0 | 1 | 3 | 1 | 0 | 0 | -1 | 5 | 2 | 2 | 4 | 3 | 0 | 5 | 6 | 0 | -1 | 4 | 3 | 6 | 1 | 3 | 8 | 4 | 4 | 4 | 4 | 4 | 0 | 4 | 47 | 3 | 2 | 1 | -1 |
| RAP1A | chr1:111706663-111706777 | 3 | 3 | 1 | 2 | 2 | 3 | 4 | 4 | 3 | 4 | 3 | 5 | 5 | 4 | 4 | 4 | 3 | 3 | 3 | 4 | 4 | 6 | 4 | 3 | 2 | 2 | 2 | 2 | 2 | 3 | 2 | 2 | 2 | 2 | 2 | 3 | 6 | 5 | 2 | 3 | 4 | 4 | 4 | 3 | 2 | 2 | 25 | 3 | 3 | 4 | 2 |
| MLIP | chr6:54190583-54194371 | 3 | 3 | 2 | 1 | 1 | 2 | 0 | 2 | 3 | 3 | 2 | 1 | 0 | 2 | 2 | 5 | 1 | 1 | 3 | 0 | 3 | 0 | 0 | 2 | 3 | 1 | 3 | 1 | 0 | 1 | 1 | -1 | 5 | 1 | 1 | 1 | 1 | 1 | 1 | 3 | 2 | 2 | 1 | 3 | -1 | 1 | 26 | 3 | 3 | 2 | 1 |
| TPK1 | chr7:144646086-144646163 | 2 | 2 | 2 | 2 | 2 | 1 | 3 | 3 | 1 | 1 | 1 | 1 | 1 | 1 | 1 | 1 | 1 | 2 | 1 | 3 | 1 | 0 | 0 | 1 | 3 | 2 | 2 | 1 | -1 | 2 | 2 | 1 | 3 | 2 | 1 | 1 | 1 | 2 | 5 | 1 | 1 | 2 | 2 | 3 | 2 | 2 | 21 | 1 | 1 | 2 | 2 |
| NECTIN4 | chr1:161076698-161076933 | 3 | 3 | -1 | 3 | 1 | 1 | 2 | 4 | -1 | -1 | 0 | -1 | 0 | 0 | 0 | -1 | 0 | 0 | 0 | -1 | 0 | 1 | -1 | 2 | 1 | 1 | 1 | 0 | 0 | 2 | 1 | 0 | 0 | 3 | 2 | 7 | 1 | 3 | 1 | 0 | 5 | 1 | 2 | 1 | -1 | 1 | 33 | 2 | 0 | 1 | 3 |
| INVS | chr9:100251968-100252093 | 5 | 5 | 2 | 8 | 7 | 9 | 6 | 7 | 4 | 3 | 6 | 6 | 7 | 6 | 5 | 3 | 5 | 6 | 4 | 4 | 4 | 13 | 8 | 8 | 7 | 7 | 4 | 6 | 10 | 7 | 5 | 5 | 6 | 7 | 4 | 7 | 6 | 13 | 9 | 7 | 8 | 7 | 6 | 8 | 7 | 6 | 53 | 7 | 12 | 8 | 4 |
| CLCF1 | chr11:67369404\|67368292-67369566 | 4 | 3 | 4 | 5 | 4 | 3 | 3 | 4 | 4 | 5 | 3 | -1 | -1 | 4 | 4 | 2 | 0 | 2 | 6 | 2 | 4 | 2 | 6 | 4 | 3 | 4 | 3 | 4 | 3 | 9 | 7 | 3 | 4 | 6 | 5 | 5 | 5 | 5 | 3 | 4 | 2 | 6 | 3 | 4 | 3 | 3 | 39 | 8 | 6 | 5 | 4 |
| SPATA5 | chr4:123019162-123019249 | 2 | 3 | 2 | 3 | 2 | 2 | 3 | 3 | 2 | 3 | 1 | 5 | 5 | 2 | 2 | 3 | 2 | 2 | 5 | 3 | 2 | 3 | 4 | 3 | 2 | 3 | 2 | 2 | 0 | 3 | 1 | 2 | 3 | 4 | 4 | 2 | 3 | 3 | 3 | 4 | 4 | 4 | 5 | 3 | 3 | 2 | 22 | 3 | 2 | 3 | 3 |
| SLC12A1 | chr15:48298526-48298681 | 0 | 0 | -1 | 0 | 0 | 0 | -1 | 0 | 0 | 0 | 0 | 2 | 6 | 2 | 1 | 0 | 2 | 2 | 1 | 0 | 0 | -1 | -1 | 0 | 0 | 0 | 0 | 0 | -1 | 0 | 0 | 1 | 0 | 0 | 0 | 0 | 0 | 0 | 0 | 1 | 0 | 0 | 0 | 0 | -1 | 0 | 21 | 0 | -1 | 0 | 0 |
| CPNE8 | chr12:38725544-38725592 | 7 | 1 | 0 | 1 | 1 | 2 | 1 | 6 | 0 | 0 | 0 | 0 | 0 | 0 | 0 | 0 | 1 | 0 | 0 | 0 | 0 | 0 | 0 | 1 | 1 | 0 | 0 | 1 | 0 | 1 | 1 | 0 | 0 | 3 | 1 | 6 | 5 | 2 | 1 | 0 | 0 | 5 | 6 | 2 | 0 | 1 | 41 | 1 | 1 | 2 | 3 |
| SNCAIP | chr5:122400986-122401126 | 5 | 2 | 1 | 1 | 1 | 1 | 0 | 3 | 1 | 1 | 1 | 2 | 2 | 1 | 1 | 0 | 1 | 1 | 1 | 1 | 0 | 1 | 4 | 2 | 1 | 2 | 1 | 2 | 0 | 2 | 1 | 0 | 0 | 3 | 0 | 1 | 2 | 1 | 11 | 0 | 2 | 3 | 4 | 0 | 2 | 1 | 39 | 1 | 2 | 1 | 2 |
| PROM2 | chr2:95280994-95281043 | 4 | 6 | 7 | 4 | 5 | 2 | 4 | 3 | 7 | 3 | 4 | 2 | 0 | 3 | 2 | 3 | 6 | 3 | 4 | 6 | 3 | 4 | 5 | 2 | 3 | 3 | 3 | 2 | 2 | 2 | 4 | 3 | 2 | 4 | 3 | 4 | 6 | 3 | 4 | 3 | 5 | 3 | 3 | 3 | 7 | 3 | 32 | 4 | 3 | 4 | 2 |
| DPP10 | chr2:115400802-115400953 | 0 | 1 | 1 | -1 | 0 | -1 | 0 | 0 | 2 | 1 | 2 | 1 | 1 | 1 | 1 | 1 | 2 | 1 | 1 | 1 | 2 | 0 | 0 | 2 | 3 | 1 | 3 | 1 | 0 | 4 | 0 | 0 | -1 | 2 | -1 | 0 | 3 | -1 | 5 | 3 | 0 | 3 | 1 | 1 | 0 | 1 | 30 | 3 | 0 | -1 | -1 |
| GINS1 | chr20:25443009-25443115 | 0 | 0 | 0 | 0 | 1 | 0 | -1 | 1 | 1 | 1 | 0 | 2 | 3 | 2 | 0 | 1 | 1 | 0 | 0 | 0 | 1 | 0 | 0 | 1 | 0 | 1 | 0 | 1 | 0 | 1 | 4 | 4 | 2 | 0 | 1 | 5 | 0 | 1 | 2 | 2 | 1 | 0 | 0 | 0 | 1 | 0 | 24 | 1 | 0 | 0 | 0 |
| CFTR | chr7:117626004-117626199 | 4 | 1 | -1 | 2 | 0 | -1 | 0 | 0 | 0 | 0 | 1 | 0 | 0 | 1 | 0 | 2 | 1 | 2 | 0 | 0 | 0 | 0 | 0 | 0 | 0 | 2 | 1 | 0 | 0 | 2 | 1 | 0 | 0 | 1 | 0 | 0 | 0 | 0 | 0 | 2 | 1 | 4 | 5 | 0 | 0 | 2 | 28 | 0 | 0 | 0 | 0 |
| KCNJ15 | chr21:38298144-38298282 | 7 | 6 | 1 | 5 | 6 | 5 | 3 | 5 | -1 | -1 | 4 | 3 | -1 | -1 | 0 | 0 | 0 | -1 | 0 | 3 | 0 | 2 | 0 | 3 | 5 | 5 | 2 | 4 | -1 | 6 | 2 | 3 | 2 | 5 | 2 | 4 | 5 | 5 | 3 | 2 | 2 | 4 | 4 | 4 | 4 | 1 | 40 | 2 | 1 | 2 | 2 |
| RCOR3 | chr1:211297728-211299705\|211297843 | 5 | 2 | 1 | 1 | 1 | 2 | 1 | 4 | 0 | 1 | 0 | 1 | 1 | 1 | 1 | 1 | 1 | 0 | 1 | 1 | 1 | 1 | 1 | 1 | 2 | 1 | 1 | 1 | 3 | 1 | 0 | 3 | 2 | 3 | 2 | 1 | 4 | 2 | 3 | 1 | 1 | 3 | 4 | 5 | 2 | 3 | 25 | 2 | 1 | 1 | 2 |
| RP11-178L8.4 | chr16:87310297-87311296 | 9 | 5 | 5 | 4 | 5 | 5 | 4 | 9 | 6 | 8 | 6 | 9 | 7 | 10 | 9 | 7 | 5 | 7 | 6 | 4 | 4 | -1 | 0 | 6 | 11 | 5 | 8 | 5 | 11 | 6 | 6 | 8 | 13 | 9 | 7 | 11 | 6 | 3 | 9 | 5 | 7 | 8 | 9 | 11 | 10 | 8 | 65 | 7 | 4 | 5 | 9 |
| SCUBE2 | chr11:9045712-9045901 | 2 | 1 | 3 | 3 | 1 | 2 | 2 | 3 | 2 | 2 | 3 | 6 | 4 | 4 | 3 | 2 | 3 | 3 | 2 | 1 | 3 | 4 | 2 | 2 | 2 | 2 | 2 | 2 | 3 | 2 | 1 | 2 | 2 | 2 | 1 | 2 | 3 | 1 | 3 | 6 | 4 | 3 | 4 | 2 | 3 | 2 | 28 | 3 | 2 | 2 | 7 |
| ZNF568 | chr19:36921751-36921965 | 4 | 3 | 2 | 2 | 2 | 2 | 4 | 4 | 1 | 1 | 1 | 2 | 1 | 1 | 2 | 1 | 1 | 1 | 1 | 2 | 0 | 1 | 4 | 2 | 1 | 2 | 3 | 2 | 3 | 2 | 2 | 5 | 6 | 3 | 4 | 2 | 2 | 3 | 3 | 2 | 3 | 4 | 4 | 4 | 4 | 3 | 26 | 3 | 2 | 1 | 7 |
| VEPH1 | chr3:157314484-157314578 | 3 | 1 | 0 | 0 | 2 | 0 | 0 | 4 | 0 | 0 | 0 | 0 | 0 | 0 | 0 | 0 | 0 | 0 | 0 | 0 | 0 | -1 | 0 | 0 | 2 | 1 | 0 | 0 | 0 | 0 | 1 | 0 | 0 | 2 | 2 | 0 | 1 | 0 | 1 | 0 | 0 | 4 | 4 | 0 | 0 | 1 | 24 | 0 | 0 | -1 | 1 |
| MYH14 | chr19:50230825-50230999 | 1 | 1 | 1 | 0 | 0 | 0 | 0 | 0 | 5 | 9 | 5 | 18 | 20 | 7 | 7 | 6 | 5 | 6 | 5 | 2 | 3 | 0 | 0 | 0 | 0 | 0 | 0 | 0 | 0 | 3 | 1 | 0 | 0 | 0 | 0 | 0 | 2 | 0 | 0 | 1 | 1 | 0 | 0 | 0 | 0 | 0 | 63 | 1 | 0 | 0 | 0 |
| YTHDF3 | chr8:63172653-63172798\|63173681 | 7 | 7 | 4 | 6 | 6 | 6 | 6 | 7 | 7 | 8 | 7 | 7 | 6 | 9 | 9 | 7 | 7 | 6 | 7 | 6 | 7 | 6 | 9 | 6 | 5 | 5 | 5 | 5 | 2 | 8 | 7 | 6 | 7 | 8 | 6 | 8 | 8 | 7 | 5 | 8 | 7 | 6 | 7 | 7 | 6 | 6 | 31 | 7 | 7 | 7 | 7 |
| SCUBE3 | chr6:35218014-35218156 | 1 | 0 | 1 | 1 | 0 | 1 | 1 | 2 | 0 | -1 | -1 | 0 | -1 | 2 | 0 | 0 | 0 | 0 | 0 | 0 | 0 | -1 | 0 | 1 | 1 | 1 | 2 | 1 | 0 | 1 | 4 | 0 | 0 | 2 | 0 | 1 | 1 | 4 | 0 | 1 | 1 | 0 | 2 | 2 | 0 | 0 | 21 | 1 | 1 | 0 | 0 |
| MACROD2 | chr20:15977454-15977534 | 1 | 3 | 1 | 3 | 1 | 2 | -1 | 3 | 2 | 2 | 2 | 1 | 1 | 2 | 2 | 2 | 3 | 2 | 2 | 2 | 2 | -1 | 0 | 2 | 2 | 2 | 3 | 2 | -1 | 2 | 2 | 5 | 6 | 3 | 2 | 2 | 2 | 2 | 7 | 2 | 3 | 1 | 2 | 2 | 4 | 2 | 32 | 3 | 5 | 2 | 2 |
| IL16 | chr15:81295332-81295520 | 7 | 7 | 3 | 4 | 5 | 4 | 4 | 7 | 11 | 1 | 7 | 6 | 6 | 4 | 3 | 20 | 7 | 7 | 3 | 5 | -1 | 3 | 1 | 6 | 7 | 6 | 5 | 6 | 6 | 6 | 8 | 7 | 11 | 6 | 7 | 7 | 6 | 8 | 6 | 7 | 5 | 7 | 7 | 5 | 4 | 6 | 39 | 6 | 5 | 4 | 3 |
| PLIN1 | chr15:89670387-89670708 | 0 | 0 | 1 | 1 | 0 | 1 | 0 | 0 | 1 | 1 | 0 | 2 | 1 | 2 | 1 | 2 | 1 | 1 | 0 | 2 | 0 | -1 | 0 | 0 | 1 | 1 | 6 | 1 | 0 | 1 | 1 | 0 | 1 | 6 | 0 | 2 | 0 | 6 | 2 | 6 | 7 | 0 | 0 | 0 | 2 | 1 | 51 | 2 | 11 | 7 | 2 |
| PCNX1 | chr14:71048990-71049145 | 13 | 12 | 4 | 10 | 9 | 10 | 6 | 13 | 11 | 8 | 10 | 7 | 6 | 9 | 7 | 12 | 10 | 8 | 10 | 11 | 10 | 12 | 7 | 8 | 8 | 7 | 8 | 6 | 4 | 10 | 5 | 12 | 11 | 16 | 9 | 5 | 14 | 12 | 8 | 12 | 8 | 14 | 12 | 13 | 14 | 9 | 67 | 10 | 9 | 10 | 13 |
| TMEM156 | chr4:38969246-38969311 | 2 | 2 | 2 | 0 | 3 | 2 | 0 | 2 | 2 | 3 | 3 | 3 | 4 | 1 | 3 | 2 | 5 | 4 | 2 | 3 | 2 | 0 | 0 | 2 | 3 | 0 | 3 | 1 | -1 | 2 | -1 | 2 | 2 | 3 | 1 | 0 | 4 | -1 | 5 | 2 | 2 | 3 | 1 | 3 | 3 | 5 | 33 | 4 | 0 | 6 | 2 |
| GTF2IP5 | chr7:65794880-65795091 | 0 | 1 | 1 | 0 | 0 | 0 | 0 | 0 | 0 | -1 | 0 | -1 | 0 | 0 | 0 | 0 | -1 | -1 | -1 | 3 | -1 | 0 | 0 | 0 | 0 | 2 | 1 | 3 | -1 | 0 | 4 | 0 | 4 | 1 | 0 | 4 | 0 | 0 | 0 | 1 | 0 | 1 | 0 | 0 | 0 | 0 | 21 | 1 | 0 | 1 | 0 |
| SCD | chr10:100352622-100352868 | 1 | 1 | 1 | 1 | 2 | 3 | 3 | 2 | 0 | 0 | 0 | 0 | 1 | 1 | 0 | 0 | 0 | 0 | 0 | 1 | 0 | 5 | 0 | 2 | 2 | 3 | 3 | 1 | 0 | 4 | 1 | 1 | 4 | 5 | 4 | 2 | 2 | 7 | 2 | 5 | 4 | 5 | 4 | 2 | 4 | 1 | 35 | 2 | 3 | 6 | 8 |
| ACE | chr17:63478349-63478647 | 1 | 1 | 1 | 1 | 1 | 1 | 1 | 2 | 2 | 3 | 5 | 3 | 4 | 5 | 3 | 3 | 6 | 3 | 4 | 3 | 4 | 0 | 2 | 1 | 1 | 1 | 2 | 1 | 4 | 3 | 2 | 4 | 2 | 2 | 2 | 1 | 2 | 3 | 3 | 10 | 2 | 1 | 1 | 1 | 1 | 1 | 31 | 2 | 2 | 2 | 3 |
| DLG2 | chr11:85625107-85625251 | 2 | 2 | 2 | 1 | 1 | 2 | -1 | 2 | 2 | 0 | 2 | 2 | 1 | 1 | 1 | 2 | 2 | 1 | 1 | 1 | 4 | 0 | 1 | 2 | 3 | 1 | 1 | 1 | 0 | 1 | 1 | 3 | 1 | 1 | 3 | 1 | 1 | 1 | 4 | 2 | 1 | 2 | 2 | 2 | 6 | 2 | 23 | 2 | 1 | 1 | 1 |
| RAD51D | chr17:35106128-35106249 | 3 | 4 | 3 | 2 | 3 | 2 | 5 | 3 | 2 | 2 | 2 | 4 | 4 | 3 | 3 | 2 | 1 | 1 | 2 | 3 | 3 | 4 | 0 | 4 | 5 | 4 | 5 | 4 | 2 | 4 | 3 | 4 | 4 | 5 | 6 | 4 | 2 | 3 | 5 | 3 | 5 | 5 | 4 | 7 | 8 | 5 | 32 | 5 | 3 | 4 | 9 |
| TBC1D19 | chr4:26740679-26740733 | 7 | 7 | 3 | 5 | 5 | 5 | 6 | 10 | 3 | 2 | 3 | 1 | 2 | 2 | 1 | 2 | 1 | 5 | 4 | 1 | 2 | 7 | 6 | 5 | 5 | 5 | 4 | 4 | 6 | 4 | 3 | 4 | 7 | 5 | 7 | 4 | 10 | 7 | 11 | 5 | 9 | 7 | 6 | 8 | 11 | 7 | 43 | 7 | 6 | 6 | 7 |
| SLC6A6 | chr3:14480120-14480238 | 2 | 3 | 1 | 2 | 2 | 2 | 0 | 2 | 0 | 1 | 0 | 1 | 0 | 1 | 0 | 1 | 1 | 0 | 1 | 0 | 1 | 1 | 0 | 2 | 2 | 2 | 1 | 1 | 3 | 1 | 1 | 2 | 5 | 4 | 2 | 1 | 2 | 1 | 2 | 1 | 2 | 2 | 2 | 3 | 4 | 2 | 25 | 2 | 1 | 2 | 7 |
| RP11-550I24.2 | chr3:157230089-157230297 | 1 | 2 | 5 | 1 | 1 | 1 | -1 | 1 | 1 | 3 | 2 | 1 | 1 | 0 | 1 | 1 | 2 | 0 | 1 | 1 | 1 | -1 | -1 | 1 | 1 | 2 | 2 | 1 | -1 | 1 | 0 | -1 | 1 | 2 | 2 | 1 | 1 | 0 | 5 | 3 | 2 | 1 | 3 | 3 | 2 | 2 | 33 | 3 | 0 | 0 | 8 |
| RP11-106M3.2 | chr15:72313664-72313741 | 6 | 6 | 5 | 7 | 7 | 8 | 7 | 7 | 5 | 2 | 5 | 3 | 4 | 3 | 3 | 3 | 2 | 4 | 5 | 5 | 4 | 7 | 6 | 8 | 7 | 7 | 6 | 7 | 6 | 5 | 6 | 7 | 7 | 6 | 6 | 6 | 6 | 7 | 8 | 4 | 6 | 7 | 7 | 8 | 8 | 7 | 23 | 6 | 7 | 5 | 6 |
| STAC | chr3:36485763-36485925 | 4 | 5 | 0 | 2 | 3 | 2 | 0 | 4 | 2 | 1 | 0 | 1 | 1 | 0 | 2 | 0 | 0 | 0 | 2 | 1 | 0 | -1 | 0 | 1 | 2 | 1 | 2 | 1 | 0 | 1 | 2 | 0 | 6 | 1 | 2 | 2 | 3 | 2 | -1 | 1 | 1 | 2 | 3 | 3 | 3 | 0 | 30 | 4 | 3 | 5 | 1 |
| CCDC153 | chr11:119193595-119193631 | 3 | 3 | 5 | 4 | 4 | 6 | -1 | 4 | 3 | 1 | 4 | 8 | 10 | 6 | 2 | 3 | 5 | 6 | 3 | 3 | 0 | -1 | -1 | 2 | 6 | 4 | 5 | 3 | 3 | 8 | 10 | 1 | 1 | 5 | 5 | 3 | 5 | 1 | 3 | 2 | 4 | 5 | 5 | 4 | 4 | 6 | 45 | 3 | 4 | 4 | 5 |
| PTAR1 | chr9:69740721-69740776 | 8 | 5 | 2 | 4 | 3 | 5 | 6 | 7 | 1 | 5 | 1 | 2 | 5 | 5 | 2 | 3 | 4 | 5 | 4 | 2 | 2 | 6 | 4 | 4 | 5 | 3 | 4 | 3 | 3 | 4 | 3 | 2 | 1 | 5 | 5 | 8 | 7 | 7 | 3 | 4 | 5 | 9 | 8 | 7 | 3 | 5 | 37 | 3 | 3 | 4 | 6 |
| SULT1A4 | chr16:29462693-29462785 | 1 | 2 | 0 | 2 | -1 | 0 | 0 | 1 | 0 | 0 | 0 | 4 | 1 | 3 | 1 | 0 | -1 | -1 | 0 | -1 | 0 | -1 | 0 | 0 | 0 | 2 | 1 | 1 | 0 | 0 | 3 | 0 | -1 | 2 | -1 | 0 | 1 | 0 | 1 | 1 | 0 | 1 | 2 | 1 | 4 | 0 | 20 | 2 | -1 | -1 | 1 |
| MACROD2 | chr20:15916742-15916843 | 1 | 2 | 2 | 1 | 1 | 1 | 5 | 3 | 0 | 0 | 0 | 0 | 1 | 0 | 0 | 1 | 0 | 0 | 0 | 1 | 1 | 0 | -1 | 1 | 2 | 1 | 2 | 1 | -1 | 1 | 1 | 0 | 3 | 4 | 3 | 1 | 1 | 0 | 1 | 1 | 2 | 2 | 2 | 0 | -1 | 1 | 23 | 1 | 1 | 2 | -1 |
| TG | chr8:133014725-133014836 | 3 | 2 | 2 | 2 | 3 | 2 | 0 | 3 | 5 | 4 | 2 | 1 | 2 | 3 | 2 | 3 | 3 | 2 | 1 | 1 | 1 | 0 | 0 | 3 | 2 | 2 | 2 | 3 | -1 | 3 | 1 | 2 | 1 | 3 | 4 | 2 | 3 | 3 | 2 | 3 | 7 | 7 | 7 | 1 | 3 | 2 | 33 | 1 | 1 | 3 | 4 |
| RP11-613M10.8 | chr9:37575522-37575661 | 1 | 3 | 3 | 3 | 2 | 1 | 1 | 1 | 2 | 2 | 5 | 3 | 3 | 2 | 4 | 3 | 4 | 3 | 1 | 4 | 3 | 0 | 0 | 2 | 3 | 2 | 2 | 3 | 0 | 3 | 2 | 6 | 9 | 3 | 4 | 3 | 2 | 1 | 4 | 5 | 3 | 2 | 1 | 2 | 4 | 3 | 31 | 3 | 3 | 2 | 2 |
| F7 | chr13:113116092-113116415 | 2 | 0 | 0 | 0 | 0 | -1 | 0 | 2 | 0 | 0 | 1 | 1 | 4 | 1 | 1 | 1 | 3 | 2 | -1 | -1 | 1 | 0 | 0 | -1 | -1 | 1 | 1 | 1 | 0 | 1 | 0 | 0 | 1 | 1 | 6 | 0 | 0 | 3 | 0 | 7 | 1 | 1 | 2 | 2 | 0 | 0 | 32 | 1 | 0 | 4 | 0 |
| NAIP | chr5:71011712-71011786 | 8 | 11 | 6 | 6 | 6 | 6 | -1 | 10 | 3 | 5 | 3 | 3 | -1 | 8 | 2 | 6 | 5 | 4 | 3 | 4 | 3 | -1 | -1 | 6 | 9 | 11 | 7 | 5 | -1 | 8 | 9 | 14 | 8 | 10 | 8 | 9 | 8 | 11 | 4 | 5 | 9 | 11 | 12 | 9 | 13 | 13 | 77 | 7 | 8 | 13 | 6 |
| BCL2L11 | chr2:111153780-111153904\|111153889\|111161550 | 20 | 16 | 17 | 14 | 15 | 19 | 13 | 16 | 13 | 13 | 17 | 23 | 24 | 22 | 26 | 16 | 14 | 11 | 18 | 14 | 16 | 12 | 13 | 14 | 14 | 15 | 13 | 14 | 18 | 19 | 18 | 12 | 17 | 17 | 13 | 27 | 20 | 16 | 14 | 17 | 16 | 18 | 22 | 12 | 14 | 14 | 77 | 17 | 13 | 13 | 7 |
| HK3 | chr5:176885812\|176884744-176885897 | 3 | 4 | 4 | 3 | 4 | 3 | 6 | 4 | 5 | 4 | 3 | 5 | 3 | 5 | 6 | 4 | 4 | 6 | 6 | 6 | 4 | 0 | -1 | 4 | 4 | 4 | 3 | 4 | 4 | 4 | 5 | 5 | 4 | 4 | 4 | 4 | 3 | 4 | 2 | 4 | 4 | 3 | 3 | 4 | 3 | 4 | 23 | 3 | 4 | 4 | 3 |
| TMEM50B | chr21:33456004-33456097 | 9 | 6 | 3 | 9 | 8 | 11 | 6 | 8 | 3 | 3 | 3 | 4 | 5 | 4 | 3 | 3 | 3 | 3 | 4 | 4 | 4 | 9 | 6 | 8 | 6 | 8 | 5 | 7 | 8 | 5 | 5 | 6 | 4 | 8 | 6 | 7 | 12 | 5 | 6 | 6 | 7 | 9 | 9 | 8 | 8 | 8 | 38 | 5 | 7 | 7 | 6 |
| CCDC15 | chr11:124991095-124991214 | 2 | 1 | 0 | 3 | 0 | 3 | 0 | 2 | 0 | 3 | 1 | 6 | 1 | 3 | 0 | 3 | 0 | -1 | 2 | 2 | 0 | -1 | 0 | 1 | 2 | 3 | 2 | 0 | 0 | 3 | 2 | -1 | 3 | 2 | 1 | 2 | 5 | 5 | 3 | 1 | 1 | 2 | 4 | 0 | 1 | 4 | 27 | 1 | 0 | 4 | 1 |
| ZDHHC3 | chr3:44932907-44932990 | 15 | 13 | 11 | 13 | 13 | 13 | 17 | 15 | 9 | 9 | 9 | 8 | 8 | 9 | 8 | 9 | 11 | 10 | 10 | 10 | 10 | 13 | 15 | 14 | 16 | 14 | 16 | 14 | 15 | 14 | 14 | 12 | 14 | 16 | 16 | 15 | 16 | 13 | 14 | 12 | 15 | 20 | 22 | 16 | 18 | 16 | 61 | 12 | 14 | 14 | 23 |
| DONSON | chr21:33576926\|33575698-33577068 | 5 | 4 | 6 | 5 | 5 | 4 | 0 | 3 | 4 | 11 | 4 | 4 | 4 | 4 | 3 | 4 | 6 | 3 | 4 | 4 | 2 | 1 | -1 | 3 | 5 | 3 | 5 | 2 | 6 | 5 | 4 | 0 | 8 | 4 | 2 | 4 | 4 | 5 | 8 | 6 | 4 | 5 | 4 | 5 | 1 | 4 | 39 | 3 | 4 | 3 | 9 |
| PDE7A | chr8:65780844-65780947 | 5 | 5 | 2 | 3 | 2 | 2 | 7 | 6 | 5 | 1 | 2 | 2 | 2 | 3 | 2 | 1 | 1 | 1 | 2 | 2 | 1 | 0 | 2 | 2 | 3 | 2 | 3 | 2 | 3 | 2 | 1 | 5 | 6 | 5 | 3 | 2 | 4 | 5 | 4 | 3 | 4 | 6 | 5 | 4 | 4 | 3 | 29 | 3 | 4 | 3 | 3 |
| PGAP2 | chr11:3822909-3822993\|3823740 | 16 | 13 | 6 | 9 | 11 | 11 | 14 | 15 | 15 | 12 | 11 | 13 | 12 | 15 | 13 | 13 | 16 | 12 | 13 | 15 | 15 | 13 | 9 | 11 | 13 | 11 | 14 | 11 | 9 | 15 | 13 | 10 | 9 | 10 | 13 | 22 | 17 | 14 | 10 | 12 | 11 | 17 | 22 | 13 | 15 | 12 | 70 | 8 | 9 | 12 | 15 |
| CEP152 | chr15:48781585-48781720 | 1 | 2 | 0 | 2 | 1 | 1 | 0 | 2 | 2 | 0 | 0 | 1 | 1 | 2 | 0 | 0 | 0 | 3 | 0 | 1 | 0 | 0 | 0 | 1 | 2 | 1 | 1 | 1 | -1 | 3 | 3 | 4 | 7 | 1 | 1 | 1 | 1 | 1 | 1 | 1 | 1 | 1 | 1 | 6 | 2 | 2 | 28 | 2 | 1 | 3 | 4 |
| CRYM | chr16:21268547-21268684 | 5 | 5 | 1 | 4 | 4 | 5 | 5 | 3 | 0 | 0 | 0 | 1 | 1 | 0 | 0 | 0 | 0 | 0 | 0 | 1 | 0 | 7 | 0 | 2 | 1 | 4 | 2 | 4 | 0 | 1 | 1 | 0 | 3 | 2 | 1 | 2 | 4 | 3 | 2 | 1 | 4 | 2 | 2 | 1 | 9 | 4 | 36 | 3 | 6 | 5 | 1 |
| HAUS6 | chr9:19079039-19079136 | 2 | 2 | 0 | 1 | 1 | 2 | -1 | 2 | 0 | 0 | 0 | 1 | 0 | 1 | 0 | 0 | 0 | 0 | 1 | 1 | 1 | 0 | -1 | 1 | 0 | 1 | 1 | 2 | 0 | 1 | 1 | -1 | 0 | 3 | 2 | 6 | 3 | 2 | 1 | 0 | 0 | 4 | 2 | 5 | 0 | 1 | 28 | 2 | 4 | 1 | 3 |
| HES7 | chr17:8123320-8124741 | 5 | 5 | 1 | 4 | 3 | 4 | 0 | 3 | 4 | 3 | 6 | 13 | 14 | 2 | 2 | 4 | 1 | 2 | 3 | 1 | 2 | -1 | 0 | 5 | 7 | 3 | 4 | 5 | -1 | 3 | 3 | -1 | -1 | 4 | 10 | 6 | 10 | 6 | -1 | 4 | 2 | 7 | 6 | 6 | 5 | 5 | 59 | 5 | 9 | 5 | 7 |
| CATSPER2 | chr15:43635727-43635826 | 14 | 10 | 6 | 10 | 8 | 8 | 9 | 12 | 1 | 0 | 1 | 0 | 1 | 0 | 1 | 1 | 1 | 1 | 3 | 1 | 2 | 3 | -1 | 4 | 6 | 4 | 7 | 7 | 15 | 4 | 7 | 13 | 7 | 7 | 9 | 2 | 9 | 5 | 8 | 3 | 6 | 11 | 17 | 10 | 10 | 7 | 68 | 8 | 7 | 7 | 6 |
| C1orf228 | chr1:44684393-44684656 | 3 | 2 | 3 | 2 | 2 | 1 | -1 | 3 | 4 | 1 | 3 | 4 | 2 | 2 | 1 | 4 | 3 | 3 | 1 | 2 | 1 | 0 | 0 | 2 | 2 | 1 | 6 | 2 | 4 | 3 | 6 | 4 | 1 | 7 | 3 | 5 | 1 | 1 | 2 | 4 | 3 | 5 | 4 | 0 | 5 | 2 | 31 | 3 | 2 | 3 | 4 |
| LINC00470 | chr18:1271611-1271804\|1271819 | 2 | 3 | 1 | 5 | 7 | 1 | 0 | 2 | -1 | 0 | 0 | -1 | -1 | 0 | -1 | 0 | 0 | 0 | 0 | 0 | 0 | 0 | -1 | 0 | 1 | 2 | 1 | 2 | 0 | 6 | 8 | 0 | 1 | 3 | 0 | -1 | 5 | -1 | 3 | 2 | 1 | 2 | 0 | 2 | 2 | 2 | 39 | 6 | 0 | 0 | -1 |
| SLC25A40 | chr7:87841932-87842041 | 4 | 3 | 2 | 4 | 4 | 3 | 5 | 5 | 1 | 1 | 2 | 2 | 3 | 2 | 1 | 2 | 1 | 2 | 1 | 2 | 1 | 2 | 6 | 4 | 4 | 4 | 2 | 3 | 5 | 3 | 2 | 2 | 2 | 4 | 4 | 2 | 6 | 6 | 2 | 3 | 3 | 4 | 4 | 5 | 6 | 3 | 24 | 4 | 4 | 4 | 3 |
| SEC24D | chr4:118766529-118766823 | 5 | 4 | 1 | 2 | 2 | 3 | 2 | 5 | 3 | 2 | 2 | 2 | 2 | 2 | 1 | 3 | 2 | 3 | 2 | 6 | 1 | 4 | 3 | 3 | 2 | 2 | 2 | 2 | 2 | 5 | 3 | 5 | 1 | 4 | 2 | 5 | 4 | 4 | 2 | 3 | 4 | 5 | 5 | 2 | 2 | 2 | 22 | 3 | 3 | 4 | 4 |

SFig5, pg13

| Gene Symbol | Exon Location (hg38, human) | Adipose - Subcutaneous | Adipose - Visceral (Omentum) | Adrenal Gland | Artery - Aorta | Artery - Coronary | Artery - Tibial | Bladder | Breast - Mammary Tissue | Brain - Amygdala | Brain - Anterior cingulate cortex (BA24) | Brain - Caudate (basal ganglia) | Brain - Cerebellar Hemisphere | Brain - Cerebellum | Brain - Cortex | Brain - Frontal Cortex (BA9) | Brain - Hippocampus | Brain - Hypothalamus | Brain - Nucleus accumbens (basal ganglia) | Brain - Putamen (basal ganglia) | Brain - Spinal cord (cervical c-1) | Brain - Substantia nigra | Cervix - Ectocervix | Cervix - Endocervix | Colon - Sigmoid | Colon - Transverse | Esophagus - Gastroesophageal Junction | Esophagus - Mucosa | Esophagus - Muscularis | Fallopian Tube | Heart - Atrial Appendage | Heart - Left Ventricle | Kidney - Cortex | Liver | Lung | Minor Salivary Gland | Muscle - Skeletal | Nerve - Tibial | Ovary | Pancreas | Pituitary | Prostate | Skin - Not Sun Exposed (Suprapubic) | Skin - Sun Exposed (Lower leg) | Small Intestine - Terminal Ileum | Spleen | Stomach | Testis | Thyroid | Uterus | Vagina | Whole Blood |
|---|---|---|---|---|---|---|---|---|---|---|---|---|---|---|---|---|---|---|---|---|---|---|---|---|---|---|---|---|---|---|---|---|---|---|---|---|---|---|---|---|---|---|---|---|---|---|---|---|---|---|---|---|
| ADGRG6 | chr6:142420998-142421084 | 2 | 1 | 3 | 2 | 2 | 2 | 2 | 2 | 4 | 1 | 1 | 0 | 4 | 1 | 2 | 1 | 1 | 1 | 1 | 1 | 3 | 5 | -1 | 5 | 4 | 3 | 1 | 3 | -1 | 2 | 2 | 2 | 3 | 1 | 2 | 2 | 4 | 3 | 2 | 1 | 3 | 2 | 2 | 2 | 2 | 2 | 22 | 2 | 2 | 2 | 1 |
| TNNI3K | chr1:74260694-74267478 | 6 | 2 | 2 | 2 | 1 | 2 | -1 | 4 | 1 | 3 | 1 | 2 | 2 | 1 | 2 | 2 | 2 | 2 | 2 | 2 | 1 | 0 | 0 | 3 | 3 | 3 | 3 | 2 | -1 | 1 | 1 | 2 | 4 | 4 | 5 | 2 | 4 | 2 | 5 | 2 | 5 | 5 | 7 | 6 | 4 | 3 | 31 | 3 | 3 | 3 | 2 |
| DPP8 | chr15:65515656-65515791 | 7 | 8 | 10 | 6 | 7 | 6 | 16 | 7 | 2 | 4 | 7 | 2 | 3 | 5 | 5 | 5 | 8 | 6 | 4 | 8 | 5 | -1 | -1 | 9 | 12 | 9 | 13 | 10 | -1 | 11 | 11 | 11 | 12 | 10 | 9 | 9 | 6 | 11 | 14 | 12 | 8 | 9 | 12 | 10 | 12 | 10 | 69 | 9 | 7 | 9 | 10 |
| ATPAF2 | chr17:18021422-18021663 | 6 | 10 | 3 | 6 | 6 | 5 | 5 | 6 | 5 | 3 | 5 | 5 | 4 | 3 | 2 | 6 | 5 | 3 | 4 | 7 | 5 | 7 | 9 | 6 | 6 | 6 | 7 | 5 | 3 | 3 | 2 | 5 | 6 | 11 | 5 | 3 | 8 | 13 | 6 | 6 | 10 | 6 | 5 | 12 | 17 | 4 | 63 | 8 | 12 | 12 | 16 |
| ZFHX2 | chr14:23530884-23531019 | 3 | 2 | 3 | 1 | 3 | 1 | 0 | 3 | 4 | 1 | 6 | 3 | 5 | 3 | 1 | 6 | 4 | 3 | 2 | 3 | 1 | 0 | -1 | 3 | 4 | 2 | 1 | 2 | 0 | 3 | 1 | 4 | -1 | 4 | 0 | 2 | 3 | 3 | 6 | 1 | 5 | 2 | 7 | 3 | -1 | 3 | 33 | 3 | 0 | -1 | 0 |
| CD24 | chr6:106973317-106974168 | 3 | 4 | 2 | 2 | 1 | 1 | 3 | 3 | 2 | 4 | 2 | 0 | -1 | 5 | 3 | 5 | 2 | 3 | -1 | 5 | 8 | 4 | 3 | 3 | 2 | 3 | 2 | 2 | -1 | 0 | 1 | 3 | 3 | 2 | 3 | 5 | 5 | 3 | 1 | 2 | 3 | 4 | 3 | 2 | 4 | 3 | 24 | 1 | 2 | 3 | 2 |
| PVRL3-AS1 | chr3:111052504-111052659 | 4 | 4 | 4 | 2 | 6 | 5 | 0 | 4 | 1 | 1 | 1 | 0 | -1 | 1 | 1 | 1 | 1 | 2 | 1 | 3 | 3 | 0 | 0 | 2 | 4 | 1 | -1 | 1 | 0 | 2 | 2 | 0 | -1 | 5 | -1 | 0 | 2 | 3 | 6 | 7 | 9 | 8 | 1 | -1 | -1 | 5 | 45 | 3 | 1 | 6 | -1 |
| BACH1 | chr21:29626126-29626276 | 5 | 6 | 10 | 4 | 4 | 2 | 0 | 5 | 4 | 4 | 5 | 6 | 8 | 4 | 5 | 2 | 1 | 6 | 4 | 3 | 11 | 0 | -1 | 3 | 4 | 4 | 8 | 4 | 0 | 7 | 6 | 2 | 8 | 5 | 5 | 3 | 7 | 6 | 12 | 10 | 8 | 4 | 5 | 4 | 9 | 4 | 52 | 7 | 3 | 5 | 10 |
| AC093818.1 | chr2:172496132-172496226 | 5 | 5 | 7 | 7 | 5 | 5 | -1 | 4 | 6 | 4 | 3 | 3 | 5 | 3 | 2 | 3 | 3 | 4 | 3 | 5 | 6 | 0 | -1 | 7 | 8 | 6 | 8 | 5 | -1 | 11 | 10 | 4 | 10 | 8 | 8 | 5 | 5 | 8 | 8 | 9 | 9 | 3 | 3 | 10 | 9 | 6 | 57 | 7 | 6 | 9 | 13 |
| SLC25A40 | chr7:87843977-87844053 | 4 | 5 | 3 | 5 | 5 | 3 | 4 | 5 | 4 | 2 | 3 | 3 | 5 | 2 | 1 | 3 | 3 | 3 | 4 | 4 | 3 | 8 | 4 | 5 | 4 | 4 | 4 | 4 | 5 | 4 | 3 | 3 | 7 | 6 | 5 | 4 | 7 | 5 | 9 | 3 | 3 | 4 | 4 | 7 | 9 | 4 | 30 | 6 | 5 | 5 | 6 |
| ALPK2 | chr18:58501684\|58503360-58503524 | 1 | 5 | -1 | 3 | 1 | 1 | 0 | 1 | 0 | 2 | 0 | -1 | -1 | 0 | -1 | 0 | -1 | 0 | 0 | 0 | 0 | -1 | -1 | 4 | 4 | 1 | 2 | 2 | -1 | 1 | 1 | 7 | 15 | 2 | 1 | 1 | 3 | 0 | 5 | -1 | 4 | 2 | 2 | 4 | 3 | 1 | 46 | 3 | 0 | 3 | 2 |
| PDS5B | chr13:32674610-32674745 | 6 | 4 | 3 | 2 | 3 | 3 | 3 | 5 | 2 | 1 | 1 | 1 | 2 | 0 | 1 | 1 | 1 | 2 | 1 | 2 | 2 | 4 | 1 | 4 | 4 | 3 | 3 | 3 | 7 | 4 | 3 | 5 | 6 | 4 | 4 | 2 | 7 | 3 | 3 | 3 | 5 | 5 | 4 | 5 | 5 | 4 | 26 | 4 | 3 | 4 | 1 |
| ALMS1 | chr2:73408622-73408747 | 2 | 4 | 1 | 4 | 2 | 2 | -1 | 3 | 3 | 7 | 2 | 3 | 2 | 3 | 5 | 3 | 2 | 4 | 3 | 3 | 2 | 0 | -1 | 3 | 5 | 6 | 4 | 3 | 0 | 4 | 4 | 3 | 4 | 3 | 5 | 2 | 2 | 3 | 4 | 4 | 5 | 2 | 2 | 3 | 3 | 3 | 28 | 3 | 5 | 2 | 5 |
| P4HA3 | chr11:74267596-74267811 | 1 | 2 | 3 | 1 | 1 | 1 | 0 | 2 | 3 | 3 | 2 | 3 | 2 | 1 | 1 | 2 | 1 | 1 | 2 | 4 | 1 | 1 | 5 | 2 | 1 | 2 | 2 | 2 | 1 | 4 | 1 | 0 | 5 | 3 | 0 | 3 | 2 | 3 | 1 | 3 | 4 | 2 | 2 | 4 | 2 | 1 | 22 | 4 | 1 | 3 | -1 |
| ICA1 | chr7:8227638-8227851 | 7 | 5 | 5 | 3 | 4 | 4 | 5 | 6 | 1 | 1 | 1 | 0 | 1 | 1 | 0 | 2 | 2 | 1 | 2 | 2 | 0 | 3 | 3 | 5 | 4 | 5 | 5 | 4 | 3 | 3 | 2 | 6 | 4 | 4 | 6 | 6 | 4 | 3 | 2 | 2 | 7 | 5 | 6 | 4 | 5 | 4 | 26 | 3 | 2 | 3 | 3 |
| RTTN | chr18:70078619\|70078609-70078738 | 2 | 1 | 0 | 1 | 1 | 1 | 0 | 1 | 0 | 2 | 1 | 4 | 5 | 2 | 1 | 1 | 1 | 1 | 0 | 1 | 1 | 0 | 0 | 1 | 1 | 1 | 2 | 1 | 0 | 2 | 1 | 3 | 3 | 2 | 1 | 7 | 2 | 1 | 2 | 1 | 1 | 1 | 3 | 1 | 2 | 1 | 22 | 1 | 1 | 0 | 1 |
| NMNAT3 | chr3:139575583-139575994 | 5 | 4 | 2 | 4 | 3 | 3 | 3 | 5 | 2 | 3 | 4 | 7 | 5 | 4 | 2 | 5 | 3 | 3 | 2 | 4 | 3 | 4 | 7 | 3 | 4 | 3 | 6 | 2 | 3 | 3 | 2 | 6 | 3 | 6 | 8 | 2 | 7 | 8 | 7 | 7 | 9 | 5 | 5 | 4 | 5 | 4 | 34 | 8 | 4 | 7 | 6 |
| UBE3D | chr6:83022647-83022820\|83022823 | 5 | 3 | 1 | 3 | 2 | 4 | 2 | 4 | 1 | 1 | 2 | 3 | 2 | 2 | 1 | 1 | 3 | 2 | 2 | 3 | 4 | -1 | 2 | 2 | 2 | 3 | 3 | 2 | 1 | 4 | 1 | 4 | 3 | 5 | 3 | 3 | 6 | 3 | 3 | 2 | 4 | 4 | 6 | 2 | 2 | 3 | 23 | 3 | 4 | 3 | 2 |
| MMP16 | chr8:88046203-88046248 | 0 | 1 | -1 | 1 | 2 | 1 | 0 | 2 | 1 | 0 | 0 | 0 | 0 | 0 | 1 | 0 | 1 | 0 | 0 | 0 | 0 | -1 | 0 | 1 | 1 | 1 | 0 | 0 | 0 | 0 | 0 | -1 | 0 | 3 | -1 | 0 | 4 | 1 | -1 | 1 | 4 | 1 | 1 | 0 | 0 | 3 | 20 | 1 | 0 | 0 | -1 |
| RNF115 | chr1:145788612-145788802 | 4 | 4 | 4 | 6 | 4 | 5 | 3 | 4 | 5 | 5 | 5 | 6 | 6 | 5 | 5 | 6 | 7 | 5 | 5 | 7 | 5 | 3 | 5 | 3 | 3 | 3 | 4 | 2 | 3 | 5 | 3 | 6 | 5 | 7 | 5 | 5 | 4 | 6 | 4 | 10 | 4 | 5 | 6 | 5 | 5 | 4 | 28 | 8 | 4 | 5 | 8 |
| AC007249.3 | chr2:10454827-10455089 | 3 | 1 | 1 | 2 | 4 | 2 | -1 | 6 | 1 | 1 | 1 | 1 | 0 | 0 | 1 | 0 | 1 | 0 | 0 | 1 | 0 | 0 | -1 | 3 | 1 | 2 | 4 | 1 | 0 | 3 | 3 | -1 | 1 | 6 | 3 | 1 | 2 | 2 | 1 | 1 | 2 | 6 | 6 | 1 | 1 | 2 | 29 | 2 | 2 | 1 | 3 |
| UBR2 | chr6:42634248-42634349\|42634294 | 10 | 7 | 2 | 6 | 5 | 6 | 7 | 11 | 4 | 2 | 2 | 1 | 1 | 3 | 2 | 3 | 3 | 2 | 3 | 6 | 4 | 8 | 4 | 6 | 6 | 4 | 5 | 4 | 8 | 8 | 4 | 6 | 3 | 5 | 7 | 3 | 11 | 7 | 8 | 8 | 8 | 9 | 8 | 8 | 6 | 6 | 41 | 8 | 7 | 6 | 4 |
| MTHFD2L | chr4:74261112-74261209 | 7 | 6 | 3 | 3 | 3 | 2 | 0 | 12 | 5 | 3 | 3 | 4 | 6 | 4 | 2 | 3 | 3 | 2 | 2 | 4 | 3 | 0 | 0 | 4 | 5 | 3 | 6 | 3 | -1 | 3 | 2 | 7 | 8 | 6 | 5 | 4 | 7 | 9 | 11 | 4 | 8 | 10 | 9 | 5 | 13 | 6 | 54 | 6 | 6 | 8 | 3 |
| TEX21P | chr14:64358769-64358859 | 5 | 7 | 0 | 4 | 2 | 4 | 0 | 2 | -1 | -1 | 15 | 10 | 10 | -1 | 6 | -1 | 3 | 3 | 3 | -1 | 0 | 0 | 0 | 4 | 4 | 5 | 6 | 3 | -1 | 8 | 5 | 0 | 2 | 4 | 10 | 5 | 5 | 1 | 3 | 3 | 2 | 2 | 3 | 4 | 6 | 5 | 42 | 3 | 2 | 1 | 2 |
| PIK3R6 | chr17:8832017-8832201 | 7 | 4 | 4 | 3 | 6 | 3 | -1 | 9 | 4 | 1 | 3 | 7 | 2 | 4 | 2 | 2 | 1 | 3 | 2 | 2 | 3 | -1 | -1 | 5 | 6 | 8 | 6 | 6 | 0 | 7 | 7 | 11 | 8 | 6 | 5 | 9 | 6 | 2 | 7 | 4 | 6 | 11 | 10 | 7 | 5 | 7 | 50 | 5 | 8 | 3 | 6 |
| LAMP5 | chr20:9515055-9515158 | 3 | 1 | 1 | 4 | 5 | 4 | -1 | 1 | 3 | 2 | 3 | 2 | 5 | 2 | 2 | 2 | 2 | 3 | 2 | 2 | 2 | 0 | 0 | 1 | 3 | 3 | 5 | 2 | 2 | 0 | 3 | 1 | -1 | 4 | 3 | 2 | 1 | 1 | 1 | 2 | 2 | 1 | 1 | 2 | 5 | 3 | 27 | 2 | 2 | 1 | 6 |
| PCIF1 | chr20:45944245-45944320\|45944537 | 4 | 5 | 5 | 3 | 4 | 3 | 7 | 5 | 5 | 5 | 6 | 7 | 8 | 6 | 5 | 5 | 5 | 7 | 7 | 3 | 5 | 3 | 5 | 5 | 7 | 4 | 6 | 4 | 5 | 5 | 4 | 6 | 8 | 3 | 6 | 3 | 5 | 5 | 7 | 7 | 7 | 4 | 5 | 8 | 8 | 6 | 27 | 6 | 4 | 4 | 4 |
| USH1C | chr11:17496499-17496709 | 5 | 5 | 1 | 2 | 0 | 3 | 1 | 2 | 3 | 2 | 1 | 5 | 3 | 3 | 4 | 4 | 3 | 2 | 2 | 4 | 3 | -1 | 0 | 2 | 1 | 2 | 1 | 3 | 0 | 5 | 5 | 1 | 1 | 3 | -1 | 10 | 0 | 4 | 1 | 3 | 5 | 1 | 3 | 1 | 0 | 1 | 34 | 1 | 1 | 2 | 0 |
| AP1AR | chr4:112258333-112258479 | 3 | 2 | 2 | 3 | 3 | 4 | 4 | 3 | 3 | 4 | 4 | 9 | 10 | 4 | 5 | 5 | 3 | 5 | 4 | 5 | 4 | 4 | 4 | 3 | 3 | 3 | 3 | 4 | 2 | 3 | 3 | 2 | 3 | 2 | 4 | 3 | 4 | 5 | 4 | 5 | 4 | 3 | 3 | 2 | 2 | 3 | 22 | 3 | 4 | 4 | 3 |
| RAVER2 | chr1:64825254-64825408\|64825391 | 4 | 4 | 2 | 3 | 3 | 4 | 4 | 7 | 5 | 5 | 5 | 9 | 9 | 8 | 6 | 6 | 5 | 5 | 5 | 4 | 5 | 3 | 5 | 4 | 4 | 3 | 4 | 3 | 5 | 4 | 3 | 5 | 8 | 6 | 4 | 6 | 7 | 7 | 7 | 4 | 5 | 5 | 7 | 5 | 3 | 4 | 27 | 5 | 5 | 5 | 5 |
| RAB38 | chr11:88173412-88174345 | 2 | 3 | 1 | 3 | 3 | 5 | 2 | 3 | -1 | -1 | 4 | -1 | -1 | -1 | -1 | -1 | 5 | -1 | 0 | 3 | 2 | 2 | 0 | 4 | 3 | 3 | 2 | 3 | 4 | 3 | 1 | 4 | 5 | 3 | 2 | 2 | 4 | 5 | 2 | 6 | 2 | 2 | 1 | 3 | 6 | 4 | 25 | 5 | 5 | 3 | 3 |
| CD200 | chr3:112347349-112347423 | 5 | 5 | 4 | 4 | 3 | 4 | 5 | 5 | 1 | 1 | 0 | 0 | 0 | 0 | 0 | 1 | 0 | 0 | 1 | 1 | 0 | 2 | 6 | 3 | 5 | 4 | 4 | 3 | 3 | 4 | 3 | 3 | 3 | 4 | 6 | 4 | 3 | 3 | 6 | 2 | 3 | 7 | 6 | 8 | 12 | 4 | 33 | 5 | 4 | 3 | 6 |
| ARID2 | chr12:45904342-45904398 | 7 | 4 | 1 | 4 | 3 | 5 | 2 | 5 | 1 | 0 | 1 | 1 | 2 | 1 | 1 | 1 | 2 | 1 | 1 | 1 | 2 | 1 | 0 | 2 | 4 | 2 | 2 | 2 | 1 | 2 | 2 | 1 | 2 | 4 | 3 | 6 | 5 | 1 | 2 | 0 | 2 | 5 | 5 | 9 | 3 | 3 | 31 | 1 | 1 | 1 | 3 |
| VWA3A | chr16:22147571-22147642 | 2 | 2 | -1 | 1 | 0 | 2 | 0 | 3 | 1 | 1 | 1 | 0 | 0 | 1 | 0 | 1 | 1 | 1 | 1 | 1 | 2 | 0 | 0 | 4 | 2 | 2 | 1 | 1 | 1 | 5 | 6 | 4 | -1 | 2 | 0 | 3 | 1 | 2 | -1 | 1 | 3 | 1 | 0 | 3 | 1 | 0 | 24 | 1 | 2 | 0 | 0 |
| SUSD4 | chr1:223289123-223289277 | 4 | 3 | 3 | 1 | 0 | 0 | 4 | 3 | 1 | 0 | 0 | 0 | 0 | 0 | 0 | 1 | 0 | 1 | 1 | 1 | 0 | 3 | -1 | 1 | 2 | 1 | 4 | 2 | -1 | 0 | 0 | 2 | 3 | 3 | 3 | 0 | 2 | 1 | 4 | 2 | 4 | 3 | 3 | 5 | 2 | 2 | 23 | 2 | 1 | 3 | 3 |
| SLC24A3 | chr20:19697989-19698076\|19698061 | 0 | 1 | 1 | 0 | 1 | 1 | 7 | 0 | 0 | 0 | 0 | 0 | 0 | 0 | 0 | 0 | 0 | 0 | 0 | 0 | 0 | 0 | 1 | 4 | 3 | 3 | 1 | 4 | 1 | 0 | 1 | 1 | 0 | 0 | 0 | 0 | 0 | 0 | 0 | 0 | 4 | 0 | 0 | 1 | 1 | 1 | 22 | 0 | 0 | 1 | 1 |
| SSR1 | chr6:7287862-7287922 | 5 | 12 | 6 | 4 | 6 | 4 | -1 | 6 | 4 | 3 | 6 | 3 | 3 | 4 | 3 | 6 | 3 | 5 | 5 | 5 | 6 | -1 | -1 | 6 | 7 | 4 | 7 | 6 | 0 | 6 | 6 | 1 | 13 | 6 | 7 | 9 | 6 | 4 | 11 | 5 | 5 | 4 | 3 | 5 | 10 | 9 | 53 | 7 | 4 | 5 | 11 |
| PWWP2B | chr10:132409762-132409879 | 4 | 9 | 12 | 6 | 7 | 6 | 8 | 6 | 6 | 6 | 7 | 8 | 10 | 6 | 6 | 5 | 8 | 7 | 6 | 9 | 7 | 7 | 9 | 8 | 9 | 6 | 8 | 7 | 13 | 8 | 8 | 10 | 8 | 10 | 8 | 5 | 6 | 8 | 10 | 10 | 10 | 4 | 3 | 8 | 11 | 10 | 40 | 13 | 7 | 8 | 8 |
| KIAA1462 | chr10:30034920-30039905\|30044823 | 4 | 3 | 3 | 2 | 3 | 2 | 1 | 4 | 2 | 3 | 3 | 2 | 3 | 3 | 3 | 3 | 3 | 2 | 2 | 3 | 4 | 0 | 0 | 2 | 2 | 2 | 2 | 2 | 1 | 2 | 3 | 5 | 5 | 4 | 4 | 3 | 3 | 2 | 4 | 3 | 2 | 4 | 7 | 3 | 7 | 2 | 24 | 3 | 1 | 2 | 3 |
| INTS2 | chr17:61884571-61884662 | 5 | 2 | 2 | 2 | 3 | 2 | 8 | 4 | 3 | 3 | 4 | 4 | 4 | 3 | 2 | 5 | 3 | 5 | 2 | 4 | 1 | -1 | -1 | 2 | 3 | 2 | 2 | 2 | -1 | 3 | 2 | 3 | 3 | 3 | 3 | 4 | 5 | 4 | 5 | 3 | 4 | 5 | 6 | 5 | 5 | 4 | 28 | 3 | 3 | 3 | 3 |
| SLC25A17 | chr22:40797284-40797336 | 4 | 4 | 1 | 2 | 2 | 2 | 5 | 3 | 2 | 1 | 1 | 0 | 1 | 0 | 0 | 1 | 0 | 1 | 1 | 1 | 1 | 3 | 1 | 3 | 3 | 2 | 2 | 2 | 3 | 3 | 3 | 3 | 4 | 4 | 4 | 4 | 3 | 2 | 4 | 1 | 3 | 4 | 3 | 7 | 6 | 4 | 22 | 3 | 2 | 3 | 5 |
| ZNF438 | chr10:30923287-30923415 | 9 | 7 | 6 | 4 | 4 | 4 | 5 | 6 | 5 | 2 | 3 | 3 | 2 | 2 | 5 | 2 | 4 | 3 | 2 | 3 | 2 | 7 | 4 | 2 | 5 | 3 | 6 | 4 | 7 | 6 | 9 | 6 | 6 | 6 | 7 | 5 | 5 | 3 | 3 | 4 | 5 | 5 | 8 | 6 | 7 | 5 | 29 | 5 | 3 | 5 | 4 |
| ADAMTS6 | chr5:65270451-65270673 | 2 | 2 | 0 | 2 | 0 | 3 | 0 | 5 | 0 | 1 | 3 | 0 | 0 | 0 | -1 | 0 | 1 | 0 | -1 | -1 | -1 | 0 | 0 | 1 | 1 | 2 | 2 | 2 | 0 | 2 | 0 | -1 | 0 | 3 | -1 | 1 | 1 | 4 | -1 | 1 | 1 | 2 | 2 | 1 | 1 | 2 | 21 | 4 | 1 | 3 | 0 |
| RSRC1 | chr3:158225657-158225733 | 6 | 4 | 1 | 3 | 3 | 3 | 5 | 6 | 2 | 1 | 1 | 2 | 2 | 1 | 1 | 1 | 1 | 1 | 1 | 2 | 1 | 2 | 2 | 3 | 4 | 3 | 4 | 3 | 2 | 3 | 2 | 2 | 3 | 6 | 4 | 3 | 6 | 4 | 5 | 4 | 5 | 9 | 10 | 6 | 7 | 4 | 32 | 4 | 4 | 6 | 9 |
| RIMBP2 | chr12:130469020-130469169 | 0 | 1 | 1 | 0 | 0 | -1 | 0 | 3 | 6 | 3 | 5 | 15 | 11 | 3 | 2 | 5 | 6 | 3 | 2 | 13 | 6 | 0 | 0 | 1 | 1 | 2 | 0 | 2 | 0 | 2 | -1 | 3 | 0 | 1 | 0 | 0 | 7 | 3 | 0 | 0 | 1 | 0 | 0 | 0 | 2 | 1 | 31 | 1 | 0 | 1 | 0 |
| STARD3NL | chr7:38179090-38179145 | 9 | 10 | 7 | 7 | 9 | 7 | 7 | 10 | 12 | 12 | 11 | 10 | 9 | 12 | 11 | 11 | 12 | 10 | 11 | 9 | 11 | 10 | 9 | 8 | 9 | 7 | 7 | 6 | 10 | 8 | 9 | 9 | 7 | 13 | 6 | 11 | 12 | 11 | 8 | 9 | 9 | 8 | 6 | 12 | 14 | 9 | 35 | 7 | 10 | 10 | 7 |
| WDFY3-AS2 | chr4:85005474-85005568 | 2 | 2 | 2 | 1 | 2 | 1 | -1 | 2 | 0 | 1 | 1 | 5 | 3 | 1 | 3 | 1 | 2 | 2 | 2 | 1 | 1 | 0 | -1 | 1 | 3 | 2 | 1 | 1 | 0 | 1 | 2 | 7 | 5 | 2 | 3 | 1 | 1 | 2 | 4 | 2 | 3 | 2 | 1 | 2 | 0 | 3 | 23 | 3 | 0 | 1 | 0 |
| CSGALNACT1 | chr8:19409238-19409424 | 3 | 4 | 2 | 2 | 3 | 2 | 1 | 3 | 6 | 5 | 5 | 5 | 4 | 5 | 5 | 6 | 5 | 6 | 4 | 4 | 4 | 5 | 4 | 4 | 3 | 3 | 3 | 2 | 2 | 3 | 3 | 3 | 6 | 3 | 2 | 3 | 5 | 6 | 4 | 4 | 6 | 3 | 3 | 2 | 3 | 3 | 20 | 3 | 4 | 5 | 2 |
| MBD1 | chr18:50275370-50275444 | 0 | 0 | 1 | 0 | 0 | 0 | 1 | 0 | 21 | 40 | 20 | 37 | 36 | 43 | 47 | 25 | 27 | 26 | 20 | 7 | 15 | 0 | 0 | 0 | 0 | 0 | 1 | 0 | 0 | 9 | 9 | 1 | 1 | 0 | 1 | 5 | 0 | 0 | 0 | 2 | 1 | 1 | 1 | 0 | 0 | 1 | 82 | 1 | 0 | 0 | 0 |
| KIAA1328 | chr18:37063521-37063633 | 2 | 3 | 3 | 1 | 1 | 1 | 4 | 3 | 1 | 4 | 3 | 4 | 3 | 2 | 3 | 1 | 1 | 4 | 2 | 2 | 1 | 1 | 0 | 3 | 3 | 2 | 3 | 2 | -1 | 2 | 2 | 4 | 8 | 4 | 2 | 1 | 2 | 2 | 5 | 6 | 3 | 2 | 2 | 4 | 6 | 3 | 26 | 4 | 1 | 2 | 2 |
| DIS3L2 | chr2:232198527-232198776 | 9 | 8 | 6 | 6 | 8 | 7 | 10 | 11 | 7 | 5 | 7 | 8 | 9 | 6 | 4 | 6 | 7 | 7 | 5 | 6 | 5 | 7 | 8 | 7 | 8 | 6 | 9 | 6 | 11 | 8 | 5 | 7 | 11 | 10 | 10 | 4 | 9 | 11 | 11 | 13 | 13 | 11 | 12 | 12 | 10 | 9 | 41 | 11 | 8 | 9 | 12 |
| IKBKE | chr1:206494786-206494969 | 4 | 3 | 3 | 3 | 3 | 4 | 3 | 3 | 3 | 3 | 4 | 7 | 9 | 3 | 3 | 4 | 4 | 4 | 3 | 2 | 3 | 4 | -1 | 2 | 2 | 3 | 2 | 3 | 3 | 3 | 2 | 2 | 2 | 4 | 3 | 4 | 4 | 2 | 3 | 4 | 2 | 3 | 3 | 2 | 2 | 2 | 21 | 3 | 3 | 2 | 3 |
| CCDC150 | chr2:196665567-196666848 | 10 | 10 | 8 | 8 | 6 | 11 | -1 | 8 | 23 | -1 | 11 | 8 | -1 | -1 | 17 | 22 | 15 | -1 | -1 | 35 | 17 | 0 | 0 | 6 | 9 | 9 | 5 | 6 | -1 | 5 | 9 | -1 | 5 | 10 | 5 | 10 | 13 | 9 | 10 | 16 | 15 | 8 | 7 | 6 | 7 | 12 | 65 | 6 | 13 | 9 | 6 |
| DDX60L | chr4:168457385-168457432 | 2 | 4 | 2 | 3 | 3 | 3 | 0 | 3 | 5 | 1 | 1 | 0 | 3 | 7 | 1 | 4 | 4 | 5 | 1 | 2 | 0 | 0 | 0 | 3 | 3 | 2 | 3 | 1 | 6 | 3 | 2 | 2 | 2 | 6 | 3 | 4 | 3 | 2 | 2 | 5 | 2 | 3 | 1 | 6 | 5 | 3 | 23 | 2 | 2 | 3 | 4 |
| GPX2 | chr14:64940117-64941526\|64941521 | 2 | 1 | 2 | 0 | 2 | 3 | 1 | 1 | 0 | -1 | 2 | 23 | 19 | 1 | -1 | -1 | 0 | 1 | 1 | 1 | -1 | 1 | 0 | 1 | 1 | 1 | 1 | 0 | -1 | 3 | 0 | 1 | 1 | 1 | 1 | 2 | 1 | 0 | 1 | 1 | 1 | 1 | 1 | 1 | 1 | 1 | 41 | 1 | 0 | 1 | 1 |
| INPP5K | chr17:1515888\|1515427\|1515569-1516118 | 5 | 5 | 3 | 2 | 3 | 2 | 3 | 5 | 7 | 5 | 6 | 11 | 10 | 5 | 5 | 6 | 6 | 5 | 5 | 6 | 6 | 4 | 2 | 4 | 5 | 3 | 5 | 3 | 5 | 9 | 11 | 7 | 6 | 7 | 5 | 12 | 5 | 4 | 5 | 6 | 5 | 5 | 5 | 6 | 5 | 4 | 35 | 5 | 4 | 4 | 7 |
| EEA1 | chr12:92821053-92821116 | 6 | 6 | 4 | 5 | 5 | 4 | 2 | 7 | 3 | 4 | 2 | 4 | 9 | 2 | 2 | 4 | 5 | 4 | 2 | 2 | 3 | 7 | -1 | 6 | 6 | 6 | 5 | 5 | -1 | 4 | 4 | 5 | 8 | 4 | 7 | 3 | 9 | 6 | 9 | 8 | 8 | 8 | 8 | 7 | 9 | 6 | 37 | 6 | 6 | 8 | 6 |
| WBP1L | chr10:102825045-102825097 | 11 | 7 | 5 | 5 | 5 | 7 | -1 | 9 | 0 | 0 | 0 | 1 | 0 | 0 | 0 | 0 | 0 | 0 | 0 | 1 | 0 | -1 | 0 | 5 | 5 | 4 | 3 | 4 | 2 | 2 | 2 | 6 | 6 | 4 | 4 | 4 | 4 | 3 | 9 | 1 | 5 | 4 | 5 | 10 | 10 | 5 | 42 | 2 | 4 | 3 | 4 |
| BBIP1 | chr10:110907680-110907836 | 5 | 6 | 3 | 4 | 4 | 4 | 4 | 5 | 2 | 1 | 2 | 3 | 3 | 2 | 2 | 2 | 2 | 2 | 2 | 2 | 2 | 7 | 9 | 5 | 5 | 4 | 5 | 3 | 7 | 3 | 2 | 4 | 5 | 6 | 6 | 2 | 5 | 5 | 6 | 4 | 6 | 7 | 7 | 7 | 7 | 5 | 30 | 6 | 5 | 5 | 12 |
| ZNF628 | chr19:55479834-55479917 | 6 | 10 | 7 | 7 | 6 | 6 | 4 | 7 | 10 | 13 | 17 | 10 | 8 | 12 | 17 | 15 | 10 | 17 | 12 | 9 | 14 | -1 | -1 | 8 | 7 | 6 | 7 | 7 | 7 | 14 | 10 | 12 | 7 | 8 | 7 | 11 | 10 | 7 | 10 | 9 | 7 | 5 | 6 | 7 | 6 | 8 | 44 | 9 | 8 | 7 | 5 |
| VWA3A | chr16:22141203-22141249 | 11 | 4 | 8 | 2 | 4 | 7 | 0 | 7 | 13 | 12 | 12 | 6 | 4 | 9 | 11 | 15 | 13 | 13 | 11 | 14 | 9 | 0 | 9 | 6 | 7 | 6 | 6 | 5 | 17 | 1 | 6 | 5 | 0 | 14 | -1 | 8 | 10 | 11 | -1 | 4 | 10 | 3 | 2 | 4 | 6 | 4 | 59 | 8 | 10 | 3 | 7 |
| EPSTI1 | chr13:42919300-42919332 | 18 | 17 | 11 | 14 | 16 | 14 | 14 | 20 | 24 | 23 | 19 | 20 | 27 | 24 | 20 | 23 | 21 | 20 | 20 | 18 | 20 | 13 | 9 | 16 | 15 | 15 | 13 | 14 | 13 | 18 | 22 | 11 | 15 | 16 | 13 | 22 | 17 | 12 | 14 | 17 | 16 | 17 | 16 | 14 | 14 | 14 | 60 | 15 | 15 | 14 | 16 |
| VRK2 | chr2:58058352\|58057862-58058488 | 8 | 6 | 5 | 5 | 7 | 7 | 6 | 8 | 10 | 10 | 10 | 9 | 6 | 7 | 8 | 9 | 11 | 10 | 7 | 10 | 10 | 7 | 4 | 6 | 5 | 5 | 3 | 6 | 8 | 8 | 7 | 9 | 3 | 7 | 6 | 8 | 8 | 13 | 5 | 13 | 9 | 6 | 6 | 6 | 5 | 6 | 34 | 5 | 6 | 5 | 3 |
| EIF4B | chr12:53022254-53022366 | 3 | 3 | 1 | 3 | 4 | 2 | 4 | 5 | 2 | 2 | 2 | 1 | 4 | 1 | 0 | 2 | 2 | 2 | 3 | 1 | 0 | 4 | 0 | 5 | 4 | 4 | 3 | 5 | 2 | 4 | 1 | 3 | 3 | 2 | 4 | 2 | 6 | 5 | 3 | 4 | 5 | 5 | 2 | 5 | 4 | 5 | 22 | 1 | 5 | 4 | 2 |
| PI4KB | chr1:151293400-151293462 | 11 | 5 | 2 | 10 | 7 | 10 | 6 | 9 | 4 | 2 | 3 | 3 | 2 | 2 | 2 | 3 | 2 | 3 | 3 | 3 | 3 | 9 | 9 | 5 | 5 | 5 | 3 | 5 | 9 | 4 | 4 | 7 | 4 | 7 | 8 | 3 | 13 | 7 | 7 | 3 | 5 | 10 | 13 | 7 | 5 | 6 | 43 | 9 | 10 | 6 | 9 |
| AGO4 | chr1:35825605-35825678 | 5 | 5 | 3 | 3 | 3 | 2 | 4 | 6 | 2 | 2 | 2 | 1 | 1 | 1 | 0 | 2 | 3 | 1 | 1 | 3 | 1 | 4 | 4 | 4 | 4 | 3 | 3 | 3 | 7 | 2 | 2 | 4 | 6 | 7 | 3 | 5 | 5 | 2 | 5 | 2 | 3 | 6 | 5 | 8 | 12 | 5 | 34 | 3 | 3 | 3 | 11 |
| SNCAIP | chr5:122452925-122452995 | 1 | 1 | 2 | 0 | 0 | 0 | -1 | 1 | 50 | 43 | 29 | 35 | 36 | 35 | 37 | 48 | 47 | 26 | 24 | 29 | 40 | 0 | 0 | 0 | 1 | 1 | 2 | 1 | 6 | 0 | 1 | 0 | 2 | 8 | 2 | 1 | 0 | 0 | 3 | 8 | 1 | 5 | 4 | 1 | 4 | 1 | 81 | 1 | 0 | 0 | 10 |
| PPP1R16A | chr8:144478586-144489164\|144478682 | 8 | 5 | 4 | 6 | 6 | 8 | 8 | 7 | 8 | 9 | 9 | 12 | 11 | 7 | 8 | 6 | 9 | 10 | 8 | 8 | 7 | 5 | 4 | 10 | 23 | 5 | 6 | 5 | 8 | 4 | 3 | 9 | 7 | 7 | 9 | 4 | 10 | 6 | 8 | 5 | 6 | 9 | 10 | 17 | 10 | 8 | 54 | 6 | 4 | 6 | 5 |
| GALNT8 | chr12:4761402-4763080 | 3 | 2 | 1 | 4 | 2 | 2 | 0 | 5 | 3 | 2 | 1 | 4 | 1 | 1 | 2 | 1 | 1 | 1 | 2 | 2 | 3 | -1 | 0 | 2 | 3 | 2 | 3 | 2 | 0 | 0 | 1 | -1 | -1 | 5 | 4 | 7 | 3 | 0 | 2 | 3 | 1 | 2 | 2 | 4 | 5 | 6 | 27 | 2 | 4 | 3 | 4 |
| NKD1 | chr16:50616049\|50617753-50617805 | 7 | 4 | 2 | 4 | 4 | 6 | 5 | 5 | 6 | 4 | 4 | 3 | 2 | 4 | 4 | 5 | 5 | 4 | 3 | 6 | 4 | 2 | 1 | 4 | 4 | 3 | 3 | 3 | 6 | 4 | 3 | 5 | 5 | 5 | 3 | 5 | 7 | 3 | 6 | 4 | 2 | 7 | 8 | 5 | 3 | 4 | 24 | 2 | 4 | 2 | 2 |
| ZNF512 | chr2:27619650\|27619105-27619816 | 3 | 5 | 3 | 6 | 5 | 3 | 2 | 3 | 1 | 1 | 1 | 4 | 5 | 2 | 1 | 2 | 2 | 1 | 1 | 2 | 1 | 4 | 3 | 3 | 5 | 3 | 6 | 3 | 2 | 3 | 3 | 6 | 8 | 5 | 6 | 2 | 4 | 2 | 7 | 4 | 5 | 4 | 4 | 7 | 9 | 5 | 25 | 5 | 4 | 4 | 6 |
| BAALC-AS1 | chr8:103267242-103268843 | 8 | 11 | 11 | 7 | 8 | 8 | 7 | 11 | 13 | 12 | 15 | 16 | 16 | 16 | 14 | 12 | 9 | 14 | 10 | 10 | 12 | -1 | -1 | 10 | 12 | 9 | 10 | 10 | -1 | 16 | 14 | 5 | -1 | 14 | 12 | 14 | 8 | 9 | 10 | 11 | 13 | 10 | 9 | 14 | 11 | 9 | 67 | 12 | 5 | 5 | -1 |
| DZANK1 | chr20:18394675\|18394462-18394858 | 9 | 9 | 8 | 4 | 6 | 5 | 12 | 8 | 5 | 4 | 5 | 12 | 9 | 5 | 4 | 6 | 7 | 3 | 5 | 3 | 4 | -1 | -1 | 7 | 10 | 9 | 9 | 7 | 10 | 6 | 10 | 6 | 9 | 12 | 8 | 8 | 9 | 11 | 11 | 6 | 8 | 9 | 10 | 8 | 8 | 8 | 43 | 7 | 11 | 11 | 7 |
| MBD5 | chr2:148505761-148505830 | 7 | 7 | 5 | 3 | 3 | 4 | 3 | 7 | 3 | 3 | 2 | 2 | 2 | 2 | 2 | 3 | 2 | 2 | 2 | 3 | 3 | 3 | 5 | 3 | 5 | 4 | 5 | 3 | 3 | 4 | 3 | 5 | 8 | 5 | 8 | 5 | 4 | 3 | 6 | 4 | 4 | 6 | 8 | 8 | 8 | 6 | 29 | 4 | 3 | 4 | 9 |
| ZNHIT6 | chr1:85679373-85679448 | 5 | 3 | 2 | 3 | 4 | 6 | 5 | 7 | 1 | 1 | 1 | 3 | 2 | 1 | 1 | 1 | 1 | 1 | 1 | 2 | 1 | 10 | 4 | 4 | 4 | 4 | 3 | 3 | 7 | 3 | 3 | 4 | 3 | 4 | 5 | 3 | 6 | 4 | 4 | 3 | 5 | 5 | 5 | 6 | 7 | 5 | 24 | 4 | 4 | 4 | 5 |
| TRIM39-RPP21 | chr6:30340272-30340361 | 7 | 6 | 4 | 4 | 5 | 5 | 8 | 7 | 2 | 2 | 2 | 1 | 1 | 2 | 2 | 3 | 3 | 2 | 3 | 3 | 2 | 3 | 3 | 7 | 7 | 6 | 6 | 7 | 5 | 4 | 5 | 5 | 11 | 9 | 5 | 3 | 5 | 4 | 7 | 3 | 6 | 7 | 6 | 11 | 15 | 8 | 40 | 4 | 4 | 7 | 12 |
| USP25 | chr21:15822288-15822411 | 7 | 7 | 5 | 6 | 7 | 6 | 5 | 9 | 9 | 9 | 10 | 14 | 19 | 11 | 8 | 9 | 11 | 10 | 11 | 8 | 9 | 18 | 12 | 7 | 9 | 6 | 6 | 5 | 11 | 7 | 5 | 9 | 8 | 8 | 7 | 3 | 10 | 14 | 9 | 15 | 11 | 8 | 7 | 9 | 8 | 8 | 51 | 10 | 10 | 10 | 7 |
| TMEM222 | chr1:27334667-27334754 | 5 | 6 | 4 | 8 | 7 | 8 | 6 | 5 | 3 | 3 | 3 | 5 | 5 | 3 | 3 | 3 | 3 | 3 | 3 | 3 | 2 | 6 | 6 | 9 | 6 | 7 | 4 | 7 | 5 | 4 | 4 | 6 | 5 | 5 | 4 | 3 | 4 | 5 | 5 | 5 | 4 | 4 | 5 | 6 | 6 | 6 | 21 | 5 | 7 | 5 | 4 |
| RP11-849H4.2 | chr11:71926418-71926510 | 5 | 2 | 1 | 2 | 1 | 1 | -1 | 2 | 1 | 1 | 2 | 2 | 1 | 0 | 0 | 0 | 3 | 2 | 1 | 4 | 2 | 0 | 0 | 2 | 4 | 1 | 3 | 1 | 0 | 2 | 1 | 5 | 3 | 4 | 1 | 4 | 3 | 2 | 4 | 0 | 2 | 4 | 3 | 5 | 7 | 2 | 25 | 2 | 2 | 1 | 6 |
| BPNT1 | chr1:220066073-220066117 | 3 | 4 | 1 | 2 | 3 | 3 | 1 | 3 | 4 | 5 | 3 | 8 | 8 | 6 | 6 | 3 | 4 | 4 | 3 | 3 | 2 | 2 | 0 | 3 | 3 | 2 | 3 | 3 | 3 | 6 | 7 | 2 | 2 | 2 | 2 | 10 | 2 | 2 | 2 | 1 | 2 | 3 | 3 | 3 | 3 | 4 | 26 | 2 | 2 | 2 | 2 |
| PHF3 | chr6:63678828-63679020 | 1 | 3 | 3 | 2 | 4 | 3 | 1 | 3 | 3 | 4 | 4 | 11 | 12 | 6 | 8 | 3 | 4 | 4 | 2 | 2 | 3 | 1 | 3 | 4 | 5 | 2 | 4 | 3 | 7 | 2 | 1 | 4 | 4 | 3 | 6 | 1 | 2 | 4 | 10 | 14 | 7 | 3 | 2 | 5 | 3 | 4 | 40 | 6 | 4 | 4 | 3 |
| MARCH1 | chr4:163612271-163613038 | 6 | 7 | 8 | 7 | 7 | 6 | -1 | 7 | 5 | 3 | 4 | 5 | 8 | 3 | 2 | 5 | 3 | 4 | 5 | 9 | 5 | -1 | 0 | 6 | 10 | 7 | 9 | 8 | -1 | 7 | 5 | 6 | 6 | 7 | 4 | 9 | 8 | 7 | 5 | 3 | 11 | 8 | 7 | 12 | 7 | 6 | 43 | 10 | 8 | 4 | 5 |
| SEPT7-AS1 | chr7:35763765-35763909 | 4 | 5 | 5 | 5 | 3 | 5 | 0 | 4 | 6 | 4 | 5 | 12 | 10 | 7 | 7 | 6 | 5 | 4 | 6 | 6 | 6 | -1 | -1 | 7 | 5 | 7 | 4 | 5 | -1 | 4 | 7 | 5 | 3 | 4 | 3 | 3 | 5 | 4 | 4 | 4 | 5 | 4 | 4 | 6 | 3 | 6 | 30 | 5 | 4 | 4 | 3 |
| SNX32 | chr11:65835506-65835673 | 4 | 4 | 6 | 4 | 6 | 4 | -1 | 6 | 5 | 4 | 5 | 7 | 7 | 6 | 4 | 6 | 6 | 6 | 5 | 5 | 4 | -1 | -1 | 9 | 5 | 7 | 3 | 8 | -1 | 5 | 13 | -1 | -1 | 6 | 2 | 6 | 8 | 6 | 4 | 7 | 9 | 3 | 4 | 9 | 4 | 6 | 48 | 8 | 9 | 12 | 5 |
| ICE1 | chr5:5453116-5453212 | 11 | 8 | 4 | 7 | 8 | 7 | 7 | 9 | 3 | 3 | 3 | 6 | 7 | 5 | 3 | 4 | 4 | 2 | 3 | 6 | 5 | -1 | -1 | 6 | 6 | 7 | 7 | 6 | 3 | 7 | 6 | 7 | 7 | 7 | 5 | 4 | 13 | 6 | 10 | 6 | 8 | 12 | 11 | 5 | 8 | 6 | 40 | 9 | 7 | 9 | 6 |
| PCLO | chr7:82840605-82840755 | 2 | 7 | 2 | 4 | 3 | 3 | -1 | 3 | 0 | 0 | 0 | 1 | 2 | 0 | 0 | 0 | 0 | 0 | 0 | 0 | 0 | -1 | 0 | 4 | 5 | 4 | 4 | 4 | -1 | 1 | 1 | 1 | -1 | 6 | 2 | 2 | 6 | 9 | 3 | 3 | 5 | 3 | 2 | 3 | 6 | 5 | 32 | 3 | 8 | 4 | 0 |
| FAM179B | chr14:45046901-45047024 | 10 | 8 | 7 | 7 | 9 | 12 | 8 | 12 | 5 | 4 | 5 | 10 | 9 | 5 | 3 | 5 | 5 | 4 | 6 | 8 | 6 | 12 | 10 | 11 | 11 | 8 | 9 | 8 | 12 | 7 | 5 | 6 | 9 | 10 | 11 | 7 | 11 | 10 | 10 | 8 | 12 | 11 | 11 | 12 | 13 | 11 | 35 | 11 | 11 | 10 | 15 |
| TBC1D32 | chr6:121139774-121139836 | 8 | 5 | 5 | 4 | 5 | 6 | -1 | 8 | 2 | 5 | 2 | 5 | 6 | 2 | 4 | 3 | 2 | 3 | 5 | 3 | 3 | -1 | -1 | 8 | 10 | 8 | 10 | 8 | -1 | 4 | 6 | 6 | 5 | 7 | 8 | 6 | 8 | 2 | 7 | 7 | 10 | 7 | 10 | 13 | 13 | 7 | 50 | 5 | 4 | 8 | 16 |
| RYR3 | chr15:33855884-33856051 | 4 | 6 | 5 | 7 | 6 | 5 | 5 | 5 | 2 | 2 | 1 | 2 | 2 | 2 | 2 | 1 | 1 | 1 | 1 | 3 | 2 | 11 | 11 | 4 | 4 | 4 | 4 | 3 | 7 | 7 | 13 | 3 | -1 | 5 | 4 | 0 | 3 | 5 | 10 | 6 | 4 | 5 | 4 | 5 | 6 | 4 | 37 | 6 | 6 | 6 | 1 |
| ACSS3 | chr12:81146302-81150017 | 4 | 2 | 1 | 5 | 4 | 4 | 6 | 4 | 2 | 2 | 2 | 2 | 3 | 2 | 1 | 2 | 3 | 2 | 2 | 3 | 2 | 7 | 5 | 6 | 5 | 4 | 5 | 6 | 3 | 7 | 6 | 3 | 3 | 5 | 4 | 9 | 12 | 5 | 6 | 7 | 6 | 6 | 7 | 5 | 4 | 4 | 28 | 5 | 7 | 7 | 2 |
| TOR1AIP2 | chr1:179848225-179848388 | 2 | 3 | 1 | 2 | 2 | 1 | 2 | 2 | 2 | 6 | 5 | 9 | 10 | 5 | 6 | 3 | 3 | 7 | 5 | 1 | 3 | -1 | -1 | 2 | 1 | 3 | 1 | 1 | -1 | 2 | 2 | 0 | 2 | 3 | 2 | 9 | 3 | 2 | 3 | 2 | 2 | 4 | 3 | 2 | 3 | 2 | 27 | 1 | 2 | 2 | 2 |
| PGAP1 | chr2:196922103-196922186 | 4 | 13 | 12 | 3 | 5 | 3 | 5 | 4 | 7 | 3 | 7 | 4 | 4 | 5 | 5 | 5 | 5 | 7 | 2 | 4 | 2 | 0 | 0 | 8 | 6 | 4 | 4 | 4 | 0 | 3 | 3 | -1 | 7 | 3 | 5 | 1 | 4 | 7 | 3 | 7 | 5 | 4 | 3 | 6 | 7 | 7 | 39 | 5 | 1 | 4 | 1 |
| RNF170 | chr8:42864617-42864693 | 5 | 5 | 1 | 8 | 6 | 8 | 9 | 5 | 5 | 2 | 2 | 7 | 9 | 3 | 2 | 3 | 2 | 3 | 2 | 3 | 2 | 5 | 8 | 7 | 3 | 8 | 3 | 7 | 6 | 7 | 7 | 2 | 3 | 5 | 3 | 9 | 6 | 6 | 3 | 4 | 6 | 3 | 3 | 5 | 5 | 5 | 30 | 3 | 7 | 4 | 3 |
| RP11-274B21.1 | chr7:128562315-128562460 | 8 | 7 | 7 | 4 | 5 | 5 | 10 | 7 | 5 | 6 | 6 | 6 | 6 | 4 | 6 | 6 | 6 | 6 | 7 | 8 | 4 | 11 | -1 | 5 | 5 | 5 | 8 | 5 | 6 | 6 | 5 | 7 | 6 | 6 | 10 | 5 | 5 | 6 | 10 | 7 | 8 | 9 | 10 | 7 | 7 | 8 | 32 | 6 | 4 | 6 | 7 |
| FBXO24 | chr7:100600031-100600201 | 16 | 15 | 10 | 22 | 17 | 22 | -1 | 21 | 14 | 11 | 21 | 17 | 17 | 15 | 15 | 13 | 15 | 22 | 20 | 13 | 18 | -1 | -1 | 15 | 14 | 24 | 11 | 24 | -1 | 15 | 14 | 15 | 8 | 14 | 12 | 17 | 19 | 20 | 16 | 14 | 17 | 11 | 14 | 19 | 13 | 16 | 89 | 16 | 13 | 18 | 13 |
| RAB18 | chr10:27531529-27531615 | 5 | 2 | 0 | 2 | 2 | 3 | 3 | 5 | 1 | 1 | 1 | 1 | 1 | 1 | 1 | 1 | 1 | 1 | 1 | 1 | 1 | 3 | 2 | 2 | 2 | 2 | 3 | 1 | 3 | 3 | 2 | 2 | 2 | 4 | 3 | 7 | 5 | 2 | 3 | 1 | 2 | 7 | 8 | 2 | 2 | 2 | 23 | 2 | 2 | 3 | 4 |
| MAGI2 | chr7:78184428-78184651 | 4 | 1 | 0 | 3 | 1 | 5 | -1 | 7 | 0 | 0 | 0 | 0 | 0 | 0 | 0 | 0 | 0 | 0 | 0 | 0 | 0 | -1 | 0 | 1 | 1 | 0 | 0 | 1 | 0 | 6 | 6 | 0 | -1 | 3 | 0 | 3 | 3 | 6 | 1 | 1 | 3 | 7 | 6 | 0 | 0 | 2 | 29 | 2 | 2 | 1 | 0 |
| AADAT | chr4:170084725-170084810\|170084999 | 3 | 4 | 2 | 2 | 2 | 2 | 3 | 3 | 1 | 1 | 1 | 0 | 1 | 1 | 0 | 1 | 1 | 1 | 1 | 1 | 1 | 0 | 1 | 2 | 3 | 2 | 3 | 1 | 3 | 1 | 1 | 3 | 2 | 5 | 3 | 1 | 3 | 1 | 9 | 2 | 4 | 2 | 2 | 4 | 4 | 4 | 20 | 3 | 2 | 1 | 4 |
| LINC01125 | chr2:97688895-97688993 | 35 | 20 | 11 | 24 | 22 | 31 | 23 | 24 | 14 | 13 | 11 | 14 | 12 | 12 | 14 | 9 | 11 | 12 | 11 | 9 | 9 | -1 | 18 | 25 | 22 | 25 | 16 | 23 | 33 | 20 | 24 | 19 | 14 | 23 | 21 | 22 | 19 | 21 | 21 | 17 | 21 | 25 | 30 | 29 | 25 | 21 | 94 | 17 | 24 | 20 | 28 |
| C9orf85 | chr9:71949462\|71952045-71952132 | 7 | 5 | 5 | 5 | 5 | 5 | 10 | 8 | 7 | 4 | 6 | 6 | 6 | 7 | 4 | 7 | 5 | 6 | 5 | 6 | 8 | 5 | 2 | 7 | 7 | 5 | 4 | 5 | 9 | 4 | 4 | 7 | 7 | 7 | 7 | 4 | 9 | 6 | 10 | 5 | 8 | 7 | 9 | 10 | 9 | 9 | 30 | 6 | 6 | 5 | 5 |
| NUCB2 | chr11:17286634-17286717\|17286713 | 12 | 11 | 10 | 10 | 10 | 14 | 12 | 12 | 20 | 23 | 22 | 18 | 18 | 25 | 27 | 19 | 19 | 24 | 23 | 18 | 20 | 12 | 11 | 15 | 12 | 15 | 15 | 15 | 14 | 12 | 14 | 13 | 12 | 11 | 10 | 16 | 14 | 12 | 11 | 14 | 14 | 13 | 13 | 12 | 9 | 11 | 37 | 11 | 13 | 13 | 7 |
| F7 | chr13:113114468-113115067 | 2 | 4 | 0 | 1 | 0 | -1 | -1 | 6 | 0 | 2 | 2 | 3 | 5 | 1 | 1 | 0 | 3 | 1 | -1 | 0 | -1 | 0 | 0 | 0 | 0 | 0 | 2 | 2 | 0 | 3 | 0 | -1 | 4 | 5 | -1 | -1 | -1 | 1 | 0 | 4 | 2 | 5 | 2 | 6 | 4 | 7 | 32 | 10 | 0 | 2 | 0 |
| SEPT7P2 | chr7:45739380-45739515 | 15 | 13 | 6 | 16 | 19 | 22 | 9 | 13 | 2 | 2 | 2 | 2 | 2 | 3 | 3 | 2 | 2 | 2 | 2 | 2 | 3 | 18 | 6 | 13 | 10 | 11 | 8 | 10 | 12 | 8 | 6 | 8 | 10 | 7 | 8 | 6 | 10 | 6 | 4 | 3 | 8 | 8 | 8 | 10 | 7 | 9 | 59 | 7 | 10 | 12 | 4 |
| MYH7B | chr20:34989324-34989367 | 2 | 0 | 2 | 1 | 3 | 2 | 0 | 1 | 1 | 2 | 0 | 0 | 0 | 0 | 0 | 1 | 1 | 1 | 0 | 0 | 0 | 0 | 0 | 0 | 1 | 0 | 3 | 1 | 0 | 5 | 4 | 1 | 1 | 2 | 7 | 7 | 1 | 0 | 6 | 0 | 5 | 1 | 1 | 2 | 1 | 2 | 24 | 2 | 0 | 0 | 5 |
| SUSD4 | chr1:223296727-223296890 | 6 | 4 | 3 | 4 | 2 | 4 | 7 | 6 | 1 | 1 | 2 | 1 | 2 | 1 | 1 | 1 | 1 | 1 | 2 | 1 | 1 | 10 | 0 | 2 | 5 | 4 | 4 | 4 | 0 | 0 | 1 | 4 | 3 | 5 | 4 | 1 | 3 | 2 | 6 | 4 | 6 | 4 | 4 | 7 | 8 | 3 | 27 | 5 | 3 | 4 | 5 |
| TAC3 | chr12:57012621-57012737 | 10 | 4 | 7 | 4 | 2 | 8 | 1 | 8 | 1 | 1 | 1 | 3 | 2 | 1 | 1 | 2 | 0 | 1 | 1 | 2 | 2 | 0 | 0 | 3 | 4 | 2 | 7 | 6 | 0 | 16 | -1 | -1 | -1 | 6 | -1 | 8 | 9 | 4 | 1 | 6 | 1 | 7 | 6 | 4 | 7 | 2 | 46 | 5 | 7 | 5 | 5 |
| TMEM260 | chr14:56619358-56619451\|56620642 | 17 | 15 | 9 | 18 | 20 | 19 | 10 | 19 | 24 | 21 | 20 | 22 | 23 | 21 | 16 | 22 | 15 | 21 | 24 | 21 | 22 | 21 | 20 | 20 | 17 | 20 | 15 | 18 | 20 | 15 | 13 | 18 | 12 | 14 | 13 | 14 | 22 | 21 | 19 | 21 | 22 | 18 | 17 | 18 | 22 | 16 | 56 | 17 | 20 | 20 | 19 |

### SFig5, pg14

| Gene Symbol | Exon Location (hg38, human) | Adipose - Subcutaneous | Adipose - Visceral (Omentum) | Adrenal Gland | Artery - Aorta | Artery - Coronary | Artery - Tibial | Bladder | Breast - Mammary Tissue | Brain - Amygdala | Brain - Anterior cingulate cortex (BA24) | Brain - Caudate (basal ganglia) | Brain - Cerebellar Hemisphere | Brain - Cerebellum | Brain - Cortex | Brain - Frontal Cortex (BA9) | Brain - Hippocampus | Brain - Hypothalamus | Brain - Nucleus accumbens (basal ganglia) | Brain - Putamen (basal ganglia) | Brain - Spinal cord (cervical c-1) | Brain - Substantia nigra | Cervix - Ectocervix | Cervix - Endocervix | Colon - Sigmoid | Colon - Transverse | Esophagus - Gastroesophageal Junction | Esophagus - Mucosa | Esophagus - Muscularis | Fallopian Tube | Heart - Atrial Appendage | Heart - Left Ventricle | Kidney - Cortex | Liver | Lung | Minor Salivary Gland | Muscle - Skeletal | Nerve - Tibial | Ovary | Pancreas | Pituitary | Prostate | Skin - Not Sun Exposed (Suprapubic) | Skin - Sun Exposed (Lower leg) | Small Intestine - Terminal Ileum | Spleen | Stomach | Testis | Thyroid | Uterus | Vagina | Whole Blood |
|---|---|---|---|---|---|---|---|---|---|---|---|---|---|---|---|---|---|---|---|---|---|---|---|---|---|---|---|---|---|---|---|---|---|---|---|---|---|---|---|---|---|---|---|---|---|---|---|---|---|---|---|---|
| MCUR1 | chr6:13810422\|13810499-13810600 | 9 | 6 | 5 | 6 | 6 | 7 | 7 | 8 | 7 | 6 | 6 | 6 | 7 | 7 | 7 | 7 | 6 | 6 | 7 | 8 | 6 | 6 | 6 | 7 | 9 | 5 | 10 | 5 | 4 | 3 | 3 | 11 | 7 | 8 | 8 | 6 | 7 | 7 | 8 | 7 | 7 | 9 | 10 | 12 | 7 | 7 | 29 | 8 | 7 | 8 | 8 |
| MAGI3 | chr1:113582123-113582233 | 4 | 2 | 2 | 1 | 2 | 2 | 0 | 5 | 2 | 0 | 0 | 2 | 1 | 0 | 1 | 1 | 1 | 1 | 2 | 2 | 0 | 0 | 0 | 2 | 3 | 2 | 3 | 2 | -1 | 2 | 1 | 1 | 1 | 3 | 4 | 1 | 3 | 1 | 6 | 2 | 1 | 5 | 5 | 6 | 2 | 4 | 23 | 2 | 2 | 3 | 7 |
| USMG5 | chr10:103395746\|103394199-103396032 | 7 | 10 | 7 | 8 | 9 | 7 | 10 | 9 | 11 | 13 | 12 | 17 | 22 | 16 | 15 | 13 | 12 | 11 | 12 | 15 | 11 | 13 | 10 | 12 | 12 | 10 | 14 | 11 | 9 | 11 | 10 | 10 | 8 | 10 | 11 | 15 | 9 | 14 | 10 | 11 | 9 | 11 | 10 | 12 | 11 | 11 | 41 | 9 | 13 | 13 | 13 |
| AQP7 | chr9:33401536-33401672 | 16 | 14 | 12 | 13 | 16 | 16 | 21 | 18 | 9 | 23 | 16 | 29 | 30 | 21 | 22 | 16 | 21 | 18 | 12 | 18 | 26 | -1 | -1 | 15 | 14 | 13 | 16 | 16 | -1 | 19 | 21 | 10 | 15 | 15 | 20 | 21 | 17 | 21 | 13 | 21 | 16 | 15 | 16 | 11 | 17 | 18 | 83 | 17 | 19 | 15 | 19 |
| GRIK3 | chr1:36821999-36822575 | 3 | 3 | 1 | 0 | 1 | 5 | 0 | 4 | 2 | 1 | 1 | 1 | 2 | 1 | 0 | 0 | 1 | 1 | 1 | 1 | 3 | 0 | 0 | 4 | 3 | 3 | 3 | 3 | 0 | 4 | 5 | -1 | -1 | 2 | 3 | 3 | 5 | 4 | 2 | 2 | 2 | 5 | 3 | 2 | 0 | 3 | 22 | 2 | -1 | 1 | 0 |
| FAM228A | chr2:24179111-24179297 | 9 | 4 | 4 | 5 | 3 | 7 | 3 | 6 | 3 | 3 | 4 | 4 | 3 | 2 | 3 | 1 | 3 | 3 | 4 | 5 | 3 | 0 | 0 | 2 | 4 | 3 | 7 | 4 | 0 | 4 | 5 | 11 | 8 | 7 | 7 | 7 | 9 | 5 | 7 | 6 | 5 | 11 | 12 | 3 | 6 | 6 | 40 | 6 | 5 | 7 | 14 |
| COL11A1 | chr1:102965161-102965365 | 5 | 8 | 6 | 5 | 2 | 2 | 0 | 4 | 1 | 1 | 1 | 2 | 1 | 1 | 1 | 1 | 1 | 1 | 2 | 1 | 0 | -1 | 2 | 1 | 3 | 0 | 2 | -1 | 11 | -1 | 0 | -1 | 16 | 2 | 4 | 4 | 3 | 5 | 3 | 3 | 2 | 2 | 2 | 4 | 5 | 3 | 42 | 10 | 4 | 1 | -1 |
| SNCAIP | chr5:122443667\|122443571-122444195 | 5 | 7 | 5 | 6 | 6 | 4 | -1 | 5 | 1 | 1 | 3 | 2 | 2 | 3 | 3 | 2 | 2 | 3 | 3 | 4 | 3 | 7 | 4 | 4 | 6 | 3 | 9 | 4 | 3 | 5 | 5 | 6 | 9 | 5 | 8 | 4 | 5 | 4 | 12 | 8 | 7 | 4 | 4 | 9 | 10 | 5 | 32 | 8 | 5 | 5 | 3 |
| CDADC1 | chr13:49269698-49269831 | 5 | 6 | 2 | 3 | 2 | 3 | 7 | 6 | 1 | 1 | 1 | 1 | 1 | 1 | 1 | 2 | 2 | 1 | 2 | 1 | 2 | 1 | 2 | 4 | 5 | 3 | 7 | 3 | 5 | 3 | 2 | 3 | 4 | 5 | 5 | 8 | 6 | 4 | 8 | 4 | 4 | 7 | 7 | 5 | 11 | 5 | 27 | 4 | 4 | 5 | 8 |
| PPP2R2B | chr5:146592049-146592196\|146592193 | 4 | 5 | 2 | 3 | 2 | 3 | 3 | 5 | 1 | 0 | 1 | 1 | 1 | 0 | 0 | 1 | 1 | 1 | 1 | 1 | 1 | -1 | -1 | 2 | 2 | 3 | 3 | 2 | 5 | 1 | 1 | 4 | 8 | 6 | 4 | 3 | 3 | 5 | 5 | 2 | 2 | 4 | 3 | 4 | 9 | 4 | 26 | 4 | 3 | 4 | 7 |
| FAM184A | chr6:118960769-118960863 | 11 | 12 | 13 | 4 | 5 | 4 | 8 | 13 | 16 | 13 | 18 | 3 | 4 | 10 | 9 | 12 | 8 | 16 | 18 | 11 | 10 | 4 | 11 | 5 | 7 | 4 | 10 | 5 | 18 | 6 | 8 | 18 | 10 | 14 | 16 | 8 | 12 | 10 | 20 | 14 | 10 | 13 | 12 | 12 | 32 | 10 | 74 | 20 | 5 | 6 | 17 |
| STKLD1 | chr9:133397164-133398055 | 10 | 14 | 6 | 10 | 6 | 15 | -1 | 12 | 4 | 4 | 7 | 7 | 7 | 10 | 3 | 10 | 4 | 6 | 3 | 4 | 2 | 0 | -1 | 16 | 16 | 16 | 8 | 10 | -1 | 10 | 4 | -1 | 5 | 7 | -1 | 20 | 11 | 10 | 16 | 4 | 11 | 7 | 11 | 11 | 8 | 9 | 67 | 15 | 9 | 8 | 4 |
| KPTN | chr19:47476115-47476231\|47476301 | 6 | 6 | 5 | 4 | 5 | 6 | 11 | 6 | 9 | 9 | 8 | 22 | 22 | 11 | 10 | 10 | 13 | 7 | 6 | 19 | 15 | 4 | 8 | 8 | 6 | 6 | 5 | 7 | 10 | 8 | 10 | 6 | 4 | 6 | 7 | 7 | 8 | 11 | 8 | 11 | 6 | 8 | 9 | 6 | 7 | 6 | 40 | 7 | 8 | 7 | 7 |
| FKBP1B | chr2:24054473-24054540\|24054616 | 9 | 8 | 5 | 9 | 8 | 8 | 6 | 9 | 9 | 8 | 9 | 10 | 10 | 7 | 7 | 8 | 6 | 8 | 10 | 7 | 10 | 7 | 6 | 8 | 7 | 7 | 6 | 8 | 8 | 10 | 7 | 7 | 6 | 9 | 7 | 10 | 9 | 12 | 9 | 6 | 8 | 8 | 9 | 8 | 10 | 7 | 27 | 8 | 11 | 9 | 4 |
| CYP2J2 | chr1:59904366-59904516 | 18 | 13 | 7 | 10 | 12 | 8 | 17 | 22 | 20 | 13 | 14 | 13 | 12 | 18 | 15 | 23 | 17 | 14 | 19 | 29 | 23 | 11 | -1 | 14 | 9 | 12 | 8 | 13 | -1 | 17 | 10 | 6 | 5 | 19 | 12 | 6 | 20 | 23 | 12 | 18 | 16 | 11 | 12 | 8 | 13 | 10 | 73 | 11 | -1 | 12 | 12 |
| CACNA1C | chr12:2550544-2550663 | 8 | 8 | 5 | 9 | 8 | 7 | 9 | 7 | 6 | 3 | 5 | 3 | 3 | 3 | 4 | 3 | 5 | 4 | 2 | 6 | 10 | -1 | -1 | 8 | 8 | 6 | 9 | 6 | 6 | 4 | 7 | 7 | -1 | 7 | 10 | 8 | 7 | 6 | 7 | 8 | 6 | 6 | 5 | 6 | 10 | 6 | 37 | 7 | 10 | 7 | -1 |
| TAC3 | chr12:57012134-57012291 | 12 | 4 | 6 | 6 | 10 | 7 | 0 | 6 | 0 | 0 | 0 | 0 | 0 | 0 | 0 | 0 | 0 | 0 | 0 | 1 | 1 | 0 | -1 | 4 | 11 | 2 | 15 | 2 | 0 | 7 | -1 | 0 | 12 | 10 | -1 | 6 | 10 | 2 | 2 | 1 | 16 | 9 | 14 | 11 | 16 | 2 | 60 | 7 | 6 | 3 | 6 |
| KSR1 | chr17:27558046-27558187 | 9 | 7 | 5 | 7 | 7 | 12 | 10 | 9 | 5 | 4 | 3 | 2 | 2 | 4 | 3 | 3 | 4 | 3 | 3 | 4 | 4 | 11 | 9 | 9 | 9 | 9 | 7 | 8 | 8 | 7 | 7 | 8 | 9 | 9 | 7 | 8 | 7 | 10 | 11 | 6 | 8 | 10 | 11 | 9 | 7 | 9 | 28 | 8 | 11 | 8 | 10 |
| MRRF | chr9:122305788-122305877 | 11 | 16 | 10 | 15 | 14 | 10 | 19 | 16 | 20 | 20 | 18 | 17 | 23 | 22 | 13 | 20 | 24 | 14 | 20 | 25 | 24 | 16 | 15 | 12 | 12 | 13 | 12 | 12 | 21 | 28 | 22 | 13 | 17 | 12 | 13 | 23 | 17 | 10 | 10 | 14 | 16 | 14 | 10 | 16 | 20 | 13 | 60 | 10 | 12 | 14 | 9 |
| TIAL1 | chr10:119593452\|119591976-119593513 | 13 | 13 | 13 | 11 | 13 | 12 | 16 | 13 | 14 | 14 | 13 | 8 | 7 | 15 | 14 | 15 | 14 | 13 | 12 | 14 | 13 | 12 | 13 | 11 | 16 | 11 | 16 | 11 | 18 | 11 | 12 | 16 | 11 | 13 | 15 | 17 | 15 | 13 | 14 | 11 | 13 | 15 | 14 | 16 | 14 | 15 | 37 | 17 | 12 | 13 | 11 |
| ZNF512 | chr2:27619650-27619816\|27619728 | 3 | 6 | 3 | 4 | 4 | 2 | 2 | 4 | 1 | 1 | 1 | 3 | 4 | 1 | 1 | 1 | 1 | 1 | 1 | 2 | 1 | 5 | 2 | 3 | 5 | 2 | 7 | 2 | 2 | 3 | 3 | 6 | 8 | 5 | 7 | 2 | 3 | 2 | 8 | 4 | 5 | 5 | 4 | 8 | 10 | 5 | 25 | 5 | 4 | 4 | 7 |
| MCTP2 | chr15:94314727-94314864 | 7 | 5 | 0 | 5 | 1 | 3 | 3 | 3 | -1 | 0 | 2 | -1 | -1 | -1 | 0 | -1 | 6 | 1 | 0 | 0 | -1 | 0 | 0 | 5 | 10 | 3 | 3 | 3 | -1 | 7 | 6 | 3 | 5 | 6 | 4 | 11 | 2 | 1 | 3 | 2 | 4 | 3 | 3 | 5 | 10 | 4 | 32 | 3 | 0 | 3 | 6 |
| GPATCH2L | chr14:76178208\|76178223-76178322 | 6 | 4 | 3 | 4 | 4 | 5 | 2 | 6 | 2 | 3 | 2 | 0 | 2 | 2 | 2 | 2 | 2 | 2 | 2 | 2 | 2 | 0 | 4 | 3 | 3 | 2 | 3 | 2 | 1 | 4 | 4 | 5 | 8 | 5 | 3 | 7 | 5 | 3 | 3 | 3 | 3 | 6 | 8 | 5 | 4 | 4 | 23 | 5 | 2 | 4 | 6 |
| AL034550.1 | chr20:32451497-32451573 | 8 | 8 | 5 | 5 | 6 | 6 | 8 | 7 | 5 | 4 | 5 | 3 | 3 | 4 | 3 | 5 | 5 | 4 | 6 | 10 | 8 | 7 | 4 | 7 | 7 | 6 | 6 | 6 | 5 | 8 | 8 | 4 | 7 | 9 | 7 | 8 | 6 | 5 | 8 | 3 | 6 | 6 | 7 | 8 | 6 | 7 | 21 | 6 | 4 | 5 | 8 |
| GBA2 | chr9:35743276-35743326 | 16 | 13 | 12 | 11 | 12 | 10 | 13 | 15 | 4 | 4 | 4 | 6 | 7 | 5 | 3 | 5 | 3 | 4 | 4 | 6 | 4 | 8 | 7 | 14 | 20 | 11 | 16 | 10 | 12 | 7 | 8 | 11 | 17 | 11 | 14 | 8 | 9 | 7 | 17 | 9 | 14 | 12 | 13 | 22 | 16 | 16 | 50 | 12 | 8 | 11 | 8 |
| ZNF815P | chr7:5849373-5849516\|5849822 | 9 | 9 | 7 | 9 | 8 | 10 | -1 | 7 | 7 | 8 | 9 | 11 | 8 | 7 | 9 | 9 | 9 | 7 | 8 | 7 | 8 | -1 | -1 | 9 | 11 | 10 | 9 | 10 | -1 | 10 | 11 | 14 | 16 | 12 | 11 | 8 | 8 | 12 | 8 | 9 | 15 | 9 | 8 | 14 | 12 | 9 | 51 | 12 | 10 | 7 | 12 |
| FAM63B | chr15:58806776-58806829 | 5 | 3 | 1 | 2 | 2 | 3 | 2 | 5 | 2 | 0 | 2 | 1 | 1 | 1 | 1 | 1 | 0 | 1 | 1 | 0 | 0 | -1 | 0 | 2 | 3 | 2 | 3 | 1 | 0 | 1 | 1 | 0 | 1 | 5 | 3 | 2 | 4 | 1 | 6 | 1 | 2 | 6 | 5 | 5 | 3 | 4 | 24 | 2 | 1 | 3 | 10 |
| SLC22A1 | chr6:160155306-160155517 | 6 | 3 | 4 | 1 | 1 | 1 | -1 | 4 | -1 | 8 | 0 | 2 | 4 | 3 | 3 | 4 | 8 | 0 | 0 | -1 | -1 | 0 | 0 | 1 | 3 | 3 | 2 | 4 | -1 | 2 | 3 | 0 | 2 | 5 | 5 | 8 | 5 | 3 | 3 | 3 | 4 | 4 | 2 | 4 | 2 | 4 | 23 | 3 | 3 | 0 | 6 |
| ZNF644 | chr1:90935408-90935530 | 3 | 4 | 6 | 6 | 6 | 4 | 6 | 3 | 12 | 19 | 14 | 10 | 11 | 19 | 20 | 11 | 12 | 15 | 11 | 9 | 9 | 3 | 3 | 4 | 4 | 5 | 5 | 6 | 8 | 4 | 5 | 4 | 4 | 5 | 4 | 4 | 4 | 4 | 5 | 8 | 4 | 4 | 3 | 4 | 5 | 5 | 22 | 3 | 3 | 3 | 5 |
| DNAH6 | chr2:84678991-84679094 | 1 | 3 | 0 | 0 | -1 | 8 | 0 | 5 | 0 | 0 | 1 | -1 | 0 | 0 | 0 | 3 | 2 | 0 | 1 | 0 | 0 | 0 | 0 | 1 | 4 | 0 | 0 | 2 | 2 | -1 | 0 | -1 | 2 | 5 | -1 | 0 | 3 | 2 | 9 | 1 | 4 | 0 | 12 | -1 | -1 | 4 | 31 | 1 | 1 | 0 | 0 |
| SCUBE3 | chr6:35228817-35228865 | 3 | 1 | 3 | 3 | 4 | 3 | 1 | 3 | 3 | 3 | 4 | 1 | 3 | 4 | 1 | 0 | 4 | 1 | 0 | -1 | -1 | -1 | -1 | 4 | 4 | 4 | 3 | 6 | 0 | 2 | 2 | 6 | -1 | 2 | 3 | 5 | 5 | 3 | 0 | 5 | 6 | 6 | 5 | 8 | 0 | 5 | 26 | 6 | 3 | 1 | -1 |
| ADTRP | chr6:11717309-11717414 | 9 | 9 | 9 | 7 | 4 | 4 | 2 | 8 | 2 | 4 | 2 | -1 | 5 | 4 | 3 | 3 | 4 | 2 | -1 | -1 | 3 | 0 | 0 | 9 | 9 | 7 | 6 | 7 | 0 | 5 | 5 | 7 | 6 | 6 | 5 | 9 | 7 | 11 | 6 | 4 | 3 | 10 | 9 | 11 | 5 | 5 | 37 | 5 | -1 | 8 | 3 |
| CCDC66 | chr3:56572337-56572428 | 12 | 14 | 12 | 14 | 14 | 11 | 14 | 14 | 25 | 26 | 24 | 26 | 28 | 27 | 24 | 25 | 24 | 26 | 30 | 27 | 24 | 20 | 8 | 15 | 15 | 14 | 15 | 14 | 20 | 18 | 20 | 18 | 13 | 13 | 17 | 18 | 17 | 16 | 11 | 26 | 19 | 12 | 11 | 15 | 18 | 13 | 56 | 19 | 19 | 16 | 11 |
| LPCAT4 | chr15:34361169\|34360754-34361229 | 10 | 8 | 5 | 8 | 9 | 9 | 11 | 10 | 3 | 3 | 3 | 6 | 6 | 4 | 3 | 3 | 3 | 2 | 3 | 7 | 5 | 9 | 8 | 9 | 8 | 9 | 9 | 9 | 7 | 11 | 11 | 10 | 8 | 14 | 9 | 9 | 8 | 6 | 6 | 7 | 9 | 10 | 11 | 11 | 11 | 9 | 27 | 8 | 6 | 9 | 12 |
| ICE2 | chr15:60445693-60445791 | 6 | 10 | 9 | 11 | 12 | 9 | 7 | 8 | 10 | 12 | 11 | 18 | 22 | 12 | 13 | 8 | 11 | 16 | 11 | 5 | 7 | 14 | 11 | 11 | 10 | 10 | 7 | 10 | 10 | 19 | 15 | 12 | 11 | 6 | 8 | 19 | 18 | 8 | 12 | 14 | 10 | 6 | 5 | 9 | 11 | 11 | 50 | 9 | 11 | 9 | 3 |
| GLUD1 | chr10:87067726-87067842 | 2 | 5 | 2 | 2 | 2 | 1 | 5 | 5 | 0 | 0 | 0 | 0 | 0 | 0 | 0 | 1 | 0 | 0 | 0 | 0 | 0 | -1 | -1 | 2 | 5 | 2 | 3 | 2 | 3 | 3 | 0 | 3 | 2 | 2 | 4 | 1 | 2 | 2 | 1 | 1 | 6 | 6 | 2 | 8 | 5 | 3 | 22 | 1 | 1 | 3 | 1 |
| BNIPL | chr1:151044880-151045049\|151045033 | 24 | 16 | 3 | 18 | 16 | 11 | 16 | 15 | -1 | 17 | 11 | 5 | 4 | 7 | 5 | 18 | 13 | 10 | -1 | -1 | -1 | 13 | -1 | 15 | 21 | 11 | 12 | 13 | -1 | 17 | 12 | 15 | 11 | 17 | 13 | 19 | 11 | 11 | 13 | 15 | 20 | 14 | 14 | 17 | 15 | 13 | 65 | 12 | 14 | 14 | 11 |
| METAP1 | chr4:99036043\|99035954-99036089 | 5 | 4 | 3 | 5 | 4 | 5 | 8 | 8 | 5 | 3 | 4 | 5 | 7 | 4 | 2 | 4 | 3 | 4 | 4 | 6 | 5 | 6 | 3 | 5 | 3 | 4 | 3 | 5 | 6 | 5 | 2 | 3 | 2 | 5 | 4 | 3 | 10 | 7 | 5 | 4 | 5 | 6 | 5 | 6 | 4 | 5 | 22 | 4 | 6 | 4 | 3 |
| ZNF542P | chr19:56369279-56369337 | 12 | 12 | 8 | 8 | 7 | 6 | 6 | 12 | 8 | 7 | 7 | 5 | 6 | 5 | 4 | 7 | 6 | 6 | 7 | 9 | 7 | -1 | 10 | 7 | 10 | 5 | 9 | 5 | 15 | 6 | 4 | 16 | 19 | 13 | 9 | 8 | 12 | 5 | 15 | 5 | 9 | 10 | 12 | 16 | 12 | 11 | 49 | 7 | 6 | 6 | 14 |
| ZCCHC17 | chr1:31318167-31318236 | 13 | 9 | 7 | 7 | 7 | 8 | 7 | 13 | 16 | 15 | 18 | 7 | 8 | 12 | 10 | 15 | 12 | 14 | 18 | 16 | 16 | 7 | 5 | 8 | 10 | 7 | 9 | 7 | 7 | 8 | 7 | 10 | 11 | 10 | 12 | 10 | 13 | 6 | 10 | 8 | 10 | 11 | 10 | 13 | 12 | 11 | 32 | 8 | 6 | 8 | 11 |
| TCERG1 | chr5:146481150-146482004\|146481200\|146481476 | 8 | 10 | 6 | 7 | 7 | 7 | 11 | 10 | 11 | 16 | 9 | 17 | 17 | 15 | 14 | 11 | 13 | 13 | 8 | 8 | 11 | 10 | 6 | 9 | 9 | 7 | 9 | 7 | 12 | 7 | 9 | 8 | 7 | 9 | 8 | 9 | 8 | 12 | 9 | 9 | 9 | 7 | 6 | 9 | 10 | 10 | 29 | 7 | 8 | 8 | 5 |
| PRKAA1 | chr5:40774928-40774972 | 18 | 13 | 9 | 11 | 12 | 12 | 10 | 16 | 5 | 7 | 6 | 4 | 5 | 8 | 6 | 8 | 8 | 8 | 9 | 10 | 8 | 17 | 7 | 14 | 15 | 14 | 14 | 14 | 15 | 12 | 13 | 14 | 14 | 11 | 15 | 12 | 15 | 10 | 17 | 7 | 10 | 17 | 18 | 21 | 19 | 17 | 47 | 13 | 10 | 11 | 11 |
| SLC7A11 | chr4:138174535-138174787 | 1 | 2 | -1 | 3 | 3 | 0 | 0 | 2 | 3 | 2 | 2 | 2 | 2 | 1 | 1 | 0 | 1 | 1 | 2 | 2 | 2 | 0 | 0 | 0 | 4 | -1 | 1 | 1 | 0 | 9 | -1 | -1 | -1 | 1 | 2 | -1 | 2 | 1 | 4 | 2 | 0 | 1 | 1 | -1 | 0 | 1 | 23 | 1 | 0 | 6 | 5 |
| NAF1 | chr4:163141931-163141966 | 5 | 4 | 2 | 2 | 2 | 2 | 1 | 5 | 5 | 2 | 2 | 1 | 1 | 3 | 2 | 3 | 5 | 3 | 4 | 5 | 5 | 1 | 5 | 2 | 5 | 1 | 2 | 1 | 2 | 2 | 1 | 8 | 3 | 2 | 6 | 2 | 8 | 2 | 3 | 3 | 3 | 5 | 4 | 5 | 2 | 3 | 21 | 3 | 2 | 1 | 2 |
| UBE2L3 | chr22:21614557-21614665 | 11 | 10 | 5 | 10 | 7 | 10 | 19 | 13 | 5 | 4 | 5 | 2 | 5 | 4 | 3 | 4 | 4 | 4 | 5 | 7 | 5 | 17 | 18 | 9 | 6 | 5 | 4 | 6 | 16 | 6 | 3 | 5 | 5 | 11 | 8 | 11 | 14 | 12 | 4 | 4 | 6 | 17 | 13 | 8 | 4 | 8 | 52 | 6 | 7 | 8 | 12 |
| CNPY2 | chr12:56310310-56310386 | 5 | 7 | 6 | 10 | 10 | 12 | -1 | 10 | 9 | 8 | 8 | 9 | 10 | 10 | 8 | 14 | 15 | 7 | 4 | 11 | 10 | -1 | -1 | 11 | 10 | 11 | 7 | 12 | -1 | 4 | 4 | 5 | 5 | 7 | 13 | 9 | 8 | 13 | 8 | 12 | 11 | 5 | 6 | 10 | 10 | 10 | 46 | 11 | 13 | 12 | 11 |
| PPP1R12A | chr12:79789598-79789680\|79789744 | 15 | 13 | 7 | 7 | 7 | 7 | 8 | 12 | 1 | 1 | 2 | 1 | 1 | 1 | 0 | 2 | 1 | 1 | 2 | 2 | 0 | 3 | 4 | 5 | 7 | 3 | 5 | 3 | 10 | 6 | 6 | 7 | 9 | 14 | 8 | 9 | 12 | 5 | 14 | 3 | 6 | 20 | 16 | 13 | 13 | 7 | 48 | 8 | 4 | 4 | 9 |
| ZNF300 | chr5:150902409-150902566 | 10 | 10 | 12 | 6 | 7 | 7 | 10 | 8 | 4 | 7 | 4 | 3 | 3 | 4 | 5 | 7 | 5 | 5 | 3 | 7 | 5 | -1 | -1 | 7 | 8 | 8 | 7 | 7 | 8 | 8 | 7 | 10 | 13 | 9 | 6 | 11 | 9 | 6 | 11 | 6 | 6 | 11 | 11 | 9 | 7 | 9 | 36 | 7 | 5 | 7 | 11 |
| NCAPD3 | chr11:134183013-134183153 | 10 | 11 | 6 | 8 | 9 | 7 | 8 | 9 | 17 | 14 | 16 | 14 | 13 | 12 | 12 | 15 | 13 | 17 | 14 | 12 | 15 | 0 | -1 | 10 | 7 | 8 | 5 | 7 | 14 | 16 | 12 | 10 | 9 | 9 | 8 | 14 | 17 | 12 | 7 | 11 | 13 | 8 | 8 | 6 | 7 | 6 | 43 | 8 | 11 | 10 | 4 |
| ANKRD20A5P | chr18:14242140-14242419 | 18 | 14 | 21 | 29 | 21 | -1 | 16 | 15 | 9 | 10 | 8 | 17 | 10 | 8 | 10 | 9 | 7 | 6 | 7 | 8 | 10 | -1 | -1 | 12 | 20 | 21 | 24 | 17 | -1 | -1 | -1 | 7 | -1 | 18 | 22 | 16 | 14 | 28 | 15 | 15 | 15 | 19 | 22 | 18 | 19 | 19 | 98 | 12 | -1 | 26 | 14 |
| RP11-156P1.3 | chr17:46985802\|46985823-46985961 | 19 | 17 | 12 | 14 | 11 | 17 | -1 | 20 | 15 | 18 | 16 | 32 | 33 | 19 | 16 | 17 | 16 | 19 | 18 | 25 | 12 | -1 | -1 | 19 | 15 | 14 | 13 | 15 | -1 | 23 | 17 | 9 | 10 | 18 | 16 | 25 | 22 | 24 | 11 | 15 | 19 | 15 | 21 | 16 | 9 | 14 | 83 | 13 | 20 | 15 | 12 |
| RAVER2 | chr1:64745583-64745748 | 2 | 2 | 2 | 2 | 2 | 2 | 0 | 2 | 4 | 6 | 4 | 8 | 7 | 4 | 3 | 6 | 3 | 5 | 4 | 3 | 3 | -1 | 0 | 2 | 4 | 2 | 2 | 1 | -1 | 2 | 1 | 1 | 9 | 2 | 3 | 1 | 4 | 5 | 4 | 4 | 4 | 3 | 2 | 5 | 6 | 3 | 21 | 2 | 4 | 1 | 1 |
| CLDN15 | chr7:101236747-101236886 | 6 | 3 | 13 | 8 | 8 | 9 | 7 | 7 | 13 | 11 | 12 | 9 | 9 | 12 | 11 | 10 | 10 | 12 | 13 | 11 | 12 | 10 | 10 | 9 | 6 | 8 | 11 | 9 | 10 | 7 | 12 | 9 | 10 | 11 | 11 | 11 | 15 | 12 | 18 | 12 | 11 | 9 | 11 | 3 | 12 | 13 | 37 | 11 | 10 | 10 | 14 |
| INTS3 | chr1:153761020-153761109 | 11 | 11 | 13 | 11 | 10 | 9 | 11 | 12 | 16 | 13 | 15 | 9 | 11 | 13 | 12 | 14 | 12 | 13 | 15 | 14 | 13 | 11 | 12 | 10 | 9 | 10 | 8 | 8 | 22 | 9 | 9 | 13 | 16 | 10 | 10 | 7 | 18 | 12 | 12 | 18 | 13 | 9 | 7 | 10 | 11 | 11 | 40 | 15 | 13 | 11 | 16 |
| CADPS | chr3:62530654-62530812 | 3 | 6 | 0 | 6 | 6 | 3 | 0 | 1 | 8 | 2 | 7 | 0 | 0 | 2 | 1 | 3 | 3 | 4 | 5 | 21 | 3 | 0 | -1 | 2 | 7 | 3 | 8 | 3 | 0 | 6 | 9 | -1 | 0 | 12 | 25 | 0 | 2 | 5 | 41 | 5 | 7 | 5 | 3 | 9 | 2 | 12 | 78 | 12 | -1 | 1 | -1 |
| NCOR1 | chr17:16087172-16087345 | 7 | 12 | 14 | 8 | 8 | 7 | 10 | 9 | 11 | 11 | 12 | 9 | 9 | 12 | 11 | 11 | 10 | 13 | 10 | 10 | 10 | 7 | 7 | 8 | 14 | 8 | 16 | 8 | 9 | 14 | 13 | 16 | 21 | 10 | 13 | 9 | 9 | 8 | 14 | 14 | 10 | 8 | 7 | 16 | 11 | 13 | 41 | 12 | 7 | 10 | 10 |
| PWRN1 | chr15:24580227-24580330 | 7 | 8 | 5 | -1 | -1 | 3 | 4 | 9 | 1 | 4 | 2 | 1 | 8 | 2 | 3 | 2 | 3 | 1 | 2 | 3 | 3 | 0 | 0 | 2 | 5 | 6 | 4 | 5 | 7 | 4 | 5 | 3 | 5 | 8 | 5 | 13 | 14 | 3 | 4 | 4 | 4 | -1 | 9 | 3 | 8 | 6 | 38 | 4 | 4 | 12 | -1 |
| ZGRF1 | chr4:112603524-112603697 | 16 | 12 | 11 | 15 | 9 | 13 | -1 | 14 | 18 | 17 | 13 | 14 | 15 | 14 | 14 | 21 | 4 | 12 | 8 | 10 | 15 | -1 | -1 | 12 | 9 | 10 | 11 | 10 | -1 | 18 | 18 | -1 | 11 | 15 | 14 | 18 | 17 | 21 | 10 | 14 | 15 | 19 | 20 | 12 | 19 | 11 | 68 | 17 | 17 | 14 | 9 |
| C7orf26 | chr7:6607537-6607597\|6607851 | 15 | 16 | 21 | 13 | 12 | 12 | 15 | 16 | 20 | 22 | 20 | 18 | 19 | 19 | 19 | 22 | 22 | 23 | 20 | 17 | 23 | 10 | 17 | 11 | 14 | 13 | 12 | 12 | 12 | 10 | 14 | 21 | 23 | 14 | 15 | 10 | 16 | 13 | 15 | 17 | 18 | 10 | 10 | 13 | 19 | 18 | 47 | 22 | 13 | 11 | 12 |
| CARS | chr11:3047753-3048001 | 10 | 13 | 13 | 12 | 11 | 10 | 12 | 11 | 9 | 8 | 11 | 9 | 11 | 9 | 7 | 10 | 7 | 9 | 8 | 12 | 9 | 11 | 11 | 10 | 13 | 10 | 16 | 10 | 9 | 13 | 15 | 13 | 20 | 14 | 10 | 17 | 12 | 11 | 10 | 11 | 11 | 14 | 13 | 12 | 11 | 9 | 35 | 13 | 12 | 13 | 18 |
| SETD5 | chr3:9454005-9454101 | 15 | 11 | 5 | 8 | 7 | 9 | 14 | 16 | 6 | 5 | 5 | 9 | 8 | 7 | 5 | 5 | 5 | 6 | 7 | 7 | 5 | 12 | 10 | 9 | 10 | 8 | 6 | 8 | 19 | 10 | 10 | 8 | 7 | 10 | 10 | 13 | 13 | 13 | 7 | 6 | 7 | 14 | 13 | 18 | 13 | 11 | 40 | 8 | 10 | 8 | 12 |
| TMED5 | chr1:93158815-93158885 | 12 | 11 | 6 | 10 | 10 | 11 | 11 | 12 | 12 | 13 | 12 | 14 | 17 | 13 | 13 | 14 | 14 | 14 | 13 | 16 | 14 | 10 | 9 | 15 | 11 | 12 | 11 | 12 | 15 | 9 | 10 | 10 | 8 | 8 | 9 | 14 | 16 | 15 | 8 | 10 | 13 | 13 | 13 | 13 | 8 | 11 | 35 | 10 | 13 | 15 | 7 |
| ADCY10P1 | chr6:41132875-41134132\|41133000 | 5 | 7 | 9 | 5 | 7 | 4 | 5 | 5 | 8 | 8 | 9 | 4 | 4 | 10 | 6 | 9 | 8 | 8 | 10 | 10 | 10 | 8 | 7 | 6 | 9 | 6 | 9 | 7 | 6 | 13 | 17 | 8 | 14 | 9 | 7 | 15 | 6 | 3 | 9 | 4 | 6 | 5 | 5 | 8 | 13 | 8 | 42 | 6 | 4 | 5 | 19 |
| SNX7 | chr1:98662697-98662816 | 6 | 9 | 7 | 8 | 8 | 8 | 9 | 8 | 12 | 11 | 13 | 16 | 14 | 9 | 11 | 14 | 12 | 13 | 14 | 15 | 13 | 5 | 11 | 8 | 7 | 8 | 6 | 8 | 11 | 10 | 10 | 9 | 6 | 9 | 7 | 11 | 9 | 11 | 10 | 11 | 10 | 7 | 6 | 7 | 8 | 8 | 25 | 9 | 10 | 8 | 6 |
| UBP1 | chr3:33396445\|33396745-33396878 | 13 | 8 | 7 | 6 | 5 | 6 | 5 | 12 | 5 | 4 | 4 | 8 | 9 | 4 | 3 | 4 | 5 | 4 | 4 | 5 | 5 | 10 | 6 | 4 | 4 | 4 | 4 | 3 | 8 | 5 | 4 | 6 | 2 | 9 | 7 | 14 | 10 | 10 | 4 | 13 | 5 | 14 | 16 | 10 | 12 | 6 | 39 | 8 | 8 | 7 | 9 |
| CUL9 | chr6:43182992-43183088 | 11 | 9 | 8 | 12 | 10 | 13 | 5 | 11 | 4 | 6 | 3 | 8 | 7 | 4 | 4 | 3 | 4 | 3 | 3 | 5 | 5 | -1 | 0 | 9 | 9 | 12 | 9 | 10 | -1 | 10 | 7 | 16 | 17 | 10 | 10 | 9 | 7 | 7 | 11 | 4 | 9 | 10 | 7 | 11 | 7 | 8 | 42 | 11 | 8 | 10 | 9 |
| TMEM39B | chr1:32095660-32095762 | 3 | 3 | 4 | 3 | 3 | 3 | 3 | 5 | 9 | 9 | 7 | 18 | 17 | 10 | 10 | 11 | 7 | 7 | 7 | 15 | 10 | 3 | 1 | 5 | 5 | 4 | 4 | 4 | 4 | 11 | 11 | 5 | 5 | 4 | 5 | 25 | 6 | 4 | 5 | 3 | 5 | 5 | 5 | 4 | 5 | 4 | 44 | 5 | 3 | 4 | 5 |
| CDC42BPA | chr1:227219263-227219420 | 1 | 1 | 2 | 0 | 0 | 0 | 2 | 1 | 7 | 9 | 7 | 20 | 25 | 9 | 9 | 5 | 7 | 10 | 8 | 5 | 3 | -1 | 0 | 0 | 1 | 1 | 4 | 1 | 0 | 2 | 1 | 5 | 2 | 2 | 3 | 0 | 3 | 3 | 2 | 4 | 2 | 2 | 2 | 1 | 3 | 3 | 39 | 3 | 1 | 1 | 5 |
| RP11-104N10.1 | chr16:89493498-89493597 | 0 | 0 | 0 | 0 | 0 | 0 | 0 | 0 | 0 | 0 | 0 | 0 | 0 | 4 | 6 | 1 | 0 | 1 | 5 | 14 | 8 | 0 | 0 | 0 | 0 | 0 | 0 | 0 | 0 | 14 | -1 | 0 | 0 | 0 | 0 | 25 | 0 | 0 | 0 | 0 | 0 | 0 | 0 | 0 | 0 | 0 | 42 | 0 | 0 | 0 | 0 |
| ZNF783 | chr7:149291689\|149290675-149291745 | 8 | 12 | 14 | 8 | 9 | 7 | 17 | 10 | 15 | 12 | 14 | 13 | 14 | 12 | 11 | 16 | 12 | 12 | 13 | 15 | 14 | 5 | 12 | 11 | 12 | 11 | 13 | 13 | 9 | 11 | 12 | 13 | 13 | 11 | 13 | 6 | 9 | 12 | 16 | 13 | 13 | 8 | 8 | 14 | 17 | 12 | 37 | 14 | 9 | 9 | 15 |
| PKIB | chr6:122701440-122701500 | 8 | 7 | 6 | 7 | 8 | 6 | -1 | 9 | 2 | 2 | 2 | 2 | 2 | 3 | 1 | 1 | 1 | 2 | 2 | 2 | 1 | 0 | 8 | 13 | 14 | 5 | 11 | 9 | 0 | 5 | 10 | 9 | 14 | 12 | 8 | 9 | 8 | 8 | 12 | 4 | 10 | 9 | 8 | 16 | 16 | 12 | 44 | 9 | 5 | 7 | 11 |
| FRMD4A | chr10:13651903-13651974 | 3 | 4 | 3 | 1 | 2 | 2 | 4 | 4 | 30 | 48 | 23 | 40 | 31 | 62 | 63 | 37 | 50 | 21 | 27 | 18 | 32 | 3 | 3 | 3 | 4 | 2 | 3 | 2 | 3 | 11 | 22 | 3 | 7 | 2 | 6 | 3 | 3 | 2 | 10 | 24 | 8 | 4 | 4 | 4 | 3 | 10 | 65 | 3 | 2 | 2 | 13 |
| ADAM9 | chr8:39076147-39076275 | 6 | 5 | 2 | 3 | 3 | 3 | 3 | 8 | 4 | 3 | 3 | 13 | 19 | 6 | 3 | 4 | 4 | 4 | 4 | 4 | 4 | 8 | 7 | 4 | 3 | 4 | 4 | 4 | 5 | 8 | 4 | 3 | 4 | 6 | 4 | 6 | 9 | 8 | 4 | 8 | 6 | 8 | 8 | 3 | 7 | 4 | 32 | 4 | 7 | 7 | 8 |
| KLHL7 | chr7:23117834-23117964 | 15 | 12 | 9 | 12 | 13 | 13 | 12 | 13 | 10 | 8 | 9 | 12 | 11 | 8 | 9 | 9 | 10 | 9 | 9 | 8 | 8 | 12 | 13 | 13 | 12 | 11 | 12 | 11 | 12 | 15 | 17 | 9 | 8 | 12 | 14 | 16 | 12 | 11 | 11 | 11 | 15 | 13 | 14 | 15 | 12 | 12 | 29 | 11 | 12 | 11 | 11 |
| RP11-983P16.4 | chr12:53044015-53044217 | 2 | 3 | 3 | 3 | 3 | 2 | -1 | 2 | 3 | 2 | 2 | 2 | 3 | 1 | 3 | 2 | 1 | 1 | 5 | 3 | 2 | -1 | 0 | 2 | 1 | 3 | 3 | 2 | -1 | 2 | 3 | 3 | 13 | 4 | 2 | 3 | 3 | 3 | 5 | 6 | 4 | 3 | 3 | 2 | 7 | 2 | 24 | 4 | 2 | 3 | -1 |
| AC007405.6 | chr2:170779793-170779888 | 17 | 15 | 18 | 10 | 12 | 12 | 8 | 15 | 3 | 4 | 3 | 6 | 5 | 5 | 3 | 5 | 3 | 4 | 4 | 7 | 5 | -1 | 9 | 15 | 16 | 13 | 17 | 12 | 15 | 15 | 19 | 12 | 19 | 19 | 23 | 24 | 15 | 15 | 34 | 11 | 13 | 19 | 20 | 16 | -1 | 26 | 83 | 12 | 11 | 12 | 40 |
| TPTE2 | chr13:19436226-19436306 | 14 | 10 | 6 | 6 | 9 | 7 | -1 | 12 | 3 | 2 | 3 | 2 | 2 | 1 | 3 | 3 | 4 | 1 | 3 | 6 | 5 | 0 | 0 | 9 | 11 | 14 | 10 | 6 | -1 | -1 | -1 | 3 | -1 | 4 | 3 | 7 | 11 | 11 | -1 | 8 | 8 | 8 | 10 | 7 | 11 | 12 | 50 | 15 | 11 | 3 | 0 |
| CAMKMT | chr2:44741236-44741424 | 7 | 4 | 3 | 4 | 2 | 4 | 1 | 6 | 4 | 3 | 3 | 5 | 6 | 3 | 3 | 4 | 3 | 5 | 3 | 4 | 2 | 5 | -1 | 5 | 4 | 4 | 3 | 5 | 3 | 4 | 3 | 3 | 4 | 5 | 4 | 10 | 7 | 5 | 5 | 6 | 6 | 6 | 8 | 5 | 6 | 4 | 22 | 5 | 6 | 5 | 3 |
| SFXN5 | chr2:72967237-72967323 | 4 | 5 | 1 | 4 | 3 | 4 | 3 | 4 | 0 | 0 | 0 | 1 | 0 | 0 | 0 | 0 | 0 | 0 | 0 | 0 | 0 | 5 | 1 | 2 | 2 | 2 | 3 | 1 | 3 | 2 | 2 | 2 | 2 | 5 | 4 | 15 | 3 | 1 | 1 | 1 | 2 | 7 | 8 | 5 | 4 | 3 | 29 | 1 | 1 | 2 | 10 |
| C7orf26 | chr7:6607537\|6602664-6607597 | 16 | 16 | 22 | 14 | 13 | 12 | 15 | 17 | 21 | 23 | 21 | 19 | 20 | 19 | 19 | 21 | 22 | 23 | 20 | 16 | 23 | 10 | 17 | 12 | 15 | 13 | 13 | 12 | 14 | 11 | 14 | 22 | 23 | 17 | 17 | 10 | 16 | 14 | 17 | 17 | 20 | 11 | 11 | 16 | 21 | 20 | 48 | 23 | 13 | 12 | 15 |
| CSNK2A1 | chr20:527933-528049 | 8 | 17 | 18 | 12 | 13 | 8 | 13 | 12 | 10 | 8 | 10 | 8 | 10 | 9 | 5 | 11 | 9 | 8 | 9 | 11 | 9 | 19 | 11 | 10 | 17 | 9 | 18 | 9 | 13 | 10 | 10 | 15 | 24 | 15 | 15 | 8 | 9 | 10 | 18 | 13 | 17 | 13 | 11 | 18 | 21 | 17 | 48 | 15 | 11 | 16 | 17 |
| DNAJC6 | chr1:65381821-65381875 | 17 | 11 | 2 | 6 | 4 | 10 | -1 | 18 | 3 | 1 | 3 | 2 | 3 | 2 | 1 | 2 | 2 | 2 | 3 | 2 | 2 | -1 | 0 | 5 | 5 | 7 | 4 | 7 | -1 | 2 | 2 | 5 | -1 | 9 | 12 | 5 | 11 | 11 | 7 | 2 | 6 | 14 | 17 | 14 | 9 | 8 | 51 | 6 | 6 | 9 | 4 |
| DDX19A | chr16:70356846-70356906 | 8 | 4 | 3 | 4 | 4 | 5 | 5 | 8 | 3 | 3 | 3 | 10 | 10 | 4 | 3 | 4 | 5 | 4 | 3 | 5 | 4 | 5 | 6 | 5 | 6 | 4 | 4 | 4 | 8 | 5 | 4 | 5 | 4 | 6 | 7 | 3 | 10 | 7 | 8 | 5 | 8 | 9 | 10 | 8 | 9 | 5 | 24 | 7 | 5 | 6 | 5 |
| NALCN | chr13:101067510-101067838 | 14 | 7 | 2 | 10 | 8 | 13 | -1 | 6 | 2 | 2 | 1 | 6 | 6 | 2 | 1 | 2 | 1 | 1 | 2 | 2 | 2 | -1 | 0 | 4 | 3 | 7 | 7 | 2 | 0 | 8 | 6 | 0 | 7 | 10 | 0 | 6 | 6 | 7 | 4 | 13 | 6 | 8 | 11 | -1 | -1 | 11 | 44 | 8 | -1 | 8 | 0 |
| KIAA1755 | chr20:38221574-38221677 | 5 | 5 | 5 | 6 | 5 | 4 | 6 | 4 | 2 | 1 | 1 | 2 | 2 | 1 | 1 | 2 | 2 | 1 | 2 | 2 | 1 | 3 | 3 | 5 | 6 | 6 | 8 | 6 | 3 | 4 | 5 | 6 | 4 | 4 | 4 | 5 | 3 | 3 | 6 | 4 | 6 | 6 | 5 | 8 | 8 | 7 | 21 | 5 | 2 | 2 | 11 |
| POU2F3 | chr11:120311295-120311400 | 3 | 3 | 1 | -1 | 0 | 5 | 2 | 5 | -1 | 0 | -1 | 0 | -1 | -1 | 0 | -1 | 0 | 0 | 0 | 0 | 0 | 3 | 0 | 5 | 5 | 3 | 2 | 2 | 0 | 5 | 1 | 2 | 0 | 3 | 5 | 5 | 3 | 1 | 2 | 4 | 7 | 3 | 4 | 4 | 5 | 1 | 21 | 5 | -1 | 3 | 7 |
| RNF34 | chr12:121402742-121402807 | 8 | 8 | 5 | 7 | 6 | 6 | 8 | 7 | 6 | 5 | 7 | 7 | 6 | 5 | 5 | 6 | 4 | 9 | 8 | 5 | 5 | 6 | 9 | 5 | 7 | 5 | 7 | 5 | 8 | 5 | 5 | 9 | 4 | 9 | 7 | 7 | 9 | 7 | 8 | 6 | 7 | 7 | 6 | 12 | 14 | 8 | 24 | 7 | 7 | 6 | 9 |
| LONRF3 | chrX:118986925-118987080 | 15 | 15 | 21 | 11 | 11 | 9 | 12 | 15 | 22 | 22 | 14 | 10 | 11 | 21 | 18 | 17 | 21 | 14 | 20 | 23 | 24 | -1 | -1 | 12 | 15 | 10 | 9 | 10 | 0 | 11 | 10 | 8 | 25 | 18 | 17 | 22 | 14 | 13 | 26 | 17 | 14 | 13 | 17 | 18 | 13 | 17 | 66 | 19 | 13 | 12 | 23 |
| ZFHX2 | chr14:23528629-23528714 | 4 | 10 | 5 | 4 | 2 | 6 | 0 | 5 | 11 | 4 | 11 | 8 | 6 | 4 | 5 | 6 | 5 | 7 | 6 | 6 | 10 | 0 | 0 | 5 | 1 | 4 | 6 | 5 | -1 | 8 | 5 | 3 | -1 | 18 | -1 | 2 | 5 | 7 | 14 | 8 | 13 | 8 | 10 | 6 | -1 | 6 | 43 | 9 | 5 | -1 | -1 |
| ARL6 | chr3:97765991-97766089 | 7 | 7 | 4 | 7 | 5 | 7 | 7 | 9 | 11 | 22 | 12 | 16 | 14 | 22 | 27 | 14 | 11 | 9 | 13 | 13 | 19 | 6 | 9 | 10 | 6 | 7 | 5 | 7 | 15 | 5 | 7 | 6 | 5 | 7 | 7 | 18 | 9 | 10 | 8 | 5 | 8 | 6 | 6 | 9 | 4 | 9 | 34 | 4 | 8 | 7 | 3 |
| P4HA3 | chr11:74298048-74298099 | 5 | 5 | 7 | 3 | 3 | 4 | 9 | 3 | -1 | 6 | 13 | 6 | 8 | 8 | -1 | -1 | 5 | 13 | 7 | -1 | -1 | -1 | 0 | 6 | 4 | 5 | 6 | 5 | 8 | 5 | 3 | 6 | 2 | 4 | 6 | 13 | 4 | 8 | 6 | 8 | 7 | 4 | 5 | 7 | 2 | 3 | 29 | 7 | 4 | 8 | -1 |
| FAM86DP | chr3:75427427-75427908 | 7 | 5 | 7 | 4 | 5 | 6 | 5 | 6 | 7 | 6 | 6 | 3 | 3 | 5 | 6 | 6 | 4 | 12 | 6 | 3 | 5 | 11 | 8 | 6 | 6 | 6 | 6 | 5 | 9 | 3 | 5 | 6 | 9 | 5 | 6 | 3 | 5 | 6 | 6 | 6 | 6 | 5 | 5 | 6 | 7 | 5 | 20 | 5 | 7 | 5 | 8 |
| ALMS1 | chr2:73620496-73620760 | 3 | 3 | 7 | 2 | 1 | 2 | -1 | 4 | 4 | 4 | 7 | 3 | 3 | 6 | 2 | 3 | 2 | 5 | 4 | 3 | 4 | -1 | 0 | 4 | 5 | 4 | 2 | 3 | -1 | 6 | 10 | 1 | 5 | 5 | 6 | 3 | 3 | 4 | 3 | 6 | 7 | 2 | 4 | 2 | 4 | 3 | 23 | 3 | 4 | 4 | 5 |
| RALGAPA1 | chr14:35682058-35682152 | 2 | 6 | 6 | 4 | 7 | 3 | -1 | 3 | 4 | 10 | 4 | 2 | 3 | 4 | 2 | -1 | 1 | 1 | -1 | 2 | -1 | 0 | 0 | 8 | 11 | 5 | 11 | 5 | -1 | 5 | 4 | -1 | 17 | 4 | 6 | 2 | 4 | 4 | 6 | 5 | 8 | 4 | 2 | 18 | 7 | 7 | 40 | 3 | 3 | 9 | 5 |
| FGL1 | chr8:17891301\|17891681-17891807 | 15 | 9 | 9 | 12 | 14 | 14 | -1 | 12 | 46 | 35 | 29 | 28 | 26 | 50 | 46 | 27 | 30 | 33 | 35 | -1 | 27 | -1 | -1 | 10 | 11 | 14 | 14 | 13 | -1 | 7 | 15 | -1 | 10 | 11 | 10 | 15 | 14 | 16 | 16 | 19 | 23 | 5 | 12 | 12 | 7 | 9 | 67 | 13 | 22 | 20 | 14 |
| ATF2 | chr2:175092543-175092616 | 18 | 8 | 4 | 8 | 8 | 12 | 13 | 17 | 2 | 2 | 3 | 1 | 2 | 2 | 1 | 2 | 2 | 2 | 3 | 3 | 3 | 9 | 9 | 9 | 10 | 8 | 7 | 8 | 8 | 10 | 10 | 6 | 6 | 10 | 9 | 14 | 18 | 7 | 6 | 6 | 7 | 18 | 20 | 14 | 13 | 8 | 46 | 5 | 7 | 9 | 16 |
| CMPK2 | chr2:6842130-6842243 | 9 | 10 | 14 | 11 | 8 | 15 | 0 | 9 | 7 | 12 | 12 | 12 | 13 | 10 | 12 | 16 | 12 | 10 | 15 | 15 | 10 | 0 | 0 | 10 | 14 | 8 | 7 | 11 | -1 | 9 | 10 | 10 | 8 | 11 | 12 | 5 | 11 | -1 | 19 | 12 | 15 | 8 | 9 | 8 | 10 | 10 | 48 | 8 | 7 | 10 | 16 |
| LINC00152 | chr2:87700398-87700466 | 4 | 3 | 6 | 4 | 5 | 3 | -1 | 5 | 3 | 5 | 3 | 1 | -1 | 4 | 5 | -1 | 7 | 0 | 3 | 0 | -1 | -1 | 0 | 4 | 7 | 2 | 5 | 4 | 0 | 4 | 3 | 2 | 4 | 4 | 4 | 2 | 4 | 8 | 6 | 3 | 6 | 9 | 4 | 4 | 6 | 4 | 23 | 3 | 4 | 4 | 4 |
| BAG1 | chr9:33262105-33262246 | 8 | 5 | 2 | 3 | 2 | 4 | 9 | 9 | 1 | 1 | 1 | 3 | 3 | 1 | 1 | 1 | 1 | 1 | 1 | 1 | 1 | 6 | 5 | 3 | 4 | 2 | 4 | 2 | 5 | 2 | 2 | 2 | 3 | 5 | 6 | 6 | 5 | 10 | 4 | 1 | 5 | 24 | 22 | 7 | 4 | 4 | 46 | 3 | 4 | 6 | 2 |
| RP9P | chr7:32933400-32933506 | 12 | 16 | 9 | 16 | 15 | 15 | 15 | 14 | 11 | 13 | 11 | 14 | 12 | 12 | 12 | 12 | 13 | 11 | 11 | 15 | 14 | 20 | 13 | 14 | 14 | 14 | 13 | 14 | 13 | 14 | 14 | 16 | 14 | 17 | 12 | 11 | 14 | 16 | 13 | 15 | 17 | 13 | 11 | 14 | 15 | 15 | 31 | 16 | 16 | 16 | 18 |
| TSC22D2 | chr3:150438198-150438354 | 11 | 11 | 9 | 6 | 7 | 6 | 7 | 11 | 10 | 8 | 9 | 4 | 4 | 7 | 4 | 9 | 8 | 9 | 8 | 11 | 9 | 15 | 9 | 8 | 11 | 7 | 9 | 7 | 14 | 6 | 4 | 10 | 9 | 8 | 11 | 2 | 14 | 11 | 14 | 11 | 11 | 8 | 8 | 12 | 12 | 10 | 34 | 13 | 8 | 10 | 7 |
| TRHDE | chr12:72653645-72653682 | 11 | 7 | 4 | 4 | 3 | 4 | 2 | 12 | 0 | 1 | 0 | 2 | 1 | 0 | 0 | 0 | 0 | 1 | 1 | 2 | 1 | 6 | 0 | 5 | 8 | 1 | 1 | 2 | -1 | 2 | 0 | 5 | 7 | 6 | 3 | 7 | 6 | 3 | 6 | 0 | 3 | 16 | 7 | 12 | 2 | 4 | 36 | 2 | 2 | 2 | 0 |
| PPM1K | chr4:88271000-88276777\|88271287\|88271258 | 6 | 6 | 5 | 5 | 4 | 4 | 3 | 8 | 4 | 5 | 6 | 3 | 4 | 5 | 4 | 4 | 4 | 5 | 4 | 2 | 3 | 9 | 5 | 4 | 5 | 4 | 7 | 4 | 5 | 5 | 3 | 11 | 6 | 6 | 6 | 4 | 4 | 8 | 6 | 11 | 9 | 6 | 6 | 6 | 6 | 6 | 24 | 12 | 5 | 5 | 3 |
| ALDH1L2 | chr12:105042192-105042313 | 3 | 3 | 2 | 3 | 3 | 2 | -1 | 3 | 1 | 4 | 2 | 8 | 8 | 5 | 4 | 2 | 3 | 5 | 1 | 4 | 4 | -1 | -1 | 3 | 5 | 4 | 4 | 3 | -1 | 4 | 4 | 4 | 7 | 3 | 3 | 3 | 4 | 4 | 9 | 8 | 4 | 4 | 3 | 3 | 8 | 6 | 24 | 3 | 5 | 4 | 5 |
| CPEB2 | chr4:15008338-15008427 | 11 | 18 | 19 | 22 | 19 | 14 | 14 | 12 | 37 | 48 | 37 | 27 | 32 | 47 | 48 | 43 | 48 | 43 | 41 | 45 | 46 | 9 | -1 | 18 | 18 | 19 | 19 | 18 | 14 | 35 | 40 | 24 | 17 | 14 | 18 | 25 | 16 | 21 | 14 | 29 | 20 | 12 | 11 | 18 | 17 | 22 | 80 | 18 | 16 | 17 | 10 |
| USP30 | chr12:109054720-109054792 | 12 | 8 | 4 | 4 | 4 | 5 | 4 | 9 | 2 | 2 | 3 | 2 | 2 | 1 | 2 | 2 | 3 | 2 | 1 | 3 | 4 | 7 | 0 | 6 | 7 | 5 | 4 | 5 | 8 | 4 | 4 | 6 | 3 | 7 | 5 | 6 | 6 | 5 | 6 | 4 | 5 | 8 | 11 | 9 | 4 | 7 | 25 | 4 | 5 | 5 | 7 |
| NAGPA | chr16:5028297-5028365\|5028398 | 12 | 11 | 13 | 12 | 12 | 12 | 18 | 15 | 20 | 25 | 21 | 22 | 22 | 29 | 27 | 28 | 23 | 18 | 21 | 17 | 24 | 19 | 8 | 19 | 16 | 21 | 15 | 22 | 9 | 13 | 13 | 15 | 14 | 12 | 13 | 9 | 19 | 17 | 12 | 12 | 21 | 18 | 17 | 20 | 15 | 19 | 47 | 14 | 13 | 14 | 8 |

SFig5, pg15

| Gene Symbol | Exon Location (hg38, human) | Adipose - Subcutaneous | Adipose - Visceral (Omentum) | Adrenal Gland | Artery - Aorta | Artery - Coronary | Artery - Tibial | Bladder | Breast - Mammary Tissue | Brain - Amygdala | Brain - Anterior cingulate cortex (BA24) | Brain - Caudate (basal ganglia) | Brain - Cerebellar Hemisphere | Brain - Cerebellum | Brain - Cortex | Brain - Frontal Cortex (BA9) | Brain - Hippocampus | Brain - Hypothalamus | Brain - Nucleus accumbens (basal ganglia) | Brain - Putamen (basal ganglia) | Brain - Spinal cord (cervical c-1) | Brain - Substantia nigra | Cervix - Ectocervix | Cervix - Endocervix | Colon - Sigmoid | Colon - Transverse | Esophagus - Gastroesophageal Junction | Esophagus - Mucosa | Esophagus - Muscularis | Fallopian Tube | Heart - Atrial Appendage | Heart - Left Ventricle | Kidney - Cortex | Liver | Lung | Minor Salivary Gland | Muscle - Skeletal | Nerve - Tibial | Ovary | Pancreas | Pituitary | Prostate | Skin - Not Sun Exposed (Suprapubic) | Skin - Sun Exposed (Lower leg) | Small Intestine - Terminal Ileum | Spleen | Stomach | Testis | Thyroid | Uterus | Vagina | Whole Blood |
|---|---|---|---|---|---|---|---|---|---|---|---|---|---|---|---|---|---|---|---|---|---|---|---|---|---|---|---|---|---|---|---|---|---|---|---|---|---|---|---|---|---|---|---|---|---|---|---|---|---|---|---|---|
| FER | chr5:108897963-108898132 | 7 | 17 | 8 | 13 | 10 | 8 | 9 | 11 | 15 | 18 | 21 | 24 | 28 | 23 | 16 | 17 | 21 | 18 | 14 | 13 | 13 | -1 | -1 | 13 | 14 | 10 | 11 | 10 | -1 | 19 | 9 | 17 | 20 | 16 | 13 | 9 | 11 | 14 | 19 | 18 | 18 | 13 | 11 | 13 | 12 | 12 | 62 | 18 | 16 | 20 | 8 |
| SIRT3 | chr11:233740\|233911-234821 | 2 | 3 | 1 | 3 | 3 | 3 | 2 | 4 | 3 | 3 | 4 | 7 | 8 | 4 | 4 | 5 | 3 | 5 | 5 | 7 | 2 | 5 | 7 | 4 | 3 | 2 | 3 | 2 | 5 | 1 | 0 | 5 | 2 | 4 | 3 | 1 | 6 | 8 | 2 | 9 | 5 | 3 | 3 | 5 | 6 | 2 | 22 | 6 | 6 | 2 | 7 |
| FREM1 | chr9:14788075-14788177\|14788085 | 7 | 6 | 6 | 3 | 4 | 3 | -1 | 6 | 28 | 16 | 20 | 16 | 16 | 15 | 23 | 17 | 16 | 26 | 31 | 21 | 19 | -1 | -1 | 6 | 6 | 6 | 6 | 4 | 6 | 6 | 4 | 7 | 6 | 3 | 4 | 5 | 9 | 9 | 9 | 7 | 7 | 6 | 5 | 9 | 8 | 4 | 31 | 4 | 5 | 6 | -1 |
| CRELD2 | chr22:49922294\|49921871\|49922297-49922443 | 6 | 6 | 4 | 7 | 6 | 6 | 11 | 6 | 5 | 6 | 5 | 10 | 10 | 6 | 5 | 6 | 5 | 5 | 6 | 6 | 6 | 5 | 4 | 10 | 7 | 8 | 5 | 9 | 8 | 9 | 10 | 6 | 7 | 6 | 6 | 10 | 6 | 5 | 5 | 7 | 8 | 6 | 6 | 7 | 5 | 7 | 22 | 9 | 8 | 8 | 5 |
| FBXO11 | chr2:47797973-47798121 | 3 | 8 | 3 | 2 | 7 | 2 | -1 | 4 | -1 | -1 | 6 | 7 | 4 | 3 | 6 | 0 | 1 | 0 | -1 | -1 | 0 | 0 | 0 | 2 | 2 | 1 | 3 | 3 | 0 | 5 | 4 | -1 | -1 | 4 | -1 | 1 | 3 | 3 | 7 | 3 | 4 | 3 | 1 | 1 | 5 | 3 | 21 | 4 | 3 | 2 | 6 |
| RP11-333E1.1 | chr17:5213015-5213216 | 6 | 8 | 5 | 8 | 7 | 8 | 8 | 5 | 12 | 13 | 16 | 8 | 8 | 16 | 14 | 14 | 12 | 14 | 9 | 12 | 8 | 3 | 5 | 7 | 7 | 11 | 6 | 8 | 5 | 10 | 10 | 9 | 3 | 8 | 5 | 12 | 7 | 12 | 5 | 6 | 9 | 4 | 4 | 8 | 7 | 6 | 25 | 6 | 8 | 10 | 6 |
| TSC2 | chr16:2060953-2061039 | 5 | 6 | 5 | 6 | 5 | 4 | 7 | 6 | 6 | 5 | 7 | 11 | 11 | 5 | 5 | 8 | 7 | 6 | 6 | 9 | 9 | 5 | 5 | 7 | 7 | 5 | 7 | 6 | 7 | 4 | 4 | 10 | 13 | 9 | 8 | 3 | 8 | 4 | 9 | 14 | 9 | 7 | 7 | 8 | 7 | 6 | 26 | 10 | 5 | 7 | 8 |
| KCNK17 | chr6:39300373-39300505 | 1 | 3 | -1 | 3 | 3 | 3 | -1 | 3 | -1 | -1 | 8 | 1 | 10 | 7 | 0 | 5 | 0 | 6 | 4 | 4 | 2 | 0 | 0 | 2 | 2 | 4 | 6 | 4 | 0 | 3 | 4 | -1 | 6 | 3 | 7 | 3 | 3 | 3 | 9 | 3 | 6 | -1 | 3 | 4 | 4 | 8 | 24 | 4 | 3 | 2 | 3 |
| CTC-432M15.3 | chr5:131440146-131441602\|131440228 | 3 | 2 | 1 | 1 | 3 | 3 | 1 | 3 | 8 | 10 | 11 | 10 | 10 | 8 | 7 | 5 | 15 | 19 | 7 | 4 | 5 | 2 | 0 | 2 | 2 | 1 | 1 | 1 | 3 | 5 | 4 | 4 | 2 | 2 | 1 | 17 | 5 | 1 | 2 | 2 | 1 | 2 | 2 | 2 | 1 | 1 | 26 | 2 | 1 | 1 | 1 |
| TRPM7 | chr15:50588191-50588250 | 9 | 11 | 9 | 8 | 9 | 7 | 7 | 9 | 6 | 4 | 5 | 3 | 2 | 3 | 4 | 3 | 3 | 4 | 6 | 4 | 6 | 6 | 4 | 6 | 10 | 6 | 9 | 6 | 9 | 5 | 6 | 10 | 13 | 8 | 9 | 6 | 7 | 7 | 11 | 5 | 6 | 7 | 6 | 14 | 12 | 11 | 28 | 8 | 6 | 7 | 8 |
| DYRK3 | chr1:206636875-206636968 | 22 | 17 | 15 | 15 | 14 | 17 | -1 | 23 | 29 | 24 | 27 | 22 | 31 | 20 | 17 | 21 | 20 | 26 | 22 | 23 | 30 | -1 | -1 | 15 | 17 | 14 | 16 | 15 | 19 | 15 | 12 | 19 | 10 | 18 | 24 | 15 | 20 | 22 | 24 | 23 | 19 | 20 | 21 | 26 | 13 | 24 | 72 | 22 | 23 | 16 | 10 |
| CSDE1 | chr1:114737471-114737563 | 7 | 10 | 9 | 12 | 13 | 10 | 12 | 8 | 14 | 17 | 15 | 16 | 18 | 18 | 19 | 15 | 13 | 16 | 13 | 16 | 14 | 11 | 12 | 13 | 13 | 11 | 13 | 11 | 10 | 16 | 13 | 14 | 11 | 10 | 11 | 9 | 10 | 12 | 12 | 10 | 14 | 9 | 7 | 13 | 9 | 12 | 30 | 13 | 12 | 13 | 8 |
| TANK | chr2:161161285-161163358\|161161412\|161163660\|161163453 | 5 | 9 | 5 | 8 | 7 | 9 | 8 | 7 | 12 | 11 | 11 | 8 | 8 | 10 | 8 | 10 | 10 | 11 | 13 | 8 | 10 | 9 | 12 | 9 | 9 | 7 | 6 | 7 | 12 | 8 | 7 | 8 | 7 | 10 | 5 | 13 | 10 | 14 | 7 | 12 | 9 | 7 | 6 | 11 | 11 | 9 | 26 | 10 | 11 | 8 | 10 |
| NPHP1 | chr2:110125189\|110124437\|110125219-110125323 | 5 | 14 | 16 | 7 | 7 | 7 | 10 | 9 | 16 | 11 | 15 | 10 | 13 | 13 | 12 | 17 | 14 | 12 | 9 | 13 | 10 | 12 | 11 | 8 | 8 | 8 | 11 | 7 | 16 | 10 | 13 | 10 | 22 | 16 | 9 | 8 | 5 | 6 | 14 | 17 | 15 | 7 | 5 | 10 | 17 | 12 | 43 | 9 | 7 | 12 | 16 |
| ACSS3 | chr12:81252679-81252804 | 12 | 5 | 1 | 8 | 6 | 10 | 11 | 10 | 8 | 8 | 7 | 11 | 10 | 6 | 7 | 8 | 9 | 11 | 7 | 5 | 6 | 11 | 10 | 11 | 7 | 7 | 8 | 7 | 14 | 14 | 10 | 3 | 3 | 8 | 6 | 16 | 27 | 12 | 7 | 9 | 10 | 14 | 17 | 8 | 6 | 7 | 45 | 7 | 11 | 10 | 8 |
| BBOF1 | chr14:74022916-74023144\|74023148 | 14 | 15 | 14 | 13 | 16 | 16 | 13 | 22 | 12 | 9 | 11 | 15 | 15 | 9 | 10 | 12 | 11 | 11 | 10 | 10 | 7 | -1 | 25 | 18 | 19 | 14 | 16 | 13 | 45 | 17 | 17 | 15 | 14 | 19 | 17 | 13 | 19 | 35 | 22 | 21 | 21 | 16 | 18 | 21 | 15 | 18 | 76 | 18 | 23 | 22 | 1 |
| PTPN3 | chr9:109450243-109450369 | 6 | 5 | 2 | 4 | 3 | 5 | 5 | 5 | 6 | 4 | 6 | 5 | 8 | 3 | 4 | 7 | 6 | 7 | -1 | -1 | 0 | -1 | -1 | 3 | 3 | 7 | 3 | 2 | -1 | 3 | 2 | 4 | 5 | 6 | 3 | 3 | 4 | 3 | 2 | 5 | 6 | 4 | 4 | 4 | 0 | 5 | 20 | 4 | 4 | 3 | -1 |
| AZIN1-AS1 | chr8:102938663\|102941539-102941698 | 19 | 20 | 11 | 13 | 14 | 14 | -1 | 19 | 12 | 8 | 13 | 11 | 14 | 12 | 10 | 10 | 8 | 12 | 14 | 11 | 11 | -1 | -1 | 15 | 15 | 15 | 20 | 11 | 21 | 23 | 23 | 22 | 14 | 23 | 14 | 17 | 16 | 21 | 22 | 14 | 19 | 16 | 16 | 24 | 24 | 21 | 69 | 16 | 17 | 14 | 33 |
| ARL6IP6 | chr2:152719898-152720004 | 4 | 4 | 12 | 3 | 4 | 3 | 6 | 5 | 3 | 3 | 3 | 8 | 8 | 4 | 4 | 4 | 3 | 5 | 6 | 4 | 3 | 5 | 7 | 6 | 4 | 6 | 3 | 8 | 6 | 4 | 4 | 6 | 6 | 4 | 4 | 3 | 5 | 6 | 6 | 8 | 6 | 4 | 4 | 8 | 6 | 7 | 21 | 6 | 6 | 3 | 1 |
| ZBTB34 | chr9:126863592-126863712 | 11 | 19 | 24 | 12 | 15 | 10 | -1 | 11 | 11 | 7 | 11 | 7 | 8 | 7 | 9 | 8 | 15 | 5 | 7 | 11 | 10 | -1 | -1 | 13 | 20 | 18 | 20 | 17 | -1 | 11 | 15 | 7 | 15 | 11 | 11 | 19 | 7 | 12 | 23 | 21 | 11 | 14 | 14 | 20 | 16 | 19 | 63 | 11 | 9 | 12 | 8 |
| FREM1 | chr9:14908764-14909061 | 10 | 5 | 5 | 2 | 3 | 2 | 0 | 6 | 3 | 9 | 19 | 4 | 5 | 14 | 12 | -1 | 35 | -1 | -1 | 19 | -1 | -1 | 0 | 4 | 2 | 5 | 7 | 5 | 0 | 4 | 11 | 2 | -1 | 3 | -1 | 5 | 9 | 2 | 3 | 6 | 1 | 5 | 4 | 5 | 0 | 2 | 27 | 5 | 5 | 8 | 0 |
| TDP2 | chr6:24653362-24653473 | 8 | 5 | 2 | 4 | 3 | 7 | 6 | 10 | 4 | 3 | 4 | 3 | 3 | 3 | 2 | 3 | 3 | 5 | 4 | 5 | 2 | 12 | 13 | 4 | 4 | 4 | 4 | 3 | 13 | 3 | 2 | 4 | 2 | 6 | 6 | 2 | 15 | 11 | 4 | 4 | 7 | 9 | 11 | 5 | 8 | 4 | 33 | 9 | 8 | 8 | 6 |
| BRD4 | chr19:15268154\|15268158-15268281 | 8 | 9 | 8 | 7 | 9 | 7 | 10 | 9 | 6 | 7 | 6 | 9 | 9 | 6 | 5 | 6 | 5 | 6 | 6 | 5 | 6 | 8 | 9 | 8 | 8 | 7 | 9 | 7 | 11 | 7 | 6 | 10 | 8 | 8 | 11 | 3 | 9 | 9 | 13 | 11 | 10 | 9 | 9 | 11 | 13 | 8 | 23 | 11 | 8 | 9 | 7 |
| NUP43 | chr6:149728379\|149727414-149730763 | 7 | 8 | 3 | 8 | 8 | 11 | 10 | 8 | 6 | 4 | 5 | 8 | 6 | 4 | 5 | 6 | 5 | 5 | 6 | 7 | 6 | 3 | 6 | 7 | 6 | 7 | 4 | 6 | 5 | 4 | 4 | 5 | 5 | 8 | 5 | 4 | 9 | 6 | 8 | 5 | 8 | 8 | 7 | 12 | 10 | 7 | 23 | 8 | 8 | 7 | 9 |
| ATXN7L1 | chr7:105649407-105649560 | 6 | 7 | 6 | 5 | 5 | 6 | 8 | 6 | 7 | 3 | 8 | 7 | 5 | 5 | 2 | 4 | 4 | 5 | 6 | 4 | 8 | -1 | -1 | 6 | 5 | 6 | 6 | 5 | -1 | 7 | 8 | 8 | 9 | 6 | 8 | 5 | 5 | 6 | 9 | 8 | 9 | 5 | 5 | 8 | 7 | 7 | 24 | 7 | 6 | 8 | 8 |
| MCF2L2 | chr3:183221427-183221588 | 8 | -1 | 1 | 4 | 0 | 6 | 0 | 2 | 0 | 0 | 0 | 1 | 1 | 0 | 0 | 0 | 0 | 0 | 0 | 1 | 0 | 0 | 0 | 2 | 0 | 5 | 2 | 3 | 0 | -1 | 1 | -1 | 0 | 6 | -1 | 3 | 2 | 0 | 0 | 1 | 1 | 6 | 6 | 6 | 1 | 3 | 21 | 2 | 3 | 3 | 2 |
| RP11-295P9.13 | chr10:13648620-13652489\|13648768\|13648698 | 5 | 8 | 11 | 6 | 5 | 4 | 6 | 5 | 4 | 10 | 7 | 5 | 6 | 5 | 8 | 7 | 4 | 6 | 5 | 5 | 7 | 4 | -1 | 6 | 8 | 6 | 9 | 4 | 7 | 9 | 8 | 11 | 12 | 9 | 11 | 9 | 4 | 7 | 8 | 7 | 9 | 4 | 4 | 8 | 10 | 9 | 28 | 9 | 5 | 5 | 13 |
| NAGPA | chr16:5028297\|5028293-5028365 | 10 | 10 | 13 | 11 | 11 | 11 | 18 | 14 | 21 | 26 | 22 | 23 | 24 | 29 | 28 | 29 | 23 | 18 | 22 | 16 | 23 | 22 | 6 | 18 | 15 | 20 | 14 | 21 | 10 | 12 | 12 | 14 | 13 | 11 | 12 | 9 | 19 | 16 | 11 | 13 | 20 | 16 | 15 | 19 | 15 | 19 | 46 | 13 | 12 | 13 | 7 |
| ABCB8 | chr7:151032844\|151031241\|151028780-151033061 | 4 | 5 | 3 | 5 | 5 | 5 | 7 | 5 | 6 | 5 | 7 | 16 | 16 | 6 | 5 | 6 | 7 | 6 | 5 | 9 | 6 | 10 | 4 | 6 | 5 | 5 | 3 | 5 | 5 | 4 | 4 | 5 | 4 | 5 | 4 | 8 | 7 | 9 | 8 | 7 | 6 | 6 | 5 | 5 | 5 | 5 | 23 | 7 | 7 | 6 | 5 |
| PIGF | chr2:46588075-46591925\|46588179\|46588126 | 17 | 16 | 14 | 21 | 20 | 25 | 28 | 20 | 11 | 13 | 10 | 11 | 12 | 13 | 13 | 12 | 13 | 11 | 9 | 13 | 11 | 16 | 21 | 19 | 19 | 16 | 17 | 17 | 20 | 16 | 18 | 17 | 19 | 19 | 18 | 27 | 19 | 18 | 20 | 16 | 21 | 19 | 18 | 24 | 23 | 19 | 43 | 20 | 18 | 16 | 17 |
| ZDHHC6 | chr10:112433541-112433638\|112433616 | 17 | 7 | 7 | 9 | 9 | 14 | 13 | 16 | 2 | 2 | 2 | 1 | 1 | 2 | 2 | 2 | 2 | 2 | 2 | 3 | 3 | 18 | 11 | 9 | 10 | 9 | 7 | 9 | 11 | 6 | 6 | 5 | 4 | 8 | 8 | 10 | 15 | 11 | 11 | 7 | 9 | 20 | 27 | 13 | 19 | 9 | 47 | 7 | 10 | 10 | 11 |
| PCYOX1L | chr5:149362967-149363016\|149363050 | 9 | 6 | 5 | 5 | 4 | 4 | 9 | 7 | 2 | 1 | 1 | 1 | 2 | 1 | 1 | 3 | 1 | 1 | 1 | 1 | 1 | 5 | 3 | 3 | 6 | 4 | 5 | 4 | 6 | 4 | 4 | 8 | 8 | 7 | 5 | 5 | 5 | 2 | 10 | 3 | 6 | 9 | 12 | 13 | 13 | 7 | 27 | 6 | 2 | 5 | 9 |
| CBWD2 | chr2:113457065-113457204\|113457209 | 11 | 6 | 5 | 4 | 4 | 5 | 4 | 11 | 2 | 2 | 2 | 5 | 5 | 3 | 3 | 3 | 3 | 3 | 3 | 6 | 3 | 6 | 7 | 4 | 5 | 4 | 4 | 4 | 6 | 3 | 1 | 4 | 6 | 8 | 6 | 3 | 12 | 9 | 6 | 7 | 7 | 12 | 14 | 8 | 8 | 5 | 27 | 7 | 6 | 6 | 8 |
| ZDHHC20 | chr13:21378661-21378738 | 7 | 17 | 21 | 18 | 18 | 15 | 16 | 11 | 12 | 21 | 15 | 33 | 42 | 26 | 26 | 7 | 15 | 22 | 9 | 3 | 4 | 12 | 23 | 23 | 20 | 21 | 15 | 20 | 13 | 13 | 13 | 12 | 35 | 17 | 15 | 6 | 6 | 12 | 31 | 35 | 20 | 7 | 5 | 19 | 21 | 22 | 78 | 21 | 14 | 17 | 17 |
| SMC6 | chr2:17752978-17753064 | 15 | 22 | 20 | 17 | 14 | 12 | 3 | 17 | 23 | 28 | 25 | 11 | 12 | 20 | 24 | 26 | 22 | 28 | 24 | 22 | 28 | 11 | -1 | 19 | 15 | 18 | 17 | 17 | -1 | 17 | 19 | 20 | 17 | 16 | 14 | 20 | 19 | 16 | 17 | 22 | 11 | 16 | 12 | 18 | 15 | 18 | 54 | 18 | 17 | 14 | 14 |
| OTUD5 | chrX:48936191-48936358\|48936355 | 6 | 9 | 4 | 6 | 6 | 7 | 7 | 8 | 2 | 1 | 2 | 9 | 11 | 2 | 1 | 2 | 1 | 2 | 2 | 2 | 2 | 9 | 10 | 9 | 7 | 8 | 4 | 8 | 8 | 13 | 12 | 7 | 6 | 9 | 6 | 8 | 9 | 11 | 5 | 7 | 9 | 5 | 4 | 10 | 12 | 7 | 26 | 6 | 11 | 9 | 13 |
| UGGT2 | chr13:95943667-95943723 | 15 | 14 | 13 | 8 | 9 | 10 | 13 | 17 | 14 | 10 | 15 | 6 | 25 | 13 | 7 | 22 | 12 | 18 | 17 | 32 | 27 | 3 | 9 | 13 | 11 | 13 | 19 | 11 | 10 | 13 | 12 | 13 | 17 | 14 | 15 | 23 | 27 | 12 | 26 | 16 | 15 | 17 | 16 | 19 | 23 | 19 | 53 | 13 | 12 | 16 | 18 |
| AC006116.20 | chr19:56302842-56302983\|56304601 | 1 | 1 | 0 | 1 | 0 | 0 | 0 | 1 | -1 | -1 | 18 | 54 | 49 | -1 | -1 | -1 | 0 | 3 | 13 | -1 | -1 | 0 | -1 | 0 | 0 | 0 | 1 | 0 | 0 | 8 | 23 | 0 | 0 | 0 | 2 | 1 | 2 | 0 | 0 | 1 | 1 | 1 | 1 | 0 | 0 | 0 | 67 | 0 | 0 | 1 | 0 |
| AGPAT4 | chr6:161165577-161165713 | 17 | 16 | 7 | 7 | 9 | 7 | 5 | 14 | 10 | 5 | 11 | 4 | 4 | 9 | 4 | 12 | 7 | 7 | 11 | 11 | 11 | 7 | 6 | 6 | 8 | 7 | 13 | 6 | 7 | 15 | 13 | 13 | 22 | 21 | 13 | 17 | 15 | 9 | 10 | 6 | 5 | 13 | 12 | 11 | 21 | 7 | 56 | 12 | 6 | 9 | 34 |
| RAB11FIP2 | chr10:118033982-118034041 | 19 | 14 | 10 | 17 | 16 | 19 | 15 | 16 | 11 | 13 | 9 | 19 | 15 | 14 | 16 | 12 | 11 | 12 | 12 | 7 | 11 | 17 | 14 | 17 | 17 | 16 | 16 | 14 | 18 | 15 | 16 | 16 | 14 | 13 | 13 | 20 | 14 | 14 | 11 | 11 | 17 | 17 | 17 | 15 | 12 | 15 | 32 | 15 | 20 | 17 | 14 |
| WSCD2 | chr12:108244304-108244581 | 21 | 16 | 15 | 7 | 8 | 10 | 18 | 20 | 5 | 3 | 3 | 23 | 24 | 4 | 3 | 4 | 9 | 4 | 4 | 13 | 5 | 3 | -1 | 10 | 9 | 10 | 19 | 12 | 11 | 11 | 10 | 11 | -1 | 18 | 28 | 17 | 17 | 7 | 11 | 19 | 11 | 22 | 21 | 12 | -1 | 12 | 63 | 22 | 13 | 11 | 12 |
| KCNK6 | chr19:38325043\|38324384-38325577 | 12 | 14 | 14 | 20 | 22 | 20 | 23 | 14 | 10 | 13 | 11 | 22 | 24 | 22 | 23 | 9 | 11 | 9 | 13 | 12 | 8 | 21 | 16 | 20 | 21 | 15 | 22 | 17 | 13 | 14 | 27 | 15 | 15 | 19 | 18 | 12 | 13 | 21 | 16 | 24 | 19 | 16 | 16 | 22 | 16 | 17 | 47 | 22 | 19 | 20 | 14 |
| KLHL3 | chr5:137629243-137629644 | 11 | 10 | 3 | 11 | 6 | 10 | -1 | 8 | 4 | 2 | 1 | 2 | 2 | 2 | 0 | 3 | 3 | 1 | 2 | 4 | 3 | -1 | -1 | 6 | 5 | 7 | 6 | 5 | -1 | 4 | 1 | 2 | 5 | 9 | 7 | 14 | 8 | 8 | 4 | 4 | 8 | 9 | 12 | 5 | 4 | 5 | 34 | 5 | 9 | 10 | 7 |
| NFATC3 | chr16:68221231\|68221196-68221284 | 7 | 10 | 9 | 7 | 7 | 5 | 7 | 8 | 5 | 4 | 5 | 9 | 9 | 5 | 5 | 4 | 4 | 5 | 4 | 5 | 4 | 5 | 5 | 8 | 9 | 8 | 9 | 8 | 10 | 7 | 9 | 8 | 12 | 10 | 8 | 7 | 5 | 7 | 10 | 5 | 8 | 6 | 6 | 12 | 13 | 9 | 22 | 7 | 7 | 6 | 9 |
| ATRNL1 | chr10:115281031-115281103 | 7 | 7 | 3 | 4 | 4 | 2 | 0 | 12 | 3 | 2 | 2 | 2 | 1 | 3 | 2 | 2 | 2 | 3 | 2 | 7 | 3 | -1 | 0 | 4 | 4 | 4 | 4 | 4 | -1 | 3 | 4 | 5 | 0 | 5 | 5 | 2 | 0 | 6 | 3 | 5 | 4 | -1 | 5 | 3 | 0 | 4 | 24 | 8 | 7 | 6 | 0 |
| MFHAS1 | chr8:8790324-8790434 | 10 | 12 | 11 | 8 | 10 | 8 | 8 | 9 | 15 | 15 | 17 | 17 | 19 | 16 | 14 | 14 | 16 | 16 | 14 | 21 | 15 | 10 | 8 | 11 | 10 | 10 | 9 | 10 | 13 | 7 | 7 | 11 | 14 | 9 | 12 | 10 | 11 | 10 | 15 | 17 | 12 | 9 | 9 | 10 | 11 | 11 | 29 | 14 | 8 | 9 | 9 |
| CXorf40B | chrX:149934377-149934573 | 9 | 4 | 3 | 5 | 3 | 7 | 6 | 6 | 3 | 4 | 3 | 5 | 3 | 3 | 4 | 4 | 3 | 2 | 5 | 6 | 4 | 6 | 4 | 5 | 4 | 6 | 3 | 5 | 5 | 4 | 4 | 7 | 4 | 8 | 6 | 6 | 7 | 4 | 7 | 3 | 5 | 8 | 14 | 5 | 4 | 4 | 21 | 8 | 5 | 6 | 5 |
| SLC38A6 | chr14:61048151-61048297\|61050074 | 22 | 13 | 8 | 13 | 14 | 15 | 17 | 20 | 19 | 16 | 16 | 19 | 19 | 16 | 15 | 17 | 18 | 19 | 19 | 20 | 12 | 22 | 14 | 14 | 14 | 13 | 12 | 13 | 24 | 22 | 20 | 15 | 12 | 16 | 17 | 25 | 24 | 17 | 10 | 19 | 18 | 22 | 26 | 16 | 19 | 12 | 49 | 14 | 15 | 16 | 12 |
| ATE1-AS1 | chr10:121949412-121949487 | 14 | 12 | 14 | 10 | 10 | 13 | -1 | 9 | 8 | 7 | 9 | 5 | 5 | 6 | 9 | 8 | 9 | 6 | 12 | 12 | 14 | -1 | -1 | 8 | 12 | 14 | 16 | 11 | -1 | 9 | 13 | 17 | 18 | 16 | 22 | 9 | 11 | 6 | 21 | 8 | 9 | 13 | 13 | 18 | 13 | 7 | 53 | 11 | 7 | 12 | 21 |
| ZNF333 | chr19:14700206\|14701581-14701897 | 12 | 15 | 10 | 12 | 12 | 12 | 13 | 13 | 13 | 10 | 10 | 16 | 17 | 11 | 9 | 12 | 10 | 13 | 11 | 13 | 12 | 14 | 17 | 15 | 15 | 14 | 16 | 14 | 10 | 13 | 13 | 12 | 14 | 16 | 13 | 16 | 14 | 17 | 21 | 16 | 19 | 13 | 14 | 14 | 16 | 15 | 30 | 15 | 17 | 15 | 15 |
| CDC42BPG | chr11:64828655-64828749 | 4 | 3 | 1 | 3 | 3 | 5 | 3 | 4 | 2 | 0 | 9 | 19 | 20 | 14 | -1 | 2 | 11 | 10 | -1 | 16 | -1 | 3 | -1 | 4 | 3 | 2 | 2 | 6 | 3 | 5 | 4 | 4 | 4 | 2 | 3 | 8 | 6 | 3 | 3 | 6 | 3 | 3 | 3 | 4 | 3 | 2 | 26 | 4 | 3 | 2 | 2 |
| WDFY3-AS2 | chr4:85000691-85000765 | 21 | 13 | 9 | 16 | 16 | 21 | 16 | 19 | 23 | 25 | 20 | 27 | 25 | 23 | 25 | 26 | 21 | 19 | 21 | 38 | 26 | -1 | -1 | 16 | 17 | 15 | 16 | 14 | -1 | 14 | 14 | 21 | 16 | 16 | 13 | 16 | 24 | 13 | 26 | 15 | 17 | 17 | 18 | 17 | 10 | 18 | 59 | 17 | 18 | 14 | 8 |
| KDM4A | chr1:43700535-43700759 | 10 | 4 | 1 | 2 | 3 | 4 | 4 | 7 | 3 | 2 | 2 | 1 | 1 | 1 | 1 | 2 | 2 | 1 | 2 | 4 | 3 | 3 | 3 | 2 | 2 | 1 | 3 | 2 | 6 | 3 | 1 | 3 | 2 | 4 | 6 | 5 | 9 | 4 | 4 | 3 | 4 | 10 | 14 | 4 | 4 | 3 | 24 | 4 | 2 | 4 | 5 |
| PHOSPHO1 | chr17:49225656-49225777 | 2 | 1 | 0 | 0 | 1 | 4 | -1 | 7 | 3 | 3 | 5 | 2 | 5 | 4 | 4 | 3 | 2 | 1 | 4 | 2 | 3 | -1 | -1 | 3 | -1 | 2 | 4 | 1 | 0 | 1 | 0 | 1 | 6 | 0 | -1 | 1 | 6 | 0 | -1 | 5 | 2 | 10 | 12 | 3 | 0 | -1 | 24 | 2 | 0 | 0 | 0 |
| INPP1 | chr2:190348888-190349031 | 17 | 15 | 12 | 14 | 13 | 17 | 14 | 16 | 11 | 11 | 11 | 16 | 13 | 10 | 10 | 11 | 10 | 10 | 10 | 12 | 13 | 13 | 10 | 15 | 15 | 15 | 13 | 15 | 9 | 16 | 18 | 13 | 21 | 17 | 14 | 23 | 13 | 12 | 21 | 13 | 15 | 17 | 19 | 16 | 14 | 15 | 37 | 15 | 15 | 15 | 23 |
| DAAM2 | chr6:39886421-39886447 | 2 | 3 | 15 | 1 | 1 | 1 | 3 | 2 | 29 | 23 | 28 | 18 | 21 | 22 | 24 | 26 | 25 | 31 | 31 | 19 | 24 | 0 | 1 | 1 | 2 | 1 | 1 | 1 | 2 | 3 | 2 | 3 | 4 | 3 | 9 | 0 | 1 | 1 | 17 | 11 | 1 | 2 | 2 | 2 | 6 | 3 | 39 | 5 | 1 | 1 | 10 |
| TNK2 | chr3:195885413-195885543\|195886774 | 9 | 10 | 6 | 9 | 10 | 9 | 11 | 11 | 4 | 3 | 4 | 5 | 5 | 3 | 3 | 5 | 5 | 4 | 4 | 10 | 6 | 9 | 10 | 9 | 10 | 8 | 12 | 8 | 14 | 11 | 10 | 12 | 12 | 13 | 12 | 15 | 12 | 8 | 9 | 10 | 13 | 15 | 15 | 15 | 17 | 9 | 29 | 11 | 9 | 12 | 14 |
| LTN1 | chr21:28961078-28964766 | 19 | 16 | 7 | 12 | 10 | 12 | 11 | 23 | 5 | 6 | 9 | 11 | 13 | 7 | 4 | 6 | 7 | 10 | 5 | 8 | 9 | 25 | -1 | 12 | 17 | 9 | 13 | 8 | 21 | 10 | 5 | 23 | 18 | 19 | 18 | 5 | 27 | 29 | 28 | 14 | 26 | 17 | 20 | 30 | 24 | 19 | 68 | 22 | 20 | 18 | 18 |
| POLG | chr15:89332218-89332374 | 11 | 17 | 12 | 16 | 15 | 12 | 14 | 12 | 12 | 10 | 15 | 17 | 18 | 12 | 12 | 12 | 14 | 15 | 12 | 11 | 13 | 11 | 18 | 17 | 14 | 13 | 13 | 14 | 14 | 14 | 10 | 17 | 14 | 12 | 15 | 3 | 13 | 18 | 26 | 28 | 21 | 11 | 10 | 14 | 18 | 15 | 46 | 23 | 15 | 16 | 11 |
| FRA10AC1 | chr10:93672042-93675701\|93672089\|93673376 | 9 | 6 | 3 | 6 | 5 | 6 | 6 | 9 | 1 | 1 | 1 | 1 | 1 | 1 | 1 | 1 | 1 | 1 | 1 | 1 | 1 | 9 | 3 | 5 | 5 | 5 | 3 | 5 | 8 | 3 | 3 | 4 | 7 | 8 | 5 | 3 | 5 | 5 | 6 | 3 | 6 | 7 | 7 | 8 | 8 | 6 | 20 | 5 | 4 | 5 | 10 |
| STRADB | chr2:201457379-201457450 | 9 | 21 | 17 | 14 | 24 | 8 | -1 | 19 | 11 | 12 | 12 | 3 | 11 | 7 | 2 | 9 | 11 | 14 | 11 | 7 | 5 | 0 | -1 | 14 | 22 | 11 | 17 | 10 | -1 | 16 | 9 | 29 | 19 | 10 | 19 | 9 | 14 | 8 | 14 | 13 | 10 | 15 | 11 | 31 | 17 | 15 | 64 | 7 | 7 | 10 | 12 |
| GTF2IP1 | chr7:75201812-75201923 | 9 | 4 | 2 | 3 | 5 | 5 | 0 | 7 | 3 | 3 | 2 | 0 | 1 | 2 | 2 | 4 | 1 | 1 | 3 | 1 | 2 | 0 | 0 | 7 | 3 | 5 | 2 | 4 | 0 | 4 | 5 | -1 | -1 | 5 | 2 | 10 | 6 | 5 | 3 | 2 | 5 | 5 | 10 | 8 | 2 | 5 | 24 | 2 | 3 | 2 | 3 |
| BRD4 | chr19:15268154-15268281\|15268240 | 10 | 10 | 8 | 9 | 9 | 9 | 11 | 11 | 7 | 8 | 6 | 10 | 11 | 7 | 6 | 7 | 6 | 7 | 7 | 6 | 6 | 11 | 9 | 9 | 10 | 8 | 11 | 8 | 13 | 8 | 7 | 12 | 9 | 10 | 12 | 4 | 12 | 10 | 15 | 12 | 11 | 11 | 12 | 13 | 14 | 10 | 25 | 13 | 10 | 11 | 8 |
| BRE | chr2:28310082-28310153 | 10 | 9 | 14 | 8 | 9 | 8 | 10 | 9 | 8 | 8 | 8 | 8 | 8 | 8 | 8 | 8 | 9 | 9 | 8 | 11 | 9 | 5 | 7 | 9 | 10 | 9 | 10 | 9 | 9 | 13 | 16 | 10 | 12 | 8 | 11 | 8 | 9 | 8 | 12 | 13 | 11 | 9 | 9 | 10 | 12 | 10 | 25 | 10 | 7 | 8 | 11 |
| BACH1 | chr21:29305204-29305322 | 17 | 12 | 7 | 10 | 9 | 12 | 9 | 16 | 12 | 7 | 11 | 8 | 8 | 9 | 14 | 9 | 10 | 13 | 12 | 9 | 13 | 7 | 11 | 9 | 8 | 8 | 8 | 8 | 13 | 10 | 7 | 11 | 8 | 17 | 10 | 12 | 17 | 11 | 10 | 9 | 8 | 17 | 18 | 11 | 9 | 12 | 33 | 10 | 9 | 9 | 16 |
| MYNN | chr3:169774031-169774097\|169774141 | 13 | 15 | 14 | 11 | 12 | 8 | 16 | 12 | 10 | 10 | 11 | 10 | 9 | 12 | 13 | 8 | 12 | 10 | 10 | 7 | 10 | 13 | 15 | 13 | 15 | 12 | 16 | 12 | 13 | 9 | 9 | 14 | 19 | 16 | 14 | 11 | 10 | 7 | 20 | 11 | 13 | 9 | 10 | 19 | 24 | 18 | 38 | 12 | 9 | 9 | 13 |
| BRE | chr2:27892383-27892448 | 9 | 7 | 5 | 5 | 6 | 6 | 5 | 8 | 8 | 5 | 6 | 4 | 4 | 5 | 5 | 6 | 5 | 4 | 5 | 8 | 7 | 6 | 6 | 5 | 7 | 5 | 7 | 5 | 6 | 5 | 5 | 10 | 10 | 9 | 10 | 12 | 7 | 5 | 13 | 5 | 8 | 8 | 9 | 8 | 7 | 9 | 21 | 8 | 5 | 6 | 8 |
| LRRC28 | chr15:99360218\|99355406-99360286 | 1 | 1 | 1 | 1 | 1 | 1 | 1 | 1 | 12 | 15 | 10 | 27 | 27 | 14 | 15 | 12 | 10 | 13 | 11 | 19 | 12 | 2 | 2 | 2 | 1 | 1 | 1 | 1 | 1 | 3 | 2 | 3 | 2 | 1 | 2 | 2 | 3 | 3 | 1 | 4 | 2 | 1 | 1 | 3 | 2 | 2 | 31 | 2 | 2 | 1 | 2 |
| MFF | chr2:227329606-227329783\|227330366 | 17 | 17 | 8 | 25 | 22 | 22 | 17 | 17 | 15 | 21 | 14 | 22 | 23 | 26 | 28 | 17 | 21 | 12 | 15 | 25 | 21 | 23 | 22 | 21 | 18 | 15 | 16 | 13 | 17 | 26 | 28 | 12 | 12 | 16 | 16 | 78 | 20 | 20 | 11 | 16 | 16 | 18 | 17 | 17 | 14 | 14 | 92 | 13 | 21 | 20 | 8 |
| PPP1R11 | chr6:30068279-30068353\|30068430 | 12 | 17 | 16 | 15 | 16 | 9 | 22 | 23 | 19 | 15 | 16 | 6 | 16 | 14 | 5 | 23 | 18 | 14 | 18 | 23 | 19 | 17 | 21 | 19 | 17 | 17 | 18 | 18 | 19 | 16 | 7 | 14 | 12 | 12 | 18 | 12 | 21 | 20 | 7 | 19 | 19 | 27 | 16 | 19 | 21 | 18 | 47 | 9 | 18 | 18 | 13 |
| PRIMA1 | chr14:93725836-93726006\|93733655\|93730434 | 10 | 12 | 14 | 16 | 23 | 11 | 8 | 9 | 69 | 60 | 37 | 16 | 14 | 64 | 62 | 75 | 61 | 36 | 41 | 69 | 66 | 12 | 15 | 19 | 15 | 17 | 14 | 16 | 10 | 19 | 28 | 12 | 21 | 11 | 8 | 25 | 21 | 8 | 18 | 26 | 19 | 7 | 6 | 14 | -1 | 13 | 57 | 16 | 9 | 12 | -1 |
| RMDN1 | chr8:86468863\|86469424-86469479 | 8 | 11 | 6 | 12 | 15 | 12 | -1 | 11 | 9 | 11 | 15 | 16 | 18 | 10 | 13 | 9 | 10 | 17 | 17 | 13 | 14 | -1 | 0 | 9 | 10 | 9 | 9 | 11 | 0 | 11 | 15 | 8 | 6 | 12 | 7 | 21 | 11 | 16 | 12 | 6 | 8 | 10 | 6 | 12 | 17 | 12 | 48 | 7 | 24 | 7 | 19 |
| TTC21A | chr3:39126261-39126390 | 31 | 27 | 30 | 24 | 21 | 27 | 26 | 30 | 18 | 13 | 15 | 10 | 12 | 12 | 8 | 18 | 16 | 11 | 9 | 12 | 16 | -1 | -1 | 23 | 29 | 22 | 32 | 25 | 39 | 26 | 33 | 30 | 35 | 44 | 31 | 25 | 20 | 17 | 25 | 18 | 21 | 33 | 32 | 41 | 34 | 33 | 94 | 21 | 18 | 20 | 49 |
| TCERG1 | chr5:146481447\|146481150-146482004 | 8 | 7 | 3 | 6 | 5 | 6 | 10 | 9 | 5 | 9 | 5 | 12 | 14 | 10 | 8 | 5 | 8 | 8 | 6 | 6 | 6 | 7 | 6 | 7 | 6 | 5 | 5 | 5 | 9 | 5 | 5 | 5 | 4 | 7 | 5 | 9 | 7 | 12 | 5 | 6 | 7 | 8 | 6 | 7 | 6 | 6 | 21 | 5 | 6 | 6 | 3 |
| TLK1 | chr2:171081640-171081708 | 11 | 11 | 8 | 9 | 9 | 9 | 8 | 12 | 12 | 9 | 10 | 8 | 9 | 8 | 6 | 11 | 7 | 10 | 11 | 14 | 12 | 12 | 8 | 9 | 12 | 8 | 10 | 8 | 13 | 10 | 9 | 11 | 9 | 11 | 9 | 12 | 12 | 10 | 12 | 10 | 10 | 12 | 12 | 13 | 15 | 10 | 22 | 9 | 9 | 10 | 11 |
| HIBCH | chr2:190260400-190260511 | 4 | 2 | 1 | 6 | 5 | 6 | 5 | 5 | 3 | 2 | 2 | 10 | 14 | 3 | 2 | 3 | 2 | 2 | 2 | 3 | 2 | 5 | 9 | 4 | 2 | 3 | 2 | 3 | 5 | 3 | 2 | 1 | 1 | 4 | 4 | 3 | 7 | 7 | 4 | 6 | 7 | 5 | 5 | 4 | 4 | 3 | 21 | 5 | 8 | 5 | 2 |
| RIMKLB | chr12:8776767\|8775331-8777061 | 6 | 9 | 8 | 6 | 8 | 6 | 5 | 6 | 11 | 11 | 9 | 8 | 9 | 13 | 10 | 9 | 10 | 10 | 9 | 16 | 10 | 8 | 6 | 5 | 6 | 5 | 8 | 6 | 3 | 16 | 10 | 7 | 13 | 12 | 4 | 14 | 6 | 5 | 7 | 10 | 6 | 7 | 6 | 6 | 8 | 7 | 27 | 5 | 7 | 7 | 13 |
| CEP112 | chr17:66043096\|66043106-66043250 | 12 | 10 | 9 | 9 | 9 | 9 | 10 | 12 | 7 | 5 | 5 | 7 | 5 | 8 | 6 | 5 | 4 | 5 | 7 | 5 | 5 | 16 | 7 | 9 | 9 | 9 | 11 | 8 | 12 | 9 | 7 | 11 | 14 | 12 | 13 | 5 | 10 | 11 | 14 | 13 | 11 | 14 | 13 | 12 | 15 | 9 | 27 | 13 | 11 | 10 | 7 |
| CDC42BPA | chr1:227051881-227051985 | 30 | 18 | 9 | 14 | 13 | 17 | 11 | 26 | 12 | 26 | 12 | 11 | 12 | 22 | 27 | 12 | 9 | 19 | 10 | 6 | 6 | 19 | 8 | 13 | 19 | 12 | 19 | 10 | 20 | 10 | 9 | 13 | 16 | 23 | 18 | 11 | 30 | 18 | 12 | 15 | 15 | 28 | 30 | 19 | 12 | 17 | 59 | 15 | 17 | 20 | 29 |
| TTC38 | chr22:46281971-46282151 | 10 | 9 | 6 | 7 | 7 | 6 | 7 | 10 | 7 | 6 | 6 | 13 | 11 | 6 | 6 | 7 | 6 | 6 | 6 | 7 | 7 | 7 | 8 | 9 | 6 | 7 | 8 | 7 | 12 | 8 | 5 | 10 | 15 | 8 | 10 | 5 | 10 | 14 | 8 | 10 | 9 | 10 | 12 | 12 | 10 | 8 | 24 | 11 | 12 | 12 | 7 |
| CEP63 | chr3:134558142-134559429\|134559305 | 27 | 18 | 10 | 17 | 18 | 28 | 20 | 27 | 12 | 9 | 12 | 11 | 12 | 9 | 7 | 12 | 12 | 11 | 10 | 10 | 9 | 36 | 26 | 18 | 18 | 14 | 16 | 13 | 33 | 8 | 5 | 14 | 13 | 20 | 20 | 10 | 28 | 23 | 16 | 15 | 22 | 32 | 33 | 24 | 20 | 17 | 71 | 15 | 25 | 25 | 35 |
| CEP63 | chr3:134558142-134559429\|134559305 | 27 | 18 | 10 | 17 | 18 | 28 | 20 | 27 | 12 | 9 | 12 | 11 | 12 | 9 | 7 | 12 | 12 | 11 | 10 | 10 | 9 | 36 | 26 | 18 | 18 | 14 | 16 | 13 | 33 | 8 | 5 | 14 | 13 | 20 | 20 | 10 | 28 | 23 | 16 | 15 | 22 | 32 | 33 | 24 | 20 | 17 | 71 | 15 | 25 | 25 | 35 |
| TCF7 | chr5:134144797-134144869\|134145340 | 5 | 7 | 5 | 6 | 4 | 5 | 3 | 3 | 61 | 68 | 53 | 59 | 56 | 73 | 72 | 56 | 52 | 64 | 69 | 54 | 53 | -1 | -1 | 5 | 4 | 3 | 1 | 4 | 2 | 44 | 50 | 9 | 3 | 6 | 2 | 8 | 10 | 4 | 0 | 11 | 3 | 3 | 2 | 3 | 2 | 2 | 92 | 5 | 3 | 4 | 6 |
| ZC2HC1A | chr8:78725142-78725276 | 9 | 10 | 10 | 10 | 11 | 8 | 0 | 8 | 15 | 9 | 10 | 9 | 5 | 14 | 4 | 8 | 7 | 13 | 18 | 6 | 9 | -1 | -1 | 9 | 7 | 6 | 4 | 8 | -1 | 7 | 8 | -1 | 5 | 6 | 5 | 5 | 16 | 13 | 6 | 9 | 8 | 4 | 5 | 6 | 10 | 9 | 32 | 9 | 9 | 6 | 2 |
| HIST1H2BN | chr6:27845227\|27843539-27845357 | 17 | 20 | 12 | 14 | 13 | 11 | -1 | 26 | -1 | -1 | 20 | 10 | -1 | 6 | 5 | 12 | 18 | 21 | -1 | 7 | 7 | 0 | 0 | 24 | 25 | 5 | 27 | 9 | -1 | 19 | 10 | 12 | 27 | 20 | 29 | 11 | 13 | 17 | 42 | 19 | 39 | 15 | 16 | 18 | 21 | 16 | 82 | 20 | 25 | 25 | 14 |
| DNAJC21 | chr5:34949543-34949677 | 12 | 11 | 11 | 8 | 9 | 8 | 9 | 11 | 14 | 16 | 14 | 12 | 15 | 16 | 14 | 13 | 15 | 14 | 14 | 15 | 13 | 14 | 15 | 7 | 8 | 8 | 9 | 8 | 10 | 16 | 15 | 11 | 12 | 10 | 10 | 13 | 12 | 15 | 9 | 16 | 8 | 11 | 12 | 8 | 11 | 8 | 27 | 9 | 10 | 10 | 12 |
| MVD | chr16:88657146-88657247 | 6 | 5 | 4 | 6 | 6 | 5 | 7 | 7 | 9 | 8 | 8 | 9 | 9 | 7 | 8 | 8 | 6 | 9 | 8 | 7 | 7 | 11 | 4 | 5 | 8 | 5 | 6 | 4 | 6 | 5 | 5 | 14 | 7 | 10 | 8 | 7 | 10 | 3 | 10 | 11 | 7 | 12 | 14 | 9 | 6 | 7 | 26 | 15 | 4 | 8 | 8 |
| EFCAB13 | chr17:47347808-47347951 | 10 | 12 | 12 | 14 | 14 | 12 | 10 | 9 | 9 | 5 | 10 | 6 | 2 | 7 | 6 | 11 | 12 | 9 | 4 | 5 | 7 | -1 | -1 | 12 | 13 | 10 | 11 | 12 | -1 | 12 | 10 | 18 | 19 | 15 | 13 | 9 | 10 | 6 | 17 | 11 | 12 | 7 | 8 | 13 | 12 | 11 | 41 | 16 | 7 | 9 | 17 |
| E2F6 | chr2:11455342-11455474 | 14 | 16 | 15 | 14 | 13 | 12 | 14 | 15 | 15 | 14 | 11 | 12 | 12 | 12 | 10 | 14 | 14 | 11 | 12 | 18 | 13 | 18 | 13 | 16 | 16 | 15 | 17 | 16 | 10 | 17 | 16 | 17 | 24 | 12 | 15 | 17 | 14 | 13 | 19 | 12 | 16 | 13 | 14 | 16 | 18 | 16 | 32 | 16 | 13 | 14 | 10 |
| CTD-2554C21.1 | chr19:37785219-37785354 | 5 | 3 | 3 | 1 | 2 | 2 | -1 | 3 | 0 | 1 | 1 | 1 | 1 | 1 | 2 | 0 | 1 | 1 | 1 | 1 | 0 | 0 | -1 | 3 | 2 | 1 | 3 | 3 | -1 | 2 | 2 | 1 | 8 | 4 | 8 | 4 | 2 | 2 | 8 | 2 | 13 | 1 | 3 | 2 | 7 | 5 | 21 | 4 | 2 | 4 | 2 |
| AC144449.1 | chr2:149843388-149843447 | 21 | 8 | 7 | 9 | 9 | 15 | -1 | 12 | 3 | 3 | 11 | 2 | 3 | 9 | 5 | 9 | 8 | 7 | 6 | 11 | 10 | -1 | -1 | 9 | 19 | 14 | 15 | 12 | 0 | 12 | 13 | 6 | 12 | 9 | 13 | 14 | 11 | 14 | -1 | 1 | 14 | 24 | 33 | 26 | 12 | 21 | 62 | 17 | 8 | 13 | -1 |
| NR1D2 | chr3:23963115\|23963183\|23963224-23963330 | 8 | 6 | 3 | 5 | 5 | 7 | 12 | 8 | 1 | 1 | 1 | 1 | 2 | 1 | 1 | 1 | 1 | 1 | 1 | 1 | 2 | 8 | 8 | 7 | 8 | 5 | 5 | 6 | 14 | 3 | 2 | 5 | 4 | 7 | 7 | 4 | 7 | 11 | 5 | 3 | 5 | 12 | 10 | 9 | 7 | 6 | 24 | 5 | 6 | 8 | 7 |
| FAM114A2 | chr5:154038293\|154038388-154038434 | 16 | 20 | 8 | 11 | 13 | 10 | 18 | 16 | 8 | 8 | 10 | 14 | 16 | 10 | 12 | 13 | 11 | 7 | 7 | 9 | 9 | 14 | 11 | 16 | 14 | 15 | 19 | 16 | 10 | 12 | 13 | 18 | 22 | 19 | 13 | 9 | 13 | 12 | 15 | 9 | 14 | 12 | 12 | 19 | 19 | 14 | 36 | 16 | 14 | 16 | 20 |
| KLF3-AS1 | chr4:38620072-38620162 | 8 | 7 | 8 | 3 | 3 | 3 | 6 | 7 | 36 | 29 | 30 | 41 | 39 | 26 | 26 | 29 | 27 | 32 | 29 | 26 | 30 | 11 | 9 | 5 | 5 | 5 | 3 | 5 | 7 | 8 | 7 | 7 | 4 | 7 | 4 | 7 | 14 | 12 | 3 | 6 | 4 | 4 | 5 | 7 | 9 | 5 | 44 | 5 | 9 | 5 | 7 |
| PLEKHA6 | chr1:204271248-204271374 | 14 | 6 | 4 | 3 | 4 | 7 | 5 | 11 | 2 | 2 | 1 | 2 | 2 | 1 | 1 | 1 | 2 | 1 | 1 | 2 | 1 | 0 | 0 | 6 | 7 | 4 | 4 | 6 | -1 | 5 | 5 | 4 | 9 | 7 | 6 | 11 | 8 | 5 | 4 | 3 | 5 | 9 | 13 | 12 | 14 | 7 | 29 | 5 | 6 | 4 | 5 |
| ENPP3 | chr6:131651500-131651609 | 2 | 5 | 1 | 4 | 4 | 7 | -1 | 4 | 0 | -1 | 10 | 6 | 6 | 8 | 15 | -1 | 12 | -1 | -1 | 0 | -1 | 0 | -1 | 2 | 1 | 4 | 4 | 5 | 0 | 2 | 2 | 1 | 2 | 2 | 1 | 1 | 6 | 4 | 16 | 0 | 5 | 2 | 2 | 1 | 0 | 6 | 22 | 2 | 3 | 4 | 1 |
| DDX60L | chr4:168426771-168426909 | 15 | 17 | 10 | 16 | 14 | 13 | 19 | 16 | 11 | 8 | 11 | 13 | 19 | 17 | 7 | 6 | 9 | 10 | 11 | 5 | 13 | 14 | -1 | 18 | 15 | 17 | 15 | 17 | 19 | 17 | 10 | 15 | 11 | 11 | 16 | 14 | 17 | 22 | 13 | 17 | 15 | 15 | 13 | 15 | 16 | 11 | 41 | 18 | 19 | 17 | 5 |
| RIC1 | chr9:5748585-5748714 | 1 | 2 | 1 | 1 | 2 | 1 | 2 | 1 | 6 | 6 | 5 | 16 | 21 | 7 | 2 | 5 | 5 | 5 | 5 | 7 | 7 | 2 | 1 | 2 | 2 | 2 | 1 | 2 | 1 | 5 | 5 | 1 | 2 | 1 | 2 | 6 | 2 | 3 | 3 | 5 | 4 | 1 | 1 | 2 | 1 | 2 | 24 | 2 | 1 | 1 | 1 |
| TBC1D19 | chr4:26722851-26722890 | 9 | 6 | 3 | 4 | 6 | 5 | 6 | 9 | 2 | 0 | 2 | 3 | 2 | 1 | 1 | 1 | 1 | 2 | 1 | 2 | 1 | 4 | 6 | 5 | 6 | 5 | 7 | 4 | 5 | 3 | 3 | 7 | 7 | 6 | 11 | 3 | 6 | 4 | 11 | 5 | 8 | 10 | 11 | 11 | 14 | 5 | 25 | 7 | 5 | 6 | 14 |
| PRKD1 | chr14:29656476-29656499 | 2 | 3 | 5 | 3 | 3 | 3 | 17 | 3 | 16 | 22 | 17 | 15 | 14 | 21 | 29 | 18 | 28 | 18 | 21 | 13 | 18 | 6 | 4 | 9 | 8 | 7 | 7 | 9 | 7 | 6 | 4 | 4 | 5 | 2 | 4 | 2 | 3 | 8 | 5 | 17 | 7 | 2 | 1 | 4 | 7 | 7 | 29 | 5 | 6 | 5 | 1 |
| ANO2 | chr12:5772361-5772420 | 12 | 8 | 3 | 8 | 8 | 11 | 0 | 9 | 7 | 4 | 4 | 5 | 2 | 8 | 7 | 2 | 9 | 3 | 3 | 7 | 6 | -1 | 0 | 6 | 7 | 7 | 6 | 10 | -1 | 6 | 6 | 2 | -1 | 7 | 7 | 10 | 9 | 9 | 6 | 4 | 6 | 10 | 14 | 7 | 8 | 6 | 29 | 8 | 6 | 5 | 5 |
| PPP6R2 | chr22:50388986\|50389034-50389111 | 6 | 10 | 13 | 7 | 9 | 7 | 11 | 9 | 15 | 14 | 16 | 10 | 10 | 13 | 14 | 12 | 13 | 14 | 15 | 11 | 15 | 5 | 7 | 8 | 12 | 7 | 13 | 8 | 9 | 7 | 8 | 14 | 21 | 11 | 10 | 6 | 7 | 7 | 15 | 14 | 11 | 8 | 7 | 11 | 10 | 11 | 30 | 14 | 8 | 11 | 11 |
| TAOK2 | chr16:29990543-29990647 | 8 | 15 | 9 | 9 | 10 | 5 | 7 | 10 | 8 | 6 | 8 | 8 | 10 | 6 | 5 | 9 | 8 | 7 | 6 | 13 | 11 | 9 | 12 | 10 | 12 | 10 | 12 | 9 | 8 | 15 | 15 | 18 | 13 | 12 | 14 | 13 | 9 | 14 | 14 | 18 | 17 | 8 | 8 | 16 | 23 | 12 | 37 | 19 | 14 | 12 | 21 |
| CDADC1 | chr13:49265879\|49261178-49266003 | 26 | 16 | 10 | 15 | 15 | 19 | 14 | 24 | 6 | 5 | 6 | 8 | 6 | 5 | 5 | 7 | 6 | 6 | 6 | 9 | 5 | 32 | 24 | 15 | 16 | 15 | 13 | 13 | 21 | 15 | 8 | 13 | 14 | 20 | 17 | 11 | 25 | 23 | 11 | 9 | 18 | 24 | 28 | 20 | 20 | 15 | 56 | 16 | 16 | 17 | 32 |
| ATAD2B | chr2:23816637-23816720 | 11 | 8 | 4 | 6 | 8 | 6 | 15 | 10 | 5 | 1 | 5 | 12 | 8 | 7 | 6 | 5 | 3 | 7 | 2 | 2 | 2 | 4 | -1 | 7 | 8 | 6 | 7 | 6 | -1 | 6 | 5 | 5 | 5 | 9 | 9 | 10 | 6 | 5 | 8 | 4 | 8 | 10 | 11 | 11 | 9 | 7 | 26 | 6 | 5 | 6 | 7 |
| FAM155A | chr13:107172529-107172574 | 13 | 6 | 6 | 7 | 7 | 7 | 6 | 9 | 7 | 7 | 7 | 6 | 5 | 6 | 7 | 5 | 8 | 8 | 6 | 8 | 7 | 0 | 0 | 6 | 9 | 6 | 8 | 5 | -1 | 7 | 5 | 3 | -1 | 6 | 9 | 9 | 14 | 7 | 5 | 7 | 5 | 6 | 10 | 4 | -1 | 7 | 27 | 7 | 3 | 8 | 0 |
| ANPEP | chr15:89802407-89802593\|89802534 | 6 | 6 | 5 | 5 | 4 | 6 | 3 | 6 | 10 | 9 | 7 | 6 | 9 | 5 | 8 | 10 | 9 | 7 | 6 | 7 | 9 | 5 | 8 | 2 | 1 | 6 | 4 | 4 | 10 | 7 | 6 | 2 | 2 | 9 | 4 | 6 | 11 | 4 | 3 | 8 | 5 | 7 | 6 | 2 | 14 | 5 | 23 | 5 | 6 | 5 | 12 |

SFig5, pg16

| Gene Symbol | Exon Location (hg38, human) | Adipose - Subcutaneous | Adipose - Visceral (Omentum) | Adrenal Gland | Artery - Aorta | Artery - Coronary | Artery - Tibial | Bladder | Breast - Mammary Tissue | Brain - Amygdala | Brain - Anterior cingulate cortex (BA24) | Brain - Caudate (basal ganglia) | Brain - Cerebellar Hemisphere | Brain - Cerebellum | Brain - Cortex | Brain - Frontal Cortex (BA9) | Brain - Hippocampus | Brain - Hypothalamus | Brain - Nucleus accumbens (basal ganglia) | Brain - Putamen (basal ganglia) | Brain - Spinal cord (cervical c-1) | Brain - Substantia nigra | Cervix - Ectocervix | Cervix - Endocervix | Colon - Sigmoid | Colon - Transverse | Esophagus - Gastroesophageal Junction | Esophagus - Mucosa | Esophagus - Muscularis | Fallopian Tube | Heart - Atrial Appendage | Heart - Left Ventricle | Kidney - Cortex | Liver | Lung | Minor Salivary Gland | Muscle - Skeletal | Nerve - Tibial | Ovary | Pancreas | Pituitary | Prostate | Skin - Not Sun Exposed (Suprapubic) | Skin - Sun Exposed (Lower leg) | Small Intestine - Terminal Ileum | Spleen | Stomach | Testis | Thyroid | Uterus | Vagina | Whole Blood |
|---|---|---|---|---|---|---|---|---|---|---|---|---|---|---|---|---|---|---|---|---|---|---|---|---|---|---|---|---|---|---|---|---|---|---|---|---|---|---|---|---|---|---|---|---|---|---|---|---|---|---|---|---|
| SYNJ1 | chr21:32670757-32670859 | 8 | 7 | 3 | 7 | 7 | 8 | 5 | 10 | 4 | 1 | 3 | 3 | 4 | 2 | 1 | 4 | 2 | 2 | 3 | 4 | 3 | -1 | -1 | 8 | 10 | 5 | 5 | 7 | -1 | 5 | 3 | 9 | 8 | 6 | 6 | 4 | 13 | 9 | 7 | 7 | 10 | 9 | 6 | 12 | 8 | 8 | 26 | 9 | 10 | 7 | 8 |
| FSD1L | chr9:105461520-105461615 | 9 | 15 | 10 | 12 | 14 | 10 | -1 | 10 | 13 | 11 | 11 | 15 | 12 | 10 | 10 | 8 | 14 | 10 | 9 | 9 | 9 | -1 | -1 | 15 | 8 | 13 | 10 | 15 | -1 | 16 | 9 | 10 | 13 | 10 | 3 | 7 | 12 | 15 | 14 | 13 | 16 | 17 | 9 | 9 | 12 | 12 | 40 | 10 | 10 | 7 | 18 |
| RP11-342M1.3 | chr1:42842497-42842685 | 14 | 7 | 5 | 9 | 5 | 7 | -1 | 11 | 5 | 6 | 3 | 9 | 5 | 4 | 6 | 3 | 5 | 3 | 2 | 2 | 8 | 0 | -1 | 7 | 8 | 11 | 8 | 7 | -1 | 7 | 5 | 8 | 7 | 5 | 9 | 5 | 6 | 7 | 6 | 4 | 9 | 11 | 18 | 8 | 10 | 10 | 30 | 8 | 8 | 7 | 9 |
| KMT2D | chr12:49024053-49024100 | 8 | 10 | 7 | 5 | 6 | 5 | 8 | 9 | 3 | 3 | 3 | 3 | 2 | 3 | 3 | 3 | 3 | 3 | 4 | 4 | 3 | 8 | 5 | 6 | 11 | 5 | 17 | 5 | 9 | 4 | 5 | 10 | 12 | 8 | 9 | 2 | 7 | 4 | 13 | 4 | 7 | 13 | 12 | 14 | 12 | 10 | 28 | 7 | 4 | 8 | 8 |
| MSH5-SAPCD1 | chr6:31745237-31745319 | 14 | 10 | 14 | 11 | 10 | 10 | 6 | 11 | 7 | 9 | 4 | 7 | 6 | 6 | 6 | 6 | 8 | 8 | 7 | 7 | 6 | 5 | 11 | 9 | 14 | 7 | 14 | 8 | 11 | 6 | 7 | 15 | 9 | 13 | 12 | 9 | 9 | 6 | 12 | 6 | 8 | 12 | 14 | 16 | 17 | 13 | 34 | 8 | 6 | 9 | 24 |
| RUFY1 | chr5:179586331-179586475\|179586563 | 10 | 7 | 5 | 6 | 7 | 7 | 10 | 12 | 12 | 11 | 13 | 26 | 26 | 11 | 11 | 13 | 11 | 13 | 12 | 12 | 10 | 10 | 9 | 9 | 8 | 9 | 7 | 9 | 9 | 13 | 12 | 11 | 8 | 8 | 11 | 11 | 14 | 17 | 11 | 13 | 16 | 11 | 11 | 11 | 10 | 8 | 33 | 14 | 11 | 11 | 3 |
| CHUK | chr10:100192459-100192693 | 10 | 7 | 3 | 6 | 6 | 9 | 8 | 10 | 16 | 21 | 15 | 69 | 69 | 25 | 22 | 21 | 21 | 18 | 14 | 17 | 15 | 21 | 14 | 9 | 4 | 7 | 6 | 6 | 13 | 20 | 15 | 4 | 2 | 6 | 8 | 23 | 18 | 23 | 5 | 10 | 11 | 13 | 14 | 4 | 4 | 6 | 78 | 10 | 13 | 13 | 3 |
| TMEM200A | chr6:130439348-130439478 | 7 | 8 | 6 | 6 | 5 | 7 | -1 | 5 | 4 | 10 | 5 | -1 | -1 | 3 | 4 | 16 | 10 | -1 | -1 | -1 | -1 | 3 | -1 | 6 | 7 | 7 | 6 | 7 | 7 | 9 | 6 | 8 | 6 | 7 | 10 | 8 | 7 | 9 | 8 | 10 | 5 | 5 | 6 | 6 | 16 | 8 | 24 | 7 | 7 | 6 | 8 |
| MYO1B | chr2:191400749-191402718 | 21 | 25 | 34 | 14 | 17 | 16 | 16 | 22 | 46 | 61 | 46 | 36 | 44 | 71 | 75 | 51 | 53 | 58 | 44 | 38 | 44 | 14 | 22 | 21 | 16 | 22 | 9 | 23 | 26 | 27 | 29 | 31 | 22 | 13 | 20 | 21 | 45 | 35 | 21 | 39 | 23 | 13 | 12 | 27 | 33 | 25 | 74 | 26 | 21 | 13 | 19 |
| ZNF470 | chr19:56573911-56573984 | 10 | 10 | 6 | 9 | 9 | 11 | 8 | 11 | 10 | 10 | 12 | 8 | 7 | 10 | 6 | 12 | 9 | 11 | 10 | 15 | 13 | -1 | -1 | 8 | 7 | 7 | 6 | 7 | 6 | 9 | 8 | 5 | 3 | 10 | 10 | 9 | 14 | 13 | 10 | 9 | 9 | 11 | 12 | 11 | 10 | 9 | 27 | 9 | 10 | 8 | 11 |
| SLC38A6 | chr14:61048151\|61046800-61048297 | 19 | 11 | 7 | 12 | 12 | 14 | 16 | 18 | 18 | 17 | 16 | 18 | 17 | 16 | 15 | 17 | 18 | 19 | 18 | 20 | 11 | 19 | 13 | 13 | 11 | 12 | 10 | 11 | 20 | 20 | 17 | 13 | 10 | 14 | 15 | 24 | 21 | 16 | 8 | 18 | 16 | 20 | 23 | 13 | 13 | 11 | 39 | 12 | 14 | 15 | 10 |
| SNX16 | chr8:81840198-81840443 | 16 | 14 | 12 | 19 | 17 | 22 | 20 | 15 | 33 | 31 | 25 | 24 | 23 | 31 | 32 | 27 | 30 | 23 | 30 | 27 | 31 | 19 | 14 | 19 | 16 | 16 | 13 | 19 | 14 | 14 | 14 | 14 | 10 | 16 | 13 | 27 | 15 | 14 | 13 | 22 | 17 | 17 | 16 | 14 | 12 | 16 | 38 | 12 | 21 | 15 | 8 |
| C19orf66 | chr19:10089111-10089378 | 10 | 9 | 6 | 6 | 6 | 5 | 12 | 11 | 4 | 3 | 3 | 5 | 5 | 4 | 3 | 4 | 3 | 3 | 3 | 5 | 4 | 8 | 7 | 6 | 10 | 6 | 10 | 6 | 8 | 8 | 9 | 12 | 19 | 11 | 10 | 10 | 10 | 5 | 11 | 6 | 10 | 11 | 13 | 15 | 12 | 9 | 27 | 10 | 5 | 8 | 13 |
| ERCC6L2 | chr9:95967459\|95967319-95967625 | 13 | 11 | 6 | 10 | 10 | 11 | 6 | 15 | 9 | 5 | 9 | 15 | 17 | 11 | 7 | 10 | 7 | 7 | 9 | 16 | 11 | 17 | 11 | 11 | 10 | 9 | 12 | 8 | 10 | 11 | 9 | 9 | 12 | 14 | 12 | 7 | 16 | 16 | 10 | 16 | 15 | 18 | 19 | 13 | 17 | 10 | 31 | 11 | 13 | 15 | 14 |
| ANKRD29 | chr18:23609224-23609291\|23609882 | 18 | 16 | 9 | 13 | 15 | 13 | 17 | 17 | 10 | 9 | 13 | 7 | 7 | 10 | 8 | 9 | 9 | 11 | 12 | 9 | 8 | 14 | 13 | 13 | 16 | 13 | 16 | 13 | 17 | 12 | 14 | 20 | 25 | 15 | 14 | 18 | 22 | 9 | 14 | 10 | 12 | 19 | 20 | 19 | 23 | 18 | 38 | 13 | 12 | 13 | 15 |
| CTDSPL2 | chr15:44501962\|44503424-44503850 | 19 | 21 | 19 | 20 | 22 | 24 | 24 | 23 | 16 | 15 | 18 | 21 | 28 | 18 | 7 | 24 | 18 | 16 | 13 | 26 | 15 | 31 | 21 | 29 | 22 | 27 | 21 | 25 | 21 | 23 | 21 | 20 | 26 | 21 | 28 | 22 | 27 | 22 | 30 | 35 | 32 | 23 | 21 | 26 | 19 | 24 | 46 | 24 | 23 | 26 | 21 |
| ZNF532 | chr18:58931646-58932528\|58933313 | 12 | 9 | 10 | 8 | 8 | 9 | 10 | 12 | 10 | 10 | 12 | 12 | 13 | 11 | 7 | 10 | 10 | 12 | 10 | 9 | 8 | 7 | 8 | 8 | 8 | 8 | 7 | 8 | 10 | 13 | 14 | 6 | 9 | 9 | 10 | 34 | 18 | 10 | 7 | 7 | 10 | 10 | 12 | 10 | 7 | 8 | 37 | 7 | 9 | 8 | 6 |
| TCP11L2 | chr12:106312293-106312434 | 10 | 13 | 9 | 11 | 12 | 12 | 10 | 11 | 19 | 22 | 22 | 23 | 24 | 26 | 21 | 24 | 22 | 26 | 22 | 22 | 18 | 10 | 10 | 14 | 10 | 14 | 8 | 11 | 17 | 15 | 8 | 12 | 10 | 10 | 9 | 5 | 20 | 21 | 12 | 23 | 14 | 6 | 6 | 11 | 9 | 12 | 37 | 13 | 17 | 11 | 7 |
| DENND3 | chr8:141150098-141150348\|141150285 | 4 | 6 | 4 | 6 | 4 | 6 | 9 | 5 | 15 | 38 | 14 | 11 | 11 | 43 | 44 | 16 | 13 | 18 | 14 | 8 | 12 | 5 | 6 | 11 | 7 | 7 | 4 | 8 | 5 | 12 | 9 | 6 | 4 | 4 | 2 | 5 | 5 | 8 | 4 | 7 | 8 | 3 | 3 | 3 | 2 | 4 | 20 | 4 | 8 | 9 | 2 |
| TCF12 | chr15:57252421-57252492 | 2 | 2 | 3 | 1 | 1 | 1 | 7 | 3 | 24 | 30 | 22 | 59 | 45 | 27 | 30 | 25 | 26 | 28 | 24 | 27 | 25 | 2 | 3 | 6 | 15 | 5 | 4 | 5 | 5 | 9 | 13 | 5 | 4 | 2 | 8 | 7 | 2 | 3 | 17 | 32 | 8 | 3 | 2 | 14 | 3 | 13 | 58 | 5 | 3 | 3 | 3 |
| PGAP1 | chr2:196877573-196877636 | 16 | 16 | 16 | 11 | 9 | 12 | 13 | 23 | 10 | 5 | 10 | 4 | 7 | 6 | 4 | 11 | 2 | 5 | 8 | 10 | 3 | -1 | -1 | 11 | 11 | 12 | 8 | 8 | 0 | 8 | 9 | 14 | 10 | 13 | 10 | 13 | 26 | 33 | 26 | 10 | 14 | 28 | 26 | 15 | 10 | 13 | 59 | 10 | 11 | 15 | 30 |
| RP11-401P9.5 | chr16:50670133-50670332 | 3 | 14 | 3 | 7 | 3 | 3 | 0 | 1 | 1 | -1 | 9 | 3 | 8 | 7 | 5 | 7 | 5 | 9 | 3 | 1 | 4 | 0 | 0 | 7 | 9 | 3 | 2 | 6 | 0 | -1 | 0 | -1 | 7 | 6 | 2 | 1 | 2 | 10 | 1 | 4 | 3 | 1 | 1 | 2 | 5 | 6 | 24 | 4 | 10 | 3 | 0 |
| MCEE | chr2:71110763\|71110633-71110858 | 5 | 4 | 1 | 4 | 4 | 5 | 9 | 5 | 2 | 1 | 2 | 4 | 5 | 2 | 2 | 2 | 1 | 1 | 2 | 1 | 1 | 12 | 10 | 6 | 4 | 3 | 2 | 4 | 9 | 6 | 5 | 1 | 2 | 6 | 3 | 20 | 6 | 8 | 3 | 2 | 3 | 6 | 5 | 5 | 5 | 3 | 26 | 2 | 6 | 3 | 2 |
| COX20 | chr1:244841944\|244836485-244842058 | 11 | 9 | 9 | 7 | 6 | 7 | 7 | 9 | 6 | 7 | 6 | 14 | 7 | 7 | 13 | 7 | 7 | 4 | 5 | 9 | 8 | 8 | 11 | 6 | 8 | 5 | 9 | 6 | 7 | 3 | 7 | 7 | 7 | 15 | 9 | 9 | 7 | 12 | 18 | 7 | 7 | 8 | 10 | 8 | 12 | 7 | 29 | 14 | 10 | 9 | 21 |
| TEAD2 | chr19:49348957-49349059 | 29 | 15 | 12 | 16 | 14 | 20 | 16 | 27 | 9 | 7 | 10 | 10 | 9 | 11 | 11 | 7 | 9 | 9 | 10 | 11 | 10 | 17 | 13 | 18 | 17 | 16 | 16 | 16 | 18 | 10 | 8 | 14 | 13 | 25 | 21 | 22 | 22 | 16 | 17 | 16 | 18 | 25 | 31 | 17 | 17 | 15 | 47 | 19 | 19 | 18 | 22 |
| EP400 | chr12:131963532-131963642 | 14 | 12 | 21 | 13 | 13 | 14 | 14 | 18 | 14 | 12 | 10 | 16 | 21 | 12 | 9 | 12 | 16 | 8 | 17 | 12 | 10 | -1 | -1 | 10 | 12 | 9 | 13 | 7 | 11 | 11 | 10 | 12 | 13 | 11 | 17 | 4 | 10 | 20 | 12 | 24 | 14 | 11 | 14 | 9 | 8 | 12 | 42 | 14 | 14 | 12 | 1 |
| NRDE2 | chr14:90317704-90317820 | 8 | 6 | 6 | 8 | 7 | 7 | 9 | 10 | 16 | 12 | 14 | 15 | 14 | 10 | 13 | 16 | 12 | 15 | 14 | 19 | 13 | 5 | 14 | 8 | 8 | 6 | 7 | 8 | 11 | 9 | 6 | 9 | 4 | 8 | 9 | 4 | 11 | 7 | 9 | 14 | 9 | 10 | 9 | 11 | 8 | 11 | 23 | 13 | 9 | 9 | 6 |
| VEPH1 | chr3:157317586-157317746 | 7 | 3 | 1 | 3 | 3 | 8 | 0 | 5 | 0 | 0 | -1 | 3 | 8 | -1 | 0 | -1 | -1 | 0 | 0 | 0 | 0 | -1 | -1 | 4 | 4 | 1 | -1 | 1 | 0 | 10 | 4 | 1 | 0 | 3 | 3 | 5 | 7 | 0 | 4 | 3 | 4 | 5 | 8 | -1 | 2 | 3 | 26 | 1 | 0 | -1 | 20 |
| SLC35B3 | chr6:8418744-8418885 | 10 | 10 | 4 | 11 | 11 | 12 | 8 | 10 | 42 | 35 | 41 | 24 | 26 | 40 | 29 | 47 | 36 | 39 | 44 | 64 | 50 | 15 | 12 | 12 | 6 | 11 | 8 | 11 | 13 | 15 | 15 | 7 | 4 | 8 | 6 | 15 | 28 | 18 | 4 | 13 | 12 | 8 | 8 | 5 | 8 | 7 | 41 | 9 | 12 | 14 | 3 |
| KLKB1 | chr4:186228871-186228989 | 5 | 7 | 9 | 9 | 5 | 4 | 0 | 5 | 4 | 3 | 3 | 4 | 5 | 2 | 2 | 4 | 3 | 4 | 6 | 6 | 4 | -1 | -1 | 4 | 6 | 5 | 4 | 3 | -1 | 4 | 2 | 9 | 4 | 8 | 11 | 3 | 6 | 4 | 14 | 8 | 7 | 9 | 6 | 15 | 11 | 11 | 28 | 11 | 4 | 2 | 7 |
| BACH2 | chr6:90062884-90062963 | 15 | 17 | 20 | 15 | 11 | 19 | -1 | 17 | 16 | 10 | 13 | 10 | 8 | 13 | 20 | 12 | 10 | 14 | 14 | 16 | 14 | -1 | -1 | 16 | 18 | 18 | 17 | 14 | -1 | 13 | 18 | -1 | 13 | 15 | 20 | 24 | 16 | 14 | 15 | 11 | 18 | 17 | 17 | 24 | 28 | 14 | 54 | 14 | 15 | 15 | 18 |
| SCYL2 | chr12:100324938-100325138\|100325172 | 10 | 6 | 4 | 4 | 4 | 5 | 3 | 11 | 4 | 5 | 5 | 4 | 3 | 4 | 2 | 5 | 4 | 5 | 4 | 3 | 5 | 5 | 6 | 3 | 3 | 1 | 3 | 1 | 4 | 5 | 2 | 6 | 7 | 9 | 4 | 3 | 11 | 7 | 5 | 10 | 5 | 7 | 9 | 4 | 7 | 4 | 21 | 5 | 4 | 5 | 12 |
| ACER2 | chr9:19424369-19424551 | 8 | 11 | 3 | 9 | 7 | 5 | 4 | 8 | 12 | 16 | 12 | 25 | 31 | 19 | 13 | 15 | 22 | 12 | 17 | 13 | 14 | 0 | 0 | 8 | 10 | 8 | 10 | 10 | 0 | 15 | 16 | -1 | 9 | 8 | 3 | 7 | 11 | 14 | 6 | 18 | 9 | 8 | 5 | 10 | 11 | 7 | 40 | 8 | 10 | 6 | 5 |
| KIF21B | chr1:201003079-201003242 | 8 | 9 | 4 | 7 | 7 | 4 | -1 | 9 | 1 | 1 | 2 | 1 | 3 | 2 | 1 | 1 | 1 | 2 | 1 | 6 | 1 | -1 | 0 | 5 | 9 | 6 | 8 | 4 | 0 | 9 | 6 | 14 | 8 | 7 | 5 | 9 | 8 | 5 | 5 | 2 | 7 | 7 | 11 | 6 | 7 | 3 | 27 | 11 | 12 | 13 | 5 |
| TESMIN | chr11:68742318-68745111 | 28 | 24 | 21 | 34 | 24 | 40 | -1 | 36 | 8 | 22 | 12 | 11 | 20 | 22 | 15 | 20 | 9 | 11 | 15 | -1 | 15 | -1 | -1 | 37 | 30 | 29 | 25 | 30 | -1 | 17 | 18 | 37 | 29 | 43 | 30 | 49 | 30 | 23 | 34 | 42 | 44 | 30 | 33 | 38 | 42 | 36 | 99 | 36 | 36 | 24 | 34 |
| KDM4C | chr9:7105402-7105500 | 7 | 11 | 19 | 12 | 12 | 11 | 15 | 10 | 33 | 37 | 38 | 24 | 25 | 34 | 33 | 33 | 33 | 42 | 31 | 28 | 27 | 10 | 10 | 13 | 19 | 11 | 11 | 9 | 15 | 8 | 10 | 21 | 16 | 9 | 17 | 12 | 16 | 17 | 23 | 34 | 21 | 8 | 7 | 17 | 12 | 22 | 46 | 18 | 13 | 10 | 9 |
| PNPLA8 | chr7:108485694-108485762 | 5 | 8 | 2 | 3 | 3 | 3 | 5 | 6 | 1 | 1 | 0 | 2 | 2 | 1 | 0 | 1 | 1 | 0 | 1 | 1 | 0 | 6 | 3 | 3 | 6 | 2 | 5 | 2 | 10 | 2 | 1 | 3 | 5 | 6 | 5 | 4 | 3 | 4 | 5 | 2 | 3 | 12 | 8 | 12 | 7 | 8 | 20 | 2 | 3 | 4 | 9 |
| DENND3 | chr8:141150098\|141146888-141150348 | 3 | 5 | 4 | 5 | 4 | 5 | 9 | 4 | 13 | 31 | 12 | 10 | 9 | 36 | 36 | 14 | 11 | 16 | 13 | 7 | 11 | 4 | 5 | 10 | 6 | 7 | 3 | 7 | 5 | 11 | 8 | 6 | 4 | 4 | 2 | 5 | 5 | 7 | 3 | 6 | 7 | 3 | 3 | 2 | 2 | 4 | 18 | 3 | 8 | 8 | 2 |
| NNT | chr5:43647924-43648182\|43648440\|43647978 | 12 | 11 | 3 | 13 | 10 | 14 | 10 | 15 | 2 | 2 | 3 | 6 | 7 | 2 | 1 | 2 | 3 | 2 | 3 | 7 | 4 | 22 | 13 | 8 | 6 | 7 | 9 | 5 | 13 | 3 | 1 | 7 | 11 | 12 | 6 | 2 | 21 | 14 | 10 | 13 | 14 | 16 | 15 | 10 | 12 | 5 | 38 | 11 | 16 | 18 | 12 |
| UBE2Q2P1 | chr15:84539704-84539788 | 15 | 14 | 12 | 15 | 15 | 15 | 16 | 13 | 10 | 7 | 7 | 9 | 8 | 8 | 8 | 9 | 8 | 11 | 12 | 9 | 12 | 12 | 21 | 12 | 13 | 10 | 12 | 11 | 13 | 14 | 13 | 18 | 9 | 14 | 15 | 14 | 16 | 17 | 16 | 12 | 11 | 15 | 15 | 13 | 15 | 11 | 28 | 15 | 13 | 12 | 13 |
| DYRK1A | chr21:37421774-37421866 | 7 | 16 | 16 | 10 | 12 | 7 | 10 | 12 | 9 | 8 | 13 | 10 | 11 | 10 | 8 | 10 | 11 | 9 | 13 | 8 | 9 | 10 | 15 | 14 | 19 | 12 | 16 | 11 | 15 | 9 | 8 | 18 | 19 | 15 | 13 | 8 | 10 | 12 | 20 | 13 | 14 | 8 | 7 | 17 | 16 | 14 | 33 | 18 | 12 | 13 | 12 |
| SLC38A11 | chr2:164909593-164909696 | 1 | 1 | 10 | 3 | 2 | 2 | -1 | 1 | 22 | 22 | 25 | 3 | 6 | 61 | 63 | 25 | 12 | 17 | 27 | 5 | 4 | 0 | -1 | 7 | 2 | 4 | 2 | 5 | -1 | 4 | 2 | 3 | 3 | 3 | 2 | 2 | 2 | 2 | 1 | 12 | 5 | 1 | 1 | 3 | 3 | 2 | 17 | 2 | 1 | 1 | 2 |
| LYST | chr1:235808079-235808283 | 8 | 10 | 10 | 8 | 6 | 7 | -1 | 9 | -1 | 7 | 5 | 3 | 5 | 4 | -1 | 5 | -1 | 4 | -1 | 7 | -1 | -1 | -1 | 7 | 9 | 8 | 7 | 7 | 0 | 8 | 6 | 6 | 4 | 8 | 6 | 6 | 9 | 8 | 8 | 9 | 8 | 8 | 6 | 10 | 11 | 8 | 23 | 6 | 4 | 7 | 6 |
| RNFT2 | chr12:116824983-116825080 | 4 | 1 | 4 | 2 | 3 | 3 | 0 | 2 | 2 | 0 | 2 | 2 | 1 | 0 | 1 | 2 | 2 | 2 | 1 | 0 | 0 | -1 | 0 | 6 | 4 | 3 | 3 | 1 | 0 | 4 | 5 | 0 | -1 | 4 | -1 | 2 | 2 | 7 | -1 | 6 | -1 | 6 | 3 | 13 | 6 | 13 | 22 | 7 | 2 | -1 | 3 |
| CPNE8 | chr12:38679578-38679714 | 21 | 12 | 5 | 6 | 6 | 10 | 10 | 20 | 2 | 2 | 2 | 5 | 3 | 3 | 1 | 2 | 3 | 2 | 2 | 3 | 3 | 15 | 6 | 11 | 9 | 8 | 7 | 7 | 9 | 6 | 9 | 7 | 11 | 15 | 12 | 21 | 16 | 7 | 10 | 6 | 9 | 22 | 22 | 15 | 10 | 12 | 39 | 6 | 10 | 11 | 10 |
| CTH | chr1:70423215-70423272 | 4 | 7 | 1 | 5 | 4 | 7 | 13 | 5 | 0 | 0 | 1 | 1 | 1 | 1 | 1 | 1 | 1 | 0 | 1 | 2 | 2 | 18 | 0 | 5 | 2 | 4 | 1 | 4 | 2 | 8 | 4 | 1 | 1 | 3 | 1 | 3 | 5 | 4 | 3 | 1 | 2 | 6 | 4 | 3 | 4 | 3 | 22 | 2 | 3 | 4 | 2 |
| FLJ21408 | chr16:27269671-27269805 | 28 | 23 | 21 | 29 | 25 | 22 | -1 | 23 | 9 | -1 | 19 | 11 | 10 | 20 | 17 | 32 | 19 | 15 | 13 | 24 | 11 | 0 | -1 | 20 | 31 | 16 | 31 | 17 | -1 | 18 | 19 | -1 | 33 | 28 | 20 | -1 | 24 | 25 | 33 | 19 | 26 | 26 | 21 | 27 | 31 | 26 | 79 | 23 | 26 | 21 | 34 |
| RP11-106M3.2 | chr15:72292198-72292265 | 12 | 10 | 12 | 13 | 12 | 13 | 10 | 11 | 7 | 4 | 9 | 5 | 6 | 5 | 3 | 6 | 5 | 7 | 9 | 11 | 7 | 9 | 16 | 11 | 11 | 11 | 10 | 12 | 11 | 12 | 12 | 11 | 13 | 14 | 12 | 14 | 14 | 9 | 12 | 8 | 13 | 10 | 11 | 12 | 15 | 11 | 22 | 10 | 10 | 10 | 15 |
| AUH | chr9:91360371\|91356889\|91360279\|91357484-91360440 | 22 | 14 | 3 | 12 | 12 | 18 | 13 | 20 | 10 | 7 | 7 | 15 | 20 | 8 | 5 | 9 | 7 | 6 | 7 | 11 | 8 | 21 | 19 | 17 | 12 | 14 | 12 | 15 | 12 | 9 | 6 | 8 | 9 | 20 | 13 | 13 | 23 | 23 | 17 | 12 | 17 | 21 | 25 | 20 | 18 | 11 | 44 | 14 | 22 | 21 | 15 |
| COL11A1 | chr1:102895365-102895464 | 6 | 5 | 6 | 5 | 7 | 5 | 0 | 7 | 9 | 11 | 9 | 9 | 8 | 11 | 10 | 9 | 8 | 8 | 10 | 9 | 11 | 0 | -1 | 10 | 9 | 9 | -1 | 8 | 0 | 11 | -1 | -1 | -1 | 8 | 4 | 8 | 10 | 5 | 2 | 9 | 6 | 3 | 5 | 6 | 6 | 8 | 25 | 6 | 9 | 7 | -1 |
| OXSR1 | chr3:38193282-38193418 | 13 | 8 | 4 | 7 | 7 | 9 | 7 | 14 | 7 | 6 | 6 | 7 | 10 | 7 | 4 | 5 | 5 | 6 | 6 | 8 | 6 | 14 | 4 | 9 | 9 | 7 | 8 | 7 | 17 | 6 | 4 | 8 | 7 | 9 | 11 | 2 | 12 | 10 | 11 | 12 | 13 | 15 | 15 | 12 | 10 | 10 | 28 | 10 | 8 | 11 | 11 |
| TNNI3K | chr1:74342083-74342286 | 21 | 13 | 7 | 12 | 11 | 13 | 11 | 21 | 17 | 21 | 18 | 22 | 20 | 20 | 23 | 17 | 17 | 23 | 22 | 17 | 14 | -1 | -1 | 11 | 11 | 11 | 11 | 11 | 18 | 8 | 4 | 13 | 25 | 15 | 18 | 12 | 29 | 14 | 20 | 17 | 15 | 19 | 25 | 15 | 20 | 12 | 48 | 13 | 14 | 17 | 16 |
| RP11-84G21.1 | chr12:79938728-79938852 | 14 | 14 | 11 | 6 | 5 | 5 | -1 | 11 | 2 | 1 | 2 | 0 | -1 | 2 | 7 | 0 | 4 | 3 | 4 | 1 | 6 | -1 | 0 | 8 | 25 | 5 | 24 | 5 | -1 | 7 | 11 | -1 | -1 | 12 | 24 | 4 | 6 | 2 | 14 | 1 | 11 | 13 | 11 | 19 | 14 | 13 | 49 | 3 | 5 | 9 | 20 |
| ATG4A | chrX:108126793-108127094 | 9 | 8 | 5 | 12 | 10 | 12 | 8 | 9 | 20 | 22 | 19 | 28 | 32 | 21 | 21 | 20 | 22 | 22 | 15 | 15 | 17 | 16 | 19 | 9 | 7 | 10 | 5 | 10 | 7 | 18 | 13 | 8 | 3 | 10 | 7 | 19 | 15 | 19 | 4 | 19 | 11 | 8 | 9 | 7 | 6 | 6 | 41 | 8 | 13 | 9 | 6 |
| PGAP3 | chr17:39675010\|39676614-39676755 | 7 | 7 | 5 | 6 | 6 | 5 | 9 | 10 | 7 | 4 | 5 | 11 | 9 | 4 | 3 | 7 | 5 | 4 | 6 | 8 | 7 | 5 | 6 | 6 | 8 | 5 | 7 | 6 | 9 | 5 | 5 | 11 | 6 | 7 | 6 | 5 | 8 | 5 | 9 | 10 | 14 | 6 | 7 | 9 | 12 | 6 | 20 | 13 | 5 | 7 | 9 |
| FUZ | chr19:49808115\|49807570-49808159 | 10 | 11 | 11 | 12 | 11 | 10 | 14 | 12 | 5 | 4 | 5 | 6 | 7 | 5 | 4 | 6 | 6 | 4 | 5 | 8 | 7 | 13 | 13 | 13 | 14 | 12 | 15 | 11 | 14 | 8 | 9 | 14 | 15 | 12 | 15 | 4 | 9 | 10 | 18 | 12 | 14 | 13 | 13 | 17 | 19 | 15 | 30 | 16 | 11 | 13 | 20 |
| DUSP18 | chr22:30664955-30665121\|30666938 | 10 | 15 | 21 | 15 | 14 | 13 | 15 | 11 | 15 | 16 | 12 | 13 | 18 | 19 | 16 | 14 | 18 | 14 | 18 | 15 | 14 | -1 | -1 | 17 | 14 | 14 | 17 | 18 | 18 | 14 | 17 | 12 | 24 | 12 | 12 | 15 | 11 | 13 | 14 | 13 | 14 | 10 | 9 | 15 | 12 | 14 | 38 | 10 | 14 | 14 | 12 |
| XKR9 | chr8:70683509-70683630 | 1 | 1 | 1 | 2 | 3 | 2 | 0 | 4 | 1 | 3 | 1 | 1 | -1 | 1 | 1 | 2 | 1 | 4 | 6 | 0 | 1 | 0 | 0 | 1 | 1 | 1 | 1 | 1 | 0 | 2 | 1 | 0 | 1 | 2 | 2 | 3 | 1 | 1 | 1 | 1 | 0 | 3 | 2 | 1 | 1 | 1 | 1 | 1 | 2 | 1 | 28 |
| FRY | chr13:32188219-32188525 | 7 | 11 | 4 | 7 | 6 | 6 | 4 | 7 | 2 | 1 | 3 | 1 | 1 | 1 | 1 | 3 | 3 | 0 | 2 | 2 | 2 | -1 | -1 | 5 | 5 | 7 | 12 | 6 | 3 | 4 | 1 | 7 | 19 | 16 | 4 | 4 | 6 | 6 | 10 | 6 | 6 | 13 | 11 | 7 | 11 | 7 | 5 | 6 | 5 | 9 | 55 |
| ANKS1B | chr12:98807844-98807915\|98807918 | 2 | 1 | 1 | 0 | 0 | 1 | 0 | 1 | 31 | 54 | 49 | 4 | 6 | 58 | 64 | 31 | 9 | 61 | 45 | 0 | 7 | 0 | -1 | 1 | 0 | 0 | 2 | 1 | 0 | 4 | 3 | 0 | -1 | 5 | 2 | 20 | 1 | 0 | 1 | 5 | 0 | 1 | 3 | 1 | -1 | 1 | 0 | 1 | 1 | 2 | 41 |
| CPS1 | chr2:210589309-210589541 | 5 | 2 | 0 | 1 | 2 | 2 | 0 | 4 | 1 | 0 | 2 | 2 | 2 | 3 | 0 | 1 | 4 | 1 | 1 | 1 | 1 | 0 | -1 | 2 | 2 | 1 | 1 | 0 | 0 | 2 | 2 | 0 | 1 | 2 | 3 | 1 | 6 | 2 | 5 | 4 | 1 | 3 | 4 | 1 | 1 | 2 | 17 | 2 | 2 | 3 | 29 |
| C14orf159 | chr14:91189715\|91189270\|91189616-91189807 | 3 | 5 | 4 | 3 | 4 | 2 | 6 | 5 | 1 | 1 | 1 | 2 | 2 | 1 | 1 | 1 | 1 | 1 | 1 | 2 | 1 | 4 | 6 | 3 | 6 | 3 | 5 | 3 | 4 | 1 | 1 | 5 | 15 | 7 | 5 | 1 | 4 | 3 | 9 | 6 | 7 | 3 | 3 | 9 | 11 | 7 | 4 | 4 | 4 | 4 | 26 |
| SELO | chr22:50212468-50212684 | 6 | 5 | 2 | 6 | 6 | 8 | 6 | 5 | 6 | 8 | 10 | 11 | 12 | 11 | 11 | 6 | 4 | 11 | 8 | 4 | 4 | 3 | 3 | 7 | 4 | 7 | 3 | 7 | 4 | 4 | 3 | 2 | 3 | 11 | 3 | 10 | 7 | 3 | 3 | 4 | 3 | 5 | 6 | 7 | 8 | 3 | 2 | 3 | 3 | 3 | 25 |
| WDR19 | chr4:39276679\|39276625-39276879 | 4 | 7 | 4 | 7 | 4 | 5 | 5 | 3 | 3 | 3 | 3 | 10 | 11 | 4 | 3 | 2 | 2 | 2 | 2 | 3 | 3 | 13 | 11 | 5 | 5 | 4 | 5 | 4 | 1 | 8 | 4 | 6 | 3 | 12 | 4 | 6 | 2 | 11 | 3 | 6 | 7 | 4 | 4 | 9 | 9 | 4 | 2 | 3 | 10 | 9 | 29 |
| FCHO2 | chr5:73050346-73051195 | 2 | 5 | 2 | 2 | 4 | 1 | 1 | 2 | 2 | 5 | 2 | 1 | 2 | 2 | 1 | 5 | 4 | 2 | 4 | 2 | 3 | 4 | -1 | 3 | 3 | 2 | 6 | 2 | 0 | 6 | 4 | 4 | 16 | 6 | 3 | 1 | 2 | 5 | 10 | 5 | 3 | 3 | 2 | 3 | 12 | 4 | 2 | 7 | 5 | 5 | 26 |
| CUBN | chr10:17025690-17025851 | 10 | 5 | 6 | 2 | 1 | 3 | 0 | 8 | 16 | 12 | 14 | 8 | 10 | 12 | 7 | 14 | 12 | 16 | 12 | 19 | -1 | 0 | 0 | 7 | 9 | 5 | 7 | 7 | -1 | 5 | 3 | 6 | -1 | 10 | 12 | 22 | 12 | 1 | 7 | 11 | 5 | 6 | 11 | 4 | 14 | 11 | 12 | 5 | 2 | 6 | 35 |
| PCNX3 | chr11:65635951\|65635931-65636031 | 7 | 10 | 5 | 6 | 6 | 4 | 5 | 6 | 3 | 2 | 3 | 2 | 1 | 2 | 2 | 3 | 3 | 2 | 3 | 5 | 5 | 8 | 8 | 7 | 8 | 6 | 9 | 6 | 4 | 10 | 7 | 7 | 7 | 13 | 8 | 7 | 7 | 6 | 6 | 8 | 7 | 6 | 7 | 10 | 15 | 7 | 10 | 5 | 6 | 9 | 21 |

**Supplemental Figure 5. Examples of human tissue-specific exons (GTEx)**

Signals of cell type-specific splicing can be diluted in RNA-Seq data obtained from whole tissues. However, analyzing human GTEx data reveals some tissue-specific splicing patterns. Generally, brain, heart, skeletal muscle, pituitary and testis exhibit the most tissue-specific alternative exons, with some exons unique to a single tissue, and other exons that are shared among this group of tissues that show high levels of tissue-specific splicing.

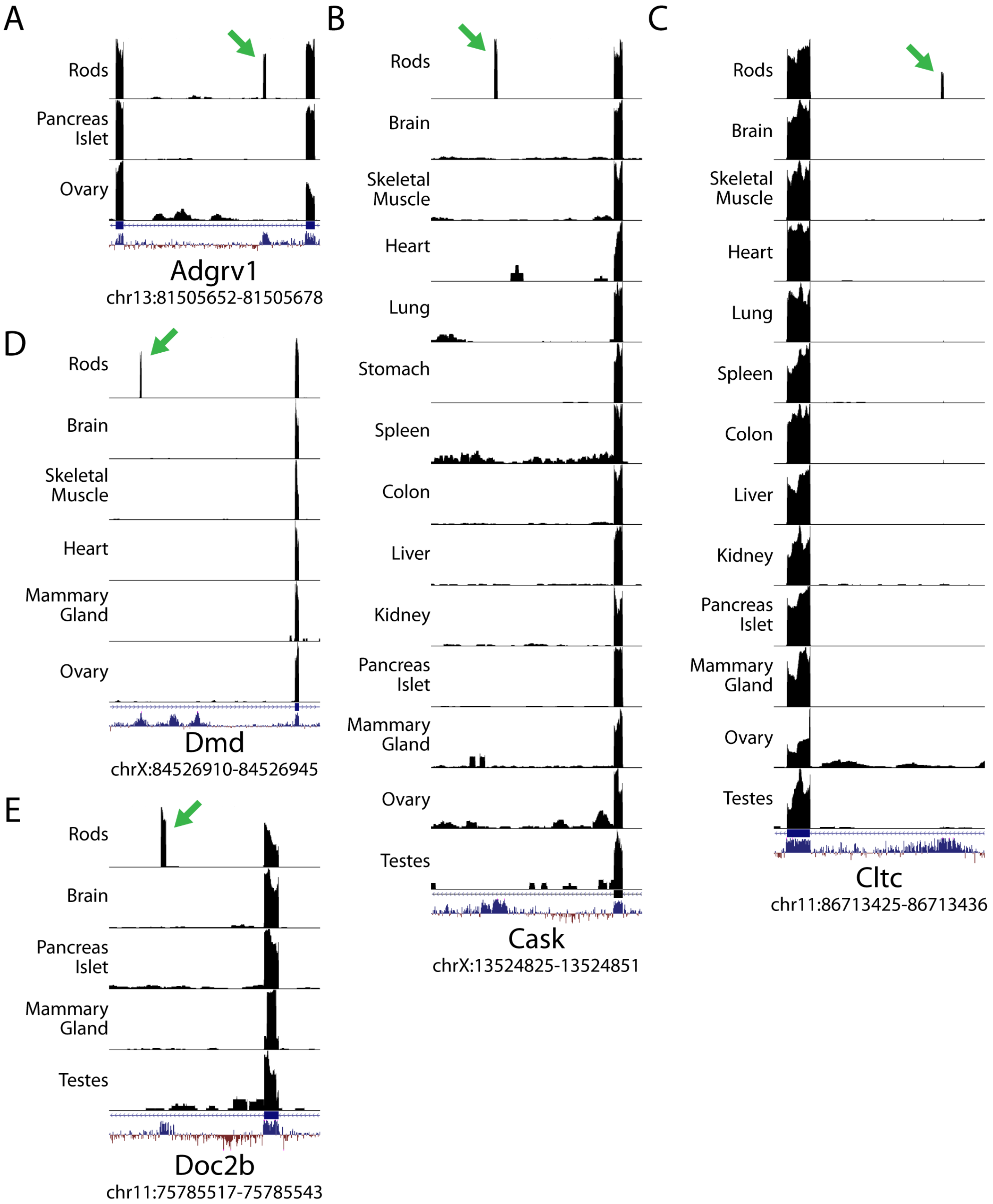
A
Rods
Pancreas Islet
Ovary
Adgrv1
chr13:81505652-81505678
B
Rods
Brain
Skeletal Muscle
Heart
Lung
Stomach
Spleen
Colon
Liver
Kidney
Pancreas Islet
Mammary Gland
Ovary
Testes
Cask
chrX:13524825-13524851
C
Rods
Brain
Skeletal Muscle
Heart
Lung
Spleen
Colon
Liver
Kidney
Pancreas Islet
Mammary Gland
Ovary
Testes
Cltc
chr11:86713425-86713436
D
Rods
Brain
Skeletal Muscle
Heart
Mammary Gland
Ovary
Dmd
chrX:84526910-84526945
E
Rods
Brain
Pancreas Islet
Mammary Gland
Testes
Doc2b
chr11:75785517-75785543

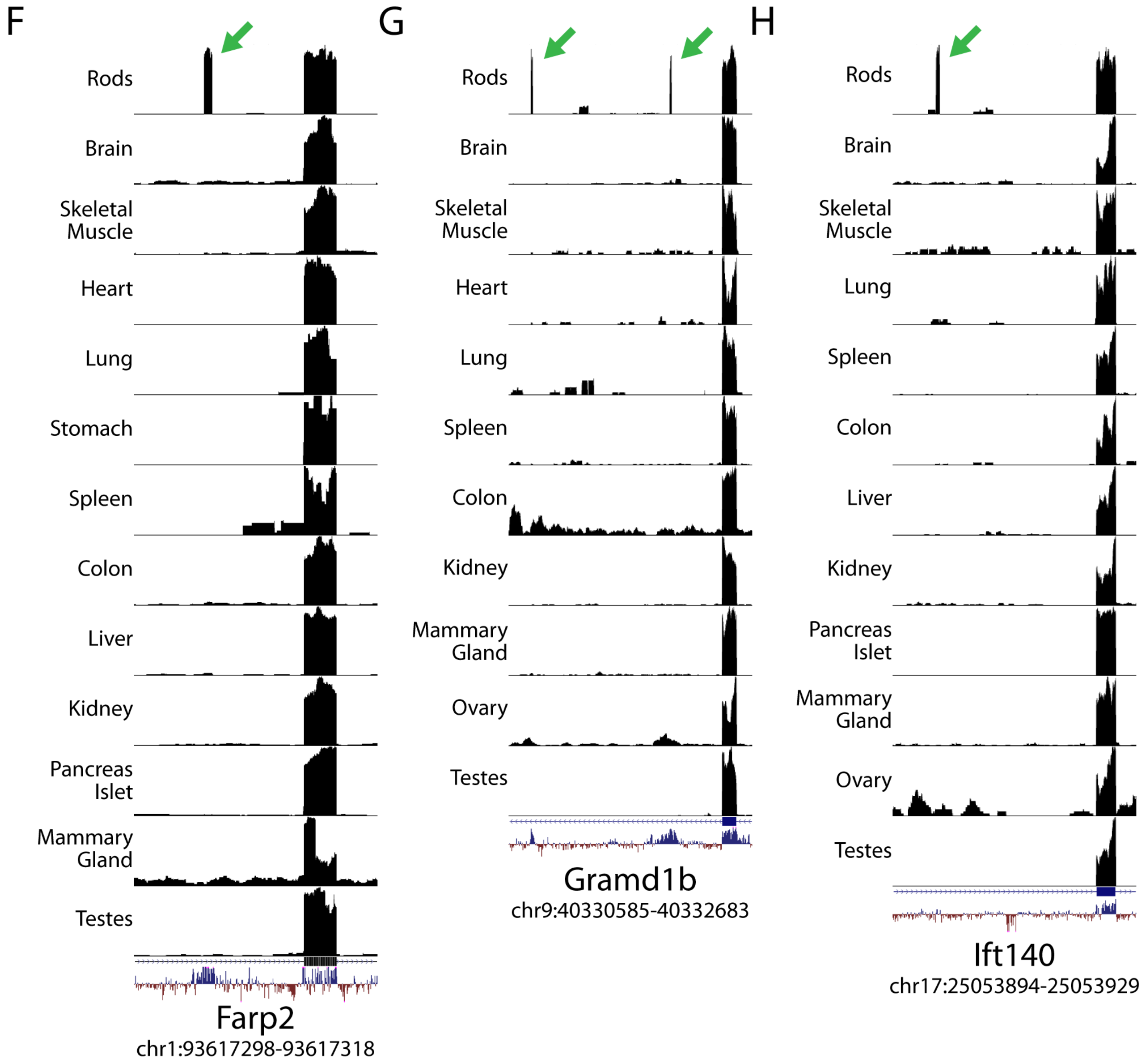
F
Rods
Brain
Skeletal Muscle
Heart
Lung
Stomach
Spleen
Colon
Liver
Kidney
Pancreas Islet
Mammary Gland
Testes
Farp2
chr1:93617298-93617318
G
Rods
Brain
Skeletal Muscle
Heart
Lung
Spleen
Colon
Kidney
Mammary Gland
Ovary
Testes
Gramd1b
chr9:40330585-40332683
H
Rods
Brain
Skeletal Muscle
Lung
Spleen
Colon
Liver
Kidney
Pancreas Islet
Mammary Gland
Ovary
Testes
Ift140
chr17:25053894-25053929

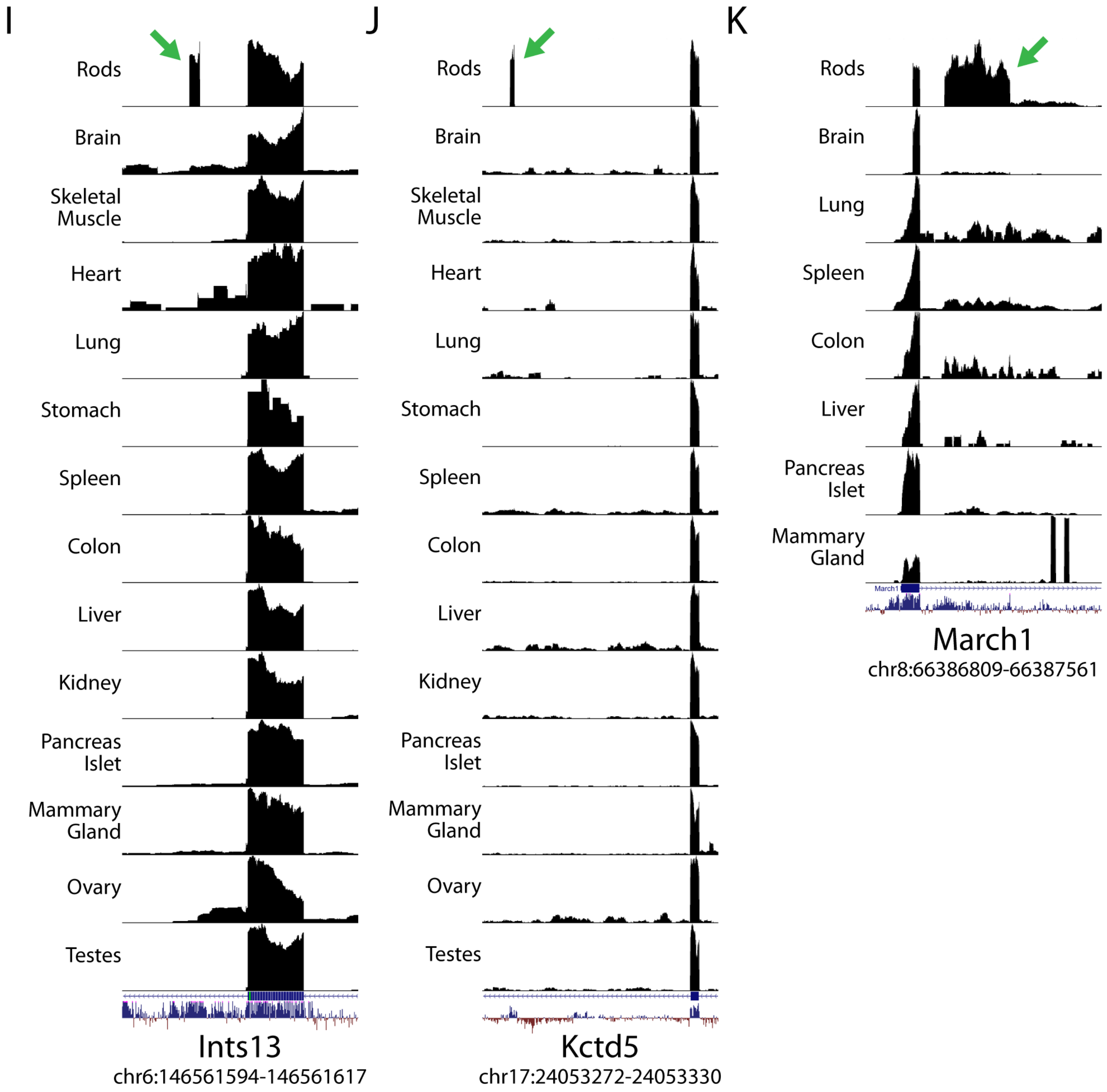
I
Rods
Brain
Skeletal Muscle
Heart
Lung
Stomach
Spleen
Colon
Liver
Kidney
Pancreas Islet
Mammary Gland
Ovary
Testes
Ints13
chr6:146561594-146561617
J
Rods
Brain
Skeletal Muscle
Heart
Lung
Stomach
Spleen
Colon
Liver
Kidney
Pancreas Islet
Mammary Gland
Ovary
Testes
Kctd5
chr17:24053272-24053330
K
Rods
Brain
Lung
Spleen
Colon
Liver
Pancreas Islet
Mammary Gland
March1
chr8:66386809-66387561

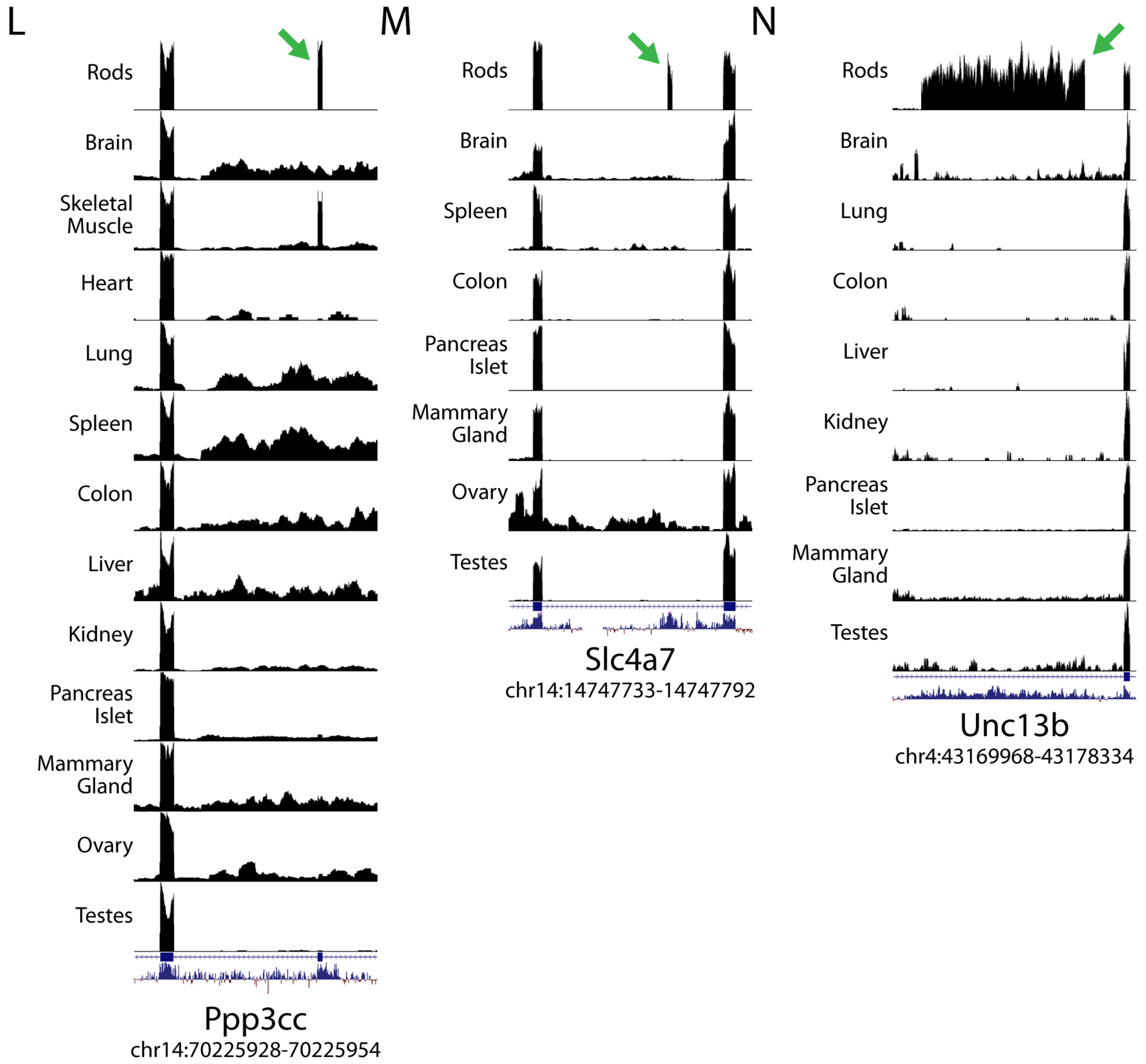
L
Rods
Brain
Skeletal Muscle
Heart
Lung
Spleen
Colon
Liver
Kidney
Pancreas Islet
Mammary Gland
Ovary
Testes
Ppp3cc
chr14:70225928-70225954
M
Rods
Brain
Spleen
Colon
Pancreas Islet
Mammary Gland
Ovary
Testes
Slc4a7
chr14:14747733-14747792
N
Rods
Brain
Lung
Colon
Liver
Kidney
Pancreas Islet
Mammary Gland
Testes
Unc13b
chr4:43169968-43178334

**Supplemental Figure 6. Rod-specific exons, both previously identified and novel**

Of the 31 rod-specific alternative splicing events that are syntenic between mouse and human, 17 have been previously reported (*Amph*[3,4], *Atp1b2*[4,5], *Plekhb1*[4,6], *Bsg*[4,7–9], *Ttc8*[4,10,11], *Impdh1*[4,12], *Cc2d2a*[4], *Kif1b*[4], *Efr3a*[4], *Hcn1*[4], *Unc13b*[4], *Kmt2d*[4], *Man2a2*[4], *Msi2*[4], *Stxbp5*[4]) while 14 novel rod-specific splicing events have not been previously reported (*Adgrv1*, *Cask*, *Cltc*, *Dmd*, *Doc2b*, *Farp2*, *Gramd1b*, *Ift140*, *Ints13*, *Kctd5*, *March1*, *Ppp3cc*, *Slc4a7*, and a second exon in *Unc13b*). Shown in this figure are UCSC bigwig visualizations of the novel rod exons comparing rods to other mouse tissues. Tissues in which the gene is not expressed are excluded from the visualization.

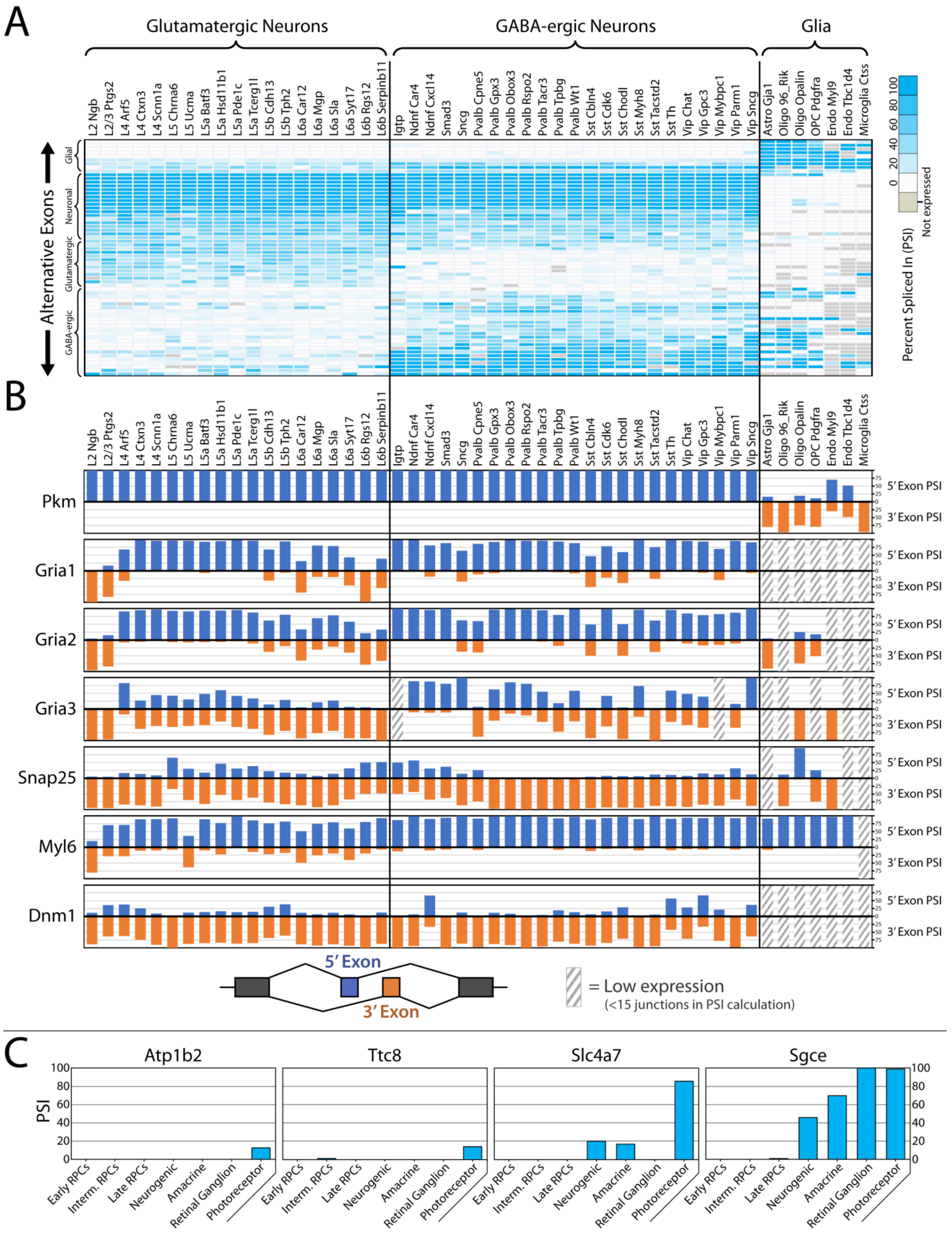
A
Glutamatergic Neurons
GABA-ergic Neurons
Glia
L2 Ngb
L2/3 Ptgs2
L4 Arf5
L4 Ctxn3
L4 Scnn1a
L5 Chrna6
L5 Ucma
L5a Batf3
L5a Hsd11b1
L5a Pde1c
L5a Tcerg1l
L5b Cdh13
L5b Tph2
L6a Car12
L6a Mgp
L6a Sla
L6a Syt17
L6b Rgs12
L6b Serpinb11
Igtp
Ndnf Car4
Ndnf Cxcl14
Smad3
Sncg
Pvalb Cpne5
Pvalb Gpx3
Pvalb Obox3
Pvalb Rspo2
Pvalb Tacr3
Pvalb Tpbg
Pvalb Wt1
Sst Cbln4
Sst Cdk6
Sst Chodl
Sst Myh8
Sst Tacstd2
Sst Th
Vip Chat
Vip Gpc3
Vip Mybpc1
Vip Parm1
Vip Sncg
Astro Gja1
Oligo 96_Rik
Oligo Opalin
OPC Pdgfra
Endo Myl9
Endo Tbc1d4
Microglia Ctss
Alternative Exons
Glial
Neuronal
Glutamatergic
GABA-ergic
100
80
60
40
20
0
Not expressed
Percent Spliced In (PSI)
B
Pkm
Gria1
Gria2
Gria3
Snap25
Myl6
Dnm1
5′ Exon PSI
3′ Exon PSI
5′ Exon
3′ Exon
= Low expression
(<15 junctions in PSI calculation)
C
Atp1b2
Ttc8
Slc4a7
Sgce
PSI
100
80
60
40
20
0
Early RPCs
Interm. RPCs
Late RPCs
Neurogenic
Amacrine
Retinal Ganglion
Photoreceptor

D

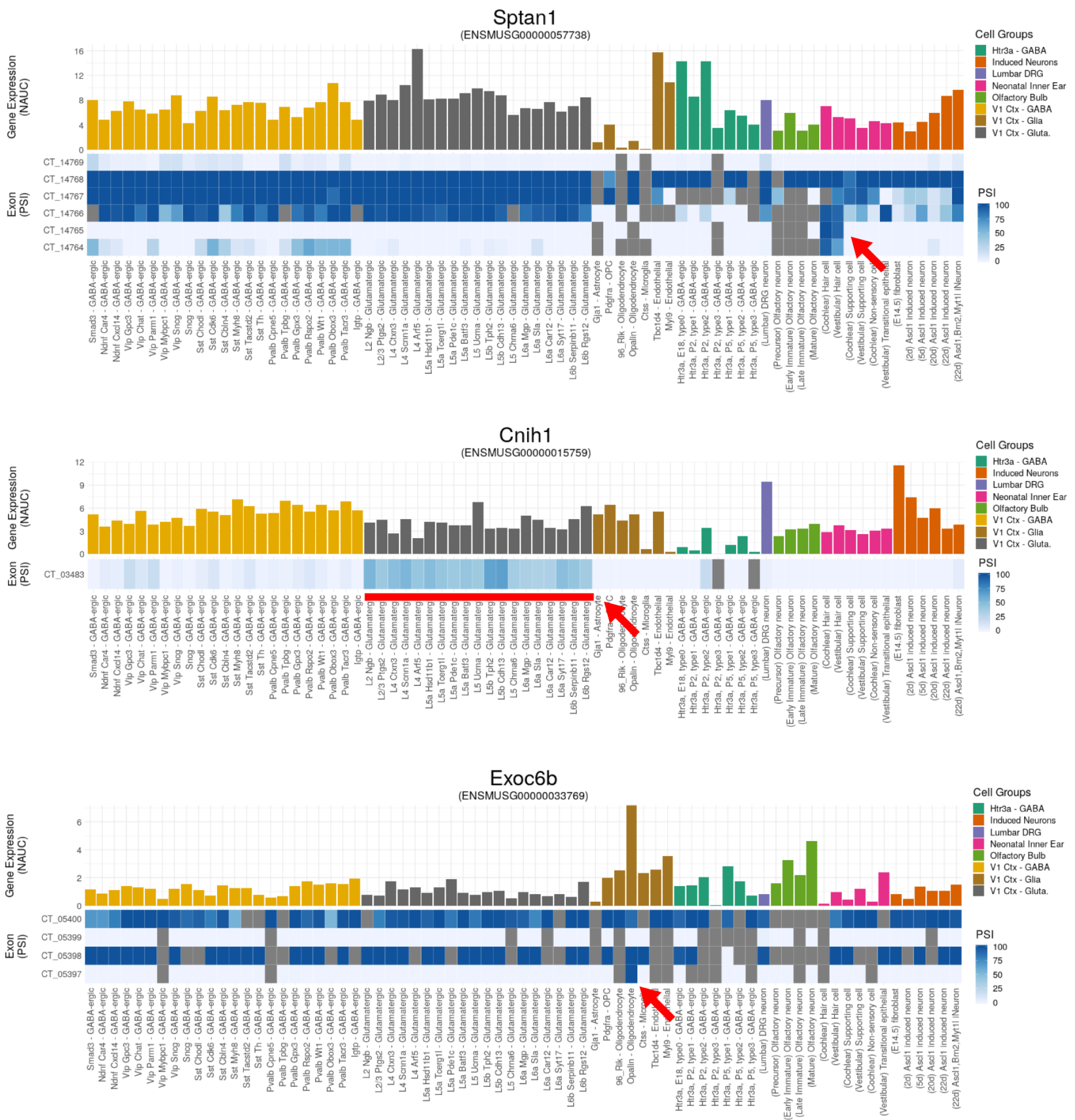

Sptan1
(ENSMUSG00000057738)
Gene Expression (NAUC)
16
12
8
4
0
Exon (PSI)
CT_14769
CT_14768
CT_14767
CT_14766
CT_14765
CT_14764
Cell Groups
Htr3a - GABA
Induced Neurons
Lumbar DRG
Neonatal Inner Ear
Olfactory Bulb
V1 Ctx - GABA
V1 Ctx - Glia
V1 Ctx - Gluta.
PSI
100
75
50
25
0
Smad3 - GABA-ergic
Ndnf Car4 - GABA-ergic
Ndnf Cxcl14 - GABA-ergic
Vip Gpc3 - GABA-ergic
Vip Chat - GABA-ergic
Vip Parm1 - GABA-ergic
Vip Mybpc1 - GABA-ergic
Vip Sncg - GABA-ergic
Sncg - GABA-ergic
Sst Chodl - GABA-ergic
Sst Cdk6 - GABA-ergic
Sst Cbln4 - GABA-ergic
Sst Myh8 - GABA-ergic
Sst Tacstd2 - GABA-ergic
Sst Th - GABA-ergic
Pvalb Cpne5 - GABA-ergic
Pvalb Tpbg - GABA-ergic
Pvalb Gpx3 - GABA-ergic
Pvalb Rspo2 - GABA-ergic
Pvalb Wt1 - GABA-ergic
Pvalb Obox3 - GABA-ergic
Pvalb Tacr3 - GABA-ergic
Igtp - GABA-ergic
L2 Ngb - Glutamatergic
L2/3 Ptgs2 - Glutamatergic
L4 Ctxn3 - Glutamatergic
L4 Scnn1a - Glutamatergic
L4 Arf5 - Glutamatergic
L5a Hsd11b1 - Glutamatergic
L5a Tcerg1l - Glutamatergic
L5a Pde1c - Glutamatergic
L5a Batf3 - Glutamatergic
L5 Ucma - Glutamatergic
L5b Tph2 - Glutamatergic
L5b Cdh13 - Glutamatergic
L5 Chrna6 - Glutamatergic
L6a Mgp - Glutamatergic
L6a Sla - Glutamatergic
L6a Car12 - Glutamatergic
L6a Syt17 - Glutamatergic
L6b Serpinb11 - Glutamatergic
L6b Rgs12 - Glutamatergic
Gja1 - Astrocyte
Pdgfra - OPC
96_Rik - Oligodendrocyte
Opalin - Oligodendrocyte
Ctss - Microglia
Tbc1d4 - Endothelial
Myl9 - Endothelial
Htr3a, E18, type0 - GABA-ergic
Htr3a, P2, type1 - GABA-ergic
Htr3a, P2, type2 - GABA-ergic
Htr3a, P2, type3 - GABA-ergic
Htr3a, P5, type1 - GABA-ergic
Htr3a, P5, type2 - GABA-ergic
Htr3a, P5, type3 - GABA-ergic
(Lumbar) DRG neuron
(Precursor) Olfactory neuron
(Early Immature) Olfactory neuron
(Late Immature) Olfactory neuron
(Mature) Olfactory neuron
(Cochlear) Hair cell
(Vestibular) Hair cell
(Cochlear) Supporting cell
(Vestibular) Supporting cell
(Cochlear) Non-sensory cell
(Vestibular) Transitional epithelial
(E14.5) fibroblast
(2d) Ascl1 induced neuron
(5d) Ascl1 induced neuron
(20d) Ascl1 induced neuron
(22d) Ascl1 induced neuron
(22d) Ascl1,Brn2,Myt1l iNeuron
Cnih1
(ENSMUSG00000015759)
Gene Expression (NAUC)
12
9
6
3
0
Exon (PSI)
CT_03483
Cell Groups
Htr3a - GABA
Induced Neurons
Lumbar DRG
Neonatal Inner Ear
Olfactory Bulb
V1 Ctx - GABA
V1 Ctx - Glia
V1 Ctx - Gluta.
PSI
100
75
50
25
0
Exoc6b
(ENSMUSG00000033769)
Gene Expression (NAUC)
6
4
2
0
Exon (PSI)
CT_05400
CT_05399
CT_05398
CT_05397
Cell Groups
Htr3a - GABA
Induced Neurons
Lumbar DRG
Neonatal Inner Ear
Olfactory Bulb
V1 Ctx - GABA
V1 Ctx - Glia
V1 Ctx - Gluta.
PSI
100
75
50
25
0

**Supplemental Figure 7. Alternative splicing analysis of full-length single-cell RNA-Seq data**. As a proof of concept, we demonstrate that our splicing analysis method can detect alternative exons in single-cell RNA-Seq data. We used data generated from V1 mouse cortex and merged junction counts based on the 49 cell types identified in Tasic et al.[13] (**A**) Splicing analysis reveals a wide degree of variability in exon PSI that can differentiate not only between glia and neuron but also between individual subtypes of glutamatergic or GABA-ergic neurons. (**B**) Furthermore, we can robustly identify mutually exclusive exon splicing patterns used by different neuronal subtypes. (**C**) Analysis of SmartSeq2 single-cell data generated from E14, E18 and P2 Chx10:GFP-positive mouse retinal progenitor cells[14] reveals that even at early stages of development, we can begin to detect photoreceptor-specific exons. Rod-specific exons in *Atp1b2*, *Ttc8*, and *Slc4a7* are primarily detected in photoreceptor precursors. Neurogenic and amacrine cells also appear to utilize the exon in *Slc4a7* at low levels. In contrast, a pan-neuronal exon in *Sgce* is fully utilized in retinal ganglion cells and photoreceptor precursors but is only moderately incorporated in neurogenic cells. In addition to V1 cortex, we have also analyzed several other single-cell studies in the CellTower compilation of ASCOT (http://ascot.cs.jhu.edu) and can identify (**D**) the same cell type-specific exons in *Sptan1*, *Cnih1*, and *Exoc6* as we found in Fig. 1B-D. The exon in *Sptan1* is only found in cochlear/vestibular hair cells, the exon in *Cnih1* is primarily enriched in glutamatergic neurons, and the exon in Exoc6 is only detected in mature oligodendrocytes.

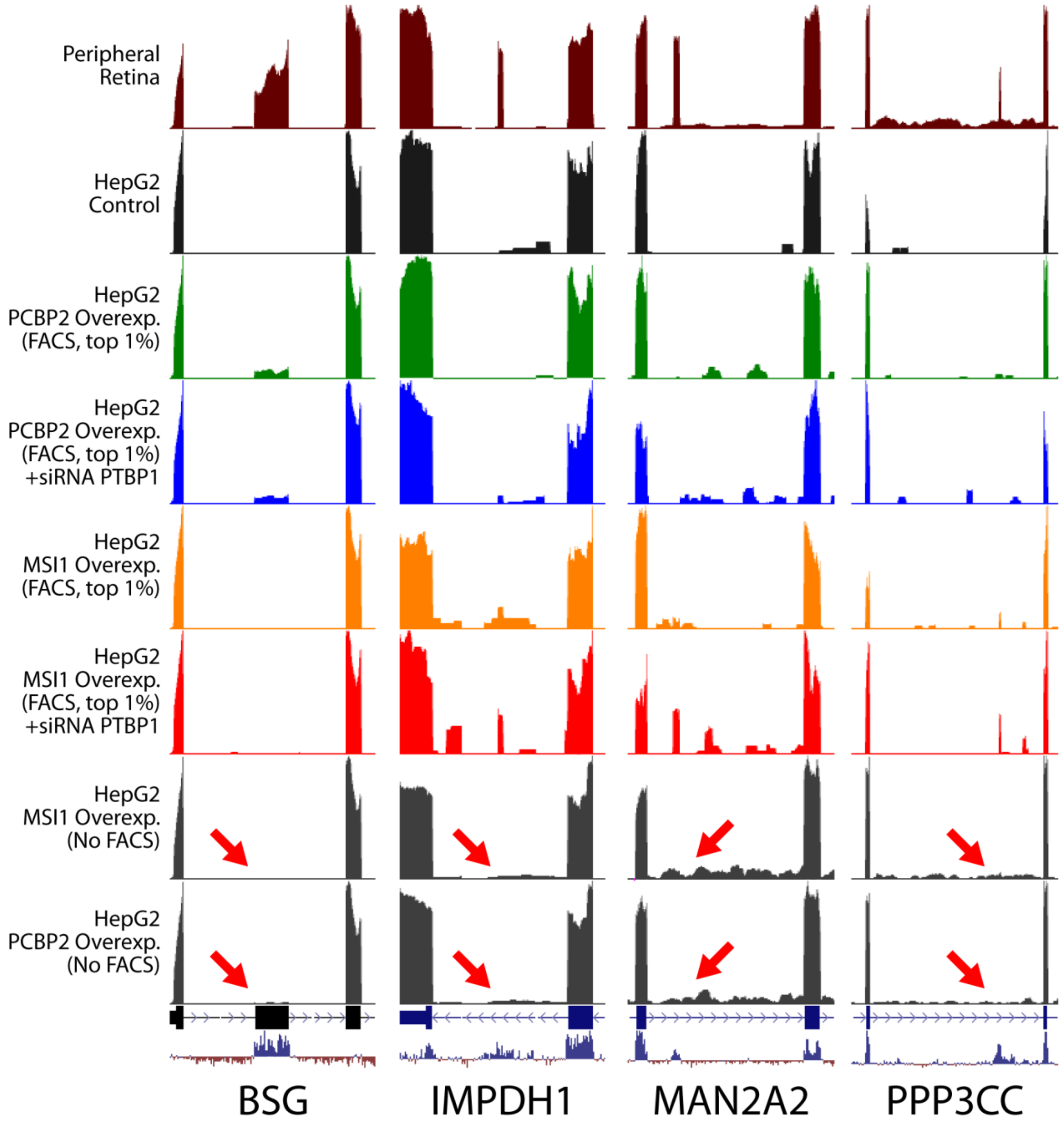


**Supplemental Figure 8. Activation of rod-specific exons in HepG2 cells requires FACS isolation of the most strongly expressing cells**

Normal transfection of MSI1 or PCBP2 is unable to strongly activate rod-specific exons in HepG2 (bottom two rows, red arrows). Rod-specific exon activation is only achieved after isolating the top expressing cells with FACS.

A

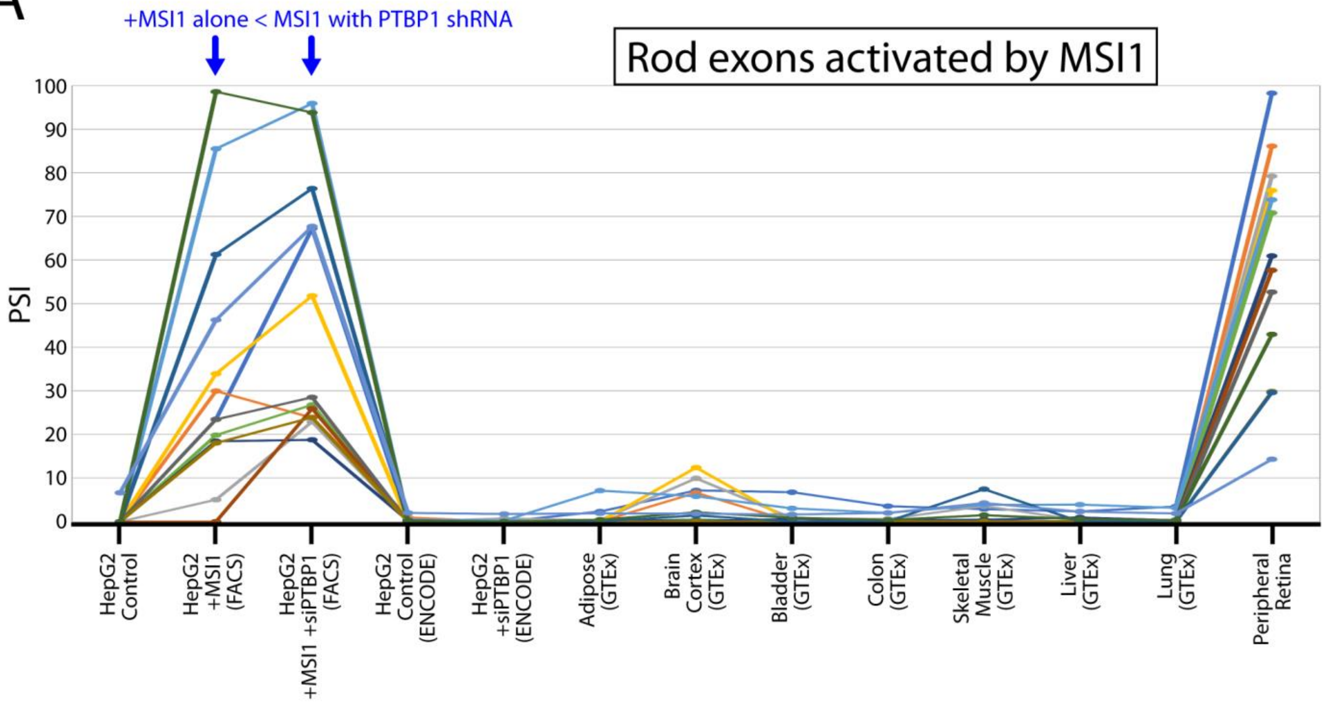

+MSI1 alone < MSI1 with PTBP1 shRNA
Rod exons activated by MSI1
PSI
100
90
80
70
60
50
40
30
20
10
0
HepG2 Control
HepG2 +MSI1 (FACS)
HepG2 +MSI1 +siPTBP1 (FACS)
HepG2 Control (ENCODE)
HepG2 +siPTBP1 (ENCODE)
Adipose (GTEx)
Brain Cortex (GTEx)
Bladder (GTEx)
Colon (GTEx)
Skeletal Muscle (GTEx)
Liver (GTEx)
Lung (GTEx)
Peripheral Retina


B

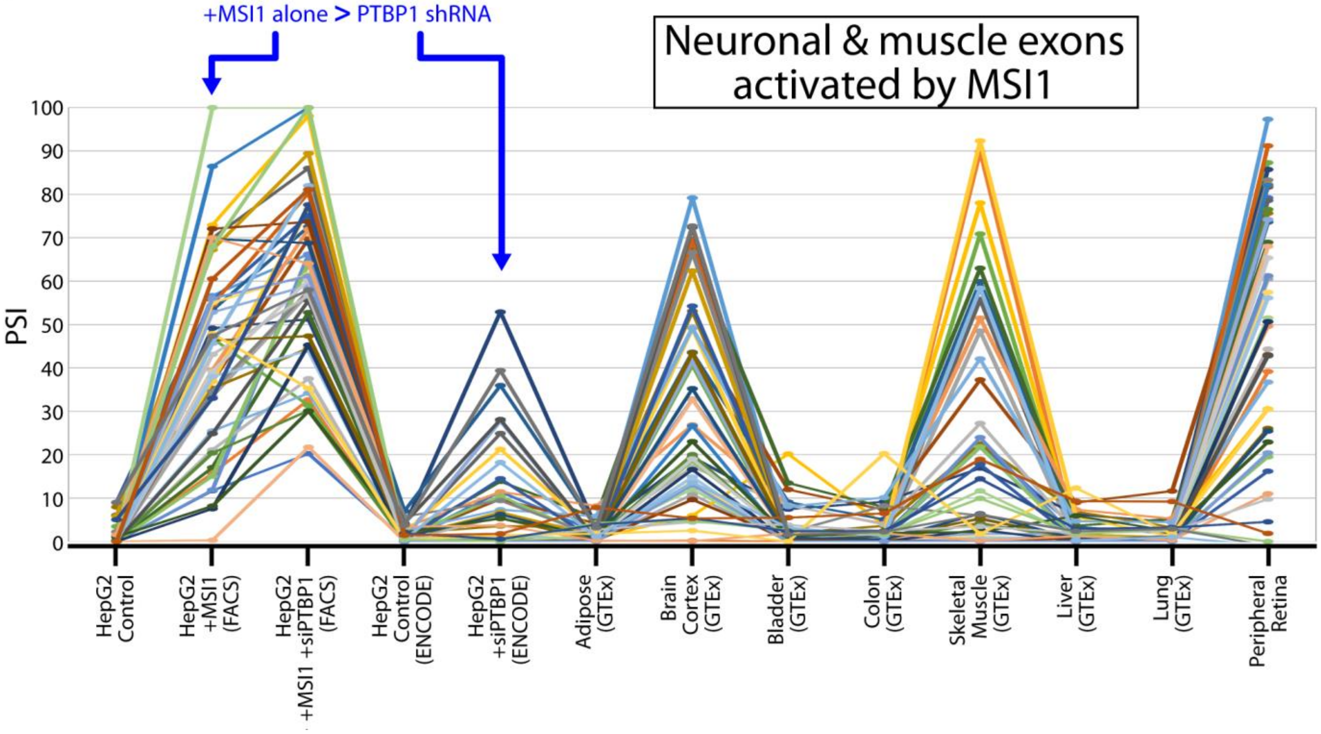

+MSI1 alone > PTBP1 shRNA
Neuronal & muscle exons activated by MSI1
PSI
100
90
80
70
60
50
40
30
20
10
0
HepG2 Control
HepG2 +MSI1 (FACS)
HepG2 +MSI1 +siPTBP1 (FACS)
HepG2 Control (ENCODE)
HepG2 +siPTBP1 (ENCODE)
Adipose (GTEx)
Brain Cortex (GTEx)
Bladder (GTEx)
Colon (GTEx)
Skeletal Muscle (GTEx)
Liver (GTEx)
Lung (GTEx)
Peripheral Retina

**Supplemental Figure 9. Increased *MSI1* expression and *PTBP1* downregulation may interact synergistically to activate rod/neuron-specific alternative exons**

Exons activated by *MSI1* overexpression in HepG2 cells can be grouped into two categories: rod-specific exons (**A**) and neuronal/muscle exons that are also regulated by *PTBP1* (**B**). Two lines of evidence suggest that *MSI1* interacts synergistically with downregulation of *PTBP1*. First, activation of rod exons is stronger when *MSI1* is combined with *PTBP1* knockdown (**A**). Second, overexpression of MSI1 alone can activate neuronal/muscle exons more effectively than *PTBP1* knockdown in HepG2 cells (**B**).

#### Upstream flanking sequences of rod-specific exons (-75 to +20 from 3' splice site)

YELLOW = PTBP1/PTBP2 binding site (repetitive CU/UC)

3'SS

| | | |
|---|---|---|
| (human) | ADGRV1 | AUUUGCUUCUUUGUUGCUUUAUUGUGUUCUCUGCUUCUUUCAUGACCUUUUAUCUUUUUCUUUUCAUGCUUUUAGAAAAGACUCAGUUAAUUCAU |
| (mouse) | Adgrv1 | AGUUUAAUAUUUAUUUUGUGAAUUCAGUGCUUGCUUCUAUUCUGACCUUUGAUCUUUUUCUUUUCAUGCUUUUAGAAAACUUAAUUCAUUACAGA |
| (human) | AMPH | AUACAGUGAAGAAAAAGCGGGGCCAUGACCACUUAAUGAAUGUUCCUUUCACUUUUUUUCAACAUUCCUCUGCAGAAUGAGGUGGUGAGUGAGCU |
| (mouse) | Amph | AAAACAAGAAGUAGGGUUACAGCCACGUAACAAAUGUCUCUUAUUUUUAUUCCUUUUUAAAACAUUCCUCUGCAGAGCGAGGUGGUGAGUUAGCU |
| (human) | AMPH | UCAUCAAAUCUUACCAUGUUCUGUUUCAUCUUUUGUUUUUGUUUUUUUUCCUUUGCUCUUCUGCAUACUUUAAAGACUGACUCUUCAGAAAGUCU |
| (mouse) | Amph | GACCUAUAUGUUUCUCAUCAAAUAUCUCAUGGUCUAUUCCAUUUUUUCCUCUUUGCUUUUCUGCAUACUUUAAAGACUGACUCUUCAAAAAGUCU |
| (human) | ATP1B2_ex1 | CUGAGUGGCUCACCUGAGUGUCCUUCAUGGUUUCUGUGUAAUUCACCUGUCUCUCCCUAUCUUCUUUGCUCCUAGAGGCCCCAUCACCAUAGAAA |
| (mouse) | Atp1b2_ex1 | CUGUCCUCUCUAGAAACUCUCCUCCUUCCCACUGAGCCUUGACUGUUUAUCUGUUCUUUCUUGGCUCUGCUCCAGAGACUGAUUCCCAAGGACGG |
| (human) | ATP1B2_ex2 | CUCCCUCCACAUCCCCUUCCUUGCUUCCUAUUCAACCCUUAAUCAUGUAUCUCUUCUUUCUUGGCUCUGCUCCAGAAACUGAUUCCUGAGGAUGG |
| (mouse) | Atp1b2_ex2 | UUACCUGCCCACCUGGUUAGACUUCCUGGUUUCUGAGUGCUUCACCCAUAUCUCCCUAUCUUUUUGUGCUUUCAGAGGCCACAGCACUAUAGGGA |
| (human) | BSG | GCGAGGCCUUCCCCAGGAGCUGCCCUCCCAAGUGCCCCAGGCACUAACAAGACCCCACGCGUGCUCUCCCCACAGCCGGCUUCGUCCAGGCGCCG |
| (mouse) | Bsg | UGUUGUGUUGGCCCCUGUGGCAGGCCCUGGUGGCUGUCCUUCCUUGUAGUAACGGUACUAACCCUUUCCCAUAGCUGGUUUCCUCAAGGCACCA |
| (human) | CASK | UAUUUUUCUUGAAUCAUCAUCUUUGCUUUGUUUUUGUUUUGCCUGUUUUCCUGCCCUUUUACUAAUUUUCCAUAGCUGCCAAAAUGGUGCAGAAA |
| (mouse) | Cask | UCUUGAAUUAUCAUCUUUGCUUUGUUUUUGUUUCUGUUUUGCCUGUUAUCCUACCUUUUUACUAAUUUUCCAUAGCUGCCAAAAUGGUGCAGAAA |
| (human) | CC2D2A | UUUUUUAAAGAUUAUCUCUGUUACUAAAGCCUGCCUCUUGUUUCUUUUUCUUUCUUUCUUUCUUGCCUCUGAUAGAUGAGUAACAUGCUUAAGAA |
| (mouse) | Cc2d2a | AUUAUUUUGGUUAAAAGCCUUUAAGCUGUUGCUAAUACUGUGUUUCUAUCAUUUCCUUUUAUUGCCCAUGGUAGAUGAGUGACAUGCUUAAGAA |
| (human) | CLTC | UCACUGUGUAUAAAUGAAUUCCUGAUGGAAACUGUUGUUUUUUGUGCUCUACCUUAUGUCUGUGGAAUGUAACAGGACAUGUAUCAGGUAAAUGC |
| (mouse) | Cltc | UCACUGUGUAUAAAUGAAUUCCUGAUGGAAACUGUUGUUUUUUGUGCUGUACCUUUUGUCUGUGGAAUAUAACAGGACAUGUAUCAGGUAAUUGC |
| (human) | DMD | GGGGGACUUAGUGAGAUUAAAGACUUCUGAUUCACUUUGUAUUUGAAGAAUUUUUUUUCCUCCAUCUUUGCUCAGCUAGUGGAAUCCAUGAUGAA |
| (mouse) | Dmd | GGGGAUUUAGUAGGAUUAAAGAGUUCUGAUUCACUUUGUAUUUGAAGUUUUUUUUUUUCCUCCACCUUUGCUCAGCUAGUGGAAUCCAUGAUGAA |
| (human) | DOC2B | CAAGCAUGUGUCAGCUCCAUGUUCUCAACUGUCCUGUUCCCUUCCCCUCACCCCAUUGCUCCCUGUGGGAGGAAGGUGCCAAAAUUGAUGUUCUC |
| (mouse) | Doc2b | CUCUGCAUUGAACCUGCGUCUGUUCCAUGGUCUCAUCCACACCGUUCCCUUCACACUGCUCCCUCUGGGAGAAAGGUGCCAAAGCUGAUGUUUUC |
| (human) | EFR3A | AAGAUUAUUAGAGCUUUAUAAUUUAAAAUAUUAAAAAAACAUUUAAAAUAUUUUACUCUUAUAAUUUGAAUUUAGCUCUCUUUCCUUCAAGCAAA |
| (mouse) | Efr3a | AUUUAAGGUUUUUUUUUUGUUUGUUUUUUUUUUUAAAUGCAUUUAAAAGAUUAAAUAAUUUUACUUGGAUUUAGCUGUCUUUUCUUCAGGCCCA |
| (human) | FARP2 | GUUCAUAAAUGAAUUGUCCAACUCCCUUUUGGAUAAUCAAAAUGCUCAACAUUUUUUCCUUUGCUUGAUUUUCAGCUAAUAGUGCUGGACCCCCC |
| (mouse) | Farp2 | AUGUUGGUCCCAGGGUGAUGAGUUUUCCAGCUCCUAGUACUGAAAUCCUCAACAUUUUCUUUGCUUGAUUUUCAGCUAAUAGUGUUGGACCCACC |
| (human) | GRAMD1B_ex1 | UUUUGAUUUCUUGUUCUUCCUCUCCUCUUCUUCCCCACCAUUUCCCAAAUACCUUCUUUGCAUGCAUGGUCUUAGUCUCACAAGCGCCUCUCAAA |
| (mouse) | Gramd1b_ex1 | CUGUGACCUUUCACCUUUUUGUUCGUUUGUUUAUUGUUUGUUUGAAUUCUAAUUUUUGUGGUUUUUUUUUCCAGUAUGACAGACUUAAUCUGAU |
| (human) | GRAMD1B_ex2 | CGCUGUGGCACUGACCCUUCUUUGACUUUCCUUUUUUUUUUUUUUCAAAUCUAAUUUUUGUGGUUUUCUUUCCAGUAUGACAGACUUAACCUGAU |
| (mouse) | Gramd1b_ex2 | GAACUUGUCUUUUGAUUUCUUGUUCUUCCUCUUCUCCCCACUGUUUCCAAACACCUUUGCAUGCAUGGUCUUAGUCUCACAAGCGCCUCUCAAA |
| (human) | HCN1 | UUCUUGAGAGAUCUGGGUCCCUAUCUAUCUCUUCUAAGUAUUCUGGGUUUAUUUAUUAAUGUCUAUUUUUGCUAGCUAAACAAACUGAUUUCCCG |
| (mouse) | Hcn1 | GUAUAGAAAGAUCCGAGGCGGUACCUGCUCCUGCUAGGUGCUCUGGAUUGAUUUGUUCAUGUCUGUUUUCACUAGCUAUGCAAUCUGAUAACCCA |
| (human) | IFT140 | UGUCUUGCACACACUCAUGCAUCUGUUCUUUUCUCUCUCAUUUCUCACCUUCCCGCAUGCGUCCACUCCUCAUAGUUGUUGUUCCAGAGAGCAGG |
| (mouse) | Ift140 | UGCAUUUCUGUUUCAUCCACACAUUUAUGCAUGCUUUCUUUUUUCUCUGCUCCCUCUCUGUGCCUGCAUUCAUAGUUGUUAUUCCACAGAAGAGG |
| (human) | IMPDH1 | CCCUGGUCAGGAGCCCUGUCCCCAGGUAUCAGCGGGCGUCUAACCUGUGUCUCUUUCCGCCCCACCCUUCCGUAGCUAUACCUUUCUGCCGUGUA |
| (mouse) | Impdh1 | UGGUCGAGAGGACCCAUCUCCCUAAUCAUAGGCGGGCAUCUAACCUGUGUCUCUUUCUGCCCCACCCUCCCAUAGCUAUACCUUUCUGCCGUGUA |
| (human) | INTS13 | AUUGUUGGACAUGUAUUAUUUAUUUUAAUUGAACCUUUACCUUUUCUCCCCAUAUCGUAUUUUCUGUUUUGCUAGCCCACAUUCAAUGUCUUUGA |
| (mouse) | Ints13 | UGUUUGCUUGAUAUGCCUUAUUCACUUUAAUUGAACCUUUACUUUUUCUCCCCAUAUCAUAUUUUGUUUUGCUAGCCCACAUUCAGUGUCUUUGA |
| (human) | KCTD5 | GGGCCCGGGGCACCCGCCCACCCGCACUGUCCUGACAGCCUGUCUUCUCCCCCACUGCGCGCUGCACGCCGCCAGAGUGACGACGAGGACCAGGG |
| (mouse) | Kctd5 | AGUCCUCCUGCAGGCCUGCACUGCUUAGGCUGGGCUGACAGCUGUCUUCUUCCUUACUUCCUGCCUUCACACCAGAGUGACCAAGAGGAGCUGAG |
| (human) | KIF1B | AUUUUUUCUGAUUCUCUCCCUUAUUUCUUUUGUUUUUUGCUGCACUAACUAAAGCCUUUUCUAUGUUUUUGCAGGCUCUUUCUUGAUAAGGAUA |

| | |
|---|---|
| (mouse) Kif1b | GUAUAUAUUUGUUUGUUUUUCUUUCUGUCUGUCUUUUGGUUGCUGGACUAACGAAGGCAUUUUCUGUUUCUGCAGGCUCUUGCUUGAUAAGGAUA |
| (human) KIF1B | CCUAUUCGCCAAAUUCCCACCCCACCCUGUGCUCACUCUCCACCCUCUGUAUUCCCCAUCUUCAUCUGCCUUCAGACCAUGAGAGAAAGAUUGAG |
| (mouse) Kif1b | CCACUUACCAAAUCCCCACCCCUGCCCUGCACUCAUUCUCCUCCCUCUACCUUUCCCAUCCCCAUCGGCCUUCAGAGCAUGAAAGAAAGAUUGAG |
| (human) KMT2D | UAAAUAGUUAGAGAUAUAAUAGAAAAAUAAUAAAACCUUACAUUUGUAUAGCACUUUAUACUUUUUUUUUCAAAGCGUUUUCACAUCCAUUAUCU |
| (mouse) Kmt2d | UAAAUAACUAGAAGUAUUCUACAGAAGCACUGCAGCCUUCCGUUUGUAUAGCACUUUAUACUUUUUUUUUCAAAGCGUUUUCACAUCCAUUAUCU |
| (human) MAN2A2 | CCUCUGCCCUGUCCCAGCAUUCUCUAAGCCUGUGCUCCAACCCCGCUCAUCUCAUGGCUUUCUUUGCCCCACCAGCCUGACUUUUUCUCCAAACU |
| (mouse) Man2a2 | UUCUCACCCCAAGCCCGAGCACUUCCUAACCUUGACACAACACUGCUCAUCUCACAGUUUUCUUUGCCUCACCAGCCUGGCUUUUUCUCCAAACU |
| (human) MARCH1 | CAUUCUCUCUCUCUGUUUUUCCAUGGCUUUCCUCUCCAUUCCCAUUUCCCAUCCUUUGCCUGCUGCUCUGUCUAGAUCUGCCAUCUUGCAUGACA |
| (mouse) March1 | UCUUUCUCUCUUUGUUUUCCCAUGACUUUCCUUUCCAUUCCCACUUCCCAUCCUUUUGCCUGUUGCCUUGUAUAGAUCUUCACUCUUGCAUGACA |
| (human) MSI2 | GGAGCUUGGAGACCUGCAGCCAGGGACAACCAUACAUUUAUUUCACCUGGAUCUUUCUUUUUCCCCCUUCAACAGCCCUAUUACCAUAUUUAAAU |
| (mouse) Msi2 | UGGGAGACUCAUUCCCAGCCUAGGAGAACCGUGUACAAUUGUUUCACCUGGAUCUUUCUUUUUCCCCUUCAACAGCCCUAUUACCAUAUUUAAAU |
| (human) PLEKHB1 | AGUCUCUUUAGGUGUAGAGGGCCUCUGACACCCCCUCCCUCCCUCCUCCUUCUUGGGUUUCUGUGGCUUGAGCAGGCCCCAGCUGGAGCCACCGU |
| (mouse) Plekhb1 | CCUAGUCUCUUAAGAUGUAAAGGGCAUCUGACAUCUCCCUCCCUCUCCCUUCUUGGGUUUCUGUGGCUGGAGCAGGCCCCAGCUGGAGCCACCGU |
| (human) PPP3CC | CCAUGUAAACAUAAAUUUCUCUUCUCAUAUUUCUGUCUUCAUCUCCUUGUUUCUUGGUGUUUCUUCUCCCUGAAGAUCACUACAUUCCAAGCUAU |
| (mouse) Ppp3cc | UCCAUGUAAACUCCCAUCAUCCUCCCAUGUGUCUGUCUUCAUCUCUUUAUUUCUUGGUGUCUCUUCUCCCUGAAGAUCACUACAUUUCAAGCUGU |
| (human) SLC4A7 | UGUUCCUCUGUCACAGUCUUACUUUUUUUUACUUGGCCUAUCUUUGUGUAUACUUCCUUGUCUCCUGUUGGGCAGGAGUCUGCCUCCUGGCACUG |
| (mouse) Slc4a7 | UUUUGUUCCUCUGUCACAGUUUUACUUUUUUACUUGGCCUAUUCAUGCAUACCCUCCCUGUCUCUUGUUGGGCAGGAGUCAGCCUCCUGGCACUG |
| (human) STXBP5 | GAUUGUUGUCCCAACCAUACCAUUCUAGUAUUUUUGUCUCCUUUUAUUUCUUCCGCUUUUCAUUGCUUCACUAAGGUGUGCUGAUUGUGACUUCU |
| (mouse) Stxbp5 | CAGCAGAAGCUCUGUUCCAACCAUGCCAUCUAGUGUCUGUCUCGUUUCUCUCCCUCCCCCUCAUGGCUCACUGAGGUAUGCUGCUUGGGACUCCU |
| (human) TTC8 | GGUUGGUCCUUAGGACUUUUUAUUUUAGAAACUGUAUUUCUUAAAACUGUUUACUGCCUUCUUAAUGCUUUCCAGGAACCAGAUCCUGAAUUGCC |
| (mouse) Ttc8 | UAAUACUCAUUAAGACUUUUGUCUUCAGAAAUUGCAUGCCUUAAAAAUGUUUACAACCUUCUUAACGCUUUCCAGGAACCAGCUCCUGAUUUGCC |
| (human) UNC13B | GUCCCUGAGGGUCUCUCUGAUGUCCCUGAUUUUUUCACUCUCUUUGUUGCUCAUUGGUGUCUGCUCUGUCACAAGCAACCACUGUCAUUCAUAUA |
| (mouse) Unc13b | AAUAGAAAGGAGGGUCUCUGGCAUCCCCGAUUUUCACACUUUUUGUUGCUCAUUGGUAUCUCUGUUUGUCACAAGCCACUUCUAUCAUUCGUAUU |
| (human) UNC13B | CUUCUUUCUUCCUUUUCCCUGUCUCUUCCUUUUUCUUUGCUUUAUUGUUCCUCCAAUUUUAUCCCUUCACAUCAGACAAGGCUCACAGGCUAAAG |
| (mouse) Unc13b | AACUCAUUUUUUCCUUUUCUCUGUUUCUCCUUUUUUUUUUUUCCAUUUUCUUUAUUCCUCUAAUUUUAACAUCAGACAAGACUGGCAGGCUAGGA |

#### Downstream flanking sequences of rod-specific exons (-20 to +75 from 5' splice site)

RED = MSI1 binding site (UAG clusters, <15bp apart)

5'SS ↓

| | | | |
|---|---|---|---|
| (human) | ADGRV1 | AAUUCAUUACAGACUGAAAG | GUAAAACCUGAGCCUUGGUACAAACUUAUCUAGAUUCCUUUGAAAAGAUGUAUUCCUAUGUUUAAACACAAACUC |
| (mouse) | Adgrv1 | AAUUCAUUACAGACUGAAAG | GUAAAACAUAACUUGGUAGAAACUUAAUUAGACCCUUCAAUAGAUAAAGUCUAAGUAUUGAUGGAGAGUGUGUGA |
| (human) | AMPH | UUCCUCUGCAGAAUGAGGUG | GUGAGUGAGCUCAGUGGCAUCUCUAUUAAGCUUAGAUUGGGGCUAUGUUUCCAUCUGAGGCCAUGAUUGCUGGCA |
| (mouse) | Amph | UUCCUCUGCAGAGCGAGGUG | GUGAGUUAGCUCUGUGGGCAAUCUCUAGACUAGAGCAUAGACCGAGCUGUUUCUAGUCAGAGGUCAUGAUUGCUG |
| (human) | AMPH | UAGAGGCUGACAAAGACGAG | GUACUUACCCUGGUCUGUAGCAUAGACUAGCCAAAGACUUCUGUAGGCUCUGUUUUUCCUCAUGUCUGUCACCUA |
| (mouse) | Amph | AGUUAGAAGAUCCUAUGUGG | GUAUGUGUGGAGACAUACCAAAGGGUAGACACUGGAGCUAUUGGUAUCAAGGCAAUAGAAGGUACUGAGGGAUUU |
| (human) | ATP1B2_ex1 | AUCACCAUAGAAACAAGGGG | GUAAGAGUGGGCUUUUGGGCUCCACUGUAGCUUGAACUCCGAGGGCCCCGCACUCCUCUUGCUUCUCUCUGGGAU |
| (mouse) | Atp1b2_ex1 | GACUGAUUCCCAAGGACGGG | GUAAGAACUUGGGGAUUGAUGGUGGAGUUAGAAGGCCCUCACCGUGUUGUCAGCACCCUUAGAAGACCUAGUCUG |
| (human) | ATP1B2_ex2 | AACUGAUUCCUGAGGAUGGG | GUAAGAACUUGGGGUAGGAGUGGAGUAGGAGGCUUUCGUGCCAUUAGCAGCCUUCAGGAGUUCCUAGAAUGAUGA |
| (mouse) | Atp1b2_ex2 | AGCACUAUAGGGACAAGGGG | GUAAGAGUGGGCGCCUAUGCAGUUUUAGCUCUAAGAGGCUCUUAGCCCUAUUGCUUCUCUCUAGGAUAAAGAGAG |
| (human) | BSG | AGUCGUGCUAGUCCUGGAAC | GUGAGUGGCGGGCACCUCCCUCCCCGCCUCCCUCAGUUUCCCUCCUGUGCCGCUCGCCUCCCGGCUCCUGCUCAG |
| (mouse) | Bsg | GAGCGUGGUGGUCCUUGAAC | GUGAGUGGCCAGGGCACCUGUUUCUGUUUACCUCUCGUGCCACCCAACCCUGGCUCCCACUUUGUAGAACCAGAU |
| (human) | CASK | AAUGGUGCAGAAAAUUGCCU | GUAAGUACCAGCUAGGAGGUGGUGAAAUACAGCCGUGUUUCUUUUAAUAGAUUUAGAAUGUAGACACUUUGUUCC |
| (mouse) | Cask | AAUGGUGCAGAAAAUUGCCU | GUAAGUACCAGCUAGGAGGUGAUGAAAUACAGUCUGUGUCUUUUAAUAGAUGUAGAAUGUAGACAUUUGGUUCCC |
| (human) | CC2D2A | UGAGUAACAUGCUUAAGAAG | GUACUAAGUAUUUUGUUUUACUAUUUUAUUGGUCAAUAGUUAUUAGGCUUUAAUUAGCUGCUUCUGUCCACUGCC |
| (mouse) | Cc2d2a | UGAGUGACAUGCUUAAGAAG | GUACCGAGCAUCUUGUUUUAUAAUCCAGCAGUCAUCAGUUAAACGCUUUUACUAGCUGCUUUUGUUUACUGCCUC |
| (human) | CLTC | UGUAACAGGACAUGUAUCAG | GUAAAUGCCCAGUACAGGUUUUACUUACCUGUCUAUAACAUCUUCUAUGUCAGUCUAUAAGUAGGAUUGUAGUUU |
| (mouse) | Cltc | UAUAACAGGACAUGUAUCAG | GUAAUUGCCCCACACAGGUCUUACCCUUACCUGCCUGUCUAUAACAUCUCUGUCAGUCUAUAAGUAGGACUCUAU |
| (human) | DMD | UGAAUUCUCAUCUCCAAGGG | GUAAGCAGUUUUUAGUAAAGCCCAGUAGCUGACUUAUGACUCCUUAGAAAUAGCAUUGAUUCCUUCCUUCUCCUG |
| (mouse) | Dmd | UGAAUUCUCAUCUCCAAGGG | GUAAGCUGUUUGUAGUAAAGCCUGAUAGCUGCCUUAGGUCUCUUUAGAAGUAGCAUCGAUUCCUUCCUUUUCCUG |
| (human) | DOC2B | AAUUGAUGUUCUCUCGCAAG | GUAAGGGCCUGCAUGGCCAGUAGAGUCUCCCUAGUAGUGCCAUGUAAGGGCCUGAGCCAGCCCAGCUCAGAUCCA |
| (mouse) | Doc2b | AGCUGAUGUUUUCCAGAAAG | GUAAGGGCCCGUGUGGCCAUUAGAGAGCCUUGUAGGACUGUGGAAAGGCCUGGGCCAGCCCAGCACACAUGGAAU |
| (human) | EFR3A | ACCCAGGCACAAGCACAAAG | GUAAAUCCAGUUGUCAGUUUGGCCUGCCUUUUCCCUUACCUGUCCUCUAGGAAAAUAGCUUAAUAAAAUGUGCCU |
| (mouse) | Efr3a | ACCCAAGCAGAAGUACAAAG | GUAACUAGCUAUAUAGCUCCCCCUGGUGUCGCCUGCCCGUCCUGUAGGGAAAGCACUCCCUGGAUGCAGUGUUUA |
| (human) | FARP2 | UAAUAGUGCUGGACCCCCCG | GUGAGUAUUUGGGUGCCCUCUGGCCCAUGGCUCACCCCCUGGGGUGCAGCUGCCAUGGCGCAGGGAUACCAGUGU |
| (mouse) | Farp2 | UAAUAGUGUUGGACCCACCG | GUGAGUAUUUCUGUGUCUCUAGACCAGACCUCAGUCACUUGGCAGCAGUUGCCAUAAUGUAGGGACACAAGUGUG |
| (human) | GRAMD1B_ex1 | CUCACAAGCGCCUCUCAAAA | GUAAGCCAUCUUUUGUCUCUGUCCUUUUAUCUCCCCUCUAGUGUCGAUCUGUCUGUAGUGCUCAGCACCCCCUCC |
| (mouse) | Gramd1b_ex1 | ACAGACUUAAUCUGAUCAAA | GUAAGACAUUGUUUUGUUUUGUUUUGAUUUUUUUUUUGCCUCCCGGCUCCAGUUUAUGCUUAAUGUAGAGCUCCU |
| (human) | GRAMD1B_ex2 | ACAGACUUAACCUGAUCAAA | GUAAGAAUUGUUUUUUAUUUCUAACUCCUACUAAAUGUAGAGCUCUCCUAGUCCCUGCCCCCCUACCUCCUCCAC |
| (mouse) | Gramd1b_ex2 | CUCACAAGCGCCUCUCAAAA | GUAAGCCAUCUUUUGUCUCUGUCUUUUAAUCUCCCCUCUAGUAUUGAAUUGUCUGUAGUGCUCAGCAUCCCUCCU |
| (human) | HCN1 | AAAACAAUUUGAAUCUUCAG | GUAAAAAGCAAUGAUAAAAAUUCUAGAUAGCAUUGAAAUAGAACUAAAAAUAUGAUAUAUGAGUCUAUAUUACAA |
| (mouse) | Hcn1 | ACUUCAGUUUGAAUCUUCAG | GUAAAAAGCACUGGUAUAAACUCUAGAUAGCACUAAAAUAGUUCUUAAUGCUAAUAUCACGAGUCUAUAUUACAU |
| (human) | IFT140 | CAGGCGUGUAUGACGCAGUC | GUAAGUACAGACGCCCAGGUCGUGUGUCAUUUGCUUCACCGGGUUGUUUUCCUGAGAUCCUUUUCCUCCCUUAUU |
| (mouse) | Ift140 | GAGGUGUCUAUGGUGCAGUC | GUAAGUAUGAAGCCCCAGGCGUGGCUUGUCACCUUGGUUUUGUCAUGUCCCUGAGAUGUCUUUCUUCUCAUUGUG |
| (human) | IMPDH1 | UAGCUAUACCUUUCUGCCGU | GUAAGUAACCGCGCUGCCCGGGCCAAGCGUAGAGUAGAGCAGGGGCAGUCUGGGUGGGACCUGGGCUGGAGGUGG |
| (mouse) | Impdh1 | UAGCUAUACCUUUCUGCCGU | GUAAGUAACUGCGCUGCCGGGGCCAGGUGUAGAAUAGAGCAGGGGCAGUCUGGGUGGGCCAUGGGCUGGGAGAAU |
| (human) | INTS13 | CAUUCAAUGUCUUUGACCAG | GUCAGUAAAUACUGAAAAGCAACCAGAAAGGAUCACAGAAUAAAAUGCUAUAUUUUUGUUUUCUAAUAAGAACAC |
| (mouse) | Ints13 | CAUUCAGUGUCUUUGAACAG | GUCAGUAAAUACUGAAAAUCAAAAAGAAAGGAUCACAGACCAAAGUAUAUUAUUACUAUUUCUAAUAAGAACAUA |
| (human) | KCTD5 | GAUUAAGUGUGAAUGUCAUG | GUGAGCUCCGUGGCCUCCUUGACCCCAUGGCCCCAGCUGACCCUUGCCACAGACCCUCCCUGUGUGUCUGUCCCU |
| (mouse) | Kctd5 | GAUUAAGCAUAAAUGUCACG | GUGAGCUCUGCAGCCCUUUAAGUAGGCCUAGCACCAUAGUCUCCUUCCCAUAUGACCCUGGUGACUUUUGCCUGC |
| (human) | KIF1B | CUUGAUAAGGAUAUUCCUAG | GUAUUGUCUGUUUGCUUUUUUGCACGUAGGAUUCUAGUUUAUCAGAAUGCUUUCUGUCUUAUGUUGCUUUGCCAU |

| | |
|---|---|
| (mouse) Kif1b | CUUGAUAAGGAUAUUCCUAGGUACUUUCUGUUUGCUUUUUUUUUUUUUUUUUUUUUAAUUUAUUUAUUAAUUAUAUGUAAGUACACUGUAGCUG |
| (human) KIF1B | GAGGGUACCCAAACGAGAUGGUUAGUAGCUGAAUUAGCUCCACCUUAGAUGUAGAACCAUCCUCAGGCGCUUGUCCUGAAGCCUGGCUCUCUGCA |
| (mouse) Kif1b | GAGGGUACCCAAGCAAUAUGGUUAGUAGCUAAACUAGCCCUGCCUUAGAUGUAGAACCAACUUGGACAUAUGUGAAGCACUGAUGUCUCUGUUUG |
| (human) KMT2D | CCUCACAACAACCUUAUGAGGUAGGUAGGGCAGGUGGUAUGACCCCCAUUUUACAGCUGAGAAAACUGAGACCCAGAAAGAUUAAAUGACUAGGC |
| (mouse) Kmt2d | CCUCACAACAACCUUAUGAGGUAGGUAGGGCAGGUGGUAUGAUCCCAUUUUAUAGCUGAGAAAACUGAGUCCCAGAAAGACUGUGUGACUAGAUA |
| (human) MAN2A2 | GCAGGAGCGGUAACCGAGAGGUAAGGGUCGGCUCUGGGAGCGGGGAACUAGUACGAGCAAGGUGUAGCUGCGCCAGAGCCCAGGUGACAGAGCCC |
| (mouse) Man2a2 | ACACAAUCAGGAGCUCAAAGGUAAGGGCCAGGGAGGGCCUUGGUGGCCUAGGGCGAGCGGCCUGGCACUGUAGUUGGAGUCUGCUGCCUAGAACC |
| (human) MARCH1 | GAUAAUUUAGAAGUAUGCAGGUCAGUCACUGUAGAGAAAGGCUUGAUGUCAUCCAUAUAUAGUUCUUUUUGCAGUGAGUUAGGUGCGUGUUUGAC |
| (mouse) March1 | GAUAAUUUCGAAGUGUGCAGGUCAGUAGCUGUAGUGAAAGACUGGACGCAACCCAAUUAUAAGUCUUCUGGAUGGGAGCUCAGUAUGUGUUUGUG |
| (human) MSI2 | AUUACCAUAUUUAAAUGCAAGUAUGUCUCUUUUCAUUGACAUUUUAAGUAGCUUAGCCUGACCUUUUUAGAGUAGCAUGUUCCCCCUUUGAACUU |
| (mouse) Msi2 | AUUACCAUAUUUAAAUGCAAGUAUGUCUCUUUUCAUUGGCAUUUUCAGUAGCUUAGCCUGACCUUUUUAGAGUAGCUCUUUGCCUUUUGACCGUC |
| (human) PLEKHB1 | AGGUUGAGAGGCGGAUCUGGGUAAGUGCUGGCUCGGCCCUCCCUGCCUCCAUACUGGCCACCAGGAUGAGCCUCCAAAACCAGGCCCAGUCCAUC |
| (mouse) Plekhb1 | AGGUUGAGAGGCGGAUCUGGGUAAGUUCAAGCUCAGCCCUUCCUGCCUCCUCUCUGACCAUCAGGAUGAGCCUCCAAAGCCACCUGUCCCAUUCC |
| (human) PPP3CC | CAUUCCAAGCUAUCAGAAAGGUAUCCGACUAAACUGUGCUAAGAGCUUAAUGCUUACUCUCUGUCCCACCAUAGUCUAUUGGUUAGACUUAAAAU |
| (mouse) Ppp3cc | CAUUUCAAGCUGUCCAAAAGGUAUCCAACUGGAAUGCACUGCCAGCUAAGUUCUUGCUGUCUGCUCUGCCACUGGUGAUUGGUUAGUCGUCUUGA |
| (human) SLC4A7 | UGGGACUCAAGAGGCCCGCUGUAAGUGCUGGUUUUACUUUUCAUAGAUAGCCCUGUAAGUGGUUGUUUAGUGACUUGUGUGAAACUGCAUAUAAA |
| (mouse) Slc4a7 | UGGGACUCAAGAAGCCGGCUGUAAGUGCUGGUUUUAUCUUUUAGUAGAUAGUGUUGUUAGUGGUUGUUUAGUAGUUUGUAUGAGCUGUGUAUGAA |
| (human) STXBP5 | ACUACGUUCUUCUUUUUUAAGUAAGUCAUUUAUGCCUUGGCAUGUUUGUUCUUUGUAAAAUCAUUGGGAGUUUUUGUGUUCAUUUUUAAAAGAGC |
| (mouse) Stxbp5 | ACUCCGCUCUUCCGUCAUAAGUAAGUCAUUUCUGCCUUGAUACUUUUGUUCUUUGUAAAAUCACUGCUGCCUUUUUAUUUUACUUACUAAAAUAC |
| (human) TTC8 | CUGAAUUGCCAGUGCAUCAGGUAAAGAAAGGUUUAGCUGCAACCUUUAGUUUAGAGAAUUCUUUAAUGCUAUUGCUUAGUUGUUGCUAUAGAUUA |
| (mouse) Ttc8 | CUGAUUUGCCAGUGUCUCAGGUAAAAAGUUAGCUGCAAACUAUGGCUUAGAGAGUAGCUUAGUUGCUGUCGCUUAUUUGCUCCUGAAGAUUGGAA |
| (human) UNC13B | AAAAGUCCUCUUUCUGGGUGGUAAGAACAUUUAGUUGUCCUUUUGCCUCUAGUCUUACACCUUUUCCCUCUUCUCCUGCUCACUGCUUCAUAGAA |
| (mouse) Unc13b | UCGAAAAGUCUUCUUUGGUGGUAAGAACUUUUGGUUGUCCCUUUUGCCUCUAGUCUUAUAUCCUUCCUUCUCUUGCUCACUGCUUUCUAGAAGAG |
| (human) UNC13B | GAGCCCAUGACCAACAAAAGGCCUGUUCUUAACUCAAGCAUCAUAAAUGGCUUUAAGGAAUUGACCAUUUAUGAUGCAAGUGACUUAGUGGUAGA |
| (mouse) Unc13b | GAAACAGUAACCAACAAAAGGCCGGUUCUCCACUCAGGCAUCAUAAAUGGUUUUCAGGAAUUGACCAUUUAUGAUGCAAUUGAUUAGUAGUAGAU |

**Supplemental Figure 10. Upstream and downstream flanking sequences for rod-specific exons**

Motif analysis of the sequences flanking the 31 syntenic rod-specific splicing events reveals motifs for both *PTBP1*/*PTBP2* and *MSI1*. We have previously shown that *PTBP1* and *PTBP2* bind to repetitive CU/UC elements upstream of the 3' splice site[15,16]. Previous work has also shown that *MSI1* preferentially binds to UAG motifs[17–19]. We observe that MSI1 overexpression and PTBP1 downregulation act synergistically (Fig. S9). Binding motifs for PTBP1 (upstream) and MSI1 (downstream) often flank the same exon, suggesting that *PTBP1* and *MSI1* may directly interact.

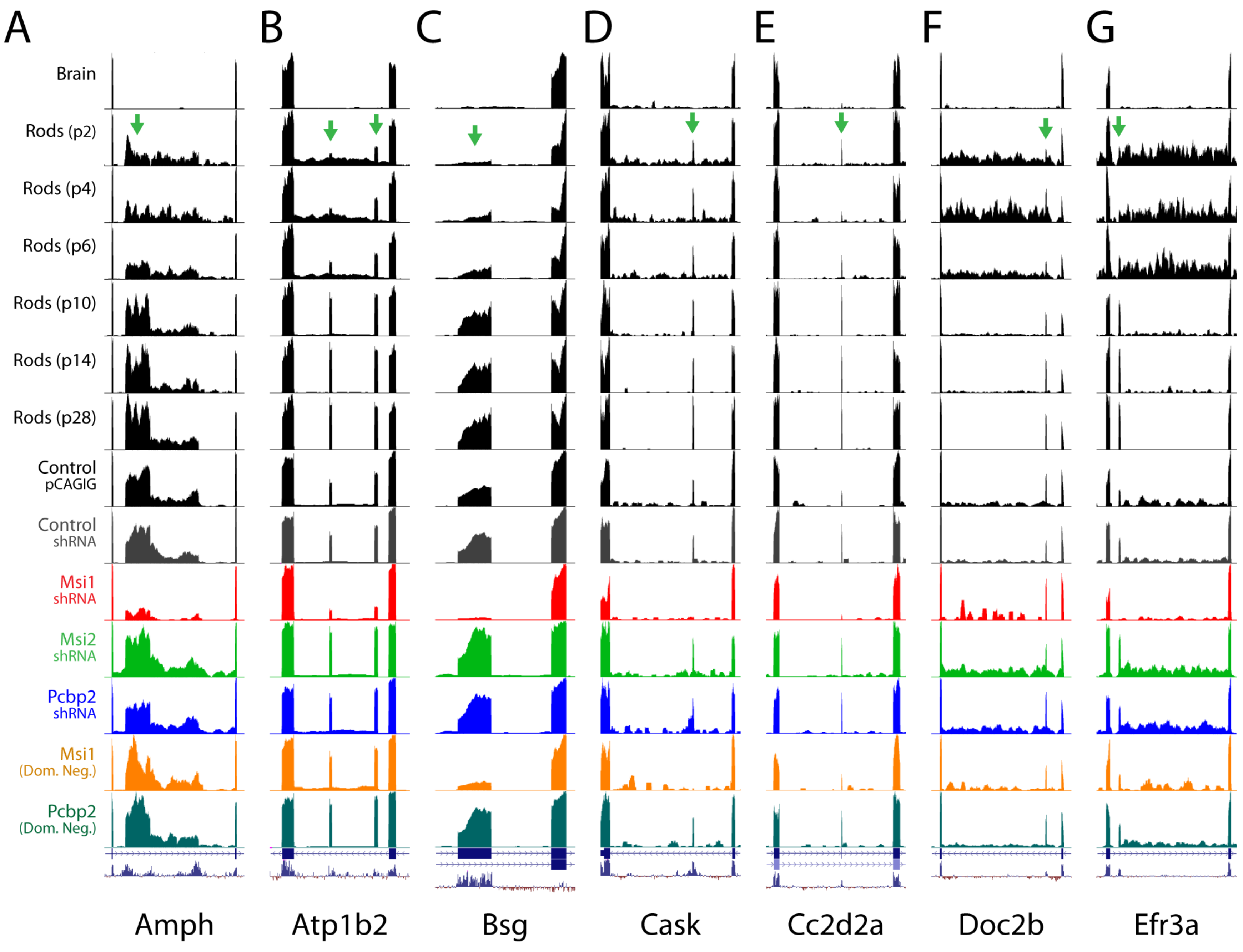
A
B
C
D
E
F
G
Brain
Rods (p2)
Rods (p4)
Rods (p6)
Rods (p10)
Rods (p14)
Rods (p28)
Control pCAGIG
Control shRNA
Msi1 shRNA
Msi2 shRNA
Pcbp2 shRNA
Msi1 (Dom. Neg.)
Pcbp2 (Dom. Neg.)
Amph
Atp1b2
Bsg
Cask
Cc2d2a
Doc2b
Efr3a

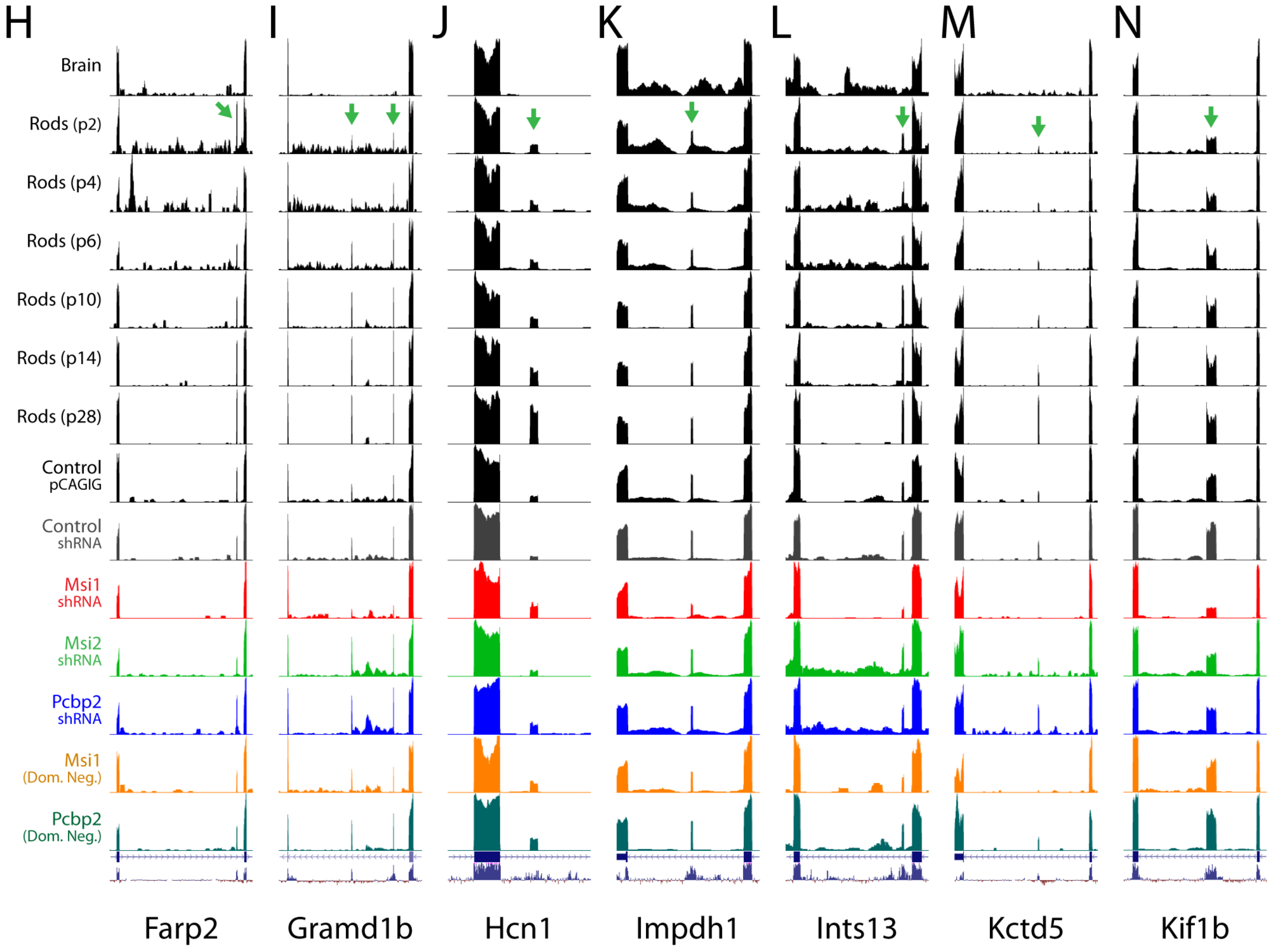
H
I
J
K
L
M
N
Brain
Rods (p2)
Rods (p4)
Rods (p6)
Rods (p10)
Rods (p14)
Rods (p28)
Control pCAGIG
Control shRNA
Msi1 shRNA
Msi2 shRNA
Pcbp2 shRNA
Msi1 (Dom. Neg.)
Pcbp2 (Dom. Neg.)
Farp2
Gramd1b
Hcn1
Impdh1
Ints13
Kctd5
Kif1b

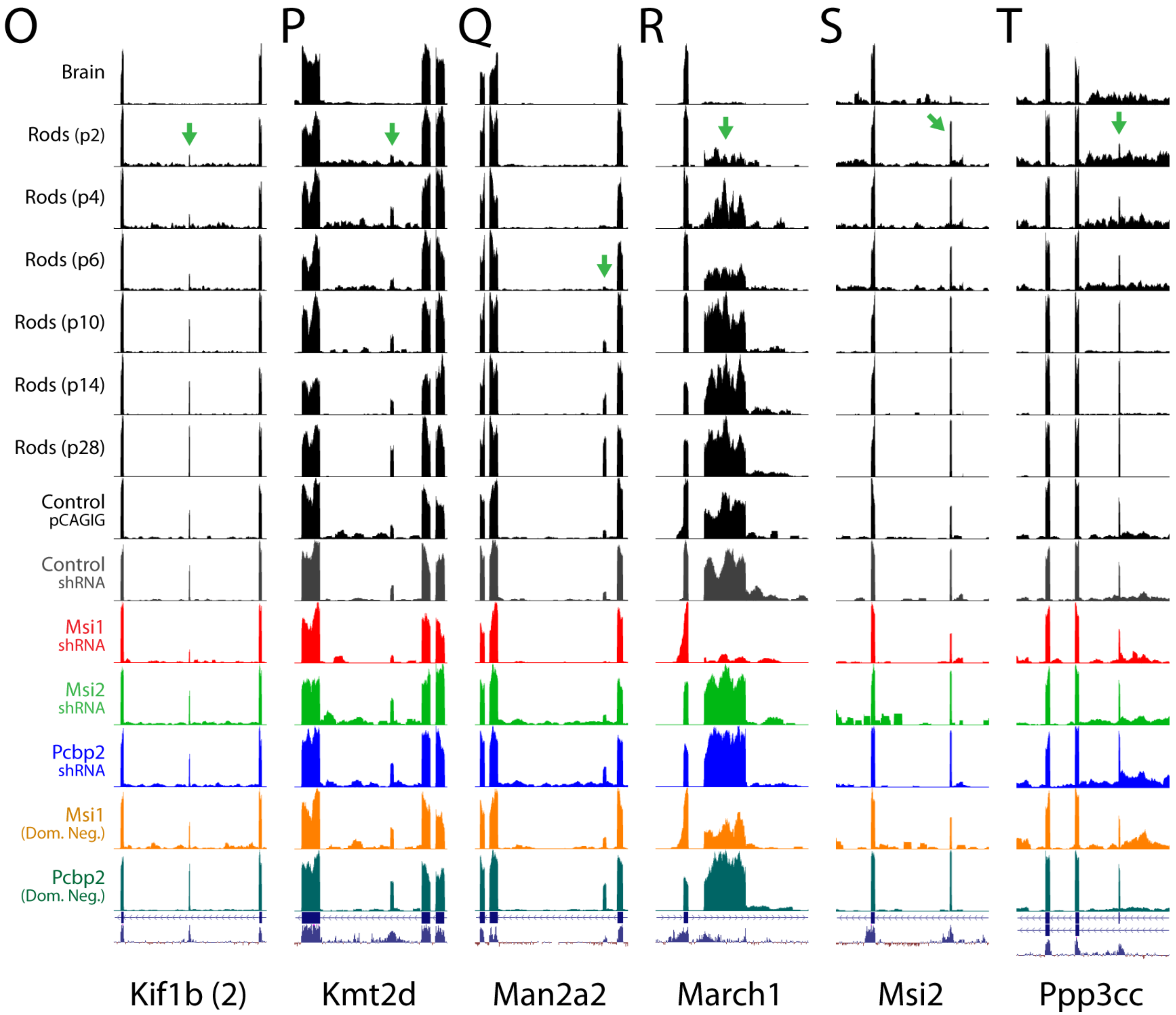
O
P
Q
R
S
T
Brain
Rods (p2)
Rods (p4)
Rods (p6)
Rods (p10)
Rods (p14)
Rods (p28)
Control pCAGIG
Control shRNA
Msi1 shRNA
Msi2 shRNA
Pcbp2 shRNA
Msi1 (Dom. Neg.)
Pcbp2 (Dom. Neg.)
Kif1b (2)
Kmt2d
Man2a2
March1
Msi2
Ppp3cc

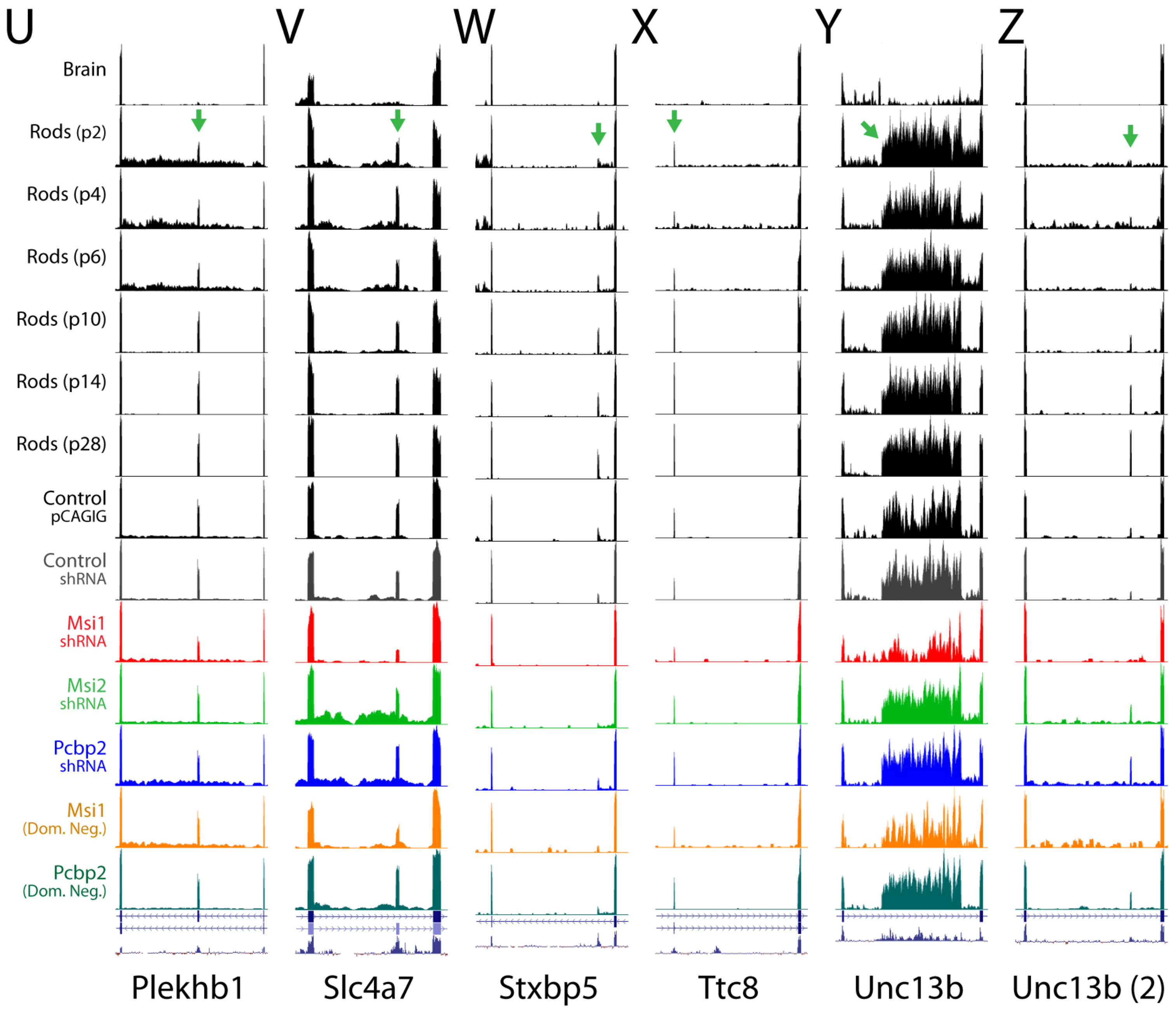
U
V
W
X
Y
Z
Brain
Rods (p2)
Rods (p4)
Rods (p6)
Rods (p10)
Rods (p14)
Rods (p28)
Control pCAGIG
Control shRNA
Msi1 shRNA
Msi2 shRNA
Pcbp2 shRNA
Msi1 (Dom. Neg.)
Pcbp2 (Dom. Neg.)
Plekhb1
Slc4a7
Stxbp5
Ttc8
Unc13b
Unc13b (2)

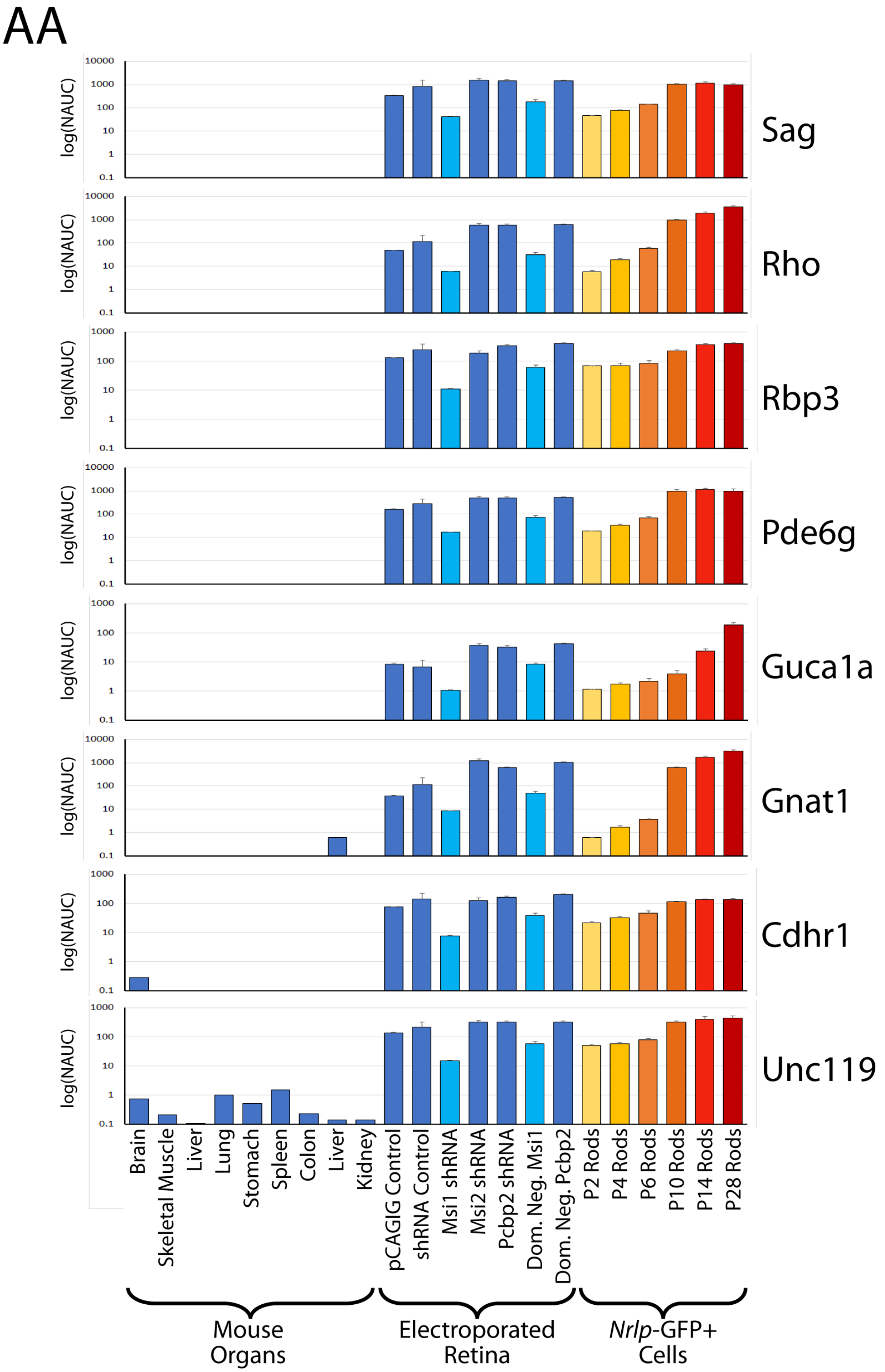

AA
log(NAUC)
Sag
Rho
Rbp3
Pde6g
Guca1a
Gnat1
Cdhr1
Unc119
0.1
1
10
100
1000
10000
Brain
Skeletal Muscle
Liver
Lung
Stomach
Spleen
Colon
Liver
Kidney
pCAGIG Control
shRNA Control
Msi1 shRNA
Msi2 shRNA
Pcbp2 shRNA
Dom. Neg. Msi1
Dom. Neg. Pcbp2
P2 Rods
P4 Rods
P6 Rods
P10 Rods
P14 Rods
P28 Rods
Mouse Organs
Electroporated Retina
Nrlp-GFP+ Cells

**Supplemental Figure 11. *Msi1* is necessary for rod photoreceptor-specific splicing**

(**A**-**Z**) Knockdown of *Msi1* in retina abolishes nearly all rod-specific splicing, while knockdown of *Msi2* and *Pcbp2* have minimal effects on photoreceptor splicing. Expression of a dominant negative *Msi1* protein phenocopies *Msi1* knockdown, although to a lesser degree. Notably, some exons remain unaffected by *Msi1* knockdown, e.g. exons in *Doc2b* (**F**), *Ppp3cc* (**T**), and *Plekhb1* (**U**), while other exons are only partially repressed. (**AA**) Analysis of genes previously associated with photoreceptor development[20] reveals that both *Msi1* knockdown and dominant negative *Msi1* may delay photoreceptor maturation.

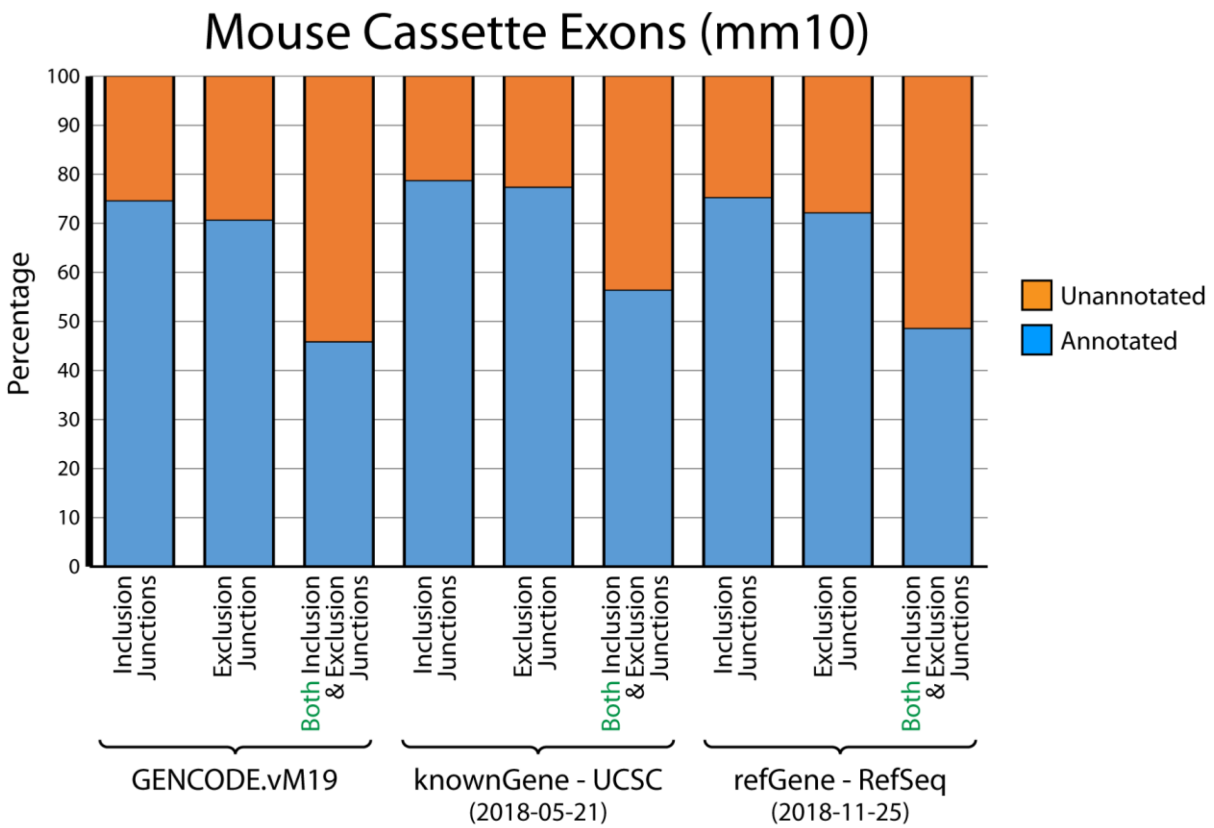

Mouse Cassette Exons (mm10)
Percentage
100
90
80
70
60
50
40
30
20
10
0
Unannotated
Annotated
Inclusion Junctions
Exclusion Junction
Both Inclusion & Exclusion Junctions
GENCODE.vM19
knownGene - UCSC
(2018-05-21)
refGene - RefSeq
(2018-11-25)


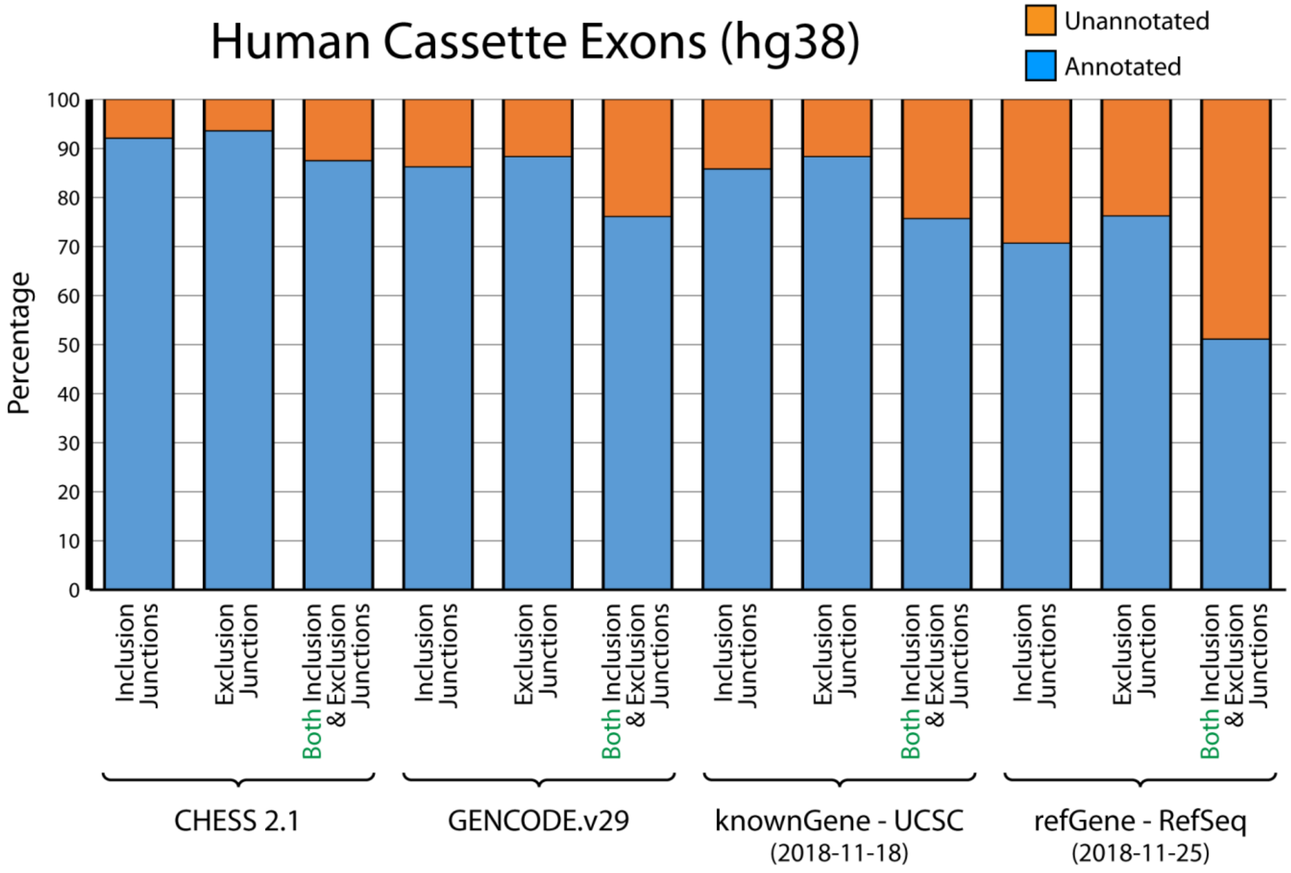

Human Cassette Exons (hg38)
Unannotated
Annotated
Percentage
100
90
80
70
60
50
40
30
20
10
0
Inclusion Junctions
Exclusion Junction
Both Inclusion & Exclusion Junctions
CHESS 2.1
GENCODE.v29
knownGene - UCSC
(2018-11-18)
refGene - RefSeq
(2018-11-25)

**Supplemental Figure 12. Many of the exons detected by ASCOT are unannotated**

To estimate the percentage of unannotated exons that are identified by ASCOT, we looked at high abundance cassette exons (exons detected in at least 75% of samples in MESA or GTEx and exons where the difference between the maximum PSI and the minimum PSI was >75). For mouse cassette exons, we compared against the GENCODE.vM19, knownGene (2018-05-21), and refGene (2018-11-25) gene reference annotations. For human cassette exons, we compared against the CHESS2.1[21], GENCODE.v29, knownGene (2018-11-18), and refGene (2018-11-25) gene reference annotations. For each gene reference annotation, we determined whether the inclusion junctions or exclusion junction of a given cassette exon was annotated and whether the cassette exon was fully annotated ("Both Inclusion & Exclusion Junctions"). We estimate that ~40-60% of mouse and ~10-30% of human cassette exons identified by ASCOT are unannotated.

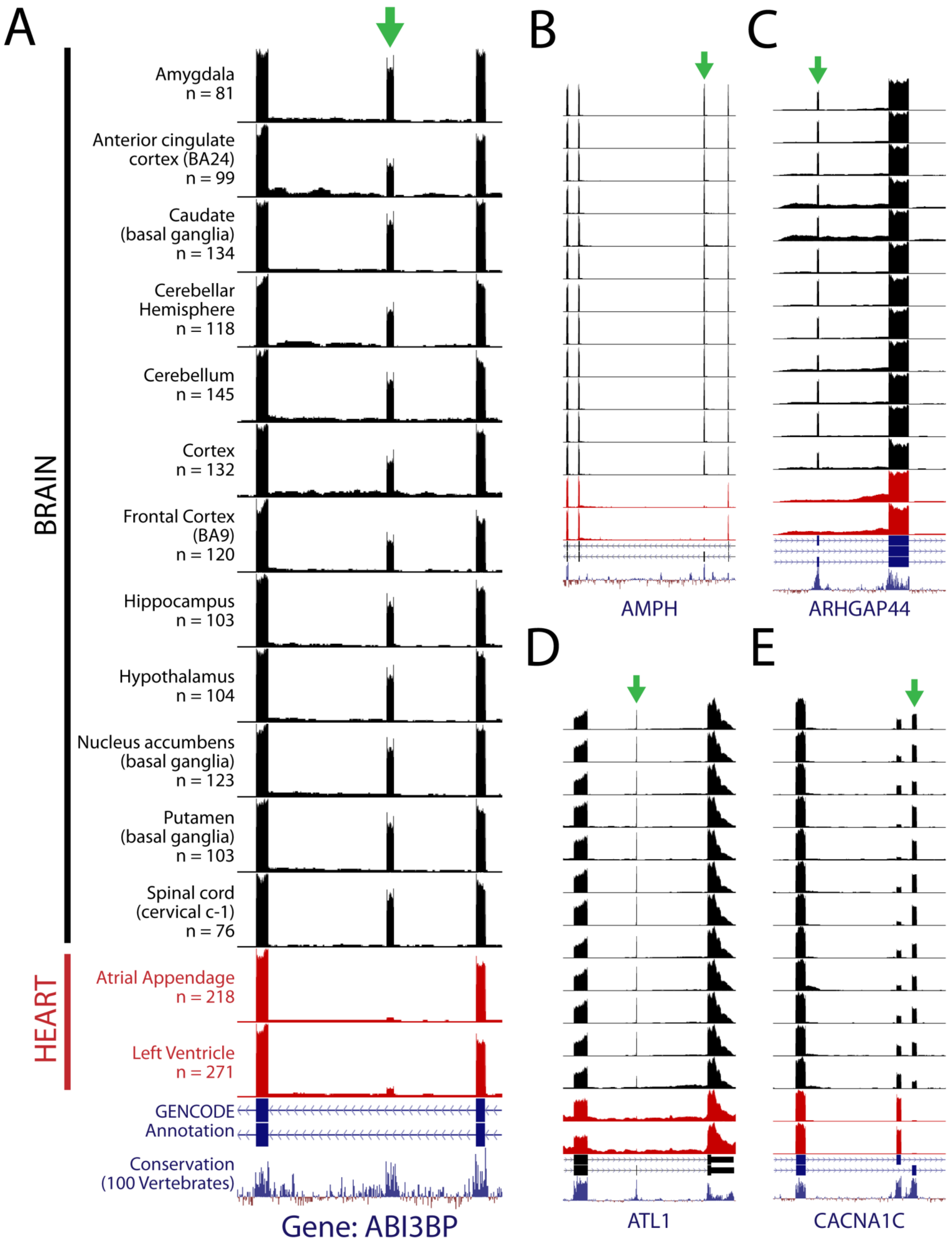

A
B
C
D
E
BRAIN
HEART
Amygdala n = 81
Anterior cingulate cortex (BA24) n = 99
Caudate (basal ganglia) n = 134
Cerebellar Hemisphere n = 118
Cerebellum n = 145
Cortex n = 132
Frontal Cortex (BA9) n = 120
Hippocampus n = 103
Hypothalamus n = 104
Nucleus accumbens (basal ganglia) n = 123
Putamen (basal ganglia) n = 103
Spinal cord (cervical c-1) n = 76
Atrial Appendage n = 218
Left Ventricle n = 271
GENCODE Annotation
Conservation (100 Vertebrates)
Gene: ABI3BP
AMPH
ARHGAP44
ATL1
CACNA1C

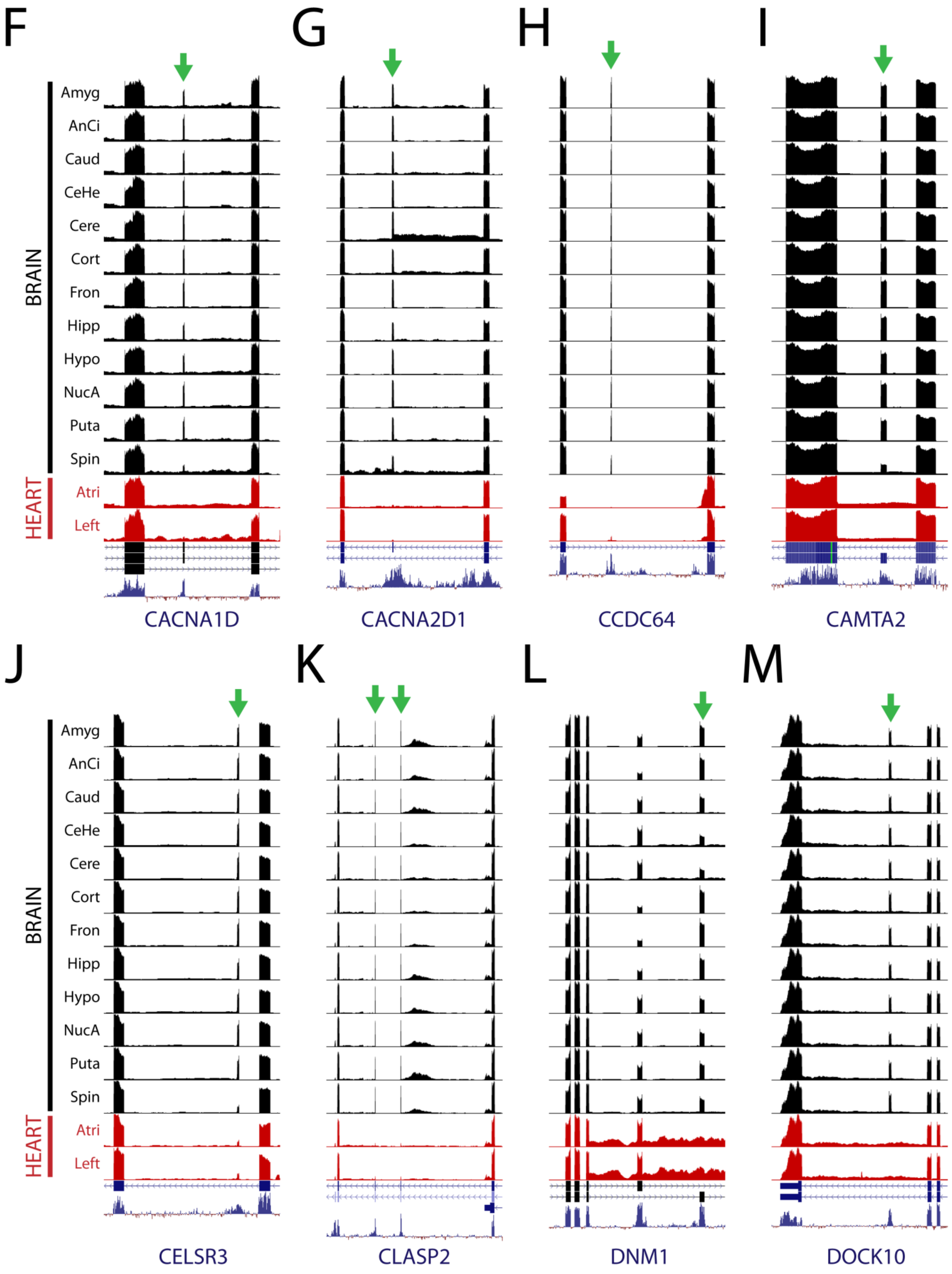
F
G
H
I
BRAIN
HEART
Amyg
AnCi
Caud
CeHe
Cere
Cort
Fron
Hipp
Hypo
NucA
Puta
Spin
Atri
Left
CACNA1D
CACNA2D1
CCDC64
CAMTA2
J
K
L
M
CELSR3
CLASP2
DNM1
DOCK10

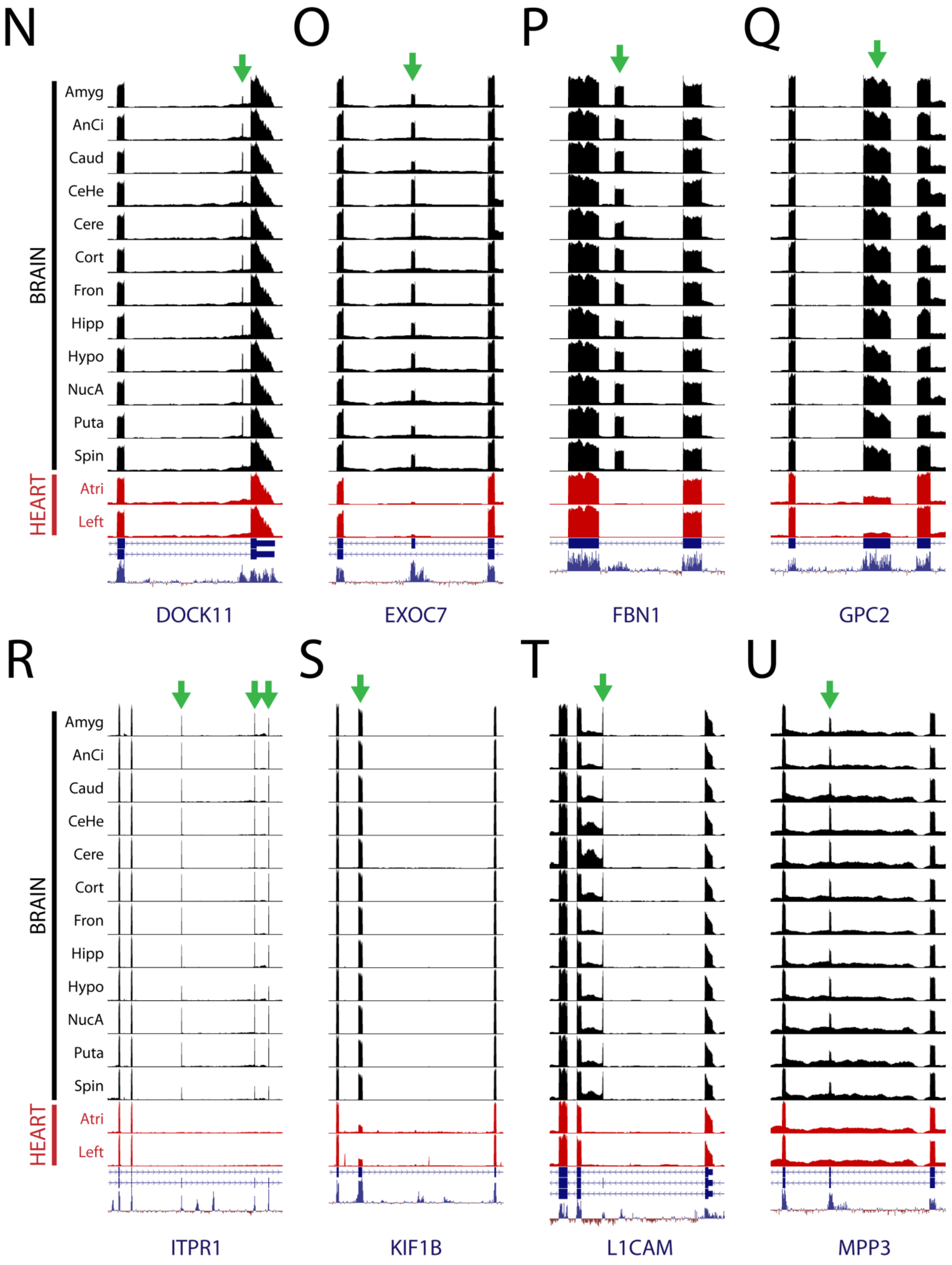
N
O
P
Q
BRAIN
HEART
Amyg
AnCi
Caud
CeHe
Cere
Cort
Fron
Hipp
Hypo
NucA
Puta
Spin
Atri
Left
DOCK11
EXOC7
FBN1
GPC2
R
S
T
U
ITPR1
KIF1B
L1CAM
MPP3

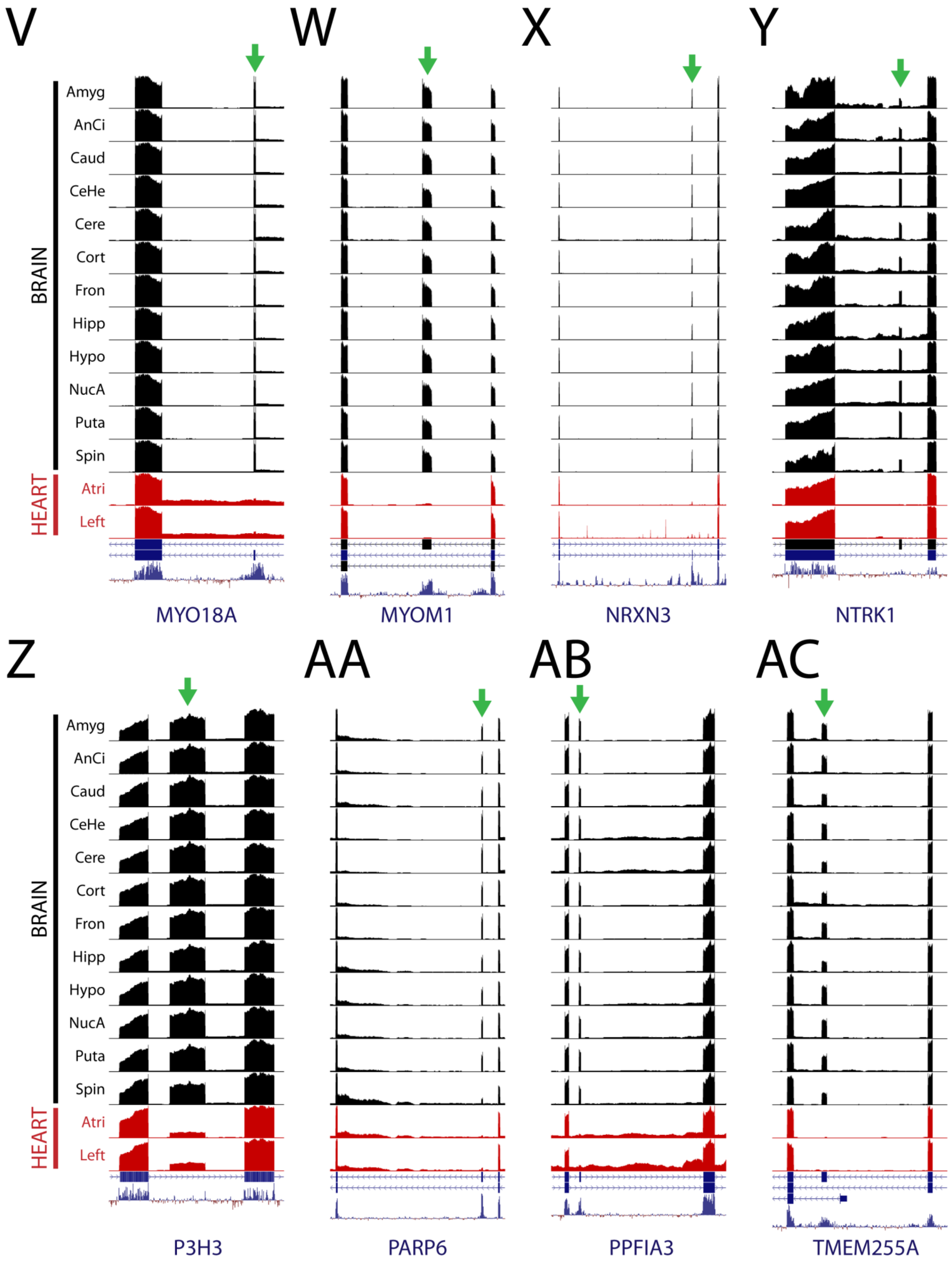

V
W
X
Y
Z
AA
AB
AC
BRAIN
HEART
Amyg
AnCi
Caud
CeHe
Cere
Cort
Fron
Hipp
Hypo
NucA
Puta
Spin
Atri
Left
MYO18A
MYOM1
NRXN3
NTRK1
P3H3
PARP6
PPFIA3
TMEM255A

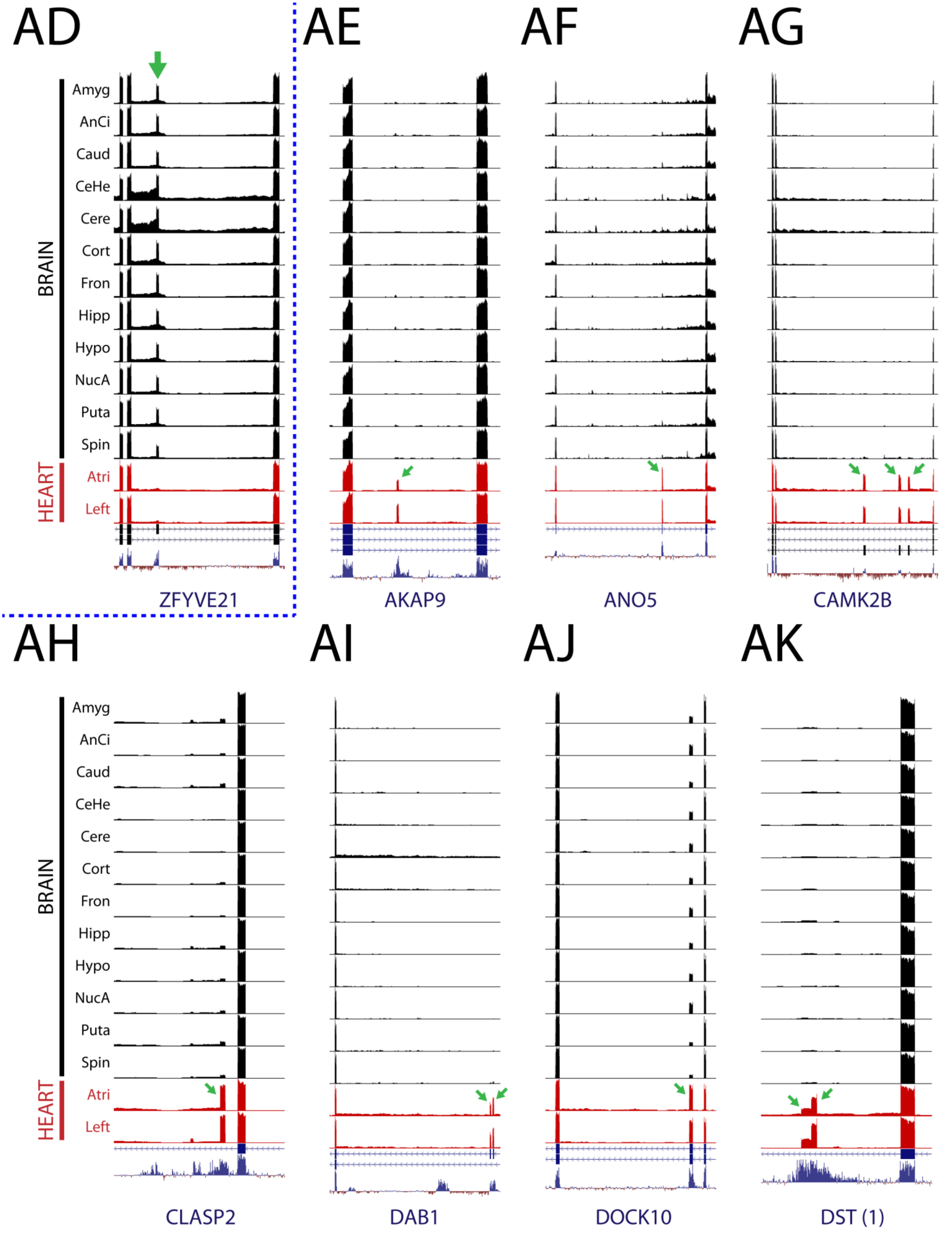
AD
AE
AF
AG
BRAIN
HEART
Amyg
AnCi
Caud
CeHe
Cere
Cort
Fron
Hipp
Hypo
NucA
Puta
Spin
Atri
Left
ZFYVE21
AKAP9
ANO5
CAMK2B
AH
AI
AJ
AK
CLASP2
DAB1
DOCK10
DST (1)

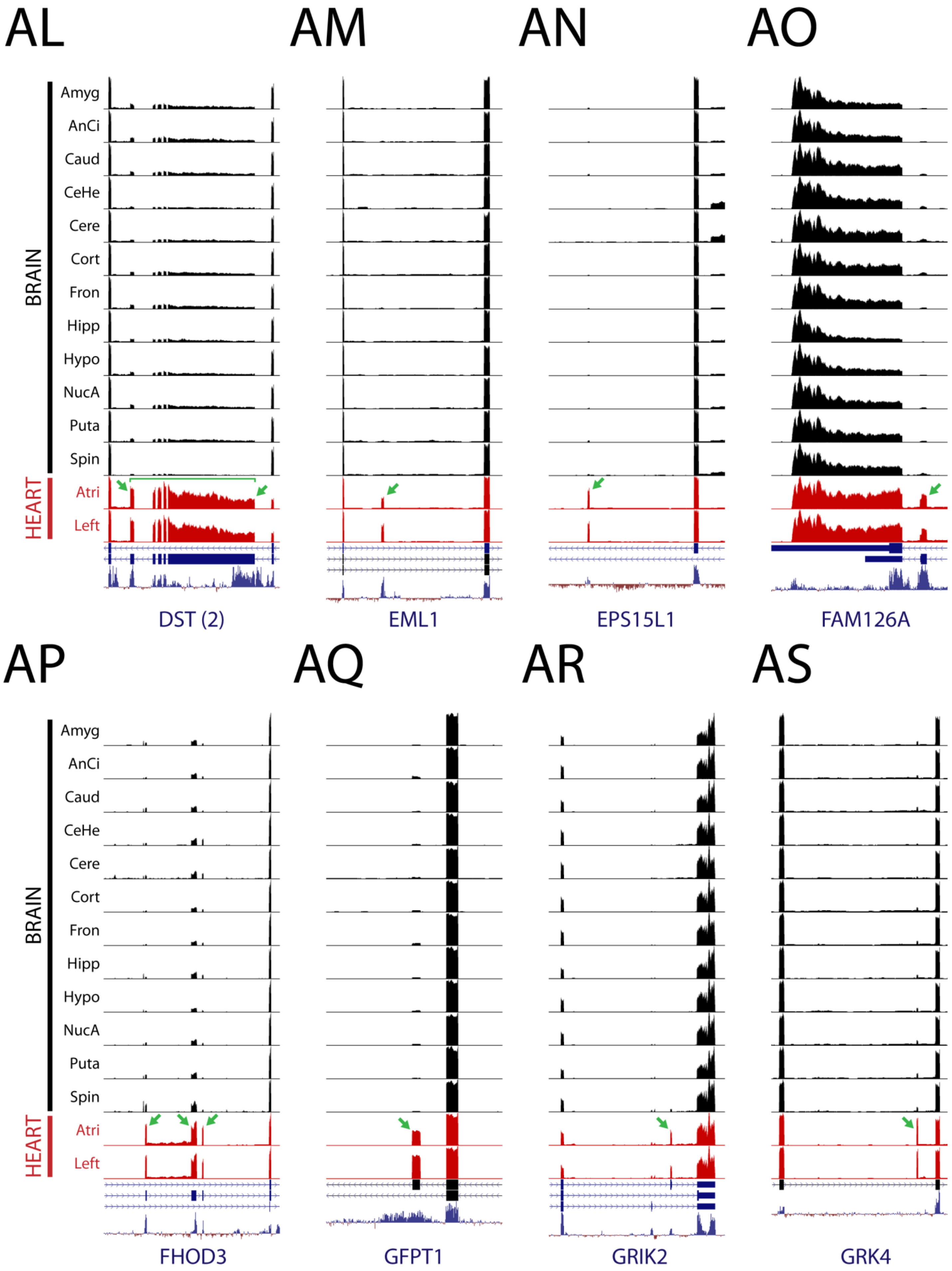
AL
AM
AN
AO
BRAIN
Amyg
AnCi
Caud
CeHe
Cere
Cort
Fron
Hipp
Hypo
NucA
Puta
Spin
HEART
Atri
Left
DST (2)
EML1
EPS15L1
FAM126A
AP
AQ
AR
AS
FHOD3
GFPT1
GRIK2
GRK4

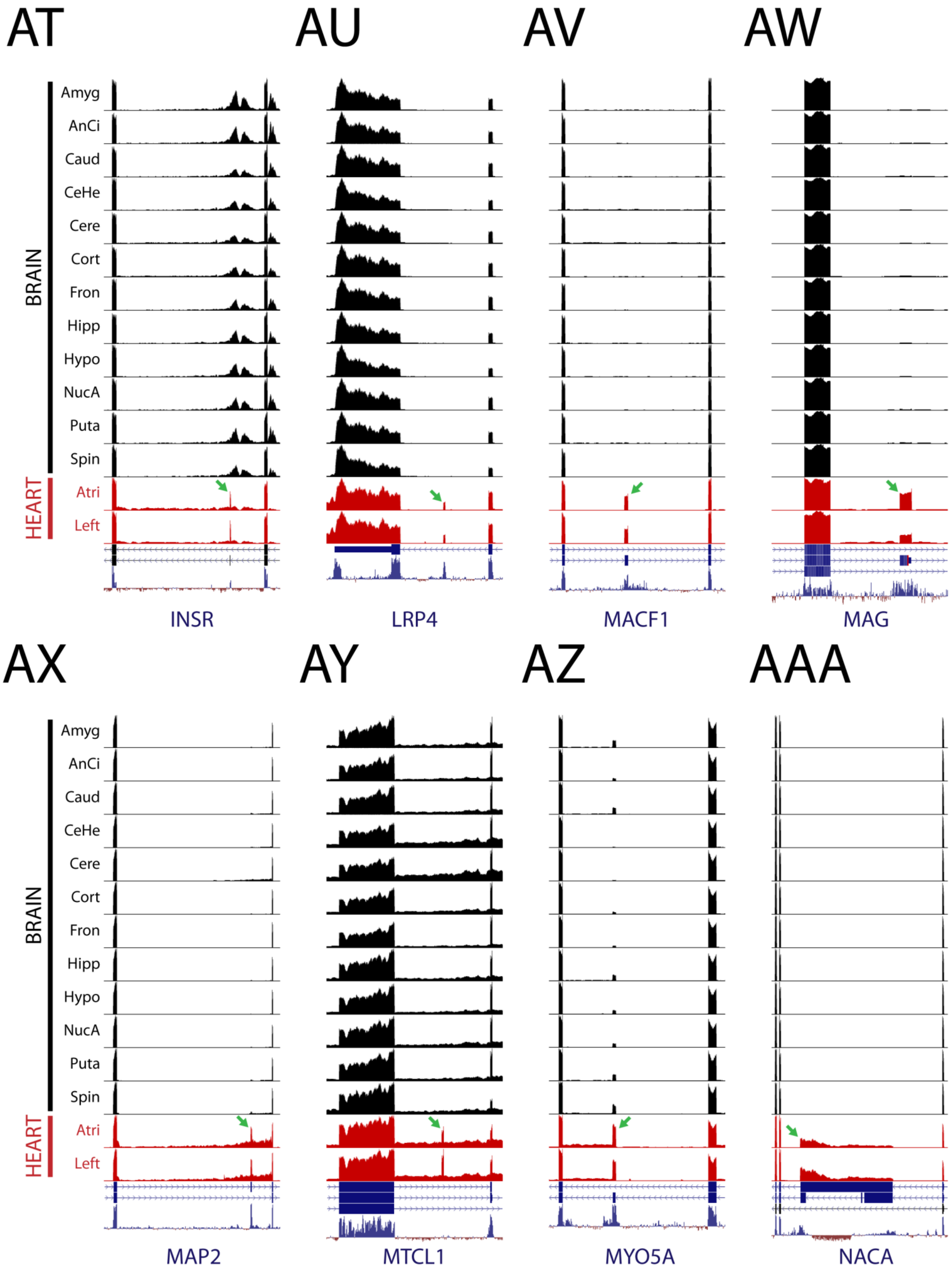

AT
AU
AV
AW
BRAIN
HEART
Amyg
AnCi
Caud
CeHe
Cere
Cort
Fron
Hipp
Hypo
NucA
Puta
Spin
Atri
Left
INSR
LRP4
MACF1
MAG
AX
AY
AZ
AAA
MAP2
MTCL1
MYO5A
NACA

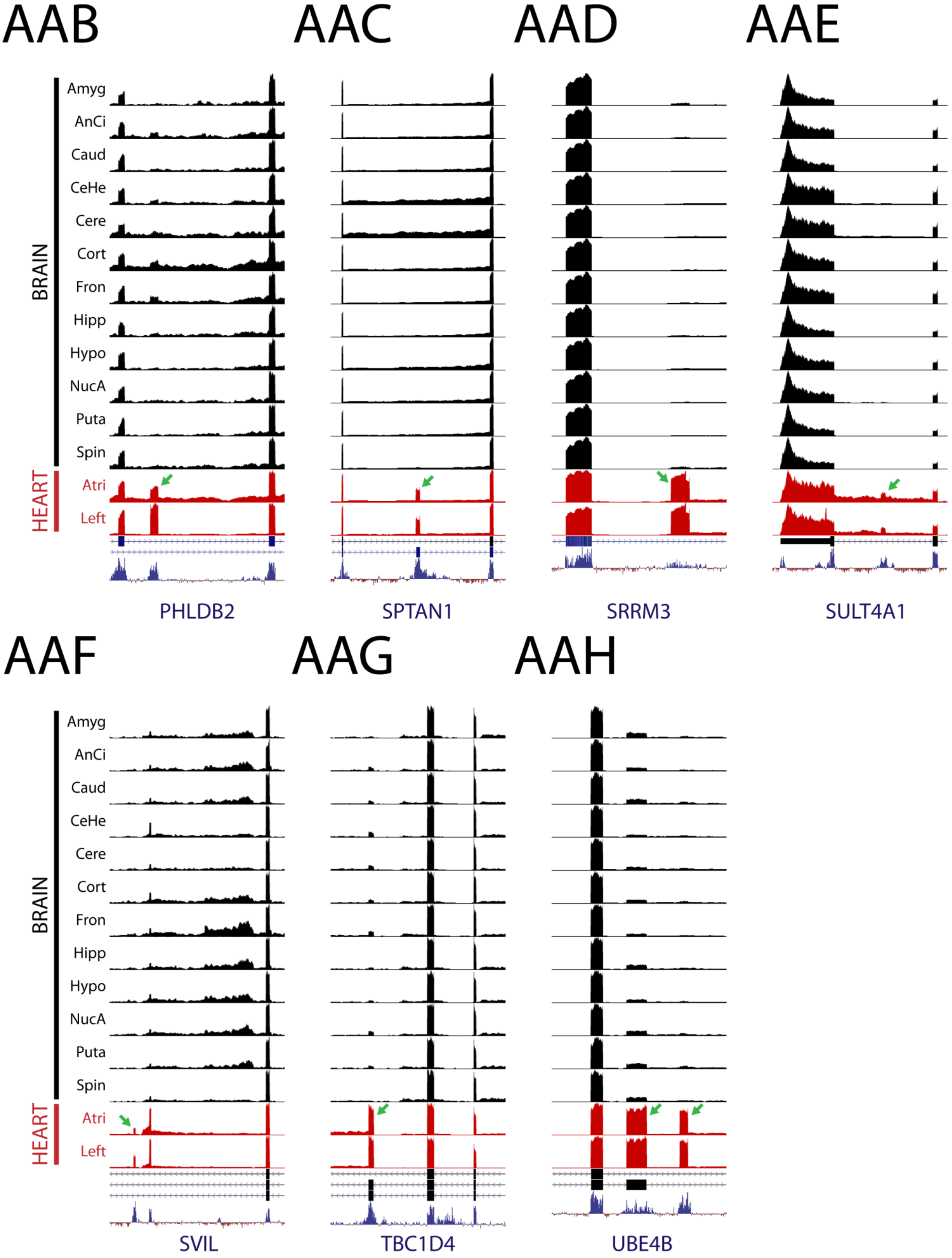
AAB
AAC
AAD
AAE
BRAIN
HEART
Amyg
AnCi
Caud
CeHe
Cere
Cort
Fron
Hipp
Hypo
NucA
Puta
Spin
Atri
Left
PHLDB2
SPTAN1
SRRM3
SULT4A1
AAF
AAG
AAH
SVIL
TBC1D4
UBE4B

**Supplemental Figure 13. Dataset clustering can improve the detection of binary splicing events**

Alternative splicing algorithms such as MAJIQ and LeafCutter can model complex splicing events that are not analyzed by ASCOT. However, ASCOT can reliably detect simple binary splicing events due to the simplified junction walking search logic. (**A** – **AAH**) 60 examples of alternative exons that are not recorded in the LeafCutter shinyapp (https://leafcutter.shinyapps.io/leafviz/) are shown; 30 exons that are highly specific to brain (**A** – **AD**) and 30 exons that are highly specific to heart (**AE** – **AAH**).

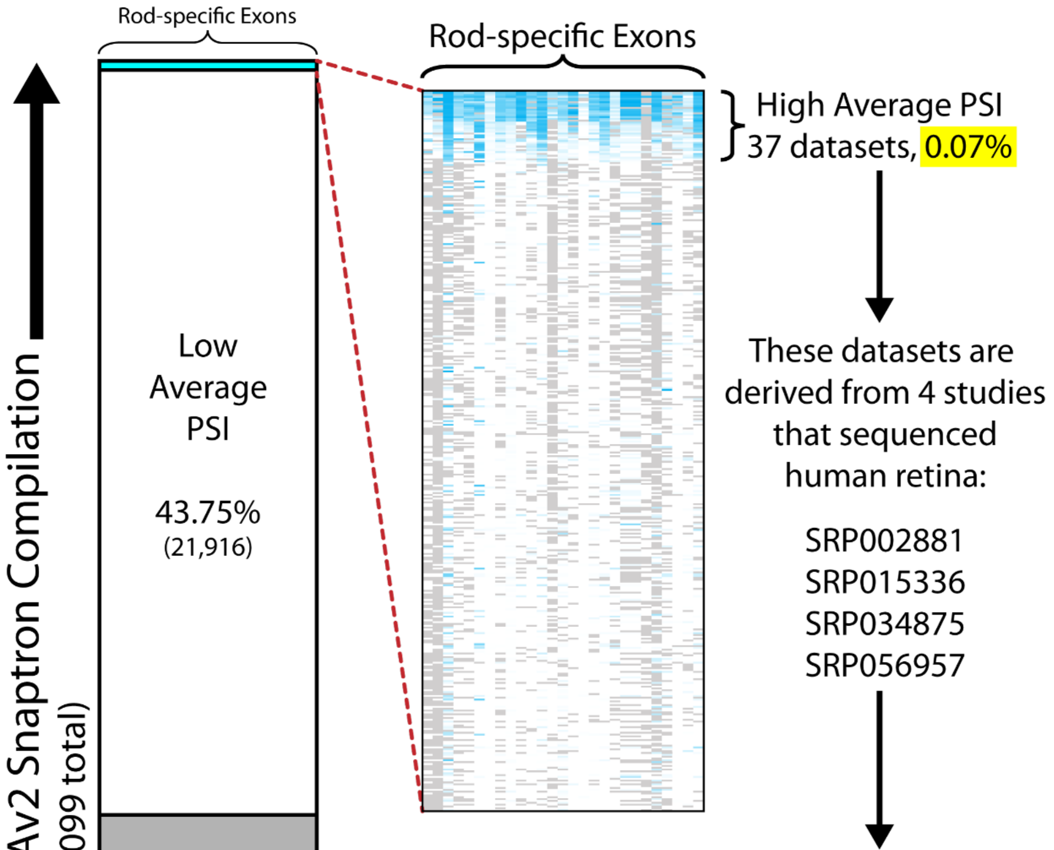


| Snaptron Rail ID | Run Accession | Study Accession | Layout | Data Source |
|---|---|---|---|---|
| 10123 | SRR060734 | SRP002881 | PAIRED | Retina |
| 31575 | SRR548613 | SRP015336 | PAIRED | Retina |
| 43088 | SRR548612 | SRP015336 | PAIRED | Retina |
| 39088 | SRR548611 | SRP015336 | PAIRED | Retina |
| 17066 | SRR1067948 | SRP034875 | PAIRED | Peripheral Retina |
| 10053 | SRR1067942 | SRP034875 | PAIRED | Peripheral Retina |
| 24070 | SRR1067928 | SRP034875 | PAIRED | Peripheral Retina |
| 25588 | SRR1067984 | SRP034875 | PAIRED | Peripheral Retina |
| 29085 | SRR1067996 | SRP034875 | PAIRED | Peripheral Retina |
| 3050 | SRR1067975 | SRP034875 | PAIRED | Peripheral Retina |
| 20571 | SRR1067936 | SRP034875 | PAIRED | Peripheral Retina |
| 48623 | SRR1067988 | SRP034875 | PAIRED | Peripheral Retina |
| 35597 | SRR1067979 | SRP034875 | PAIRED | Macular Retina |
| 44111 | SRR1067970 | SRP034875 | PAIRED | Macular Retina |
| 24570 | SRR1067932 | SRP034875 | PAIRED | Macular Retina |
| 12067 | SRR1068000 | SRP034875 | PAIRED | Macular Retina |
| 38603 | SRR1067986 | SRP034875 | PAIRED | Macular Retina |
| 28597 | SRR1067939 | SRP034875 | PAIRED | Macular Retina |
| 13085 | SRR1067991 | SRP034875 | PAIRED | Macular Retina |
| 8544 | SRR1067946 | SRP034875 | PAIRED | Macular RPE Choroid Sclera |
| 5028 | SRR1067973 | SRP034875 | PAIRED | Macular RPE Choroid Sclera |
| 7661 | SRR1956813 | SRP056957 | SINGLE | hES derived neural retina |
| 1144 | SRR1956814 | SRP056957 | SINGLE | hES derived neural retina |
| 30183 | SRR1956815 | SRP056957 | SINGLE | hES derived neural retina |
| 37191 | SRR1956816 | SRP056957 | SINGLE | hES derived neural retina |
| 43723 | SRR1956817 | SRP056957 | SINGLE | hES derived neural retina |
| 15141 | SRR1956818 | SRP056957 | SINGLE | hES derived neural retina |
| 29172 | SRR1956819 | SRP056957 | SINGLE | hES derived neural retina |
| 19171 | SRR1956820 | SRP056957 | SINGLE | hES derived neural retina |
| 25693 | SRR1956821 | SRP056957 | SINGLE | hES derived neural retina |
| 26676 | SRR1956822 | SRP056957 | SINGLE | hES derived neural retina |
| 42208 | SRR1956823 | SRP056957 | SINGLE | hES derived neural retina |
| 37201 | SRR1956824 | SRP056957 | SINGLE | hES derived neural retina |
| 15134 | SRR1956829 | SRP056957 | SINGLE | hES derived neural retina |
| 17174 | SRR1956830 | SRP056957 | SINGLE | hES derived neural retina |
| 27175 | SRR1956831 | SRP056957 | SINGLE | hES derived neural retina |
| 11154 | SRR1956832 | SRP056957 | SINGLE | hES derived neural retina |

**Supplemental Figure 14. Rod-specific exons in the SRAv2 Snaptron compilation**

Screening the SRA for rod-specific exons, we identify 37 datasets (~0.07%) in the SRAv2 Snaptron compilation with rod-specific exons that originated from human retina RNA-Seq.

**Personal clarification statement from Christopher Wilks**

The use of the terms "synteny" and "conserved", and the PhyloP-derived coverage track in this work are understood by this author to only mean areas of the genome that are similar or fully equivalent across certain groups of species (e.g. mammals) without reference to the origin of that similarity. Use of these terms here does not imply that this author agrees with the idea that all of life derived from a single ancestral organism, which he does not. This view is only this author's view and not shared by his coauthors. The results in this paper have not been affected in any material way by this difference of understanding.

#### Supplemental File Bibliography


1. Wilks, C., Gaddipati, P., Nellore, A. & Langmead, B. Snaptron: querying splicing patterns across tens of thousands of RNA-seq samples. *Bioinformatics* **34,** 114–116 (2018).
2. Vaquero-Garcia, J. *et al.* A new view of transcriptome complexity and regulation through the lens of local splicing variations. *Elife* **5,** e11752 (2016).
3. Terada, Y. *et al.* Novel splice variants of amphiphysin I are expressed in retina. *FEBS Lett.* **519,** 185–90 (2002).
4. Murphy, D., Cieply, B., Carstens, R., Ramamurthy, V. & Stoilov, P. The Musashi 1 Controls the Splicing of Photoreceptor-Specific Exons in the Vertebrate Retina. *PLoS Genet.* **12,** e1006256 (2016).
5. Friedrich, U. *et al.* The Na/K-ATPase is obligatory for membrane anchorage of retinoschisin, the protein involved in the pathogenesis of X-linked juvenile retinoschisis. *Hum. Mol. Genet.* **20,** 1132–42 (2011).
6. Wan, J. *et al.* Dynamic usage of alternative splicing exons during mouse retina development. *Nucleic Acids Res.* **39,** 7920–30 (2011).
7. Ochrietor, J. D. *et al.* Retina-specific expression of 5A11/Basigin-2, a member of the immunoglobulin gene superfamily. *Invest. Ophthalmol. Vis. Sci.* **44,** 4086–96 (2003).
8. Ochrietor, J. D. & Linser, P. J. 5A11/Basigin gene products are necessary for proper maturation and function of the retina. *Dev. Neurosci.* **26,** 380–7 (2004).
9. Clamp, M. F., Ochrietor, J. D., Moroz, T. P. & Linser, P. J. Developmental analyses of 5A11/Basigin, 5A11/Basigin-2 and their putative binding partner MCT1 in the mouse eye. *Exp. Eye Res.* **78,** 777–89 (2004).
10. Riazuddin, S. A. *et al.* A splice-site mutation in a retina-specific exon of BBS8 causes nonsyndromic retinitis pigmentosa. *Am. J. Hum. Genet.* **86,** 805–12 (2010).
11. Murphy, D., Singh, R., Kolandaivelu, S., Ramamurthy, V. & Stoilov, P. Alternative Splicing Shapes the Phenotype of a Mutation in BBS8 To Cause Nonsyndromic Retinitis Pigmentosa. *Mol. Cell. Biol.* **35,** 1860–70 (2015).
12. Bowne, S. J. *et al.* Why do mutations in the ubiquitously expressed housekeeping gene IMPDH1 cause retina-specific photoreceptor degeneration? *Invest. Ophthalmol. Vis. Sci.* **47,** 3754–65 (2006).
13. Tasic, B. *et al.* Adult mouse cortical cell taxonomy revealed by single cell transcriptomics. *Nat. Neurosci.* **19,** 335–46 (2016).
